# Somatic hypomethylation of pericentromeric SST1 repeats and tetraploidization in human colorectal cancer cells

**DOI:** 10.1101/2021.02.19.431645

**Authors:** Beatriz González, Maria Navarro-Jiménez, María José Alonso-De Gennaro, Sanne Marcia Jansen, Isabel Granada, Manuel Perucho, Sergio Alonso

**Author notes:** To whom correspondence should be addressed: Sergio Alonso. equal contribution (Beatriz González, Maria Navarro-Jiménez).

## Abstract

Somatic DNA hypomethylation and aneuploidy are hallmarks of cancer, and there is evidence for a causal relationship between them in knockout mice, but not in human cancer. The non-mobile pericentromeric repetitive elements SST1 are hypomethylated in about 17% of human colorectal cancers (CRC) with some 5-7% exhibiting a more severe age-independent demethylation. Tetraploidy is a common and early event in solid tumors generating subsequent aneuploidy. We compared the relative frequency of chromosomal variations during culture of randomly selected single cell clones of diploid LS174T human CRC cells differing in their levels of SST1 demethylation. Diploid cells underwent frequent genome reduplication events generating tetraploid clones that correlated with SST1 demethylation. In primary CRC, severe SST1 hypomethylation was significantly associated with global genomic hypomethylation and mutations in *TP53*. This work uncovers the association of the naturally occurring demethylation of the SST1 pericentromeric repeat with the onset of spontaneous tetraploidization in human CRC cells in culture, and with *TP53* mutations in primary CRCs. Altogether, our findings provide further support for an oncogenic pathway linking somatic epigenetic and genetic alterations in a subset of human CRC.

## 1. Introduction

### DNA methylation alterations in cancer

DNA methylation is essential for the establishment and maintenance of cell-type specific transcriptional profiles during cell differentiation, conferring cell type identity. It is also involved in suppressing the potentially harmful mobilization of endogenous transposable elements [1,2]. In humans, DNA methylation takes place almost exclusively in the cytosine residues within CpG dinucleotides. In differentiated somatic cells, around 90% of the CpG sites are methylated, with some variation among different cell types. During the lifespan of an individual, the level of genomic methylation gradually declines in all dividing tissues, at a rate proportional to their proliferative potential [3–5]. While the overall level of methylation declines with aging, some *loci* become more methylated [3].

Cancer cells exhibit an accelerated epigenetic aging pace [6], leading to a substantial loss of genome-wide methylation (global hypomethylation) and aberrant hypermethylation of some particular *loci [7,8]*. Genomic DNA hypomethylation, particularly in repeated sequences, can disrupt mitotic processes and cause deletions, translocations, and chromosomal rearrangements [9,10]. Global DNA hypomethylation leads to tumor formation in mice with knocked out methyltransferases, accompanied by chromosomal instability [11]. Genomic hypomethylation mainly reflects the loss of methylation of interspersed repetitive DNA elements that account for up 45% of our genome and over 90% of the methylated CpG sites [12–14]. Among these, LINE-1 elements (together with Alu elements) have been widely employed as surrogate markers of global DNA methylation due to their high CpG-content and abundance [15,16].

We previously found that global hypomethylation in gastric and colorectal cancers increased with patient age and correlated with genomic damage [17]. Based on these findings, a “wear and tear” model, linking aging to cancer development, was proposed. It postulated that changes in methylation - particularly hypomethylation - occur because of the inevitable accumulation of errors during replication of the genome in stem cells or their descendants during aging. When a minimum tolerable demethylation level is reached, or some particular sequences are affected, proper mitosis is compromised and errors in chromosomal segregation occur [17].

### Hypomethylation of pericentromeric repeat SST1 sequences in cancer

One of the *loci* found frequently hypomethylated in gastrointestinal tumors corresponded to pericentromeric repeated elements SST1 [17–19], also known as NBL2 [20]. These 1.4kb long sequences are moderately repeated in tandem clusters, mainly located near the centromeres of the short arm of acrocentric chromosomes 13, 14, 15 and 21. A severe hypomethylation of these sequences occurred in some CRC patients younger than the average, suggesting the existence of some specific demethylation mechanism other than the age-dependent stochastic process [17].

We also previously found that SST1 demethylation was associated with changes in the histone code of these sequences and with down-regulation of *HELLS* (Helicase Lymphoid-Specific Enzyme) [18], a DNA-helicase that plays an essential role in the maintenance of proper levels of methylation of repetitive DNA elements, for which it has been dubbed “the epigenetic guardian of repetitive elements” [21]. More recently, we reported that SST1 demethylation led to the expression of a novel long non coding RNA (tumor-associated NBL2 transcript, TNBL), stable throughout the mitotic cycle, that forms a perinucleolar aggregate during interphase [22]. Despite the fact that aberrant hypomethylation of SST1 sequences has been found in CRC and other cancer types, its phenotypic effects are unknown.

### The role of aneuploidy in cancer

Cancer cells exhibit numerous types of genetic alterations, i.e. chromosomal rearrangements, amplifications, deletions, point mutations, etc. An abnormal number of chromosomes, or aneuploidy, is one of the earliest discovered and most prevalent characteristics of cancer cells. Around 90% of solid tumors exhibit some degree of aneuploidy. More than one century ago, Theodor Boveri proposed that aneuploidy could, in fact, promote malignant transformation [23,24]. In the seventies, before the era of cellular oncogenes and tumor suppressor genes, there was a general support for this view [25]. Whether aneuploidy is the driving force of malignant transformation is still a hot debated topic [26].

Aneuploidy is detrimental to the fitness of non-malignant cells during development or when introduced experimentally. However, it is tolerated by cancer cells and, in some contexts, it might even contribute to their fitness [27]. Aneuploidy can be generated by different mechanisms. After chromosomal alignment in metaphase, sister chromatids are segregated during anaphase. In most cells, cytokinesis occurs simultaneously causing the cytoplasm to divide into two daughter cells accommodating the newly segregated chromosomes [28]. Failure of these processes can affect genomic duplication fidelity. Furthermore, failures during cytokinesis can lead to complete genome duplication, or tetraploidy [29].

### The importance of tetraploidy in cancer

Genome-doubling and consequently tetraploidy is perhaps one of the most underestimated precursors of aneuploidy and chromosomal instability in human cancers [26]. Tetraploid cells are substantially more tolerant to deleterious mutations, since they harbor multiple copies of every gene, thus facilitating cell survival in a mutation-prone environment like that associated with many types of cancers. Also, tetraploidy may trigger chromosomal instability (CIN), probably due to centrosome abnormalities and duplicated chromosome mass, leading to aneuploidy [30,31]. There is experimental evidence directly linking tetraploidy with the onset of tumorigenesis in mice [31,32].

Analyses of chromosome copy number alterations of human cancers have shown that in the ontogenic genealogy of around one third of all solid tumors, a prior doubling event of the genome can be detected. These cancers have a post genome-doubling relative higher rate of chromosome gain and losses [33,34]. Tetraploid cell lines tolerate chromosomal segregation errors better than their diploid precursors [34,35]. While aneuploidy generates allelic imbalances that can be straightforwardly detected using molecular techniques, such as fingerprinting [reviewed in 36], comparative genome hybridization arrays (aCGH) [37,reviewed in 38], SNP arrays [39] and even methylation arrays [40], tetraploidy evades detection using these techniques because the relative chromosomal content, allele balance and presumably methylation profiles remain unaltered.

### *TP53* and aneuploidy in cancer

The protein of Tumor Suppressor Gene P53 (*TP53*) monitors several aspects of DNA repair and cell division, and contributes to suppress tumor progression by hindering the division of cells with damaged DNA. Because of this, *TP53* has been named the “guardian of the genome” [41]. *TP53* mutations are very common in cancer [42] and according to TCGA data they are the only cancer gene mutations significantly associated with CIN [43]. They also associate with tumors displaying evidence for genome duplication [44]. Moreover, it has been shown that wild type p53 blocks the growth of tetraploid cells, through G1 arrest [31,45].

In this work we provide experimental evidence for a link between demethylation of the pericentromeric SST1 repeats and the onset of tetraploidy in human CRC cells in culture, and for the association of drastic demethylation of SST1 with mutations in *TP53* in primary CRC.

## 2. Results

### Isolation of LS174T single cell-derived clones

To test the hypothesis that demethylation of the SST1 pericentromeric repetitive sequences could increase the probability of undergoing mitotic errors [17], we designed a strategy to study the effects of variations in SST1 demethylation level in chromosomal changes in isogenic cells. Among 14 CRC cell lines, LST174T cells exhibited the highest level of SST1 demethylation and also highest range of variation of SST1 methylation [18]. This suggested that it would be feasible to isolate single cell clones with relatively high and low levels of SST1 demethylation. Furthermore, we hypothesized that SST1 demethylation could be accompanied by observable alterations in karyotype during increasing culture time. The approach was facilitated because this cell line, derived from a female CRC patient, is diploid (technically near-diploid, 45 X, after the loss of one of the X chromosomes). LS174T tumor cells present deficiency in DNA mismatch repair (MMR) and microsatellite instability (MSI) [46]. To this end, we generated a collection of randomly chosen 14 single cell-derived clones (C1 to C14) using the limited dilution method, as detailed in materials and methods. To rule out possible contaminations with other cell lines, the identity of these individual clones as well as the original LS174T cells was confirmed by short tandem repeat (STR) analysis (supplemental material).

### LS174T is a mixture of regular and large-nucleus cells

We intended to subject these LS174T-derived single cell clones to long term growth, that we reckoned necessary for observing possible chromosomal variations. However, we unexpectedly observed noticeable differences in cell size among the single cell clones during their initial expansion. We then investigated whether the difference in cell size was reflecting larger nuclei, the existence of multinucleated cells, or just larger cytoplasms. Cell staining with phalloidin and DAPI revealed that cells from clones C5-C8 had substantially larger nuclei in interphase (figure 1A). No multinucleated cells were found in any of the clones. We measured the nuclei area of LS174T and derived clones on microphotographs of cells stained with DAPI (200X magnification, 200∼600 nuclei per sample). The parental cell line displayed an average nuclei area of 89.7±37.8 μm^2^. Clustering of the clones based on the differences of nuclei area showed the existence of two distinct groups: a group comprising clones C1-C4 and C9-C14, with nuclei area of 72.5±26.2 μm^2^; and a group comprising clones C5-C8, with nuclei area of 132.1±50 μm^2^ (figure 1B).

**Figure 1.**
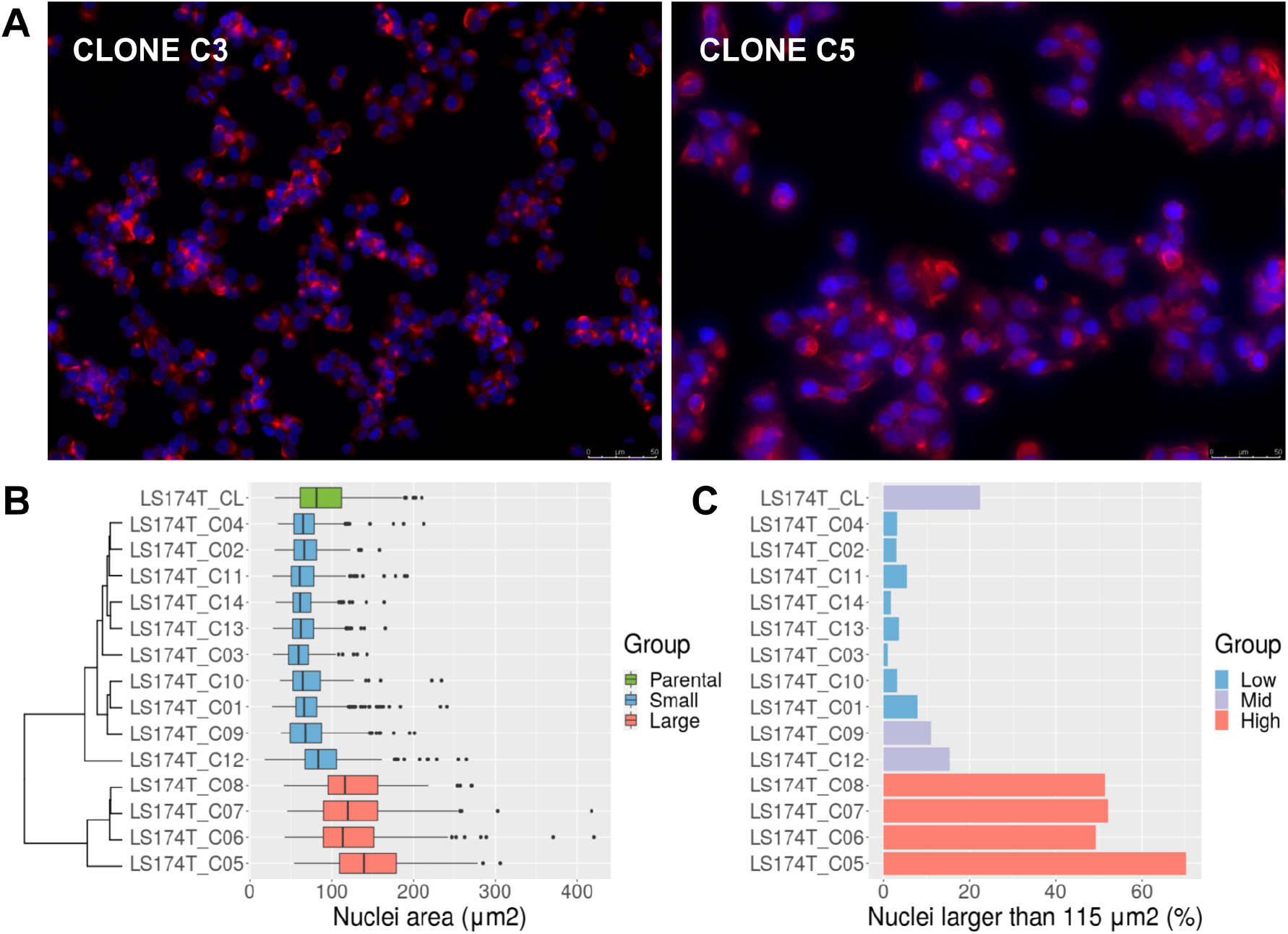
**A)** Morphological analysis of LS174T clones. The figure shows a small cell size clone (C3) and a large cell size clone (C5) stained with phalloidin (red, cytoskeleton) and DAPI (blue, nuclei). Pictures were taken at 200X magnification. Scale bar 50 μm is shown in the bottom right corner of the pictures. **B)** Nuclei area of LS174T and derived clones, measured by microscopy after DAPI staining. The parental cell line (CL) in green. Clones are colored according to the average nuclei size into small (blue) and large (red), and classified by unsupervised clustering based on the difference of their average nuclei size (dendrogram on the left). **C)** Percentage of nuclei with areas larger than 115 μm^2^ in LS174T and derived clones. Most of the clones exhibited a low proportion (less than 10%, low group, blue) of nuclei above 115 μm^2^. The parental cell line (CL) and clones C9 and C12 exhibited a larger proportion (10%∼25%, mid group, in purple). Clones C5-C8 exhibited around 50% of nuclei larger than that value (high group, in red).

The difference in nuclei size among clones suggested that the parental cell line was a mixed population of small and large nuclei cells, which could reflect different degrees of ploidy. Indeed, LS174T nuclei size exhibited a trimodal distribution, with peaks at approximately 65, 115 and 180 μm^2^ (supplemental figure S1) that could not be explained just by the coexistence of cells in G0/G1 vs. cells in S/G2. Doubling of a nucleus volume corresponds to a ∼2^2/3^≈1.6 times increase of its area. Thus, nuclei with areas of 115 μm^2^ would correspond to roughly 2 times the volume of nuclei with areas of 65 μm^2^ and nuclei with areas of 180 μm^2^ over 4 times that.

To estimate the proportion of cells with large nuclei, a threshold of 115 μm^2^ was set based on the nuclei distribution of the parental LS174T cells. About 22% of the LS174T cells had a nucleus larger than 115 μm^2^. Over 50% of the cells in clones C5-C8 had a nucleus larger than that. Clones C1-C4, C10-C11 and C13-C14 had less than 10% of nuclei over 115 μm^2^. Two LS174T-derived clones classified as small according to their average nuclei area exhibited a higher proportion of nuclei larger than 115 μm^2^, i.e. C9 (11%) and C12 (15%) (figure 1C).

To expedite the process and analyze a larger number of nuclei per sample, genomic content was measured in LS174T and its derived clones using flow cytometry (FCM). In agreement with the DAPI staining, FCM analysis confirmed that clones C5-C8 had approximately two times more genomic content than clones C1-C4 and C9-C14 (supplemental figure S2).

### LS174T is composed of a mixture of cells with 45, 46 and 90 chromosomes

To analyze the chromosomal content, the LS174T parental cell line and its derived clones were karyotyped. LS174T exhibited a mixture of near-diploid (45,X) and near-tetraploid (90,XX) cells, with sporadic appearance of 46,X cells with trisomy of chromosome chr7 (figure 2A). In agreement with the nuclei size and cytometry results, clones C5, C6, C7 and C8 were composed by near-tetraploid cells (90,XX). Clones C1, C3, C4, C10, C11 exhibited almost exclusively near-diploid nuclei (45,X). Clones C2, C13 and C14 exhibited cells with 46,X and trisomy of chromosome 7 (figure 2B).

**Figure 2.**
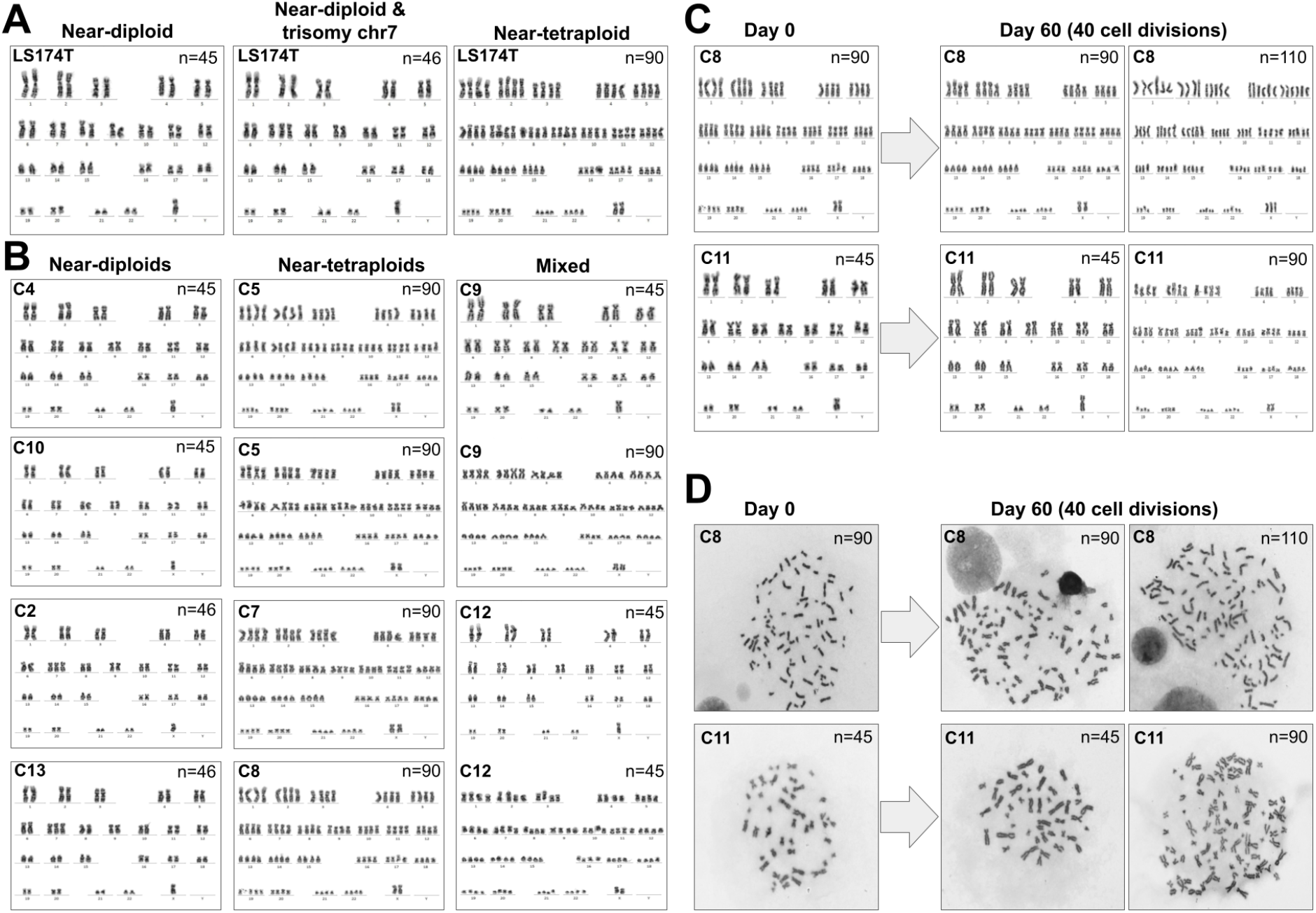
**A)** Karyotypes of the parental cell line LS174T exhibited a mixture of near-diploid (n=45, left), near-diploid with trisomy of chromosome 7 (n=46, middle), and near-tetraploid cells (n=90, right). **B)** left, karyotypes of near-diploid clones C4 and C10 (n=45), and near-diploid clones C2 and C13 with trisomy of chr7 (n=46); middle, near-tetraploid clones C5, C6, C7 and C8; right, clones with mixture of karyotypes C9 and C12. **C)** Karyotypes of LS174T clones C8 and C11. At time 0 (left) C8 cells were pure near-tetraploids and C11 cells were almost pure near-diploids. After 60 days of culture (right) C8 cells remained mostly near-tetraploid with the occasional appearance of higher ploidy cells, while C11 cells exhibited a 13.3% of near-tetraploid cells. **D)** Metaphases of the karyotypes presented in panel C.

As mentioned above, clones C9 and C12 showed a substantial proportion of large nuclei. The karyotyping analysis confirmed that they were a mixture of near-diploid and near-tetraploid cells. An obvious interpretation was that some diploid cells underwent spontaneous tetraploidization during the culturing time after isolation, suggesting that tetraploidization is a dynamic process in LS174T cells and not the result of a single past event. However, it was also possible that clones C9 and C12 were derived from two (or more) cells with different chromosomal content and substantially different proliferation rates favoring a larger proportion of near-diploid cells after several rounds of cell division.

To investigate this possibility, we selected clones C8 and C11 as representative of pure near-tetraploid and near-diploid cells, respectively, and determined their karyotype after 60 days of culture, equivalent to over 40 cell divisions considering an experimentally estimated doubling time of 35h (supplemental figure S3). Clone C8 cells exhibited an almost stable near-tetraploid karyotype. No near-diploid cells were found at any time point in this clone. At time 0, 100% of the metaphases were near-tetraploid (97 out of 97). After 60 days of culture we found three mitotic cells out of 72 with n>90 chromosomes (figure 2C). On the other hand, clone C11 cells exhibited pure near-diploid karyotype at time 0, with only one near-tetraploid metaphase out of 95 mitoses analyzed (1.03%), but we found 10 out of 75 mitoses (13.3%) with near-tetraploid metaphases after 60 days of culture (*p*=0.003, Fisher’s exact test) (figure 2C). Since the individual cells analyzed for their karyotype originated from a clone that was homogeneously diploid, and the mitoses were typical of single cells and not mixtures of different cells (figure 2D), we concluded that these mitotic cells underwent tetraploidy from a prior parent diploid cell. Modeling the generation of tetraploid cells in clone C11, we estimated that approximately 2.67% of the mitosis resulted in spontaneous tetraploidization (supplemental figure S4).

### Analysis of other cancer cell lines

In an effort to extend these observations, we analyzed the degree of anisonucleosis, i.e. variability in cell nuclei size, calculating the coefficient of variation (CV) of the nuclei area as a proxy for genomic content in HCT116 and DLD-1 CRC cell lines. The level of anisonucleosis in LS174T (CV=42%) was higher than in HCT116 (CV=33%, asymptotic test *p*=4.2×10^−7^) and DLD-1 (CV=38%, asymptotic test *p*=0.036), despite the three cell lines being MSI and originally described as mainly diploid. Moreover, the proportion of cells with large nuclei (over 3 times the modal nucleus volume, where the modal nucleus volume was assumed to correspond to diploid cells in G0/G1) was significantly higher in LS174T (12.7%) than in DLD-1 (7%, Fisher’s test *p*=0.004) and HCT116 (1%, Fisher’s test *p*=6.3×10^−16^) (supplemental figure S5). The original publication of the isolation of LS174T reported the presence of 6.1% of tetraploid cells [47]. The higher incidence of polyploidy in our cultures of LS174 cells is likely due to the accumulation of tetraploid cells during the culture passages. Due to the absence of significant SST1 demethylation in DLD-1 and HCT116 cell lines (supplemental figure S5), a similar study to isolate single cell clones with substantially different levels of SST1 methylation was not feasible.

We then analyzed an ovarian cancer cell line, OV-90, that also displayed considerable SST1 demethylation. OV-90 was originally described as mainly diploid albeit displaying complex karyotypic changes involving chromosomal rearrangements, translocations, deletions, and duplications [48]. In contrast with LS174T, OV-90 is microsatellite stable (MSS) and harbors a mutant *TP53* gene. Karyotype analysis of single cell clones of OV-90 showed aberrant and very variable chromosomal content with an overall distribution of hypertriploidy-hypopentaploidy. We also found numerous multipolar spindle mitoses, indicative of active chromosomal instability (supplemental figure S6). In addition to several chromosomes with two and three copies, the karyotypes presented several chromosomes with four copies (for instance, the karyotype of supplemental figure S6 shows chrs. 2, 9, 19, 20, 21 and X). Thus, these results are consistent with an ancestral tetraploid cell lineage. However, the high chromosomal instability of these cells precluded detecting methylation-associated chromosomal changes in the time frame of the cultures (data not shown).

### Chromosomal content correlates with SST1 methylation level in LS174T cells

We analyzed the methylation level of SST1 sequences in the 14 LS174T-derived clones as well as LS174T parental cell line. We used bisulfite sequencing, following our previously reported methodology that interrogates 27-28 CpG sites (some of them polymorphic) within the SST1 CpG-rich region originally found to undergo hypomethylation in cancer [18]. The methylation of individual SST1 elements within each single-cell derived clone was highly variable (figure 3A). The ANOVA analysis of the methylation levels of the 14 single-cell clones revealed that only 10.7% of the methylation variance was explained by differences among them, while 89.2% reflected intra-clonal variability that we interpret as differences in methylation among SST1 elements within every cell. Despite this variability, the near-tetraploid clones (C5-C8) exhibited statistically significant lower average methylation than the near-diploid clones (30.1±2.8% vs 43.6±4.0%, t-test *p*=1.0×10^−4^). The clones with mixed populations of near-diploid and near-tetraploid cells (C9 and C12) also exhibited low levels of methylation (30.5±3.8%). These results indicated a significant inverse correlation between SST1 methylation and tetraploidization in LS174T cells (r^2^=0.53, *p*=0.002, figure 3B).

**Figure 3.**
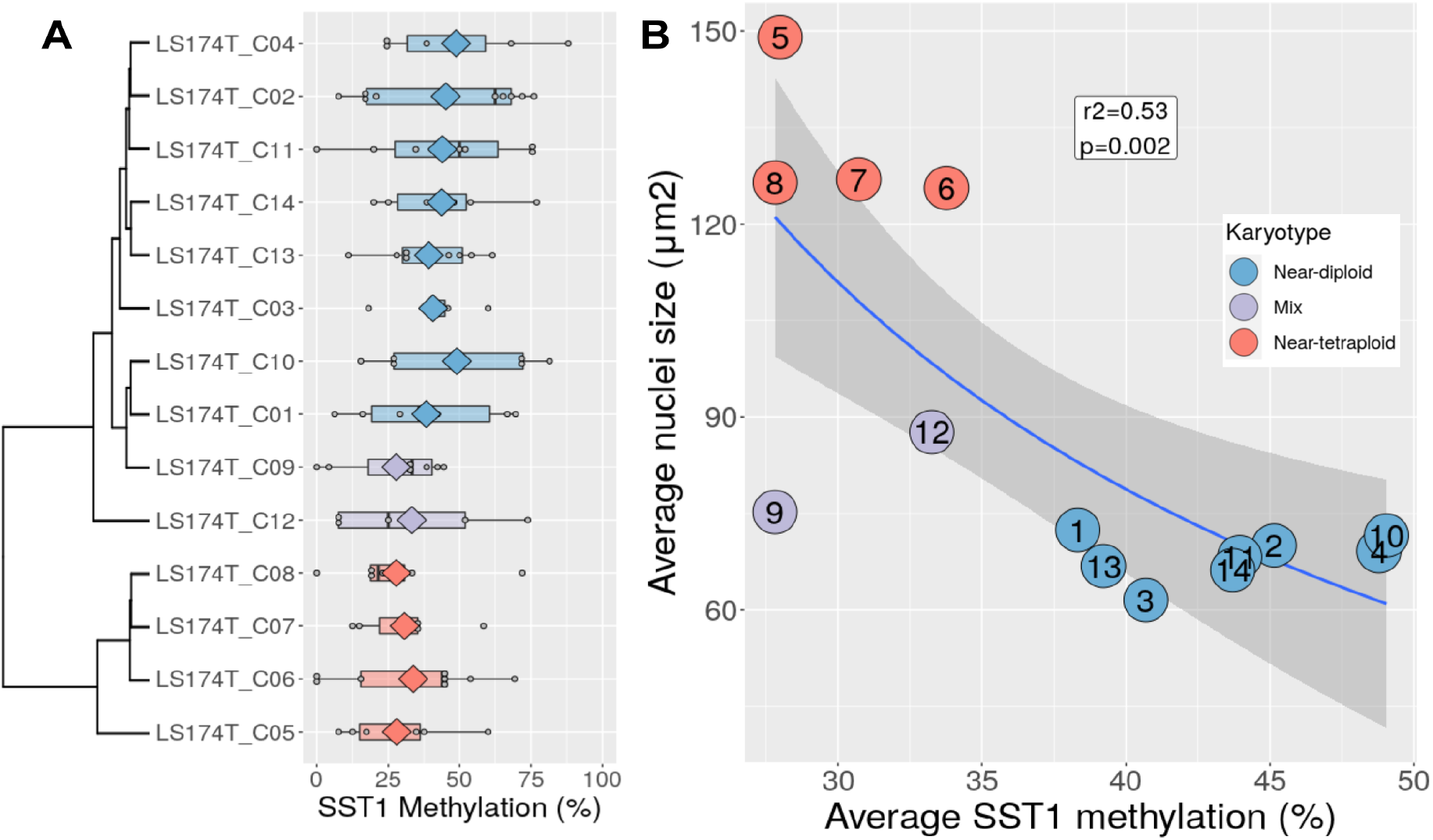
**A)** SST1 methylation levels measured by bisulfite sequencing in LS174T-derived single cell clones. Clones are ordered according to their nuclei size-based clustering (see figure 1), and colored according to their karyotype. Every grey dot represents the average methylation of an individually sequenced SST1 molecule. Diamonds indicate the average methylation level of all SST1 the molecules sequenced in every clone. **B)** SST1 average methylation (x-axis) vs. average nuclei size (y-axis) in LS174T-derived single cell clones. Every dot represents the value of a LS174T-derived cell clone, colored according to its karyotype. The regression line (nuclei area ∼ 1/methylation) and its 95% confidence interval are depicted in blue and dark grey, respectively. The inverse correlation between nuclei area and methylation was r=0.73, r²=0.53, p=0.002.

All together, these results show that severe demethylation of SST1 sequences is associated with tetraploidization in LS174T cells, a process that appears to be spontaneous and active with a relatively high frequency.

### Transcriptional profile differences between near-diploid and near-tetraploid LS174T cells

We compared the transcriptional profile of near-tetraploid (clones C5-C8) vs. near-diploid (clones C3, C10, C11, and C14) LS174T cells, using Affymetrix Clariom™ S arrays. Although there were no dramatic differences in gene expression, 0.4% (84/21448) of the probes displayed statistically significant differences (p<0.01, t-test, supplemental figure S7 and supplemental information). Of note, *HDLBP* was the most significant down-regulated gene in the near-tetraploid cells (supplemental figure S7). This gene encodes vigilin, a multi-functional protein that, among other functions, plays an essential role in maintaining genome ploidy [49]. Thus, down-regulation of *HDLBP* might be functionally related to the spontaneous tetraploidization observed in LS174T cells. We also identified downregulated *TBC1D16* and *GNAO1*, whose expression have been reported to be altered through DNA methylation in different types of cancers [50,51]. Moreover, comparing Hallmark gene signatures using Gene set enrichment analysis (GSEA) [52], overexpression of E2F targets, MYC targets and G2M checkpoint genes were significatively enriched in the near-tetraploid cells (supplemental figure S8), all consistent with the fact that the control of the cell cycle is altered in these cells [53].

### SST1 demethylation associates with LINE1 demethylation in primary CRCs

We then analyzed 148 primary CRC tumors and their matching normal tissues for SST1 demethylation status and correlated it with genotype and phenotype. Clinicopathological, mutational and DNA methylation data from these 148 CRC is shown in supplemental table 1. To expedite the number of samples analyzed and obtain quantitative measurements, we designed a novel assay named methyl-sensitive QPCR (MS-QPCR), very similar in concept to MethyLight [16]. SST1 MS-QPCR estimates the proportion of completely (or near-completely) unmethylated SST1 elements relative to the total number of amplifiable elements. This value was termed *relative demethylation level* (RDL). Hence, higher RDL indicates a higher proportion of unmethylated molecules, i.e. lower levels of methylation. We employed a similar approach to analyze LINE-1 methylation in 107 of these cases. A detailed explanation of SST1 and LINE-1 MS-QPCR is provided in the methods section. To benchmark MS-QPCR, forty of these cases were chosen among those previously analyzed by bisulfite sequencing. We obtained very concordant results between both techniques (r=0.73, p=8×10^−8^, supplemental figure S9).

**Table 1.** Association of SST1 severe demethylation with CRC genotype and phenotype.

| | Not demethylated<br>SST1 ΔRDL<br>< 1.0 | Moderately demethylated<br>SST1 ΔRDL<br>> 1.0 & < 3.0 | Severely demethylated<br>SST1 ΔRDL<br>> 3.0 | Fisher's test<br>p-value |
| --- | --- | --- | --- | --- |
| Gender | Women (n=59)<br>Men (n=64) | Women (n=6)<br>Men (n=12) | Women (n=2)<br>Men (n=5) | p1=0.19<br>p2=0.46 |
| Age | < 66 yo (n=57)<br>> 66 yo (n=66) | < 66 yo (n=10)<br>> 66 yo (n=8) | < 66 yo (n=5)<br>> 66 yo (n=2) | p1=0.27<br>p2=0.27 |
| Ethnic | Caucasian (n=94)<br>Afr.Am. (n=18) | Caucasian (n=9)<br>Afr.Am. (n=6) | Caucasian (n=3)<br>Afr.Am. (n=1) | p1=0.05<br>p2=0.58 |
| Tumor Location | Proximal (n=69)<br>Distal (n=50) | Proximal (n=10)<br>Distal (n=8) | Proximal (n=2)<br>Distal (n=5) | p1=0.38<br>p2=0.24 |
| Dukes' stage | IS-A-B (n=52)<br>C-D-M (n=71) | IS-A-B (n=9)<br>C-D-M (n=9) | IS-A-B (n=4)<br>C-D-M (n=3) | p1=0.39<br>p2=0.70 |
| MSI status | MSS (n=102)<br>MSI (n=21) | MSS (n=17)<br>MSI (n=1) | MSS (n=6)<br>MSI (n=1) | p1=0.37<br>p2=1.00 |
| *TP53* | WT (n=44)<br>MUT (n=32) | WT (n=5)<br>MUT (n=11) | WT (n=0)<br>MUT (n=7) | p1=**0.0037\*\***<br>p2=**0.012\*** |
| *KRAS* | WT (n=75)<br>MUT (n=48) | WT (n=12)<br>MUT (n=6) | WT (n=7)<br>MUT (n=0) | p1=0.18<br>p2=**0.048\*** |
| *BRAF* | WT (n=112)<br>MUT (n=10) | WT (n=16)<br>MUT (n=2) | WT (n=4)<br>MUT (n=2) | p1=0.25<br>p2=0.10 |

*p1*: Fisher's test p-value comparing not-demethylated tumors vs the rest (demethylated and severely demethylated).
*p2*: Fisher's test p-value comparing severely demethylated tumors vs the rest. \*: p-value < 0.05. \*\*: p-value < 0.01.
In parenthesis, the number of informative cases in each category. In Ethnic, Afr.Am.: African American. In Duke's stage, tumors were grouped into in-situ (IS, n=1), Duke's A (n=7) and Duke's B (B, n=54) vs. Duke's C (n=58), Duke's D (n=23) and metastases (M, n=2). WT: wild-type. MUT: mutant. Some of the total number of samples do not match the total 148 cases sample due to incomplete information.

SST1 RDL in tumors and matching normals tissues strongly correlated (r=0.64, p=3.9×10^−18^, figure 4A). Most tumors exhibited a degree of SST1 methylation very similar to that of their matching normal tissues, and the average methylation level was very similar in both groups (paired t-test p=0.07, figure 4B). Seven tumors, however, deviated from this trend and exhibited SST1 RDL values higher than the 95% confidence interval of the regression model prediction. These tumors were classified as SST1 severely demethylated (figure 4A and 4B). Methylation of LINE-1 exhibited a very different behaviour. We found no significant correlation between tumor and matching normal tissues (r=-0.06, p=0.56, figure 4C), and tumors overwhelmingly exhibited higher demethylation than normal tissues (paired t-test p=6×10^−23^, figure 4D).

**Figure 4.**
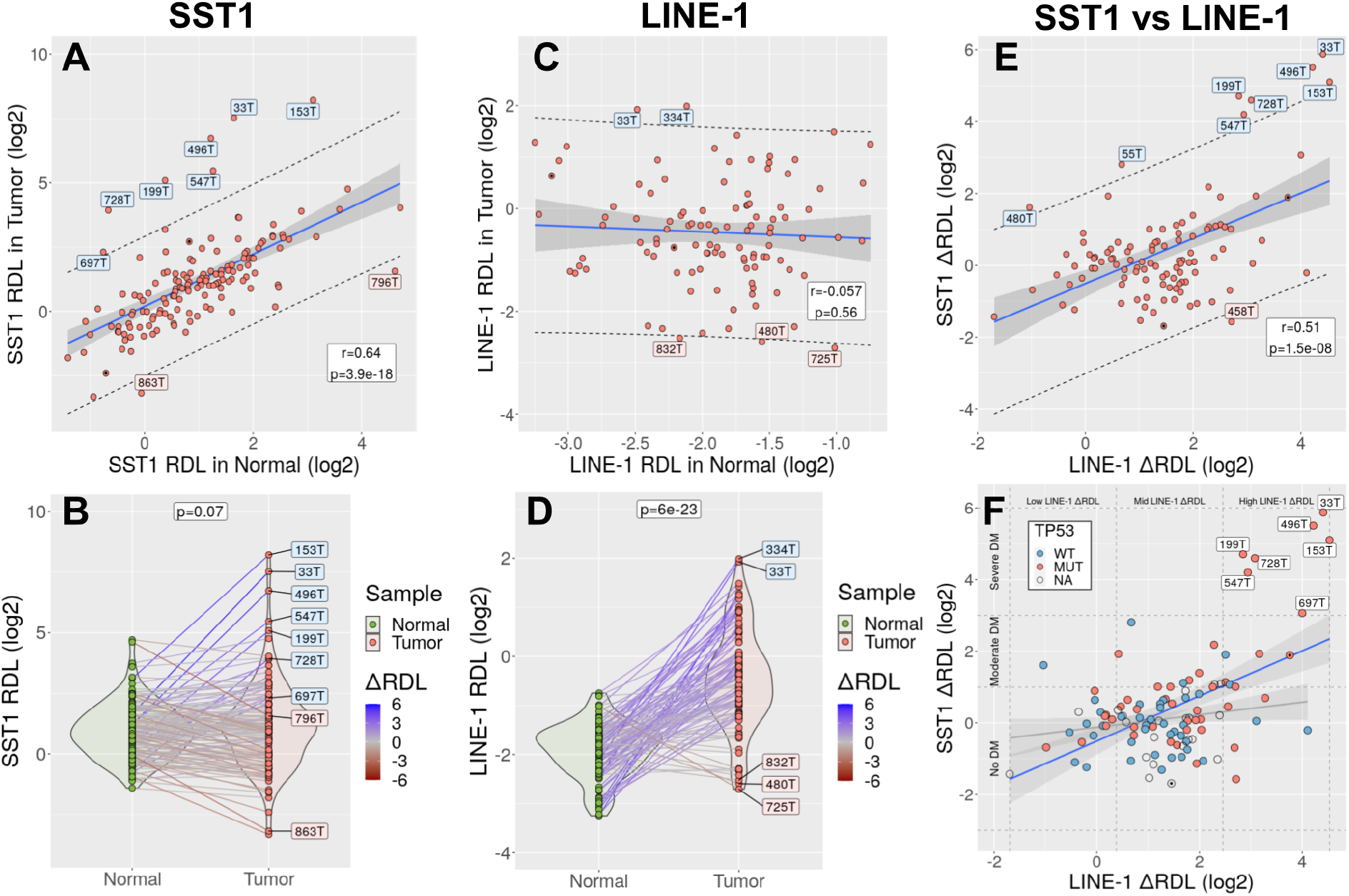
**A)** Correlation between SST1 RDL in 148 CRC tumors (y-axis) and matched normal tissues (x-axis). Every dot represents a tumor. In blue, the regression line. Shaded in grey, the 95% CI of the slope. The dashed lines indicate the 95% CI of the values predicted by the regression model. Labeled in blue and red, tumors with SST1 RDL above (severely demethylated) and below the predicted upper limit of the 95% CI, respectively. Two metastases are indicated with an internal black dot. **B)** SST1 RDL values in normal tissues (green) and tumors (red). Tumor and matched normal samples are connected by solid lines, colored according to their difference in SST1 RDL (somatic demethylation, ΔRDL). Tumors with SST1 RDL values outside the 95% CI of the regression model prediction (shown in panel A) are labeled. **C)** Correlation between LINE-1 RDL in 107 tumors and matched normal tissues. Symbols as in panel A. **D)** LINE-1 RDL values in normal tissues and tumors. Symbols as in panel B. **E)** Correlation between SST1 ΔRDL and LINE-1 ΔRDL. Symbols as in panels A and C. **F)** Correlation between LINE-1 and SST1 somatic demethylation in 107 CRCs. Tumors are colored according to their TP53 mutational status (WT: wild type; MUT: mutant; NA: not analyzed). The graph is divided with dashed lines in 9 areas according to the levels of LINE-1 ΔRDL (horizontal axis, divided in low, mid and high demethylated) and SST1 ΔRDL (vertical axis, divided into no demethylated, moderately demethylated and severely demethylated). Two regression models are depicted, one with all the samples (blue line) and the other excluding the SST1 severely demethylated cases (grey line). The dark grey areas indicate the 95% CI of the slope.

We defined the somatic demethylation value (ΔRDL) as the difference between RDL from the tumors and their matching normal samples, after log2 transformation. Thus, ΔRDL = 0 indicates the same proportion of demethylated SST1 molecules in the tumor and the matching normal sample, ΔRDL = 1 indicates that the proportion of demethylated SST1 molecules is two times larger in the tumor than in the normal, etc. All the seven outlier cases classified as SST1 severely demethylated exhibited SST1 ΔRDL > 3.

SST1 somatic demethylation correlated with LINE-1 somatic demethylation (r=0.51, CI95%=[0.36-0.64], *p*=1.5×10^−8^, figure 4E). This correlation was strongly influenced by the few cases with severe SST1 somatic demethylation, all of them exhibiting high levels of LINE-1 somatic demethylation. Excluding the cases with severe SST1 demethylation, the correlation between SST1 ΔRDL and LINE-1 ΔRDL was much lower, albeit still positive (r=0.2, CI95%=[0.009-0.39]).

Genome-wide methylation profiling of 30 CRCs and their matching normal samples with Illumina HM450K arrays showed that LINE-1 somatic demethylation correlated with genome-wide demethylation, particularly in CpG sites not associated with CpG islands (those located 2000bp or farther apart from their closest CpG island). This association was expected, since LINE-1 methylation is widely considered as a proxy of global genome demethylation. SST1 demethylation, on the other hand, did not correlate with genome-wide demethylation regardless of the analyzed subset of CpG sites (supplemental figure S10). Therefore, we interpret our data as indicative that SST1 severe demethylation is a somatic process occurring in an overall genome demethylation background, but by a hitherto unknown mechanism.

### SST1 severe demethylation is associated with *TP53* mutations in CRC

To analyze the association of SST1 demethylation with clinical and mutational characteristics of the CRCs, cases were classified into three categories based on their SST1 ΔRDL: tumors with severe (ΔRDL > 3, n=7, 4.7%), moderate (1 < ΔRDL < 3, n=18, 12.2%), and without (ΔRDL < 1, n=123, 83.1%) somatic demethylation. For every parameter we performed two comparisons, i.e. tumors with moderate or severe demethylation vs. rest, and tumors with severe demethylation vs. rest. Associations between SST1 somatic demethylation and clinicopathological characteristics of the 148 CRCs are shown in table 1. SST1 somatic demethylation correlated with *TP53* mutations in both comparisons (moderate or severe demethylation vs. rest, *p*=0.0037; and severe demethylation vs. rest, *p*=0.012). Severe demethylation also exhibited a borderline significant correlation with wild-type *KRAS* (*p*=0.048). We also studied the association of the clinical and mutational characteristics with SST1 ΔRDL as a continuous variable, i.e. without performing a categorical classification of the cases (supplemental figure S11). There was a significant correlation of SST1 ΔRDL with *TP53* mutations (*p*=0.0015), with *KRAS* wild-type tumors (*p*=0.017) and with African Americans (*p*=0.007). In a multivariate regression analysis, only *TP53* mutations remained statistically significant (*p*=0.026).

## 3. Discussion

The original aim of this work was to explore the link between DNA demethylation of SST1 pericentromeric repeat elements and chromosomal alterations in human CRC. The approach employed LS174T, a CRC cell line with demethylated SST1 elements [18]. Due to the variability of demethylation of this cell line, we reasoned that isolation of single cell clones with significant differences in levels of methylation could be feasible without any prior selection. The isolation of single cell clones allows the comparison of methylation differences in the same genetic background and in a measurable number of mitotic cycles during cell culture. We hypothesized that LS174T cells with SST1 demethylation could have an increased rate of chromosomal alterations that could be measurable by monitoring the karyotype of single-cell derived clones during an increasing number of cell divisions. The comparison of clones with high and low SST1 demethylation could test our epigenetic → genetic (demethylation → chromosomal alteration) hypothesis. We used a similar approach to isolate single cell clones and compare the mutator consequences of different spontaneous mutations in DNA mismatch repair (MMR) genes in CRC cell lines with microsatellite instability (MSI) [54].

The experimental strategy was facilitated because LS174T is an MSI and mainly diploid cell line. We randomly isolated 14 single cell clones (C1-C14) for further expansion and subsequent analysis of changes in chromosomal content. Unexpectedly, we found that right after isolation, some LS174T clones had a much larger average cell and nuclei size (clones C5 to C8). This was due to the presence of near-tetraploid karyotypes. We observed the presence of near-tetraploid cells in originally near-diploid clones after very few cell divisions (clones C9 and C12, figure 2B). Longer culturing time (∼40 cell divisions) of pure near-diploid cells (clone C11) also generated near-tetraploid cells (figure 2C). This showed that spontaneous tetraploidization was not the result of a single past event, but the result of an intrinsic and active chromosomal segregation defect taking place with an estimated rate of 1 in 37 cells entering mitosis (2.67% of the mitoses, supplemental figure S4).

Tetraploid cells were stable and did not revert to the diploid status, at least within the studied timeframe. They, however, grew at approximately 80% the proliferation rate of the diploid cells (supplemental figure S3) likely due to the extra time required to duplicate a genome that is twice the size of diploid cells. A mathematical model of the proportion of diploid and tetraploid cells in a mixed cell population, considering their different growth rates the rate of spontaneous tetraploidization predicted that the proportion of tetraploid cells would stabilize at around 13% of the population (supplemental figure S4). This prediction is in line with the proportion of cells with nuclei size above 3x de modal volume in the parental LS174T cell population (12.8%, supplemental figure S5).

In agreement with the initial hypothesis, tetraploidy in LS174T single cell-derived clones strongly associated with lower levels of methylation of SST1 elements (r^2^=0.53, p=0.002, figure 3B). The results with the ovarian cancer cell line OV-90 also suggested that drastic SST1 demethylation results in a high risk for tetraploidization. Altogether, our results suggest that drastic demethylation of SST1 elements induces tetraploidy in human cancer cells.

The link between SST1 demethylation and spontaneous tetraploidization in LS174T reinforces the hypothesis that the underlying reason for the association between SST1 severe demethylation and *TP53* mutations is that SST1-associated tetraploidization would require inactivation of *TP53* to evade tetraploidy-triggered cell cycle arrest [31,45,55]. While LS174 cells are *TP53* wild-type, they harbor a homozygous frameshift mutation in *BAX*, and also in other pro-apoptotic genes. *BAX* is a common MSI target because it contains a mutational hotspot (G)_8_ mononucleotide tract in its coding sequence [56,57]. It has been shown that inactivation of *BAX* facilitates the survival of tetraploid cells at least as efficiently as the p53 or p21 knockouts [58]. Our experiments indicate that upon tetraploidization, LS174T cells activated the G2M checkpoint genes (supplementary figure S8), but nevertheless they continued to proliferate and did not enter apoptosis or cell cycle arrest. When p53 and BAX are functional, the cells undergo mitotic arrest and apoptosis. If the cell cycle arrest and/or apoptosis networks are derailed, either by somatic *TP53* mutations in MSS tumors, or mutations in the *BAX* and other apoptotic genes in MSI tumors, the cells undergoing genome reduplication are able to survive.

In MSS tumors, tetraploid cells are prone to subsequent chromosomal changes ending in the aneuploidy typical of these solid tumors that underlie the unmasking of recessive mutations by LOH and amplification of dominant cancer genes. In MSI tumors this is similarly achieved by their mutator phenotype causing many point mutations with similar phenotypic effect, thus substituting aneuploidy. In MSI tumors, tetraploidization is also tolerated, but due to the growth disadvantage of tetraploid cells, the tumor may progress retaining their overall near diploidy (see our model in figure 5).

**Figure 5.**
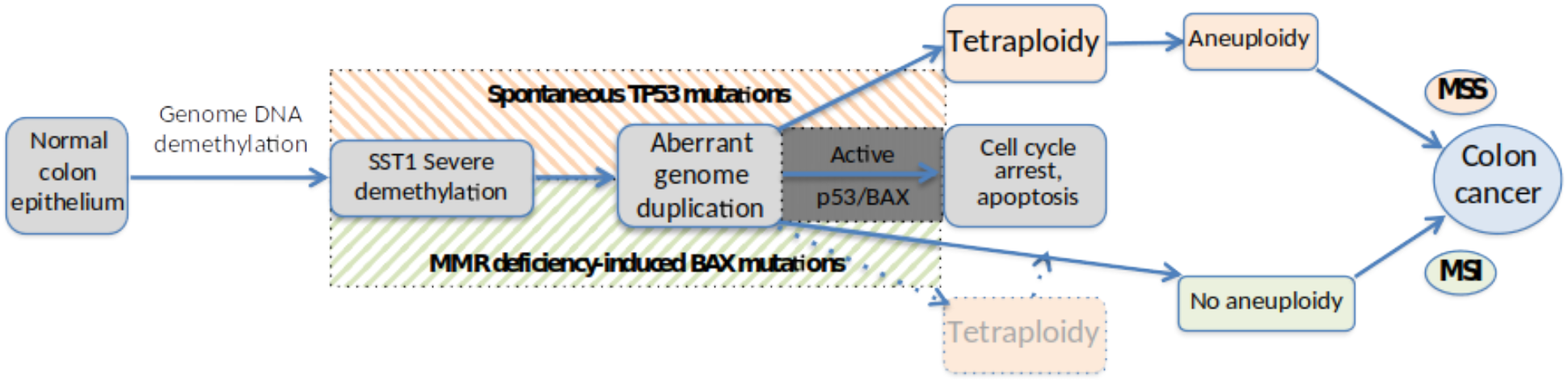
Model for a link between SST1 somatic demethylation, tetraploidy and TP53 and BAX mutations in CRC. In a genomic background of somatic DNA demethylation, severe demethylation of SST1 pericentromeric elements leads to genome duplication and consequent tetraploidy in CRC precursor cells. When cell cycle arrest or apoptosis networks are functional, the resultant tetraploid cells do not proliferate. When TP53 mutations in MSS CRC spontaneously occur before or during the presumably gradual demethylation of these (or other) sequences, the tetraploid cells survive initiating a pathway for aneuploidy and eventually CRC. In a less frequent MSI pathway for CRC, the tetraploid cells can evade apoptosis if there are somatic mutations in BAX and/or other pro-apoptotic proteins. In MSI CRC the SST1 demethylation-associated tetraploidy is not essential for tumor progression, and the MSI mutator phenotype may lead to a pseudo-diploid tumor.

In this model, SST1 severe demethylation takes place in a globally demethylated genome, resulting in an increased probability of spontaneous tetraploidization that in turn triggers p38-p53-p21 mediated cell cycle arrest. Therefore, cells with SST1 severe demethylation can only thrive if the p38-p53-p21 pathway is previously inactivated either by mutations in *TP53* or by inactivation of essential downstream effectors such as *BAX* [45,55].

*We speculate that TP53* mutations are spontaneous, occurring during tumorigenesis, before or during the gradual demethylation events that eventually lead to the drastic demethylation triggering tetraploidization, because they would be under a strong selective pressure during tumor clonal evolution. In contrast, inactivating frameshift mutations of *BAX* and other pro-apoptosis can be readily explained by the previous mutator phenotype of MSI tumors [56], before or simultaneously during tetraploidization.

The mechanisms underlying the drastic SST1 demethylation in some tumors, or why this demethylation may increase the risk for genome duplication, remain to be elucidated. The cases with severe SST1 somatic demethylation tend to also have very high levels of global genomic hypomethylation in primary CRCs (figure 4F). However, there were fundamental differences in the hypomethylation affecting SST1 and LINE-1 sequences: while SST1 methylation in tumor samples mainly reflected the methylation of their matching normal tissue, LINE-1 elements were generally hypomethylated somatically only in tumors (figure 4). Therefore, the events that lead to drastic demethylation in SST1 elements, while may occur in a genetic background of global genome demethylation, represent a different mechanism. We do not consider mutant p53 to directly underlie the drastic demethylation of SST1 elements because the spectrum of *TP53* mutations in the cases with severe demethylation is not atypical, with common single point mutations in typical codons (supplemental figure S12). In this context, we previously reported that the demethylation of SST1 was associated with downregulation of *HELLS*, a protein that has been denominated the epigenetic guardian of the repetitive elements [21].

Despite the abundant evidence on the involvement of demethylation of the heavily methylated pericentromeric repeat elements and an increased risk for undergoing mitotic errors, the precise underlying mechanisms remain elusive [13]. In our case there is an added uncertainty on how to explain that demethylation of SST1 elements, that are localized in only some acrocentric chromosomes may lead to tetraploidization. The repetitive nature of SST1 elements makes it difficult to determine the precise chromosomal location of those elements that undergo demethylation in cancer. We cannot rule out that demethylation of some of these elements, but no others, might have different phenotypic effects. Therefore, it is possible that the drastic demethylation of SST elements is not *per se* the direct cause of tetraploidization, but a surrogate diagnostic marker for the underlying epigenetic defect that leads to the genetic alteration. We recently reported that demethylation of SST1 precedes the expression of a long non-coding RNA that forms a perinucleolar aggregate with RNA binding proteins [22]. However, we found no association between the expression of this lncRNA and the demethylation-associated tetraploidization in LS174T-derived clones (data not shown). The down-regulation of *HDLBP* revealed by transcriptomic analyses provides a role-playing candidate for tetraploidization.

We have not found definitive evidence linking demethylation and tetraploidization in primary CRC. To test whether the tumors positive for SST1 severe demethylation possess putative tetraploid karyotypes, or have had a past tetraploid stage, remains a task ahead. Preliminary analysis by CGH arrays of a subset of these tumors did not allow reaching solid conclusions. Unfortunately, we cannot compare the data from our cohort with that of larger more standard cohorts, for instance the TCGA COAD and READ datasets, because those samples have not been analyzed for SST1 methylation, and the HM450K methylation arrays that were employed to profile methylation on those samples do not cover these repetitive elements. Thus, the validation of our results has to be done by analyzing additional fresh samples. SNP array or point mutation based methods that can infer tetraploid lineage in mixed cell populations may return definitive evidence [39,59].

Despite these limitations, our report provides novel clues relative to the role of somatic DNA demethylation in human cancer. The causal relationship between DNA demethylation of SST1 pericentromeric repetitive elements and chromosomal instability is supported by its association with mutational inactivation of p53 or pro-apoptotic proteins in a subset of human primary CRC. The correlation between global demethylation and copy number alterations has been reported by us and many other researchers [13,17,60]. There is also solid mechanistic evidence for the oncogenic effect of genome-wide demethylation in CRC cell lines as well as in mice in which the DNA methyltransferases (DNMTs) have been genetically disrupted [9,11,61]. Lastly, the association between LINE-1 hypomethylation and hyperploidy has been described in ovarian cancers [62]. However, to our knowledge, we present the first evidence linking naturally occurring DNA demethylation in human CRC cells with the onset of tetraploidy, which is likely involved in the early steps of tumorigenesis [30,63].

## 4. Materials and Methods

### Cell lines and tumor samples

Colorectal cancer cell lines LS174T (CL-188), HCT116 (CCL-247), DLD-1 (CCL-221) and ovarian cell line OV-90 (CRL-11732) were obtained from the American Type Culture Collection (ATCC). Primary CRCs with matched normal tissue from 148 patients were obtained as frozen specimens from the Cooperative Human Tissue Network (CHTN, https://www.chtn.org/). Informed consent was obtained and managed by the CHTN at the time of sample collection, following US regulations and the guidelines of the Declaration of Helsinki [64]. Approval to collect samples, extract DNA and perform analyses was obtained from the Institutional Review Board (IRB) of the Sanford-Burnham-Prebys (SBP) Medical Discovery Institute. Additional approval was obtained from the IRB of the IGTP to further analyze DNA samples of the study.

The identity of all the cells used in this work as well as their derived clones, was validated by STR analysis using AmpFLSTR^™^ Identifiler^™^ Plus PCR Amplification Kit (Cat n:4486467, ThermoFisher Scientific) using the provided protocol. Sequencing results were analyzed using GeneMapper ID v3.2 (Applied Biosystems) and analyzed with Expasy Cellosaurus tool CLASTR 1.4.4 [65]. Data is provided in supplemental materials. All cell lines were confirmed to be mycoplasma-free by PCR analysis, prior to the initiation of the experiments and then regularly during the experimental timeframe.

### Cell culture conditions

LS174T, HCT116 and DLD-1 cell lines were cultured in DMEM:F12 medium supplemented with 10% fetal bovine serum (FBS), 2mM L-glutamine, 1mM sodium pyruvate (NaPyr) and Antibiotic-Antimycotic. Ovarian cell line OV-90 was cultured in ATCC recommended media, 1:1 MCDB 105 medium containing a final concentration of 1.5 g/L sodium bicarbonate and Medium 199 containing a final concentration of 2.2 g/L sodium bicarbonate supplemented with 15% FBS and antibiotic and antimycotic. All cells were cultured in ø100 mm culture plates, or in 24 well plates with coverslips, in a 37 °C incubator with 5% CO_2_. Cells were grown until reaching 80–90% confluency before collection or passage.

### Isolation of single cell-derived LS174T cell clones

LS174T and OV-90 cell line subcloning was performed using the limiting dilution method [66], as we have already performed in previous works for the isolation of spontaneous frameshift mutations in MMR genes in CRC cells with microsatellite instability (MSI) and study their phenotype consequences [54]. Cells were counted after the addition of Trypan Blue 0.4% 1:1, and diluted in complete medium to a concentration of 5 cells/ml (0.5 cells/100µl). 100µl of this dilution were seeded into each well of 96 plates, and incubated at 37ºC. Cell growth was monitored by microscopy. The occasional presence of more than a single growth focus during culture, was used as indication of non-clonality, and those cultures were discarded.

### Nuclei size measurement

Cell morphology was studied by fluorescence microscopy after phalloidin and DAPI staining (Phalloidin-iFluor 594 Reagent, ab176757 - Abcam). Cells were cultured in 24-well plates with glass coverslips inside until they reached 70-80% confluence. Then, the culture medium was carefully removed and washed once with PBS. Cells were fixed by incubation with 3-4% formaldehyde in PBS at room temperature for 20 minutes. The fixation solution was aspirated and fixed cells were washed 3 times with PBS. To increase permeability, cells were incubated in PBS + 0.1% Triton X-100 for 5 minutes and washed with PBS 3 times. Next, 100µl of 1x Phalloidin conjugate (1µl Phalloidin-iFluor 594 1000x in 1ml PBS + 1% BSA) was added to each coverslip. They were incubated at room temperature for 60 minutes, protected from light. Finally, 100µl of PBS + 1µg/ml DAPI (Cat N.: D1306 - Thermo Fisher Scientific) were added to stain and visualize the nuclei. Lastly, 3 washes with PBS were performed to remove DAPI excess. The samples were dehydrated by immersion in 95% ethanol during 15 seconds, and left to dry for 10 minutes in total darkness and mounted with ProLong Gold antifade reagent (Invitrogen). Samples were observed and photographed by fluorescence microscopy (LEICA DM1600B Microscope - LAS X Software). Fluorescence microscopy images (200x) were analyzed with ImageJ Software [67]. The area of at least 200 nuclei was quantified per sample.

### Karyotyping

Karyotypes were prepared from mitotic cells arrested in metaphase using colcemid, lysed in hypotonic solution (KCl 0.075M), fixed with Carnoy’s solution (methanol:glacial acetic acid, 3:1) and stained according to the G-banding pattern, following standard procedures. The analyses were done with cells of the parental cell lines and derived single cell clones at different time points of the culture after single cell clone isolation. Karyotypes with tetraploidy were true tetraploid and not mixtures of two diploid cells, as determined by the mitotic patterns under microscopy, as well as the sparse cell densities in the cultures treated with colcemid for mitotic arrest.

### DNA and RNA extraction

Genomic DNA from cell lines was isolated using Maxwell^®^ 16 Instrument and the Maxwell^®^ 16 DNA Purification Kit (Promega Cat.# AS1020) following the protocol provided. DNA concentration was measured by NanoDrop™-1000 UV/Vis Spectrophotometer (Thermo Fisher Scientific) and visualized in an agarose gel. Total RNA from cell lines was isolated using Maxwell® 16 Instrument and the Maxwell® 16 LEV simplyRNA Cells Kit (Promega Cat.# AS1270) using the protocol provided. RNA quality control and integrity was measured using Agilent 2100 Bioanalyzer (Agilent Technologies, Santa Clara, USA). All samples had an RNA Integrity Number (RIN) above 9.

### LINE-1 MS-QPCR

LINE-1 MS-QPCR is a technique developed to estimate the ratio between unmethylated vs methylated LINE-1 elements. This technique employs two sets of primers, one to amplify methylated (LINE1-mF and LINE1-mR) and the other unmethylated (LINE1-uF and LINE1-uR) LINE-1 molecules. Primer sequences are detailed in supplemental figure S13. Two separate reactions were performed in duplicate for every sample. Conditions per reactions were: 5µl of Roche MasterMix SYBR-Green 2X, 100ng of bisulfite-transformed DNA, primers at 0.4µM each, 10µl final volume, 1 cycle of 95ºC 10’ followed by 40 cycles of 95ºC 10’’, 60ºC 10’’ and 72ºC 12’’, 1 cycle of 95ºC 5’’, 60ºC 1’ and 97ºC. Quantification was made with LightCycler® 480 Software, using a relative quantification program with standard settings and calculating Cts by double derivative method. All measures were normalized vs. genomic DNA from DLD-1 cells. LINE-1 MS-QPCR provide the relative demethylation level of LINE-1 elements (LINE-1 RDL), i.e. an estimation of the ratio of demethylated vs methylated LINE-1 sequences in the studied sample relative to that of DLD-1 genomic DNA. Therefore, higher RDL indicates higher *demethylation*, i.e. lower level of methylation. For the statistical analysis, LINE-1 RDL was log2 transformed to reduce the skewness of the distribution.

Albeit very similar in concept to LINE-1 MethyLight [16], LINE-1 MS-QPCR provides the ratio of unmethylated vs. methylated LINE-1 molecules from the same sample without relying on the amplification of any other type of repetitive elements, such as Alu elements. Thus, there is no effect of possible differences in Alu vs. LINE-1 abundance among samples due to germline or somatic copy number differences. Moreover, while LINE-1 MethyLight employs specific primers and internal probes for the methylated and unmethylated PCR reactions to increase specificity, LINE-1 MS-QPCR employs just specific primers but no internal fluorescent probes. We purposely selected this approach to increase the range of detected molecules in the unmethylated and in the methylated reactions. We estimated the specificity of the amplification by sequencing clones from the LINE-1 methylated reaction and from the LINE-1 unmethylated reaction (supplemental figure S13). From the methylated LINE-1 reaction, the proportion of methylated CpG sites ranged from 58% and 100% (median value 91%). Nine out of 10 clones from the unmethylated reaction showed complete demethylation, and one 22% methylation (2 CpG sites methylated and 7 unmethylated, on a sequence that exhibited 4 polymorphisms in the 13 interrogated CpG sites), for a median value of 0% methylation. Thus, LINE-1 MS-QPCR is not restricted to fully unmethylated or methylated sequences but instead provides the ratio of sequences with 0%∼22% methylation (unmethylated sequences) vs 60%-100% methylation (methylated sequences).

### SST1 MS-QPCR

SST1 MS-QPCR is similar to the LINE-1 MS-QPCR technique described above, but in this case providing the ratio between unmethylated SST1 vs amplifiable methylation-independent SST1 sequences by using two sets of primers, one specific for unmethylated SST1 elements and the other amplifying SST1 elements irrespective of their methylation status (primer sequences are detailed in figure S9). The primers for the methylation-independent SST1 sequences were SST1-F and SST1-R, that we employed for bisulfite sequencing in a previous publication [18]. These primers amplify a 317bp internal region of the consensus SST1 sequence containing 27-28 CpG sites, some of them polymorphic, and including the *Not*I site that we originally found hypomethylated by MS-AFLP DNA fingerprinting [17]. Analyzing our bisulfite sequencing data from LS174T cells, we determined the CpG sites that better estimated the overall methylation level of this region (supplemental figure S9). We designed the primer SST1-U-F to amplify SST1 sequences with unmethylated CpG3 (avoiding both CpG4, with low correlation with overall methylation, and CpG5, highly polymorphic), and primer SST1-U-R to amplify unmethylated CpG17 and CpG18 that exhibited the highest correlation with the overall methylation value (r=0.77 and r=0.79, respectively, supplemental figure S9). Specificity of SST1-U-F and SST1-U-R primers was estimated by bisulfite sequencing (supplemental figure S9). QPCR conditions and calculation of SST1 RDL were identical to LINE1 MS-QPCR, also using DLD-1 genomic DNA as reference sample (see above). For the statistical analysis, SST1 RDL was log2 transformed to reduce the skewness of the distribution.

SST1 MS-QPCR exhibited strong concordance with our previous results using bisulfite sequencing (correlation=0.73, CI95%=[0.54,0.85], Pearson’s test p=7.9×10^−8^, supplemental figure S9). Considering bisulfite sequencing (originally used to classify cases into no-change / demethylated or severely demethylated [18]) as gold standard, MS-QPCR was very accurate to detect severely demethylated samples (max accuracy = 0.96, AUC=0.79, *p*=0.01). Samples with moderate changes in SST1 methylation were not detected by MS-QPCR, very likely because they do not have very low methylated SST1 molecules that are targeted by SST1-U-F and SST1-U-R primers.

### Bisulfite sequencing

Genomic DNA was treated with sodium bisulfite using EZ DNA Methylation Kit (Zymo Research, Orange, Cat.#D5001). Bisulfite treated samples were used as a template for PCR and QPCR amplification of SST1/LINE-1 sequences.

A SST1-specific PCR was designed to cover a total 317bp, including 27-28 CpGs including the NotI site of the band previously identified by MS-AFLP. The PCR Master Mix contained: 1x Buffer, 0.125nM dNTPs, 0.25 units of Taq DNA Polymerase (Roche) and 0.4mM of primers SST1-F and SST1-R (supplemental figure S9). The PCR program consisted of 1 cycle at 95ºC for 5 min, followed by 35 cycles of denaturation at 95ºC for 15 s, annealing at 55ºC for 30 s, and extension at 72ºC for 60 s, ending at 72ºC for 5 min to complete extension.

For bisulfite sequencing, 1 μL of SST1-F/R PCR product was cloned into pSC-A-amp/kan vector using Strataclone™ PCR Cloning Kit (Agilent, Cat.#240205) following manufacturer’s instructions and transformed into *E. coli* Statataclone SoloPack competent cells. Transformed cells were selected onto LB plates containing Ampicillin (50 μg/mL) and X-Gal (40 μg/mL). To identify bacterial colonies harboring the vector with the cloned PCR product, colony PCRs were performed using universal primers T3 and T7 [68]. Amplicon size was analyzed by gel electrophoresis and those reactions that rendered a correct size fragment were sequenced (GATC Services, Eurofins Genomics) and analyzed with CLC Main Workbench 5.6.1 program.

### Illumina HM450K methylation arrays

Genomic DNA from tumors and matching normal tissues from 30 CRC patients were previously analyzed using Illumina HM450K arrays, following standard procedures [69]. The generated idat files were preprocessed and filtered using RnBeads [70], discarding data from X and Y chromosomes. Type-I vs Type-II probe bias was corrected using the BMIQ method from the wateRmelon package [71].

### Gene expression arrays and gene set enrichment analysis

RNA was extracted from cultured cells using Maxwell® RSC simplyRNA Cells Kit (Promega) on a Maxwell AS2000 (Promega) equipment. Quantity and quality of RNA was verified on an Agilent 2100 Bioanalyzer. cDNA synthesis, labeling and hybridization on Clariom S Arrays from Thermo Fisher Scientific was performed by the Genomics Unit of Josep Carreras Leukaemia Research Institute. Raw data files were preprocessed using RMA (oligo R package). Gene set enrichment analysis was performed using GSEA, comparing with MSigDB hall mark gene set [52]. Full input dataset and output html, specifying the running parameters, are provided as supplemental material.

### Statistical analyses

All statistical analyses were performed using R [72]. Normality was tested using Shapiro’s test. Variables normally distributed were analyzed using parametric tests (e.g. t-test, ANOVA). Variables not normally distributed were analyzed using Wilcox’s exact test. When appropriate, multiple hypothesis correction was performed using the false discovery rate (FDR) method implemented in *p*.*adjust* function. Comparison of coefficient of variation (*CV*=σ/μ) was performed using the asymptotic Feltz-Miller’s test [73].

## 4. Conclusions

LS174T cells undergo spontaneous tetraploidization *in vitro*, associated with demethylation of SST1 repetitive elements. To the best of our knowledge, this is the first evidence linking naturally occurring DNA demethylation in human CRC cells with the onset of tetraploidy, which is likely involved in the early steps of tumorigenesis

SST1 methylation in primary tumors reflects, in general, the methylation found in their matched normal tissues. SST1 demethylation occurs in ∼17% of colorectal tumors (12% with moderate demethylation and 5% with severe demethylation), associated with mutations in *TP53*.

## Supporting information

STR analysis of cells used in the study

GSEA results

## List of abbreviations

CRC: colorectal cancer
LINE-1: long insterspected nuclear element 1
MSI: microsatellite instability
MSS: microsatellite stability
CIN: chromosomal instability
MMR: DNA mismatch repair
CGH: comparative genome hybridization
STR: short tandem repeat
MS-QPCR: methylation sensitive quantitative PCR
DAPI: 4′,6-diamidino-2-phenylindole
FCM: flow cytometry
CV: coefficient of variation (CV=σ/μ, where σ is the standard deviation and μ the mean)
RDL: relative demethylation ratio, i.e. the ratio between unmethylated vs methylated molecules relative to a normalizing sample
ΔRDL: differential relative demethylation ratio, also termed somatic demethylation, i.e. the difference of RDL between tumor and matching normal samples.

## 5. Supplemental information

We provide 5 files of supplemental information

| File name | Content |
| --- | --- |
| STR | zip-compressed file with the STR analyses of all the cell lines and derived clones analyzed in this study |
| Supplemental_Figures | pdf file with supplemental figures and methods |
| Supplemental_table_1 | zip-compressed tsv file with the clinical and molecular data of the 148 CRCs analyzed in this study |
| Supplemental_table_2 | zip-compressed tsv file with the normalized and log2 transformed gene expression values of tetraploid vs diploid LS174T clones, measured with Affymetrix Clariom™ S arrays. It also includes a chip file with the probe to gene information, required for the GSEA analysis. |
| GSEA | zip-compressed results of the GSEA analysis output, in HTML |

## 6. Acknowledgments

We thank Dr. M.A. Fernández Sanmartín and G. Requena Fernández from the IGTP cytometry unit (http://www.germanstrias.org/technology-services/cytometry/) for their support and advice on the FCM experiments. Microarrays studies were performed in the Microarrays Unit of Josep Carreras Leukaemia Research Institute. We are indebted to all members of Cytogenetics Platform, Hematology Department at Germans Trias i Pujol Hospital, Catalan Institute of Oncology, Josep Carreras Leukaemia Research Institute (Badalona, Spain) for providing the karyotype studies and images. Tissue samples were provided by the Cooperative Human Tissue Network (CHTN), which is funded by the National Cancer Institute. Other investigators may have received specimens from the same subjects. This work has been supported by a grant to MP and SA from the Instituto de Salud Carlos III (PI18/01484), Spanish Ministry of Science, Innovation and Universities, FEDER.

## 7. Author Contributions

SJ and SA set up the LINE-1 MS-QPCR technique, and applied it to analyze primary colorectal cancers. BG and MNJ set up the SST1 MS-QPCR technique, generated the LS174T clones and performed all the analyses in LS174T clones and primary colorectal tumors. MJAdG and BG performed the analysis of the ovarian cancer cells. SA performed the statistical analyses, and the transcriptome and HM450K analyses. IG led the karyotyping analysis. SA, MP and BG designed the experimental strategy. SA and BG supervised the work. BG, MNJ, MP and SA interpreted the results and wrote the manuscript.

