## Supplemental Figures for "Somatic hypomethylation of pericentromeric SST1 repeats and tetraploidization in human colorectal cancer cells"

### Supplemental Material


**1. LS174T nuclei size distribution** **2**

**2. Genomic content of LS174T and derived clones measured by flow cytometry.** **2**

**3. Growth rate of LS174T and derived clones.** **4**

**4. Modeling the tetraploidization of LS174T cells.** **5**

**5. SST1 methylation and nuclei size in CRC MSI cell lines.** **6**

**6. Karyotype of OV-90 ovarian cancer cells.** **7**

**7. Transcriptional profiling of near-diploid and near-tetraploid LS174T clones.** **8**

**8. Gene Set Enrichment Analysis of the transcriptional profile of near-tetraploid LS174T clones.** **9**

**9. SST1 MS-QPCR vs Bisulfite sequencing** **10**

**10. SST1 and LINE-1 ΔRDL vs global methylation levels.** **12**

**11. Association of SST1 ΔRDL with clinicopathological and mutational characteristics of CRCs.** **13**

**12. TP53 mutations in SST1 severely demethylated cases.** **14**

**13. LINE-1 MS-QPCR.** **15**

#### 1. LS174T nuclei size distribution

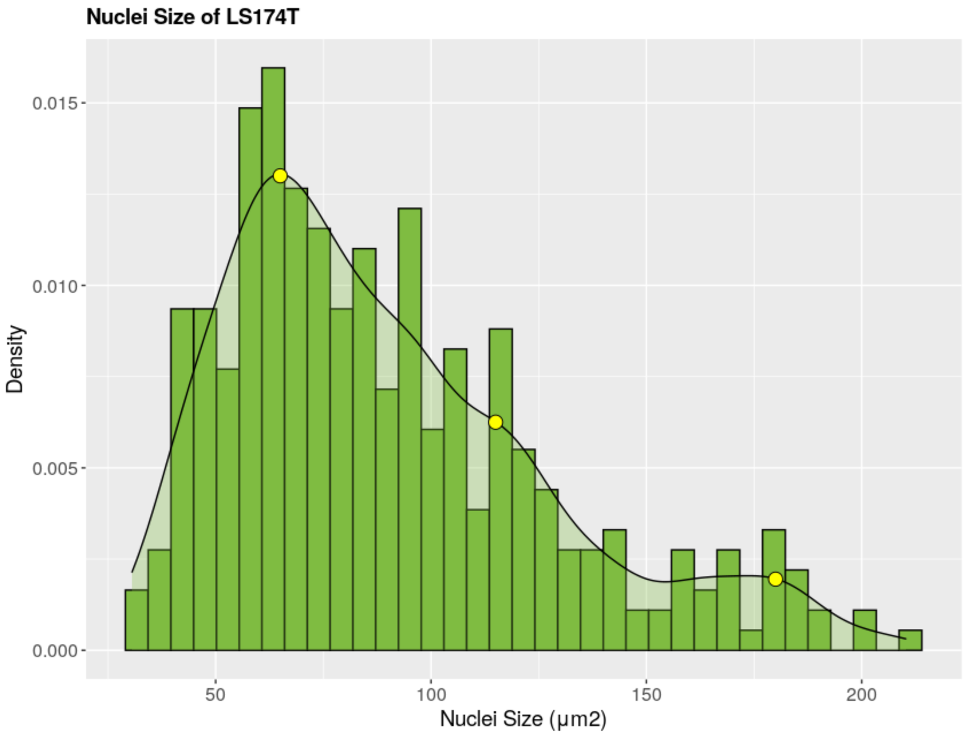


**Supplemental figure S1.** Histogram and density plot of LS174T nuclei size distribution, measured on DAPI-stained fixed cells. The three modal points at approximately 65, 115 and 180 µm$^2$ are indicated in yellow.

#### 2. Genomic content of LS174T and derived clones measured by flow cytometry.

Cell cycle and total DNA content were analyzed by flow cytometry (FCM) of cells stained with DAPI [74]. We collected cells at 70-80% confluence by trypsin-EDTA treatment. Cell density was measured using a Countess Automated Cell Counter apparatus from Invitrogen. $10^6$ to $10^7$ cells were washed with PBS and subsequently collected by centrifugation at 1200rpm for 5 min. After removing PBS, the cell pellet was resuspended in 0.5ml 1x PBS. Cells were then transferred to tubes with 4.5ml of 70% ethanol at 4ºC for at least 2 hours. For DAPI staining, fixed cells were first centrifuged for 5 minutes at 1200 rpm and the remaining ethanol was decanted. The pellet was resuspended in 5ml 1x PBS to wash the cells. The cell pellet was resuspended in 1ml of 1µg/ml DAPI/Triton X-100 solution and kept 30 minutes in total darkness at room temperature. Samples were analyzed using Flow Cytometer FACSCanto II (BD Biosciences) and the results were analyzed with FlowJo v10 software. Results are shown in supplemental figure S2.

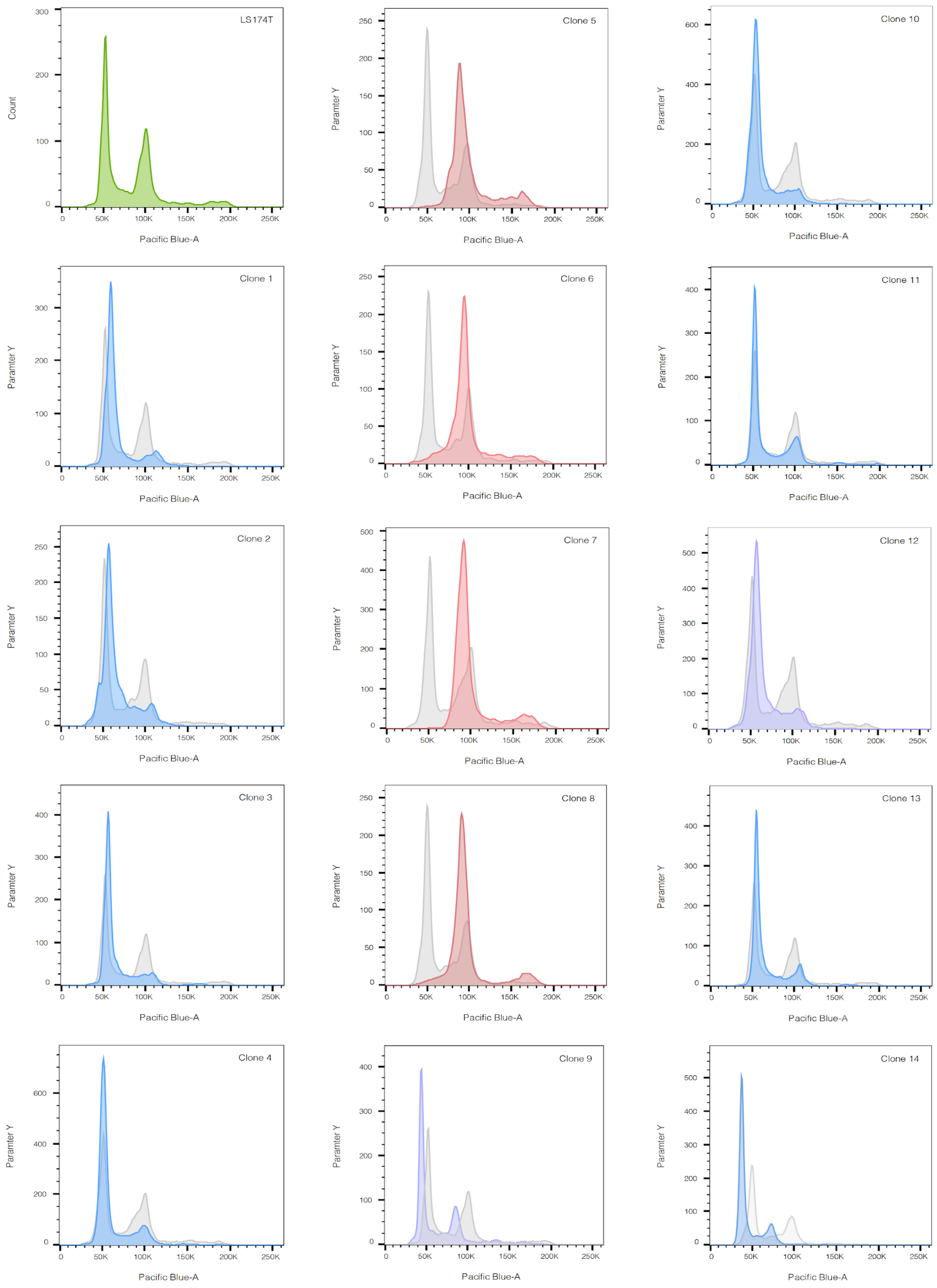


**Supplemental figure S2.** Cell cycle analysis by flow cytometry of cells stained with DAPI. In green, LS174T parental cell line used as a reference in all panels (gray). In blue, the near-diploid clones confirmed by karyotyping. In red, the near-tetraploid clones confirmed by karyotyping. In purple, clone 9 and 12 which presented a mixture of near-diploid and near-tetraploid cells.

#### 3. Growth rate of LS174T and derived clones.

Cell proliferation was measured using a colorimetric test based on the capability of metabolic active cells to cleavage the tetrazolium salt (XTT) to form a soluble formazan dye tetrazolium salt (Cell Proliferation Kit II XTT, Roche), with different absorption wavelength. Briefly, the cells were plated in 96-well microtiter plates at a density of 3000 cells in 100 µl of Dulbecco's modified Eagle's medium (DMEM:F12) medium with 10% FBS, 2mM L-glutamine, 1mM sNaPyr and Antibiotic-Antimycotic and cultured at 37ºC for 0, 12, 24, 48, 72 h and 96h in a humidified atmosphere of 5 % CO2. Then, 50 µl of XTT was added to each well and incubated for up to 4 h at 37°C in the presence of 5 % CO2. After the incubation with XTT, optical density was measured at 492 and 690 nm using a plate reader. The amount of metabolic active cells was estimated by subtracting the OD690nm value to the OD492nm value, as indicated by the kit manufacturer.

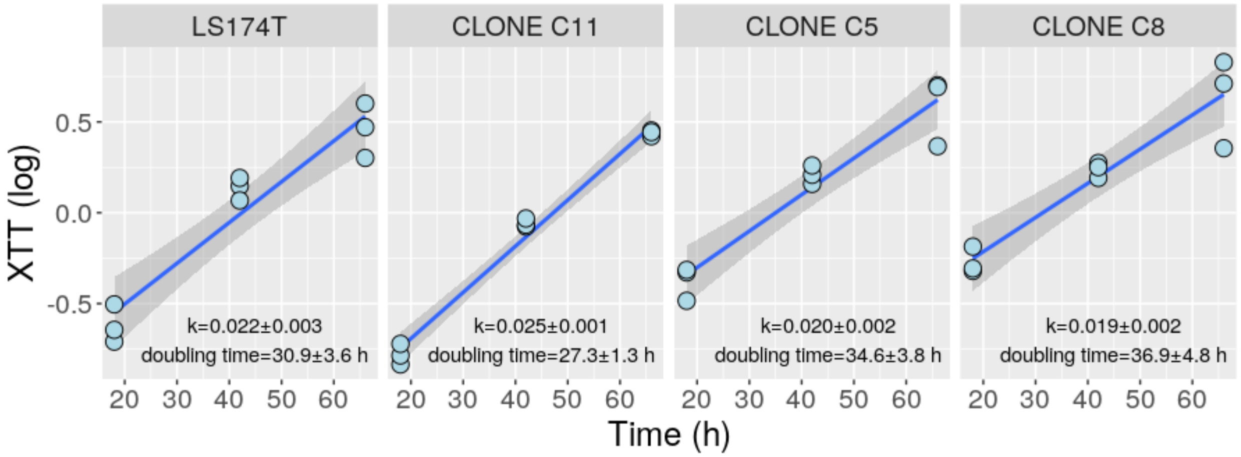


**Supplemental figure S3.** Proliferation of LS174T cells and its derived near-diploid (C11) or near-tetraploid (C5 and C8) subclones, under our in vitro cell culture conditions. Cell growth was monitored using XTT method (see above) at times 18, 42 and 66h. Every experiment included 3 independent internal replicas. XTT reaction was measured after 4h of incubation. Growth was modeled according to the exponential growth equation ($N_t=N_0 \cdot e^{kt}$). Exponential growth rate (k) and doubling time are shown. The doubling time ranged between 27 and 37h, very similar to that previously reported of 30-40h, according to DSMZ: https://www.dsmz.de/collection/catalogue/details/culture/ACC-759.

**4. Modeling the tetraploidization of LS174T cells.**

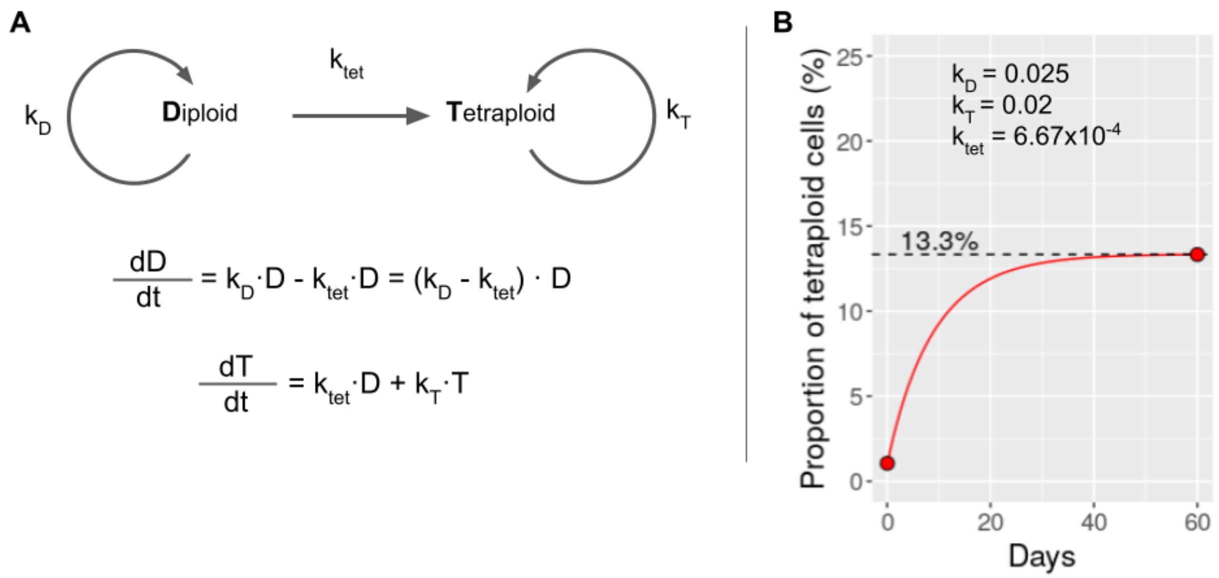


**Supplemental figure S4. A)** Mathematical model of LS174T cells spontaneous tetraploidization. In this model, diploid cells duplicate at a rate of $k_D$ and undergo spontaneous tetraploidization at a rate of $k_{tet}$, generating tetraploid cells that divide at a rate of $k_T$. Thus, the increment in diploid cells by unit of time (dD/dt) is the number of newly generated diploid cells by regular cell division ($k_D \cdot D$) minus those that have undergone tetraploidization ($k_{tet} \cdot D$). The increment of tetraploid cells by unit of time (dT/dt) is the number of diploid cells undergoing tetraploidization ($k_{tet} \cdot D$) plus the number of tetraploid cells generated from other tetraploid cells by regular cell division ($k_T \cdot T$). Since tetraploid cells do not revert to diploid cells, the diploid cell population will asymptotically tend to disappear unless $k_D > k_T$. **B)** Proportion of tetraploid cells in LS174T clone C11 during 60 days of culture. Growth rate of diploid ($k_D$) and tetraploid cells ($k_T$) were estimated from the culture of LS174T clones C11 (diploid) and C5 (tetraploid) shown in supplemental figure S3. At time zero, the proportion of tetraploid cells in LS174T clone C11 was 1.05% (1/95). After sixty days of culture, the proportion of tetraploids increased to 13.3% (10/75) that, according to the mathematical model, would require a $k_{tet}$ of $6.67 \times 10^{-4}$ ($k_{tet} \simeq k_D \times 0.0267$, i.e. $k_{tet}$ is 2.67% of the mitotic rate).

#### 5. SST1 methylation and nuclei size in CRC MSI cell lines.

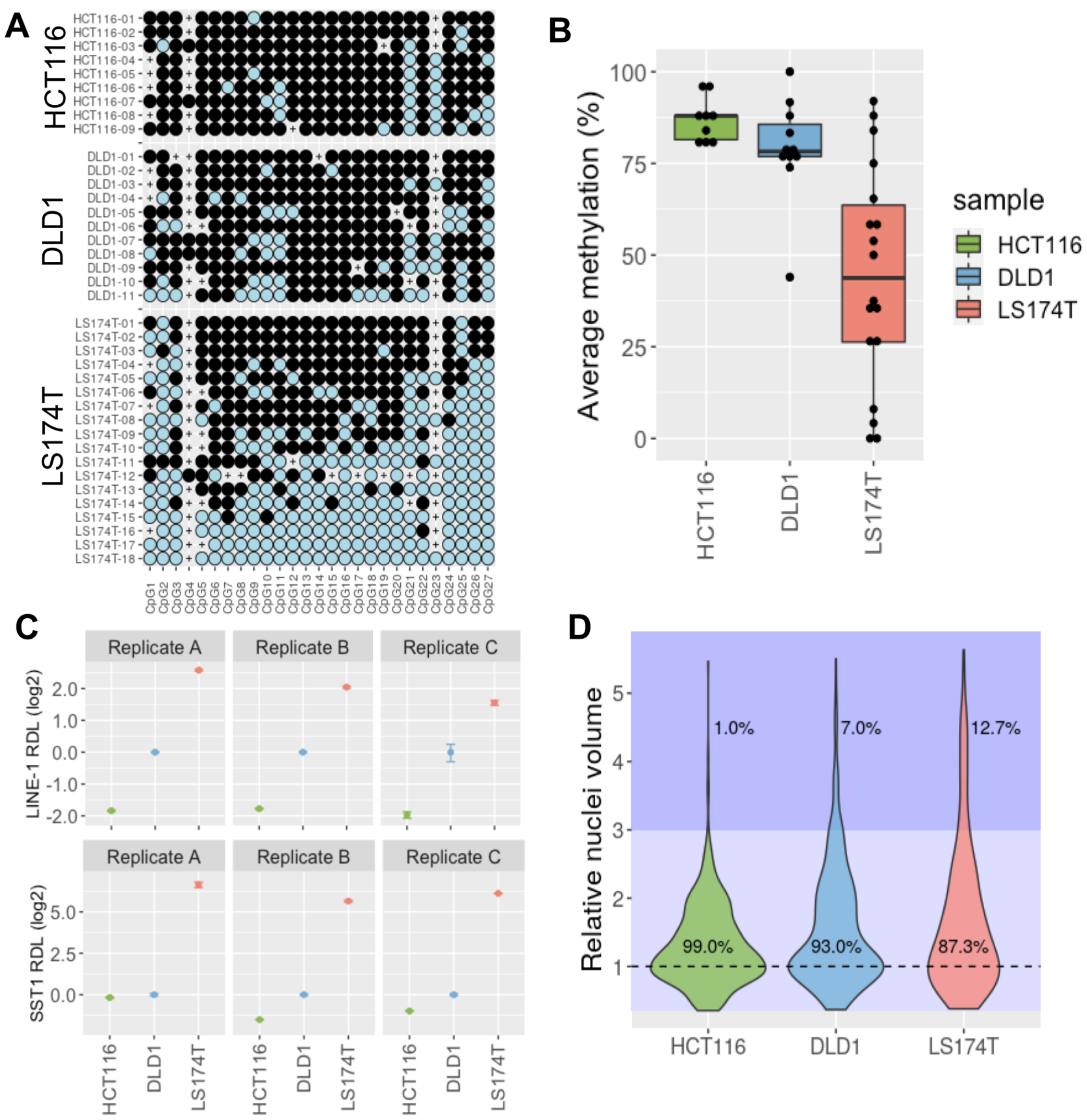


**Supplemental figure S5. A)** Bisulfite sequencing of SST1 in HCT116, DLD1 and LS174T. Every line represents an individual sequence containing 27 CpG sites. Unmethylated CpG sites in blue, and methylated CpG sites in black. Crosses indicate polymorphisms. **B)** Average methylation of individual SST1 sequences in HCT116, DLD1 and LS174T, calculated from the data shown in a. **C)** LINE-1 and SST1 RDL measured by MS-QPCR in three separate experiments (replicates A to C). Every experiment included two internal replicas for every reaction. **D)** Distribution of nuclei volume of HCT116, DLD1 and LS174T, normalized to their respective modal nuclei volumes (diploid cells in G0/1). The percentage of nuclei above (dark blue area) and below (light blue area) three times the modal volume is indicated.

**6. Karyotype of OV-90 ovarian cancer cells.**

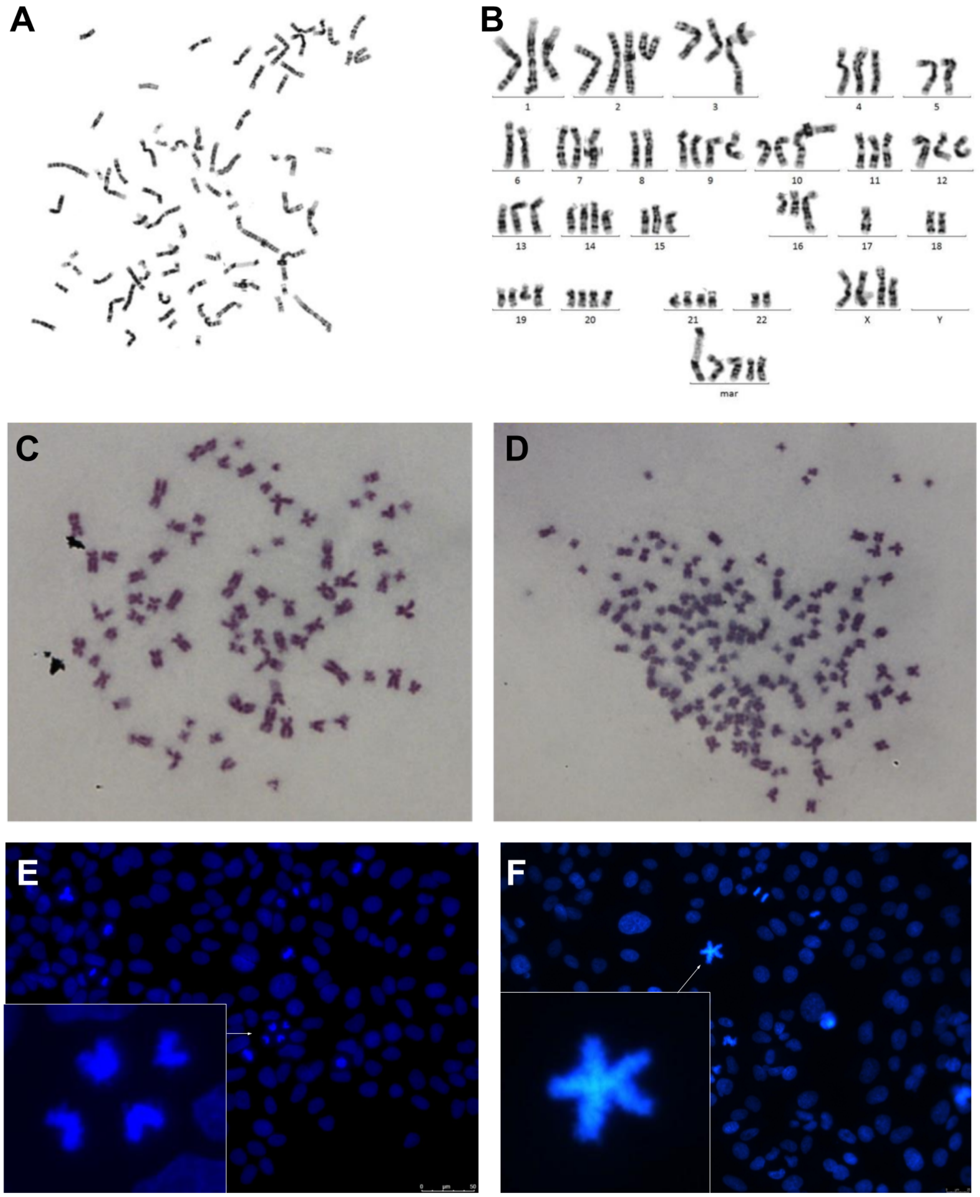


**Supplemental figure S6.** Top row, a modal hyper-triploid metaphase of OV-90 cells stained according to the G band pattern (**A**) and its karyotype (**B**). Middle row, hyper-triploid (**C**) and hypo-pentaploid (**D**) metaphases of OV-90 cells. Bottom row, examples of tetrapolar (**E**) and pentapolar (**F**) mitoses in OV-90 cells stained with DAPI (200x magnification).

#### 7. Transcriptional profiling of near-diploid and near-tetraploid LS174T clones.

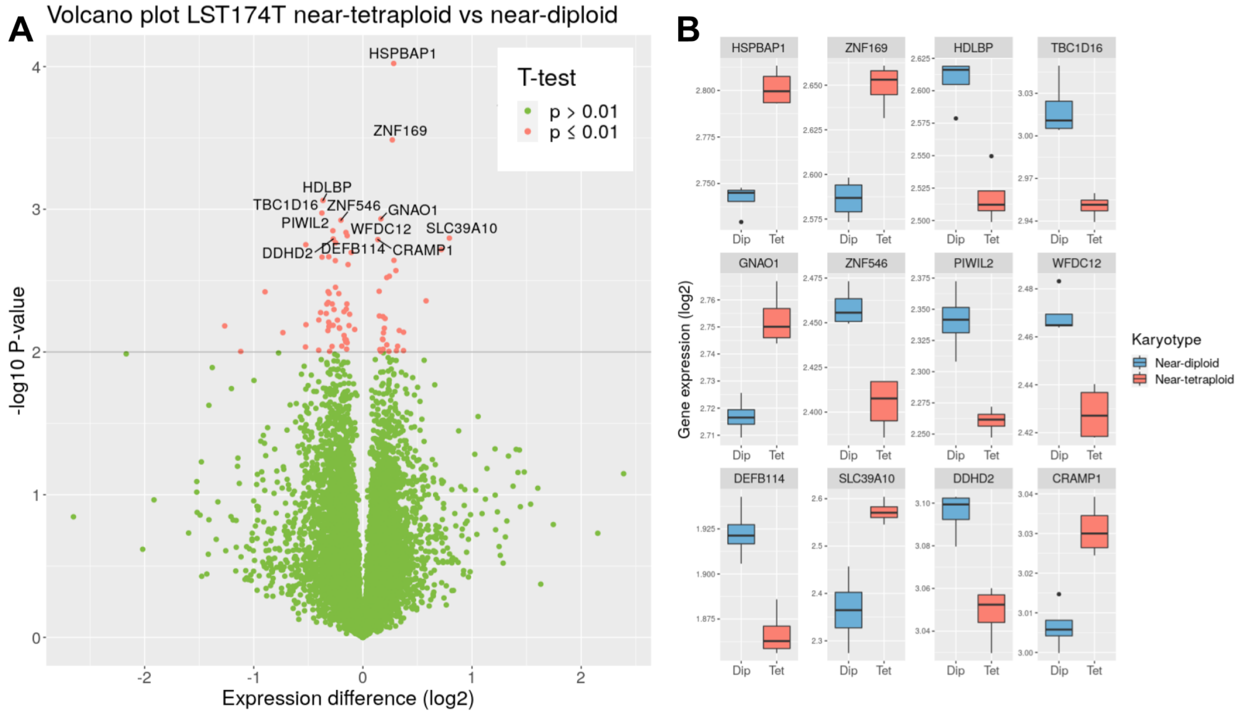


**Supplemental figure S7. A)** Volcano plot of the differential gene expression comparing near-tetraploid LS174T clones (C5-C8) vs near-diploid clones (C3, C10, C11, and C14). In green, genes that did not reach statistical significance (99.61%, n=21364). In red, genes that reached statistical significance (0.39%, n=84). The twelve genes whose differential expression exhibited the highest statistical significance are labeled. **B)** Expression of the 12 most statistically significant differentially expressed genes. In blue, expression level in the near-diploid clones. In red, expression level in near-tetraploid clones.

#### 8. Gene Set Enrichment Analysis of the transcriptional profile of near-tetraploid LS174T clones.

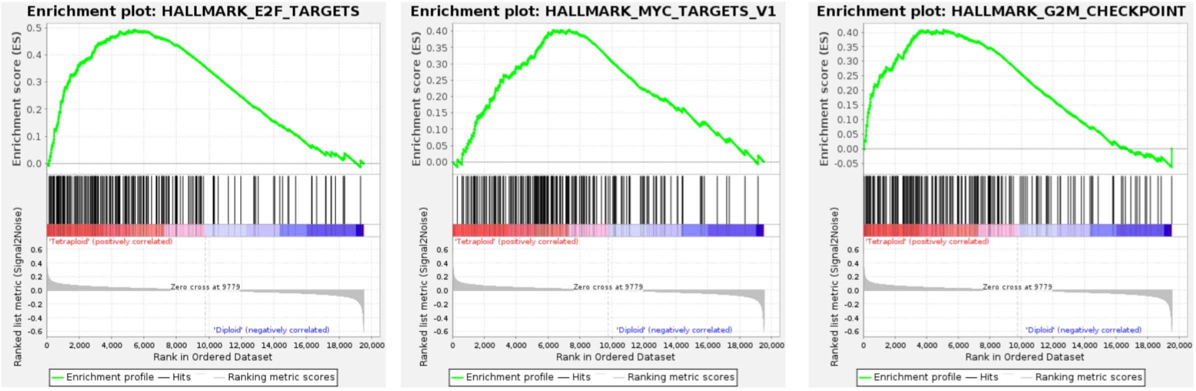


| NAME | SIZE | ES | NES | NOM p-val | FDR q-val | FWER p-val |
|---|---|---|---|---|---|---|
| HALLMARK_E2F_TARGETS | 195 | 0.49 | 2.20 | <0.0001 | <0.0001 | <0.0001 |
| HALLMARK_MYC_TARGETS_V2 | 57 | 0.56 | 2.05 | <0.0001 | 0.0007 | 0.0010 |
| HALLMARK_MYC_TARGETS_V1 | 194 | 0.40 | 1.80 | <0.0001 | 0.0021 | 0.0040 |
| HALLMARK_G2M_CHECKPOINT | 189 | 0.41 | 1.78 | <0.0001 | 0.0020 | 0.0050 |

**Supplemental figure S8.** Gene set enrichment analysis (GSEA) of genes differentially expressed in near-tetraploid vs near-diploid LS174T clones (see supplemental figure S7). On top, three most relevant Hallmark gene signatures over-expressed in near-tetraploid clones: E2F targets (genes encoding cell cycle related targets of E2F transcription factors), MYC targets (a subgroup of genes regulated by MYC) and G2M checkpoint (genes involved in the G2/M checkpoint, as in progression through the cell division cycle), defined by the Molecular Signatures Database [52]. Below, a summary table of these gene signatures indicating the size of the genes in every signature (size), the enrichment score (ES), the normalized enrichment score (NES), the nominal p-value (NOM p-value), the false discovery rate multi-hypothesis corrected p-value (FDR q-val) and the family-wise error corrected p-value (FWER p-val). Input data, analysis parameters and complete output are provided in supplemental file GSEA.

#### 9. SST1 MS-QPCR vs. Bisulfite sequencing

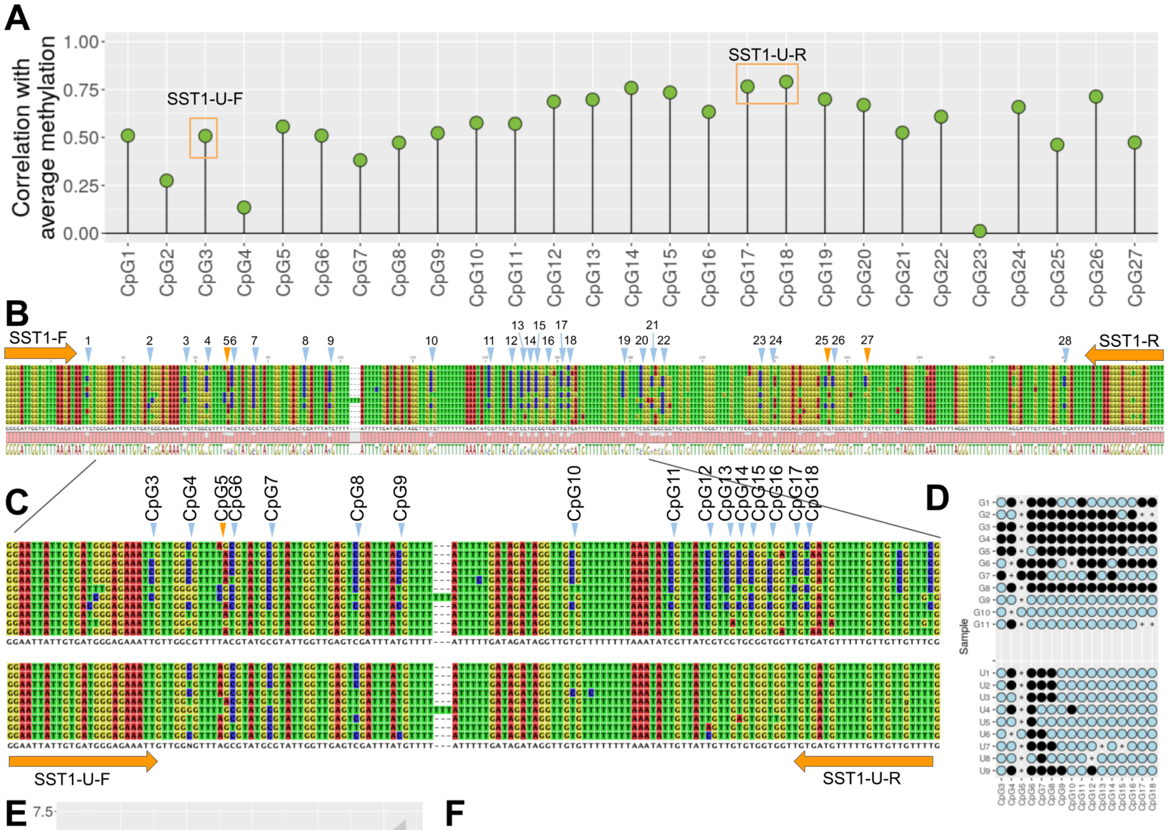


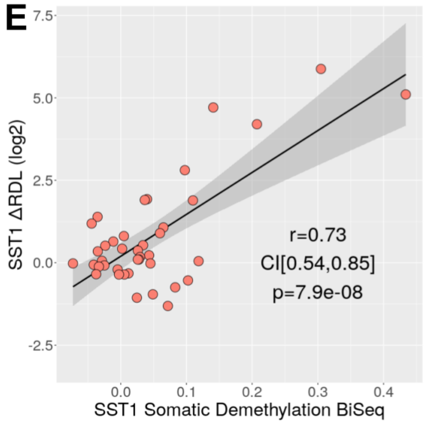


**F**

| Primer | Sequence | Reaction |
|---|---|---|
| SST1-F | 5'GGGGATTGGTGTTTAAGATA3' | Methylation-independent SST1 MS-QPCR |
| SST1-R | 5'AAAACTCCCCCTCCCTTAATAA3' | Methylation-independent SST1 MS-QPCR |
| SST1-U-F | 5'GGAATTATTGTGATGGGAGAAATTG3' | Unmethylated SST1 MS-QPCR |
| SST1-U-R | 5'CAAAACAACAACAAAAACATCACA3' | Unmethylated SST1 MS-QPCR |

**Supplemental figure S9. A)** Correlation of the methylation status of individual CpG sites within SST1 sequences, and the overall methylation of the studied region. The CpG sites interrogated by MS-QPCR primers SST1-U-F and SST1-U-R are indicated with orange boxes. **B)** Bisulfite sequencing of SST1 elements from LS174T using methylation-independent primers SST1-F and SST1-R. Adenines in red, cytosines in blue, guanines in yellow and thymines in red. CpG sites are indicated with blue triangles. Very polymorphic CpG sites are indicated with orange triangles. Methylated CpG sites appear as CG, while unmethylated CpG sites appear as TG. **C)** Bisulfite sequencing of an internal region of SST1 in LS174T using SST1-F and SST1-R primers (top) or the methylation specific SST1-U-F and SST1-U-R primers (bottom). **D)** Methylation plot of the results shown in panel C. In blue, unmethylated CpG sites. In black, methylated CpG sites. Crosses indicate polymorphisms (the sequence shows other than CG or TG) **E)** Correlation between SST1 somatic demethylation measured by bisulfite sequencing (x-axis) and log2 transformed SST1 ΔRDL measured by SST1 MS-QPCR in 40 CRC cases chosen to benchmark MS-QPCR technology. Every dot represents a CRC. In black, the regression line. In grey, the 95% CI of the slope. **F)** Primer sequences.

#### 10. SST1 and LINE-1 ΔRDL vs. global methylation levels.

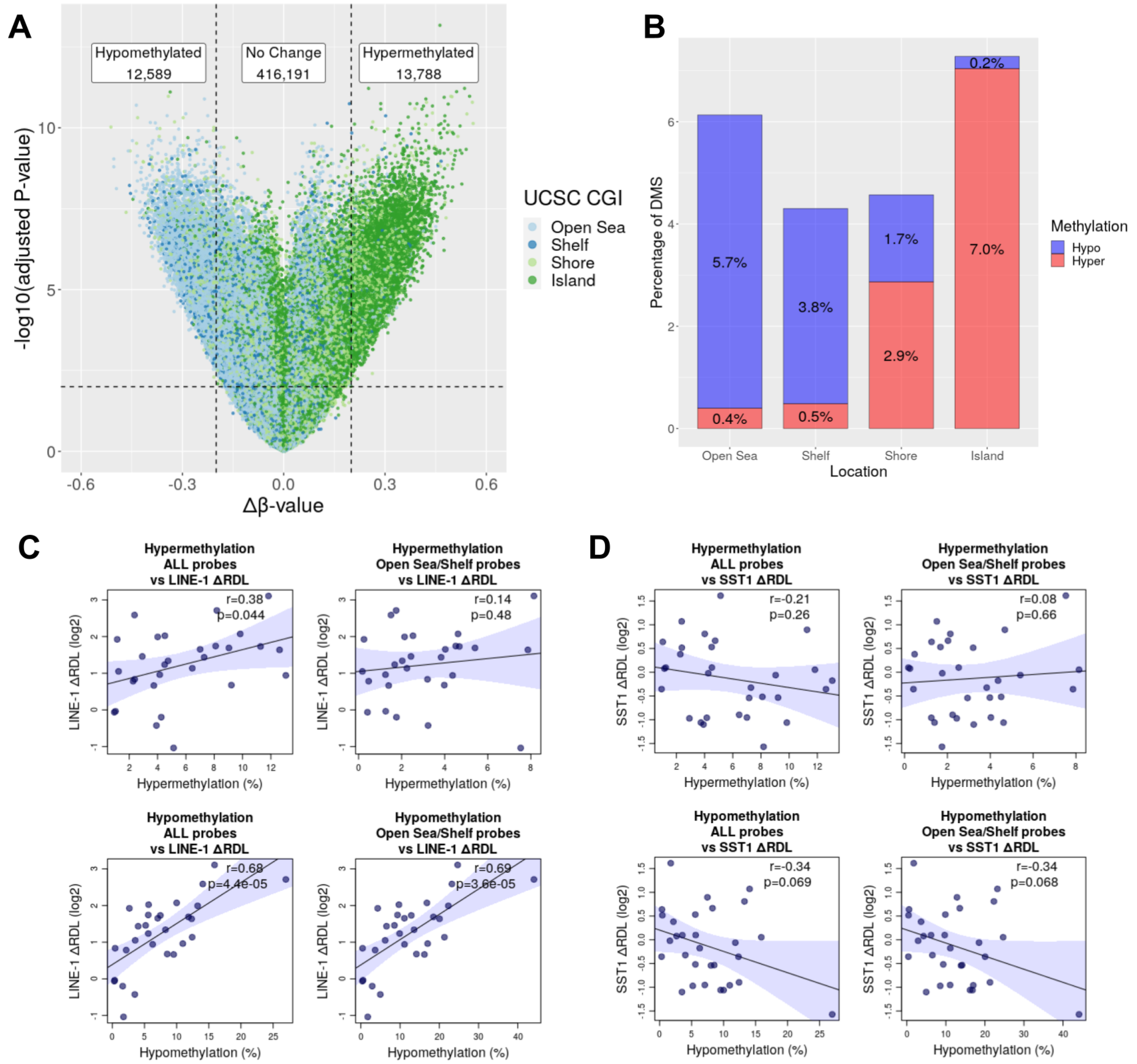


**Supplemental figure S10. A)** Volcano plot of differentially methylated CpG sites (DMS) in 30 CRCs profiled with Illumina HM450K arrays. Only autosomal CpG sites were considered (n=442,568). The Δβ-value indicates the average difference in methylation between tumors and matched normal tissues. P-values were calculated by paired t-test, and corrected using FDR method. Every dot represents a CpG site, colored according to the distance to the nearest CpG island: open sea (>4000bp, n=154,993), shelf (2000-4000bp, n=41,770), shore (<2000bp, n=103,803) or island (inside CpG island, n=142,002). Sites with a |Δβ-value| > 0.2 and FDR-corrected p-value < 0.01 were considered to be differentially methylated. 12,589 CpG sites (2.8%) were hypomethylated and 13,788 CpG sites (3.11%) were hypermethylated. **B)** Proportion of DMS according to their distance to CpG islands. CpG sites in open sea and shelves predominantly became hypomethylated (5.7% and 3.8%, respectively), while CpG sites in islands and, to a lesser extent, in shores, predominantly became hypermethylated (7% and 2.9%, respectively). **C)** Correlation of LINE-1 ΔRDL with hypermethylation (upper row) or hypomethylation (lower row), considering all CpG sites (n=442,568, left column) or just those in open sea or shelves (n=196,763, right column). Every dot represents a CRC case. For every tumor-normal pair, genome-wide hypermethylation was calculated as the percentage of probes with Δβ-value > 0.2, while genome-wide hypomethylation was calculated as the percentage of probes with Δβ-value < -0.2. The regression lines are in solid blue, and the 95% confidence interval of the slope is indicated by the blue shaded areas. The correlation coefficients and p-values are indicated. **D)** Correlation of SST1 ΔRDL with genomewide hypomethylation (upper row) or hypomethylation (lower row), considering all CpG sites (left) or those in open sea or shelves (right). Graphs and symbols as in panel C.

**11. Association of SST1 ΔRDL with clinicopathological and mutational characteristics of CRCs.**

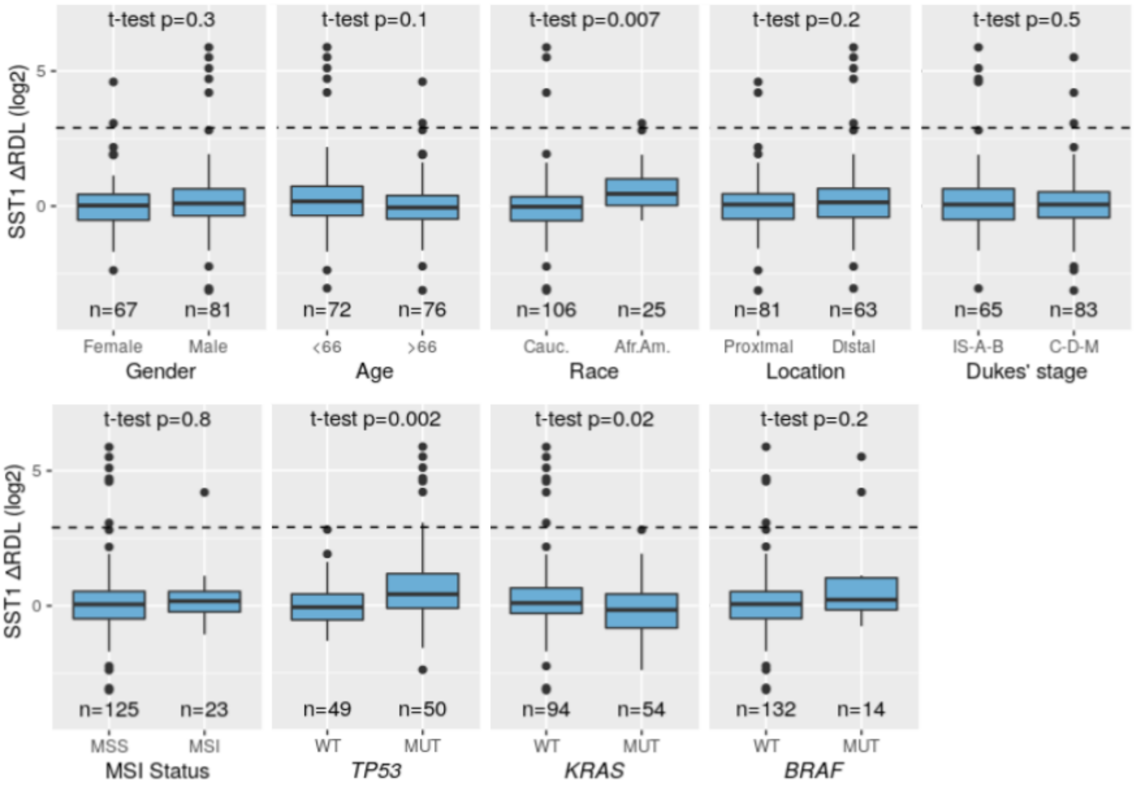


**Supplemental figure S11.** Association of SST1 ΔRDL with clinical, pathological and molecular characteristics of the CRC patients and tumors. For every analyzed parameter, patients were divided in two groups and a t-test comparing the average SST1 ΔRDL was performed. The number of informative cases in every group is indicated below each box. For age, cases were divided in two categories according to the median age of 66 years.

#### 12. *TP53* mutations in SST1 severely demethylated cases.

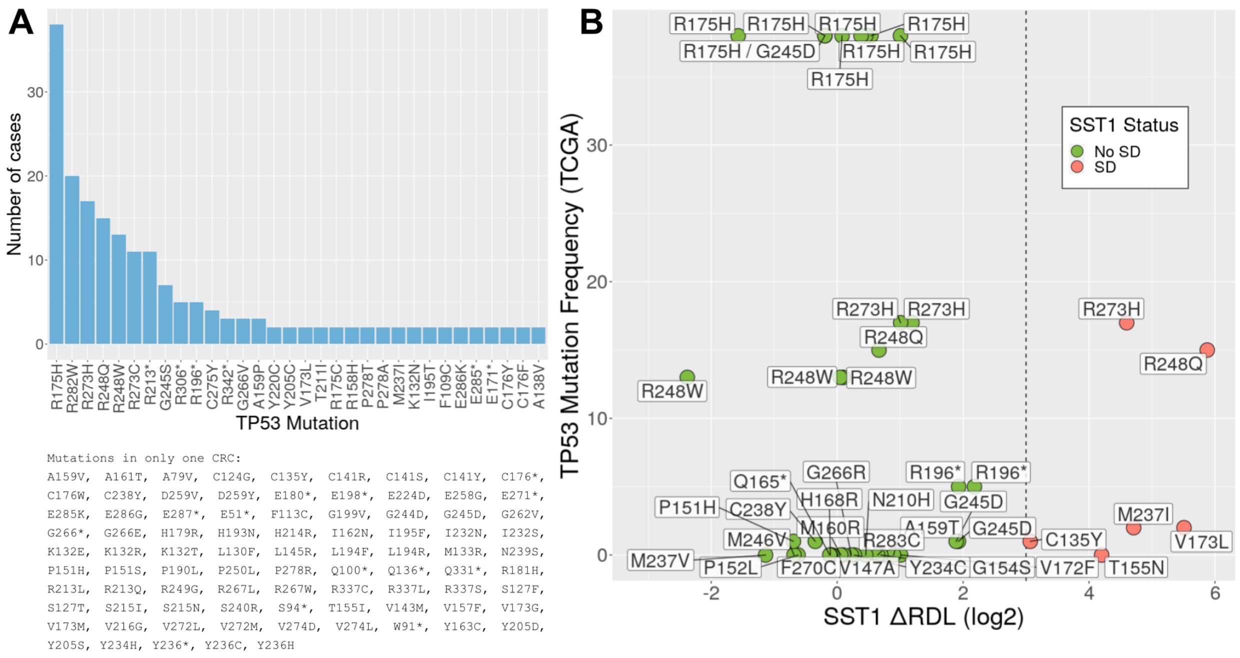


**Supplemental figure S12**. **A)** *TP53* mutations ordered according to their frequency in 535 CRC from the TCGA COAD and READ datasets. Only mutations occurring in at least two different tumors are shown in the graph. Below, mutations that occur in only one tumor. **B)** Association of SST1 somatic demethylation (x-axis) with *TP53* mutations identified in 50 out of 99 analyzed CRCs from our collection. The frequency of the mutation (y-axis) was calculated from the TCGA data (panel A). In green, *TP53* mutant tumors without SST1 severe demethylation (No SD). In red, *TP53* mutant tumors with SST1 severe demethylation (SD). The type of mutation found in every individual tumor is labeled. Severely demethylated case 153 harbored a complex multiple TP53 mutation and it is not represented in the graph. There was no clear association between SST1 severe methylation and the *TP53* mutation type frequency (Fisher's test $p$=0.32).

**13. LINE-1 MS-QPCR.**

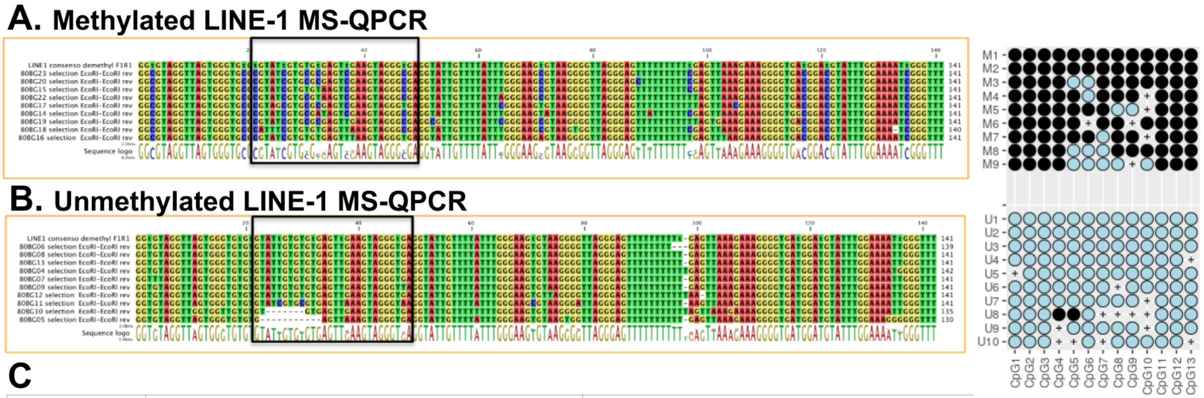


**C**

| Primer | Sequence | Reaction |
|---|---|---|
| LINE1-M-F | 5'GGCGTAGGTTAGTGGGTGCGC3' | Methylated LINE-1 MS-QPCR |
| LINE1-M-R | 5'AAACCCGATTTTCCAAATAGGTCCG3' | Methylated LINE-1 MS-QPCR |
| LINE1-U-F | 5'GGTGTAGGTTAGTGGGTGTGT3' | Unmethylated LINE-1 MS-QPCR |
| LINE1-U-R | 5'AAACCCAATTTTCCAAATACATCCA3' | Unmethylated LINE-1 MS-QPCR |

**Supplemental figure S13.** Bisulfite sequencing of LINE-1 elements amplified in DLD-1 with primers LINE1-M-F and LINE-M-R **(A)**, specific for methylated LINE-1 elements, and LINE1-U-F and LINE-U-R **(B)**, specific for methylated LINE-1 elements. Adenines in red, cytosines in blue, guanines in yellow and thymines in red. The consensus demethylated LINE-1 sequence is shown on top of both alignments. Every line correspond to an individual LINE-1 element. The black boxes indicate the region recognized by the internal probe in the original LINE-1 MethyLight technique, not employed in the derived technique LINE-1 MS-QPCR. On the right, the methylation plot of the individual LINE-1 elements shown in the alignments (9 for the methylated LINE-1 reaction and 10 for the unmethylated LINE-1 reaction). n blue, unmethylated CpG sites. In black, methylated CpG sites. Crosses indicate polymorphisms (the sequence shows other than CG or TG). **C)** Primer sequences.
