## Supplementary material for "Somatic hypomethylation of pericentromeric SST1 repeats and tetraploidization in human colorectal cancer cells": GSEA results: gsea_report_for_Diploid_1608738186968.html

Report for Diploid 1608738186968 [GSEA]

| GS  follow link to MSigDB | GS DETAILS | SIZE | ES | NES | NOM p-val | FDR q-val | FWER p-val | RANK AT MAX | LEADING EDGE || 1 | HALLMARK\_UNFOLDED\_PROTEIN\_RESPONSE | Details ... | 108 | -0.47 | -1.78 | 0.000 | 0.007 | 0.010 | 3259 | tags=33%, list=17%, signal=40% |
| 2 | HALLMARK\_PROTEIN\_SECRETION | Details ... | 95 | -0.46 | -1.72 | 0.000 | 0.009 | 0.026 | 2143 | tags=24%, list=11%, signal=27% |
| 3 | HALLMARK\_COMPLEMENT | Details ... | 199 | -0.37 | -1.51 | 0.002 | 0.071 | 0.255 | 1460 | tags=14%, list=7%, signal=15% |
| 4 | HALLMARK\_WNT\_BETA\_CATENIN\_SIGNALING | Details ... | 42 | -0.45 | -1.45 | 0.061 | 0.103 | 0.437 | 1256 | tags=17%, list=6%, signal=18% |
| 5 | HALLMARK\_PI3K\_AKT\_MTOR\_SIGNALING | Details ... | 103 | -0.35 | -1.32 | 0.065 | 0.264 | 0.853 | 1915 | tags=14%, list=10%, signal=15% |
| 6 | HALLMARK\_EPITHELIAL\_MESENCHYMAL\_TRANSITION | Details ... | 197 | -0.31 | -1.29 | 0.044 | 0.283 | 0.912 | 2104 | tags=18%, list=11%, signal=20% |
| 7 | HALLMARK\_COAGULATION | Details ... | 138 | -0.32 | -1.27 | 0.083 | 0.280 | 0.934 | 1779 | tags=16%, list=9%, signal=17% |
| 8 | HALLMARK\_HYPOXIA | Details ... | 193 | -0.30 | -1.24 | 0.082 | 0.301 | 0.965 | 3530 | tags=29%, list=18%, signal=35% |
| 9 | HALLMARK\_INTERFERON\_ALPHA\_RESPONSE | Details ... | 94 | -0.33 | -1.24 | 0.145 | 0.279 | 0.967 | 3429 | tags=27%, list=18%, signal=32% |
| 10 | HALLMARK\_XENOBIOTIC\_METABOLISM | Details ... | 198 | -0.30 | -1.23 | 0.091 | 0.261 | 0.970 | 2436 | tags=17%, list=12%, signal=19% |
| 11 | HALLMARK\_APOPTOSIS | Details ... | 159 | -0.30 | -1.21 | 0.120 | 0.270 | 0.976 | 2381 | tags=17%, list=12%, signal=19% |
| 12 | HALLMARK\_HEME\_METABOLISM | Details ... | 191 | -0.28 | -1.16 | 0.165 | 0.356 | 0.997 | 4268 | tags=29%, list=22%, signal=36% |
| 13 | HALLMARK\_INTERFERON\_GAMMA\_RESPONSE | Details ... | 196 | -0.28 | -1.15 | 0.166 | 0.346 | 0.997 | 1585 | tags=14%, list=8%, signal=15% |
| 14 | HALLMARK\_MTORC1\_SIGNALING | Details ... | 195 | -0.28 | -1.14 | 0.203 | 0.337 | 0.998 | 2843 | tags=20%, list=15%, signal=23% |
| 15 | HALLMARK\_GLYCOLYSIS | Details ... | 199 | -0.27 | -1.12 | 0.222 | 0.359 | 1.000 | 3081 | tags=22%, list=16%, signal=25% |
| 16 | HALLMARK\_KRAS\_SIGNALING\_UP | Details ... | 198 | -0.27 | -1.10 | 0.247 | 0.378 | 1.000 | 3271 | tags=25%, list=17%, signal=30% |
| 17 | HALLMARK\_UV\_RESPONSE\_UP | Details ... | 155 | -0.27 | -1.07 | 0.281 | 0.422 | 1.000 | 1503 | tags=12%, list=8%, signal=12% |
| 18 | HALLMARK\_BILE\_ACID\_METABOLISM | Details ... | 112 | -0.26 | -1.00 | 0.439 | 0.574 | 1.000 | 2941 | tags=21%, list=15%, signal=25% |
| 19 | HALLMARK\_IL6\_JAK\_STAT3\_SIGNALING | Details ... | 87 | -0.27 | -0.99 | 0.477 | 0.570 | 1.000 | 2219 | tags=18%, list=11%, signal=21% |
| 20 | HALLMARK\_ANDROGEN\_RESPONSE | Details ... | 97 | -0.26 | -0.98 | 0.482 | 0.572 | 1.000 | 1354 | tags=10%, list=7%, signal=11% |
| 21 | HALLMARK\_UV\_RESPONSE\_DN |  | 139 | -0.24 | -0.93 | 0.598 | 0.663 | 1.000 | 2334 | tags=16%, list=12%, signal=18% |
| 22 | HALLMARK\_MYOGENESIS |  | 197 | -0.22 | -0.93 | 0.645 | 0.648 | 1.000 | 2459 | tags=14%, list=13%, signal=16% |
| 23 | HALLMARK\_APICAL\_JUNCTION |  | 194 | -0.20 | -0.82 | 0.866 | 0.840 | 1.000 | 3271 | tags=18%, list=17%, signal=21% |
Table: Gene sets enriched in phenotype **Diploid (4 samples) **[plain text format]****

  
