## Supplementary material for "Somatic hypomethylation of pericentromeric SST1 repeats and tetraploidization in human colorectal cancer cells": GSEA results: gsea_report_for_Tetraploid_1608738186968.html

Report for Tetraploid 1608738186968 [GSEA]

| GS  follow link to MSigDB | GS DETAILS | SIZE | ES | NES | NOM p-val | FDR q-val | FWER p-val | RANK AT MAX | LEADING EDGE || 1 | HALLMARK\_E2F\_TARGETS | Details ... | 195 | 0.49 | 2.20 | 0.000 | 0.000 | 0.000 | 5455 | tags=55%, list=28%, signal=76% |
| 2 | HALLMARK\_MYC\_TARGETS\_V2 | Details ... | 57 | 0.56 | 2.05 | 0.000 | 0.001 | 0.001 | 6340 | tags=67%, list=32%, signal=98% |
| 3 | HALLMARK\_MYC\_TARGETS\_V1 | Details ... | 194 | 0.40 | 1.80 | 0.000 | 0.002 | 0.004 | 7315 | tags=60%, list=37%, signal=95% |
| 4 | HALLMARK\_G2M\_CHECKPOINT | Details ... | 189 | 0.41 | 1.78 | 0.000 | 0.002 | 0.005 | 5073 | tags=44%, list=26%, signal=59% |
| 5 | HALLMARK\_CHOLESTEROL\_HOMEOSTASIS | Details ... | 73 | 0.46 | 1.76 | 0.000 | 0.002 | 0.007 | 1905 | tags=38%, list=10%, signal=42% |
| 6 | HALLMARK\_INFLAMMATORY\_RESPONSE | Details ... | 199 | 0.36 | 1.59 | 0.000 | 0.018 | 0.069 | 1066 | tags=15%, list=5%, signal=15% |
| 7 | HALLMARK\_P53\_PATHWAY | Details ... | 190 | 0.35 | 1.53 | 0.000 | 0.029 | 0.126 | 2036 | tags=22%, list=10%, signal=24% |
| 8 | HALLMARK\_ESTROGEN\_RESPONSE\_LATE | Details ... | 197 | 0.30 | 1.34 | 0.015 | 0.138 | 0.531 | 2500 | tags=26%, list=13%, signal=29% |
| 9 | HALLMARK\_ESTROGEN\_RESPONSE\_EARLY | Details ... | 196 | 0.30 | 1.32 | 0.024 | 0.152 | 0.607 | 2439 | tags=23%, list=12%, signal=26% |
| 10 | HALLMARK\_ANGIOGENESIS | Details ... | 36 | 0.39 | 1.28 | 0.138 | 0.176 | 0.704 | 1363 | tags=14%, list=7%, signal=15% |
| 11 | HALLMARK\_ALLOGRAFT\_REJECTION | Details ... | 193 | 0.28 | 1.26 | 0.035 | 0.185 | 0.756 | 1838 | tags=16%, list=9%, signal=18% |
| 12 | HALLMARK\_IL2\_STAT5\_SIGNALING | Details ... | 195 | 0.28 | 1.24 | 0.068 | 0.194 | 0.809 | 2544 | tags=21%, list=13%, signal=23% |
| 13 | HALLMARK\_KRAS\_SIGNALING\_DN | Details ... | 195 | 0.27 | 1.24 | 0.039 | 0.185 | 0.820 | 3313 | tags=23%, list=17%, signal=27% |
| 14 | HALLMARK\_FATTY\_ACID\_METABOLISM | Details ... | 155 | 0.28 | 1.21 | 0.082 | 0.210 | 0.876 | 4084 | tags=30%, list=21%, signal=37% |
| 15 | HALLMARK\_TNFA\_SIGNALING\_VIA\_NFKB | Details ... | 198 | 0.26 | 1.17 | 0.137 | 0.272 | 0.952 | 1606 | tags=16%, list=8%, signal=17% |
| 16 | HALLMARK\_TGF\_BETA\_SIGNALING | Details ... | 54 | 0.31 | 1.13 | 0.239 | 0.339 | 0.981 | 1775 | tags=13%, list=9%, signal=14% |
| 17 | HALLMARK\_APICAL\_SURFACE | Details ... | 43 | 0.29 | 0.99 | 0.499 | 0.751 | 1.000 | 2421 | tags=21%, list=12%, signal=24% |
| 18 | HALLMARK\_HEDGEHOG\_SIGNALING | Details ... | 36 | 0.30 | 0.97 | 0.493 | 0.777 | 1.000 | 1363 | tags=11%, list=7%, signal=12% |
| 19 | HALLMARK\_DNA\_REPAIR | Details ... | 148 | 0.22 | 0.95 | 0.552 | 0.812 | 1.000 | 4840 | tags=31%, list=25%, signal=41% |
| 20 | HALLMARK\_SPERMATOGENESIS | Details ... | 133 | 0.23 | 0.94 | 0.589 | 0.813 | 1.000 | 3673 | tags=24%, list=19%, signal=29% |
| 21 | HALLMARK\_ADIPOGENESIS |  | 192 | 0.20 | 0.85 | 0.878 | 1.000 | 1.000 | 3939 | tags=19%, list=20%, signal=24% |
| 22 | HALLMARK\_PANCREAS\_BETA\_CELLS |  | 40 | 0.25 | 0.84 | 0.756 | 1.000 | 1.000 | 2751 | tags=20%, list=14%, signal=23% |
| 23 | HALLMARK\_OXIDATIVE\_PHOSPHORYLATION |  | 183 | 0.19 | 0.82 | 0.926 | 1.000 | 1.000 | 6935 | tags=44%, list=36%, signal=67% |
| 24 | HALLMARK\_MITOTIC\_SPINDLE |  | 197 | 0.18 | 0.81 | 0.933 | 0.999 | 1.000 | 3741 | tags=21%, list=19%, signal=25% |
| 25 | HALLMARK\_REACTIVE\_OXYGEN\_SPECIES\_PATHWAY |  | 47 | 0.22 | 0.76 | 0.873 | 1.000 | 1.000 | 6109 | tags=43%, list=31%, signal=62% |
| 26 | HALLMARK\_NOTCH\_SIGNALING |  | 32 | 0.22 | 0.71 | 0.922 | 1.000 | 1.000 | 2091 | tags=16%, list=11%, signal=17% |
| 27 | HALLMARK\_PEROXISOME |  | 104 | 0.17 | 0.70 | 0.983 | 0.977 | 1.000 | 3691 | tags=20%, list=19%, signal=25% |
Table: Gene sets enriched in phenotype **Tetraploid (4 samples) **[plain text format]****

  
