## Supplementary material for "Somatic hypomethylation of pericentromeric SST1 repeats and tetraploidization in human colorectal cancer cells": GSEA results: HALLMARK_ALLOGRAFT_REJECTION.html

Details for gene set HALLMARK\_ALLOGRAFT\_REJECTION[GSEA]

|  || Dataset | eset\_byprobe\_collapsed\_to\_symbols.Diploid\_vs\_Tetraploid.cls #Tetraploid\_versus\_Diploid.Diploid\_vs\_Tetraploid.cls #Tetraploid\_versus\_Diploid\_repos |
| Phenotype | Diploid\_vs\_Tetraploid.cls#Tetraploid\_versus\_Diploid\_repos |
| Upregulated in class | Tetraploid |
| GeneSet | HALLMARK\_ALLOGRAFT\_REJECTION |
| Enrichment Score (ES) | 0.28330335 |
| Normalized Enrichment Score (NES) | 1.2627382 |
| Nominal p-value | 0.034985423 |
| FDR q-value | 0.18454063 |
| FWER p-Value | 0.756 |
Table: GSEA Results Summary

  

Fig 1: Enrichment plot: HALLMARK\_ALLOGRAFT\_REJECTION      
 Profile of the Running ES Score & Positions of GeneSet Members on the Rank Ordered List

  

| SYMBOL | TITLE | RANK IN GENE LIST | RANK METRIC SCORE | RUNNING ES | CORE ENRICHMENT || 1 | HLA-DMA | major histocompatibility complex, class II, DM alpha | 15 | 0.392 | 0.0386 | Yes |
| 2 | CCND2 | cyclin D2 | 34 | 0.313 | 0.0691 | Yes |
| 3 | IL15 | interleukin 15 | 64 | 0.261 | 0.0938 | Yes |
| 4 | FAS | Fas cell surface death receptor | 186 | 0.161 | 0.1037 | Yes |
| 5 | APBB1 | amyloid beta (A4) precursor protein-binding, family B, member 1 (Fe65) | 224 | 0.150 | 0.1169 | Yes |
| 6 | IRF7 | interferon regulatory factor 7 | 230 | 0.148 | 0.1314 | Yes |
| 7 | IL27RA | interleukin 27 receptor, alpha | 234 | 0.147 | 0.1461 | Yes |
| 8 | TIMP1 | TIMP metallopeptidase inhibitor 1 | 254 | 0.141 | 0.1592 | Yes |
| 9 | IL2RG | interleukin 2 receptor, gamma | 268 | 0.137 | 0.1724 | Yes |
| 10 | GBP2 | guanylate binding protein 2, interferon-inducible | 302 | 0.133 | 0.1841 | Yes |
| 11 | CD47 | CD47 molecule | 349 | 0.125 | 0.1943 | Yes |
| 12 | LIF | leukemia inhibitory factor | 423 | 0.117 | 0.2022 | Yes |
| 13 | IRF8 | interferon regulatory factor 8 | 434 | 0.115 | 0.2133 | Yes |
| 14 | CD3E | CD3e molecule, epsilon (CD3-TCR complex) | 670 | 0.099 | 0.2111 | Yes |
| 15 | SRGN | serglycin | 807 | 0.092 | 0.2133 | Yes |
| 16 | TRAF2 | TNF receptor-associated factor 2 | 907 | 0.088 | 0.2170 | Yes |
| 17 | SIT1 | signaling threshold regulating transmembrane adaptor 1 | 941 | 0.087 | 0.2239 | Yes |
| 18 | FASLG | Fas ligand (TNF superfamily, member 6) | 1039 | 0.083 | 0.2272 | Yes |
| 19 | CSF1 | colony stimulating factor 1 (macrophage) | 1040 | 0.083 | 0.2356 | Yes |
| 20 | IL4R | interleukin 4 receptor | 1066 | 0.082 | 0.2425 | Yes |
| 21 | BRCA1 | breast cancer 1, early onset | 1119 | 0.080 | 0.2479 | Yes |
| 22 | CD96 | CD96 molecule | 1252 | 0.077 | 0.2488 | Yes |
| 23 | CDKN2A | cyclin-dependent kinase inhibitor 2A | 1257 | 0.077 | 0.2564 | Yes |
| 24 | IL7 | interleukin 7 | 1267 | 0.077 | 0.2637 | Yes |
| 25 | GLMN | glomulin, FKBP associated protein | 1308 | 0.076 | 0.2692 | Yes |
| 26 | EGFR | epidermal growth factor receptor | 1396 | 0.074 | 0.2722 | Yes |
| 27 | CD4 | CD4 molecule | 1530 | 0.071 | 0.2724 | Yes |
| 28 | ICAM1 | intercellular adhesion molecule 1 | 1606 | 0.069 | 0.2754 | Yes |
| 29 | NCR1 | natural cytotoxicity triggering receptor 1 | 1655 | 0.068 | 0.2797 | Yes |
| 30 | IFNG | interferon, gamma | 1797 | 0.065 | 0.2789 | Yes |
| 31 | HLA-DOB | major histocompatibility complex, class II, DO beta | 1838 | 0.064 | 0.2833 | Yes |
| 32 | IKBKB | inhibitor of kappa light polypeptide gene enhancer in B-cells, kinase beta | 2093 | 0.060 | 0.2762 | No |
| 33 | IL11 | interleukin 11 | 2352 | 0.056 | 0.2685 | No |
| 34 | MRPL3 | mitochondrial ribosomal protein L3 | 2527 | 0.053 | 0.2648 | No |
| 35 | PF4 | platelet factor 4 | 2778 | 0.050 | 0.2569 | No |
| 36 | NPM1 | nucleophosmin (nucleolar phosphoprotein B23, numatrin) | 2890 | 0.049 | 0.2561 | No |
| 37 | GALNT1 | polypeptide N-acetylgalactosaminyltransferase 1 | 3015 | 0.047 | 0.2544 | No |
| 38 | TNF | tumor necrosis factor | 3210 | 0.045 | 0.2489 | No |
| 39 | DYRK3 | dual specificity tyrosine-(Y)-phosphorylation regulated kinase 3 | 3539 | 0.042 | 0.2362 | No |
| 40 | CFP | complement factor properdin | 3594 | 0.042 | 0.2376 | No |
| 41 | IFNAR2 | interferon (alpha, beta and omega) receptor 2 | 3642 | 0.041 | 0.2393 | No |
| 42 | TGFB2 | transforming growth factor beta 2 | 3684 | 0.041 | 0.2413 | No |
| 43 | MMP9 | matrix metallopeptidase 9 | 3774 | 0.040 | 0.2407 | No |
| 44 | CD79A | CD79a molecule, immunoglobulin-associated alpha | 4078 | 0.037 | 0.2287 | No |
| 45 | IGSF6 | immunoglobulin superfamily, member 6 | 4187 | 0.036 | 0.2267 | No |
| 46 | RPS3A | ribosomal protein S3A | 4397 | 0.034 | 0.2193 | No |
| 47 | NME1 | NME/NM23 nucleoside diphosphate kinase 1 | 4477 | 0.033 | 0.2186 | No |
| 48 | LTB | lymphotoxin beta (TNF superfamily, member 3) | 4496 | 0.033 | 0.2210 | No |
| 49 | GZMB | granzyme B | 4513 | 0.033 | 0.2235 | No |
| 50 | CXCR3 | chemokine (C-X-C motif) receptor 3 | 4526 | 0.033 | 0.2262 | No |
| 51 | THY1 | Thy-1 cell surface antigen | 4641 | 0.032 | 0.2235 | No |
| 52 | UBE2N | ubiquitin conjugating enzyme E2N | 4992 | 0.029 | 0.2083 | No |
| 53 | GPR65 | G protein-coupled receptor 65 | 5142 | 0.028 | 0.2034 | No |
| 54 | LCP2 | lymphocyte cytosolic protein 2 | 5232 | 0.027 | 0.2016 | No |
| 55 | MTIF2 | mitochondrial translational initiation factor 2 | 5241 | 0.027 | 0.2039 | No |
| 56 | CAPG | capping protein (actin filament), gelsolin-like | 5399 | 0.026 | 0.1985 | No |
| 57 | PTPRC | protein tyrosine phosphatase, receptor type, C | 5462 | 0.026 | 0.1979 | No |
| 58 | CCL2 | chemokine (C-C motif) ligand 2 | 5479 | 0.026 | 0.1996 | No |
| 59 | PRF1 | perforin 1 (pore forming protein) | 5547 | 0.025 | 0.1987 | No |
| 60 | MBL2 | mannose-binding lectin (protein C) 2, soluble | 5624 | 0.025 | 0.1972 | No |
| 61 | CD86 | CD86 molecule | 5650 | 0.024 | 0.1983 | No |
| 62 | CD74 | CD74 molecule, major histocompatibility complex, class II invariant chain | 5852 | 0.023 | 0.1903 | No |
| 63 | EREG | epiregulin | 5902 | 0.023 | 0.1900 | No |
| 64 | CD8B | CD8b molecule | 5933 | 0.022 | 0.1907 | No |
| 65 | HLA-A | major histocompatibility complex, class I, A | 5955 | 0.022 | 0.1919 | No |
| 66 | NLRP3 | NLR family, pyrin domain containing 3 | 6017 | 0.022 | 0.1909 | No |
| 67 | LCK | LCK proto-oncogene, Src family tyrosine kinase | 6061 | 0.022 | 0.1909 | No |
| 68 | IL10 | interleukin 10 | 6071 | 0.022 | 0.1926 | No |
| 69 | HLA-DQA1 | major histocompatibility complex, class II, DQ alpha 1 | 6155 | 0.021 | 0.1904 | No |
| 70 | TAP1 | transporter 1, ATP-binding cassette, sub-family B (MDR/TAP) | 6169 | 0.021 | 0.1918 | No |
| 71 | RPL9 | ribosomal protein L9 | 6223 | 0.021 | 0.1911 | No |
| 72 | HLA-G | major histocompatibility complex, class I, G | 6286 | 0.020 | 0.1900 | No |
| 73 | CCL13 | chemokine (C-C motif) ligand 13 | 6395 | 0.020 | 0.1864 | No |
| 74 | MAP3K7 | mitogen-activated protein kinase kinase kinase 7 | 6582 | 0.018 | 0.1786 | No |
| 75 | C2 | complement component 2 | 6731 | 0.017 | 0.1727 | No |
| 76 | CCL19 | chemokine (C-C motif) ligand 19 | 6924 | 0.016 | 0.1644 | No |
| 77 | HDAC9 | histone deacetylase 9 | 7032 | 0.016 | 0.1605 | No |
| 78 | B2M | beta-2-microglobulin | 7058 | 0.015 | 0.1607 | No |
| 79 | IRF4 | interferon regulatory factor 4 | 7069 | 0.015 | 0.1617 | No |
| 80 | ACHE | acetylcholinesterase (Yt blood group) | 7104 | 0.015 | 0.1615 | No |
| 81 | CCL7 | chemokine (C-C motif) ligand 7 | 7135 | 0.015 | 0.1615 | No |
| 82 | ITK | IL2-inducible T-cell kinase | 7267 | 0.014 | 0.1562 | No |
| 83 | RPS19 | ribosomal protein S19 | 7403 | 0.013 | 0.1505 | No |
| 84 | ABCE1 | ATP binding cassette subfamily E member 1 | 7513 | 0.013 | 0.1462 | No |
| 85 | KLRD1 | killer cell lectin-like receptor subfamily D, member 1 | 7636 | 0.012 | 0.1411 | No |
| 86 | EIF5A | eukaryotic translation initiation factor 5A | 7652 | 0.012 | 0.1415 | No |
| 87 | NCF4 | neutrophil cytosolic factor 4 | 7776 | 0.011 | 0.1363 | No |
| 88 | BCL10 | B-cell CLL/lymphoma 10 | 7789 | 0.011 | 0.1368 | No |
| 89 | CD28 | CD28 molecule | 7826 | 0.011 | 0.1360 | No |
| 90 | IL18 | interleukin 18 | 7966 | 0.010 | 0.1298 | No |
| 91 | NCK1 | NCK adaptor protein 1 | 8042 | 0.010 | 0.1269 | No |
| 92 | F2 | coagulation factor II (thrombin) | 8109 | 0.009 | 0.1244 | No |
| 93 | CCL11 | chemokine (C-C motif) ligand 11 | 8174 | 0.009 | 0.1220 | No |
| 94 | HCLS1 | hematopoietic cell-specific Lyn substrate 1 | 8198 | 0.009 | 0.1217 | No |
| 95 | TLR1 | toll-like receptor 1 | 8229 | 0.008 | 0.1210 | No |
| 96 | CSK | c-src tyrosine kinase | 8262 | 0.008 | 0.1201 | No |
| 97 | IFNGR1 | interferon gamma receptor 1 | 8413 | 0.007 | 0.1131 | No |
| 98 | CCL5 | chemokine (C-C motif) ligand 5 | 8596 | 0.006 | 0.1044 | No |
| 99 | IL16 | interleukin 16 | 8645 | 0.006 | 0.1025 | No |
| 100 | SOCS5 | suppressor of cytokine signaling 5 | 8835 | 0.005 | 0.0932 | No |
| 101 | IL12RB1 | interleukin 12 receptor, beta 1 | 8957 | 0.004 | 0.0874 | No |
| 102 | IFNGR2 | interferon gamma receptor 2 (interferon gamma transducer 1) | 9041 | 0.004 | 0.0835 | No |
| 103 | TPD52 | tumor protein D52 | 9288 | 0.003 | 0.0710 | No |
| 104 | IL13 | interleukin 13 | 9330 | 0.002 | 0.0691 | No |
| 105 | CD40 | CD40 molecule, TNF receptor superfamily member 5 | 9416 | 0.002 | 0.0649 | No |
| 106 | CXCL9 | chemokine (C-X-C motif) ligand 9 | 9517 | 0.001 | 0.0599 | No |
| 107 | EIF3D | eukaryotic translation initiation factor 3, subunit D | 9586 | 0.001 | 0.0565 | No |
| 108 | HLA-DOA | major histocompatibility complex, class II, DO alpha | 9784 | -0.000 | 0.0463 | No |
| 109 | HLA-E | major histocompatibility complex, class I, E | 9785 | -0.000 | 0.0463 | No |
| 110 | IL18RAP | interleukin 18 receptor accessory protein | 9873 | -0.000 | 0.0418 | No |
| 111 | FLNA | Jeck2013 ANTISENSE, CDS, coding, INTERNAL, OVCODE, OVEXON best transcript NM\_001110556 | 9902 | -0.001 | 0.0404 | No |
| 112 | CD1D | CD1d molecule | 9911 | -0.001 | 0.0401 | No |
| 113 | IL2RB | interleukin 2 receptor, beta | 10056 | -0.001 | 0.0328 | No |
| 114 | ITGB2 | Memczak2013 ANTISENSE, CDS, coding, INTERNAL best transcript NM\_001127491 | 10383 | -0.003 | 0.0163 | No |
| 115 | AKT1 | v-akt murine thymoma viral oncogene homolog 1 | 10680 | -0.005 | 0.0015 | No |
| 116 | PRKCB | protein kinase C, beta | 10864 | -0.006 | -0.0073 | No |
| 117 | WAS | Wiskott-Aldrich syndrome | 10901 | -0.006 | -0.0086 | No |
| 118 | IL9 | interleukin 9 | 11135 | -0.008 | -0.0198 | No |
| 119 | RPL39 | ribosomal protein L39 | 11145 | -0.008 | -0.0195 | No |
| 120 | CD80 | CD80 molecule | 11264 | -0.009 | -0.0247 | No |
| 121 | PRKCG | protein kinase C, gamma | 11383 | -0.009 | -0.0299 | No |
| 122 | IL6 | interleukin 6 | 11825 | -0.012 | -0.0515 | No |
| 123 | INHBB | inhibin beta B | 12019 | -0.013 | -0.0602 | No |
| 124 | RPS9 | ribosomal protein S9 | 12110 | -0.014 | -0.0635 | No |
| 125 | CRTAM | cytotoxic and regulatory T-cell molecule | 12137 | -0.014 | -0.0635 | No |
| 126 | CD40LG | CD40 ligand | 12277 | -0.015 | -0.0692 | No |
| 127 | TGFB1 | transforming growth factor beta 1 | 12464 | -0.016 | -0.0772 | No |
| 128 | SOCS1 | suppressor of cytokine signaling 1 | 12611 | -0.017 | -0.0831 | No |
| 129 | RIPK2 | receptor-interacting serine-threonine kinase 2 | 12627 | -0.017 | -0.0822 | No |
| 130 | FGR | FGR proto-oncogene, Src family tyrosine kinase | 12735 | -0.018 | -0.0859 | No |
| 131 | IL12A | interleukin 12A | 12739 | -0.018 | -0.0843 | No |
| 132 | EIF3A | eukaryotic translation initiation factor 3, subunit A | 12805 | -0.018 | -0.0859 | No |
| 133 | STAT1 | signal transducer and activator of transcription 1 | 12830 | -0.018 | -0.0853 | No |
| 134 | JAK2 | Janus kinase 2 | 13031 | -0.019 | -0.0937 | No |
| 135 | F2R | coagulation factor II (thrombin) receptor | 13235 | -0.021 | -0.1021 | No |
| 136 | EIF3J | eukaryotic translation initiation factor 3, subunit J | 13279 | -0.021 | -0.1022 | No |
| 137 | CD7 | CD7 molecule | 13289 | -0.021 | -0.1006 | No |
| 138 | RPL3L | ribosomal protein L3-like | 13315 | -0.021 | -0.0997 | No |
| 139 | IL4 | interleukin 4 | 13374 | -0.022 | -0.1006 | No |
| 140 | UBE2D1 | ubiquitin conjugating enzyme E2D 1 | 13523 | -0.023 | -0.1059 | No |
| 141 | ABI1 | abl-interactor 1 | 13706 | -0.024 | -0.1129 | No |
| 142 | PTPN6 | protein tyrosine phosphatase, non-receptor type 6 | 13725 | -0.024 | -0.1114 | No |
| 143 | ETS1 | v-ets avian erythroblastosis virus E26 oncogene homolog 1 | 13826 | -0.025 | -0.1141 | No |
| 144 | TAP2 | transporter 2, ATP-binding cassette, sub-family B (MDR/TAP) | 13911 | -0.026 | -0.1158 | No |
| 145 | ACVR2A | activin A receptor type IIA | 14013 | -0.026 | -0.1184 | No |
| 146 | TAPBP | TAP binding protein (tapasin) | 14083 | -0.027 | -0.1193 | No |
| 147 | HLA-DMB | major histocompatibility complex, class II, DM beta | 14394 | -0.030 | -0.1323 | No |
| 148 | LYN | LYN proto-oncogene, Src family tyrosine kinase | 14438 | -0.030 | -0.1316 | No |
| 149 | ITGAL | integrin alpha L | 14573 | -0.031 | -0.1354 | No |
| 150 | CCND3 | cyclin D3 | 14958 | -0.034 | -0.1518 | No |
| 151 | INHBA | inhibin beta A | 15087 | -0.035 | -0.1549 | No |
| 152 | PSMB10 | proteasome subunit beta 10 | 15143 | -0.036 | -0.1542 | No |
| 153 | CD8A | CD8a molecule | 15182 | -0.036 | -0.1525 | No |
| 154 | FCGR2B | Fc fragment of IgG, low affinity IIb, receptor (CD32) | 15298 | -0.037 | -0.1548 | No |
| 155 | HLA-DRA | major histocompatibility complex, class II, DR alpha | 15300 | -0.037 | -0.1511 | No |
| 156 | STAB1 | stabilin 1 | 15306 | -0.037 | -0.1476 | No |
| 157 | IL2 | interleukin 2 | 15633 | -0.040 | -0.1605 | No |
| 158 | BCL3 | B-cell CLL/lymphoma 3 | 15800 | -0.042 | -0.1649 | No |
| 159 | IL2RA | interleukin 2 receptor, alpha | 16231 | -0.046 | -0.1825 | No |
| 160 | CD3G | CD3g molecule, gamma (CD3-TCR complex) | 16331 | -0.047 | -0.1828 | No |
| 161 | TLR2 | toll-like receptor 2 | 16448 | -0.049 | -0.1839 | No |
| 162 | LY86 | lymphocyte antigen 86 | 16508 | -0.050 | -0.1819 | No |
| 163 | CXCL13 | chemokine (C-X-C motif) ligand 13 | 16578 | -0.051 | -0.1804 | No |
| 164 | CTSS | cathepsin S | 16657 | -0.052 | -0.1792 | No |
| 165 | SPI1 | Spi-1 proto-oncogene | 16721 | -0.053 | -0.1771 | No |
| 166 | NOS2 | nitric oxide synthase 2, inducible | 16738 | -0.053 | -0.1726 | No |
| 167 | ICOSLG | inducible T-cell co-stimulator ligand | 16929 | -0.056 | -0.1769 | No |
| 168 | HIF1A | hypoxia inducible factor 1, alpha subunit (basic helix-loop-helix transcription factor) | 16936 | -0.056 | -0.1716 | No |
| 169 | EIF4G3 | eukaryotic translation initiation factor 4 gamma, 3 | 16968 | -0.056 | -0.1675 | No |
| 170 | BCAT1 | branched chain amino-acid transaminase 1, cytosolic | 17138 | -0.059 | -0.1704 | No |
| 171 | CD2 | CD2 molecule | 17207 | -0.060 | -0.1678 | No |
| 172 | ELF4 | E74-like factor 4 (ets domain transcription factor) | 17291 | -0.062 | -0.1659 | No |
| 173 | KRT1 | keratin 1, type II | 17418 | -0.064 | -0.1660 | No |
| 174 | CD247 | CD247 molecule | 17434 | -0.064 | -0.1604 | No |
| 175 | CD3D | CD3d molecule, delta (CD3-TCR complex) | 17649 | -0.068 | -0.1646 | No |
| 176 | DEGS1 | delta(4)-desaturase, sphingolipid 1 | 17711 | -0.070 | -0.1607 | No |
| 177 | MAP4K1 | mitogen-activated protein kinase kinase kinase kinase 1 | 17834 | -0.073 | -0.1597 | No |
| 178 | CARTPT | CART prepropeptide | 17856 | -0.073 | -0.1534 | No |
| 179 | STAT4 | signal transducer and activator of transcription 4 | 17951 | -0.076 | -0.1507 | No |
| 180 | TRAT1 | T cell receptor associated transmembrane adaptor 1 | 17976 | -0.076 | -0.1442 | No |
| 181 | ELANE | elastase, neutrophil expressed | 18070 | -0.079 | -0.1411 | No |
| 182 | CCL22 | chemokine (C-C motif) ligand 22 | 18217 | -0.083 | -0.1404 | No |
| 183 | IL12B | interleukin 12B | 18360 | -0.087 | -0.1389 | No |
| 184 | GCNT1 | glucosaminyl (N-acetyl) transferase 1, core 2 | 18387 | -0.088 | -0.1314 | No |
| 185 | CCR1 | chemokine (C-C motif) receptor 1 | 18734 | -0.105 | -0.1388 | No |
| 186 | ZAP70 | zeta chain of T cell receptor associated protein kinase 70kDa | 18776 | -0.108 | -0.1301 | No |
| 187 | TLR6 | toll-like receptor 6 | 18913 | -0.118 | -0.1252 | No |
| 188 | IL1B | interleukin 1 beta | 19023 | -0.130 | -0.1178 | No |
| 189 | ST8SIA4 | ST8 alpha-N-acetyl-neuraminide alpha-2,8-sialyltransferase 4 | 19177 | -0.154 | -0.1102 | No |
| 190 | GZMA | granzyme A | 19327 | -0.200 | -0.0979 | No |
| 191 | CCR5 | chemokine (C-C motif) receptor 5 (gene/pseudogene) | 19474 | -0.331 | -0.0722 | No |
| 192 | CCR2 | chemokine (C-C motif) receptor 2 | 19477 | -0.333 | -0.0389 | No |
| 193 | TLR3 | toll-like receptor 3 | 19502 | -0.411 | 0.0011 | No |
Table: GSEA details [plain text format]

  

Fig 2: HALLMARK\_ALLOGRAFT\_REJECTION      
 Blue-Pink O' Gram in the Space of the Analyzed GeneSet

  

Fig 3: HALLMARK\_ALLOGRAFT\_REJECTION: Random ES distribution      
 Gene set null distribution of ES for **HALLMARK\_ALLOGRAFT\_REJECTION**

  
