## Supplementary material for "Somatic hypomethylation of pericentromeric SST1 repeats and tetraploidization in human colorectal cancer cells": GSEA results: HALLMARK_ANDROGEN_RESPONSE.html

Details for gene set HALLMARK\_ANDROGEN\_RESPONSE[GSEA]

|  || Dataset | eset\_byprobe\_collapsed\_to\_symbols.Diploid\_vs\_Tetraploid.cls #Tetraploid\_versus\_Diploid.Diploid\_vs\_Tetraploid.cls #Tetraploid\_versus\_Diploid\_repos |
| Phenotype | Diploid\_vs\_Tetraploid.cls#Tetraploid\_versus\_Diploid\_repos |
| Upregulated in class | Diploid |
| GeneSet | HALLMARK\_ANDROGEN\_RESPONSE |
| Enrichment Score (ES) | -0.26096466 |
| Normalized Enrichment Score (NES) | -0.9788245 |
| Nominal p-value | 0.48248407 |
| FDR q-value | 0.5723843 |
| FWER p-Value | 1.0 |
Table: GSEA Results Summary

  

Fig 1: Enrichment plot: HALLMARK\_ANDROGEN\_RESPONSE      
 Profile of the Running ES Score & Positions of GeneSet Members on the Rank Ordered List

  

| SYMBOL | TITLE | RANK IN GENE LIST | RANK METRIC SCORE | RUNNING ES | CORE ENRICHMENT || 1 | HPGD | hydroxyprostaglandin dehydrogenase 15-(NAD) | 62 | 0.267 | 0.0512 | No |
| 2 | KRT19 | keratin 19, type I | 157 | 0.170 | 0.0809 | No |
| 3 | INPP4B | inositol polyphosphate-4-phosphatase type II B | 262 | 0.140 | 0.1040 | No |
| 4 | FADS1 | fatty acid desaturase 1 | 296 | 0.134 | 0.1297 | No |
| 5 | ZBTB10 | zinc finger and BTB domain containing 10 | 326 | 0.129 | 0.1544 | No |
| 6 | HMGCS1 | 3-hydroxy-3-methylglutaryl-CoA synthase 1 (soluble) | 387 | 0.121 | 0.1759 | No |
| 7 | ELL2 | elongation factor, RNA polymerase II, 2 | 918 | 0.087 | 0.1664 | No |
| 8 | KLK2 | kallikrein related peptidase 2 | 1372 | 0.075 | 0.1583 | No |
| 9 | HMGCR | 3-hydroxy-3-methylglutaryl-CoA reductase | 1821 | 0.064 | 0.1484 | No |
| 10 | FKBP5 | FK506 binding protein 5 | 1840 | 0.064 | 0.1605 | No |
| 11 | DBI | diazepam binding inhibitor (GABA receptor modulator, acyl-CoA binding protein) | 1874 | 0.064 | 0.1718 | No |
| 12 | IDI1 | isopentenyl-diphosphate delta isomerase 1 | 1905 | 0.063 | 0.1831 | No |
| 13 | MAP7 | microtubule-associated protein 7 | 2428 | 0.055 | 0.1674 | No |
| 14 | SORD | sorbitol dehydrogenase | 2500 | 0.054 | 0.1747 | No |
| 15 | DHCR24 | 24-dehydrocholesterol reductase | 2597 | 0.052 | 0.1804 | No |
| 16 | IQGAP2 | IQ motif containing GTPase activating protein 2 | 2654 | 0.052 | 0.1880 | No |
| 17 | STK39 | serine threonine kinase 39 | 2848 | 0.049 | 0.1881 | No |
| 18 | SCD | stearoyl-CoA desaturase (delta-9-desaturase) | 3227 | 0.045 | 0.1779 | No |
| 19 | INSIG1 | insulin induced gene 1 | 3412 | 0.044 | 0.1773 | No |
| 20 | CENPN | centromere protein N | 3649 | 0.041 | 0.1736 | No |
| 21 | KRT8 | keratin 8, type II | 3950 | 0.038 | 0.1658 | No |
| 22 | MYL12A | myosin light chain 12A | 4217 | 0.035 | 0.1594 | No |
| 23 | XRCC5 | X-ray repair complementing defective repair in Chinese hamster cells 5 (double-strand-break rejoining) | 4524 | 0.033 | 0.1503 | No |
| 24 | CCND1 | cyclin D1 | 4585 | 0.032 | 0.1539 | No |
| 25 | ABHD2 | abhydrolase domain containing 2 | 4774 | 0.031 | 0.1505 | No |
| 26 | HSD17B14 | hydroxysteroid (17-beta) dehydrogenase 14 | 5014 | 0.029 | 0.1442 | No |
| 27 | ARID5B | AT rich interactive domain 5B (MRF1-like) | 5331 | 0.027 | 0.1333 | No |
| 28 | VAPA | VAMP associated protein A | 5442 | 0.026 | 0.1330 | No |
| 29 | ELOVL5 | ELOVL fatty acid elongase 5 | 5798 | 0.023 | 0.1194 | No |
| 30 | ACTN1 | actinin, alpha 1 | 5826 | 0.023 | 0.1228 | No |
| 31 | SMS | spermine synthase | 6236 | 0.021 | 0.1059 | No |
| 32 | TMPRSS2 | transmembrane protease, serine 2 | 6244 | 0.021 | 0.1098 | No |
| 33 | PA2G4 | proliferation-associated 2G4 | 6269 | 0.020 | 0.1127 | No |
| 34 | SGK1 | serum/glucocorticoid regulated kinase 1 | 6290 | 0.020 | 0.1158 | No |
| 35 | RRP12 | Zhang2013 ALT\_ACCEPTOR, ALT\_DONOR, coding, INTERNAL, intronic best transcript NM\_015179 | 6340 | 0.020 | 0.1174 | No |
| 36 | PLPP1 | phospholipid phosphatase 1 | 6641 | 0.018 | 0.1056 | No |
| 37 | ADAMTS1 | ADAM metallopeptidase with thrombospondin type 1 motif 1 | 6653 | 0.018 | 0.1087 | No |
| 38 | PDLIM5 | PDZ and LIM domain 5 | 6773 | 0.017 | 0.1061 | No |
| 39 | TMEM50A | transmembrane protein 50A | 6828 | 0.017 | 0.1067 | No |
| 40 | ELK4 | ELK4, ETS-domain protein (SRF accessory protein 1) | 6969 | 0.016 | 0.1028 | No |
| 41 | B2M | beta-2-microglobulin | 7058 | 0.015 | 0.1014 | No |
| 42 | RAB4A | RAB4A, member RAS oncogene family | 7729 | 0.012 | 0.0693 | No |
| 43 | TNFAIP8 | tumor necrosis factor, alpha-induced protein 8 | 8321 | 0.008 | 0.0405 | No |
| 44 | XRCC6 | X-ray repair complementing defective repair in Chinese hamster cells 6 | 8395 | 0.007 | 0.0382 | No |
| 45 | NDRG1 | N-myc downstream regulated 1 | 9078 | 0.004 | 0.0039 | No |
| 46 | TPD52 | tumor protein D52 | 9288 | 0.003 | -0.0064 | No |
| 47 | NKX3-1 | NK3 homeobox 1 | 9663 | 0.001 | -0.0255 | No |
| 48 | PTPN21 | protein tyrosine phosphatase, non-receptor type 21 | 9681 | 0.001 | -0.0263 | No |
| 49 | PMEPA1 | prostate transmembrane protein, androgen induced 1 | 9736 | 0.000 | -0.0290 | No |
| 50 | GSR | glutathione reductase | 9874 | -0.000 | -0.0359 | No |
| 51 | ADRM1 | adhesion regulating molecule 1 | 10136 | -0.002 | -0.0490 | No |
| 52 | CAMKK2 | calcium/calmodulin-dependent protein kinase kinase 2, beta | 10423 | -0.004 | -0.0630 | No |
| 53 | AKT1 | v-akt murine thymoma viral oncogene homolog 1 | 10680 | -0.005 | -0.0751 | No |
| 54 | ALDH1A3 | aldehyde dehydrogenase 1 family, member A3 | 10720 | -0.005 | -0.0760 | No |
| 55 | RPS6KA3 | ribosomal protein S6 kinase, 90kDa, polypeptide 3 | 10766 | -0.006 | -0.0771 | No |
| 56 | GNAI3 | guanine nucleotide binding protein (G protein), alpha inhibiting activity polypeptide 3 | 10833 | -0.006 | -0.0793 | No |
| 57 | UBE2I | ubiquitin conjugating enzyme E2I | 10871 | -0.006 | -0.0799 | No |
| 58 | CDK6 | cyclin-dependent kinase 6 | 10904 | -0.007 | -0.0802 | No |
| 59 | LIFR | leukemia inhibitory factor receptor alpha | 10934 | -0.007 | -0.0803 | No |
| 60 | ACSL3 | acyl-CoA synthetase long-chain family member 3 | 11087 | -0.008 | -0.0866 | No |
| 61 | GPD1L | glycerol-3-phosphate dehydrogenase 1-like | 11488 | -0.010 | -0.1052 | No |
| 62 | KLK3 | kallikrein related peptidase 3 | 12104 | -0.013 | -0.1341 | No |
| 63 | PIAS1 | protein inhibitor of activated STAT 1 | 12419 | -0.016 | -0.1471 | No |
| 64 | ZMIZ1 | zinc finger, MIZ-type containing 1 | 12589 | -0.017 | -0.1524 | No |
| 65 | MAF | v-maf avian musculoaponeurotic fibrosarcoma oncogene homolog | 13016 | -0.019 | -0.1704 | No |
| 66 | NCOA4 | nuclear receptor coactivator 4 | 13253 | -0.021 | -0.1783 | No |
| 67 | AZGP1 | alpha-2-glycoprotein 1, zinc-binding | 13489 | -0.022 | -0.1858 | No |
| 68 | NGLY1 | N-glycanase 1 | 13512 | -0.023 | -0.1823 | No |
| 69 | AKAP12 | A kinase (PRKA) anchor protein 12 | 13539 | -0.023 | -0.1790 | No |
| 70 | APPBP2 | amyloid beta precursor protein (cytoplasmic tail) binding protein 2 | 13843 | -0.025 | -0.1895 | No |
| 71 | TSC22D1 | TSC22 domain family, member 1 | 14022 | -0.026 | -0.1933 | No |
| 72 | STEAP4 | STEAP family member 4 | 14727 | -0.032 | -0.2229 | No |
| 73 | SLC38A2 | solute carrier family 38, member 2 | 14825 | -0.033 | -0.2212 | No |
| 74 | CDC14B | cell division cycle 14B | 14922 | -0.034 | -0.2193 | No |
| 75 | CCND3 | cyclin D3 | 14958 | -0.034 | -0.2141 | No |
| 76 | SRF | serum response factor | 15271 | -0.037 | -0.2227 | No |
| 77 | HERC3 | HECT and RLD domain containing E3 ubiquitin protein ligase 3 | 15566 | -0.039 | -0.2298 | No |
| 78 | UAP1 | UDP-N-acetylglucosamine pyrophosphorylase 1 | 15846 | -0.042 | -0.2356 | No |
| 79 | SLC26A2 | solute carrier family 26 (anion exchanger), member 2 | 15913 | -0.043 | -0.2302 | No |
| 80 | ITGAV | integrin alpha V | 16105 | -0.045 | -0.2309 | No |
| 81 | PTK2B | protein tyrosine kinase 2 beta | 16127 | -0.045 | -0.2228 | No |
| 82 | ANKH | ANKH inorganic pyrophosphate transport regulator | 16325 | -0.047 | -0.2233 | No |
| 83 | B4GALT1 | UDP-Gal:betaGlcNAc beta 1,4- galactosyltransferase, polypeptide 1 | 16445 | -0.049 | -0.2194 | No |
| 84 | SPCS3 | signal peptidase complex subunit 3 | 16529 | -0.050 | -0.2134 | No |
| 85 | SRP19 | signal recognition particle 19kDa | 16910 | -0.055 | -0.2217 | No |
| 86 | BMPR1B | bone morphogenetic protein receptor type IB | 16952 | -0.056 | -0.2124 | No |
| 87 | UBE2J1 | ubiquitin-conjugating enzyme E2, J1 | 17690 | -0.069 | -0.2362 | No |
| 88 | SPDEF | SAM pointed domain containing ETS transcription factor | 18172 | -0.082 | -0.2443 | Yes |
| 89 | SAT1 | spermidine/spermine N1-acetyltransferase 1 | 18295 | -0.085 | -0.2332 | Yes |
| 90 | MAK | male germ cell-associated kinase | 18556 | -0.095 | -0.2272 | Yes |
| 91 | LMAN1 | lectin, mannose-binding, 1 | 18623 | -0.097 | -0.2108 | Yes |
| 92 | PGM3 | phosphoglucomutase 3 | 19104 | -0.142 | -0.2066 | Yes |
| 93 | ABCC4 | ATP binding cassette subfamily C member 4 | 19150 | -0.149 | -0.1784 | Yes |
| 94 | HOMER2 | homer scaffolding protein 2 | 19317 | -0.197 | -0.1469 | Yes |
| 95 | DNAJB9 | DnaJ (Hsp40) homolog, subfamily B, member 9 | 19366 | -0.220 | -0.1046 | Yes |
| 96 | SEC24D | SEC24 homolog D, COPII coat complex component | 19384 | -0.228 | -0.0590 | Yes |
| 97 | MERTK | MER proto-oncogene, tyrosine kinase | 19468 | -0.325 | 0.0029 | Yes |
Table: GSEA details [plain text format]

  

Fig 2: HALLMARK\_ANDROGEN\_RESPONSE      
 Blue-Pink O' Gram in the Space of the Analyzed GeneSet

  

Fig 3: HALLMARK\_ANDROGEN\_RESPONSE: Random ES distribution      
 Gene set null distribution of ES for **HALLMARK\_ANDROGEN\_RESPONSE**

  
