## Supplementary material for "Somatic hypomethylation of pericentromeric SST1 repeats and tetraploidization in human colorectal cancer cells": GSEA results: HALLMARK_ANGIOGENESIS.html

Details for gene set HALLMARK\_ANGIOGENESIS[GSEA]

|  || Dataset | eset\_byprobe\_collapsed\_to\_symbols.Diploid\_vs\_Tetraploid.cls #Tetraploid\_versus\_Diploid.Diploid\_vs\_Tetraploid.cls #Tetraploid\_versus\_Diploid\_repos |
| Phenotype | Diploid\_vs\_Tetraploid.cls#Tetraploid\_versus\_Diploid\_repos |
| Upregulated in class | Tetraploid |
| GeneSet | HALLMARK\_ANGIOGENESIS |
| Enrichment Score (ES) | 0.39035353 |
| Normalized Enrichment Score (NES) | 1.2824036 |
| Nominal p-value | 0.13793103 |
| FDR q-value | 0.17610374 |
| FWER p-Value | 0.704 |
Table: GSEA Results Summary

  

Fig 1: Enrichment plot: HALLMARK\_ANGIOGENESIS      
 Profile of the Running ES Score & Positions of GeneSet Members on the Rank Ordered List

  

| SYMBOL | TITLE | RANK IN GENE LIST | RANK METRIC SCORE | RUNNING ES | CORE ENRICHMENT || 1 | S100A4 | S100 calcium binding protein A4 | 5 | 0.471 | 0.1995 | Yes |
| 2 | CCND2 | cyclin D2 | 34 | 0.313 | 0.3306 | Yes |
| 3 | TIMP1 | TIMP metallopeptidase inhibitor 1 | 254 | 0.141 | 0.3792 | Yes |
| 4 | JAG1 | jagged 1 | 962 | 0.085 | 0.3792 | Yes |
| 5 | NRP1 | neuropilin 1 | 1363 | 0.075 | 0.3904 | Yes |
| 6 | KCNJ8 | potassium channel, inwardly rectifying subfamily J, member 8 | 2165 | 0.059 | 0.3742 | No |
| 7 | PF4 | platelet factor 4 | 2778 | 0.050 | 0.3641 | No |
| 8 | STC1 | stanniocalcin 1 | 3174 | 0.046 | 0.3632 | No |
| 9 | SERPINA5 | serpin peptidase inhibitor, clade A (alpha-1 antiproteinase, antitrypsin), member 5 | 3551 | 0.042 | 0.3617 | No |
| 10 | VTN | vitronectin | 3798 | 0.040 | 0.3659 | No |
| 11 | VAV2 | vav 2 guanine nucleotide exchange factor | 4415 | 0.034 | 0.3486 | No |
| 12 | PDGFA | platelet-derived growth factor alpha polypeptide | 5403 | 0.026 | 0.3091 | No |
| 13 | SLCO2A1 | solute carrier organic anion transporter family, member 2A1 | 5751 | 0.024 | 0.3013 | No |
| 14 | APP | amyloid beta (A4) precursor protein | 6318 | 0.020 | 0.2808 | No |
| 15 | LRPAP1 | LDL receptor related protein associated protein 1 | 6616 | 0.018 | 0.2733 | No |
| 16 | PTK2 | protein tyrosine kinase 2 | 6854 | 0.017 | 0.2683 | No |
| 17 | APOH | apolipoprotein H (beta-2-glycoprotein I) | 7891 | 0.010 | 0.2195 | No |
| 18 | OLR1 | oxidized low density lipoprotein (lectin-like) receptor 1 | 8290 | 0.008 | 0.2025 | No |
| 19 | JAG2 | jagged 2 | 10208 | -0.002 | 0.1052 | No |
| 20 | PRG2 | proteoglycan 2, bone marrow (natural killer cell activator, eosinophil granule major basic protein) | 10857 | -0.006 | 0.0746 | No |
| 21 | THBD | thrombomodulin | 11144 | -0.008 | 0.0633 | No |
| 22 | SPP1 | secreted phosphoprotein 1 | 11575 | -0.010 | 0.0455 | No |
| 23 | COL3A1 | collagen, type III, alpha 1 | 11694 | -0.011 | 0.0442 | No |
| 24 | CXCL6 | chemokine (C-X-C motif) ligand 6 | 11900 | -0.012 | 0.0388 | No |
| 25 | COL5A2 | collagen, type V, alpha 2 | 12761 | -0.018 | 0.0022 | No |
| 26 | LPL | lipoprotein lipase | 13793 | -0.025 | -0.0402 | No |
| 27 | FSTL1 | follistatin like 1 | 13956 | -0.026 | -0.0375 | No |
| 28 | FGFR1 | fibroblast growth factor receptor 1 | 15489 | -0.039 | -0.0997 | No |
| 29 | TNFRSF21 | tumor necrosis factor receptor superfamily, member 21 | 15802 | -0.042 | -0.0980 | No |
| 30 | ITGAV | integrin alpha V | 16105 | -0.045 | -0.0944 | No |
| 31 | MSX1 | msh homeobox 1 | 16206 | -0.046 | -0.0801 | No |
| 32 | PGLYRP1 | peptidoglycan recognition protein 1 | 16294 | -0.047 | -0.0646 | No |
| 33 | VEGFA | vascular endothelial growth factor A | 16659 | -0.052 | -0.0612 | No |
| 34 | VCAN | versican | 18696 | -0.103 | -0.1222 | No |
| 35 | POSTN | periostin, osteoblast specific factor | 18928 | -0.119 | -0.0834 | No |
| 36 | LUM | lumican | 19432 | -0.269 | 0.0047 | No |
Table: GSEA details [plain text format]

  

Fig 2: HALLMARK\_ANGIOGENESIS      
 Blue-Pink O' Gram in the Space of the Analyzed GeneSet

  

Fig 3: HALLMARK\_ANGIOGENESIS: Random ES distribution      
 Gene set null distribution of ES for **HALLMARK\_ANGIOGENESIS**

  
