## Supplementary material for "Somatic hypomethylation of pericentromeric SST1 repeats and tetraploidization in human colorectal cancer cells": GSEA results: HALLMARK_APICAL_SURFACE.html

Details for gene set HALLMARK\_APICAL\_SURFACE[GSEA]

|  || Dataset | eset\_byprobe\_collapsed\_to\_symbols.Diploid\_vs\_Tetraploid.cls #Tetraploid\_versus\_Diploid.Diploid\_vs\_Tetraploid.cls #Tetraploid\_versus\_Diploid\_repos |
| Phenotype | Diploid\_vs\_Tetraploid.cls#Tetraploid\_versus\_Diploid\_repos |
| Upregulated in class | Tetraploid |
| GeneSet | HALLMARK\_APICAL\_SURFACE |
| Enrichment Score (ES) | 0.28885508 |
| Normalized Enrichment Score (NES) | 0.9896641 |
| Nominal p-value | 0.49878934 |
| FDR q-value | 0.75086987 |
| FWER p-Value | 1.0 |
Table: GSEA Results Summary

  

Fig 1: Enrichment plot: HALLMARK\_APICAL\_SURFACE      
 Profile of the Running ES Score & Positions of GeneSet Members on the Rank Ordered List

  

| SYMBOL | TITLE | RANK IN GENE LIST | RANK METRIC SCORE | RUNNING ES | CORE ENRICHMENT || 1 | SULF2 | Memczak2013 ANTISENSE, coding, INTERNAL, intronic best transcript NM\_001161841 | 135 | 0.185 | 0.0756 | Yes |
| 2 | AKAP7 | A kinase (PRKA) anchor protein 7 | 145 | 0.176 | 0.1540 | Yes |
| 3 | IL2RG | interleukin 2 receptor, gamma | 268 | 0.137 | 0.2093 | Yes |
| 4 | HSPB1 | heat shock 27kDa protein 1 | 832 | 0.091 | 0.2210 | Yes |
| 5 | BRCA1 | breast cancer 1, early onset | 1119 | 0.080 | 0.2423 | Yes |
| 6 | PLAUR | plasminogen activator, urokinase receptor | 1355 | 0.075 | 0.2638 | Yes |
| 7 | CX3CL1 | chemokine (C-X3-C motif) ligand 1 | 1648 | 0.068 | 0.2790 | Yes |
| 8 | RTN4RL1 | reticulon 4 receptor-like 1 | 2386 | 0.055 | 0.2660 | Yes |
| 9 | ATP8B1 | ATPase, aminophospholipid transporter, class I, type 8B, member 1 | 2421 | 0.055 | 0.2889 | Yes |
| 10 | GSTM3 | glutathione S-transferase mu 3 (brain) | 3265 | 0.045 | 0.2657 | No |
| 11 | DCBLD2 | discoidin, CUB and LCCL domain containing 2 | 3862 | 0.039 | 0.2525 | No |
| 12 | THY1 | Thy-1 cell surface antigen | 4641 | 0.032 | 0.2269 | No |
| 13 | GATA3 | GATA binding protein 3 | 5072 | 0.029 | 0.2177 | No |
| 14 | SLC2A4 | solute carrier family 2 (facilitated glucose transporter), member 4 | 5764 | 0.024 | 0.1928 | No |
| 15 | NTNG1 | netrin G1 | 6033 | 0.022 | 0.1888 | No |
| 16 | APP | amyloid beta (A4) precursor protein | 6318 | 0.020 | 0.1833 | No |
| 17 | TMEM8B | transmembrane protein 8B | 6457 | 0.019 | 0.1848 | No |
| 18 | SHROOM2 | shroom family member 2 | 7112 | 0.015 | 0.1581 | No |
| 19 | SLC22A12 | solute carrier family 22 (organic anion/urate transporter), member 12 | 7260 | 0.014 | 0.1570 | No |
| 20 | PCSK9 | proprotein convertase subtilisin/kexin type 9 | 7351 | 0.014 | 0.1586 | No |
| 21 | FLOT2 | Memczak2013 ANTISENSE, coding, INTERNAL, UTR3 best transcript NM\_004475 | 7502 | 0.013 | 0.1566 | No |
| 22 | ADIPOR2 | adiponectin receptor 2 | 8746 | 0.005 | 0.0952 | No |
| 23 | EPHB4 | EPH receptor B4 | 9714 | 0.000 | 0.0458 | No |
| 24 | CD160 | CD160 molecule | 9741 | 0.000 | 0.0445 | No |
| 25 | MDGA1 | MAM domain containing glycosylphosphatidylinositol anchor 1 | 9871 | -0.000 | 0.0381 | No |
| 26 | IL2RB | interleukin 2 receptor, beta | 10056 | -0.001 | 0.0294 | No |
| 27 | GHRL | ghrelin/obestatin prepropeptide | 10232 | -0.002 | 0.0215 | No |
| 28 | ADAM10 | ADAM metallopeptidase domain 10 | 10365 | -0.003 | 0.0162 | No |
| 29 | ATP6V0A4 | ATPase, H+ transporting, lysosomal V0 subunit a4 | 10508 | -0.004 | 0.0108 | No |
| 30 | MAL | mal, T-cell differentiation protein | 10821 | -0.006 | -0.0025 | No |
| 31 | GAS1 | growth arrest-specific 1 | 11303 | -0.009 | -0.0233 | No |
| 32 | CROCC | Zhang2013 ALT\_ACCEPTOR, ALT\_DONOR, coding, INTERNAL, intronic best transcript NM\_014675 | 11337 | -0.009 | -0.0210 | No |
| 33 | RHCG | Rh family, C glycoprotein | 11430 | -0.010 | -0.0214 | No |
| 34 | NCOA6 | nuclear receptor coactivator 6 | 12234 | -0.014 | -0.0562 | No |
| 35 | LYN | LYN proto-oncogene, Src family tyrosine kinase | 14438 | -0.030 | -0.1560 | No |
| 36 | SRPX | sushi-repeat containing protein, X-linked | 15588 | -0.040 | -0.1972 | No |
| 37 | SLC34A3 | solute carrier family 34 (type II sodium/phosphate cotransporter), member 3 | 16222 | -0.046 | -0.2091 | No |
| 38 | LYPD3 | LY6/PLAUR domain containing 3 | 16257 | -0.047 | -0.1899 | No |
| 39 | B4GALT1 | UDP-Gal:betaGlcNAc beta 1,4- galactosyltransferase, polypeptide 1 | 16445 | -0.049 | -0.1776 | No |
| 40 | AFAP1L2 | actin filament associated protein 1-like 2 | 16865 | -0.055 | -0.1747 | No |
| 41 | SCUBE1 | signal peptide, CUB domain, EGF-like 1 | 18021 | -0.077 | -0.1993 | No |
| 42 | PKHD1 | polycystic kidney and hepatic disease 1 (autosomal recessive) | 18461 | -0.091 | -0.1811 | No |
| 43 | EFNA5 | ephrin-A5 | 19522 | -0.527 | 0.0001 | No |
Table: GSEA details [plain text format]

  

Fig 2: HALLMARK\_APICAL\_SURFACE      
 Blue-Pink O' Gram in the Space of the Analyzed GeneSet

  

Fig 3: HALLMARK\_APICAL\_SURFACE: Random ES distribution      
 Gene set null distribution of ES for **HALLMARK\_APICAL\_SURFACE**

  
