## Supplementary material for "Somatic hypomethylation of pericentromeric SST1 repeats and tetraploidization in human colorectal cancer cells": GSEA results: HALLMARK_APOPTOSIS.html

Details for gene set HALLMARK\_APOPTOSIS[GSEA]

|  || Dataset | eset\_byprobe\_collapsed\_to\_symbols.Diploid\_vs\_Tetraploid.cls #Tetraploid\_versus\_Diploid.Diploid\_vs\_Tetraploid.cls #Tetraploid\_versus\_Diploid\_repos |
| Phenotype | Diploid\_vs\_Tetraploid.cls#Tetraploid\_versus\_Diploid\_repos |
| Upregulated in class | Diploid |
| GeneSet | HALLMARK\_APOPTOSIS |
| Enrichment Score (ES) | -0.3042963 |
| Normalized Enrichment Score (NES) | -1.2141395 |
| Nominal p-value | 0.11958147 |
| FDR q-value | 0.26956424 |
| FWER p-Value | 0.976 |
Table: GSEA Results Summary

  

Fig 1: Enrichment plot: HALLMARK\_APOPTOSIS      
 Profile of the Running ES Score & Positions of GeneSet Members on the Rank Ordered List

  

| SYMBOL | TITLE | RANK IN GENE LIST | RANK METRIC SCORE | RUNNING ES | CORE ENRICHMENT || 1 | CCND2 | cyclin D2 | 34 | 0.313 | 0.0323 | No |
| 2 | FAS | Fas cell surface death receptor | 186 | 0.161 | 0.0420 | No |
| 3 | IL1A | interleukin 1 alpha | 237 | 0.146 | 0.0553 | No |
| 4 | TIMP1 | TIMP metallopeptidase inhibitor 1 | 254 | 0.141 | 0.0698 | No |
| 5 | RNASEL | ribonuclease L (2,5-oligoisoadenylate synthetase-dependent) | 259 | 0.140 | 0.0848 | No |
| 6 | BTG2 | BTG family, member 2 | 281 | 0.136 | 0.0985 | No |
| 7 | SMAD7 | SMAD family member 7 | 372 | 0.123 | 0.1072 | No |
| 8 | BTG3 | BTG family, member 3 | 465 | 0.113 | 0.1148 | No |
| 9 | CD44 | CD44 molecule (Indian blood group) | 577 | 0.104 | 0.1204 | No |
| 10 | CD38 | CD38 molecule | 689 | 0.097 | 0.1253 | No |
| 11 | CDKN1A | cyclin-dependent kinase inhibitor 1A (p21, Cip1) | 690 | 0.097 | 0.1359 | No |
| 12 | HSPB1 | heat shock 27kDa protein 1 | 832 | 0.091 | 0.1385 | No |
| 13 | SPTAN1 | spectrin, alpha, non-erythrocytic 1 | 923 | 0.087 | 0.1433 | No |
| 14 | EMP1 | epithelial membrane protein 1 | 967 | 0.085 | 0.1504 | No |
| 15 | TNFSF10 | tumor necrosis factor (ligand) superfamily, member 10 | 1006 | 0.084 | 0.1576 | No |
| 16 | FASLG | Fas ligand (TNF superfamily, member 6) | 1039 | 0.083 | 0.1650 | No |
| 17 | EBP | emopamil binding protein (sterol isomerase) | 1093 | 0.081 | 0.1710 | No |
| 18 | BRCA1 | breast cancer 1, early onset | 1119 | 0.080 | 0.1785 | No |
| 19 | FDXR | ferredoxin reductase | 1184 | 0.079 | 0.1838 | No |
| 20 | BIRC3 | Memczak2013 ANTISENSE, CDS, coding, INTERNAL best transcript NM\_182962 | 1400 | 0.074 | 0.1807 | No |
| 21 | DNAJA1 | DnaJ (Hsp40) homolog, subfamily A, member 1 | 1418 | 0.074 | 0.1878 | No |
| 22 | TGFBR3 | transforming growth factor beta receptor III | 2120 | 0.060 | 0.1581 | No |
| 23 | CTH | cystathionine gamma-lyase | 2192 | 0.058 | 0.1608 | No |
| 24 | PDGFRB | platelet-derived growth factor receptor, beta polypeptide | 2343 | 0.056 | 0.1592 | No |
| 25 | BCL2L1 | BCL2-like 1 | 2466 | 0.054 | 0.1588 | No |
| 26 | CDK2 | cyclin-dependent kinase 2 | 2666 | 0.051 | 0.1541 | No |
| 27 | LMNA | lamin A/C | 2699 | 0.051 | 0.1580 | No |
| 28 | CASP6 | caspase 6 | 2891 | 0.049 | 0.1534 | No |
| 29 | ENO2 | enolase 2 (gamma, neuronal) | 3086 | 0.047 | 0.1485 | No |
| 30 | BAX | BCL2-associated X protein | 3143 | 0.046 | 0.1506 | No |
| 31 | TNF | tumor necrosis factor | 3210 | 0.045 | 0.1521 | No |
| 32 | HMOX1 | heme oxygenase 1 | 3502 | 0.043 | 0.1417 | No |
| 33 | BID | BH3 interacting domain death agonist | 3546 | 0.042 | 0.1441 | No |
| 34 | AIFM3 | apoptosis-inducing factor, mitochondrion-associated, 3 | 3582 | 0.042 | 0.1468 | No |
| 35 | NEDD9 | neural precursor cell expressed, developmentally down-regulated 9 | 3625 | 0.041 | 0.1491 | No |
| 36 | TGFB2 | transforming growth factor beta 2 | 3684 | 0.041 | 0.1506 | No |
| 37 | KRT18 | keratin 18, type I | 3794 | 0.040 | 0.1493 | No |
| 38 | PDCD4 | programmed cell death 4 (neoplastic transformation inhibitor) | 3801 | 0.040 | 0.1533 | No |
| 39 | LGALS3 | lectin, galactoside-binding, soluble, 3 | 3918 | 0.038 | 0.1514 | No |
| 40 | PLCB2 | phospholipase C, beta 2 | 4167 | 0.036 | 0.1425 | No |
| 41 | WEE1 | WEE1 G2 checkpoint kinase | 4267 | 0.035 | 0.1412 | No |
| 42 | GPX1 | glutathione peroxidase 1 | 4413 | 0.034 | 0.1374 | No |
| 43 | CCND1 | cyclin D1 | 4585 | 0.032 | 0.1321 | No |
| 44 | HGF | hepatocyte growth factor (hepapoietin A; scatter factor) | 4645 | 0.032 | 0.1326 | No |
| 45 | CASP3 | caspase 3 | 4699 | 0.032 | 0.1333 | No |
| 46 | RHOT2 | ras homolog family member T2 | 4853 | 0.030 | 0.1287 | No |
| 47 | TIMP3 | TIMP metallopeptidase inhibitor 3 | 4889 | 0.030 | 0.1301 | No |
| 48 | SOD1 | superoxide dismutase 1, soluble | 4956 | 0.030 | 0.1300 | No |
| 49 | PPT1 | palmitoyl-protein thioesterase 1 | 5049 | 0.029 | 0.1284 | No |
| 50 | SATB1 | SATB homeobox 1 | 5090 | 0.029 | 0.1294 | No |
| 51 | PRF1 | perforin 1 (pore forming protein) | 5547 | 0.025 | 0.1086 | No |
| 52 | GADD45B | growth arrest and DNA-damage-inducible, beta | 5568 | 0.025 | 0.1103 | No |
| 53 | EREG | epiregulin | 5902 | 0.023 | 0.0955 | No |
| 54 | GSTM1 | glutathione S-transferase mu 1 | 5992 | 0.022 | 0.0933 | No |
| 55 | GPX4 | glutathione peroxidase 4 | 6109 | 0.021 | 0.0897 | No |
| 56 | TAP1 | transporter 1, ATP-binding cassette, sub-family B (MDR/TAP) | 6169 | 0.021 | 0.0889 | No |
| 57 | APP | amyloid beta (A4) precursor protein | 6318 | 0.020 | 0.0835 | No |
| 58 | MGMT | O-6-methylguanine-DNA methyltransferase | 6514 | 0.019 | 0.0754 | No |
| 59 | BCL2L11 | BCL2-like 11 (apoptosis facilitator) | 6584 | 0.018 | 0.0739 | No |
| 60 | TSPO | translocator protein (18kDa) | 6706 | 0.018 | 0.0696 | No |
| 61 | TNFRSF12A | tumor necrosis factor receptor superfamily, member 12A | 6787 | 0.017 | 0.0673 | No |
| 62 | CASP7 | caspase 7 | 6811 | 0.017 | 0.0679 | No |
| 63 | PTK2 | protein tyrosine kinase 2 | 6854 | 0.017 | 0.0676 | No |
| 64 | TOP2A | topoisomerase (DNA) II alpha | 6917 | 0.016 | 0.0662 | No |
| 65 | DAP3 | death associated protein 3 | 6953 | 0.016 | 0.0661 | No |
| 66 | CASP2 | caspase 2 | 7131 | 0.015 | 0.0586 | No |
| 67 | CREBBP | CREB binding protein | 7206 | 0.015 | 0.0564 | No |
| 68 | ROCK1 | Rho-associated, coiled-coil containing protein kinase 1 | 7223 | 0.015 | 0.0572 | No |
| 69 | GADD45A | growth arrest and DNA-damage-inducible, alpha | 7290 | 0.014 | 0.0553 | No |
| 70 | NEFH | neurofilament, heavy polypeptide | 7329 | 0.014 | 0.0549 | No |
| 71 | BMF | Bcl2 modifying factor | 7405 | 0.013 | 0.0525 | No |
| 72 | MCL1 | myeloid cell leukemia 1 | 7519 | 0.013 | 0.0480 | No |
| 73 | IFITM3 | interferon induced transmembrane protein 3 | 7579 | 0.012 | 0.0463 | No |
| 74 | DPYD | dihydropyrimidine dehydrogenase | 7629 | 0.012 | 0.0451 | No |
| 75 | DNM1L | dynamin 1-like | 7773 | 0.011 | 0.0390 | No |
| 76 | BCL10 | B-cell CLL/lymphoma 10 | 7789 | 0.011 | 0.0394 | No |
| 77 | HMGB2 | high mobility group box 2 | 7921 | 0.010 | 0.0338 | No |
| 78 | IL18 | interleukin 18 | 7966 | 0.010 | 0.0326 | No |
| 79 | BCAP31 | B-cell receptor-associated protein 31 | 7967 | 0.010 | 0.0337 | No |
| 80 | F2 | coagulation factor II (thrombin) | 8109 | 0.009 | 0.0274 | No |
| 81 | IFNB1 | interferon, beta 1, fibroblast | 8176 | 0.009 | 0.0250 | No |
| 82 | IFNGR1 | interferon gamma receptor 1 | 8413 | 0.007 | 0.0136 | No |
| 83 | SOD2 | superoxide dismutase 2, mitochondrial | 8450 | 0.007 | 0.0125 | No |
| 84 | VDAC2 | voltage-dependent anion channel 2 | 8459 | 0.007 | 0.0129 | No |
| 85 | BNIP3L | BCL2/adenovirus E1B 19kDa interacting protein 3-like | 8648 | 0.006 | 0.0038 | No |
| 86 | ERBB2 | Salzman2013 ANNOTATED, coding, OVEXON, UTR5 best transcript NM\_001005862 | 9057 | 0.004 | -0.0169 | No |
| 87 | CFLAR | CASP8 and FADD like apoptosis regulator | 9278 | 0.003 | -0.0279 | No |
| 88 | ADD1 | adducin 1 (alpha) | 9577 | 0.001 | -0.0432 | No |
| 89 | GSR | glutathione reductase | 9874 | -0.000 | -0.0584 | No |
| 90 | SC5D | sterol-C5-desaturase | 9987 | -0.001 | -0.0641 | No |
| 91 | RARA | retinoic acid receptor, alpha | 10083 | -0.002 | -0.0688 | No |
| 92 | DIABLO | diablo, IAP-binding mitochondrial protein | 10516 | -0.004 | -0.0907 | No |
| 93 | PSEN1 | presenilin 1 | 10566 | -0.004 | -0.0927 | No |
| 94 | FEZ1 | fasciculation and elongation protein zeta 1 | 10684 | -0.005 | -0.0982 | No |
| 95 | PLAT | plasminogen activator, tissue | 10760 | -0.006 | -0.1014 | No |
| 96 | SLC20A1 | solute carrier family 20 (phosphate transporter), member 1 | 10841 | -0.006 | -0.1049 | No |
| 97 | CTNNB1 | catenin (cadherin-associated protein), beta 1 | 11452 | -0.010 | -0.1354 | No |
| 98 | CASP8 | caspase 8, apoptosis-related cysteine peptidase | 11632 | -0.011 | -0.1434 | No |
| 99 | IL6 | interleukin 6 | 11825 | -0.012 | -0.1521 | No |
| 100 | ETF1 | eukaryotic translation termination factor 1 | 12626 | -0.017 | -0.1916 | No |
| 101 | GCH1 | GTP cyclohydrolase 1 | 12793 | -0.018 | -0.1982 | No |
| 102 | GUCY2D | guanylate cyclase 2D, membrane (retina-specific) | 12929 | -0.019 | -0.2031 | No |
| 103 | SQSTM1 | sequestosome 1 | 13032 | -0.019 | -0.2063 | No |
| 104 | F2R | coagulation factor II (thrombin) receptor | 13235 | -0.021 | -0.2144 | No |
| 105 | BCL2L2 | BCL2-like 2 | 13343 | -0.022 | -0.2176 | No |
| 106 | DFFA | DNA fragmentation factor, 45kDa, alpha polypeptide | 13346 | -0.022 | -0.2154 | No |
| 107 | CD69 | CD69 molecule | 13361 | -0.022 | -0.2138 | No |
| 108 | CD14 | CD14 molecule | 13724 | -0.024 | -0.2298 | No |
| 109 | GPX3 | glutathione peroxidase 3 | 13735 | -0.024 | -0.2277 | No |
| 110 | BCL2L10 | BCL2-like 10 (apoptosis facilitator) | 13778 | -0.025 | -0.2271 | No |
| 111 | CCNA1 | cyclin A1 | 13813 | -0.025 | -0.2262 | No |
| 112 | GNA15 | guanine nucleotide binding protein (G protein), alpha 15 (Gq class) | 13912 | -0.026 | -0.2284 | No |
| 113 | CDC25B | cell division cycle 25B | 13972 | -0.026 | -0.2287 | No |
| 114 | IER3 | immediate early response 3 | 14076 | -0.027 | -0.2310 | No |
| 115 | EGR3 | early growth response 3 | 14802 | -0.033 | -0.2649 | No |
| 116 | CYLD | cylindromatosis (turban tumor syndrome) | 15093 | -0.035 | -0.2761 | No |
| 117 | ERBB3 | erb-b2 receptor tyrosine kinase 3 | 15110 | -0.035 | -0.2730 | No |
| 118 | MADD | MAP-kinase activating death domain | 15161 | -0.036 | -0.2717 | No |
| 119 | DCN | decorin | 15284 | -0.037 | -0.2740 | No |
| 120 | RELA | v-rel avian reticuloendotheliosis viral oncogene homolog A | 15578 | -0.039 | -0.2849 | No |
| 121 | PPP3R1 | protein phosphatase 3, regulatory subunit B, alpha | 15701 | -0.041 | -0.2867 | No |
| 122 | PPP2R5B | protein phosphatase 2, regulatory subunit B, beta | 15844 | -0.042 | -0.2895 | No |
| 123 | XIAP | X-linked inhibitor of apoptosis, E3 ubiquitin protein ligase | 15992 | -0.044 | -0.2923 | No |
| 124 | CAV1 | caveolin 1 | 16000 | -0.044 | -0.2879 | No |
| 125 | TXNIP | thioredoxin interacting protein | 16097 | -0.045 | -0.2880 | No |
| 126 | ANKH | ANKH inorganic pyrophosphate transport regulator | 16325 | -0.047 | -0.2946 | No |
| 127 | ISG20 | interferon stimulated exonuclease gene 20kDa | 16332 | -0.047 | -0.2897 | No |
| 128 | CDKN1B | cyclin-dependent kinase inhibitor 1B (p27, Kip1) | 16336 | -0.047 | -0.2847 | No |
| 129 | BMP2 | bone morphogenetic protein 2 | 16350 | -0.048 | -0.2802 | No |
| 130 | GSN | gelsolin | 16387 | -0.048 | -0.2768 | No |
| 131 | CASP9 | caspase 9 | 16638 | -0.052 | -0.2841 | No |
| 132 | PEA15 | phosphoprotein enriched in astrocytes 15 | 16790 | -0.054 | -0.2860 | No |
| 133 | DAP | death-associated protein | 17145 | -0.059 | -0.2979 | Yes |
| 134 | AVPR1A | arginine vasopressin receptor 1A | 17181 | -0.060 | -0.2932 | Yes |
| 135 | CD2 | CD2 molecule | 17207 | -0.060 | -0.2879 | Yes |
| 136 | BIK | BCL2-interacting killer (apoptosis-inducing) | 17482 | -0.065 | -0.2950 | Yes |
| 137 | PSEN2 | presenilin 2 | 17539 | -0.066 | -0.2907 | Yes |
| 138 | IGFBP6 | insulin like growth factor binding protein 6 | 17555 | -0.066 | -0.2843 | Yes |
| 139 | PAK1 | p21 protein (Cdc42/Rac)-activated kinase 1 | 17576 | -0.067 | -0.2781 | Yes |
| 140 | RETSAT | retinol saturase (all-trans-retinol 13,14-reductase) | 17683 | -0.069 | -0.2760 | Yes |
| 141 | IGF2R | insulin-like growth factor 2 receptor | 17794 | -0.072 | -0.2739 | Yes |
| 142 | ANXA1 | annexin A1 | 17825 | -0.073 | -0.2675 | Yes |
| 143 | PMAIP1 | phorbol-12-myristate-13-acetate-induced protein 1 | 17862 | -0.073 | -0.2614 | Yes |
| 144 | SAT1 | spermidine/spermine N1-acetyltransferase 1 | 18295 | -0.085 | -0.2744 | Yes |
| 145 | IRF1 | interferon regulatory factor 1 | 18746 | -0.106 | -0.2861 | Yes |
| 146 | RHOB | ras homolog family member B | 18827 | -0.112 | -0.2781 | Yes |
| 147 | IL1B | interleukin 1 beta | 19023 | -0.130 | -0.2741 | Yes |
| 148 | CASP4 | caspase 4 | 19027 | -0.131 | -0.2600 | Yes |
| 149 | DNAJC3 | DnaJ (Hsp40) homolog, subfamily C, member 3 | 19028 | -0.131 | -0.2458 | Yes |
| 150 | JUN | jun proto-oncogene | 19037 | -0.132 | -0.2319 | Yes |
| 151 | MMP2 | matrix metallopeptidase 2 | 19166 | -0.152 | -0.2219 | Yes |
| 152 | CLU | clusterin | 19343 | -0.205 | -0.2087 | Yes |
| 153 | PLPPR4 | phospholipid phosphatase related 4 | 19396 | -0.238 | -0.1855 | Yes |
| 154 | LEF1 | lymphoid enhancer-binding factor 1 | 19413 | -0.250 | -0.1591 | Yes |
| 155 | ATF3 | activating transcription factor 3 | 19421 | -0.254 | -0.1318 | Yes |
| 156 | LUM | lumican | 19432 | -0.269 | -0.1031 | Yes |
| 157 | DDIT3 | DNA-damage-inducible transcript 3 | 19440 | -0.282 | -0.0727 | Yes |
| 158 | CASP1 | caspase 1 | 19457 | -0.309 | -0.0400 | Yes |
| 159 | BGN | biglycan | 19498 | -0.399 | 0.0013 | Yes |
Table: GSEA details [plain text format]

  

Fig 2: HALLMARK\_APOPTOSIS      
 Blue-Pink O' Gram in the Space of the Analyzed GeneSet

  

Fig 3: HALLMARK\_APOPTOSIS: Random ES distribution      
 Gene set null distribution of ES for **HALLMARK\_APOPTOSIS**

  
