## Supplementary material for "Somatic hypomethylation of pericentromeric SST1 repeats and tetraploidization in human colorectal cancer cells": GSEA results: HALLMARK_BILE_ACID_METABOLISM.html

Details for gene set HALLMARK\_BILE\_ACID\_METABOLISM[GSEA]

|  || Dataset | eset\_byprobe\_collapsed\_to\_symbols.Diploid\_vs\_Tetraploid.cls #Tetraploid\_versus\_Diploid.Diploid\_vs\_Tetraploid.cls #Tetraploid\_versus\_Diploid\_repos |
| Phenotype | Diploid\_vs\_Tetraploid.cls#Tetraploid\_versus\_Diploid\_repos |
| Upregulated in class | Diploid |
| GeneSet | HALLMARK\_BILE\_ACID\_METABOLISM |
| Enrichment Score (ES) | -0.26138258 |
| Normalized Enrichment Score (NES) | -1.0010406 |
| Nominal p-value | 0.4390625 |
| FDR q-value | 0.57425636 |
| FWER p-Value | 1.0 |
Table: GSEA Results Summary

  

Fig 1: Enrichment plot: HALLMARK\_BILE\_ACID\_METABOLISM      
 Profile of the Running ES Score & Positions of GeneSet Members on the Rank Ordered List

  

| SYMBOL | TITLE | RANK IN GENE LIST | RANK METRIC SCORE | RUNNING ES | CORE ENRICHMENT || 1 | LIPE | lipase, hormone-sensitive | 236 | 0.147 | 0.0157 | No |
| 2 | FADS1 | fatty acid desaturase 1 | 296 | 0.134 | 0.0380 | No |
| 3 | FADS2 | fatty acid desaturase 2 | 315 | 0.131 | 0.0618 | No |
| 4 | ABCA3 | ATP binding cassette subfamily A member 3 | 321 | 0.129 | 0.0860 | No |
| 5 | SLC29A1 | solute carrier family 29 (equilibrative nucleoside transporter), member 1 | 410 | 0.118 | 0.1037 | No |
| 6 | BBOX1 | butyrobetaine (gamma), 2-oxoglutarate dioxygenase (gamma-butyrobetaine hydroxylase) 1 | 773 | 0.093 | 0.1028 | No |
| 7 | ABCG4 | ATP binding cassette subfamily G member 4 | 791 | 0.093 | 0.1194 | No |
| 8 | FDXR | ferredoxin reductase | 1184 | 0.079 | 0.1141 | No |
| 9 | HSD3B7 | hydroxy-delta-5-steroid dehydrogenase, 3 beta- and steroid delta-isomerase 7 | 1321 | 0.076 | 0.1215 | No |
| 10 | PEX11G | peroxisomal biogenesis factor 11 gamma | 1366 | 0.075 | 0.1333 | No |
| 11 | CYP46A1 | cytochrome P450, family 46, subfamily A, polypeptide 1 | 1399 | 0.074 | 0.1457 | No |
| 12 | NUDT12 | nudix hydrolase 12 | 1455 | 0.073 | 0.1566 | No |
| 13 | HSD3B1 | hydroxy-delta-5-steroid dehydrogenase, 3 beta- and steroid delta-isomerase 1 | 1615 | 0.068 | 0.1613 | No |
| 14 | IDI1 | isopentenyl-diphosphate delta isomerase 1 | 1905 | 0.063 | 0.1584 | No |
| 15 | CYP7B1 | cytochrome P450, family 7, subfamily B, polypeptide 1 | 2130 | 0.060 | 0.1581 | No |
| 16 | PEX7 | peroxisomal biogenesis factor 7 | 2240 | 0.058 | 0.1634 | No |
| 17 | ACSL5 | acyl-CoA synthetase long-chain family member 5 | 2576 | 0.053 | 0.1561 | No |
| 18 | DHCR24 | 24-dehydrocholesterol reductase | 2597 | 0.052 | 0.1650 | No |
| 19 | RXRG | retinoid X receptor gamma | 3043 | 0.047 | 0.1510 | No |
| 20 | NR1H4 | nuclear receptor subfamily 1, group H, member 4 | 3094 | 0.047 | 0.1572 | No |
| 21 | GNPAT | glyceronephosphate O-acyltransferase | 3177 | 0.046 | 0.1616 | No |
| 22 | SLC27A5 | solute carrier family 27 (fatty acid transporter), member 5 | 3371 | 0.044 | 0.1600 | No |
| 23 | ALDH1A1 | aldehyde dehydrogenase 1 family, member A1 | 3520 | 0.042 | 0.1604 | No |
| 24 | MLYCD | malonyl-CoA decarboxylase | 4013 | 0.037 | 0.1421 | No |
| 25 | ABCA9 | ATP binding cassette subfamily A member 9 | 4236 | 0.035 | 0.1374 | No |
| 26 | DIO1 | deiodinase, iodothyronine, type I | 4242 | 0.035 | 0.1438 | No |
| 27 | PXMP2 | peroxisomal membrane protein 2 | 4607 | 0.032 | 0.1311 | No |
| 28 | PAOX | polyamine oxidase (exo-N4-amino) | 4688 | 0.032 | 0.1330 | No |
| 29 | SOD1 | superoxide dismutase 1, soluble | 4956 | 0.030 | 0.1249 | No |
| 30 | CYP7A1 | cytochrome P450, family 7, subfamily A, polypeptide 1 | 5043 | 0.029 | 0.1259 | No |
| 31 | ISOC1 | isochorismatase domain containing 1 | 5132 | 0.028 | 0.1268 | No |
| 32 | ABCG8 | ATP binding cassette subfamily G member 8 | 5200 | 0.028 | 0.1286 | No |
| 33 | ABCD2 | Transcript Identified by AceView, Entrez Gene ID(s) 225 | 5228 | 0.028 | 0.1324 | No |
| 34 | ALDH9A1 | aldehyde dehydrogenase 9 family, member A1 | 5592 | 0.025 | 0.1183 | No |
| 35 | PEX16 | peroxisomal biogenesis factor 16 | 5713 | 0.024 | 0.1167 | No |
| 36 | HSD17B6 | hydroxysteroid (17-beta) dehydrogenase 6 | 5970 | 0.022 | 0.1077 | No |
| 37 | LCK | LCK proto-oncogene, Src family tyrosine kinase | 6061 | 0.022 | 0.1072 | No |
| 38 | IDH2 | isocitrate dehydrogenase 2 (NADP+), mitochondrial | 6235 | 0.021 | 0.1022 | No |
| 39 | ABCD3 | ATP binding cassette subfamily D member 3 | 6577 | 0.018 | 0.0881 | No |
| 40 | PEX11A | peroxisomal biogenesis factor 11 alpha | 6645 | 0.018 | 0.0881 | No |
| 41 | SLC27A2 | solute carrier family 27 (fatty acid transporter), member 2 | 6687 | 0.018 | 0.0893 | No |
| 42 | HSD17B11 | hydroxysteroid (17-beta) dehydrogenase 11 | 7455 | 0.013 | 0.0523 | No |
| 43 | GC | group-specific component (vitamin D binding protein) | 7563 | 0.013 | 0.0492 | No |
| 44 | CYP27A1 | cytochrome P450, family 27, subfamily A, polypeptide 1 | 7603 | 0.012 | 0.0495 | No |
| 45 | RXRA | retinoid X receptor alpha | 7732 | 0.011 | 0.0451 | No |
| 46 | ABCA2 | ATP binding cassette subfamily A member 2 | 7772 | 0.011 | 0.0452 | No |
| 47 | CH25H | cholesterol 25-hydroxylase | 7941 | 0.010 | 0.0384 | No |
| 48 | SLCO1A2 | solute carrier organic anion transporter family, member 1A2 | 8285 | 0.008 | 0.0223 | No |
| 49 | GNMT | glycine N-methyltransferase | 8445 | 0.007 | 0.0155 | No |
| 50 | PFKM | phosphofructokinase, muscle | 8889 | 0.005 | -0.0064 | No |
| 51 | CYP8B1 | cytochrome P450, family 8, subfamily B, polypeptide 1 | 9102 | 0.003 | -0.0167 | No |
| 52 | SCP2 | sterol carrier protein 2 | 9698 | 0.000 | -0.0473 | No |
| 53 | NEDD4 | neural precursor cell expressed, developmentally down-regulated 4, E3 ubiquitin protein ligase | 9825 | -0.000 | -0.0537 | No |
| 54 | ACSL1 | acyl-CoA synthetase long-chain family member 1 | 9852 | -0.000 | -0.0550 | No |
| 55 | LONP2 | lon peptidase 2, peroxisomal | 10172 | -0.002 | -0.0710 | No |
| 56 | PEX26 | peroxisomal biogenesis factor 26 | 10209 | -0.002 | -0.0724 | No |
| 57 | AGXT | alanine-glyoxylate aminotransferase | 10289 | -0.003 | -0.0760 | No |
| 58 | SERPINA6 | serpin peptidase inhibitor, clade A (alpha-1 antiproteinase, antitrypsin), member 6 | 10647 | -0.005 | -0.0934 | No |
| 59 | ABCD1 | ATP binding cassette subfamily D member 1 | 10738 | -0.006 | -0.0970 | No |
| 60 | SLC35B2 | solute carrier family 35 (adenosine 3-phospho 5-phosphosulfate transporter), member B2 | 10827 | -0.006 | -0.1003 | No |
| 61 | ABCA8 | ATP binding cassette subfamily A member 8 | 10837 | -0.006 | -0.0996 | No |
| 62 | HACL1 | 2-hydroxyacyl-CoA lyase 1 | 11435 | -0.010 | -0.1286 | No |
| 63 | PEX12 | peroxisomal biogenesis factor 12 | 11466 | -0.010 | -0.1283 | No |
| 64 | NR1I2 | nuclear receptor subfamily 1, group I, member 2 | 11525 | -0.010 | -0.1294 | No |
| 65 | KLF1 | Kruppel-like factor 1 (erythroid) | 11528 | -0.010 | -0.1276 | No |
| 66 | NR0B2 | nuclear receptor subfamily 0, group B, member 2 | 11794 | -0.012 | -0.1391 | No |
| 67 | GSTK1 | glutathione S-transferase kappa 1 | 11816 | -0.012 | -0.1379 | No |
| 68 | RBP1 | retinol binding protein 1, cellular | 11879 | -0.012 | -0.1388 | No |
| 69 | CAT | catalase | 12431 | -0.016 | -0.1642 | No |
| 70 | HSD17B4 | hydroxysteroid (17-beta) dehydrogenase 4 | 12798 | -0.018 | -0.1797 | No |
| 71 | OPTN | optineurin | 13509 | -0.023 | -0.2120 | No |
| 72 | PEX13 | peroxisomal biogenesis factor 13 | 13577 | -0.023 | -0.2111 | No |
| 73 | PEX6 | peroxisomal biogenesis factor 6 | 13791 | -0.025 | -0.2174 | No |
| 74 | PEX19 | peroxisomal biogenesis factor 19 | 13802 | -0.025 | -0.2132 | No |
| 75 | PEX1 | peroxisomal biogenesis factor 1 | 14165 | -0.028 | -0.2266 | No |
| 76 | TTR | transthyretin | 14235 | -0.028 | -0.2248 | No |
| 77 | GCLM | glutamate-cysteine ligase, modifier subunit | 14668 | -0.032 | -0.2411 | No |
| 78 | AR | androgen receptor | 14739 | -0.032 | -0.2386 | No |
| 79 | AKR1D1 | aldo-keto reductase family 1, member D1 | 14778 | -0.033 | -0.2343 | No |
| 80 | IDH1 | isocitrate dehydrogenase 1 (NADP+) | 14950 | -0.034 | -0.2367 | No |
| 81 | SOAT2 | sterol O-acyltransferase 2 | 14999 | -0.034 | -0.2327 | No |
| 82 | ALDH8A1 | aldehyde dehydrogenase 8 family, member A1 | 15008 | -0.035 | -0.2265 | No |
| 83 | BMP6 | bone morphogenetic protein 6 | 15013 | -0.035 | -0.2202 | No |
| 84 | SULT1B1 | sulfotransferase family 1B member 1 | 15132 | -0.036 | -0.2196 | No |
| 85 | DIO2 | deiodinase, iodothyronine, type II | 15760 | -0.041 | -0.2440 | No |
| 86 | APOA1 | apolipoprotein A-I | 15892 | -0.043 | -0.2427 | No |
| 87 | ABCA1 | ATP binding cassette subfamily A member 1 | 15924 | -0.043 | -0.2362 | No |
| 88 | ABCA4 | ATP binding cassette subfamily A member 4 | 16234 | -0.046 | -0.2434 | No |
| 89 | TFCP2L1 | transcription factor CP2-like 1 | 16585 | -0.051 | -0.2517 | Yes |
| 90 | NPC1 | Niemann-Pick disease, type C1 | 16604 | -0.051 | -0.2429 | Yes |
| 91 | SLC23A1 | solute carrier family 23 (ascorbic acid transporter), member 1 | 16629 | -0.052 | -0.2344 | Yes |
| 92 | AMACR | alpha-methylacyl-CoA racemase | 16704 | -0.053 | -0.2282 | Yes |
| 93 | PRDX5 | peroxiredoxin 5 | 16745 | -0.053 | -0.2203 | Yes |
| 94 | PHYH | phytanoyl-CoA 2-hydroxylase | 16804 | -0.054 | -0.2131 | Yes |
| 95 | BCAR3 | breast cancer anti-estrogen resistance 3 | 16807 | -0.054 | -0.2029 | Yes |
| 96 | HAO1 | hydroxyacid oxidase (glycolate oxidase) 1 | 16922 | -0.056 | -0.1983 | Yes |
| 97 | ABCA6 | ATP binding cassette subfamily A member 6 | 17154 | -0.059 | -0.1990 | Yes |
| 98 | SULT2B1 | sulfotransferase family 2B member 1 | 17511 | -0.066 | -0.2049 | Yes |
| 99 | SLC22A18 | solute carrier family 22, member 18 | 17665 | -0.069 | -0.1998 | Yes |
| 100 | RETSAT | retinol saturase (all-trans-retinol 13,14-reductase) | 17683 | -0.069 | -0.1876 | Yes |
| 101 | EFHC1 | EF-hand domain (C-terminal) containing 1 | 17875 | -0.074 | -0.1835 | Yes |
| 102 | ABCA5 | ATP binding cassette subfamily A member 5 | 18148 | -0.081 | -0.1822 | Yes |
| 103 | AQP9 | aquaporin 9 | 18219 | -0.083 | -0.1701 | Yes |
| 104 | NR3C2 | nuclear receptor subfamily 3, group C, member 2 | 18271 | -0.085 | -0.1567 | Yes |
| 105 | EPHX2 | epoxide hydrolase 2, cytoplasmic | 18373 | -0.088 | -0.1453 | Yes |
| 106 | SLC23A2 | solute carrier family 23 (ascorbic acid transporter), member 2 | 18659 | -0.100 | -0.1411 | Yes |
| 107 | PNPLA8 | patatin-like phospholipase domain containing 8 | 18942 | -0.120 | -0.1329 | Yes |
| 108 | ATXN1 | ataxin 1 | 19050 | -0.133 | -0.1131 | Yes |
| 109 | PIPOX | pipecolic acid oxidase | 19131 | -0.147 | -0.0895 | Yes |
| 110 | CYP39A1 | cytochrome P450, family 39, subfamily A, polypeptide 1 | 19213 | -0.165 | -0.0626 | Yes |
| 111 | PECR | peroxisomal trans-2-enoyl-CoA reductase | 19295 | -0.192 | -0.0305 | Yes |
| 112 | CROT | carnitine O-octanoyltransferase | 19375 | -0.223 | 0.0077 | Yes |
Table: GSEA details [plain text format]

  

Fig 2: HALLMARK\_BILE\_ACID\_METABOLISM      
 Blue-Pink O' Gram in the Space of the Analyzed GeneSet

  

Fig 3: HALLMARK\_BILE\_ACID\_METABOLISM: Random ES distribution      
 Gene set null distribution of ES for **HALLMARK\_BILE\_ACID\_METABOLISM**

  
