## Supplementary material for "Somatic hypomethylation of pericentromeric SST1 repeats and tetraploidization in human colorectal cancer cells": GSEA results: HALLMARK_CHOLESTEROL_HOMEOSTASIS.html

Details for gene set HALLMARK\_CHOLESTEROL\_HOMEOSTASIS[GSEA]

|  || Dataset | eset\_byprobe\_collapsed\_to\_symbols.Diploid\_vs\_Tetraploid.cls #Tetraploid\_versus\_Diploid.Diploid\_vs\_Tetraploid.cls #Tetraploid\_versus\_Diploid\_repos |
| Phenotype | Diploid\_vs\_Tetraploid.cls#Tetraploid\_versus\_Diploid\_repos |
| Upregulated in class | Tetraploid |
| GeneSet | HALLMARK\_CHOLESTEROL\_HOMEOSTASIS |
| Enrichment Score (ES) | 0.4552118 |
| Normalized Enrichment Score (NES) | 1.7590891 |
| Nominal p-value | 0.0 |
| FDR q-value | 0.0022598063 |
| FWER p-Value | 0.007 |
Table: GSEA Results Summary

  

Fig 1: Enrichment plot: HALLMARK\_CHOLESTEROL\_HOMEOSTASIS      
 Profile of the Running ES Score & Positions of GeneSet Members on the Rank Ordered List

  

| SYMBOL | TITLE | RANK IN GENE LIST | RANK METRIC SCORE | RUNNING ES | CORE ENRICHMENT || 1 | ACAT2 | acetyl-CoA acetyltransferase 2 | 133 | 0.185 | 0.0304 | Yes |
| 2 | SQLE | squalene epoxidase | 167 | 0.166 | 0.0621 | Yes |
| 3 | GLDC | glycine dehydrogenase (decarboxylating) | 283 | 0.136 | 0.0834 | Yes |
| 4 | CHKA | choline kinase alpha | 287 | 0.135 | 0.1105 | Yes |
| 5 | FASN | fatty acid synthase | 293 | 0.135 | 0.1373 | Yes |
| 6 | FADS2 | fatty acid desaturase 2 | 315 | 0.131 | 0.1625 | Yes |
| 7 | HMGCS1 | 3-hydroxy-3-methylglutaryl-CoA synthase 1 (soluble) | 387 | 0.121 | 0.1832 | Yes |
| 8 | ALDOC | aldolase C, fructose-bisphosphate | 407 | 0.118 | 0.2060 | Yes |
| 9 | HSD17B7 | hydroxysteroid (17-beta) dehydrogenase 7 | 425 | 0.117 | 0.2285 | Yes |
| 10 | ANXA13 | annexin A13 | 558 | 0.106 | 0.2430 | Yes |
| 11 | FDPS | farnesyl diphosphate synthase | 701 | 0.097 | 0.2551 | Yes |
| 12 | ALCAM | activated leukocyte cell adhesion molecule | 762 | 0.094 | 0.2709 | Yes |
| 13 | FDFT1 | farnesyl-diphosphate farnesyltransferase 1 | 793 | 0.093 | 0.2880 | Yes |
| 14 | MVD | mevalonate (diphospho) decarboxylase | 856 | 0.090 | 0.3029 | Yes |
| 15 | JAG1 | jagged 1 | 962 | 0.085 | 0.3147 | Yes |
| 16 | EBP | emopamil binding protein (sterol isomerase) | 1093 | 0.081 | 0.3243 | Yes |
| 17 | NSDHL | NAD(P) dependent steroid dehydrogenase-like | 1100 | 0.081 | 0.3403 | Yes |
| 18 | ANTXR2 | anthrax toxin receptor 2 | 1121 | 0.080 | 0.3554 | Yes |
| 19 | PLAUR | plasminogen activator, urokinase receptor | 1355 | 0.075 | 0.3585 | Yes |
| 20 | LSS | lanosterol synthase (2,3-oxidosqualene-lanosterol cyclase) | 1369 | 0.075 | 0.3728 | Yes |
| 21 | TMEM97 | transmembrane protein 97 | 1371 | 0.075 | 0.3878 | Yes |
| 22 | TM7SF2 | transmembrane 7 superfamily member 2 | 1402 | 0.074 | 0.4011 | Yes |
| 23 | FABP5 | fatty acid binding protein 5 (psoriasis-associated) | 1511 | 0.071 | 0.4098 | Yes |
| 24 | PCYT2 | phosphate cytidylyltransferase 2, ethanolamine | 1547 | 0.070 | 0.4222 | Yes |
| 25 | HMGCR | 3-hydroxy-3-methylglutaryl-CoA reductase | 1821 | 0.064 | 0.4211 | Yes |
| 26 | PPARG | peroxisome proliferator-activated receptor gamma | 1849 | 0.064 | 0.4326 | Yes |
| 27 | DHCR7 | 7-dehydrocholesterol reductase | 1877 | 0.064 | 0.4439 | Yes |
| 28 | IDI1 | isopentenyl-diphosphate delta isomerase 1 | 1905 | 0.063 | 0.4552 | Yes |
| 29 | S100A11 | S100 calcium binding protein A11 | 2852 | 0.049 | 0.4165 | No |
| 30 | SCD | stearoyl-CoA desaturase (delta-9-desaturase) | 3227 | 0.045 | 0.4064 | No |
| 31 | GSTM2 | glutathione S-transferase mu 2 (muscle) | 3243 | 0.045 | 0.4147 | No |
| 32 | MVK | mevalonate kinase | 3563 | 0.042 | 0.4067 | No |
| 33 | ETHE1 | ethylmalonic encephalopathy 1 | 3693 | 0.041 | 0.4082 | No |
| 34 | LGALS3 | lectin, galactoside-binding, soluble, 3 | 3918 | 0.038 | 0.4044 | No |
| 35 | ACSS2 | acyl-CoA synthetase short-chain family member 2 | 4076 | 0.037 | 0.4038 | No |
| 36 | ANXA5 | annexin A5 | 5731 | 0.024 | 0.3235 | No |
| 37 | PLSCR1 | phospholipid scramblase 1 | 5829 | 0.023 | 0.3232 | No |
| 38 | FBXO6 | F-box protein 6 | 5894 | 0.023 | 0.3245 | No |
| 39 | GUSB | glucuronidase, beta | 6020 | 0.022 | 0.3224 | No |
| 40 | CYP51A1 | cytochrome P450, family 51, subfamily A, polypeptide 1 | 6319 | 0.020 | 0.3112 | No |
| 41 | ECH1 | enoyl-CoA hydratase 1, peroxisomal | 6583 | 0.018 | 0.3014 | No |
| 42 | TP53INP1 | tumor protein p53 inducible nuclear protein 1 | 6693 | 0.018 | 0.2993 | No |
| 43 | TNFRSF12A | tumor necrosis factor receptor superfamily, member 12A | 6787 | 0.017 | 0.2980 | No |
| 44 | CD9 | CD9 molecule | 6905 | 0.016 | 0.2953 | No |
| 45 | LGMN | legumain | 7134 | 0.015 | 0.2866 | No |
| 46 | ATXN2 | ataxin 2 | 7261 | 0.014 | 0.2830 | No |
| 47 | ACTG1 | actin gamma 1 | 7506 | 0.013 | 0.2731 | No |
| 48 | ABCA2 | ATP binding cassette subfamily A member 2 | 7772 | 0.011 | 0.2617 | No |
| 49 | STARD4 | StAR-related lipid transfer domain containing 4 | 8485 | 0.007 | 0.2265 | No |
| 50 | LDLR | low density lipoprotein receptor | 9447 | 0.002 | 0.1774 | No |
| 51 | ADH4 | alcohol dehydrogenase 4 (class II), pi polypeptide | 9592 | 0.001 | 0.1702 | No |
| 52 | CXCL16 | chemokine (C-X-C motif) ligand 16 | 9619 | 0.001 | 0.1691 | No |
| 53 | SC5D | sterol-C5-desaturase | 9987 | -0.001 | 0.1504 | No |
| 54 | PMVK | phosphomevalonate kinase | 10094 | -0.002 | 0.1453 | No |
| 55 | ERRFI1 | ERBB receptor feedback inhibitor 1 | 10474 | -0.004 | 0.1266 | No |
| 56 | CTNNB1 | catenin (cadherin-associated protein), beta 1 | 11452 | -0.010 | 0.0784 | No |
| 57 | MAL2 | mal, T-cell differentiation protein 2 (gene/pseudogene) | 12533 | -0.016 | 0.0261 | No |
| 58 | SREBF2 | sterol regulatory element binding transcription factor 2 | 13267 | -0.021 | -0.0074 | No |
| 59 | LPL | lipoprotein lipase | 13793 | -0.025 | -0.0294 | No |
| 60 | PNRC1 | proline-rich nuclear receptor coactivator 1 | 15332 | -0.037 | -0.1009 | No |
| 61 | CBS | cystathionine-beta-synthase | 16810 | -0.054 | -0.1660 | No |
| 62 | STX5 | syntaxin 5 | 16918 | -0.055 | -0.1604 | No |
| 63 | AVPR1A | arginine vasopressin receptor 1A | 17181 | -0.060 | -0.1618 | No |
| 64 | SEMA3B | sema domain, immunoglobulin domain (Ig), short basic domain, secreted, (semaphorin) 3B | 17591 | -0.067 | -0.1694 | No |
| 65 | PDK3 | pyruvate dehydrogenase kinase, isozyme 3 | 18210 | -0.083 | -0.1845 | No |
| 66 | NFIL3 | nuclear factor, interleukin 3 regulated | 18475 | -0.092 | -0.1797 | No |
| 67 | CPEB2 | cytoplasmic polyadenylation element binding protein 2 | 18574 | -0.096 | -0.1655 | No |
| 68 | ATF5 | activating transcription factor 5 | 18851 | -0.114 | -0.1568 | No |
| 69 | TRIB3 | tribbles pseudokinase 3 | 18870 | -0.115 | -0.1345 | No |
| 70 | GPX8 | glutathione peroxidase 8 (putative) | 18960 | -0.123 | -0.1145 | No |
| 71 | CLU | clusterin | 19343 | -0.205 | -0.0929 | No |
| 72 | GNAI1 | guanine nucleotide binding protein (G protein), alpha inhibiting activity polypeptide 1 | 19420 | -0.254 | -0.0458 | No |
| 73 | ATF3 | activating transcription factor 3 | 19421 | -0.254 | 0.0053 | No |
Table: GSEA details [plain text format]

  

Fig 2: HALLMARK\_CHOLESTEROL\_HOMEOSTASIS      
 Blue-Pink O' Gram in the Space of the Analyzed GeneSet

  

Fig 3: HALLMARK\_CHOLESTEROL\_HOMEOSTASIS: Random ES distribution      
 Gene set null distribution of ES for **HALLMARK\_CHOLESTEROL\_HOMEOSTASIS**

  
