## Supplementary material for "Somatic hypomethylation of pericentromeric SST1 repeats and tetraploidization in human colorectal cancer cells": GSEA results: HALLMARK_COAGULATION.html

Details for gene set HALLMARK\_COAGULATION[GSEA]

|  || Dataset | eset\_byprobe\_collapsed\_to\_symbols.Diploid\_vs\_Tetraploid.cls #Tetraploid\_versus\_Diploid.Diploid\_vs\_Tetraploid.cls #Tetraploid\_versus\_Diploid\_repos |
| Phenotype | Diploid\_vs\_Tetraploid.cls#Tetraploid\_versus\_Diploid\_repos |
| Upregulated in class | Diploid |
| GeneSet | HALLMARK\_COAGULATION |
| Enrichment Score (ES) | -0.32241604 |
| Normalized Enrichment Score (NES) | -1.2722856 |
| Nominal p-value | 0.0830721 |
| FDR q-value | 0.28016058 |
| FWER p-Value | 0.934 |
Table: GSEA Results Summary

  

Fig 1: Enrichment plot: HALLMARK\_COAGULATION      
 Profile of the Running ES Score & Positions of GeneSet Members on the Rank Ordered List

  

| SYMBOL | TITLE | RANK IN GENE LIST | RANK METRIC SCORE | RUNNING ES | CORE ENRICHMENT || 1 | CTSE | cathepsin E | 9 | 0.419 | 0.0492 | No |
| 2 | F3 | coagulation factor III (thromboplastin, tissue factor) | 119 | 0.191 | 0.0661 | No |
| 3 | C3 | complement component 3 | 143 | 0.179 | 0.0862 | No |
| 4 | CFB | complement factor B | 164 | 0.167 | 0.1049 | No |
| 5 | GDA | guanine deaminase | 174 | 0.164 | 0.1238 | No |
| 6 | THBS1 | thrombospondin 1 | 192 | 0.158 | 0.1416 | No |
| 7 | TIMP1 | TIMP metallopeptidase inhibitor 1 | 254 | 0.141 | 0.1552 | No |
| 8 | PROS1 | protein S (alpha) | 556 | 0.106 | 0.1522 | No |
| 9 | GP9 | glycoprotein IX (platelet) | 580 | 0.104 | 0.1633 | No |
| 10 | DUSP6 | dual specificity phosphatase 6 | 638 | 0.101 | 0.1723 | No |
| 11 | MASP2 | mannan-binding lectin serine peptidase 2 | 988 | 0.085 | 0.1644 | No |
| 12 | GP1BA | glycoprotein Ib (platelet), alpha polypeptide | 1000 | 0.084 | 0.1738 | No |
| 13 | C9 | complement component 9 | 1024 | 0.083 | 0.1825 | No |
| 14 | HNF4A | hepatocyte nuclear factor 4, alpha | 1031 | 0.083 | 0.1920 | No |
| 15 | F11 | coagulation factor XI | 1205 | 0.078 | 0.1923 | No |
| 16 | F10 | coagulation factor X | 1340 | 0.075 | 0.1943 | No |
| 17 | KLF7 | Kruppel-like factor 7 (ubiquitous) | 1362 | 0.075 | 0.2021 | No |
| 18 | A2M | alpha-2-macroglobulin | 1412 | 0.074 | 0.2083 | No |
| 19 | S100A13 | S100 calcium binding protein A13 | 1525 | 0.071 | 0.2109 | No |
| 20 | ACOX2 | acyl-CoA oxidase 2, branched chain | 1687 | 0.067 | 0.2105 | No |
| 21 | TMPRSS6 | transmembrane protease, serine 6 | 1757 | 0.066 | 0.2148 | No |
| 22 | CFD | complement factor D (adipsin) | 1759 | 0.066 | 0.2225 | No |
| 23 | LEFTY2 | left-right determination factor 2 | 1775 | 0.065 | 0.2294 | No |
| 24 | CTSH | cathepsin H | 1817 | 0.064 | 0.2350 | No |
| 25 | CAPN2 | calpain 2, (m/II) large subunit | 2066 | 0.060 | 0.2293 | No |
| 26 | SERPING1 | serpin peptidase inhibitor, clade G (C1 inhibitor), member 1 | 2131 | 0.059 | 0.2331 | No |
| 27 | HTRA1 | HtrA serine peptidase 1 | 2295 | 0.057 | 0.2314 | No |
| 28 | CTSV | cathepsin V | 2384 | 0.056 | 0.2334 | No |
| 29 | PF4 | platelet factor 4 | 2778 | 0.050 | 0.2191 | No |
| 30 | MAFF | v-maf avian musculoaponeurotic fibrosarcoma oncogene homolog F | 3075 | 0.047 | 0.2094 | No |
| 31 | CFH | complement factor H | 3276 | 0.045 | 0.2043 | No |
| 32 | F13B | coagulation factor XIII, B polypeptide | 3312 | 0.045 | 0.2078 | No |
| 33 | MMP9 | matrix metallopeptidase 9 | 3774 | 0.040 | 0.1888 | No |
| 34 | ISCU | iron-sulfur cluster assembly enzyme | 4109 | 0.037 | 0.1759 | No |
| 35 | CPN1 | carboxypeptidase N, polypeptide 1 | 4269 | 0.035 | 0.1718 | No |
| 36 | HPN | hepsin | 4274 | 0.035 | 0.1757 | No |
| 37 | PLG | plasminogen | 4359 | 0.034 | 0.1754 | No |
| 38 | C8G | complement component 8, gamma polypeptide | 4410 | 0.034 | 0.1769 | No |
| 39 | F2RL2 | coagulation factor II (thrombin) receptor-like 2 | 4564 | 0.033 | 0.1729 | No |
| 40 | ITIH1 | inter-alpha-trypsin inhibitor heavy chain 1 | 4587 | 0.032 | 0.1756 | No |
| 41 | TIMP3 | TIMP metallopeptidase inhibitor 3 | 4889 | 0.030 | 0.1636 | No |
| 42 | MMP15 | matrix metallopeptidase 15 (membrane-inserted) | 5146 | 0.028 | 0.1537 | No |
| 43 | PLAU | plasminogen activator, urokinase | 5343 | 0.027 | 0.1468 | No |
| 44 | MBL2 | mannose-binding lectin (protein C) 2, soluble | 5624 | 0.025 | 0.1352 | No |
| 45 | PREP | prolyl endopeptidase | 5645 | 0.024 | 0.1371 | No |
| 46 | SERPINB2 | serpin peptidase inhibitor, clade B (ovalbumin), member 2 | 6047 | 0.022 | 0.1190 | No |
| 47 | KLK8 | kallikrein related peptidase 8 | 6224 | 0.021 | 0.1124 | No |
| 48 | VWF | von Willebrand factor | 6320 | 0.020 | 0.1099 | No |
| 49 | PROC | protein C (inactivator of coagulation factors Va and VIIIa) | 6402 | 0.020 | 0.1080 | No |
| 50 | C2 | complement component 2 | 6731 | 0.017 | 0.0932 | No |
| 51 | CD9 | CD9 molecule | 6905 | 0.016 | 0.0862 | No |
| 52 | MSRB2 | methionine sulfoxide reductase B2 | 7001 | 0.016 | 0.0832 | No |
| 53 | LGMN | legumain | 7134 | 0.015 | 0.0781 | No |
| 54 | LTA4H | leukotriene A4 hydrolase | 7229 | 0.015 | 0.0750 | No |
| 55 | DUSP14 | dual specificity phosphatase 14 | 7231 | 0.015 | 0.0767 | No |
| 56 | COMP | cartilage oligomeric matrix protein | 7536 | 0.013 | 0.0625 | No |
| 57 | BMP1 | bone morphogenetic protein 1 | 7539 | 0.013 | 0.0639 | No |
| 58 | FYN | FYN proto-oncogene, Src family tyrosine kinase | 7712 | 0.012 | 0.0564 | No |
| 59 | F2 | coagulation factor II (thrombin) | 8109 | 0.009 | 0.0371 | No |
| 60 | OLR1 | oxidized low density lipoprotein (lectin-like) receptor 1 | 8290 | 0.008 | 0.0288 | No |
| 61 | HMGCS2 | 3-hydroxy-3-methylglutaryl-CoA synthase 2 (mitochondrial) | 8539 | 0.007 | 0.0168 | No |
| 62 | S100A1 | S100 calcium binding protein A1 | 8573 | 0.006 | 0.0158 | No |
| 63 | HRG | histidine-rich glycoprotein | 8796 | 0.005 | 0.0050 | No |
| 64 | RGN | regucalcin | 8876 | 0.005 | 0.0015 | No |
| 65 | CSRP1 | cysteine and glycine-rich protein 1 | 8988 | 0.004 | -0.0038 | No |
| 66 | C1R | complement component 1, r subcomponent | 9028 | 0.004 | -0.0053 | No |
| 67 | MMP14 | matrix metallopeptidase 14 (membrane-inserted) | 9110 | 0.003 | -0.0091 | No |
| 68 | P2RY1 | purinergic receptor P2Y, G-protein coupled, 1 | 9564 | 0.001 | -0.0323 | No |
| 69 | LRP1 | LDL receptor related protein 1 | 9659 | 0.001 | -0.0371 | No |
| 70 | MMP8 | matrix metallopeptidase 8 | 9856 | -0.000 | -0.0471 | No |
| 71 | SIRT2 | sirtuin 2 | 10011 | -0.001 | -0.0549 | No |
| 72 | PLEK | pleckstrin | 10164 | -0.002 | -0.0625 | No |
| 73 | RAC1 | ras-related C3 botulinum toxin substrate 1 (rho family, small GTP binding protein Rac1) | 10275 | -0.003 | -0.0679 | No |
| 74 | APOC3 | apolipoprotein C-III | 10379 | -0.003 | -0.0728 | No |
| 75 | F8 | coagulation factor VIII, procoagulant component | 10407 | -0.004 | -0.0738 | No |
| 76 | C1QA | complement component 1, q subcomponent, A chain | 10470 | -0.004 | -0.0765 | No |
| 77 | CTSK | cathepsin K | 10505 | -0.004 | -0.0778 | No |
| 78 | CPQ | carboxypeptidase Q | 10570 | -0.004 | -0.0805 | No |
| 79 | TF | transferrin | 10665 | -0.005 | -0.0848 | No |
| 80 | MMP3 | matrix metallopeptidase 3 | 10688 | -0.005 | -0.0853 | No |
| 81 | PLAT | plasminogen activator, tissue | 10760 | -0.006 | -0.0883 | No |
| 82 | C8A | complement component 8, alpha polypeptide | 10861 | -0.006 | -0.0927 | No |
| 83 | SH2B2 | SH2B adaptor protein 2 | 11007 | -0.007 | -0.0993 | No |
| 84 | THBD | thrombomodulin | 11144 | -0.008 | -0.1054 | No |
| 85 | ADAM9 | ADAM metallopeptidase domain 9 | 11696 | -0.011 | -0.1325 | No |
| 86 | APOC2 | apolipoprotein C-II | 12395 | -0.015 | -0.1667 | No |
| 87 | C8B | complement component 8, beta polypeptide | 12554 | -0.016 | -0.1729 | No |
| 88 | WDR1 | Jeck2013 ANTISENSE, CDS, coding, INTERNAL, intronic, OVCODE, OVEXON best transcript NM\_017491 | 12674 | -0.017 | -0.1770 | No |
| 89 | PRSS23 | protease, serine, 23 | 12809 | -0.018 | -0.1818 | No |
| 90 | ITGB3 | integrin beta 3 | 12821 | -0.018 | -0.1802 | No |
| 91 | CPB2 | carboxypeptidase B2 (plasma) | 12886 | -0.018 | -0.1813 | No |
| 92 | GNB2 | guanine nucleotide binding protein (G protein), beta polypeptide 2 | 12915 | -0.019 | -0.1806 | No |
| 93 | SERPINE1 | serpin peptidase inhibitor, clade E (nexin, plasminogen activator inhibitor type 1), member 1 | 12974 | -0.019 | -0.1813 | No |
| 94 | APOC1 | apolipoprotein C-I | 13064 | -0.020 | -0.1836 | No |
| 95 | RABIF | RAB interacting factor | 13273 | -0.021 | -0.1918 | No |
| 96 | F12 | coagulation factor XII (Hageman factor) | 13443 | -0.022 | -0.1979 | No |
| 97 | C1S | complement component 1, s subcomponent | 13614 | -0.023 | -0.2039 | No |
| 98 | MMP10 | matrix metallopeptidase 10 | 13819 | -0.025 | -0.2115 | No |
| 99 | CFI | complement factor I | 13875 | -0.025 | -0.2113 | No |
| 100 | PROZ | protein Z, vitamin K-dependent plasma glycoprotein | 14407 | -0.030 | -0.2352 | No |
| 101 | TFPI2 | tissue factor pathway inhibitor 2 | 14489 | -0.030 | -0.2357 | No |
| 102 | KLKB1 | kallikrein B1 | 14519 | -0.030 | -0.2336 | No |
| 103 | CTSB | cathepsin B | 14720 | -0.032 | -0.2401 | No |
| 104 | ARF4 | ADP-ribosylation factor 4 | 15140 | -0.036 | -0.2575 | No |
| 105 | ITGA2 | integrin, alpha 2 (CD49B, alpha 2 subunit of VLA-2 receptor) | 15498 | -0.039 | -0.2714 | No |
| 106 | PDGFB | platelet-derived growth factor beta polypeptide | 15682 | -0.041 | -0.2760 | No |
| 107 | APOA1 | apolipoprotein A-I | 15892 | -0.043 | -0.2817 | No |
| 108 | GNG12 | guanine nucleotide binding protein (G protein), gamma 12 | 16021 | -0.044 | -0.2831 | No |
| 109 | MMP7 | matrix metallopeptidase 7 | 16218 | -0.046 | -0.2878 | No |
| 110 | PECAM1 | platelet/endothelial cell adhesion molecule 1 | 16255 | -0.047 | -0.2841 | No |
| 111 | FBN1 | fibrillin 1 | 16292 | -0.047 | -0.2804 | No |
| 112 | GSN | gelsolin | 16387 | -0.048 | -0.2795 | No |
| 113 | SERPINC1 | serpin peptidase inhibitor, clade C (antithrombin), member 1 | 16408 | -0.048 | -0.2748 | No |
| 114 | CASP9 | caspase 9 | 16638 | -0.052 | -0.2805 | No |
| 115 | F9 | coagulation factor IX | 17281 | -0.061 | -0.3063 | No |
| 116 | CRIP2 | cysteine-rich protein 2 | 17549 | -0.066 | -0.3123 | No |
| 117 | LAMP2 | lysosomal-associated membrane protein 2 | 17747 | -0.071 | -0.3140 | Yes |
| 118 | ANXA1 | annexin A1 | 17825 | -0.073 | -0.3094 | Yes |
| 119 | PEF1 | penta-EF-hand domain containing 1 | 17850 | -0.073 | -0.3020 | Yes |
| 120 | CAPN5 | calpain 5 | 17885 | -0.074 | -0.2950 | Yes |
| 121 | FURIN | furin (paired basic amino acid cleaving enzyme) | 18023 | -0.077 | -0.2929 | Yes |
| 122 | MEP1A | meprin A, alpha (PABA peptide hydrolase) | 18452 | -0.091 | -0.3042 | Yes |
| 123 | RAPGEF3 | Rap guanine nucleotide exchange factor 3 | 18527 | -0.094 | -0.2969 | Yes |
| 124 | SPARC | secreted protein, acidic, cysteine-rich (osteonectin) | 18622 | -0.097 | -0.2902 | Yes |
| 125 | MMP11 | matrix metallopeptidase 11 | 18635 | -0.098 | -0.2792 | Yes |
| 126 | FGA | fibrinogen alpha chain | 18667 | -0.101 | -0.2689 | Yes |
| 127 | MST1 | macrophage stimulating 1 | 18726 | -0.104 | -0.2595 | Yes |
| 128 | USP11 | ubiquitin specific peptidase 11 | 18992 | -0.126 | -0.2583 | Yes |
| 129 | DCT | dopachrome tautomerase | 19008 | -0.128 | -0.2439 | Yes |
| 130 | MMP2 | matrix metallopeptidase 2 | 19166 | -0.152 | -0.2339 | Yes |
| 131 | MMP1 | matrix metallopeptidase 1 | 19272 | -0.181 | -0.2179 | Yes |
| 132 | SERPINA1 | serpin peptidase inhibitor, clade A (alpha-1 antiproteinase, antitrypsin), member 1 | 19288 | -0.189 | -0.1962 | Yes |
| 133 | DPP4 | dipeptidyl-peptidase 4 | 19294 | -0.192 | -0.1738 | Yes |
| 134 | CLU | clusterin | 19343 | -0.205 | -0.1520 | Yes |
| 135 | FN1 | fibronectin 1 | 19439 | -0.280 | -0.1237 | Yes |
| 136 | FGG | fibrinogen gamma chain | 19473 | -0.330 | -0.0863 | Yes |
| 137 | CTSO | cathepsin O | 19481 | -0.342 | -0.0462 | Yes |
| 138 | ANG | angiogenin, ribonuclease, RNase A family, 5 | 19501 | -0.408 | 0.0012 | Yes |
Table: GSEA details [plain text format]

  

Fig 2: HALLMARK\_COAGULATION      
 Blue-Pink O' Gram in the Space of the Analyzed GeneSet

  

Fig 3: HALLMARK\_COAGULATION: Random ES distribution      
 Gene set null distribution of ES for **HALLMARK\_COAGULATION**

  
