## Supplementary material for "Somatic hypomethylation of pericentromeric SST1 repeats and tetraploidization in human colorectal cancer cells": GSEA results: HALLMARK_COMPLEMENT.html

Details for gene set HALLMARK\_COMPLEMENT[GSEA]

|  || Dataset | eset\_byprobe\_collapsed\_to\_symbols.Diploid\_vs\_Tetraploid.cls #Tetraploid\_versus\_Diploid.Diploid\_vs\_Tetraploid.cls #Tetraploid\_versus\_Diploid\_repos |
| Phenotype | Diploid\_vs\_Tetraploid.cls#Tetraploid\_versus\_Diploid\_repos |
| Upregulated in class | Diploid |
| GeneSet | HALLMARK\_COMPLEMENT |
| Enrichment Score (ES) | -0.36690617 |
| Normalized Enrichment Score (NES) | -1.5116382 |
| Nominal p-value | 0.0015432099 |
| FDR q-value | 0.07056172 |
| FWER p-Value | 0.255 |
Table: GSEA Results Summary

  

Fig 1: Enrichment plot: HALLMARK\_COMPLEMENT      
 Profile of the Running ES Score & Positions of GeneSet Members on the Rank Ordered List

  

| SYMBOL | TITLE | RANK IN GENE LIST | RANK METRIC SCORE | RUNNING ES | CORE ENRICHMENT || 1 | CA2 | carbonic anhydrase II | 69 | 0.250 | 0.0187 | No |
| 2 | F3 | coagulation factor III (thromboplastin, tissue factor) | 119 | 0.191 | 0.0332 | No |
| 3 | HSPA1A | heat shock 70kDa protein 1A | 129 | 0.186 | 0.0494 | No |
| 4 | C3 | complement component 3 | 143 | 0.179 | 0.0647 | No |
| 5 | CFB | complement factor B | 164 | 0.167 | 0.0785 | No |
| 6 | IRF7 | interferon regulatory factor 7 | 230 | 0.148 | 0.0883 | No |
| 7 | TIMP1 | TIMP metallopeptidase inhibitor 1 | 254 | 0.141 | 0.0997 | No |
| 8 | GPD2 | glycerol-3-phosphate dehydrogenase 2 | 534 | 0.107 | 0.0949 | No |
| 9 | GP9 | glycoprotein IX (platelet) | 580 | 0.104 | 0.1019 | No |
| 10 | DUSP6 | dual specificity phosphatase 6 | 638 | 0.101 | 0.1079 | No |
| 11 | FCER1G | Fc fragment of IgE, high affinity I, receptor for; gamma polypeptide | 654 | 0.100 | 0.1161 | No |
| 12 | CXCL1 | chemokine (C-X-C motif) ligand 1 (melanoma growth stimulating activity, alpha) | 672 | 0.098 | 0.1240 | No |
| 13 | GP1BA | glycoprotein Ib (platelet), alpha polypeptide | 1000 | 0.084 | 0.1146 | No |
| 14 | C9 | complement component 9 | 1024 | 0.083 | 0.1208 | No |
| 15 | HNF4A | hepatocyte nuclear factor 4, alpha | 1031 | 0.083 | 0.1279 | No |
| 16 | IRF2 | interferon regulatory factor 2 | 1032 | 0.083 | 0.1354 | No |
| 17 | F10 | coagulation factor X | 1340 | 0.075 | 0.1262 | No |
| 18 | PLAUR | plasminogen activator, urokinase receptor | 1355 | 0.075 | 0.1322 | No |
| 19 | S100A13 | S100 calcium binding protein A13 | 1525 | 0.071 | 0.1297 | No |
| 20 | APOA4 | apolipoprotein A-IV | 1692 | 0.067 | 0.1271 | No |
| 21 | TMPRSS6 | transmembrane protease, serine 6 | 1757 | 0.066 | 0.1296 | No |
| 22 | CTSH | cathepsin H | 1817 | 0.064 | 0.1323 | No |
| 23 | SERPING1 | serpin peptidase inhibitor, clade G (C1 inhibitor), member 1 | 2131 | 0.059 | 0.1215 | No |
| 24 | CDA | cytidine deaminase | 2157 | 0.059 | 0.1254 | No |
| 25 | MSRB1 | methionine sulfoxide reductase B1 | 2209 | 0.058 | 0.1280 | No |
| 26 | SIRT6 | sirtuin 6 | 2289 | 0.057 | 0.1290 | No |
| 27 | CTSV | cathepsin V | 2384 | 0.056 | 0.1291 | No |
| 28 | DOCK9 | dedicator of cytokinesis 9 | 2391 | 0.055 | 0.1337 | No |
| 29 | MAFF | v-maf avian musculoaponeurotic fibrosarcoma oncogene homolog F | 3075 | 0.047 | 0.1025 | No |
| 30 | SCG3 | secretogranin III | 3098 | 0.047 | 0.1056 | No |
| 31 | ZEB1 | zinc finger E-box binding homeobox 1 | 3146 | 0.046 | 0.1072 | No |
| 32 | ATOX1 | antioxidant 1 copper chaperone | 3164 | 0.046 | 0.1105 | No |
| 33 | CFH | complement factor H | 3276 | 0.045 | 0.1087 | No |
| 34 | S100A9 | S100 calcium binding protein A9 | 3554 | 0.042 | 0.0981 | No |
| 35 | VCPIP1 | valosin containing protein (p97)/p47 complex interacting protein 1 | 3762 | 0.040 | 0.0910 | No |
| 36 | DOCK10 | dedicator of cytokinesis 10 | 3850 | 0.039 | 0.0900 | No |
| 37 | LGALS3 | lectin, galactoside-binding, soluble, 3 | 3918 | 0.038 | 0.0899 | No |
| 38 | DGKG | diacylglycerol kinase gamma | 3927 | 0.038 | 0.0929 | No |
| 39 | CALM3 | calmodulin 3 (phosphorylase kinase, delta) | 4098 | 0.037 | 0.0874 | No |
| 40 | ITGAM | integrin, alpha M (complement component 3 receptor 3 subunit) | 4320 | 0.035 | 0.0790 | No |
| 41 | PLG | plasminogen | 4359 | 0.034 | 0.0801 | No |
| 42 | APOBEC3G | apolipoprotein B mRNA editing enzyme, catalytic polypeptide-like 3G | 4468 | 0.033 | 0.0775 | No |
| 43 | COL4A2 | collagen, type IV, alpha 2 | 4487 | 0.033 | 0.0796 | No |
| 44 | GZMB | granzyme B | 4513 | 0.033 | 0.0812 | No |
| 45 | ITIH1 | inter-alpha-trypsin inhibitor heavy chain 1 | 4587 | 0.032 | 0.0803 | No |
| 46 | PRSS36 | protease, serine 36 | 4657 | 0.032 | 0.0796 | No |
| 47 | CASP3 | caspase 3 | 4699 | 0.032 | 0.0803 | No |
| 48 | CR1 | complement component (3b/4b) receptor 1 (Knops blood group) | 4808 | 0.031 | 0.0775 | No |
| 49 | PRSS3 | protease, serine, 3 | 4890 | 0.030 | 0.0760 | No |
| 50 | GATA3 | GATA binding protein 3 | 5072 | 0.029 | 0.0692 | No |
| 51 | MMP15 | matrix metallopeptidase 15 (membrane-inserted) | 5146 | 0.028 | 0.0679 | No |
| 52 | CDK5R1 | cyclin-dependent kinase 5, regulatory subunit 1 (p35) | 5166 | 0.028 | 0.0694 | No |
| 53 | LCP2 | lymphocyte cytosolic protein 2 | 5232 | 0.027 | 0.0685 | No |
| 54 | GNGT2 | guanine nucleotide binding protein (G protein), gamma transducing activity polypeptide 2 | 5494 | 0.025 | 0.0573 | No |
| 55 | CALM1 | calmodulin 1 (phosphorylase kinase, delta) | 5557 | 0.025 | 0.0563 | No |
| 56 | RASGRP1 | RAS guanyl releasing protein 1 (calcium and DAG-regulated) | 5558 | 0.025 | 0.0585 | No |
| 57 | PREP | prolyl endopeptidase | 5645 | 0.024 | 0.0563 | No |
| 58 | ANXA5 | annexin A5 | 5731 | 0.024 | 0.0540 | No |
| 59 | PLSCR1 | phospholipid scramblase 1 | 5829 | 0.023 | 0.0510 | No |
| 60 | SERPINB2 | serpin peptidase inhibitor, clade B (ovalbumin), member 2 | 6047 | 0.022 | 0.0418 | No |
| 61 | LCK | LCK proto-oncogene, Src family tyrosine kinase | 6061 | 0.022 | 0.0430 | No |
| 62 | SPOCK2 | sparc/osteonectin, cwcv and kazal-like domains proteoglycan (testican) 2 | 6154 | 0.021 | 0.0401 | No |
| 63 | CD59 | CD59 molecule, complement regulatory protein | 6466 | 0.019 | 0.0258 | No |
| 64 | C2 | complement component 2 | 6731 | 0.017 | 0.0137 | No |
| 65 | CP | ceruloplasmin (ferroxidase) | 6778 | 0.017 | 0.0128 | No |
| 66 | CASP7 | caspase 7 | 6811 | 0.017 | 0.0127 | No |
| 67 | PIK3R5 | phosphoinositide-3-kinase, regulatory subunit 5 | 6911 | 0.016 | 0.0090 | No |
| 68 | LGMN | legumain | 7134 | 0.015 | -0.0011 | No |
| 69 | LTA4H | leukotriene A4 hydrolase | 7229 | 0.015 | -0.0047 | No |
| 70 | PFN1 | profilin 1 | 7240 | 0.015 | -0.0039 | No |
| 71 | CPM | carboxypeptidase M | 7295 | 0.014 | -0.0054 | No |
| 72 | PCSK9 | proprotein convertase subtilisin/kexin type 9 | 7351 | 0.014 | -0.0070 | No |
| 73 | RAF1 | Raf-1 proto-oncogene, serine/threonine kinase | 7383 | 0.014 | -0.0074 | No |
| 74 | F5 | coagulation factor V (proaccelerin, labile factor) | 7447 | 0.013 | -0.0095 | No |
| 75 | MT3 | metallothionein 3 | 7540 | 0.013 | -0.0131 | No |
| 76 | KCNIP2 | Kv channel interacting protein 2 | 7553 | 0.013 | -0.0126 | No |
| 77 | GNB4 | guanine nucleotide binding protein (G protein), beta polypeptide 4 | 7647 | 0.012 | -0.0164 | No |
| 78 | PPP4C | protein phosphatase 4, catalytic subunit | 7648 | 0.012 | -0.0153 | No |
| 79 | FYN | FYN proto-oncogene, Src family tyrosine kinase | 7712 | 0.012 | -0.0175 | No |
| 80 | EHD1 | EH domain containing 1 | 7793 | 0.011 | -0.0207 | No |
| 81 | NOTCH4 | notch 4 | 8040 | 0.010 | -0.0326 | No |
| 82 | F2 | coagulation factor II (thrombin) | 8109 | 0.009 | -0.0353 | No |
| 83 | PSMB9 | proteasome subunit beta 9 | 8144 | 0.009 | -0.0362 | No |
| 84 | CTSC | cathepsin C | 8167 | 0.009 | -0.0366 | No |
| 85 | OLR1 | oxidized low density lipoprotein (lectin-like) receptor 1 | 8290 | 0.008 | -0.0421 | No |
| 86 | CCL5 | chemokine (C-C motif) ligand 5 | 8596 | 0.006 | -0.0574 | No |
| 87 | CSRP1 | cysteine and glycine-rich protein 1 | 8988 | 0.004 | -0.0772 | No |
| 88 | C1R | complement component 1, r subcomponent | 9028 | 0.004 | -0.0789 | No |
| 89 | MMP14 | matrix metallopeptidase 14 (membrane-inserted) | 9110 | 0.003 | -0.0828 | No |
| 90 | AKAP10 | A kinase (PRKA) anchor protein 10 | 9123 | 0.003 | -0.0831 | No |
| 91 | FCN1 | ficolin (collagen/fibrinogen domain containing) 1 | 9130 | 0.003 | -0.0831 | No |
| 92 | HPCAL4 | hippocalcin like 4 | 9325 | 0.002 | -0.0930 | No |
| 93 | XPNPEP1 | X-prolyl aminopeptidase (aminopeptidase P) 1, soluble | 9532 | 0.001 | -0.1035 | No |
| 94 | KYNU | kynureninase | 9602 | 0.001 | -0.1070 | No |
| 95 | LRP1 | LDL receptor related protein 1 | 9659 | 0.001 | -0.1098 | No |
| 96 | MMP8 | matrix metallopeptidase 8 | 9856 | -0.000 | -0.1199 | No |
| 97 | USP14 | ubiquitin specific peptidase 14 (tRNA-guanine transglycosylase) | 9929 | -0.001 | -0.1236 | No |
| 98 | LTF | lactotransferrin | 10005 | -0.001 | -0.1273 | No |
| 99 | PLEK | pleckstrin | 10164 | -0.002 | -0.1353 | No |
| 100 | SH2B3 | SH2B adaptor protein 3 | 10192 | -0.002 | -0.1365 | No |
| 101 | CTSL | cathepsin L | 10220 | -0.002 | -0.1377 | No |
| 102 | F8 | coagulation factor VIII, procoagulant component | 10407 | -0.004 | -0.1470 | No |
| 103 | ADRA2B | adrenoceptor alpha 2B | 10413 | -0.004 | -0.1470 | No |
| 104 | RHOG | ras homolog family member G | 10457 | -0.004 | -0.1488 | No |
| 105 | C1QA | complement component 1, q subcomponent, A chain | 10470 | -0.004 | -0.1491 | No |
| 106 | PSEN1 | presenilin 1 | 10566 | -0.004 | -0.1536 | No |
| 107 | CPQ | carboxypeptidase Q | 10570 | -0.004 | -0.1534 | No |
| 108 | PLAT | plasminogen activator, tissue | 10760 | -0.006 | -0.1627 | No |
| 109 | PIK3CA | phosphatidylinositol-4,5-bisphosphate 3-kinase, catalytic subunit alpha | 10812 | -0.006 | -0.1648 | No |
| 110 | GNAI3 | guanine nucleotide binding protein (G protein), alpha inhibiting activity polypeptide 3 | 10833 | -0.006 | -0.1652 | No |
| 111 | WAS | Wiskott-Aldrich syndrome | 10901 | -0.006 | -0.1681 | No |
| 112 | RNF4 | ring finger protein 4 | 11292 | -0.009 | -0.1875 | No |
| 113 | PIM1 | Pim-1 proto-oncogene, serine/threonine kinase | 11480 | -0.010 | -0.1963 | No |
| 114 | PRCP | prolylcarboxypeptidase | 11568 | -0.010 | -0.1999 | No |
| 115 | ADAM9 | ADAM metallopeptidase domain 9 | 11696 | -0.011 | -0.2055 | No |
| 116 | IL6 | interleukin 6 | 11825 | -0.012 | -0.2111 | No |
| 117 | GRB2 | growth factor receptor bound protein 2 | 11880 | -0.012 | -0.2128 | No |
| 118 | ACTN2 | actinin, alpha 2 | 12029 | -0.013 | -0.2193 | No |
| 119 | STX4 | syntaxin 4 | 12045 | -0.013 | -0.2189 | No |
| 120 | CD55 | CD55 molecule, decay accelerating factor for complement (Cromer blood group) | 12082 | -0.013 | -0.2196 | No |
| 121 | LAP3 | leucine aminopeptidase 3 | 12270 | -0.015 | -0.2280 | No |
| 122 | CD40LG | CD40 ligand | 12277 | -0.015 | -0.2270 | No |
| 123 | GNG2 | guanine nucleotide binding protein (G protein), gamma 2 | 12470 | -0.016 | -0.2355 | No |
| 124 | FDX1 | ferredoxin 1 | 12606 | -0.017 | -0.2410 | No |
| 125 | C4BPB | complement component 4 binding protein, beta | 12745 | -0.018 | -0.2465 | No |
| 126 | L3MBTL4 | Memczak2013 ANTISENSE, coding, INTERNAL, intronic best transcript NM\_173464 | 12890 | -0.018 | -0.2523 | No |
| 127 | GNB2 | guanine nucleotide binding protein (G protein), beta polypeptide 2 | 12915 | -0.019 | -0.2519 | No |
| 128 | SERPINE1 | serpin peptidase inhibitor, clade E (nexin, plasminogen activator inhibitor type 1), member 1 | 12974 | -0.019 | -0.2532 | No |
| 129 | C1QC | complement component 1, q subcomponent, C chain | 13018 | -0.019 | -0.2537 | No |
| 130 | JAK2 | Janus kinase 2 | 13031 | -0.019 | -0.2526 | No |
| 131 | F7 | coagulation factor VII (serum prothrombin conversion accelerator) | 13057 | -0.020 | -0.2521 | No |
| 132 | APOC1 | apolipoprotein C-I | 13064 | -0.020 | -0.2507 | No |
| 133 | CD36 | CD36 molecule (thrombospondin receptor) | 13245 | -0.021 | -0.2582 | No |
| 134 | RABIF | RAB interacting factor | 13273 | -0.021 | -0.2577 | No |
| 135 | PPP2CB | protein phosphatase 2, catalytic subunit, beta isozyme | 13359 | -0.022 | -0.2601 | No |
| 136 | KIF2A | kinesin heavy chain member 2A | 13362 | -0.022 | -0.2583 | No |
| 137 | TNFAIP3 | tumor necrosis factor, alpha-induced protein 3 | 13363 | -0.022 | -0.2564 | No |
| 138 | PIK3CG | phosphatidylinositol-4,5-bisphosphate 3-kinase, catalytic subunit gamma | 13370 | -0.022 | -0.2548 | No |
| 139 | C1S | complement component 1, s subcomponent | 13614 | -0.023 | -0.2652 | No |
| 140 | PRKCD | protein kinase C, delta | 13658 | -0.024 | -0.2653 | No |
| 141 | RCE1 | Ras converting CAAX endopeptidase 1 | 14025 | -0.026 | -0.2819 | No |
| 142 | DYRK2 | dual specificity tyrosine-(Y)-phosphorylation regulated kinase 2 | 14237 | -0.028 | -0.2903 | No |
| 143 | CEBPB | CCAAT/enhancer binding protein (C/EBP), beta | 14355 | -0.029 | -0.2938 | No |
| 144 | LYN | LYN proto-oncogene, Src family tyrosine kinase | 14438 | -0.030 | -0.2953 | No |
| 145 | TFPI2 | tissue factor pathway inhibitor 2 | 14489 | -0.030 | -0.2952 | No |
| 146 | KLKB1 | kallikrein B1 | 14519 | -0.030 | -0.2940 | No |
| 147 | CTSB | cathepsin B | 14720 | -0.032 | -0.3015 | No |
| 148 | MMP12 | matrix metallopeptidase 12 | 14749 | -0.032 | -0.3000 | No |
| 149 | GZMK | granzyme K | 14831 | -0.033 | -0.3013 | No |
| 150 | DUSP5 | dual specificity phosphatase 5 | 15010 | -0.035 | -0.3074 | No |
| 151 | GMFB | glia maturation factor, beta | 15279 | -0.037 | -0.3180 | No |
| 152 | USP15 | ubiquitin specific peptidase 15 | 15307 | -0.037 | -0.3161 | No |
| 153 | CBLB | Cbl proto-oncogene B, E3 ubiquitin protein ligase | 15444 | -0.038 | -0.3197 | No |
| 154 | PCLO | piccolo presynaptic cytomatrix protein | 15538 | -0.039 | -0.3210 | No |
| 155 | PDGFB | platelet-derived growth factor beta polypeptide | 15682 | -0.041 | -0.3248 | No |
| 156 | S100A12 | S100 calcium binding protein A12 | 15686 | -0.041 | -0.3213 | No |
| 157 | CASP10 | caspase 10 | 15755 | -0.041 | -0.3212 | No |
| 158 | GNAI2 | guanine nucleotide binding protein (G protein), alpha inhibiting activity polypeptide 2 | 16269 | -0.047 | -0.3435 | No |
| 159 | BRPF3 | bromodomain and PHD finger containing 3 | 16289 | -0.047 | -0.3403 | No |
| 160 | SERPINC1 | serpin peptidase inhibitor, clade C (antithrombin), member 1 | 16408 | -0.048 | -0.3421 | No |
| 161 | APOBEC3F | apolipoprotein B mRNA editing enzyme, catalytic polypeptide-like 3F | 16475 | -0.049 | -0.3411 | No |
| 162 | USP16 | ubiquitin specific peptidase 16 | 16484 | -0.050 | -0.3371 | No |
| 163 | RBSN | rabenosyn, RAB effector | 16497 | -0.050 | -0.3333 | No |
| 164 | PDP1 | pyruvate dehyrogenase phosphatase catalytic subunit 1 | 16531 | -0.050 | -0.3305 | No |
| 165 | CASP9 | caspase 9 | 16638 | -0.052 | -0.3313 | No |
| 166 | CTSS | cathepsin S | 16657 | -0.052 | -0.3276 | No |
| 167 | DGKH | diacylglycerol kinase, eta | 16739 | -0.053 | -0.3271 | No |
| 168 | SRC | SRC proto-oncogene, non-receptor tyrosine kinase | 17165 | -0.059 | -0.3438 | No |
| 169 | CD46 | CD46 molecule, complement regulatory protein | 17186 | -0.060 | -0.3395 | No |
| 170 | PRDM4 | PR domain containing 4 | 17234 | -0.060 | -0.3365 | No |
| 171 | CR2 | complement component (3d/Epstein Barr virus) receptor 2 | 17684 | -0.069 | -0.3536 | No |
| 172 | LAMP2 | lysosomal-associated membrane protein 2 | 17747 | -0.071 | -0.3505 | No |
| 173 | ME1 | malic enzyme 1, NADP(+)-dependent, cytosolic | 18066 | -0.079 | -0.3599 | Yes |
| 174 | USP8 | ubiquitin specific peptidase 8 | 18077 | -0.079 | -0.3533 | Yes |
| 175 | PLA2G7 | phospholipase A2, group VII (platelet-activating factor acetylhydrolase, plasma) | 18182 | -0.082 | -0.3514 | Yes |
| 176 | ZFPM2 | zinc finger protein, FOG family member 2 | 18250 | -0.084 | -0.3474 | Yes |
| 177 | CDH13 | cadherin 13 | 18362 | -0.087 | -0.3453 | Yes |
| 178 | HSPA5 | heat shock 70kDa protein 5 (glucose-regulated protein, 78kDa) | 18430 | -0.090 | -0.3408 | Yes |
| 179 | KCNIP3 | Kv channel interacting protein 3, calsenilin | 18489 | -0.092 | -0.3355 | Yes |
| 180 | CTSD | cathepsin D | 18557 | -0.095 | -0.3305 | Yes |
| 181 | GCA | grancalcin, EF-hand calcium binding protein | 18616 | -0.097 | -0.3248 | Yes |
| 182 | IRF1 | interferon regulatory factor 1 | 18746 | -0.106 | -0.3220 | Yes |
| 183 | CASP4 | caspase 4 | 19027 | -0.131 | -0.3249 | Yes |
| 184 | PHEX | phosphate regulating endopeptidase homolog, X-linked | 19063 | -0.136 | -0.3146 | Yes |
| 185 | MMP13 | matrix metallopeptidase 13 | 19091 | -0.140 | -0.3034 | Yes |
| 186 | KLK1 | kallikrein 1 | 19151 | -0.150 | -0.2932 | Yes |
| 187 | ERAP2 | endoplasmic reticulum aminopeptidase 2 | 19270 | -0.180 | -0.2832 | Yes |
| 188 | SERPINA1 | serpin peptidase inhibitor, clade A (alpha-1 antiproteinase, antitrypsin), member 1 | 19288 | -0.189 | -0.2672 | Yes |
| 189 | DPP4 | dipeptidyl-peptidase 4 | 19294 | -0.192 | -0.2503 | Yes |
| 190 | GZMA | granzyme A | 19327 | -0.200 | -0.2341 | Yes |
| 191 | DOCK4 | dedicator of cytokinesis 4 | 19330 | -0.200 | -0.2164 | Yes |
| 192 | CLU | clusterin | 19343 | -0.205 | -0.1987 | Yes |
| 193 | PLA2G4A | phospholipase A2, group IVA (cytosolic, calcium-dependent) | 19411 | -0.248 | -0.1800 | Yes |
| 194 | FN1 | fibronectin 1 | 19439 | -0.280 | -0.1564 | Yes |
| 195 | CASP1 | caspase 1 | 19457 | -0.309 | -0.1297 | Yes |
| 196 | CTSO | cathepsin O | 19481 | -0.342 | -0.1003 | Yes |
| 197 | LIPA | Transcript Identified by AceView, Entrez Gene ID(s) 3988 | 19482 | -0.350 | -0.0691 | Yes |
| 198 | CASP5 | caspase 5 | 19497 | -0.389 | -0.0351 | Yes |
| 199 | ANG | angiogenin, ribonuclease, RNase A family, 5 | 19501 | -0.408 | 0.0012 | Yes |
Table: GSEA details [plain text format]

  

Fig 2: HALLMARK\_COMPLEMENT      
 Blue-Pink O' Gram in the Space of the Analyzed GeneSet

  

Fig 3: HALLMARK\_COMPLEMENT: Random ES distribution      
 Gene set null distribution of ES for **HALLMARK\_COMPLEMENT**

  
