## Supplementary material for "Somatic hypomethylation of pericentromeric SST1 repeats and tetraploidization in human colorectal cancer cells": GSEA results: HALLMARK_DNA_REPAIR.html

Details for gene set HALLMARK\_DNA\_REPAIR[GSEA]

|  || Dataset | eset\_byprobe\_collapsed\_to\_symbols.Diploid\_vs\_Tetraploid.cls #Tetraploid\_versus\_Diploid.Diploid\_vs\_Tetraploid.cls #Tetraploid\_versus\_Diploid\_repos |
| Phenotype | Diploid\_vs\_Tetraploid.cls#Tetraploid\_versus\_Diploid\_repos |
| Upregulated in class | Tetraploid |
| GeneSet | HALLMARK\_DNA\_REPAIR |
| Enrichment Score (ES) | 0.22218734 |
| Normalized Enrichment Score (NES) | 0.95189756 |
| Nominal p-value | 0.5524079 |
| FDR q-value | 0.8124363 |
| FWER p-Value | 1.0 |
Table: GSEA Results Summary

  

Fig 1: Enrichment plot: HALLMARK\_DNA\_REPAIR      
 Profile of the Running ES Score & Positions of GeneSet Members on the Rank Ordered List

  

| SYMBOL | TITLE | RANK IN GENE LIST | RANK METRIC SCORE | RUNNING ES | CORE ENRICHMENT || 1 | HPRT1 | hypoxanthine phosphoribosyltransferase 1 | 311 | 0.132 | 0.0080 | Yes |
| 2 | NPR2 | natriuretic peptide receptor 2 | 409 | 0.118 | 0.0244 | Yes |
| 3 | LIG1 | ligase I, DNA, ATP-dependent | 473 | 0.112 | 0.0416 | Yes |
| 4 | POLH | polymerase (DNA directed), eta | 545 | 0.106 | 0.0573 | Yes |
| 5 | TYMS | thymidylate synthetase | 694 | 0.097 | 0.0673 | Yes |
| 6 | RBX1 | ring-box 1, E3 ubiquitin protein ligase | 1157 | 0.080 | 0.0579 | Yes |
| 7 | SNAPC4 | small nuclear RNA activating complex polypeptide 4 | 1376 | 0.075 | 0.0603 | Yes |
| 8 | PNP | purine nucleoside phosphorylase | 1405 | 0.074 | 0.0722 | Yes |
| 9 | NELFCD | negative elongation factor complex member C/D | 1498 | 0.072 | 0.0805 | Yes |
| 10 | ZNRD1 | zinc ribbon domain containing 1 | 1506 | 0.071 | 0.0931 | Yes |
| 11 | GTF2H5 | general transcription factor IIH subunit 5 | 1730 | 0.066 | 0.0936 | Yes |
| 12 | DDB2 | damage-specific DNA binding protein 2 | 1815 | 0.065 | 0.1010 | Yes |
| 13 | GTF2A2 | general transcription factor IIA 2 | 1875 | 0.064 | 0.1095 | Yes |
| 14 | ADA | adenosine deaminase | 1881 | 0.063 | 0.1208 | Yes |
| 15 | POLA1 | polymerase (DNA directed), alpha 1, catalytic subunit | 1923 | 0.063 | 0.1301 | Yes |
| 16 | RPA3 | replication protein A3 | 1940 | 0.062 | 0.1406 | Yes |
| 17 | RAD51 | RAD51 recombinase | 2121 | 0.060 | 0.1422 | Yes |
| 18 | MPG | N-methylpurine DNA glycosylase | 2129 | 0.060 | 0.1527 | Yes |
| 19 | NELFE | negative elongation factor complex member E | 2135 | 0.059 | 0.1632 | Yes |
| 20 | CDA | cytidine deaminase | 2157 | 0.059 | 0.1729 | Yes |
| 21 | POLD4 | polymerase (DNA-directed), delta 4, accessory subunit | 2787 | 0.050 | 0.1495 | Yes |
| 22 | POLR2K | polymerase (RNA) II (DNA directed) polypeptide K, 7.0kDa | 2797 | 0.050 | 0.1581 | Yes |
| 23 | POLR2H | polymerase (RNA) II (DNA directed) polypeptide H | 2849 | 0.049 | 0.1644 | Yes |
| 24 | UMPS | uridine monophosphate synthetase | 2899 | 0.049 | 0.1708 | Yes |
| 25 | IMPDH2 | IMP (inosine 5-monophosphate) dehydrogenase 2 | 2988 | 0.048 | 0.1749 | Yes |
| 26 | CMPK2 | cytidine monophosphate (UMP-CMP) kinase 2, mitochondrial | 3051 | 0.047 | 0.1802 | Yes |
| 27 | PCNA | proliferating cell nuclear antigen | 3328 | 0.044 | 0.1741 | Yes |
| 28 | POLA2 | polymerase (DNA directed), alpha 2, accessory subunit | 3348 | 0.044 | 0.1811 | Yes |
| 29 | ADCY6 | adenylate cyclase 6 | 3378 | 0.044 | 0.1876 | Yes |
| 30 | POLR1C | polymerase (RNA) I polypeptide C | 3439 | 0.043 | 0.1924 | Yes |
| 31 | NUDT9 | nudix hydrolase 9 | 3622 | 0.041 | 0.1905 | Yes |
| 32 | PRIM1 | primase, DNA, polypeptide 1 (49kDa) | 3626 | 0.041 | 0.1979 | Yes |
| 33 | RFC5 | replication factor C subunit 5 | 3900 | 0.039 | 0.1908 | Yes |
| 34 | MRPL40 | mitochondrial ribosomal protein L40 | 4045 | 0.037 | 0.1902 | Yes |
| 35 | VPS37D | vacuolar protein sorting 37 homolog D (S. cerevisiae) | 4062 | 0.037 | 0.1960 | Yes |
| 36 | AK1 | adenylate kinase 1 | 4140 | 0.036 | 0.1987 | Yes |
| 37 | SSRP1 | structure specific recognition protein 1 | 4293 | 0.035 | 0.1971 | Yes |
| 38 | POLD1 | polymerase (DNA directed), delta 1, catalytic subunit | 4423 | 0.034 | 0.1966 | Yes |
| 39 | SUPT4H1 | SPT4 homolog, DSIF elongation factor subunit | 4445 | 0.034 | 0.2017 | Yes |
| 40 | SF3A3 | splicing factor 3a subunit 3 | 4451 | 0.034 | 0.2075 | Yes |
| 41 | NME1 | NME/NM23 nucleoside diphosphate kinase 1 | 4477 | 0.033 | 0.2123 | Yes |
| 42 | SNAPC5 | small nuclear RNA activating complex polypeptide 5 | 4579 | 0.032 | 0.2130 | Yes |
| 43 | SAC3D1 | SAC3 domain containing 1 | 4676 | 0.032 | 0.2138 | Yes |
| 44 | PDE6G | phosphodiesterase 6G, cGMP-specific, rod, gamma | 4799 | 0.031 | 0.2131 | Yes |
| 45 | POLR2I | polymerase (RNA) II (DNA directed) polypeptide I, 14.5kDa | 4839 | 0.030 | 0.2166 | Yes |
| 46 | ERCC2 | excision repair cross-complementation group 2 | 4840 | 0.030 | 0.2222 | Yes |
| 47 | RFC3 | replication factor C subunit 3 | 5366 | 0.026 | 0.1999 | No |
| 48 | RRM2B | ribonucleotide reductase M2 B (TP53 inducible) | 5380 | 0.026 | 0.2040 | No |
| 49 | POLR2D | polymerase (RNA) II (DNA directed) polypeptide D | 5467 | 0.026 | 0.2043 | No |
| 50 | POLR2E | polymerase (RNA) II (DNA directed) polypeptide E, 25kDa | 5653 | 0.024 | 0.1992 | No |
| 51 | GTF3C5 | general transcription factor IIIC subunit 5 | 5714 | 0.024 | 0.2004 | No |
| 52 | ALYREF | Aly/REF export factor | 5795 | 0.023 | 0.2006 | No |
| 53 | GUK1 | guanylate kinase 1 | 5796 | 0.023 | 0.2048 | No |
| 54 | RFC4 | replication factor C subunit 4 | 5802 | 0.023 | 0.2088 | No |
| 55 | NT5C | 5, 3-nucleotidase, cytosolic | 5848 | 0.023 | 0.2107 | No |
| 56 | BOLA2 | bolA family member 2 | 5866 | 0.023 | 0.2140 | No |
| 57 | GPX4 | glutathione peroxidase 4 | 6109 | 0.021 | 0.2054 | No |
| 58 | NUDT21 | nudix hydrolase 21 | 6126 | 0.021 | 0.2084 | No |
| 59 | RAD52 | RAD52 homolog, DNA repair protein | 6212 | 0.021 | 0.2078 | No |
| 60 | FEN1 | flap structure-specific endonuclease 1 | 6460 | 0.019 | 0.1985 | No |
| 61 | ZWINT | ZW10 interacting kinetochore protein | 6566 | 0.019 | 0.1965 | No |
| 62 | RPA2 | replication protein A2 | 6755 | 0.017 | 0.1899 | No |
| 63 | APRT | adenine phosphoribosyltransferase | 6838 | 0.017 | 0.1887 | No |
| 64 | ERCC8 | excision repair cross-complementation group 8 | 7031 | 0.016 | 0.1817 | No |
| 65 | NME4 | NME/NM23 nucleoside diphosphate kinase 4 | 7098 | 0.015 | 0.1811 | No |
| 66 | RFC2 | replication factor C subunit 2 | 7355 | 0.014 | 0.1704 | No |
| 67 | POLR2F | polymerase (RNA) II (DNA directed) polypeptide F | 7390 | 0.014 | 0.1711 | No |
| 68 | CSTF3 | cleavage stimulation factor, 3 pre-RNA, subunit 3 | 7464 | 0.013 | 0.1697 | No |
| 69 | RAE1 | ribonucleic acid export 1 | 7466 | 0.013 | 0.1720 | No |
| 70 | XPC | xeroderma pigmentosum, complementation group C | 7484 | 0.013 | 0.1735 | No |
| 71 | CETN2 | centrin 2 | 7936 | 0.010 | 0.1521 | No |
| 72 | BCAP31 | B-cell receptor-associated protein 31 | 7967 | 0.010 | 0.1523 | No |
| 73 | TAF1C | TATA box binding protein (TBP)-associated factor, RNA polymerase I, C, 110kDa | 8113 | 0.009 | 0.1465 | No |
| 74 | AAAS | achalasia, adrenocortical insufficiency, alacrimia | 8187 | 0.009 | 0.1444 | No |
| 75 | HCLS1 | hematopoietic cell-specific Lyn substrate 1 | 8198 | 0.009 | 0.1454 | No |
| 76 | NCBP2 | nuclear cap binding protein subunit 2 | 8274 | 0.008 | 0.1431 | No |
| 77 | GTF2H3 | general transcription factor IIH subunit 3 | 8418 | 0.007 | 0.1370 | No |
| 78 | ITPA | inosine triphosphatase (nucleoside triphosphate pyrophosphatase) | 8461 | 0.007 | 0.1361 | No |
| 79 | TARBP2 | TAR (HIV-1) RNA binding protein 2 | 8465 | 0.007 | 0.1373 | No |
| 80 | NME3 | NME/NM23 nucleoside diphosphate kinase 3 | 9040 | 0.004 | 0.1083 | No |
| 81 | REV3L | REV3 like, DNA directed polymerase zeta catalytic subunit | 9150 | 0.003 | 0.1033 | No |
| 82 | NELFB | negative elongation factor complex member B | 9274 | 0.003 | 0.0974 | No |
| 83 | DAD1 | defender against cell death 1 | 9287 | 0.003 | 0.0973 | No |
| 84 | EIF1B | eukaryotic translation initiation factor 1B | 9343 | 0.002 | 0.0949 | No |
| 85 | TAF12 | TAF12 RNA polymerase II, TATA box binding protein (TBP)-associated factor, 20kDa | 9782 | -0.000 | 0.0723 | No |
| 86 | ZNF707 | zinc finger protein 707 | 9891 | -0.001 | 0.0668 | No |
| 87 | ADRM1 | adhesion regulating molecule 1 | 10136 | -0.002 | 0.0545 | No |
| 88 | DDB1 | damage-specific DNA binding protein 1 | 10240 | -0.003 | 0.0497 | No |
| 89 | POLR2G | polymerase (RNA) II (DNA directed) polypeptide G | 10804 | -0.006 | 0.0217 | No |
| 90 | RALA | v-ral simian leukemia viral oncogene homolog A (ras related) | 11268 | -0.009 | -0.0006 | No |
| 91 | STX3 | syntaxin 3 | 11410 | -0.009 | -0.0062 | No |
| 92 | GTF2H1 | general transcription factor IIH subunit 1 | 11860 | -0.012 | -0.0272 | No |
| 93 | POLL | polymerase (DNA directed), lambda | 11921 | -0.012 | -0.0280 | No |
| 94 | ERCC1 | excision repair cross-complementation group 1 | 12020 | -0.013 | -0.0307 | No |
| 95 | CCNO | cyclin O | 12022 | -0.013 | -0.0284 | No |
| 96 | ERCC3 | excision repair cross-complementation group 3 | 12092 | -0.013 | -0.0296 | No |
| 97 | SMAD5 | SMAD family member 5 | 12225 | -0.014 | -0.0338 | No |
| 98 | GMPR2 | guanosine monophosphate reductase 2 | 12230 | -0.014 | -0.0314 | No |
| 99 | TAF9 | TAF9 RNA polymerase II, TATA box binding protein (TBP)-associated factor, 32kDa | 12231 | -0.014 | -0.0287 | No |
| 100 | UPF3B | UPF3 regulator of nonsense transcripts homolog B (yeast) | 12416 | -0.016 | -0.0354 | No |
| 101 | SRSF6 | serine/arginine-rich splicing factor 6 | 12418 | -0.016 | -0.0326 | No |
| 102 | SURF1 | surfeit 1 | 12513 | -0.016 | -0.0345 | No |
| 103 | ERCC5 | excision repair cross-complementation group 5 | 12684 | -0.017 | -0.0402 | No |
| 104 | BRF2 | BRF2, RNA polymerase III transcription initiation factor 50 kDa subunit | 12722 | -0.017 | -0.0389 | No |
| 105 | TAF10 | TAF10 RNA polymerase II, TATA box binding protein (TBP)-associated factor, 30kDa | 12734 | -0.018 | -0.0363 | No |
| 106 | BCAM | basal cell adhesion molecule (Lutheran blood group) | 12938 | -0.019 | -0.0433 | No |
| 107 | ERCC4 | excision repair cross-complementation group 4 | 13098 | -0.020 | -0.0479 | No |
| 108 | ELL | elongation factor RNA polymerase II | 13445 | -0.022 | -0.0618 | No |
| 109 | TSG101 | tumor susceptibility 101 | 13515 | -0.023 | -0.0612 | No |
| 110 | TMED2 | transmembrane p24 trafficking protein 2 | 13702 | -0.024 | -0.0664 | No |
| 111 | POLR3GL | polymerase (RNA) III (DNA directed) polypeptide G (32kD)-like | 13775 | -0.025 | -0.0656 | No |
| 112 | MPC2 | mitochondrial pyruvate carrier 2 | 13841 | -0.025 | -0.0644 | No |
| 113 | VPS37B | vacuolar protein sorting 37 homolog B (S. cerevisiae) | 14014 | -0.026 | -0.0685 | No |
| 114 | TP53 | tumor protein p53 | 14048 | -0.027 | -0.0653 | No |
| 115 | ARL6IP1 | ADP-ribosylation factor like GTPase 6 interacting protein 1 | 14255 | -0.028 | -0.0708 | No |
| 116 | SUPT5H | SPT5 homolog, DSIF elongation factor subunit | 14268 | -0.028 | -0.0663 | No |
| 117 | CANT1 | calcium activated nucleotidase 1 | 14424 | -0.030 | -0.0688 | No |
| 118 | NT5C3A | 5-nucleotidase, cytosolic IIIA | 14520 | -0.030 | -0.0682 | No |
| 119 | TAF13 | TAF13 RNA polymerase II, TATA box binding protein (TBP)-associated factor, 18kDa | 14649 | -0.032 | -0.0691 | No |
| 120 | DGUOK | deoxyguanosine kinase | 14722 | -0.032 | -0.0669 | No |
| 121 | EDF1 | endothelial differentiation-related factor 1 | 14755 | -0.032 | -0.0627 | No |
| 122 | GTF2F1 | general transcription factor IIF subunit 1 | 14919 | -0.034 | -0.0649 | No |
| 123 | POLR1D | polymerase (RNA) I polypeptide D | 14933 | -0.034 | -0.0594 | No |
| 124 | TK2 | thymidine kinase 2, mitochondrial | 14943 | -0.034 | -0.0537 | No |
| 125 | NFX1 | nuclear transcription factor, X-box binding 1 | 14946 | -0.034 | -0.0476 | No |
| 126 | AK3 | adenylate kinase 3 | 15257 | -0.037 | -0.0570 | No |
| 127 | POLR2C | polymerase (RNA) II (DNA directed) polypeptide C, 33kDa | 15294 | -0.037 | -0.0521 | No |
| 128 | COX17 | COX17 cytochrome c oxidase copper chaperone | 15316 | -0.037 | -0.0464 | No |
| 129 | POLR2J | polymerase (RNA) II (DNA directed) polypeptide J, 13.3kDa | 15410 | -0.038 | -0.0443 | No |
| 130 | POLR2A | polymerase (RNA) II (DNA directed) polypeptide A, 220kDa | 15461 | -0.038 | -0.0399 | No |
| 131 | POM121 | POM121 transmembrane nucleoporin | 15476 | -0.039 | -0.0336 | No |
| 132 | POLB | polymerase (DNA directed), beta | 15530 | -0.039 | -0.0292 | No |
| 133 | DCTN4 | dynactin 4 (p62) | 16114 | -0.045 | -0.0511 | No |
| 134 | DGCR8 | DGCR8 microprocessor complex subunit | 16116 | -0.045 | -0.0430 | No |
| 135 | SDCBP | syndecan binding protein | 16135 | -0.045 | -0.0357 | No |
| 136 | AGO4 | argonaute RISC catalytic component 4 | 16171 | -0.046 | -0.0292 | No |
| 137 | VPS28 | vacuolar protein sorting 28 homolog (S. cerevisiae) | 16534 | -0.050 | -0.0387 | No |
| 138 | POLR3C | polymerase (RNA) III (DNA directed) polypeptide C (62kD) | 16610 | -0.051 | -0.0332 | No |
| 139 | CLP1 | Transcript Identified by AceView, Entrez Gene ID(s) 10978 | 17106 | -0.059 | -0.0481 | No |
| 140 | TAF6 | TAF6 RNA polymerase II, TATA box binding protein (TBP)-associated factor, 80kDa | 17172 | -0.060 | -0.0406 | No |
| 141 | POLD3 | polymerase (DNA-directed), delta 3, accessory subunit | 17176 | -0.060 | -0.0299 | No |
| 142 | GTF2B | general transcription factor IIB | 17483 | -0.065 | -0.0339 | No |
| 143 | DUT | deoxyuridine triphosphatase | 18343 | -0.087 | -0.0625 | No |
| 144 | RNMT | RNA (guanine-7-) methyltransferase | 18744 | -0.106 | -0.0638 | No |
| 145 | SEC61A1 | Sec61 translocon alpha 1 subunit | 18812 | -0.111 | -0.0472 | No |
| 146 | USP11 | ubiquitin specific peptidase 11 | 18992 | -0.126 | -0.0335 | No |
| 147 | PDE4B | phosphodiesterase 4B, cAMP-specific | 19061 | -0.135 | -0.0124 | No |
| 148 | POLE4 | polymerase (DNA-directed), epsilon 4, accessory subunit | 19323 | -0.199 | 0.0104 | No |
Table: GSEA details [plain text format]

  

Fig 2: HALLMARK\_DNA\_REPAIR      
 Blue-Pink O' Gram in the Space of the Analyzed GeneSet

  

Fig 3: HALLMARK\_DNA\_REPAIR: Random ES distribution      
 Gene set null distribution of ES for **HALLMARK\_DNA\_REPAIR**

  
