## Supplementary material for "Somatic hypomethylation of pericentromeric SST1 repeats and tetraploidization in human colorectal cancer cells": GSEA results: HALLMARK_E2F_TARGETS.html

Details for gene set HALLMARK\_E2F\_TARGETS[GSEA]

|  || Dataset | eset\_byprobe\_collapsed\_to\_symbols.Diploid\_vs\_Tetraploid.cls #Tetraploid\_versus\_Diploid.Diploid\_vs\_Tetraploid.cls #Tetraploid\_versus\_Diploid\_repos |
| Phenotype | Diploid\_vs\_Tetraploid.cls#Tetraploid\_versus\_Diploid\_repos |
| Upregulated in class | Tetraploid |
| GeneSet | HALLMARK\_E2F\_TARGETS |
| Enrichment Score (ES) | 0.49202254 |
| Normalized Enrichment Score (NES) | 2.2049901 |
| Nominal p-value | 0.0 |
| FDR q-value | 0.0 |
| FWER p-Value | 0.0 |
Table: GSEA Results Summary

  

Fig 1: Enrichment plot: HALLMARK\_E2F\_TARGETS      
 Profile of the Running ES Score & Positions of GeneSet Members on the Rank Ordered List

  

| SYMBOL | TITLE | RANK IN GENE LIST | RANK METRIC SCORE | RUNNING ES | CORE ENRICHMENT || 1 | EED | embryonic ectoderm development | 152 | 0.170 | 0.0120 | Yes |
| 2 | CHEK2 | checkpoint kinase 2 | 208 | 0.153 | 0.0271 | Yes |
| 3 | BRCA2 | breast cancer 2, early onset | 273 | 0.137 | 0.0397 | Yes |
| 4 | CDC25A | cell division cycle 25A | 295 | 0.134 | 0.0543 | Yes |
| 5 | MLH1 | mutL homolog 1 | 332 | 0.128 | 0.0674 | Yes |
| 6 | MYBL2 | v-myb avian myeloblastosis viral oncogene homolog-like 2 | 400 | 0.119 | 0.0778 | Yes |
| 7 | PDS5B | PDS5 cohesin associated factor B | 452 | 0.114 | 0.0885 | Yes |
| 8 | TIMELESS | timeless circadian clock | 468 | 0.113 | 0.1009 | Yes |
| 9 | LIG1 | ligase I, DNA, ATP-dependent | 473 | 0.112 | 0.1138 | Yes |
| 10 | ORC6 | origin recognition complex subunit 6 | 480 | 0.112 | 0.1266 | Yes |
| 11 | RNASEH2A | ribonuclease H2, subunit A | 611 | 0.103 | 0.1319 | Yes |
| 12 | ATAD2 | ATPase family, AAA domain containing 2 | 641 | 0.101 | 0.1421 | Yes |
| 13 | CDKN1A | cyclin-dependent kinase inhibitor 1A (p21, Cip1) | 690 | 0.097 | 0.1510 | Yes |
| 14 | PRIM2 | primase, DNA, polypeptide 2 (58kDa) | 728 | 0.096 | 0.1602 | Yes |
| 15 | CCNE1 | cyclin E1 | 738 | 0.095 | 0.1709 | Yes |
| 16 | TRIP13 | thyroid hormone receptor interactor 13 | 774 | 0.093 | 0.1800 | Yes |
| 17 | SLBP | stem-loop binding protein | 790 | 0.093 | 0.1900 | Yes |
| 18 | NUP107 | nucleoporin 107kDa | 859 | 0.090 | 0.1970 | Yes |
| 19 | CTPS1 | CTP synthase 1 | 860 | 0.090 | 0.2074 | Yes |
| 20 | DSCC1 | DNA replication and sister chromatid cohesion 1 | 875 | 0.089 | 0.2171 | Yes |
| 21 | MELK | maternal embryonic leucine zipper kinase | 882 | 0.089 | 0.2272 | Yes |
| 22 | STMN1 | stathmin 1 | 951 | 0.086 | 0.2337 | Yes |
| 23 | CKS2 | CDC28 protein kinase regulatory subunit 2 | 1016 | 0.084 | 0.2402 | Yes |
| 24 | CDC20 | cell division cycle 20 | 1022 | 0.084 | 0.2497 | Yes |
| 25 | TK1 | thymidine kinase 1, soluble | 1063 | 0.082 | 0.2572 | Yes |
| 26 | DCTPP1 | dCTP pyrophosphatase 1 | 1076 | 0.081 | 0.2661 | Yes |
| 27 | BRCA1 | breast cancer 1, early onset | 1119 | 0.080 | 0.2733 | Yes |
| 28 | DIAPH3 | diaphanous-related formin 3 | 1136 | 0.080 | 0.2818 | Yes |
| 29 | CDKN2A | cyclin-dependent kinase inhibitor 2A | 1257 | 0.077 | 0.2846 | Yes |
| 30 | KIF22 | kinesin family member 22 | 1286 | 0.077 | 0.2921 | Yes |
| 31 | MCM4 | minichromosome maintenance complex component 4 | 1325 | 0.076 | 0.2990 | Yes |
| 32 | GINS1 | GINS complex subunit 1 (Psf1 homolog) | 1375 | 0.075 | 0.3052 | Yes |
| 33 | XPO1 | exportin 1 | 1444 | 0.073 | 0.3102 | Yes |
| 34 | MCM3 | minichromosome maintenance complex component 3 | 1450 | 0.073 | 0.3184 | Yes |
| 35 | LYAR | Ly1 antibody reactive | 1468 | 0.072 | 0.3260 | Yes |
| 36 | CENPM | centromere protein M | 1560 | 0.070 | 0.3295 | Yes |
| 37 | LMNB1 | lamin B1 | 1705 | 0.066 | 0.3298 | Yes |
| 38 | BUB1B | BUB1 mitotic checkpoint serine/threonine kinase B | 1762 | 0.066 | 0.3345 | Yes |
| 39 | MCM5 | minichromosome maintenance complex component 5 | 1829 | 0.064 | 0.3386 | Yes |
| 40 | CDCA3 | cell division cycle associated 3 | 1844 | 0.064 | 0.3454 | Yes |
| 41 | PPM1D | protein phosphatase, Mg2+/Mn2+ dependent, 1D | 1919 | 0.063 | 0.3489 | Yes |
| 42 | SPC25 | SPC25, NDC80 kinetochore complex component | 1922 | 0.063 | 0.3561 | Yes |
| 43 | RPA3 | replication protein A3 | 1940 | 0.062 | 0.3625 | Yes |
| 44 | DNMT1 | DNA (cytosine-5-)-methyltransferase 1 | 2038 | 0.061 | 0.3646 | Yes |
| 45 | RANBP1 | RAN binding protein 1 | 2070 | 0.060 | 0.3700 | Yes |
| 46 | MAD2L1 | MAD2 mitotic arrest deficient-like 1 (yeast) | 2149 | 0.059 | 0.3729 | Yes |
| 47 | UNG | uracil DNA glycosylase | 2227 | 0.058 | 0.3757 | Yes |
| 48 | NAP1L1 | nucleosome assembly protein 1-like 1 | 2338 | 0.056 | 0.3766 | Yes |
| 49 | DCLRE1B | DNA cross-link repair 1B | 2368 | 0.056 | 0.3816 | Yes |
| 50 | HELLS | helicase, lymphoid-specific | 2432 | 0.055 | 0.3847 | Yes |
| 51 | HMMR | hyaluronan-mediated motility receptor (RHAMM) | 2515 | 0.053 | 0.3867 | Yes |
| 52 | BARD1 | BRCA1 associated RING domain 1 | 2585 | 0.053 | 0.3893 | Yes |
| 53 | RAD51C | RAD51 paralog C | 2687 | 0.051 | 0.3900 | Yes |
| 54 | ASF1B | anti-silencing function 1B histone chaperone | 2733 | 0.051 | 0.3936 | Yes |
| 55 | TACC3 | transforming, acidic coiled-coil containing protein 3 | 2794 | 0.050 | 0.3964 | Yes |
| 56 | RAD1 | RAD1 checkpoint DNA exonuclease | 2803 | 0.050 | 0.4018 | Yes |
| 57 | LBR | lamin B receptor | 2818 | 0.050 | 0.4068 | Yes |
| 58 | E2F8 | E2F transcription factor 8 | 2851 | 0.049 | 0.4109 | Yes |
| 59 | CSE1L | CSE1 chromosome segregation 1-like (yeast) | 2926 | 0.048 | 0.4128 | Yes |
| 60 | CENPE | centromere protein E | 2957 | 0.048 | 0.4168 | Yes |
| 61 | SRSF2 | serine/arginine-rich splicing factor 2 | 2978 | 0.048 | 0.4213 | Yes |
| 62 | UBR7 | ubiquitin protein ligase E3 component n-recognin 7 (putative) | 3001 | 0.047 | 0.4258 | Yes |
| 63 | HNRNPD | heterogeneous nuclear ribonucleoprotein D | 3023 | 0.047 | 0.4302 | Yes |
| 64 | GINS4 | GINS complex subunit 4 (Sld5 homolog) | 3060 | 0.047 | 0.4338 | Yes |
| 65 | RRM2 | ribonucleotide reductase M2 | 3128 | 0.046 | 0.4357 | Yes |
| 66 | SMC4 | structural maintenance of chromosomes 4 | 3153 | 0.046 | 0.4399 | Yes |
| 67 | MYC | v-myc avian myelocytomatosis viral oncogene homolog | 3194 | 0.046 | 0.4431 | Yes |
| 68 | PCNA | proliferating cell nuclear antigen | 3328 | 0.044 | 0.4414 | Yes |
| 69 | POLA2 | polymerase (DNA directed), alpha 2, accessory subunit | 3348 | 0.044 | 0.4456 | Yes |
| 70 | NASP | nuclear autoantigenic sperm protein (histone-binding) | 3425 | 0.043 | 0.4468 | Yes |
| 71 | MKI67 | marker of proliferation Ki-67 | 3496 | 0.043 | 0.4481 | Yes |
| 72 | MCM6 | minichromosome maintenance complex component 6 | 3558 | 0.042 | 0.4499 | Yes |
| 73 | PSIP1 | PC4 and SFRS1 interacting protein 1 | 3663 | 0.041 | 0.4493 | Yes |
| 74 | AURKB | aurora kinase B | 3671 | 0.041 | 0.4537 | Yes |
| 75 | MSH2 | mutS homolog 2 | 3691 | 0.041 | 0.4575 | Yes |
| 76 | KIF18B | kinesin family member 18B | 3739 | 0.040 | 0.4597 | Yes |
| 77 | NCAPD2 | non-SMC condensin I complex subunit D2 | 3883 | 0.039 | 0.4568 | Yes |
| 78 | POLE | polymerase (DNA directed), epsilon, catalytic subunit | 3885 | 0.039 | 0.4613 | Yes |
| 79 | NOP56 | NOP56 ribonucleoprotein | 3937 | 0.038 | 0.4631 | Yes |
| 80 | UBE2T | ubiquitin conjugating enzyme E2T | 3942 | 0.038 | 0.4674 | Yes |
| 81 | MMS22L | MMS22-like, DNA repair protein | 4040 | 0.037 | 0.4667 | Yes |
| 82 | CHEK1 | checkpoint kinase 1 | 4083 | 0.037 | 0.4688 | Yes |
| 83 | PAICS | phosphoribosylaminoimidazole carboxylase, phosphoribosylaminoimidazole succinocarboxamide synthetase | 4227 | 0.035 | 0.4655 | Yes |
| 84 | WEE1 | WEE1 G2 checkpoint kinase | 4267 | 0.035 | 0.4676 | Yes |
| 85 | SSRP1 | structure specific recognition protein 1 | 4293 | 0.035 | 0.4704 | Yes |
| 86 | CDKN2C | cyclin-dependent kinase inhibitor 2C (p18, inhibits CDK4) | 4353 | 0.034 | 0.4713 | Yes |
| 87 | TRA2B | transformer 2 beta homolog (Drosophila) | 4395 | 0.034 | 0.4732 | Yes |
| 88 | POLD1 | polymerase (DNA directed), delta 1, catalytic subunit | 4423 | 0.034 | 0.4757 | Yes |
| 89 | NME1 | NME/NM23 nucleoside diphosphate kinase 1 | 4477 | 0.033 | 0.4769 | Yes |
| 90 | PRPS1 | phosphoribosyl pyrophosphate synthetase 1 | 4489 | 0.033 | 0.4802 | Yes |
| 91 | DCK | deoxycytidine kinase | 4497 | 0.033 | 0.4837 | Yes |
| 92 | RAD51AP1 | RAD51 associated protein 1 | 4516 | 0.033 | 0.4866 | Yes |
| 93 | SMC6 | Transcript Identified by AceView, Entrez Gene ID(s) 79677 | 4721 | 0.031 | 0.4797 | Yes |
| 94 | CDK1 | cyclin-dependent kinase 1 | 4823 | 0.031 | 0.4781 | Yes |
| 95 | RPA1 | replication protein A1 | 4848 | 0.030 | 0.4804 | Yes |
| 96 | SMC1A | structural maintenance of chromosomes 1A | 4903 | 0.030 | 0.4811 | Yes |
| 97 | RBBP7 | retinoblastoma binding protein 7 | 4910 | 0.030 | 0.4843 | Yes |
| 98 | HMGB3 | high mobility group box 3 | 4965 | 0.030 | 0.4850 | Yes |
| 99 | AURKA | aurora kinase A | 5067 | 0.029 | 0.4831 | Yes |
| 100 | ILF3 | interleukin enhancer binding factor 3 | 5069 | 0.029 | 0.4864 | Yes |
| 101 | HMGA1 | high mobility group AT-hook 1 | 5073 | 0.029 | 0.4896 | Yes |
| 102 | GSPT1 | G1 to S phase transition 1 | 5217 | 0.028 | 0.4855 | Yes |
| 103 | PSMC3IP | PSMC3 interacting protein | 5303 | 0.027 | 0.4842 | Yes |
| 104 | DDX39A | DEAD (Asp-Glu-Ala-Asp) box polypeptide 39A | 5309 | 0.027 | 0.4871 | Yes |
| 105 | MCM2 | minichromosome maintenance complex component 2 | 5326 | 0.027 | 0.4894 | Yes |
| 106 | RFC3 | replication factor C subunit 3 | 5366 | 0.026 | 0.4905 | Yes |
| 107 | NUP153 | nucleoporin 153kDa | 5453 | 0.026 | 0.4891 | Yes |
| 108 | TCF19 | transcription factor 19 | 5455 | 0.026 | 0.4920 | Yes |
| 109 | TUBG1 | tubulin, gamma 1 | 5617 | 0.025 | 0.4866 | No |
| 110 | PLK4 | polo-like kinase 4 | 5691 | 0.024 | 0.4856 | No |
| 111 | PRDX4 | peroxiredoxin 4 | 5730 | 0.024 | 0.4864 | No |
| 112 | ZW10 | zw10 kinetochore protein | 5810 | 0.023 | 0.4851 | No |
| 113 | SNRPB | small nuclear ribonucleoprotein polypeptides B and B1 | 5820 | 0.023 | 0.4873 | No |
| 114 | LUC7L3 | LUC7-like 3 pre-mRNA splicing factor | 5935 | 0.022 | 0.4840 | No |
| 115 | NOLC1 | nucleolar and coiled-body phosphoprotein 1 | 5963 | 0.022 | 0.4852 | No |
| 116 | EZH2 | enhancer of zeste 2 polycomb repressive complex 2 subunit | 5989 | 0.022 | 0.4865 | No |
| 117 | SRSF1 | serine/arginine-rich splicing factor 1 | 6000 | 0.022 | 0.4886 | No |
| 118 | NUDT21 | nudix hydrolase 21 | 6126 | 0.021 | 0.4846 | No |
| 119 | ANP32E | acidic nuclear phosphoprotein 32 family member E | 6152 | 0.021 | 0.4858 | No |
| 120 | KPNA2 | karyopherin alpha 2 (RAG cohort 1, importin alpha 1) | 6200 | 0.021 | 0.4857 | No |
| 121 | PA2G4 | proliferation-associated 2G4 | 6269 | 0.020 | 0.4846 | No |
| 122 | NAA38 | N(alpha)-acetyltransferase 38, NatC auxiliary subunit | 6467 | 0.019 | 0.4767 | No |
| 123 | PHF5A | PHD finger protein 5A | 6754 | 0.017 | 0.4639 | No |
| 124 | RPA2 | replication protein A2 | 6755 | 0.017 | 0.4659 | No |
| 125 | RAD21 | RAD21 cohesin complex component | 6756 | 0.017 | 0.4679 | No |
| 126 | SHMT1 | serine hydroxymethyltransferase 1 (soluble) | 6801 | 0.017 | 0.4676 | No |
| 127 | TOP2A | topoisomerase (DNA) II alpha | 6917 | 0.016 | 0.4636 | No |
| 128 | PNN | pinin, desmosome associated protein | 6937 | 0.016 | 0.4645 | No |
| 129 | RACGAP1 | Rac GTPase activating protein 1 | 7079 | 0.015 | 0.4590 | No |
| 130 | MXD3 | MAX dimerization protein 3 | 7083 | 0.015 | 0.4607 | No |
| 131 | TUBB | tubulin, beta class I | 7107 | 0.015 | 0.4612 | No |
| 132 | RAN | RAN, member RAS oncogene family | 7181 | 0.015 | 0.4592 | No |
| 133 | CDK4 | cyclin-dependent kinase 4 | 7198 | 0.015 | 0.4601 | No |
| 134 | SYNCRIP | synaptotagmin binding, cytoplasmic RNA interacting protein | 7248 | 0.015 | 0.4593 | No |
| 135 | RFC2 | replication factor C subunit 2 | 7355 | 0.014 | 0.4554 | No |
| 136 | PRKDC | protein kinase, DNA-activated, catalytic polypeptide | 7386 | 0.014 | 0.4554 | No |
| 137 | POLD2 | polymerase (DNA directed), delta 2, accessory subunit | 7486 | 0.013 | 0.4518 | No |
| 138 | CTCF | CCCTC-binding factor (zinc finger protein) | 7528 | 0.013 | 0.4512 | No |
| 139 | AK2 | adenylate kinase 2 | 7542 | 0.013 | 0.4520 | No |
| 140 | DEK | DEK proto-oncogene | 7764 | 0.011 | 0.4419 | No |
| 141 | HMGB2 | high mobility group box 2 | 7921 | 0.010 | 0.4350 | No |
| 142 | BIRC5 | baculoviral IAP repeat containing 5 | 7953 | 0.010 | 0.4346 | No |
| 143 | MCM7 | minichromosome maintenance complex component 7 | 7985 | 0.010 | 0.4341 | No |
| 144 | SUV39H1 | suppressor of variegation 3-9 homolog 1 (Drosophila) | 7990 | 0.010 | 0.4351 | No |
| 145 | ESPL1 | extra spindle pole bodies like 1, separase | 7999 | 0.010 | 0.4358 | No |
| 146 | DLGAP5 | discs, large (Drosophila) homolog-associated protein 5 | 8007 | 0.010 | 0.4366 | No |
| 147 | CCNB2 | cyclin B2 | 8020 | 0.010 | 0.4372 | No |
| 148 | TMPO | thymopoietin | 8045 | 0.010 | 0.4370 | No |
| 149 | STAG1 | stromal antigen 1 | 8239 | 0.008 | 0.4280 | No |
| 150 | XRCC6 | X-ray repair complementing defective repair in Chinese hamster cells 6 | 8395 | 0.007 | 0.4209 | No |
| 151 | PLK1 | polo-like kinase 1 | 8734 | 0.006 | 0.4041 | No |
| 152 | DEPDC1 | DEP domain containing 1 | 8741 | 0.005 | 0.4044 | No |
| 153 | SMC3 | structural maintenance of chromosomes 3 | 8842 | 0.005 | 0.3998 | No |
| 154 | TFRC | transferrin receptor | 8948 | 0.004 | 0.3949 | No |
| 155 | CBX5 | chromobox homolog 5 | 9037 | 0.004 | 0.3908 | No |
| 156 | WDR90 | WD repeat domain 90 | 9177 | 0.003 | 0.3839 | No |
| 157 | DONSON | downstream neighbor of SON | 9214 | 0.003 | 0.3824 | No |
| 158 | PAN2 | PAN2 poly(A) specific ribonuclease subunit | 9223 | 0.003 | 0.3823 | No |
| 159 | IPO7 | importin 7 | 9261 | 0.003 | 0.3807 | No |
| 160 | CDKN3 | cyclin-dependent kinase inhibitor 3 | 9369 | 0.002 | 0.3754 | No |
| 161 | GINS3 | GINS complex subunit 3 (Psf3 homolog) | 9436 | 0.002 | 0.3722 | No |
| 162 | PTTG1 | pituitary tumor-transforming 1 | 9499 | 0.001 | 0.3692 | No |
| 163 | PPP1R8 | protein phosphatase 1, regulatory subunit 8 | 9526 | 0.001 | 0.3680 | No |
| 164 | BRMS1L | breast cancer metastasis-suppressor 1-like | 9656 | 0.001 | 0.3614 | No |
| 165 | CKS1B | CDC28 protein kinase regulatory subunit 1B | 10258 | -0.003 | 0.3306 | No |
| 166 | EIF2S1 | eukaryotic translation initiation factor 2, subunit 1 alpha, 35kDa | 10272 | -0.003 | 0.3303 | No |
| 167 | TIPIN | TIMELESS interacting protein | 10301 | -0.003 | 0.3292 | No |
| 168 | RAD50 | RAD50 homolog, double strand break repair protein | 10328 | -0.003 | 0.3282 | No |
| 169 | RFC1 | replication factor C subunit 1 | 10561 | -0.004 | 0.3167 | No |
| 170 | SPC24 | SPC24, NDC80 kinetochore complex component | 11187 | -0.008 | 0.2854 | No |
| 171 | KIF2C | kinesin family member 2C | 11587 | -0.010 | 0.2659 | No |
| 172 | NBN | nibrin | 11962 | -0.013 | 0.2480 | No |
| 173 | USP1 | ubiquitin specific peptidase 1 | 12752 | -0.018 | 0.2093 | No |
| 174 | TBRG4 | transforming growth factor beta regulator 4 | 12791 | -0.018 | 0.2094 | No |
| 175 | POP7 | POP7 homolog, ribonuclease P/MRP subunit | 12871 | -0.018 | 0.2075 | No |
| 176 | CIT | citron rho-interacting serine/threonine kinase | 13026 | -0.019 | 0.2018 | No |
| 177 | CDC25B | cell division cycle 25B | 13972 | -0.026 | 0.1559 | No |
| 178 | TP53 | tumor protein p53 | 14048 | -0.027 | 0.1551 | No |
| 179 | HUS1 | HUS1 checkpoint clamp component | 14352 | -0.029 | 0.1429 | No |
| 180 | KIF4A | kinesin family member 4A | 14527 | -0.031 | 0.1375 | No |
| 181 | UBE2S | ubiquitin-conjugating enzyme E2S | 14865 | -0.033 | 0.1239 | No |
| 182 | CDCA8 | cell division cycle associated 8 | 14908 | -0.034 | 0.1257 | No |
| 183 | ASF1A | anti-silencing function 1A histone chaperone | 15331 | -0.037 | 0.1082 | No |
| 184 | PMS2 | PMS1 homolog 2, mismatch repair system component | 16312 | -0.047 | 0.0630 | No |
| 185 | CDKN1B | cyclin-dependent kinase inhibitor 1B (p27, Kip1) | 16336 | -0.047 | 0.0674 | No |
| 186 | EXOSC8 | exosome component 8 | 16621 | -0.052 | 0.0587 | No |
| 187 | POLD3 | polymerase (DNA-directed), delta 3, accessory subunit | 17176 | -0.060 | 0.0370 | No |
| 188 | ORC2 | origin recognition complex subunit 2 | 17197 | -0.060 | 0.0430 | No |
| 189 | SPAG5 | sperm associated antigen 5 | 17519 | -0.066 | 0.0341 | No |
| 190 | ING3 | inhibitor of growth family member 3 | 17520 | -0.066 | 0.0417 | No |
| 191 | CCP110 | centriolar coiled coil protein 110kDa | 17778 | -0.071 | 0.0368 | No |
| 192 | MTHFD2 | methylenetetrahydrofolate dehydrogenase (NADP+ dependent) 2, methenyltetrahydrofolate cyclohydrolase | 18255 | -0.084 | 0.0220 | No |
| 193 | NUP205 | nucleoporin 205kDa | 18334 | -0.086 | 0.0281 | No |
| 194 | DUT | deoxyuridine triphosphatase | 18343 | -0.087 | 0.0378 | No |
| 195 | POLE4 | polymerase (DNA-directed), epsilon 4, accessory subunit | 19323 | -0.199 | 0.0104 | No |
Table: GSEA details [plain text format]

  

Fig 2: HALLMARK\_E2F\_TARGETS      
 Blue-Pink O' Gram in the Space of the Analyzed GeneSet

  

Fig 3: HALLMARK\_E2F\_TARGETS: Random ES distribution      
 Gene set null distribution of ES for **HALLMARK\_E2F\_TARGETS**

  
