## Supplementary material for "Somatic hypomethylation of pericentromeric SST1 repeats and tetraploidization in human colorectal cancer cells": GSEA results: HALLMARK_EPITHELIAL_MESENCHYMAL_TRANSITION.html

Details for gene set HALLMARK\_EPITHELIAL\_MESENCHYMAL\_TRANSITION[GSEA]

|  || Dataset | eset\_byprobe\_collapsed\_to\_symbols.Diploid\_vs\_Tetraploid.cls #Tetraploid\_versus\_Diploid.Diploid\_vs\_Tetraploid.cls #Tetraploid\_versus\_Diploid\_repos |
| Phenotype | Diploid\_vs\_Tetraploid.cls#Tetraploid\_versus\_Diploid\_repos |
| Upregulated in class | Diploid |
| GeneSet | HALLMARK\_EPITHELIAL\_MESENCHYMAL\_TRANSITION |
| Enrichment Score (ES) | -0.31272915 |
| Normalized Enrichment Score (NES) | -1.2894069 |
| Nominal p-value | 0.04427481 |
| FDR q-value | 0.28269485 |
| FWER p-Value | 0.912 |
Table: GSEA Results Summary

  

Fig 1: Enrichment plot: HALLMARK\_EPITHELIAL\_MESENCHYMAL\_TRANSITION      
 Profile of the Running ES Score & Positions of GeneSet Members on the Rank Ordered List

  

| SYMBOL | TITLE | RANK IN GENE LIST | RANK METRIC SCORE | RUNNING ES | CORE ENRICHMENT || 1 | NT5E | 5-nucleotidase, ecto (CD73) | 37 | 0.307 | 0.0243 | No |
| 2 | CDH11 | cadherin 11, type 2, OB-cadherin (osteoblast) | 56 | 0.276 | 0.0469 | No |
| 3 | IL15 | interleukin 15 | 64 | 0.261 | 0.0688 | No |
| 4 | DKK1 | dickkopf WNT signaling pathway inhibitor 1 | 106 | 0.204 | 0.0841 | No |
| 5 | GREM1 | gremlin 1, DAN family BMP antagonist | 170 | 0.165 | 0.0949 | No |
| 6 | SERPINE2 | serpin peptidase inhibitor, clade E (nexin, plasminogen activator inhibitor type 1), member 2 | 171 | 0.165 | 0.1090 | No |
| 7 | FAS | Fas cell surface death receptor | 186 | 0.161 | 0.1220 | No |
| 8 | THBS1 | thrombospondin 1 | 192 | 0.158 | 0.1352 | No |
| 9 | TIMP1 | TIMP metallopeptidase inhibitor 1 | 254 | 0.141 | 0.1441 | No |
| 10 | CXCL8 | chemokine (C-X-C motif) ligand 8 | 350 | 0.125 | 0.1498 | No |
| 11 | VIM | vimentin | 489 | 0.111 | 0.1522 | No |
| 12 | CD44 | CD44 molecule (Indian blood group) | 577 | 0.104 | 0.1566 | No |
| 13 | IGFBP3 | insulin like growth factor binding protein 3 | 582 | 0.104 | 0.1653 | No |
| 14 | CXCL1 | chemokine (C-X-C motif) ligand 1 (melanoma growth stimulating activity, alpha) | 672 | 0.098 | 0.1691 | No |
| 15 | TGM2 | transglutaminase 2 | 714 | 0.096 | 0.1752 | No |
| 16 | ACTA2 | actin, alpha 2, smooth muscle, aorta | 746 | 0.095 | 0.1816 | No |
| 17 | IGFBP4 | insulin like growth factor binding protein 4 | 792 | 0.093 | 0.1872 | No |
| 18 | ITGB5 | integrin beta 5 | 810 | 0.092 | 0.1942 | No |
| 19 | ECM2 | extracellular matrix protein 2, female organ and adipocyte specific | 1051 | 0.082 | 0.1888 | No |
| 20 | FBLN1 | fibulin 1 | 1105 | 0.081 | 0.1929 | No |
| 21 | FBLN5 | fibulin 5 | 1120 | 0.080 | 0.1991 | No |
| 22 | TGFBI | transforming growth factor, beta-induced, 68kDa | 1130 | 0.080 | 0.2054 | No |
| 23 | ID2 | inhibitor of DNA binding 2, dominant negative helix-loop-helix protein | 1150 | 0.080 | 0.2113 | No |
| 24 | NID2 | nidogen 2 (osteonidogen) | 1228 | 0.078 | 0.2139 | No |
| 25 | PLAUR | plasminogen activator, urokinase receptor | 1355 | 0.075 | 0.2138 | No |
| 26 | BDNF | brain-derived neurotrophic factor | 1505 | 0.071 | 0.2122 | No |
| 27 | SLIT3 | slit guidance ligand 3 | 1612 | 0.069 | 0.2126 | No |
| 28 | ELN | elastin | 1766 | 0.066 | 0.2102 | No |
| 29 | FSTL3 | follistatin-like 3 (secreted glycoprotein) | 1974 | 0.062 | 0.2048 | No |
| 30 | NOTCH2 | notch 2 | 2091 | 0.060 | 0.2039 | No |
| 31 | TGFBR3 | transforming growth factor beta receptor III | 2120 | 0.060 | 0.2076 | No |
| 32 | HTRA1 | HtrA serine peptidase 1 | 2295 | 0.057 | 0.2034 | No |
| 33 | PDGFRB | platelet-derived growth factor receptor, beta polypeptide | 2343 | 0.056 | 0.2058 | No |
| 34 | PTX3 | pentraxin 3, long | 2893 | 0.049 | 0.1815 | No |
| 35 | ENO2 | enolase 2 (gamma, neuronal) | 3086 | 0.047 | 0.1756 | No |
| 36 | SPOCK1 | sparc/osteonectin, cwcv and kazal-like domains proteoglycan (testican) 1 | 3906 | 0.038 | 0.1365 | No |
| 37 | MATN3 | matrilin 3 | 3941 | 0.038 | 0.1380 | No |
| 38 | LRRC15 | leucine rich repeat containing 15 | 4102 | 0.037 | 0.1328 | No |
| 39 | FBN2 | fibrillin 2 | 4127 | 0.036 | 0.1347 | No |
| 40 | GEM | GTP binding protein overexpressed in skeletal muscle | 4213 | 0.035 | 0.1333 | No |
| 41 | TPM1 | tropomyosin 1 (alpha) | 4297 | 0.035 | 0.1320 | No |
| 42 | COL4A2 | collagen, type IV, alpha 2 | 4487 | 0.033 | 0.1250 | No |
| 43 | THY1 | Thy-1 cell surface antigen | 4641 | 0.032 | 0.1199 | No |
| 44 | TIMP3 | TIMP metallopeptidase inhibitor 3 | 4889 | 0.030 | 0.1097 | No |
| 45 | LAMA1 | laminin, alpha 1 | 5151 | 0.028 | 0.0986 | No |
| 46 | RGS4 | regulator of G-protein signaling 4 | 5282 | 0.027 | 0.0941 | No |
| 47 | SDC1 | syndecan 1 | 5305 | 0.027 | 0.0953 | No |
| 48 | PCOLCE2 | procollagen C-endopeptidase enhancer 2 | 5344 | 0.027 | 0.0956 | No |
| 49 | CAPG | capping protein (actin filament), gelsolin-like | 5399 | 0.026 | 0.0951 | No |
| 50 | SLIT2 | slit guidance ligand 2 | 5509 | 0.025 | 0.0916 | No |
| 51 | GADD45B | growth arrest and DNA-damage-inducible, beta | 5568 | 0.025 | 0.0907 | No |
| 52 | COL6A2 | collagen, type VI, alpha 2 | 5593 | 0.025 | 0.0916 | No |
| 53 | GLIPR1 | GLI pathogenesis-related 1 | 5605 | 0.025 | 0.0931 | No |
| 54 | COL4A1 | collagen, type IV, alpha 1 | 5717 | 0.024 | 0.0894 | No |
| 55 | LOXL1 | lysyl oxidase-like 1 | 5981 | 0.022 | 0.0777 | No |
| 56 | DPYSL3 | dihydropyrimidinase-like 3 | 6038 | 0.022 | 0.0767 | No |
| 57 | OXTR | oxytocin receptor | 6074 | 0.022 | 0.0767 | No |
| 58 | LAMA2 | laminin, alpha 2 | 6270 | 0.020 | 0.0684 | No |
| 59 | CD59 | CD59 molecule, complement regulatory protein | 6466 | 0.019 | 0.0599 | No |
| 60 | MAGEE1 | MAGE family member E1 | 6491 | 0.019 | 0.0603 | No |
| 61 | IL32 | interleukin 32 | 6502 | 0.019 | 0.0614 | No |
| 62 | PTHLH | parathyroid hormone-like hormone | 6661 | 0.018 | 0.0547 | No |
| 63 | NTM | neurotrimin | 6751 | 0.017 | 0.0516 | No |
| 64 | TNFRSF12A | tumor necrosis factor receptor superfamily, member 12A | 6787 | 0.017 | 0.0513 | No |
| 65 | PMP22 | peripheral myelin protein 22 | 7003 | 0.016 | 0.0415 | No |
| 66 | SGCB | sarcoglycan beta | 7047 | 0.016 | 0.0406 | No |
| 67 | DAB2 | Dab, mitogen-responsive phosphoprotein, homolog 2 (Drosophila) | 7140 | 0.015 | 0.0371 | No |
| 68 | GADD45A | growth arrest and DNA-damage-inducible, alpha | 7290 | 0.014 | 0.0306 | No |
| 69 | COL1A1 | Jeck2013 ANTISENSE, coding, INTERNAL, intronic best transcript NM\_000088 | 7296 | 0.014 | 0.0316 | No |
| 70 | CXCL12 | chemokine (C-X-C motif) ligand 12 | 7452 | 0.013 | 0.0247 | No |
| 71 | MGP | matrix Gla protein | 7495 | 0.013 | 0.0236 | No |
| 72 | COL7A1 | collagen, type VII, alpha 1 | 7508 | 0.013 | 0.0241 | No |
| 73 | COMP | cartilage oligomeric matrix protein | 7536 | 0.013 | 0.0238 | No |
| 74 | BMP1 | bone morphogenetic protein 1 | 7539 | 0.013 | 0.0247 | No |
| 75 | COL1A2 | collagen, type I, alpha 2 | 7583 | 0.012 | 0.0236 | No |
| 76 | CDH2 | cadherin 2, type 1, N-cadherin (neuronal) | 7774 | 0.011 | 0.0147 | No |
| 77 | MYL9 | myosin light chain 9 | 7810 | 0.011 | 0.0138 | No |
| 78 | PCOLCE | procollagen C-endopeptidase enhancer | 7847 | 0.011 | 0.0129 | No |
| 79 | ECM1 | extracellular matrix protein 1 | 7942 | 0.010 | 0.0089 | No |
| 80 | MCM7 | minichromosome maintenance complex component 7 | 7985 | 0.010 | 0.0076 | No |
| 81 | FZD8 | frizzled class receptor 8 | 8074 | 0.009 | 0.0038 | No |
| 82 | SGCD | sarcoglycan delta | 8224 | 0.008 | -0.0032 | No |
| 83 | COL11A1 | collagen, type XI, alpha 1 | 8244 | 0.008 | -0.0034 | No |
| 84 | CADM1 | cell adhesion molecule 1 | 8577 | 0.006 | -0.0201 | No |
| 85 | COLGALT1 | collagen beta(1-O)galactosyltransferase 1 | 8595 | 0.006 | -0.0204 | No |
| 86 | ITGA5 | integrin alpha 5 | 8613 | 0.006 | -0.0207 | No |
| 87 | MYLK | myosin light chain kinase | 8846 | 0.005 | -0.0323 | No |
| 88 | MMP14 | matrix metallopeptidase 14 (membrane-inserted) | 9110 | 0.003 | -0.0456 | No |
| 89 | TPM4 | tropomyosin 4 | 9464 | 0.002 | -0.0638 | No |
| 90 | COL6A3 | collagen, type VI, alpha 3 | 9469 | 0.002 | -0.0638 | No |
| 91 | LRP1 | LDL receptor related protein 1 | 9659 | 0.001 | -0.0735 | No |
| 92 | PMEPA1 | prostate transmembrane protein, androgen induced 1 | 9736 | 0.000 | -0.0775 | No |
| 93 | FLNA | Jeck2013 ANTISENSE, CDS, coding, INTERNAL, OVCODE, OVEXON best transcript NM\_001110556 | 9902 | -0.001 | -0.0859 | No |
| 94 | SCG2 | secretogranin II | 10022 | -0.001 | -0.0920 | No |
| 95 | MMP3 | matrix metallopeptidase 3 | 10688 | -0.005 | -0.1259 | No |
| 96 | SLC6A8 | solute carrier family 6 (neurotransmitter transporter), member 8 | 10735 | -0.006 | -0.1278 | No |
| 97 | PLOD1 | procollagen-lysine, 2-oxoglutarate 5-dioxygenase 1 | 10966 | -0.007 | -0.1392 | No |
| 98 | GAS1 | growth arrest-specific 1 | 11303 | -0.009 | -0.1558 | No |
| 99 | COL16A1 | collagen, type XVI, alpha 1 | 11406 | -0.009 | -0.1603 | No |
| 100 | WIPF1 | WAS/WASL interacting protein family, member 1 | 11506 | -0.010 | -0.1645 | No |
| 101 | SPP1 | secreted phosphoprotein 1 | 11575 | -0.010 | -0.1672 | No |
| 102 | GPC1 | glypican 1 | 11601 | -0.010 | -0.1676 | No |
| 103 | COL3A1 | collagen, type III, alpha 1 | 11694 | -0.011 | -0.1714 | No |
| 104 | P3H1 | prolyl 3-hydroxylase 1 | 11724 | -0.011 | -0.1720 | No |
| 105 | IL6 | interleukin 6 | 11825 | -0.012 | -0.1761 | No |
| 106 | CXCL6 | chemokine (C-X-C motif) ligand 6 | 11900 | -0.012 | -0.1789 | No |
| 107 | PFN2 | profilin 2 | 12101 | -0.013 | -0.1881 | No |
| 108 | TPM2 | tropomyosin 2 (beta) | 12172 | -0.014 | -0.1906 | No |
| 109 | FERMT2 | fermitin family member 2 | 12188 | -0.014 | -0.1901 | No |
| 110 | FMOD | fibromodulin | 12274 | -0.015 | -0.1933 | No |
| 111 | TAGLN | transgelin | 12350 | -0.015 | -0.1959 | No |
| 112 | PDLIM4 | PDZ and LIM domain 4 | 12385 | -0.015 | -0.1963 | No |
| 113 | APLP1 | amyloid beta (A4) precursor-like protein 1 | 12392 | -0.015 | -0.1953 | No |
| 114 | LAMA3 | laminin, alpha 3 | 12397 | -0.015 | -0.1942 | No |
| 115 | TGFB1 | transforming growth factor beta 1 | 12464 | -0.016 | -0.1963 | No |
| 116 | FGF2 | fibroblast growth factor 2 (basic) | 12468 | -0.016 | -0.1951 | No |
| 117 | ADAM12 | ADAM metallopeptidase domain 12 | 12524 | -0.016 | -0.1965 | No |
| 118 | COL5A2 | collagen, type V, alpha 2 | 12761 | -0.018 | -0.2072 | No |
| 119 | ITGB3 | integrin beta 3 | 12821 | -0.018 | -0.2087 | No |
| 120 | CRLF1 | cytokine receptor-like factor 1 | 12865 | -0.018 | -0.2094 | No |
| 121 | SERPINE1 | serpin peptidase inhibitor, clade E (nexin, plasminogen activator inhibitor type 1), member 1 | 12974 | -0.019 | -0.2134 | No |
| 122 | FUCA1 | fucosidase, alpha-L- 1, tissue | 13114 | -0.020 | -0.2188 | No |
| 123 | MATN2 | matrilin 2 | 13189 | -0.020 | -0.2209 | No |
| 124 | TNC | tenascin C | 13324 | -0.021 | -0.2260 | No |
| 125 | TNFAIP3 | tumor necrosis factor, alpha-induced protein 3 | 13363 | -0.022 | -0.2261 | No |
| 126 | CAP2 | CAP, adenylate cyclase-associated protein, 2 (yeast) | 13575 | -0.023 | -0.2351 | No |
| 127 | ABI3BP | ABI family, member 3 (NESH) binding protein | 13728 | -0.024 | -0.2409 | No |
| 128 | FSTL1 | follistatin like 1 | 13956 | -0.026 | -0.2504 | No |
| 129 | FOXC2 | forkhead box C2 | 14118 | -0.027 | -0.2564 | No |
| 130 | PVR | poliovirus receptor | 14143 | -0.027 | -0.2553 | No |
| 131 | SNAI2 | snail family zinc finger 2 | 14230 | -0.028 | -0.2574 | No |
| 132 | VCAM1 | vascular cell adhesion molecule 1 | 14397 | -0.030 | -0.2634 | No |
| 133 | TFPI2 | tissue factor pathway inhibitor 2 | 14489 | -0.030 | -0.2655 | No |
| 134 | SNTB1 | syntrophin, beta 1 (dystrophin-associated protein A1, 59kDa, basic component 1) | 14515 | -0.030 | -0.2642 | No |
| 135 | COL5A3 | collagen, type V, alpha 3 | 14588 | -0.031 | -0.2653 | No |
| 136 | NNMT | nicotinamide N-methyltransferase | 14699 | -0.032 | -0.2683 | No |
| 137 | COL8A2 | collagen, type VIII, alpha 2 | 14793 | -0.033 | -0.2703 | No |
| 138 | SERPINH1 | serpin peptidase inhibitor, clade H (heat shock protein 47), member 1, (collagen binding protein 1) | 14845 | -0.033 | -0.2701 | No |
| 139 | SFRP4 | secreted frizzled-related protein 4 | 14940 | -0.034 | -0.2721 | No |
| 140 | EFEMP2 | EGF containing fibulin-like extracellular matrix protein 2 | 15043 | -0.035 | -0.2744 | No |
| 141 | INHBA | inhibin beta A | 15087 | -0.035 | -0.2736 | No |
| 142 | BASP1 | brain abundant, membrane attached signal protein 1 | 15107 | -0.035 | -0.2716 | No |
| 143 | DCN | decorin | 15284 | -0.037 | -0.2775 | No |
| 144 | THBS2 | thrombospondin 2 | 15353 | -0.037 | -0.2779 | No |
| 145 | ITGA2 | integrin, alpha 2 (CD49B, alpha 2 subunit of VLA-2 receptor) | 15498 | -0.039 | -0.2820 | No |
| 146 | GPX7 | glutathione peroxidase 7 | 15589 | -0.040 | -0.2833 | No |
| 147 | PLOD3 | procollagen-lysine, 2-oxoglutarate 5-dioxygenase 3 | 15596 | -0.040 | -0.2802 | No |
| 148 | EDIL3 | EGF-like repeats and discoidin I-like domains 3 | 15639 | -0.040 | -0.2789 | No |
| 149 | MXRA5 | matrix-remodelling associated 5 | 15690 | -0.041 | -0.2781 | No |
| 150 | CTHRC1 | collagen triple helix repeat containing 1 | 15779 | -0.042 | -0.2791 | No |
| 151 | LAMC2 | laminin, gamma 2 | 15989 | -0.044 | -0.2862 | No |
| 152 | FBLN2 | fibulin 2 | 16030 | -0.044 | -0.2845 | No |
| 153 | ITGAV | integrin alpha V | 16105 | -0.045 | -0.2845 | No |
| 154 | MSX1 | msh homeobox 1 | 16206 | -0.046 | -0.2857 | No |
| 155 | FBN1 | fibrillin 1 | 16292 | -0.047 | -0.2861 | No |
| 156 | CALD1 | caldesmon 1 | 16317 | -0.047 | -0.2833 | No |
| 157 | PPIB | peptidylprolyl isomerase B (cyclophilin B) | 16361 | -0.048 | -0.2814 | No |
| 158 | AREG | amphiregulin | 16601 | -0.051 | -0.2894 | No |
| 159 | VEGFA | vascular endothelial growth factor A | 16659 | -0.052 | -0.2879 | No |
| 160 | EMP3 | epithelial membrane protein 3 | 16695 | -0.052 | -0.2853 | No |
| 161 | ANPEP | alanyl (membrane) aminopeptidase | 17087 | -0.058 | -0.3005 | No |
| 162 | VEGFC | vascular endothelial growth factor C | 17126 | -0.059 | -0.2975 | No |
| 163 | LOX | lysyl oxidase | 17422 | -0.064 | -0.3073 | Yes |
| 164 | LAMC1 | laminin, gamma 1 (formerly LAMB2) | 17462 | -0.065 | -0.3038 | Yes |
| 165 | PRSS2 | protease, serine, 2 (trypsin 2) | 17554 | -0.066 | -0.3028 | Yes |
| 166 | PLOD2 | procollagen-lysine, 2-oxoglutarate 5-dioxygenase 2 | 17575 | -0.067 | -0.2982 | Yes |
| 167 | QSOX1 | quiescin Q6 sulfhydryl oxidase 1 | 17654 | -0.069 | -0.2964 | Yes |
| 168 | ITGB1 | integrin beta 1 | 17737 | -0.070 | -0.2946 | Yes |
| 169 | WNT5A | wingless-type MMTV integration site family, member 5A | 17896 | -0.074 | -0.2964 | Yes |
| 170 | MFAP5 | microfibrillar associated protein 5 | 18013 | -0.077 | -0.2958 | Yes |
| 171 | COPA | coatomer protein complex subunit alpha | 18029 | -0.078 | -0.2900 | Yes |
| 172 | SDC4 | syndecan 4 | 18151 | -0.081 | -0.2893 | Yes |
| 173 | SFRP1 | secreted frizzled-related protein 1 | 18263 | -0.084 | -0.2879 | Yes |
| 174 | SGCG | sarcoglycan gamma | 18273 | -0.085 | -0.2811 | Yes |
| 175 | SAT1 | spermidine/spermine N1-acetyltransferase 1 | 18295 | -0.085 | -0.2749 | Yes |
| 176 | SPARC | secreted protein, acidic, cysteine-rich (osteonectin) | 18622 | -0.097 | -0.2835 | Yes |
| 177 | VCAN | versican | 18696 | -0.103 | -0.2785 | Yes |
| 178 | CALU | calumenin | 18707 | -0.103 | -0.2702 | Yes |
| 179 | RHOB | ras homolog family member B | 18827 | -0.112 | -0.2668 | Yes |
| 180 | PRRX1 | paired related homeobox 1 | 18904 | -0.118 | -0.2607 | Yes |
| 181 | POSTN | periostin, osteoblast specific factor | 18928 | -0.119 | -0.2517 | Yes |
| 182 | LGALS1 | lectin, galactoside-binding, soluble, 1 | 18961 | -0.123 | -0.2429 | Yes |
| 183 | MEST | mesoderm specific transcript | 18998 | -0.127 | -0.2340 | Yes |
| 184 | COL12A1 | collagen, type XII, alpha 1 | 18999 | -0.127 | -0.2232 | Yes |
| 185 | JUN | jun proto-oncogene | 19037 | -0.132 | -0.2138 | Yes |
| 186 | FAP | fibroblast activation protein alpha | 19074 | -0.138 | -0.2039 | Yes |
| 187 | TNFRSF11B | tumor necrosis factor receptor superfamily, member 11b | 19143 | -0.148 | -0.1948 | Yes |
| 188 | CDH6 | cadherin 6, type 2, K-cadherin (fetal kidney) | 19155 | -0.151 | -0.1825 | Yes |
| 189 | MMP2 | matrix metallopeptidase 2 | 19166 | -0.152 | -0.1700 | Yes |
| 190 | IGFBP2 | insulin like growth factor binding protein 2 | 19180 | -0.155 | -0.1574 | Yes |
| 191 | MMP1 | matrix metallopeptidase 1 | 19272 | -0.181 | -0.1466 | Yes |
| 192 | GJA1 | gap junction protein alpha 1 | 19360 | -0.217 | -0.1326 | Yes |
| 193 | LUM | lumican | 19432 | -0.269 | -0.1134 | Yes |
| 194 | FN1 | fibronectin 1 | 19439 | -0.280 | -0.0897 | Yes |
| 195 | COL5A1 | collagen, type V, alpha 1 | 19466 | -0.322 | -0.0636 | Yes |
| 196 | DST | dystonin | 19494 | -0.380 | -0.0325 | Yes |
| 197 | BGN | biglycan | 19498 | -0.399 | 0.0013 | Yes |
Table: GSEA details [plain text format]

  

Fig 2: HALLMARK\_EPITHELIAL\_MESENCHYMAL\_TRANSITION      
 Blue-Pink O' Gram in the Space of the Analyzed GeneSet

  

Fig 3: HALLMARK\_EPITHELIAL\_MESENCHYMAL\_TRANSITION: Random ES distribution      
 Gene set null distribution of ES for **HALLMARK\_EPITHELIAL\_MESENCHYMAL\_TRANSITION**

  
