## Supplementary material for "Somatic hypomethylation of pericentromeric SST1 repeats and tetraploidization in human colorectal cancer cells": GSEA results: HALLMARK_ESTROGEN_RESPONSE_EARLY.html

Details for gene set HALLMARK\_ESTROGEN\_RESPONSE\_EARLY[GSEA]

|  || Dataset | eset\_byprobe\_collapsed\_to\_symbols.Diploid\_vs\_Tetraploid.cls #Tetraploid\_versus\_Diploid.Diploid\_vs\_Tetraploid.cls #Tetraploid\_versus\_Diploid\_repos |
| Phenotype | Diploid\_vs\_Tetraploid.cls#Tetraploid\_versus\_Diploid\_repos |
| Upregulated in class | Tetraploid |
| GeneSet | HALLMARK\_ESTROGEN\_RESPONSE\_EARLY |
| Enrichment Score (ES) | 0.29826438 |
| Normalized Enrichment Score (NES) | 1.3154887 |
| Nominal p-value | 0.023809524 |
| FDR q-value | 0.15206209 |
| FWER p-Value | 0.607 |
Table: GSEA Results Summary

  

Fig 1: Enrichment plot: HALLMARK\_ESTROGEN\_RESPONSE\_EARLY      
 Profile of the Running ES Score & Positions of GeneSet Members on the Rank Ordered List

  

| SYMBOL | TITLE | RANK IN GENE LIST | RANK METRIC SCORE | RUNNING ES | CORE ENRICHMENT || 1 | CALB2 | calbindin 2 | 55 | 0.277 | 0.0255 | Yes |
| 2 | KLK10 | kallikrein related peptidase 10 | 99 | 0.209 | 0.0447 | Yes |
| 3 | WFS1 | Wolfram syndrome 1 (wolframin) | 150 | 0.172 | 0.0597 | Yes |
| 4 | KRT19 | keratin 19, type I | 157 | 0.170 | 0.0768 | Yes |
| 5 | FASN | fatty acid synthase | 293 | 0.135 | 0.0836 | Yes |
| 6 | MYB | v-myb avian myeloblastosis viral oncogene homolog | 306 | 0.133 | 0.0965 | Yes |
| 7 | ABCA3 | ATP binding cassette subfamily A member 3 | 321 | 0.129 | 0.1090 | Yes |
| 8 | TOB1 | transducer of ERBB2, 1 | 343 | 0.127 | 0.1209 | Yes |
| 9 | CLIC3 | chloride intracellular channel 3 | 510 | 0.109 | 0.1235 | Yes |
| 10 | MYBL1 | v-myb avian myeloblastosis viral oncogene homolog-like 1 | 518 | 0.108 | 0.1342 | Yes |
| 11 | KCNK5 | potassium channel, two pore domain subfamily K, member 5 | 550 | 0.106 | 0.1434 | Yes |
| 12 | TFAP2C | transcription factor AP-2 gamma (activating enhancer binding protein 2 gamma) | 567 | 0.105 | 0.1533 | Yes |
| 13 | CD44 | CD44 molecule (Indian blood group) | 577 | 0.104 | 0.1636 | Yes |
| 14 | TGM2 | transglutaminase 2 | 714 | 0.096 | 0.1664 | Yes |
| 15 | IGFBP4 | insulin like growth factor binding protein 4 | 792 | 0.093 | 0.1719 | Yes |
| 16 | FDFT1 | farnesyl-diphosphate farnesyltransferase 1 | 793 | 0.093 | 0.1813 | Yes |
| 17 | FKBP4 | FK506 binding protein 4 | 973 | 0.085 | 0.1808 | Yes |
| 18 | RHOBTB3 | Rho-related BTB domain containing 3 | 1028 | 0.083 | 0.1865 | Yes |
| 19 | CHPT1 | choline phosphotransferase 1 | 1092 | 0.081 | 0.1916 | Yes |
| 20 | FOXC1 | forkhead box C1 | 1152 | 0.080 | 0.1967 | Yes |
| 21 | BAG1 | BCL2-associated athanogene | 1176 | 0.079 | 0.2036 | Yes |
| 22 | ADCY9 | adenylate cyclase 9 | 1177 | 0.079 | 0.2116 | Yes |
| 23 | ISG20L2 | interferon stimulated exonuclease gene 20kDa like 2 | 1180 | 0.079 | 0.2196 | Yes |
| 24 | ZNF185 | zinc finger protein 185 (LIM domain) | 1295 | 0.076 | 0.2215 | Yes |
| 25 | SYNGR1 | synaptogyrin 1 | 1328 | 0.076 | 0.2276 | Yes |
| 26 | MYOF | myoferlin | 1329 | 0.076 | 0.2353 | Yes |
| 27 | TUBB2B | tubulin, beta 2B class IIb | 1597 | 0.069 | 0.2286 | Yes |
| 28 | ANXA9 | annexin A9 | 1673 | 0.067 | 0.2316 | Yes |
| 29 | SLC1A1 | solute carrier family 1 (neuronal/epithelial high affinity glutamate transporter, system Xag), member 1 | 1678 | 0.067 | 0.2382 | Yes |
| 30 | MSMB | microseminoprotein, beta- | 1770 | 0.065 | 0.2402 | Yes |
| 31 | FKBP5 | FK506 binding protein 5 | 1840 | 0.064 | 0.2432 | Yes |
| 32 | RAPGEFL1 | Rap guanine nucleotide exchange factor like 1 | 1872 | 0.064 | 0.2481 | Yes |
| 33 | DHCR7 | 7-dehydrocholesterol reductase | 1877 | 0.064 | 0.2544 | Yes |
| 34 | FLNB | filamin B, beta | 1902 | 0.063 | 0.2596 | Yes |
| 35 | NXT1 | nuclear transport factor 2-like export factor 1 | 1914 | 0.063 | 0.2655 | Yes |
| 36 | PTGES | prostaglandin E synthase | 1996 | 0.061 | 0.2676 | Yes |
| 37 | TGIF2 | TGFB-induced factor homeobox 2 | 2001 | 0.061 | 0.2736 | Yes |
| 38 | SFN | stratifin | 2005 | 0.061 | 0.2798 | Yes |
| 39 | GLA | galactosidase, alpha | 2101 | 0.060 | 0.2810 | Yes |
| 40 | KRT13 | keratin 13, type I | 2215 | 0.058 | 0.2811 | Yes |
| 41 | GREB1 | growth regulation by estrogen in breast cancer 1 | 2312 | 0.057 | 0.2819 | Yes |
| 42 | LAD1 | ladinin 1 | 2341 | 0.056 | 0.2862 | Yes |
| 43 | TIAM1 | T-cell lymphoma invasion and metastasis 1 | 2375 | 0.056 | 0.2902 | Yes |
| 44 | TMEM164 | transmembrane protein 164 | 2378 | 0.056 | 0.2958 | Yes |
| 45 | CA12 | carbonic anhydrase XII | 2439 | 0.055 | 0.2983 | Yes |
| 46 | KAZN | kazrin, periplakin interacting protein | 2559 | 0.053 | 0.2975 | No |
| 47 | NADSYN1 | NAD synthetase 1 | 2674 | 0.051 | 0.2969 | No |
| 48 | IL17RB | interleukin 17 receptor B | 2765 | 0.050 | 0.2974 | No |
| 49 | SLC16A1 | solute carrier family 16 (monocarboxylate transporter), member 1 | 2936 | 0.048 | 0.2935 | No |
| 50 | MYC | v-myc avian myelocytomatosis viral oncogene homolog | 3194 | 0.046 | 0.2849 | No |
| 51 | SLC19A2 | solute carrier family 19 (thiamine transporter), member 2 | 3211 | 0.045 | 0.2887 | No |
| 52 | SLC9A3R1 | solute carrier family 9, subfamily A (NHE3, cation proton antiporter 3), member 3 regulator 1 | 3344 | 0.044 | 0.2864 | No |
| 53 | MAST4 | microtubule associated serine/threonine kinase family member 4 | 3552 | 0.042 | 0.2800 | No |
| 54 | KRT18 | keratin 18, type I | 3794 | 0.040 | 0.2716 | No |
| 55 | KRT8 | keratin 8, type II | 3950 | 0.038 | 0.2674 | No |
| 56 | OVOL2 | ovo-like zinc finger 2 | 3978 | 0.038 | 0.2699 | No |
| 57 | KDM4B | lysine (K)-specific demethylase 4B | 4437 | 0.034 | 0.2496 | No |
| 58 | NRIP1 | nuclear receptor interacting protein 1 | 4528 | 0.033 | 0.2483 | No |
| 59 | GFRA1 | GDNF family receptor alpha 1 | 4572 | 0.033 | 0.2494 | No |
| 60 | CCND1 | cyclin D1 | 4585 | 0.032 | 0.2521 | No |
| 61 | FCMR | Fc fragment of IgM receptor | 4591 | 0.032 | 0.2552 | No |
| 62 | TPD52L1 | tumor protein D52-like 1 | 4623 | 0.032 | 0.2569 | No |
| 63 | ABHD2 | abhydrolase domain containing 2 | 4774 | 0.031 | 0.2523 | No |
| 64 | RET | ret proto-oncogene | 4931 | 0.030 | 0.2473 | No |
| 65 | TBC1D30 | TBC1 domain family, member 30 | 4970 | 0.030 | 0.2483 | No |
| 66 | ELF3 | E74-like factor 3 (ets domain transcription factor, epithelial-specific ) | 5265 | 0.027 | 0.2359 | No |
| 67 | HES1 | hes family bHLH transcription factor 1 | 5409 | 0.026 | 0.2312 | No |
| 68 | SNX24 | sorting nexin 24 | 5477 | 0.026 | 0.2303 | No |
| 69 | ADCY1 | adenylate cyclase 1 (brain) | 5528 | 0.025 | 0.2303 | No |
| 70 | MREG | melanoregulin | 5542 | 0.025 | 0.2322 | No |
| 71 | RASGRP1 | RAS guanyl releasing protein 1 (calcium and DAG-regulated) | 5558 | 0.025 | 0.2340 | No |
| 72 | ELOVL5 | ELOVL fatty acid elongase 5 | 5798 | 0.023 | 0.2240 | No |
| 73 | SH3BP5 | SH3-domain binding protein 5 (BTK-associated) | 5841 | 0.023 | 0.2242 | No |
| 74 | BLVRB | biliverdin reductase B | 5918 | 0.023 | 0.2226 | No |
| 75 | RAB31 | Transcript Identified by AceView, Entrez Gene ID(s) 11031 | 6174 | 0.021 | 0.2116 | No |
| 76 | RRP12 | Zhang2013 ALT\_ACCEPTOR, ALT\_DONOR, coding, INTERNAL, intronic best transcript NM\_015179 | 6340 | 0.020 | 0.2051 | No |
| 77 | OLFM1 | olfactomedin 1 | 6358 | 0.020 | 0.2062 | No |
| 78 | SYT12 | synaptotagmin XII | 6363 | 0.020 | 0.2081 | No |
| 79 | PEX11A | peroxisomal biogenesis factor 11 alpha | 6645 | 0.018 | 0.1954 | No |
| 80 | SLC27A2 | solute carrier family 27 (fatty acid transporter), member 2 | 6687 | 0.018 | 0.1951 | No |
| 81 | ABAT | 4-aminobutyrate aminotransferase | 6739 | 0.017 | 0.1942 | No |
| 82 | MAPT | microtubule associated protein tau | 6748 | 0.017 | 0.1956 | No |
| 83 | TMPRSS3 | transmembrane protease, serine 3 | 6794 | 0.017 | 0.1950 | No |
| 84 | PPIF | peptidylprolyl isomerase F | 7224 | 0.015 | 0.1743 | No |
| 85 | REEP1 | receptor accessory protein 1 | 7450 | 0.013 | 0.1640 | No |
| 86 | CXCL12 | chemokine (C-X-C motif) ligand 12 | 7452 | 0.013 | 0.1653 | No |
| 87 | MLPH | melanophilin | 7512 | 0.013 | 0.1635 | No |
| 88 | ELOVL2 | ELOVL fatty acid elongase 2 | 7670 | 0.012 | 0.1566 | No |
| 89 | MYBBP1A | MYB binding protein (P160) 1a | 7769 | 0.011 | 0.1527 | No |
| 90 | CALCR | calcitonin receptor | 7944 | 0.010 | 0.1448 | No |
| 91 | ESRP2 | epithelial splicing regulatory protein 2 | 7965 | 0.010 | 0.1448 | No |
| 92 | ELF1 | E74-like factor 1 (ets domain transcription factor) | 8011 | 0.010 | 0.1434 | No |
| 93 | FHL2 | four and a half LIM domains 2 | 8060 | 0.010 | 0.1419 | No |
| 94 | SLC22A5 | solute carrier family 22 (organic cation/carnitine transporter), member 5 | 8098 | 0.009 | 0.1410 | No |
| 95 | MED24 | mediator complex subunit 24 | 8183 | 0.009 | 0.1375 | No |
| 96 | BCL11B | B-cell CLL/lymphoma 11B (zinc finger protein) | 8629 | 0.006 | 0.1151 | No |
| 97 | SEC14L2 | SEC14-like lipid binding 2 | 8776 | 0.005 | 0.1081 | No |
| 98 | AKAP1 | A kinase (PRKA) anchor protein 1 | 8810 | 0.005 | 0.1069 | No |
| 99 | AQP3 | aquaporin 3 (Gill blood group) | 8812 | 0.005 | 0.1074 | No |
| 100 | MICB | MHC class I polypeptide-related sequence B | 8905 | 0.005 | 0.1031 | No |
| 101 | SCARB1 | scavenger receptor class B, member 1 | 8987 | 0.004 | 0.0994 | No |
| 102 | ASB13 | ankyrin repeat and SOCS box containing 13 | 9174 | 0.003 | 0.0900 | No |
| 103 | SLC39A6 | solute carrier family 39 (zinc transporter), member 6 | 9281 | 0.003 | 0.0848 | No |
| 104 | OPN3 | opsin 3 | 9365 | 0.002 | 0.0808 | No |
| 105 | THSD4 | thrombospondin type 1 domain containing 4 | 9489 | 0.002 | 0.0745 | No |
| 106 | DYNLT3 | dynein, light chain, Tctex-type 3 | 9497 | 0.001 | 0.0743 | No |
| 107 | SLC37A1 | solute carrier family 37 (glucose-6-phosphate transporter), member 1 | 10048 | -0.001 | 0.0460 | No |
| 108 | RARA | retinoic acid receptor, alpha | 10083 | -0.002 | 0.0444 | No |
| 109 | OLFML3 | olfactomedin like 3 | 10345 | -0.003 | 0.0313 | No |
| 110 | CBFA2T3 | core-binding factor, runt domain, alpha subunit 2; translocated to, 3 | 10641 | -0.005 | 0.0165 | No |
| 111 | TIPARP | TCDD-inducible poly(ADP-ribose) polymerase | 11003 | -0.007 | -0.0014 | No |
| 112 | SLC2A1 | solute carrier family 2 (facilitated glucose transporter), member 1 | 11178 | -0.008 | -0.0096 | No |
| 113 | RAB17 | RAB17, member RAS oncogene family | 11278 | -0.009 | -0.0138 | No |
| 114 | KRT15 | keratin 15, type I | 11281 | -0.009 | -0.0131 | No |
| 115 | SVIL | supervillin | 11496 | -0.010 | -0.0231 | No |
| 116 | SCNN1A | sodium channel, non voltage gated 1 alpha subunit | 11540 | -0.010 | -0.0243 | No |
| 117 | CELSR1 | cadherin, EGF LAG seven-pass G-type receptor 1 | 11656 | -0.011 | -0.0292 | No |
| 118 | TFF1 | trefoil factor 1 | 11711 | -0.011 | -0.0308 | No |
| 119 | DHRS3 | dehydrogenase/reductase (SDR family) member 3 | 11753 | -0.011 | -0.0318 | No |
| 120 | CLDN7 | claudin 7 | 11834 | -0.012 | -0.0347 | No |
| 121 | MUC1 | mucin 1, cell surface associated | 11994 | -0.013 | -0.0416 | No |
| 122 | INHBB | inhibin beta B | 12019 | -0.013 | -0.0416 | No |
| 123 | CELSR2 | cadherin, EGF LAG seven-pass G-type receptor 2 | 12052 | -0.013 | -0.0419 | No |
| 124 | TFF3 | trefoil factor 3 | 12114 | -0.014 | -0.0436 | No |
| 125 | NPY1R | neuropeptide Y receptor Y1 | 12162 | -0.014 | -0.0447 | No |
| 126 | NCOR2 | nuclear receptor corepressor 2 | 12339 | -0.015 | -0.0522 | No |
| 127 | UGCG | UDP-glucose ceramide glucosyltransferase | 12746 | -0.018 | -0.0714 | No |
| 128 | PRSS23 | protease, serine, 23 | 12809 | -0.018 | -0.0728 | No |
| 129 | PODXL | podocalyxin-like | 12859 | -0.018 | -0.0734 | No |
| 130 | NAV2 | neuron navigator 2 | 12912 | -0.019 | -0.0742 | No |
| 131 | JAK2 | Janus kinase 2 | 13031 | -0.019 | -0.0783 | No |
| 132 | WWC1 | WW and C2 domain containing 1 | 13116 | -0.020 | -0.0807 | No |
| 133 | DEPTOR | DEP domain containing MTOR-interacting protein | 13355 | -0.022 | -0.0908 | No |
| 134 | SIAH2 | siah E3 ubiquitin protein ligase 2 | 13372 | -0.022 | -0.0894 | No |
| 135 | IGF1R | insulin-like growth factor 1 receptor | 13395 | -0.022 | -0.0883 | No |
| 136 | ENDOD1 | endonuclease domain containing 1 | 13399 | -0.022 | -0.0862 | No |
| 137 | PDLIM3 | PDZ and LIM domain 3 | 13450 | -0.022 | -0.0865 | No |
| 138 | NBL1 | neuroblastoma 1, DAN family BMP antagonist | 13497 | -0.023 | -0.0866 | No |
| 139 | KCNK15 | potassium channel, two pore domain subfamily K, member 15 | 13558 | -0.023 | -0.0873 | No |
| 140 | SLC24A3 | solute carrier family 24 (sodium/potassium/calcium exchanger), member 3 | 13559 | -0.023 | -0.0850 | No |
| 141 | FARP1 | FERM, ARH/RhoGEF and pleckstrin domain protein 1 | 14015 | -0.026 | -0.1058 | No |
| 142 | UNC119 | unc-119 lipid binding chaperone | 14123 | -0.027 | -0.1085 | No |
| 143 | CANT1 | calcium activated nucleotidase 1 | 14424 | -0.030 | -0.1210 | No |
| 144 | BCL2 | B-cell CLL/lymphoma 2 | 14503 | -0.030 | -0.1220 | No |
| 145 | SYBU | syntabulin (syntaxin-interacting) | 14555 | -0.031 | -0.1215 | No |
| 146 | DLC1 | DLC1 Rho GTPase activating protein | 14666 | -0.032 | -0.1239 | No |
| 147 | AR | androgen receptor | 14739 | -0.032 | -0.1243 | No |
| 148 | ARL3 | ADP-ribosylation factor like GTPase 3 | 14799 | -0.033 | -0.1240 | No |
| 149 | EGR3 | early growth response 3 | 14802 | -0.033 | -0.1208 | No |
| 150 | CYP26B1 | cytochrome P450, family 26, subfamily B, polypeptide 1 | 14910 | -0.034 | -0.1229 | No |
| 151 | SLC7A2 | solute carrier family 7 (cationic amino acid transporter, y+ system), member 2 | 14954 | -0.034 | -0.1216 | No |
| 152 | TJP3 | tight junction protein 3 | 15002 | -0.034 | -0.1205 | No |
| 153 | FRK | fyn-related Src family tyrosine kinase | 15076 | -0.035 | -0.1207 | No |
| 154 | SOX3 | SRY box 3 | 15166 | -0.036 | -0.1216 | No |
| 155 | RHOD | ras homolog family member D | 15233 | -0.036 | -0.1213 | No |
| 156 | ABLIM1 | actin binding LIM protein 1 | 15414 | -0.038 | -0.1267 | No |
| 157 | MPPED2 | metallophosphoesterase domain containing 2 | 15452 | -0.038 | -0.1247 | No |
| 158 | MED13L | mediator complex subunit 13-like | 15502 | -0.039 | -0.1233 | No |
| 159 | XBP1 | X-box binding protein 1 | 15622 | -0.040 | -0.1254 | No |
| 160 | CISH | cytokine inducible SH2-containing protein | 15667 | -0.040 | -0.1235 | No |
| 161 | AFF1 | AF4/FMR2 family, member 1 | 15691 | -0.041 | -0.1205 | No |
| 162 | INPP5F | inositol polyphosphate-5-phosphatase F | 15826 | -0.042 | -0.1232 | No |
| 163 | SLC26A2 | solute carrier family 26 (anion exchanger), member 2 | 15913 | -0.043 | -0.1232 | No |
| 164 | ADD3 | adducin 3 (gamma) | 16088 | -0.045 | -0.1276 | No |
| 165 | AMFR | autocrine motility factor receptor, E3 ubiquitin protein ligase | 16444 | -0.049 | -0.1410 | No |
| 166 | B4GALT1 | UDP-Gal:betaGlcNAc beta 1,4- galactosyltransferase, polypeptide 1 | 16445 | -0.049 | -0.1360 | No |
| 167 | ITPK1 | inositol-tetrakisphosphate 1-kinase | 16518 | -0.050 | -0.1346 | No |
| 168 | AREG | amphiregulin | 16601 | -0.051 | -0.1336 | No |
| 169 | RBBP8 | retinoblastoma binding protein 8 | 16689 | -0.052 | -0.1327 | No |
| 170 | TPBG | trophoblast glycoprotein | 17065 | -0.058 | -0.1462 | No |
| 171 | P2RY2 | purinergic receptor P2Y, G-protein coupled, 2 | 17123 | -0.059 | -0.1431 | No |
| 172 | STC2 | stanniocalcin 2 | 17326 | -0.062 | -0.1472 | No |
| 173 | TTC39A | tetratricopeptide repeat domain 39A | 17428 | -0.064 | -0.1459 | No |
| 174 | SULT2B1 | sulfotransferase family 2B member 1 | 17511 | -0.066 | -0.1434 | No |
| 175 | FOS | FBJ murine osteosarcoma viral oncogene homolog | 17528 | -0.066 | -0.1375 | No |
| 176 | GAB2 | GRB2-associated binding protein 2 | 17550 | -0.066 | -0.1318 | No |
| 177 | SEMA3B | sema domain, immunoglobulin domain (Ig), short basic domain, secreted, (semaphorin) 3B | 17591 | -0.067 | -0.1270 | No |
| 178 | FAM102A | Memczak2013 ANTISENSE, CDS, coding, INTERNAL, intronic, UTR3 best transcript NM\_203305 | 17638 | -0.068 | -0.1225 | No |
| 179 | KLF10 | Kruppel-like factor 10 | 17645 | -0.068 | -0.1158 | No |
| 180 | TSKU | tsukushi, small leucine rich proteoglycan | 17746 | -0.071 | -0.1137 | No |
| 181 | SLC7A5 | solute carrier family 7 (amino acid transporter light chain, L system), member 5 | 17849 | -0.073 | -0.1115 | No |
| 182 | PMAIP1 | phorbol-12-myristate-13-acetate-induced protein 1 | 17862 | -0.073 | -0.1046 | No |
| 183 | ALDH3B1 | aldehyde dehydrogenase 3 family, member B1 | 17969 | -0.076 | -0.1023 | No |
| 184 | KLF4 | Kruppel-like factor 4 (gut) | 17974 | -0.076 | -0.0947 | No |
| 185 | PAPSS2 | 3-phosphoadenosine 5-phosphosulfate synthase 2 | 18188 | -0.082 | -0.0973 | No |
| 186 | PGR | progesterone receptor | 18358 | -0.087 | -0.0971 | No |
| 187 | PDZK1 | PDZ domain containing 1 | 18426 | -0.090 | -0.0914 | No |
| 188 | HR | hair growth associated | 18647 | -0.099 | -0.0927 | No |
| 189 | SLC1A4 | solute carrier family 1 (glutamate/neutral amino acid transporter), member 4 | 18809 | -0.110 | -0.0897 | No |
| 190 | RPS6KA2 | ribosomal protein S6 kinase, 90kDa, polypeptide 2 | 18861 | -0.114 | -0.0806 | No |
| 191 | IL6ST | interleukin 6 signal transducer | 19134 | -0.147 | -0.0797 | No |
| 192 | HSPB8 | heat shock 22kDa protein 8 | 19194 | -0.160 | -0.0664 | No |
| 193 | BHLHE40 | basic helix-loop-helix family, member e40 | 19252 | -0.175 | -0.0514 | No |
| 194 | LRIG1 | leucine-rich repeats and immunoglobulin-like domains 1 | 19335 | -0.203 | -0.0349 | No |
| 195 | GJA1 | gap junction protein alpha 1 | 19360 | -0.217 | -0.0139 | No |
| 196 | DHRS2 | dehydrogenase/reductase (SDR family) member 2 | 19363 | -0.218 | 0.0083 | No |
Table: GSEA details [plain text format]

  

Fig 2: HALLMARK\_ESTROGEN\_RESPONSE\_EARLY      
 Blue-Pink O' Gram in the Space of the Analyzed GeneSet

  

Fig 3: HALLMARK\_ESTROGEN\_RESPONSE\_EARLY: Random ES distribution      
 Gene set null distribution of ES for **HALLMARK\_ESTROGEN\_RESPONSE\_EARLY**

  
