## Supplementary material for "Somatic hypomethylation of pericentromeric SST1 repeats and tetraploidization in human colorectal cancer cells": GSEA results: HALLMARK_ESTROGEN_RESPONSE_LATE.html

Details for gene set HALLMARK\_ESTROGEN\_RESPONSE\_LATE[GSEA]

|  || Dataset | eset\_byprobe\_collapsed\_to\_symbols.Diploid\_vs\_Tetraploid.cls #Tetraploid\_versus\_Diploid.Diploid\_vs\_Tetraploid.cls #Tetraploid\_versus\_Diploid\_repos |
| Phenotype | Diploid\_vs\_Tetraploid.cls#Tetraploid\_versus\_Diploid\_repos |
| Upregulated in class | Tetraploid |
| GeneSet | HALLMARK\_ESTROGEN\_RESPONSE\_LATE |
| Enrichment Score (ES) | 0.3011058 |
| Normalized Enrichment Score (NES) | 1.3432605 |
| Nominal p-value | 0.014836796 |
| FDR q-value | 0.13831808 |
| FWER p-Value | 0.531 |
Table: GSEA Results Summary

  

Fig 1: Enrichment plot: HALLMARK\_ESTROGEN\_RESPONSE\_LATE      
 Profile of the Running ES Score & Positions of GeneSet Members on the Rank Ordered List

  

| SYMBOL | TITLE | RANK IN GENE LIST | RANK METRIC SCORE | RUNNING ES | CORE ENRICHMENT || 1 | CA2 | carbonic anhydrase II | 69 | 0.250 | 0.0185 | Yes |
| 2 | KLK10 | kallikrein related peptidase 10 | 99 | 0.209 | 0.0355 | Yes |
| 3 | NMU | neuromedin U | 105 | 0.206 | 0.0535 | Yes |
| 4 | WFS1 | Wolfram syndrome 1 (wolframin) | 150 | 0.172 | 0.0664 | Yes |
| 5 | KRT19 | keratin 19, type I | 157 | 0.170 | 0.0811 | Yes |
| 6 | GINS2 | GINS complex subunit 2 (Psf2 homolog) | 190 | 0.158 | 0.0935 | Yes |
| 7 | MYB | v-myb avian myeloblastosis viral oncogene homolog | 306 | 0.133 | 0.0993 | Yes |
| 8 | HPRT1 | hypoxanthine phosphoribosyltransferase 1 | 311 | 0.132 | 0.1108 | Yes |
| 9 | ABCA3 | ATP binding cassette subfamily A member 3 | 321 | 0.129 | 0.1217 | Yes |
| 10 | TOB1 | transducer of ERBB2, 1 | 343 | 0.127 | 0.1318 | Yes |
| 11 | SLC29A1 | solute carrier family 29 (equilibrative nucleoside transporter), member 1 | 410 | 0.118 | 0.1389 | Yes |
| 12 | BTG3 | BTG family, member 3 | 465 | 0.113 | 0.1461 | Yes |
| 13 | CLIC3 | chloride intracellular channel 3 | 510 | 0.109 | 0.1535 | Yes |
| 14 | KCNK5 | potassium channel, two pore domain subfamily K, member 5 | 550 | 0.106 | 0.1608 | Yes |
| 15 | TFAP2C | transcription factor AP-2 gamma (activating enhancer binding protein 2 gamma) | 567 | 0.105 | 0.1693 | Yes |
| 16 | CD44 | CD44 molecule (Indian blood group) | 577 | 0.104 | 0.1780 | Yes |
| 17 | GAL | galanin/GMAP prepropeptide | 581 | 0.104 | 0.1871 | Yes |
| 18 | RNASEH2A | ribonuclease H2, subunit A | 611 | 0.103 | 0.1947 | Yes |
| 19 | IGFBP4 | insulin like growth factor binding protein 4 | 792 | 0.093 | 0.1936 | Yes |
| 20 | FDFT1 | farnesyl-diphosphate farnesyltransferase 1 | 793 | 0.093 | 0.2018 | Yes |
| 21 | TNNC1 | troponin C type 1 (slow) | 953 | 0.086 | 0.2012 | Yes |
| 22 | FKBP4 | FK506 binding protein 4 | 973 | 0.085 | 0.2077 | Yes |
| 23 | CDC20 | cell division cycle 20 | 1022 | 0.084 | 0.2126 | Yes |
| 24 | HSPA4L | heat shock 70kDa protein 4-like | 1070 | 0.082 | 0.2174 | Yes |
| 25 | CHPT1 | choline phosphotransferase 1 | 1092 | 0.081 | 0.2235 | Yes |
| 26 | ID2 | inhibitor of DNA binding 2, dominant negative helix-loop-helix protein | 1150 | 0.080 | 0.2276 | Yes |
| 27 | FOXC1 | forkhead box C1 | 1152 | 0.080 | 0.2346 | Yes |
| 28 | BAG1 | BCL2-associated athanogene | 1176 | 0.079 | 0.2404 | Yes |
| 29 | MYOF | myoferlin | 1329 | 0.076 | 0.2392 | Yes |
| 30 | BATF | basic leucine zipper transcription factor, ATF-like | 1397 | 0.074 | 0.2423 | Yes |
| 31 | FABP5 | fatty acid binding protein 5 (psoriasis-associated) | 1511 | 0.071 | 0.2428 | Yes |
| 32 | METTL3 | methyltransferase like 3 | 1524 | 0.071 | 0.2484 | Yes |
| 33 | UNC13B | unc-13 homolog B (C. elegans) | 1596 | 0.069 | 0.2508 | Yes |
| 34 | ANXA9 | annexin A9 | 1673 | 0.067 | 0.2529 | Yes |
| 35 | ACOX2 | acyl-CoA oxidase 2, branched chain | 1687 | 0.067 | 0.2581 | Yes |
| 36 | FKBP5 | FK506 binding protein 5 | 1840 | 0.064 | 0.2559 | Yes |
| 37 | RAPGEFL1 | Rap guanine nucleotide exchange factor like 1 | 1872 | 0.064 | 0.2599 | Yes |
| 38 | DHCR7 | 7-dehydrocholesterol reductase | 1877 | 0.064 | 0.2653 | Yes |
| 39 | FLNB | filamin B, beta | 1902 | 0.063 | 0.2697 | Yes |
| 40 | NXT1 | nuclear transport factor 2-like export factor 1 | 1914 | 0.063 | 0.2747 | Yes |
| 41 | FGFR3 | fibroblast growth factor receptor 3 | 1953 | 0.062 | 0.2782 | Yes |
| 42 | PTGES | prostaglandin E synthase | 1996 | 0.061 | 0.2815 | Yes |
| 43 | SFN | stratifin | 2005 | 0.061 | 0.2865 | Yes |
| 44 | GLA | galactosidase, alpha | 2101 | 0.060 | 0.2869 | Yes |
| 45 | CDC6 | cell division cycle 6 | 2205 | 0.058 | 0.2867 | Yes |
| 46 | KRT13 | keratin 13, type I | 2215 | 0.058 | 0.2914 | Yes |
| 47 | DUSP2 | dual specificity phosphatase 2 | 2355 | 0.056 | 0.2891 | Yes |
| 48 | TIAM1 | T-cell lymphoma invasion and metastasis 1 | 2375 | 0.056 | 0.2931 | Yes |
| 49 | CA12 | carbonic anhydrase XII | 2439 | 0.055 | 0.2946 | Yes |
| 50 | COX6C | cytochrome c oxidase subunit VIc | 2491 | 0.054 | 0.2968 | Yes |
| 51 | SORD | sorbitol dehydrogenase | 2500 | 0.054 | 0.3011 | Yes |
| 52 | IL17RB | interleukin 17 receptor B | 2765 | 0.050 | 0.2919 | No |
| 53 | IMPA2 | inositol(myo)-1(or 4)-monophosphatase 2 | 2809 | 0.050 | 0.2941 | No |
| 54 | ATP2B4 | ATPase, Ca++ transporting, plasma membrane 4 | 2874 | 0.049 | 0.2951 | No |
| 55 | SLC16A1 | solute carrier family 16 (monocarboxylate transporter), member 1 | 2936 | 0.048 | 0.2962 | No |
| 56 | ST6GALNAC2 | ST6 (alpha-N-acetyl-neuraminyl-2,3-beta-galactosyl-1,3)-N-acetylgalactosaminide alpha-2,6-sialyltransferase 2 | 3028 | 0.047 | 0.2957 | No |
| 57 | EMP2 | epithelial membrane protein 2 | 3062 | 0.047 | 0.2981 | No |
| 58 | GJB3 | gap junction protein beta 3 | 3144 | 0.046 | 0.2980 | No |
| 59 | SLC9A3R1 | solute carrier family 9, subfamily A (NHE3, cation proton antiporter 3), member 3 regulator 1 | 3344 | 0.044 | 0.2916 | No |
| 60 | SERPINA5 | serpin peptidase inhibitor, clade A (alpha-1 antiproteinase, antitrypsin), member 5 | 3551 | 0.042 | 0.2847 | No |
| 61 | S100A9 | S100 calcium binding protein A9 | 3554 | 0.042 | 0.2883 | No |
| 62 | TRIM29 | tripartite motif containing 29 | 3749 | 0.040 | 0.2818 | No |
| 63 | PDCD4 | programmed cell death 4 (neoplastic transformation inhibitor) | 3801 | 0.040 | 0.2827 | No |
| 64 | ETFB | electron-transfer-flavoprotein, beta polypeptide | 3934 | 0.038 | 0.2792 | No |
| 65 | TH | tyrosine hydroxylase | 3966 | 0.038 | 0.2810 | No |
| 66 | OVOL2 | ovo-like zinc finger 2 | 3978 | 0.038 | 0.2837 | No |
| 67 | CPE | carboxypeptidase E | 4003 | 0.038 | 0.2858 | No |
| 68 | NRIP1 | nuclear receptor interacting protein 1 | 4528 | 0.033 | 0.2616 | No |
| 69 | CCND1 | cyclin D1 | 4585 | 0.032 | 0.2616 | No |
| 70 | TPD52L1 | tumor protein D52-like 1 | 4623 | 0.032 | 0.2625 | No |
| 71 | ABHD2 | abhydrolase domain containing 2 | 4774 | 0.031 | 0.2575 | No |
| 72 | RET | ret proto-oncogene | 4931 | 0.030 | 0.2521 | No |
| 73 | DCXR | dicarbonyl/L-xylulose reductase | 5337 | 0.027 | 0.2335 | No |
| 74 | PLK4 | polo-like kinase 4 | 5691 | 0.024 | 0.2173 | No |
| 75 | ELOVL5 | ELOVL fatty acid elongase 5 | 5798 | 0.023 | 0.2139 | No |
| 76 | BLVRB | biliverdin reductase B | 5918 | 0.023 | 0.2098 | No |
| 77 | CKB | creatine kinase, brain | 5948 | 0.022 | 0.2102 | No |
| 78 | RAB31 | Transcript Identified by AceView, Entrez Gene ID(s) 11031 | 6174 | 0.021 | 0.2004 | No |
| 79 | IDH2 | isocitrate dehydrogenase 2 (NADP+), mitochondrial | 6235 | 0.021 | 0.1992 | No |
| 80 | SGK1 | serum/glucocorticoid regulated kinase 1 | 6290 | 0.020 | 0.1982 | No |
| 81 | OLFM1 | olfactomedin 1 | 6358 | 0.020 | 0.1965 | No |
| 82 | XRCC3 | X-ray repair complementing defective repair in Chinese hamster cells 3 | 6397 | 0.020 | 0.1962 | No |
| 83 | NAB2 | NGFI-A binding protein 2 (EGR1 binding protein 2) | 6638 | 0.018 | 0.1854 | No |
| 84 | SLC27A2 | solute carrier family 27 (fatty acid transporter), member 2 | 6687 | 0.018 | 0.1845 | No |
| 85 | MAPT | microtubule associated protein tau | 6748 | 0.017 | 0.1830 | No |
| 86 | TMPRSS3 | transmembrane protease, serine 3 | 6794 | 0.017 | 0.1821 | No |
| 87 | LLGL2 | lethal giant larvae homolog 2 (Drosophila) | 6901 | 0.016 | 0.1781 | No |
| 88 | CD9 | CD9 molecule | 6905 | 0.016 | 0.1794 | No |
| 89 | TOP2A | topoisomerase (DNA) II alpha | 6917 | 0.016 | 0.1803 | No |
| 90 | CDH1 | cadherin 1, type 1 | 7180 | 0.015 | 0.1680 | No |
| 91 | PPIF | peptidylprolyl isomerase F | 7224 | 0.015 | 0.1671 | No |
| 92 | CXCL12 | chemokine (C-X-C motif) ligand 12 | 7452 | 0.013 | 0.1565 | No |
| 93 | UGDH | UDP-glucose 6-dehydrogenase | 7487 | 0.013 | 0.1559 | No |
| 94 | GALE | UDP-galactose-4-epimerase | 7614 | 0.012 | 0.1505 | No |
| 95 | MOCS2 | molybdenum cofactor synthesis 2 | 7659 | 0.012 | 0.1492 | No |
| 96 | STIL | SCL/TAL1 interrupting locus | 7923 | 0.010 | 0.1366 | No |
| 97 | CALCR | calcitonin receptor | 7944 | 0.010 | 0.1364 | No |
| 98 | PKP3 | plakophilin 3 | 7950 | 0.010 | 0.1371 | No |
| 99 | SLC22A5 | solute carrier family 22 (organic cation/carnitine transporter), member 5 | 8098 | 0.009 | 0.1303 | No |
| 100 | HMGCS2 | 3-hydroxy-3-methylglutaryl-CoA synthase 2 (mitochondrial) | 8539 | 0.007 | 0.1081 | No |
| 101 | MICB | MHC class I polypeptide-related sequence B | 8905 | 0.005 | 0.0896 | No |
| 102 | SCARB1 | scavenger receptor class B, member 1 | 8987 | 0.004 | 0.0858 | No |
| 103 | CHST8 | carbohydrate (N-acetylgalactosamine 4-0) sulfotransferase 8 | 9095 | 0.003 | 0.0806 | No |
| 104 | CACNA2D2 | calcium channel, voltage-dependent, alpha 2/delta subunit 2 | 9169 | 0.003 | 0.0771 | No |
| 105 | LSR | lipolysis stimulated lipoprotein receptor | 9320 | 0.002 | 0.0695 | No |
| 106 | OPN3 | opsin 3 | 9365 | 0.002 | 0.0674 | No |
| 107 | DYNLT3 | dynein, light chain, Tctex-type 3 | 9497 | 0.001 | 0.0608 | No |
| 108 | PLXNB1 | plexin B1 | 9668 | 0.001 | 0.0520 | No |
| 109 | PTGER3 | prostaglandin E receptor 3 (subtype EP3) | 9995 | -0.001 | 0.0353 | No |
| 110 | LTF | lactotransferrin | 10005 | -0.001 | 0.0349 | No |
| 111 | DLG5 | discs, large homolog 5 (Drosophila) | 10202 | -0.002 | 0.0250 | No |
| 112 | KIF20A | kinesin family member 20A | 10416 | -0.004 | 0.0143 | No |
| 113 | ALDH3A2 | aldehyde dehydrogenase 3 family, member A2 | 10449 | -0.004 | 0.0130 | No |
| 114 | DNAJC1 | DnaJ (Hsp40) homolog, subfamily C, member 1 | 10464 | -0.004 | 0.0126 | No |
| 115 | TSTA3 | tissue specific transplantation antigen P35B | 10774 | -0.006 | -0.0029 | No |
| 116 | SNX10 | sorting nexin 10 | 11020 | -0.007 | -0.0149 | No |
| 117 | RABEP1 | rabaptin, RAB GTPase binding effector protein 1 | 11117 | -0.008 | -0.0192 | No |
| 118 | MAPK13 | mitogen-activated protein kinase 13 | 11298 | -0.009 | -0.0277 | No |
| 119 | SCNN1A | sodium channel, non voltage gated 1 alpha subunit | 11540 | -0.010 | -0.0393 | No |
| 120 | TFF1 | trefoil factor 1 | 11711 | -0.011 | -0.0471 | No |
| 121 | CELSR2 | cadherin, EGF LAG seven-pass G-type receptor 2 | 12052 | -0.013 | -0.0636 | No |
| 122 | TFF3 | trefoil factor 3 | 12114 | -0.014 | -0.0655 | No |
| 123 | NPY1R | neuropeptide Y receptor Y1 | 12162 | -0.014 | -0.0667 | No |
| 124 | NCOR2 | nuclear receptor corepressor 2 | 12339 | -0.015 | -0.0745 | No |
| 125 | PRKAR2B | protein kinase, cAMP-dependent, regulatory, type II, beta | 12568 | -0.017 | -0.0848 | No |
| 126 | PRSS23 | protease, serine, 23 | 12809 | -0.018 | -0.0957 | No |
| 127 | JAK2 | Janus kinase 2 | 13031 | -0.019 | -0.1054 | No |
| 128 | ASS1 | argininosuccinate synthase 1 | 13106 | -0.020 | -0.1074 | No |
| 129 | SIAH2 | siah E3 ubiquitin protein ligase 2 | 13372 | -0.022 | -0.1192 | No |
| 130 | PDLIM3 | PDZ and LIM domain 3 | 13450 | -0.022 | -0.1212 | No |
| 131 | NBL1 | neuroblastoma 1, DAN family BMP antagonist | 13497 | -0.023 | -0.1216 | No |
| 132 | SLC24A3 | solute carrier family 24 (sodium/potassium/calcium exchanger), member 3 | 13559 | -0.023 | -0.1227 | No |
| 133 | PTPN6 | protein tyrosine phosphatase, non-receptor type 6 | 13725 | -0.024 | -0.1291 | No |
| 134 | CCNA1 | cyclin A1 | 13813 | -0.025 | -0.1314 | No |
| 135 | FARP1 | FERM, ARH/RhoGEF and pleckstrin domain protein 1 | 14015 | -0.026 | -0.1395 | No |
| 136 | ST14 | suppression of tumorigenicity 14 (colon carcinoma) | 14096 | -0.027 | -0.1412 | No |
| 137 | PERP | PERP, TP53 apoptosis effector | 14176 | -0.028 | -0.1429 | No |
| 138 | SCUBE2 | signal peptide, CUB domain, EGF-like 2 | 14273 | -0.029 | -0.1453 | No |
| 139 | TFPI2 | tissue factor pathway inhibitor 2 | 14489 | -0.030 | -0.1538 | No |
| 140 | BCL2 | B-cell CLL/lymphoma 2 | 14503 | -0.030 | -0.1517 | No |
| 141 | SERPINA3 | serpin peptidase inhibitor, clade A (alpha-1 antiproteinase, antitrypsin), member 3 | 14750 | -0.032 | -0.1616 | No |
| 142 | ARL3 | ADP-ribosylation factor like GTPase 3 | 14799 | -0.033 | -0.1612 | No |
| 143 | EGR3 | early growth response 3 | 14802 | -0.033 | -0.1584 | No |
| 144 | CYP26B1 | cytochrome P450, family 26, subfamily B, polypeptide 1 | 14910 | -0.034 | -0.1609 | No |
| 145 | TJP3 | tight junction protein 3 | 15002 | -0.034 | -0.1626 | No |
| 146 | FRK | fyn-related Src family tyrosine kinase | 15076 | -0.035 | -0.1633 | No |
| 147 | SOX3 | SRY box 3 | 15166 | -0.036 | -0.1647 | No |
| 148 | PRLR | prolactin receptor | 15454 | -0.038 | -0.1762 | No |
| 149 | XBP1 | X-box binding protein 1 | 15622 | -0.040 | -0.1813 | No |
| 150 | TST | thiosulfate sulfurtransferase (rhodanese) | 15640 | -0.040 | -0.1786 | No |
| 151 | CISH | cytokine inducible SH2-containing protein | 15667 | -0.040 | -0.1764 | No |
| 152 | AFF1 | AF4/FMR2 family, member 1 | 15691 | -0.041 | -0.1739 | No |
| 153 | SLC26A2 | solute carrier family 26 (anion exchanger), member 2 | 15913 | -0.043 | -0.1816 | No |
| 154 | LAMC2 | laminin, gamma 2 | 15989 | -0.044 | -0.1816 | No |
| 155 | CAV1 | caveolin 1 | 16000 | -0.044 | -0.1783 | No |
| 156 | ADD3 | adducin 3 (gamma) | 16088 | -0.045 | -0.1788 | No |
| 157 | ISG20 | interferon stimulated exonuclease gene 20kDa | 16332 | -0.047 | -0.1872 | No |
| 158 | AMFR | autocrine motility factor receptor, E3 ubiquitin protein ligase | 16444 | -0.049 | -0.1886 | No |
| 159 | ITPK1 | inositol-tetrakisphosphate 1-kinase | 16518 | -0.050 | -0.1879 | No |
| 160 | AREG | amphiregulin | 16601 | -0.051 | -0.1876 | No |
| 161 | RBBP8 | retinoblastoma binding protein 8 | 16689 | -0.052 | -0.1875 | No |
| 162 | AGR2 | anterior gradient 2, protein disulphide isomerase family member | 16744 | -0.053 | -0.1856 | No |
| 163 | TPBG | trophoblast glycoprotein | 17065 | -0.058 | -0.1970 | No |
| 164 | TSPAN13 | tetraspanin 13 | 17370 | -0.063 | -0.2072 | No |
| 165 | JAK1 | Janus kinase 1 | 17394 | -0.063 | -0.2028 | No |
| 166 | TPSAB1 | tryptase alpha/beta 1 | 17445 | -0.064 | -0.1997 | No |
| 167 | KLK11 | kallikrein related peptidase 11 | 17478 | -0.065 | -0.1956 | No |
| 168 | SULT2B1 | sulfotransferase family 2B member 1 | 17511 | -0.066 | -0.1915 | No |
| 169 | FOS | FBJ murine osteosarcoma viral oncogene homolog | 17528 | -0.066 | -0.1865 | No |
| 170 | SEMA3B | sema domain, immunoglobulin domain (Ig), short basic domain, secreted, (semaphorin) 3B | 17591 | -0.067 | -0.1837 | No |
| 171 | FAM102A | Memczak2013 ANTISENSE, CDS, coding, INTERNAL, intronic, UTR3 best transcript NM\_203305 | 17638 | -0.068 | -0.1801 | No |
| 172 | CYP4F11 | cytochrome P450, family 4, subfamily F, polypeptide 11 | 17745 | -0.071 | -0.1793 | No |
| 173 | SLC7A5 | solute carrier family 7 (amino acid transporter light chain, L system), member 5 | 17849 | -0.073 | -0.1782 | No |
| 174 | GPER1 | G protein-coupled estrogen receptor 1 | 17930 | -0.075 | -0.1757 | No |
| 175 | ZFP36 | ZFP36 ring finger protein | 17953 | -0.076 | -0.1701 | No |
| 176 | ALDH3B1 | aldehyde dehydrogenase 3 family, member B1 | 17969 | -0.076 | -0.1641 | No |
| 177 | KLF4 | Kruppel-like factor 4 (gut) | 17974 | -0.076 | -0.1576 | No |
| 178 | CXCL14 | chemokine (C-X-C motif) ligand 14 | 18090 | -0.079 | -0.1565 | No |
| 179 | PAPSS2 | 3-phosphoadenosine 5-phosphosulfate synthase 2 | 18188 | -0.082 | -0.1542 | No |
| 180 | SLC2A8 | solute carrier family 2 (facilitated glucose transporter), member 8 | 18306 | -0.086 | -0.1527 | No |
| 181 | PGR | progesterone receptor | 18358 | -0.087 | -0.1476 | No |
| 182 | PDZK1 | PDZ domain containing 1 | 18426 | -0.090 | -0.1432 | No |
| 183 | HR | hair growth associated | 18647 | -0.099 | -0.1458 | No |
| 184 | IGSF1 | immunoglobulin superfamily, member 1 | 18652 | -0.099 | -0.1372 | No |
| 185 | SLC1A4 | solute carrier family 1 (glutamate/neutral amino acid transporter), member 4 | 18809 | -0.110 | -0.1355 | No |
| 186 | RPS6KA2 | ribosomal protein S6 kinase, 90kDa, polypeptide 2 | 18861 | -0.114 | -0.1280 | No |
| 187 | MEST | mesoderm specific transcript | 18998 | -0.127 | -0.1239 | No |
| 188 | PLAC1 | placenta specific 1 | 19002 | -0.127 | -0.1127 | No |
| 189 | ASCL1 | achaete-scute family bHLH transcription factor 1 | 19116 | -0.144 | -0.1058 | No |
| 190 | PCP4 | Purkinje cell protein 4 | 19121 | -0.145 | -0.0932 | No |
| 191 | IL6ST | interleukin 6 signal transducer | 19134 | -0.147 | -0.0808 | No |
| 192 | HSPB8 | heat shock 22kDa protein 8 | 19194 | -0.160 | -0.0697 | No |
| 193 | DNAJC12 | DnaJ (Hsp40) homolog, subfamily C, member 12 | 19268 | -0.180 | -0.0576 | No |
| 194 | SERPINA1 | serpin peptidase inhibitor, clade A (alpha-1 antiproteinase, antitrypsin), member 1 | 19288 | -0.189 | -0.0418 | No |
| 195 | MDK | midkine (neurite growth-promoting factor 2) | 19304 | -0.194 | -0.0254 | No |
| 196 | HOMER2 | homer scaffolding protein 2 | 19317 | -0.197 | -0.0086 | No |
| 197 | DHRS2 | dehydrogenase/reductase (SDR family) member 2 | 19363 | -0.218 | 0.0083 | No |
Table: GSEA details [plain text format]

  

Fig 2: HALLMARK\_ESTROGEN\_RESPONSE\_LATE      
 Blue-Pink O' Gram in the Space of the Analyzed GeneSet

  

Fig 3: HALLMARK\_ESTROGEN\_RESPONSE\_LATE: Random ES distribution      
 Gene set null distribution of ES for **HALLMARK\_ESTROGEN\_RESPONSE\_LATE**

  
