## Supplementary material for "Somatic hypomethylation of pericentromeric SST1 repeats and tetraploidization in human colorectal cancer cells": GSEA results: HALLMARK_FATTY_ACID_METABOLISM.html

Details for gene set HALLMARK\_FATTY\_ACID\_METABOLISM[GSEA]

|  || Dataset | eset\_byprobe\_collapsed\_to\_symbols.Diploid\_vs\_Tetraploid.cls #Tetraploid\_versus\_Diploid.Diploid\_vs\_Tetraploid.cls #Tetraploid\_versus\_Diploid\_repos |
| Phenotype | Diploid\_vs\_Tetraploid.cls#Tetraploid\_versus\_Diploid\_repos |
| Upregulated in class | Tetraploid |
| GeneSet | HALLMARK\_FATTY\_ACID\_METABOLISM |
| Enrichment Score (ES) | 0.28012568 |
| Normalized Enrichment Score (NES) | 1.2138832 |
| Nominal p-value | 0.08238637 |
| FDR q-value | 0.20998858 |
| FWER p-Value | 0.876 |
Table: GSEA Results Summary

  

Fig 1: Enrichment plot: HALLMARK\_FATTY\_ACID\_METABOLISM      
 Profile of the Running ES Score & Positions of GeneSet Members on the Rank Ordered List

  

| SYMBOL | TITLE | RANK IN GENE LIST | RANK METRIC SCORE | RUNNING ES | CORE ENRICHMENT || 1 | HPGD | hydroxyprostaglandin dehydrogenase 15-(NAD) | 62 | 0.267 | 0.0328 | Yes |
| 2 | CA2 | carbonic anhydrase II | 69 | 0.250 | 0.0662 | Yes |
| 3 | FABP1 | fatty acid binding protein 1, liver | 75 | 0.234 | 0.0975 | Yes |
| 4 | ACAT2 | acetyl-CoA acetyltransferase 2 | 133 | 0.185 | 0.1195 | Yes |
| 5 | FASN | fatty acid synthase | 293 | 0.135 | 0.1295 | Yes |
| 6 | HMGCS1 | 3-hydroxy-3-methylglutaryl-CoA synthase 1 (soluble) | 387 | 0.121 | 0.1410 | Yes |
| 7 | HSD17B7 | hydroxysteroid (17-beta) dehydrogenase 7 | 425 | 0.117 | 0.1548 | Yes |
| 8 | GPD2 | glycerol-3-phosphate dehydrogenase 2 | 534 | 0.107 | 0.1638 | Yes |
| 9 | ACAA2 | acetyl-CoA acyltransferase 2 | 744 | 0.095 | 0.1658 | Yes |
| 10 | CPOX | coproporphyrinogen oxidase | 767 | 0.094 | 0.1773 | Yes |
| 11 | S100A10 | S100 calcium binding protein A10 | 886 | 0.089 | 0.1832 | Yes |
| 12 | METAP1 | methionyl aminopeptidase 1 | 1009 | 0.084 | 0.1882 | Yes |
| 13 | GCDH | glutaryl-CoA dehydrogenase | 1085 | 0.081 | 0.1953 | Yes |
| 14 | ACADS | acyl-CoA dehydrogenase, C-2 to C-3 short chain | 1091 | 0.081 | 0.2060 | Yes |
| 15 | NSDHL | NAD(P) dependent steroid dehydrogenase-like | 1100 | 0.081 | 0.2165 | Yes |
| 16 | TDO2 | tryptophan 2,3-dioxygenase | 1202 | 0.079 | 0.2219 | Yes |
| 17 | ACSL4 | acyl-CoA synthetase long-chain family member 4 | 1389 | 0.074 | 0.2223 | Yes |
| 18 | NTHL1 | nth-like DNA glycosylase 1 | 1546 | 0.070 | 0.2238 | Yes |
| 19 | BPHL | biphenyl hydrolase-like (serine hydrolase) | 1755 | 0.066 | 0.2219 | Yes |
| 20 | HSDL2 | hydroxysteroid dehydrogenase like 2 | 1869 | 0.064 | 0.2247 | Yes |
| 21 | IDI1 | isopentenyl-diphosphate delta isomerase 1 | 1905 | 0.063 | 0.2314 | Yes |
| 22 | HSPH1 | heat shock 105kDa/110kDa protein 1 | 1948 | 0.062 | 0.2376 | Yes |
| 23 | CA6 | carbonic anhydrase VI | 1971 | 0.062 | 0.2448 | Yes |
| 24 | HADH | hydroxyacyl-CoA dehydrogenase | 2078 | 0.060 | 0.2475 | Yes |
| 25 | SUCLG2 | succinate-CoA ligase, GDP-forming, beta subunit | 2153 | 0.059 | 0.2516 | Yes |
| 26 | GPD1 | glycerol-3-phosphate dehydrogenase 1 | 2415 | 0.055 | 0.2456 | Yes |
| 27 | KMT5A | lysine (K)-specific methyltransferase 5A | 2552 | 0.053 | 0.2457 | Yes |
| 28 | ACSL5 | acyl-CoA synthetase long-chain family member 5 | 2576 | 0.053 | 0.2516 | Yes |
| 29 | DHCR24 | 24-dehydrocholesterol reductase | 2597 | 0.052 | 0.2576 | Yes |
| 30 | HSP90AA1 | heat shock protein 90kDa alpha (cytosolic), class A member 1 | 2892 | 0.049 | 0.2490 | Yes |
| 31 | NCAPH2 | non-SMC condensin II complex subunit H2 | 2920 | 0.048 | 0.2542 | Yes |
| 32 | REEP6 | receptor accessory protein 6 | 2923 | 0.048 | 0.2606 | Yes |
| 33 | ENO2 | enolase 2 (gamma, neuronal) | 3086 | 0.047 | 0.2585 | Yes |
| 34 | AADAT | aminoadipate aminotransferase | 3112 | 0.046 | 0.2635 | Yes |
| 35 | HIBCH | 3-hydroxyisobutyryl-CoA hydrolase | 3260 | 0.045 | 0.2620 | Yes |
| 36 | FH | fumarate hydratase | 3269 | 0.045 | 0.2677 | Yes |
| 37 | PRDX6 | peroxiredoxin 6 | 3370 | 0.044 | 0.2684 | Yes |
| 38 | ECI2 | enoyl-CoA delta isomerase 2 | 3498 | 0.043 | 0.2676 | Yes |
| 39 | ALDH1A1 | aldehyde dehydrogenase 1 family, member A1 | 3520 | 0.042 | 0.2722 | Yes |
| 40 | PDHA1 | pyruvate dehydrogenase (lipoamide) alpha 1 | 3617 | 0.041 | 0.2729 | Yes |
| 41 | PDHB | pyruvate dehydrogenase (lipoamide) beta | 3715 | 0.040 | 0.2733 | Yes |
| 42 | CA4 | carbonic anhydrase IV | 3876 | 0.039 | 0.2703 | Yes |
| 43 | IDH3B | isocitrate dehydrogenase 3 (NAD+) beta | 3915 | 0.038 | 0.2735 | Yes |
| 44 | DECR1 | 2,4-dienoyl-CoA reductase 1, mitochondrial | 3939 | 0.038 | 0.2775 | Yes |
| 45 | MLYCD | malonyl-CoA decarboxylase | 4013 | 0.037 | 0.2788 | Yes |
| 46 | DLST | dihydrolipoamide S-succinyltransferase (E2 component of 2-oxo-glutarate complex) | 4084 | 0.037 | 0.2801 | Yes |
| 47 | SDHA | succinate dehydrogenase complex subunit A, flavoprotein (Fp) | 4313 | 0.035 | 0.2730 | No |
| 48 | APEX1 | APEX nuclease (multifunctional DNA repair enzyme) 1 | 4609 | 0.032 | 0.2622 | No |
| 49 | PSME1 | proteasome activator subunit 1 | 4737 | 0.031 | 0.2598 | No |
| 50 | MIF | macrophage migration inhibitory factor (glycosylation-inhibiting factor) | 4925 | 0.030 | 0.2542 | No |
| 51 | MCEE | methylmalonyl CoA epimerase | 5028 | 0.029 | 0.2529 | No |
| 52 | BCKDHB | branched chain keto acid dehydrogenase E1, beta polypeptide | 5042 | 0.029 | 0.2561 | No |
| 53 | PTPRG | protein tyrosine phosphatase, receptor type, G | 5171 | 0.028 | 0.2533 | No |
| 54 | CRYZ | crystallin zeta | 5181 | 0.028 | 0.2566 | No |
| 55 | GLUL | glutamate-ammonia ligase | 5189 | 0.028 | 0.2600 | No |
| 56 | ALDH9A1 | aldehyde dehydrogenase 9 family, member A1 | 5592 | 0.025 | 0.2426 | No |
| 57 | YWHAH | tyrosine 3-monooxygenase/tryptophan 5-monooxygenase activation protein, eta | 5640 | 0.024 | 0.2434 | No |
| 58 | MDH1 | malate dehydrogenase 1 | 5641 | 0.024 | 0.2467 | No |
| 59 | ELOVL5 | ELOVL fatty acid elongase 5 | 5798 | 0.023 | 0.2419 | No |
| 60 | CYP4A11 | cytochrome P450, family 4, subfamily A, polypeptide 11 | 5813 | 0.023 | 0.2443 | No |
| 61 | LDHA | lactate dehydrogenase A | 6019 | 0.022 | 0.2366 | No |
| 62 | ADSL | adenylosuccinate lyase | 6023 | 0.022 | 0.2394 | No |
| 63 | SUCLA2 | succinate-CoA ligase, ADP-forming, beta subunit | 6053 | 0.022 | 0.2409 | No |
| 64 | PCBD1 | pterin-4 alpha-carbinolamine dehydratase/dimerization cofactor of hepatocyte nuclear factor 1 alpha | 6150 | 0.021 | 0.2388 | No |
| 65 | CCDC58 | coiled-coil domain containing 58 | 6163 | 0.021 | 0.2410 | No |
| 66 | BLVRA | biliverdin reductase A | 6229 | 0.021 | 0.2404 | No |
| 67 | SMS | spermine synthase | 6236 | 0.021 | 0.2429 | No |
| 68 | SDHD | succinate dehydrogenase complex subunit D, integral membrane protein | 6287 | 0.020 | 0.2430 | No |
| 69 | ALDOA | aldolase A, fructose-bisphosphate | 6361 | 0.020 | 0.2420 | No |
| 70 | ECH1 | enoyl-CoA hydratase 1, peroxisomal | 6583 | 0.018 | 0.2330 | No |
| 71 | SUCLG1 | succinate-CoA ligase, alpha subunit | 6705 | 0.018 | 0.2292 | No |
| 72 | ACAA1 | acetyl-CoA acyltransferase 1 | 6834 | 0.017 | 0.2248 | No |
| 73 | ECHS1 | enoyl-CoA hydratase, short chain, 1, mitochondrial | 6902 | 0.016 | 0.2236 | No |
| 74 | CYP1A1 | cytochrome P450, family 1, subfamily A, polypeptide 1 | 6941 | 0.016 | 0.2238 | No |
| 75 | UROD | uroporphyrinogen decarboxylase | 7213 | 0.015 | 0.2118 | No |
| 76 | SDHC | succinate dehydrogenase complex, subunit C, integral membrane protein, 15kDa | 7238 | 0.015 | 0.2126 | No |
| 77 | HSD17B11 | hydroxysteroid (17-beta) dehydrogenase 11 | 7455 | 0.013 | 0.2032 | No |
| 78 | UGDH | UDP-glucose 6-dehydrogenase | 7487 | 0.013 | 0.2033 | No |
| 79 | HADHB | hydroxyacyl-CoA dehydrogenase/3-ketoacyl-CoA thiolase/enoyl-CoA hydratase (trifunctional protein), beta subunit | 7561 | 0.013 | 0.2012 | No |
| 80 | UROS | uroporphyrinogen III synthase | 7623 | 0.012 | 0.1997 | No |
| 81 | GAPDHS | glyceraldehyde-3-phosphate dehydrogenase, spermatogenic | 7644 | 0.012 | 0.2003 | No |
| 82 | ACOX1 | acyl-CoA oxidase 1, palmitoyl | 7694 | 0.012 | 0.1994 | No |
| 83 | SLC22A5 | solute carrier family 22 (organic cation/carnitine transporter), member 5 | 8098 | 0.009 | 0.1798 | No |
| 84 | HCCS | holocytochrome c synthase | 8387 | 0.008 | 0.1660 | No |
| 85 | GRHPR | glyoxylate reductase/hydroxypyruvate reductase | 8421 | 0.007 | 0.1653 | No |
| 86 | GAD2 | glutamate decarboxylase 2 | 8511 | 0.007 | 0.1616 | No |
| 87 | HMGCS2 | 3-hydroxy-3-methylglutaryl-CoA synthase 2 (mitochondrial) | 8539 | 0.007 | 0.1611 | No |
| 88 | ADIPOR2 | adiponectin receptor 2 | 8746 | 0.005 | 0.1512 | No |
| 89 | D2HGDH | D-2-hydroxyglutarate dehydrogenase | 9143 | 0.003 | 0.1312 | No |
| 90 | ACADM | acyl-CoA dehydrogenase, C-4 to C-12 straight chain | 9153 | 0.003 | 0.1312 | No |
| 91 | ECI1 | enoyl-CoA delta isomerase 1 | 9234 | 0.003 | 0.1274 | No |
| 92 | ACSL1 | acyl-CoA synthetase long-chain family member 1 | 9852 | -0.000 | 0.0956 | No |
| 93 | CD1D | CD1d molecule | 9911 | -0.001 | 0.0927 | No |
| 94 | INMT | indolethylamine N-methyltransferase | 10043 | -0.001 | 0.0861 | No |
| 95 | AUH | AU RNA binding protein/enoyl-CoA hydratase | 10198 | -0.002 | 0.0785 | No |
| 96 | ENO3 | enolase 3 (beta, muscle) | 10237 | -0.003 | 0.0769 | No |
| 97 | ALDH3A2 | aldehyde dehydrogenase 3 family, member A2 | 10449 | -0.004 | 0.0665 | No |
| 98 | ACO2 | aconitase 2, mitochondrial | 10537 | -0.004 | 0.0626 | No |
| 99 | HMGCL | 3-hydroxymethyl-3-methylglutaryl-CoA lyase | 10543 | -0.004 | 0.0629 | No |
| 100 | EHHADH | enoyl-CoA, hydratase/3-hydroxyacyl CoA dehydrogenase | 10756 | -0.006 | 0.0527 | No |
| 101 | RDH11 | retinol dehydrogenase 11 (all-trans/9-cis/11-cis) | 11501 | -0.010 | 0.0157 | No |
| 102 | AQP7 | aquaporin 7 | 11648 | -0.011 | 0.0096 | No |
| 103 | CEL | carboxyl ester lipase | 11847 | -0.012 | 0.0009 | No |
| 104 | NBN | nibrin | 11962 | -0.013 | -0.0032 | No |
| 105 | ACADL | acyl-CoA dehydrogenase, long chain | 12422 | -0.016 | -0.0248 | No |
| 106 | ETFDH | electron-transferring-flavoprotein dehydrogenase | 12478 | -0.016 | -0.0255 | No |
| 107 | DLD | dihydrolipoamide dehydrogenase | 12560 | -0.016 | -0.0275 | No |
| 108 | CBR1 | carbonyl reductase 1 | 12637 | -0.017 | -0.0291 | No |
| 109 | HSD17B4 | hydroxysteroid (17-beta) dehydrogenase 4 | 12798 | -0.018 | -0.0350 | No |
| 110 | EPHX1 | epoxide hydrolase 1, microsomal (xenobiotic) | 12904 | -0.019 | -0.0379 | No |
| 111 | ODC1 | ornithine decarboxylase 1 | 13133 | -0.020 | -0.0469 | No |
| 112 | CD36 | CD36 molecule (thrombospondin receptor) | 13245 | -0.021 | -0.0499 | No |
| 113 | CPT2 | carnitine palmitoyltransferase 2 | 13285 | -0.021 | -0.0490 | No |
| 114 | IDH3G | isocitrate dehydrogenase 3 (NAD+) gamma | 13328 | -0.021 | -0.0483 | No |
| 115 | TP53INP2 | tumor protein p53 inducible nuclear protein 2 | 13342 | -0.022 | -0.0461 | No |
| 116 | OSTC | oligosaccharyltransferase complex subunit (non-catalytic) | 13492 | -0.023 | -0.0507 | No |
| 117 | PPARA | peroxisome proliferator-activated receptor alpha | 13832 | -0.025 | -0.0648 | No |
| 118 | HSD17B10 | hydroxysteroid (17-beta) dehydrogenase 10 | 13918 | -0.026 | -0.0657 | No |
| 119 | ACOT2 | acyl-CoA thioesterase 2 | 14037 | -0.027 | -0.0683 | No |
| 120 | ACADVL | acyl-CoA dehydrogenase, very long chain | 14065 | -0.027 | -0.0660 | No |
| 121 | PTS | 6-pyruvoyltetrahydropterin synthase | 14234 | -0.028 | -0.0709 | No |
| 122 | ERP29 | endoplasmic reticulum protein 29 | 14282 | -0.029 | -0.0695 | No |
| 123 | AOC3 | amine oxidase, copper containing 3 | 14402 | -0.030 | -0.0716 | No |
| 124 | LTC4S | leukotriene C4 synthase | 14464 | -0.030 | -0.0707 | No |
| 125 | MDH2 | malate dehydrogenase 2 | 14465 | -0.030 | -0.0667 | No |
| 126 | ADH1C | alcohol dehydrogenase 1C (class I), gamma polypeptide | 14631 | -0.031 | -0.0709 | No |
| 127 | ACSS1 | acyl-CoA synthetase short-chain family member 1 | 14659 | -0.032 | -0.0680 | No |
| 128 | CIDEA | cell death-inducing DFFA-like effector a | 14801 | -0.033 | -0.0709 | No |
| 129 | IDH1 | isocitrate dehydrogenase 1 (NADP+) | 14950 | -0.034 | -0.0739 | No |
| 130 | ACSM3 | acyl-CoA synthetase medium-chain family member 3 | 15283 | -0.037 | -0.0861 | No |
| 131 | ACOT8 | acyl-CoA thioesterase 8 | 15328 | -0.037 | -0.0834 | No |
| 132 | SERINC1 | serine incorporator 1 | 15340 | -0.037 | -0.0789 | No |
| 133 | MGLL | monoglyceride lipase | 15364 | -0.038 | -0.0750 | No |
| 134 | CRAT | carnitine O-acetyltransferase | 15464 | -0.038 | -0.0749 | No |
| 135 | GSTZ1 | glutathione S-transferase zeta 1 | 15549 | -0.039 | -0.0740 | No |
| 136 | RAP1GDS1 | RAP1, GTP-GDP dissociation stimulator 1 | 15647 | -0.040 | -0.0735 | No |
| 137 | FMO1 | flavin containing monooxygenase 1 | 16324 | -0.047 | -0.1020 | No |
| 138 | HAO2 | hydroxyacid oxidase 2 (long chain) | 16537 | -0.050 | -0.1062 | No |
| 139 | CPT1A | carnitine palmitoyltransferase 1A (liver) | 16763 | -0.053 | -0.1106 | No |
| 140 | ADH7 | alcohol dehydrogenase 7 (class IV), mu or sigma polypeptide | 16808 | -0.054 | -0.1056 | No |
| 141 | BMPR1B | bone morphogenetic protein receptor type IB | 16952 | -0.056 | -0.1054 | No |
| 142 | CYP4A22 | cytochrome P450, family 4, subfamily A, polypeptide 22 | 17166 | -0.059 | -0.1084 | No |
| 143 | VNN1 | vanin 1 | 17657 | -0.069 | -0.1244 | No |
| 144 | RETSAT | retinol saturase (all-trans-retinol 13,14-reductase) | 17683 | -0.069 | -0.1164 | No |
| 145 | ALAD | aminolevulinate dehydratase | 17796 | -0.072 | -0.1124 | No |
| 146 | ME1 | malic enzyme 1, NADP(+)-dependent, cytosolic | 18066 | -0.079 | -0.1157 | No |
| 147 | ALDH3A1 | aldehyde dehydrogenase 3 family, member A1 | 18146 | -0.081 | -0.1088 | No |
| 148 | UBE2L6 | ubiquitin-conjugating enzyme E2L 6 | 18293 | -0.085 | -0.1049 | No |
| 149 | G0S2 | G0/G1 switch 2 | 18709 | -0.103 | -0.1124 | No |
| 150 | RDH16 | retinol dehydrogenase 16 (all-trans) | 18885 | -0.117 | -0.1057 | No |
| 151 | LGALS1 | lectin, galactoside-binding, soluble, 1 | 18961 | -0.123 | -0.0930 | No |
| 152 | MAOA | monoamine oxidase A | 19140 | -0.148 | -0.0822 | No |
| 153 | CBR3 | carbonyl reductase 3 | 19241 | -0.172 | -0.0642 | No |
| 154 | GABARAPL1 | GABA(A) receptor-associated protein like 1 | 19244 | -0.172 | -0.0411 | No |
| 155 | FABP2 | fatty acid binding protein 2, intestinal | 19503 | -0.411 | 0.0011 | No |
Table: GSEA details [plain text format]

  

Fig 2: HALLMARK\_FATTY\_ACID\_METABOLISM      
 Blue-Pink O' Gram in the Space of the Analyzed GeneSet

  

Fig 3: HALLMARK\_FATTY\_ACID\_METABOLISM: Random ES distribution      
 Gene set null distribution of ES for **HALLMARK\_FATTY\_ACID\_METABOLISM**

  
