## Supplementary material for "Somatic hypomethylation of pericentromeric SST1 repeats and tetraploidization in human colorectal cancer cells": GSEA results: HALLMARK_G2M_CHECKPOINT.html

Details for gene set HALLMARK\_G2M\_CHECKPOINT[GSEA]

|  || Dataset | eset\_byprobe\_collapsed\_to\_symbols.Diploid\_vs\_Tetraploid.cls #Tetraploid\_versus\_Diploid.Diploid\_vs\_Tetraploid.cls #Tetraploid\_versus\_Diploid\_repos |
| Phenotype | Diploid\_vs\_Tetraploid.cls#Tetraploid\_versus\_Diploid\_repos |
| Upregulated in class | Tetraploid |
| GeneSet | HALLMARK\_G2M\_CHECKPOINT |
| Enrichment Score (ES) | 0.40832755 |
| Normalized Enrichment Score (NES) | 1.782756 |
| Nominal p-value | 0.0 |
| FDR q-value | 0.0020001838 |
| FWER p-Value | 0.005 |
Table: GSEA Results Summary

  

Fig 1: Enrichment plot: HALLMARK\_G2M\_CHECKPOINT      
 Profile of the Running ES Score & Positions of GeneSet Members on the Rank Ordered List

  

| SYMBOL | TITLE | RANK IN GENE LIST | RANK METRIC SCORE | RUNNING ES | CORE ENRICHMENT || 1 | ATRX | alpha thalassemia/mental retardation syndrome X-linked | 60 | 0.271 | 0.0296 | Yes |
| 2 | SQLE | squalene epoxidase | 167 | 0.166 | 0.0442 | Yes |
| 3 | GINS2 | GINS complex subunit 2 (Psf2 homolog) | 190 | 0.158 | 0.0621 | Yes |
| 4 | POLQ | polymerase (DNA directed), theta | 198 | 0.156 | 0.0806 | Yes |
| 5 | EXO1 | exonuclease 1 | 271 | 0.137 | 0.0934 | Yes |
| 6 | BRCA2 | breast cancer 2, early onset | 273 | 0.137 | 0.1098 | Yes |
| 7 | CDC25A | cell division cycle 25A | 295 | 0.134 | 0.1250 | Yes |
| 8 | RPS6KA5 | ribosomal protein S6 kinase, 90kDa, polypeptide 5 | 360 | 0.124 | 0.1366 | Yes |
| 9 | MYBL2 | v-myb avian myeloblastosis viral oncogene homolog-like 2 | 400 | 0.119 | 0.1490 | Yes |
| 10 | PDS5B | PDS5 cohesin associated factor B | 452 | 0.114 | 0.1601 | Yes |
| 11 | RAD54L | RAD54-like (S. cerevisiae) | 460 | 0.113 | 0.1734 | Yes |
| 12 | ORC6 | origin recognition complex subunit 6 | 480 | 0.112 | 0.1859 | Yes |
| 13 | CDC45 | cell division cycle 45 | 603 | 0.103 | 0.1920 | Yes |
| 14 | CASP8AP2 | caspase 8 associated protein 2 | 652 | 0.100 | 0.2016 | Yes |
| 15 | CENPA | centromere protein A | 673 | 0.098 | 0.2124 | Yes |
| 16 | PRIM2 | primase, DNA, polypeptide 2 (58kDa) | 728 | 0.096 | 0.2212 | Yes |
| 17 | KIF15 | kinesin family member 15 | 913 | 0.088 | 0.2222 | Yes |
| 18 | STMN1 | stathmin 1 | 951 | 0.086 | 0.2307 | Yes |
| 19 | CKS2 | CDC28 protein kinase regulatory subunit 2 | 1016 | 0.084 | 0.2375 | Yes |
| 20 | CDC20 | cell division cycle 20 | 1022 | 0.084 | 0.2473 | Yes |
| 21 | PBK | PDZ binding kinase | 1090 | 0.081 | 0.2536 | Yes |
| 22 | TTK | TTK protein kinase | 1229 | 0.078 | 0.2559 | Yes |
| 23 | KIF22 | kinesin family member 22 | 1286 | 0.077 | 0.2622 | Yes |
| 24 | CDC7 | cell division cycle 7 | 1352 | 0.075 | 0.2679 | Yes |
| 25 | XPO1 | exportin 1 | 1444 | 0.073 | 0.2720 | Yes |
| 26 | MCM3 | minichromosome maintenance complex component 3 | 1450 | 0.073 | 0.2805 | Yes |
| 27 | LMNB1 | lamin B1 | 1705 | 0.066 | 0.2754 | Yes |
| 28 | MCM5 | minichromosome maintenance complex component 5 | 1829 | 0.064 | 0.2768 | Yes |
| 29 | NUP50 | nucleoporin 50kDa | 1867 | 0.064 | 0.2825 | Yes |
| 30 | UCK2 | Jeck2013 ALT\_ACCEPTOR, ALT\_DONOR, coding, INTERNAL, intronic best transcript NM\_012474 | 1906 | 0.063 | 0.2882 | Yes |
| 31 | DTYMK | deoxythymidylate kinase | 1938 | 0.062 | 0.2941 | Yes |
| 32 | SLC12A2 | solute carrier family 12 (sodium/potassium/chloride transporter), member 2 | 2023 | 0.061 | 0.2971 | Yes |
| 33 | MAPK14 | mitogen-activated protein kinase 14 | 2039 | 0.061 | 0.3037 | Yes |
| 34 | NOTCH2 | notch 2 | 2091 | 0.060 | 0.3083 | Yes |
| 35 | MAD2L1 | MAD2 mitotic arrest deficient-like 1 (yeast) | 2149 | 0.059 | 0.3125 | Yes |
| 36 | DBF4 | DBF4 zinc finger | 2187 | 0.058 | 0.3176 | Yes |
| 37 | CDC6 | cell division cycle 6 | 2205 | 0.058 | 0.3238 | Yes |
| 38 | HMMR | hyaluronan-mediated motility receptor (RHAMM) | 2515 | 0.053 | 0.3142 | Yes |
| 39 | KMT5A | lysine (K)-specific methyltransferase 5A | 2552 | 0.053 | 0.3188 | Yes |
| 40 | PRMT5 | protein arginine methyltransferase 5 | 2560 | 0.053 | 0.3248 | Yes |
| 41 | BARD1 | BRCA1 associated RING domain 1 | 2585 | 0.053 | 0.3299 | Yes |
| 42 | FANCC | Fanconi anemia complementation group C | 2618 | 0.052 | 0.3345 | Yes |
| 43 | MEIS1 | Meis homeobox 1 | 2689 | 0.051 | 0.3371 | Yes |
| 44 | HSPA8 | heat shock 70kDa protein 8 | 2704 | 0.051 | 0.3425 | Yes |
| 45 | PRC1 | protein regulator of cytokinesis 1 | 2775 | 0.050 | 0.3449 | Yes |
| 46 | TACC3 | transforming, acidic coiled-coil containing protein 3 | 2794 | 0.050 | 0.3500 | Yes |
| 47 | LBR | lamin B receptor | 2818 | 0.050 | 0.3548 | Yes |
| 48 | EGF | epidermal growth factor | 2875 | 0.049 | 0.3578 | Yes |
| 49 | CENPE | centromere protein E | 2957 | 0.048 | 0.3594 | Yes |
| 50 | SRSF2 | serine/arginine-rich splicing factor 2 | 2978 | 0.048 | 0.3641 | Yes |
| 51 | HNRNPD | heterogeneous nuclear ribonucleoprotein D | 3023 | 0.047 | 0.3676 | Yes |
| 52 | AMD1 | adenosylmethionine decarboxylase 1 | 3049 | 0.047 | 0.3719 | Yes |
| 53 | KATNA1 | katanin p60 (ATPase containing) subunit A 1 | 3069 | 0.047 | 0.3766 | Yes |
| 54 | SMC4 | structural maintenance of chromosomes 4 | 3153 | 0.046 | 0.3779 | Yes |
| 55 | CCNA2 | cyclin A2 | 3183 | 0.046 | 0.3819 | Yes |
| 56 | MYC | v-myc avian myelocytomatosis viral oncogene homolog | 3194 | 0.046 | 0.3869 | Yes |
| 57 | TOP1 | topoisomerase (DNA) I | 3306 | 0.045 | 0.3865 | Yes |
| 58 | POLA2 | polymerase (DNA directed), alpha 2, accessory subunit | 3348 | 0.044 | 0.3897 | Yes |
| 59 | FBXO5 | F-box protein 5 | 3395 | 0.044 | 0.3926 | Yes |
| 60 | NASP | nuclear autoantigenic sperm protein (histone-binding) | 3425 | 0.043 | 0.3963 | Yes |
| 61 | MKI67 | marker of proliferation Ki-67 | 3496 | 0.043 | 0.3979 | Yes |
| 62 | CHAF1A | chromatin assembly factor 1, subunit A (p150) | 3549 | 0.042 | 0.4002 | Yes |
| 63 | MCM6 | minichromosome maintenance complex component 6 | 3558 | 0.042 | 0.4049 | Yes |
| 64 | AURKB | aurora kinase B | 3671 | 0.041 | 0.4040 | Yes |
| 65 | E2F3 | E2F transcription factor 3 | 3729 | 0.040 | 0.4060 | Yes |
| 66 | POLE | polymerase (DNA directed), epsilon, catalytic subunit | 3885 | 0.039 | 0.4026 | Yes |
| 67 | E2F2 | E2F transcription factor 2 | 3971 | 0.038 | 0.4028 | Yes |
| 68 | KIF23 | kinesin family member 23 | 4035 | 0.037 | 0.4040 | Yes |
| 69 | CHEK1 | checkpoint kinase 1 | 4083 | 0.037 | 0.4060 | Yes |
| 70 | INCENP | inner centromere protein | 4350 | 0.034 | 0.3964 | Yes |
| 71 | CDKN2C | cyclin-dependent kinase inhibitor 2C (p18, inhibits CDK4) | 4353 | 0.034 | 0.4004 | Yes |
| 72 | TRA2B | transformer 2 beta homolog (Drosophila) | 4395 | 0.034 | 0.4024 | Yes |
| 73 | DKC1 | dyskeratosis congenita 1, dyskerin | 4547 | 0.033 | 0.3985 | Yes |
| 74 | CCND1 | cyclin D1 | 4585 | 0.032 | 0.4005 | Yes |
| 75 | KIF11 | kinesin family member 11 | 4605 | 0.032 | 0.4035 | Yes |
| 76 | CUL4A | cullin 4A | 4631 | 0.032 | 0.4060 | Yes |
| 77 | CDK1 | cyclin-dependent kinase 1 | 4823 | 0.031 | 0.3999 | Yes |
| 78 | SMC1A | structural maintenance of chromosomes 1A | 4903 | 0.030 | 0.3994 | Yes |
| 79 | HMGB3 | high mobility group box 3 | 4965 | 0.030 | 0.3998 | Yes |
| 80 | AURKA | aurora kinase A | 5067 | 0.029 | 0.3981 | Yes |
| 81 | ILF3 | interleukin enhancer binding factor 3 | 5069 | 0.029 | 0.4015 | Yes |
| 82 | UPF1 | UPF1 regulator of nonsense transcripts homolog (yeast) | 5071 | 0.029 | 0.4049 | Yes |
| 83 | HMGA1 | high mobility group AT-hook 1 | 5073 | 0.029 | 0.4083 | Yes |
| 84 | GSPT1 | G1 to S phase transition 1 | 5217 | 0.028 | 0.4043 | No |
| 85 | DDX39A | DEAD (Asp-Glu-Ala-Asp) box polypeptide 39A | 5309 | 0.027 | 0.4028 | No |
| 86 | MCM2 | minichromosome maintenance complex component 2 | 5326 | 0.027 | 0.4052 | No |
| 87 | RBM14 | RNA binding motif protein 14 | 5474 | 0.026 | 0.4007 | No |
| 88 | SMARCC1 | SWI/SNF related, matrix associated, actin dependent regulator of chromatin, subfamily c, member 1 | 5534 | 0.025 | 0.4007 | No |
| 89 | NCL | nucleolin | 5618 | 0.025 | 0.3994 | No |
| 90 | PLK4 | polo-like kinase 4 | 5691 | 0.024 | 0.3986 | No |
| 91 | NOLC1 | nucleolar and coiled-body phosphoprotein 1 | 5963 | 0.022 | 0.3872 | No |
| 92 | BUB1 | BUB1 mitotic checkpoint serine/threonine kinase | 5988 | 0.022 | 0.3887 | No |
| 93 | EZH2 | enhancer of zeste 2 polycomb repressive complex 2 subunit | 5989 | 0.022 | 0.3913 | No |
| 94 | SRSF1 | serine/arginine-rich splicing factor 1 | 6000 | 0.022 | 0.3935 | No |
| 95 | PRPF4B | pre-mRNA processing factor 4B | 6054 | 0.022 | 0.3933 | No |
| 96 | KPNA2 | karyopherin alpha 2 (RAG cohort 1, importin alpha 1) | 6200 | 0.021 | 0.3883 | No |
| 97 | MT2A | metallothionein 2A | 6247 | 0.021 | 0.3884 | No |
| 98 | NDC80 | NDC80 kinetochore complex component | 6461 | 0.019 | 0.3797 | No |
| 99 | RBL1 | retinoblastoma-like 1 | 6463 | 0.019 | 0.3820 | No |
| 100 | SNRPD1 | small nuclear ribonucleoprotein D1 polypeptide | 6481 | 0.019 | 0.3834 | No |
| 101 | TFDP1 | transcription factor Dp-1 | 6725 | 0.018 | 0.3730 | No |
| 102 | RPA2 | replication protein A2 | 6755 | 0.017 | 0.3736 | No |
| 103 | RAD21 | RAD21 cohesin complex component | 6756 | 0.017 | 0.3757 | No |
| 104 | KIF5B | Memczak2013 ANTISENSE, CDS, coding, INTERNAL, UTR3 best transcript NM\_004521 | 6866 | 0.017 | 0.3720 | No |
| 105 | TOP2A | topoisomerase (DNA) II alpha | 6917 | 0.016 | 0.3714 | No |
| 106 | DMD | dystrophin | 7020 | 0.016 | 0.3680 | No |
| 107 | E2F4 | E2F transcription factor 4, p107/p130-binding | 7053 | 0.016 | 0.3682 | No |
| 108 | RACGAP1 | Rac GTPase activating protein 1 | 7079 | 0.015 | 0.3688 | No |
| 109 | SMC2 | structural maintenance of chromosomes 2 | 7150 | 0.015 | 0.3670 | No |
| 110 | CDK4 | cyclin-dependent kinase 4 | 7198 | 0.015 | 0.3663 | No |
| 111 | SYNCRIP | synaptotagmin binding, cytoplasmic RNA interacting protein | 7248 | 0.015 | 0.3656 | No |
| 112 | PURA | purine-rich element binding protein A | 7257 | 0.014 | 0.3669 | No |
| 113 | SMAD3 | SMAD family member 3 | 7263 | 0.014 | 0.3684 | No |
| 114 | G3BP1 | GTPase activating protein (SH3 domain) binding protein 1 | 7286 | 0.014 | 0.3690 | No |
| 115 | SRSF10 | serine/arginine-rich splicing factor 10 | 7444 | 0.013 | 0.3624 | No |
| 116 | CTCF | CCCTC-binding factor (zinc finger protein) | 7528 | 0.013 | 0.3597 | No |
| 117 | CCNF | cyclin F | 7608 | 0.012 | 0.3571 | No |
| 118 | KPNB1 | karyopherin (importin) beta 1 | 7709 | 0.012 | 0.3533 | No |
| 119 | SS18 | synovial sarcoma translocation, chromosome 18 | 7906 | 0.010 | 0.3444 | No |
| 120 | STIL | SCL/TAL1 interrupting locus | 7923 | 0.010 | 0.3448 | No |
| 121 | BIRC5 | baculoviral IAP repeat containing 5 | 7953 | 0.010 | 0.3446 | No |
| 122 | NUSAP1 | nucleolar and spindle associated protein 1 | 7955 | 0.010 | 0.3457 | No |
| 123 | SUV39H1 | suppressor of variegation 3-9 homolog 1 (Drosophila) | 7990 | 0.010 | 0.3452 | No |
| 124 | ESPL1 | extra spindle pole bodies like 1, separase | 7999 | 0.010 | 0.3459 | No |
| 125 | CCNB2 | cyclin B2 | 8020 | 0.010 | 0.3461 | No |
| 126 | TMPO | thymopoietin | 8045 | 0.010 | 0.3460 | No |
| 127 | ODF2 | outer dense fiber of sperm tails 2 | 8057 | 0.010 | 0.3466 | No |
| 128 | MTF2 | metal response element binding transcription factor 2 | 8147 | 0.009 | 0.3431 | No |
| 129 | STAG1 | stromal antigen 1 | 8239 | 0.008 | 0.3394 | No |
| 130 | SFPQ | splicing factor proline/glutamine-rich | 8319 | 0.008 | 0.3363 | No |
| 131 | SAP30 | Sin3A associated protein 30kDa | 8495 | 0.007 | 0.3280 | No |
| 132 | UBE2C | ubiquitin-conjugating enzyme E2C | 8543 | 0.007 | 0.3264 | No |
| 133 | HNRNPU | heterogeneous nuclear ribonucleoprotein U (scaffold attachment factor A) | 8561 | 0.007 | 0.3263 | No |
| 134 | RASAL2 | RAS protein activator like 2 | 8590 | 0.006 | 0.3257 | No |
| 135 | PLK1 | polo-like kinase 1 | 8734 | 0.006 | 0.3189 | No |
| 136 | ORC5 | origin recognition complex subunit 5 | 8805 | 0.005 | 0.3159 | No |
| 137 | HMGN2 | high mobility group nucleosomal binding domain 2 | 9015 | 0.004 | 0.3056 | No |
| 138 | CUL3 | cullin 3 | 9256 | 0.003 | 0.2935 | No |
| 139 | TROAP | trophinin associated protein | 9268 | 0.003 | 0.2933 | No |
| 140 | TRAIP | TRAF interacting protein | 9362 | 0.002 | 0.2887 | No |
| 141 | CDKN3 | cyclin-dependent kinase inhibitor 3 | 9369 | 0.002 | 0.2887 | No |
| 142 | PTTG1 | pituitary tumor-transforming 1 | 9499 | 0.001 | 0.2822 | No |
| 143 | E2F1 | Memczak2013 ANTISENSE, coding, INTERNAL, intronic best transcript NM\_005225 | 9936 | -0.001 | 0.2597 | No |
| 144 | MEIS2 | Meis homeobox 2 | 10042 | -0.001 | 0.2545 | No |
| 145 | BUB3 | BUB3 mitotic checkpoint protein | 10144 | -0.002 | 0.2495 | No |
| 146 | CKS1B | CDC28 protein kinase regulatory subunit 1B | 10258 | -0.003 | 0.2440 | No |
| 147 | NUP98 | Transcript Identified by AceView, Entrez Gene ID(s) 4928 | 10376 | -0.003 | 0.2383 | No |
| 148 | NUMA1 | nuclear mitotic apparatus protein 1 | 10687 | -0.005 | 0.2229 | No |
| 149 | TPX2 | TPX2, microtubule-associated | 10963 | -0.007 | 0.2095 | No |
| 150 | PML | promyelocytic leukemia | 10973 | -0.007 | 0.2099 | No |
| 151 | CCNT1 | cyclin T1 | 11023 | -0.007 | 0.2082 | No |
| 152 | MNAT1 | MNAT CDK-activating kinase assembly factor 1 | 11112 | -0.008 | 0.2046 | No |
| 153 | TNPO2 | transportin 2 | 11247 | -0.008 | 0.1987 | No |
| 154 | KIF2C | kinesin family member 2C | 11587 | -0.010 | 0.1824 | No |
| 155 | NEK2 | NIMA-related kinase 2 | 11616 | -0.011 | 0.1822 | No |
| 156 | YTHDC1 | YTH domain containing 1 | 11622 | -0.011 | 0.1832 | No |
| 157 | ARID4A | AT rich interactive domain 4A (RBP1-like) | 11684 | -0.011 | 0.1814 | No |
| 158 | TLE3 | Memczak2013 ANTISENSE, coding, INTERNAL, intronic best transcript NM\_001105192 | 11746 | -0.011 | 0.1796 | No |
| 159 | CHMP1A | charged multivesicular body protein 1A | 11884 | -0.012 | 0.1740 | No |
| 160 | RAD23B | RAD23 homolog B, nucleotide excision repair protein | 12313 | -0.015 | 0.1537 | No |
| 161 | TGFB1 | transforming growth factor beta 1 | 12464 | -0.016 | 0.1478 | No |
| 162 | CENPF | centromere protein F | 12491 | -0.016 | 0.1484 | No |
| 163 | CDC27 | cell division cycle 27 | 12969 | -0.019 | 0.1260 | No |
| 164 | ODC1 | ornithine decarboxylase 1 | 13133 | -0.020 | 0.1200 | No |
| 165 | EWSR1 | EWS RNA binding protein 1 | 13307 | -0.021 | 0.1136 | No |
| 166 | DR1 | down-regulator of transcription 1 | 13404 | -0.022 | 0.1113 | No |
| 167 | KIF20B | kinesin family member 20B | 13803 | -0.025 | 0.0937 | No |
| 168 | CDC25B | cell division cycle 25B | 13972 | -0.026 | 0.0882 | No |
| 169 | HUS1 | HUS1 checkpoint clamp component | 14352 | -0.029 | 0.0721 | No |
| 170 | PAFAH1B1 | platelet-activating factor acetylhydrolase 1b, regulatory subunit 1 (45kDa) | 14439 | -0.030 | 0.0713 | No |
| 171 | WRN | Werner syndrome, RecQ helicase-like | 14454 | -0.030 | 0.0742 | No |
| 172 | KIF4A | kinesin family member 4A | 14527 | -0.031 | 0.0741 | No |
| 173 | UBE2S | ubiquitin-conjugating enzyme E2S | 14865 | -0.033 | 0.0607 | No |
| 174 | CUL1 | cullin 1 | 15794 | -0.042 | 0.0178 | No |
| 175 | BCL3 | B-cell CLL/lymphoma 3 | 15800 | -0.042 | 0.0225 | No |
| 176 | SLC38A1 | solute carrier family 38, member 1 | 15857 | -0.042 | 0.0247 | No |
| 177 | CDKN1B | cyclin-dependent kinase inhibitor 1B (p27, Kip1) | 16336 | -0.047 | 0.0057 | No |
| 178 | ABL1 | ABL proto-oncogene 1, non-receptor tyrosine kinase | 16435 | -0.049 | 0.0066 | No |
| 179 | CUL5 | cullin 5 | 16586 | -0.051 | 0.0050 | No |
| 180 | HIF1A | hypoxia inducible factor 1, alpha subunit (basic helix-loop-helix transcription factor) | 16936 | -0.056 | -0.0063 | No |
| 181 | LIG3 | ligase III, DNA, ATP-dependent | 17230 | -0.060 | -0.0142 | No |
| 182 | CBX1 | chromobox homolog 1 | 17331 | -0.062 | -0.0119 | No |
| 183 | SLC7A1 | solute carrier family 7 (cationic amino acid transporter, y+ system), member 1 | 17491 | -0.065 | -0.0122 | No |
| 184 | SLC7A5 | solute carrier family 7 (amino acid transporter light chain, L system), member 5 | 17849 | -0.073 | -0.0219 | No |
| 185 | HOXC10 | homeobox C10 | 17937 | -0.076 | -0.0173 | No |
| 186 | MARCKS | myristoylated alanine-rich protein kinase C substrate | 18167 | -0.082 | -0.0192 | No |
| 187 | FOXN3 | forkhead box N3 | 18657 | -0.100 | -0.0325 | No |
| 188 | ATF5 | activating transcription factor 5 | 18851 | -0.114 | -0.0288 | No |
| 189 | EFNA5 | ephrin-A5 | 19522 | -0.527 | 0.0001 | No |
Table: GSEA details [plain text format]

  

Fig 2: HALLMARK\_G2M\_CHECKPOINT      
 Blue-Pink O' Gram in the Space of the Analyzed GeneSet

  

Fig 3: HALLMARK\_G2M\_CHECKPOINT: Random ES distribution      
 Gene set null distribution of ES for **HALLMARK\_G2M\_CHECKPOINT**

  
