## Supplementary material for "Somatic hypomethylation of pericentromeric SST1 repeats and tetraploidization in human colorectal cancer cells": GSEA results: HALLMARK_GLYCOLYSIS.html

Details for gene set HALLMARK\_GLYCOLYSIS[GSEA]

|  || Dataset | eset\_byprobe\_collapsed\_to\_symbols.Diploid\_vs\_Tetraploid.cls #Tetraploid\_versus\_Diploid.Diploid\_vs\_Tetraploid.cls #Tetraploid\_versus\_Diploid\_repos |
| Phenotype | Diploid\_vs\_Tetraploid.cls#Tetraploid\_versus\_Diploid\_repos |
| Upregulated in class | Diploid |
| GeneSet | HALLMARK\_GLYCOLYSIS |
| Enrichment Score (ES) | -0.26923916 |
| Normalized Enrichment Score (NES) | -1.123537 |
| Nominal p-value | 0.22189349 |
| FDR q-value | 0.3593222 |
| FWER p-Value | 1.0 |
Table: GSEA Results Summary

  

Fig 1: Enrichment plot: HALLMARK\_GLYCOLYSIS      
 Profile of the Running ES Score & Positions of GeneSet Members on the Rank Ordered List

  

| SYMBOL | TITLE | RANK IN GENE LIST | RANK METRIC SCORE | RUNNING ES | CORE ENRICHMENT || 1 | NT5E | 5-nucleotidase, ecto (CD73) | 37 | 0.307 | 0.0293 | No |
| 2 | RRAGD | Ras-related GTP binding D | 54 | 0.280 | 0.0569 | No |
| 3 | CHST4 | carbohydrate (N-acetylglucosamine 6-O) sulfotransferase 4 | 84 | 0.222 | 0.0780 | No |
| 4 | GNE | glucosamine (UDP-N-acetyl)-2-epimerase/N-acetylmannosamine kinase | 225 | 0.150 | 0.0860 | No |
| 5 | EFNA3 | ephrin-A3 | 448 | 0.114 | 0.0861 | No |
| 6 | TPST1 | tyrosylprotein sulfotransferase 1 | 453 | 0.114 | 0.0975 | No |
| 7 | CD44 | CD44 molecule (Indian blood group) | 577 | 0.104 | 0.1018 | No |
| 8 | IGFBP3 | insulin like growth factor binding protein 3 | 582 | 0.104 | 0.1121 | No |
| 9 | ADORA2B | adenosine A2b receptor | 596 | 0.104 | 0.1220 | No |
| 10 | CENPA | centromere protein A | 673 | 0.098 | 0.1281 | No |
| 11 | STMN1 | stathmin 1 | 951 | 0.086 | 0.1225 | No |
| 12 | FKBP4 | FK506 binding protein 4 | 973 | 0.085 | 0.1300 | No |
| 13 | IRS2 | insulin receptor substrate 2 | 1045 | 0.083 | 0.1348 | No |
| 14 | NSDHL | NAD(P) dependent steroid dehydrogenase-like | 1100 | 0.081 | 0.1402 | No |
| 15 | TGFBI | transforming growth factor, beta-induced, 68kDa | 1130 | 0.080 | 0.1469 | No |
| 16 | DPYSL4 | dihydropyrimidinase-like 4 | 1272 | 0.077 | 0.1474 | No |
| 17 | FBP2 | fructose-1,6-bisphosphatase 2 | 1356 | 0.075 | 0.1507 | No |
| 18 | EGFR | epidermal growth factor receptor | 1396 | 0.074 | 0.1562 | No |
| 19 | PGAM2 | phosphoglycerate mutase 2 (muscle) | 1477 | 0.072 | 0.1594 | No |
| 20 | GOT2 | glutamic-oxaloacetic transaminase 2, mitochondrial | 1538 | 0.070 | 0.1635 | No |
| 21 | SLC16A3 | solute carrier family 16 (monocarboxylate transporter), member 3 | 1698 | 0.067 | 0.1620 | No |
| 22 | CLDN3 | claudin 3 | 1706 | 0.066 | 0.1684 | No |
| 23 | ANGPTL4 | angiopoietin like 4 | 1873 | 0.064 | 0.1663 | No |
| 24 | PYGL | phosphorylase, glycogen, liver | 1921 | 0.063 | 0.1702 | No |
| 25 | CTH | cystathionine gamma-lyase | 2192 | 0.058 | 0.1622 | No |
| 26 | SDC2 | syndecan 2 | 2442 | 0.055 | 0.1549 | No |
| 27 | HMMR | hyaluronan-mediated motility receptor (RHAMM) | 2515 | 0.053 | 0.1566 | No |
| 28 | PKM | pyruvate kinase, muscle | 2596 | 0.052 | 0.1578 | No |
| 29 | UGP2 | UDP-glucose pyrophosphorylase 2 | 2703 | 0.051 | 0.1575 | No |
| 30 | CASP6 | caspase 6 | 2891 | 0.049 | 0.1527 | No |
| 31 | GYS2 | glycogen synthase 2 (liver) | 3022 | 0.047 | 0.1508 | No |
| 32 | NOL3 | nucleolar protein 3 (apoptosis repressor with CARD domain) | 3042 | 0.047 | 0.1546 | No |
| 33 | ENO2 | enolase 2 (gamma, neuronal) | 3086 | 0.047 | 0.1572 | No |
| 34 | STC1 | stanniocalcin 1 | 3174 | 0.046 | 0.1573 | No |
| 35 | PKP2 | plakophilin 2 | 3282 | 0.045 | 0.1563 | No |
| 36 | ABCB6 | ATP binding cassette subfamily B member 6 (Langereis blood group) | 3410 | 0.044 | 0.1542 | No |
| 37 | NASP | nuclear autoantigenic sperm protein (histone-binding) | 3425 | 0.043 | 0.1579 | No |
| 38 | POLR3K | polymerase (RNA) III (DNA directed) polypeptide K, 12.3 kDa | 3708 | 0.041 | 0.1474 | No |
| 39 | ALG1 | ALG1, chitobiosyldiphosphodolichol beta-mannosyltransferase | 3911 | 0.038 | 0.1408 | No |
| 40 | SLC37A4 | solute carrier family 37 (glucose-6-phosphate transporter), member 4 | 3916 | 0.038 | 0.1445 | No |
| 41 | GAL3ST1 | galactose-3-O-sulfotransferase 1 | 4079 | 0.037 | 0.1399 | No |
| 42 | AGRN | agrin | 4105 | 0.037 | 0.1423 | No |
| 43 | NDUFV3 | Transcript Identified by AceView, Entrez Gene ID(s) 4731 | 4152 | 0.036 | 0.1436 | No |
| 44 | GPR87 | G protein-coupled receptor 87 | 4244 | 0.035 | 0.1425 | No |
| 45 | TGFA | transforming growth factor alpha | 4449 | 0.034 | 0.1353 | No |
| 46 | MPI | mannose phosphate isomerase | 4461 | 0.033 | 0.1382 | No |
| 47 | PRPS1 | phosphoribosyl pyrophosphate synthetase 1 | 4489 | 0.033 | 0.1402 | No |
| 48 | TPI1 | triosephosphate isomerase 1 | 4710 | 0.031 | 0.1320 | No |
| 49 | CDK1 | cyclin-dependent kinase 1 | 4823 | 0.031 | 0.1293 | No |
| 50 | MIF | macrophage migration inhibitory factor (glycosylation-inhibiting factor) | 4925 | 0.030 | 0.1271 | No |
| 51 | SOD1 | superoxide dismutase 1, soluble | 4956 | 0.030 | 0.1286 | No |
| 52 | AURKA | aurora kinase A | 5067 | 0.029 | 0.1258 | No |
| 53 | EGLN3 | egl-9 family hypoxia-inducible factor 3 | 5195 | 0.028 | 0.1221 | No |
| 54 | LHX9 | LIM homeobox 9 | 5255 | 0.027 | 0.1218 | No |
| 55 | ELF3 | E74-like factor 3 (ets domain transcription factor, epithelial-specific ) | 5265 | 0.027 | 0.1241 | No |
| 56 | SDC1 | syndecan 1 | 5305 | 0.027 | 0.1248 | No |
| 57 | VLDLR | very low density lipoprotein receptor | 5549 | 0.025 | 0.1148 | No |
| 58 | ALDH9A1 | aldehyde dehydrogenase 9 family, member A1 | 5592 | 0.025 | 0.1151 | No |
| 59 | PGAM1 | phosphoglycerate mutase 1 (brain) | 5597 | 0.025 | 0.1174 | No |
| 60 | MDH1 | malate dehydrogenase 1 | 5641 | 0.024 | 0.1177 | No |
| 61 | ANKZF1 | ankyrin repeat and zinc finger domain containing 1 | 5647 | 0.024 | 0.1199 | No |
| 62 | TXN | thioredoxin | 5710 | 0.024 | 0.1191 | No |
| 63 | CHST6 | carbohydrate (N-acetylglucosamine 6-O) sulfotransferase 6 | 5772 | 0.024 | 0.1184 | No |
| 64 | LDHA | lactate dehydrogenase A | 6019 | 0.022 | 0.1079 | No |
| 65 | GUSB | glucuronidase, beta | 6020 | 0.022 | 0.1101 | No |
| 66 | CHST2 | carbohydrate (N-acetylglucosamine-6-O) sulfotransferase 2 | 6022 | 0.022 | 0.1123 | No |
| 67 | MET | MET proto-oncogene, receptor tyrosine kinase | 6216 | 0.021 | 0.1044 | No |
| 68 | B3GALT6 | UDP-Gal:betaGal beta 1,3-galactosyltransferase 6 | 6311 | 0.020 | 0.1016 | No |
| 69 | ALDOA | aldolase A, fructose-bisphosphate | 6361 | 0.020 | 0.1011 | No |
| 70 | MXI1 | MAX interactor 1, dimerization protein | 6513 | 0.019 | 0.0952 | No |
| 71 | SDHC | succinate dehydrogenase complex, subunit C, integral membrane protein, 15kDa | 7238 | 0.015 | 0.0592 | No |
| 72 | GLCE | glucuronic acid epimerase | 7373 | 0.014 | 0.0537 | No |
| 73 | GALE | UDP-galactose-4-epimerase | 7614 | 0.012 | 0.0425 | No |
| 74 | MIOX | myo-inositol oxygenase | 7642 | 0.012 | 0.0423 | No |
| 75 | GAPDHS | glyceraldehyde-3-phosphate dehydrogenase, spermatogenic | 7644 | 0.012 | 0.0435 | No |
| 76 | PHKA2 | phosphorylase kinase, alpha 2 (liver) | 7997 | 0.010 | 0.0263 | No |
| 77 | MED24 | mediator complex subunit 24 | 8183 | 0.009 | 0.0176 | No |
| 78 | PC | pyruvate carboxylase | 8281 | 0.008 | 0.0134 | No |
| 79 | HS6ST2 | heparan sulfate 6-O-sulfotransferase 2 | 8304 | 0.008 | 0.0131 | No |
| 80 | COG2 | component of oligomeric golgi complex 2 | 8314 | 0.008 | 0.0134 | No |
| 81 | PFKP | phosphofructokinase, platelet | 8326 | 0.008 | 0.0137 | No |
| 82 | LDHC | lactate dehydrogenase C | 8359 | 0.008 | 0.0128 | No |
| 83 | TALDO1 | transaldolase 1 | 8396 | 0.007 | 0.0117 | No |
| 84 | SAP30 | Sin3A associated protein 30kDa | 8495 | 0.007 | 0.0073 | No |
| 85 | HK2 | hexokinase 2 | 8575 | 0.006 | 0.0039 | No |
| 86 | SLC25A10 | solute carrier family 25 (mitochondrial carrier; dicarboxylate transporter), member 10 | 8612 | 0.006 | 0.0026 | No |
| 87 | PPIA | peptidylprolyl isomerase A (cyclophilin A) | 8627 | 0.006 | 0.0025 | No |
| 88 | ALDOB | aldolase B, fructose-bisphosphate | 8657 | 0.006 | 0.0017 | No |
| 89 | PGLS | 6-phosphogluconolactonase | 8706 | 0.006 | -0.0003 | No |
| 90 | DEPDC1 | DEP domain containing 1 | 8741 | 0.005 | -0.0015 | No |
| 91 | CXCR4 | chemokine (C-X-C motif) receptor 4 | 8789 | 0.005 | -0.0034 | No |
| 92 | B4GALT2 | UDP-Gal:betaGlcNAc beta 1,4- galactosyltransferase, polypeptide 2 | 8942 | 0.004 | -0.0108 | No |
| 93 | ALDH7A1 | aldehyde dehydrogenase 7 family, member A1 | 9127 | 0.003 | -0.0200 | No |
| 94 | SLC35A3 | solute carrier family 35 (UDP-N-acetylglucosamine (UDP-GlcNAc) transporter), member A3 | 9168 | 0.003 | -0.0217 | No |
| 95 | RPE | ribulose-5-phosphate-3-epimerase | 9326 | 0.002 | -0.0296 | No |
| 96 | SPAG4 | sperm associated antigen 4 | 9990 | -0.001 | -0.0638 | No |
| 97 | AGL | amylo-alpha-1, 6-glucosidase, 4-alpha-glucanotransferase | 10036 | -0.001 | -0.0660 | No |
| 98 | CHST1 | carbohydrate (keratan sulfate Gal-6) sulfotransferase 1 | 10102 | -0.002 | -0.0692 | No |
| 99 | SOX9 | SRY box 9 | 10277 | -0.003 | -0.0779 | No |
| 100 | PGM2 | phosphoglucomutase 2 | 10296 | -0.003 | -0.0785 | No |
| 101 | ECD | ecdysoneless homolog (Drosophila) | 10389 | -0.003 | -0.0829 | No |
| 102 | KIF20A | kinesin family member 20A | 10416 | -0.004 | -0.0839 | No |
| 103 | CLN6 | ceroid-lipofuscinosis, neuronal 6, late infantile, variant | 10458 | -0.004 | -0.0856 | No |
| 104 | ME2 | malic enzyme 2, NAD(+)-dependent, mitochondrial | 10643 | -0.005 | -0.0946 | No |
| 105 | FAM162A | family with sequence similarity 162, member A | 10663 | -0.005 | -0.0951 | No |
| 106 | NDST3 | N-deacetylase/N-sulfotransferase (heparan glucosaminyl) 3 | 10704 | -0.005 | -0.0966 | No |
| 107 | TSTA3 | tissue specific transplantation antigen P35B | 10774 | -0.006 | -0.0996 | No |
| 108 | BPNT1 | 3(2), 5-bisphosphate nucleotidase 1 | 10796 | -0.006 | -0.1001 | No |
| 109 | XYLT2 | xylosyltransferase II | 10858 | -0.006 | -0.1026 | No |
| 110 | FUT8 | fucosyltransferase 8 (alpha (1,6) fucosyltransferase) | 10949 | -0.007 | -0.1066 | No |
| 111 | PLOD1 | procollagen-lysine, 2-oxoglutarate 5-dioxygenase 1 | 10966 | -0.007 | -0.1067 | No |
| 112 | GOT1 | glutamic-oxaloacetic transaminase 1, soluble | 11039 | -0.007 | -0.1097 | No |
| 113 | PGK1 | phosphoglycerate kinase 1 | 11128 | -0.008 | -0.1134 | No |
| 114 | PYGB | phosphorylase, glycogen; brain | 11215 | -0.008 | -0.1170 | No |
| 115 | B4GALT7 | xylosylprotein beta 1,4-galactosyltransferase, polypeptide 7 | 11228 | -0.008 | -0.1168 | No |
| 116 | PAXIP1 | PAX interacting (with transcription-activation domain) protein 1 | 11428 | -0.010 | -0.1261 | No |
| 117 | HOMER1 | homer scaffolding protein 1 | 11561 | -0.010 | -0.1319 | No |
| 118 | GPC1 | glypican 1 | 11601 | -0.010 | -0.1329 | No |
| 119 | ENO1 | Memczak2013 ANTISENSE, coding, INTERNAL, UTR3 best transcript NM\_001428 | 11613 | -0.010 | -0.1324 | No |
| 120 | TFF3 | trefoil factor 3 | 12114 | -0.014 | -0.1569 | No |
| 121 | DLD | dihydrolipoamide dehydrogenase | 12560 | -0.016 | -0.1783 | No |
| 122 | SRD5A3 | steroid 5 alpha-reductase 3 | 12647 | -0.017 | -0.1810 | No |
| 123 | PSMC4 | proteasome 26S subunit, ATPase 4 | 12803 | -0.018 | -0.1872 | No |
| 124 | CHST12 | carbohydrate (chondroitin 4) sulfotransferase 12 | 12838 | -0.018 | -0.1871 | No |
| 125 | GMPPB | GDP-mannose pyrophosphorylase B | 12894 | -0.018 | -0.1881 | No |
| 126 | ARTN | artemin | 12966 | -0.019 | -0.1898 | No |
| 127 | G6PD | glucose-6-phosphate dehydrogenase | 13330 | -0.021 | -0.2064 | No |
| 128 | PPP2CB | protein phosphatase 2, catalytic subunit, beta isozyme | 13359 | -0.022 | -0.2056 | No |
| 129 | KIF2A | kinesin heavy chain member 2A | 13362 | -0.022 | -0.2035 | No |
| 130 | CYB5A | cytochrome b5 type A (microsomal) | 13471 | -0.022 | -0.2069 | No |
| 131 | GALK2 | galactokinase 2 | 13550 | -0.023 | -0.2086 | No |
| 132 | DSC2 | desmocollin 2 | 13555 | -0.023 | -0.2064 | No |
| 133 | P4HA1 | prolyl 4-hydroxylase, alpha polypeptide I | 13604 | -0.023 | -0.2065 | No |
| 134 | PAM | peptidylglycine alpha-amidating monooxygenase | 13616 | -0.023 | -0.2047 | No |
| 135 | ERO1A | endoplasmic reticulum oxidoreductase alpha | 13862 | -0.025 | -0.2148 | No |
| 136 | IER3 | immediate early response 3 | 14076 | -0.027 | -0.2231 | No |
| 137 | ARPP19 | cAMP-regulated phosphoprotein 19kDa | 14107 | -0.027 | -0.2219 | No |
| 138 | GYS1 | glycogen synthase 1 (muscle) | 14325 | -0.029 | -0.2302 | No |
| 139 | GNPDA1 | glucosamine-6-phosphate deaminase 1 | 14449 | -0.030 | -0.2335 | No |
| 140 | MDH2 | malate dehydrogenase 2 | 14465 | -0.030 | -0.2312 | No |
| 141 | GALK1 | galactokinase 1 | 14731 | -0.032 | -0.2417 | No |
| 142 | PPFIA4 | protein tyrosine phosphatase, receptor type, f polypeptide (PTPRF), interacting protein (liprin), alpha 4 | 14744 | -0.032 | -0.2390 | No |
| 143 | NANP | N-acetylneuraminic acid phosphatase | 14759 | -0.032 | -0.2364 | No |
| 144 | IDH1 | isocitrate dehydrogenase 1 (NADP+) | 14950 | -0.034 | -0.2428 | No |
| 145 | AK3 | adenylate kinase 3 | 15257 | -0.037 | -0.2549 | No |
| 146 | DCN | decorin | 15284 | -0.037 | -0.2525 | No |
| 147 | B3GNT3 | UDP-GlcNAc:betaGal beta-1,3-N-acetylglucosaminyltransferase 3 | 15333 | -0.037 | -0.2512 | No |
| 148 | SLC25A13 | solute carrier family 25 (aspartate/glutamate carrier), member 13 | 15423 | -0.038 | -0.2519 | No |
| 149 | PMM2 | phosphomannomutase 2 | 15626 | -0.040 | -0.2583 | No |
| 150 | IDUA | iduronidase, alpha-L- | 15706 | -0.041 | -0.2582 | No |
| 151 | ZNF292 | zinc finger protein 292 | 15825 | -0.042 | -0.2601 | No |
| 152 | EXT2 | exostosin glycosyltransferase 2 | 15864 | -0.042 | -0.2577 | No |
| 153 | AKR1A1 | aldo-keto reductase family 1, member A1 (aldehyde reductase) | 15918 | -0.043 | -0.2561 | No |
| 154 | P4HA2 | prolyl 4-hydroxylase, alpha polypeptide II | 15996 | -0.044 | -0.2557 | No |
| 155 | HS2ST1 | heparan sulfate 2-O-sulfotransferase 1 | 16210 | -0.046 | -0.2620 | No |
| 156 | ISG20 | interferon stimulated exonuclease gene 20kDa | 16332 | -0.047 | -0.2634 | No |
| 157 | B4GALT1 | UDP-Gal:betaGlcNAc beta 1,4- galactosyltransferase, polypeptide 1 | 16445 | -0.049 | -0.2643 | Yes |
| 158 | GLRX | glutaredoxin | 16461 | -0.049 | -0.2600 | Yes |
| 159 | GPC3 | glypican 3 | 16605 | -0.051 | -0.2622 | Yes |
| 160 | VEGFA | vascular endothelial growth factor A | 16659 | -0.052 | -0.2597 | Yes |
| 161 | RBCK1 | RanBP-type and C3HC4-type zinc finger containing 1 | 16750 | -0.053 | -0.2589 | Yes |
| 162 | B3GAT3 | beta-1,3-glucuronyltransferase 3 | 16803 | -0.054 | -0.2561 | Yes |
| 163 | AK4 | adenylate kinase 4 | 16854 | -0.055 | -0.2532 | Yes |
| 164 | PFKFB1 | 6-phosphofructo-2-kinase/fructose-2,6-biphosphatase 1 | 16893 | -0.055 | -0.2495 | Yes |
| 165 | TPBG | trophoblast glycoprotein | 17065 | -0.058 | -0.2525 | Yes |
| 166 | HDLBP | Jeck2013 ANTISENSE, coding, INTERNAL, intronic best transcript NM\_005336 | 17127 | -0.059 | -0.2497 | Yes |
| 167 | STC2 | stanniocalcin 2 | 17326 | -0.062 | -0.2536 | Yes |
| 168 | LCT | lactase | 17328 | -0.062 | -0.2473 | Yes |
| 169 | BIK | BCL2-interacting killer (apoptosis-inducing) | 17482 | -0.065 | -0.2486 | Yes |
| 170 | DDIT4 | DNA damage inducible transcript 4 | 17557 | -0.066 | -0.2457 | Yes |
| 171 | PLOD2 | procollagen-lysine, 2-oxoglutarate 5-dioxygenase 2 | 17575 | -0.067 | -0.2398 | Yes |
| 172 | HAX1 | HCLS1 associated protein X-1 | 17593 | -0.067 | -0.2339 | Yes |
| 173 | QSOX1 | quiescin Q6 sulfhydryl oxidase 1 | 17654 | -0.069 | -0.2300 | Yes |
| 174 | COPB2 | coatomer protein complex subunit beta 2 (beta prime) | 17673 | -0.069 | -0.2239 | Yes |
| 175 | B4GALT4 | UDP-Gal:betaGlcNAc beta 1,4- galactosyltransferase, polypeptide 4 | 17701 | -0.070 | -0.2183 | Yes |
| 176 | KDELR3 | KDEL (Lys-Asp-Glu-Leu) endoplasmic reticulum protein retention receptor 3 | 17705 | -0.070 | -0.2113 | Yes |
| 177 | TKTL1 | transketolase-like 1 | 17783 | -0.072 | -0.2080 | Yes |
| 178 | GPC4 | glypican 4 | 17805 | -0.072 | -0.2018 | Yes |
| 179 | LHPP | phospholysine phosphohistidine inorganic pyrophosphate phosphatase | 17841 | -0.073 | -0.1962 | Yes |
| 180 | GMPPA | GDP-mannose pyrophosphorylase A | 17855 | -0.073 | -0.1894 | Yes |
| 181 | CAPN5 | calpain 5 | 17885 | -0.074 | -0.1833 | Yes |
| 182 | GCLC | glutamate-cysteine ligase, catalytic subunit | 17981 | -0.077 | -0.1805 | Yes |
| 183 | ME1 | malic enzyme 1, NADP(+)-dependent, cytosolic | 18066 | -0.079 | -0.1768 | Yes |
| 184 | CLDN9 | claudin 9 | 18115 | -0.080 | -0.1712 | Yes |
| 185 | IL13RA1 | interleukin 13 receptor, alpha 1 | 18154 | -0.081 | -0.1649 | Yes |
| 186 | PDK3 | pyruvate dehydrogenase kinase, isozyme 3 | 18210 | -0.083 | -0.1593 | Yes |
| 187 | SDC3 | syndecan 3 | 18233 | -0.084 | -0.1519 | Yes |
| 188 | CITED2 | Cbp/p300-interacting transactivator, with Glu/Asp rich carboxy-terminal domain, 2 | 18283 | -0.085 | -0.1459 | Yes |
| 189 | GFPT1 | glutamine--fructose-6-phosphate transaminase 1 | 18405 | -0.089 | -0.1431 | Yes |
| 190 | HSPA5 | heat shock 70kDa protein 5 (glucose-regulated protein, 78kDa) | 18430 | -0.090 | -0.1352 | Yes |
| 191 | EXT1 | exostosin glycosyltransferase 1 | 18542 | -0.094 | -0.1314 | Yes |
| 192 | B3GAT1 | beta-1,3-glucuronyltransferase 1 | 18600 | -0.096 | -0.1245 | Yes |
| 193 | VCAN | versican | 18696 | -0.103 | -0.1190 | Yes |
| 194 | CHPF2 | chondroitin polymerizing factor 2 | 18777 | -0.108 | -0.1122 | Yes |
| 195 | CHPF | chondroitin polymerizing factor | 19020 | -0.129 | -0.1116 | Yes |
| 196 | CACNA1H | calcium channel, voltage-dependent, T type, alpha 1H subunit | 19446 | -0.296 | -0.1034 | Yes |
| 197 | COL5A1 | collagen, type V, alpha 1 | 19466 | -0.322 | -0.0716 | Yes |
| 198 | MERTK | MER proto-oncogene, tyrosine kinase | 19468 | -0.325 | -0.0387 | Yes |
| 199 | ANG | angiogenin, ribonuclease, RNase A family, 5 | 19501 | -0.408 | 0.0012 | Yes |
Table: GSEA details [plain text format]

  

Fig 2: HALLMARK\_GLYCOLYSIS      
 Blue-Pink O' Gram in the Space of the Analyzed GeneSet

  

Fig 3: HALLMARK\_GLYCOLYSIS: Random ES distribution      
 Gene set null distribution of ES for **HALLMARK\_GLYCOLYSIS**

  
