## Supplementary material for "Somatic hypomethylation of pericentromeric SST1 repeats and tetraploidization in human colorectal cancer cells": GSEA results: HALLMARK_HEDGEHOG_SIGNALING.html

Details for gene set HALLMARK\_HEDGEHOG\_SIGNALING[GSEA]

|  || Dataset | eset\_byprobe\_collapsed\_to\_symbols.Diploid\_vs\_Tetraploid.cls #Tetraploid\_versus\_Diploid.Diploid\_vs\_Tetraploid.cls #Tetraploid\_versus\_Diploid\_repos |
| Phenotype | Diploid\_vs\_Tetraploid.cls#Tetraploid\_versus\_Diploid\_repos |
| Upregulated in class | Tetraploid |
| GeneSet | HALLMARK\_HEDGEHOG\_SIGNALING |
| Enrichment Score (ES) | 0.2998275 |
| Normalized Enrichment Score (NES) | 0.9723793 |
| Nominal p-value | 0.49299064 |
| FDR q-value | 0.7766785 |
| FWER p-Value | 1.0 |
Table: GSEA Results Summary

  

Fig 1: Enrichment plot: HALLMARK\_HEDGEHOG\_SIGNALING      
 Profile of the Running ES Score & Positions of GeneSet Members on the Rank Ordered List

  

| SYMBOL | TITLE | RANK IN GENE LIST | RANK METRIC SCORE | RUNNING ES | CORE ENRICHMENT || 1 | NRCAM | neuronal cell adhesion molecule | 27 | 0.351 | 0.2010 | Yes |
| 2 | TLE1 | transducin-like enhancer of split 1 (E(sp1) homolog, Drosophila) | 331 | 0.128 | 0.2592 | Yes |
| 3 | CNTFR | ciliary neurotrophic factor receptor | 920 | 0.087 | 0.2794 | Yes |
| 4 | NRP1 | neuropilin 1 | 1363 | 0.075 | 0.2998 | Yes |
| 5 | ETS2 | v-ets avian erythroblastosis virus E26 oncogene homolog 2 | 2456 | 0.054 | 0.2752 | No |
| 6 | MYH9 | Jeck2013 ANTISENSE, CDS, coding, INTERNAL, intronic, OVCODE, OVEXON best transcript NM\_002473 | 3583 | 0.042 | 0.2414 | No |
| 7 | NKX6-1 | NK6 homeobox 1 | 4052 | 0.037 | 0.2387 | No |
| 8 | PLG | plasminogen | 4359 | 0.034 | 0.2428 | No |
| 9 | SLIT1 | slit guidance ligand 1 | 4577 | 0.032 | 0.2504 | No |
| 10 | THY1 | Thy-1 cell surface antigen | 4641 | 0.032 | 0.2656 | No |
| 11 | CDK5R1 | cyclin-dependent kinase 5, regulatory subunit 1 (p35) | 5166 | 0.028 | 0.2549 | No |
| 12 | VLDLR | very low density lipoprotein receptor | 5549 | 0.025 | 0.2497 | No |
| 13 | ACHE | acetylcholinesterase (Yt blood group) | 7104 | 0.015 | 0.1788 | No |
| 14 | GLI1 | GLI family zinc finger 1 | 7287 | 0.014 | 0.1776 | No |
| 15 | L1CAM | L1 cell adhesion molecule | 7300 | 0.014 | 0.1852 | No |
| 16 | LDB1 | LIM domain binding 1 | 8125 | 0.009 | 0.1482 | No |
| 17 | NRP2 | neuropilin 2 | 9297 | 0.002 | 0.0896 | No |
| 18 | RASA1 | RAS p21 protein activator (GTPase activating protein) 1 | 9673 | 0.001 | 0.0706 | No |
| 19 | SCG2 | secretogranin II | 10022 | -0.001 | 0.0536 | No |
| 20 | PTCH1 | patched 1 | 10420 | -0.004 | 0.0353 | No |
| 21 | DPYSL2 | dihydropyrimidinase-like 2 | 10784 | -0.006 | 0.0200 | No |
| 22 | CDK6 | cyclin-dependent kinase 6 | 10904 | -0.007 | 0.0177 | No |
| 23 | PML | promyelocytic leukemia | 10973 | -0.007 | 0.0182 | No |
| 24 | CRMP1 | collapsin response mediator protein 1 | 11185 | -0.008 | 0.0121 | No |
| 25 | CELSR1 | cadherin, EGF LAG seven-pass G-type receptor 1 | 11656 | -0.011 | -0.0059 | No |
| 26 | TLE3 | Memczak2013 ANTISENSE, coding, INTERNAL, intronic best transcript NM\_001105192 | 11746 | -0.011 | -0.0039 | No |
| 27 | RTN1 | reticulon 1 | 13491 | -0.023 | -0.0804 | No |
| 28 | UNC5C | unc-5 netrin receptor C | 14386 | -0.030 | -0.1093 | No |
| 29 | NF1 | neurofibromin 1 | 15828 | -0.042 | -0.1591 | No |
| 30 | VEGFA | vascular endothelial growth factor A | 16659 | -0.052 | -0.1717 | No |
| 31 | OPHN1 | oligophrenin 1 | 17629 | -0.068 | -0.1822 | No |
| 32 | ADGRG1 | adhesion G protein-coupled receptor G1 | 17890 | -0.074 | -0.1528 | No |
| 33 | HEY1 | hes-related family bHLH transcription factor with YRPW motif 1 | 18270 | -0.085 | -0.1235 | No |
| 34 | SHH | sonic hedgehog | 18497 | -0.093 | -0.0816 | No |
| 35 | AMOT | angiomotin | 18871 | -0.116 | -0.0342 | No |
| 36 | HEY2 | hes-related family bHLH transcription factor with YRPW motif 2 | 18894 | -0.118 | 0.0323 | No |
Table: GSEA details [plain text format]

  

Fig 2: HALLMARK\_HEDGEHOG\_SIGNALING      
 Blue-Pink O' Gram in the Space of the Analyzed GeneSet

  

Fig 3: HALLMARK\_HEDGEHOG\_SIGNALING: Random ES distribution      
 Gene set null distribution of ES for **HALLMARK\_HEDGEHOG\_SIGNALING**

  
