## Supplementary material for "Somatic hypomethylation of pericentromeric SST1 repeats and tetraploidization in human colorectal cancer cells": GSEA results: HALLMARK_HEME_METABOLISM.html

Details for gene set HALLMARK\_HEME\_METABOLISM[GSEA]

|  || Dataset | eset\_byprobe\_collapsed\_to\_symbols.Diploid\_vs\_Tetraploid.cls #Tetraploid\_versus\_Diploid.Diploid\_vs\_Tetraploid.cls #Tetraploid\_versus\_Diploid\_repos |
| Phenotype | Diploid\_vs\_Tetraploid.cls#Tetraploid\_versus\_Diploid\_repos |
| Upregulated in class | Diploid |
| GeneSet | HALLMARK\_HEME\_METABOLISM |
| Enrichment Score (ES) | -0.28365332 |
| Normalized Enrichment Score (NES) | -1.1602432 |
| Nominal p-value | 0.16512346 |
| FDR q-value | 0.3559384 |
| FWER p-Value | 0.997 |
Table: GSEA Results Summary

  

Fig 1: Enrichment plot: HALLMARK\_HEME\_METABOLISM      
 Profile of the Running ES Score & Positions of GeneSet Members on the Rank Ordered List

  

| SYMBOL | TITLE | RANK IN GENE LIST | RANK METRIC SCORE | RUNNING ES | CORE ENRICHMENT || 1 | CTSE | cathepsin E | 9 | 0.419 | 0.0471 | No |
| 2 | CA2 | carbonic anhydrase II | 69 | 0.250 | 0.0723 | No |
| 3 | RHD | Rh blood group, D antigen | 118 | 0.192 | 0.0916 | No |
| 4 | C3 | complement component 3 | 143 | 0.179 | 0.1107 | No |
| 5 | BTG2 | BTG family, member 2 | 281 | 0.136 | 0.1191 | No |
| 6 | TMCC2 | transmembrane and coiled-coil domain family 2 | 354 | 0.125 | 0.1295 | No |
| 7 | PDZK1IP1 | PDZK1 interacting protein 1 | 399 | 0.119 | 0.1407 | No |
| 8 | SELENBP1 | selenium binding protein 1 | 563 | 0.105 | 0.1442 | No |
| 9 | UBAC1 | UBA domain containing 1 | 754 | 0.094 | 0.1451 | No |
| 10 | CPOX | coproporphyrinogen oxidase | 767 | 0.094 | 0.1551 | No |
| 11 | CTNS | cystinosin, lysosomal cystine transporter | 781 | 0.093 | 0.1650 | No |
| 12 | ELL2 | elongation factor, RNA polymerase II, 2 | 918 | 0.087 | 0.1679 | No |
| 13 | DMTN | dematin actin binding protein | 1303 | 0.076 | 0.1566 | No |
| 14 | SLC4A1 | solute carrier family 4 (anion exchanger), member 1 (Diego blood group) | 1313 | 0.076 | 0.1648 | No |
| 15 | SLC25A37 | solute carrier family 25 (mitochondrial iron transporter), member 37 | 1638 | 0.068 | 0.1557 | No |
| 16 | KAT2B | K(lysine) acetyltransferase 2B | 2929 | 0.048 | 0.0945 | No |
| 17 | HTRA2 | HtrA serine peptidase 2 | 3007 | 0.047 | 0.0959 | No |
| 18 | SNCA | synuclein alpha | 3009 | 0.047 | 0.1012 | No |
| 19 | HBZ | hemoglobin, zeta | 3017 | 0.047 | 0.1062 | No |
| 20 | GMPS | guanine monophosphate synthase | 3161 | 0.046 | 0.1040 | No |
| 21 | CDR2 | cerebellar degeneration related protein 2 | 3233 | 0.045 | 0.1055 | No |
| 22 | TOP1 | topoisomerase (DNA) I | 3306 | 0.045 | 0.1068 | No |
| 23 | ABCB6 | ATP binding cassette subfamily B member 6 (Langereis blood group) | 3410 | 0.044 | 0.1064 | No |
| 24 | GATA1 | GATA binding protein 1 (globin transcription factor 1) | 3535 | 0.042 | 0.1048 | No |
| 25 | E2F2 | E2F transcription factor 2 | 3971 | 0.038 | 0.0866 | No |
| 26 | RBM5 | RNA binding motif protein 5 | 4064 | 0.037 | 0.0860 | No |
| 27 | OSBP2 | oxysterol binding protein 2 | 4153 | 0.036 | 0.0856 | No |
| 28 | SLC22A4 | solute carrier family 22 (organic cation/zwitterion transporter), member 4 | 4180 | 0.036 | 0.0883 | No |
| 29 | GYPC | glycophorin C (Gerbich blood group) | 4447 | 0.034 | 0.0783 | No |
| 30 | NR3C1 | nuclear receptor subfamily 3, group C, member 1 (glucocorticoid receptor) | 4470 | 0.033 | 0.0810 | No |
| 31 | FN3K | fructosamine 3 kinase | 4629 | 0.032 | 0.0764 | No |
| 32 | ACKR1 | atypical chemokine receptor 1 (Duffy blood group) | 4812 | 0.031 | 0.0705 | No |
| 33 | ICAM4 | intercellular adhesion molecule 4 (Landsteiner-Wiener blood group) | 4895 | 0.030 | 0.0697 | No |
| 34 | RCL1 | RNA terminal phosphate cyclase-like 1 | 5135 | 0.028 | 0.0605 | No |
| 35 | RHCE | Rh blood group, CcEe antigens | 5334 | 0.027 | 0.0533 | No |
| 36 | SLC11A2 | solute carrier family 11 (proton-coupled divalent metal ion transporter), member 2 | 5392 | 0.026 | 0.0534 | No |
| 37 | BLVRB | biliverdin reductase B | 5918 | 0.023 | 0.0288 | No |
| 38 | ACP5 | acid phosphatase 5, tartrate resistant | 5930 | 0.022 | 0.0307 | No |
| 39 | PRDX2 | peroxiredoxin 2 | 6057 | 0.022 | 0.0267 | No |
| 40 | BLVRA | biliverdin reductase A | 6229 | 0.021 | 0.0202 | No |
| 41 | ACSL6 | acyl-CoA synthetase long-chain family member 6 | 6278 | 0.020 | 0.0200 | No |
| 42 | FBXO34 | F-box protein 34 | 6295 | 0.020 | 0.0215 | No |
| 43 | HBQ1 | hemoglobin, theta 1 | 6390 | 0.020 | 0.0189 | No |
| 44 | MXI1 | MAX interactor 1, dimerization protein | 6513 | 0.019 | 0.0147 | No |
| 45 | HMBS | hydroxymethylbilane synthase | 6994 | 0.016 | -0.0083 | No |
| 46 | UROD | uroporphyrinogen decarboxylase | 7213 | 0.015 | -0.0179 | No |
| 47 | SLC10A3 | solute carrier family 10, member 3 | 7384 | 0.014 | -0.0252 | No |
| 48 | AHSP | alpha hemoglobin stabilizing protein | 7418 | 0.013 | -0.0254 | No |
| 49 | EZH1 | enhancer of zeste 1 polycomb repressive complex 2 subunit | 7593 | 0.012 | -0.0330 | No |
| 50 | UROS | uroporphyrinogen III synthase | 7623 | 0.012 | -0.0331 | No |
| 51 | ISCA1 | iron-sulfur cluster assembly 1 | 7643 | 0.012 | -0.0327 | No |
| 52 | TNS1 | tensin 1 | 7662 | 0.012 | -0.0323 | No |
| 53 | SLC30A10 | solute carrier family 30, member 10 | 7788 | 0.011 | -0.0375 | No |
| 54 | LRP10 | LDL receptor related protein 10 | 7878 | 0.011 | -0.0409 | No |
| 55 | XK | X-linked Kx blood group | 8101 | 0.009 | -0.0513 | No |
| 56 | FOXO3 | forkhead box O3 | 8135 | 0.009 | -0.0520 | No |
| 57 | PPOX | protoporphyrinogen oxidase | 8217 | 0.009 | -0.0552 | No |
| 58 | PC | pyruvate carboxylase | 8281 | 0.008 | -0.0576 | No |
| 59 | GLRX5 | glutaredoxin 5 | 8411 | 0.007 | -0.0634 | No |
| 60 | CCDC28A | coiled-coil domain containing 28A | 8621 | 0.006 | -0.0735 | No |
| 61 | BNIP3L | BCL2/adenovirus E1B 19kDa interacting protein 3-like | 8648 | 0.006 | -0.0742 | No |
| 62 | NNT | nicotinamide nucleotide transhydrogenase | 8653 | 0.006 | -0.0737 | No |
| 63 | PGLS | 6-phosphogluconolactonase | 8706 | 0.006 | -0.0758 | No |
| 64 | NFE2 | nuclear factor, erythroid 2 | 8759 | 0.005 | -0.0778 | No |
| 65 | BTRC | beta-transducin repeat containing E3 ubiquitin protein ligase | 8777 | 0.005 | -0.0781 | No |
| 66 | AQP3 | aquaporin 3 (Gill blood group) | 8812 | 0.005 | -0.0793 | No |
| 67 | EPB42 | erythrocyte membrane protein band 4.2 | 8929 | 0.004 | -0.0848 | No |
| 68 | TFRC | transferrin receptor | 8948 | 0.004 | -0.0852 | No |
| 69 | HEBP1 | heme binding protein 1 | 9017 | 0.004 | -0.0883 | No |
| 70 | ADD1 | adducin 1 (alpha) | 9577 | 0.001 | -0.1171 | No |
| 71 | MARK3 | MAP/microtubule affinity-regulating kinase 3 | 9629 | 0.001 | -0.1196 | No |
| 72 | AGPAT4 | 1-acylglycerol-3-phosphate O-acyltransferase 4 | 9632 | 0.001 | -0.1196 | No |
| 73 | GYPA | glycophorin A (MNS blood group) | 9711 | 0.000 | -0.1236 | No |
| 74 | EPOR | erythropoietin receptor | 9763 | 0.000 | -0.1263 | No |
| 75 | GYPE | glycophorin E (MNS blood group) | 9768 | 0.000 | -0.1264 | No |
| 76 | NEK7 | NIMA-related kinase 7 | 9818 | -0.000 | -0.1290 | No |
| 77 | ALAS2 | 5-aminolevulinate synthase 2 | 9835 | -0.000 | -0.1297 | No |
| 78 | NARF | nuclear prelamin A recognition factor | 9878 | -0.001 | -0.1319 | No |
| 79 | HDGF | hepatoma-derived growth factor | 10182 | -0.002 | -0.1473 | No |
| 80 | HAGH | hydroxyacylglutathione hydrolase | 10193 | -0.002 | -0.1476 | No |
| 81 | MAP2K3 | mitogen-activated protein kinase kinase 3 | 10451 | -0.004 | -0.1604 | No |
| 82 | ADIPOR1 | adiponectin receptor 1 | 10576 | -0.005 | -0.1663 | No |
| 83 | SLC6A8 | solute carrier family 6 (neurotransmitter transporter), member 8 | 10735 | -0.006 | -0.1739 | No |
| 84 | ALDH1L1 | aldehyde dehydrogenase 1 family, member L1 | 10749 | -0.006 | -0.1739 | No |
| 85 | VEZF1 | vascular endothelial zinc finger 1 | 11172 | -0.008 | -0.1948 | No |
| 86 | SLC2A1 | solute carrier family 2 (facilitated glucose transporter), member 1 | 11178 | -0.008 | -0.1941 | No |
| 87 | TFDP2 | transcription factor Dp-2 (E2F dimerization partner 2) | 11180 | -0.008 | -0.1932 | No |
| 88 | TRIM10 | tripartite motif containing 10 | 11282 | -0.009 | -0.1975 | No |
| 89 | EPB41 | erythrocyte membrane protein band 4.1 | 11320 | -0.009 | -0.1984 | No |
| 90 | KLF1 | Kruppel-like factor 1 (erythroid) | 11528 | -0.010 | -0.2080 | No |
| 91 | MPP1 | membrane protein, palmitoylated 1 | 11671 | -0.011 | -0.2141 | No |
| 92 | FECH | ferrochelatase | 11686 | -0.011 | -0.2136 | No |
| 93 | DCAF10 | DDB1 and CUL4 associated factor 10 | 11743 | -0.011 | -0.2152 | No |
| 94 | DCAF11 | DDB1 and CUL4 associated factor 11 | 11875 | -0.012 | -0.2206 | No |
| 95 | TNRC6B | trinucleotide repeat containing 6B | 11883 | -0.012 | -0.2196 | No |
| 96 | MFHAS1 | malignant fibrous histiocytoma amplified sequence 1 | 12203 | -0.014 | -0.2345 | No |
| 97 | SMOX | spermine oxidase | 12281 | -0.015 | -0.2368 | No |
| 98 | PSMD9 | proteasome 26S subunit, non-ATPase 9 | 12336 | -0.015 | -0.2379 | No |
| 99 | SLC25A38 | solute carrier family 25, member 38 | 12370 | -0.015 | -0.2379 | No |
| 100 | SIDT2 | SID1 transmembrane family, member 2 | 12374 | -0.015 | -0.2363 | No |
| 101 | CAT | catalase | 12431 | -0.016 | -0.2374 | No |
| 102 | KLF3 | Kruppel-like factor 3 (basic) | 12507 | -0.016 | -0.2395 | No |
| 103 | FBXO9 | F-box protein 9 | 12534 | -0.016 | -0.2390 | No |
| 104 | FOXJ2 | forkhead box J2 | 12862 | -0.018 | -0.2538 | No |
| 105 | XPO7 | exportin 7 | 12908 | -0.019 | -0.2540 | No |
| 106 | BCAM | basal cell adhesion molecule (Lutheran blood group) | 12938 | -0.019 | -0.2534 | No |
| 107 | MINPP1 | multiple inositol-polyphosphate phosphatase 1 | 12942 | -0.019 | -0.2514 | No |
| 108 | CDC27 | cell division cycle 27 | 12969 | -0.019 | -0.2506 | No |
| 109 | ATP6V0A1 | ATPase, H+ transporting, lysosomal V0 subunit a1 | 12972 | -0.019 | -0.2485 | No |
| 110 | MGST3 | microsomal glutathione S-transferase 3 | 13062 | -0.020 | -0.2509 | No |
| 111 | RHAG | Rh-associated glycoprotein | 13123 | -0.020 | -0.2517 | No |
| 112 | GYPB | glycophorin B (MNS blood group) | 13166 | -0.020 | -0.2516 | No |
| 113 | FTCD | formimidoyltransferase cyclodeaminase | 13185 | -0.020 | -0.2502 | No |
| 114 | ABCG2 | ATP binding cassette subfamily G member 2 (Junior blood group) | 13213 | -0.021 | -0.2493 | No |
| 115 | NCOA4 | nuclear receptor coactivator 4 | 13253 | -0.021 | -0.2489 | No |
| 116 | ENDOD1 | endonuclease domain containing 1 | 13399 | -0.022 | -0.2539 | No |
| 117 | RAD23A | RAD23 homolog A, nucleotide excision repair protein | 13485 | -0.022 | -0.2558 | No |
| 118 | OPTN | optineurin | 13509 | -0.023 | -0.2544 | No |
| 119 | EIF2AK1 | eukaryotic translation initiation factor 2-alpha kinase 1 | 13648 | -0.024 | -0.2589 | No |
| 120 | NUDT4 | nudix hydrolase 4 | 13650 | -0.024 | -0.2562 | No |
| 121 | SEC14L1 | SEC14-like lipid binding 1 | 13926 | -0.026 | -0.2675 | No |
| 122 | MYL4 | myosin light chain 4 | 13996 | -0.026 | -0.2681 | No |
| 123 | GAPVD1 | GTPase activating protein and VPS9 domains 1 | 14125 | -0.027 | -0.2716 | No |
| 124 | ANK1 | ankyrin 1, erythrocytic | 14162 | -0.028 | -0.2704 | No |
| 125 | RBM38 | RNA binding motif protein 38 | 14196 | -0.028 | -0.2689 | No |
| 126 | SPTA1 | spectrin, alpha, erythrocytic 1 | 14299 | -0.029 | -0.2709 | No |
| 127 | BMP2K | BMP2 inducible kinase | 14375 | -0.029 | -0.2715 | No |
| 128 | FBXO7 | F-box protein 7 | 14468 | -0.030 | -0.2728 | No |
| 129 | RIOK3 | RIO kinase 3 | 14596 | -0.031 | -0.2759 | No |
| 130 | LMO2 | LIM domain only 2 (rhombotin-like 1) | 14614 | -0.031 | -0.2732 | No |
| 131 | GCLM | glutamate-cysteine ligase, modifier subunit | 14668 | -0.032 | -0.2723 | No |
| 132 | CTSB | cathepsin B | 14720 | -0.032 | -0.2713 | No |
| 133 | HBB | hemoglobin, beta | 14794 | -0.033 | -0.2714 | No |
| 134 | TAL1 | T-cell acute lymphocytic leukemia 1 | 14871 | -0.033 | -0.2716 | No |
| 135 | CCND3 | cyclin D3 | 14958 | -0.034 | -0.2721 | No |
| 136 | BSG | basigin (Ok blood group) | 14969 | -0.034 | -0.2688 | No |
| 137 | PICALM | phosphatidylinositol binding clathrin assembly protein | 15258 | -0.037 | -0.2795 | Yes |
| 138 | DCUN1D1 | DCN1, defective in cullin neddylation 1, domain containing 1 | 15291 | -0.037 | -0.2770 | Yes |
| 139 | USP15 | ubiquitin specific peptidase 15 | 15307 | -0.037 | -0.2736 | Yes |
| 140 | TRAK2 | trafficking protein, kinesin binding 2 | 15368 | -0.038 | -0.2724 | Yes |
| 141 | SPTB | spectrin, beta, erythrocytic | 15433 | -0.038 | -0.2714 | Yes |
| 142 | BPGM | 2,3-bisphosphoglycerate mutase | 15645 | -0.040 | -0.2777 | Yes |
| 143 | TRIM58 | tripartite motif containing 58 | 15668 | -0.041 | -0.2743 | Yes |
| 144 | BACH1 | BTB and CNC homology 1, basic leucine zipper transcription factor 1 | 15744 | -0.041 | -0.2735 | Yes |
| 145 | PPP2R5B | protein phosphatase 2, regulatory subunit B, beta | 15844 | -0.042 | -0.2738 | Yes |
| 146 | RNF123 | ring finger protein 123 | 15951 | -0.043 | -0.2744 | Yes |
| 147 | P4HA2 | prolyl 4-hydroxylase, alpha polypeptide II | 15996 | -0.044 | -0.2717 | Yes |
| 148 | CIR1 | corepressor interacting with RBPJ, 1 | 16011 | -0.044 | -0.2675 | Yes |
| 149 | KHNYN | KH and NYN domain containing | 16042 | -0.044 | -0.2640 | Yes |
| 150 | SDCBP | syndecan binding protein | 16135 | -0.045 | -0.2636 | Yes |
| 151 | ADD2 | adducin 2 (beta) | 16286 | -0.047 | -0.2661 | Yes |
| 152 | DAAM1 | dishevelled associated activator of morphogenesis 1 | 16418 | -0.049 | -0.2673 | Yes |
| 153 | MOCOS | molybdenum cofactor sulfurase | 16430 | -0.049 | -0.2624 | Yes |
| 154 | CAST | calpastatin | 16477 | -0.050 | -0.2591 | Yes |
| 155 | SYNJ1 | synaptojanin 1 | 16562 | -0.051 | -0.2577 | Yes |
| 156 | MKRN1 | makorin ring finger protein 1 | 16563 | -0.051 | -0.2520 | Yes |
| 157 | GDE1 | glycerophosphodiester phosphodiesterase 1 | 16687 | -0.052 | -0.2524 | Yes |
| 158 | TCEA1 | transcription elongation factor A (SII), 1 | 16811 | -0.054 | -0.2526 | Yes |
| 159 | KDM7A | lysine (K)-specific demethylase 7A | 16812 | -0.054 | -0.2465 | Yes |
| 160 | MOSPD1 | motile sperm domain containing 1 | 16835 | -0.054 | -0.2415 | Yes |
| 161 | CLCN3 | chloride channel, voltage-sensitive 3 | 17046 | -0.058 | -0.2458 | Yes |
| 162 | YPEL5 | yippee like 5 | 17113 | -0.059 | -0.2426 | Yes |
| 163 | LPIN2 | lipin 2 | 17229 | -0.060 | -0.2417 | Yes |
| 164 | ARHGEF12 | Rho guanine nucleotide exchange factor (GEF) 12 | 17283 | -0.061 | -0.2374 | Yes |
| 165 | NFE2L1 | nuclear factor, erythroid 2-like 1 | 17306 | -0.062 | -0.2316 | Yes |
| 166 | SLC30A1 | solute carrier family 30 (zinc transporter), member 1 | 17588 | -0.067 | -0.2385 | Yes |
| 167 | IGSF3 | immunoglobulin superfamily, member 3 | 17610 | -0.068 | -0.2319 | Yes |
| 168 | LAMP2 | lysosomal-associated membrane protein 2 | 17747 | -0.071 | -0.2309 | Yes |
| 169 | ALAD | aminolevulinate dehydratase | 17796 | -0.072 | -0.2253 | Yes |
| 170 | TSPO2 | translocator protein 2 | 17968 | -0.076 | -0.2255 | Yes |
| 171 | GCLC | glutamate-cysteine ligase, catalytic subunit | 17981 | -0.077 | -0.2174 | Yes |
| 172 | TSPAN5 | tetraspanin 5 | 18033 | -0.078 | -0.2112 | Yes |
| 173 | RANBP10 | RAN binding protein 10 | 18034 | -0.078 | -0.2024 | Yes |
| 174 | HTATIP2 | HIV-1 Tat interactive protein 2 | 18156 | -0.081 | -0.1994 | Yes |
| 175 | TMEM9B | TMEM9 domain family, member B | 18296 | -0.085 | -0.1970 | Yes |
| 176 | SLC6A9 | solute carrier family 6 (neurotransmitter transporter, glycine), member 9 | 18355 | -0.087 | -0.1901 | Yes |
| 177 | UCP2 | uncoupling protein 2 (mitochondrial, proton carrier) | 18548 | -0.094 | -0.1893 | Yes |
| 178 | ARL2BP | ADP-ribosylation factor like GTPase 2 binding protein | 18580 | -0.096 | -0.1801 | Yes |
| 179 | ALDH6A1 | aldehyde dehydrogenase 6 family, member A1 | 18640 | -0.099 | -0.1720 | Yes |
| 180 | SLC7A11 | solute carrier family 7 (anionic amino acid transporter light chain, xc- system), member 11 | 18650 | -0.099 | -0.1612 | Yes |
| 181 | HBD | hemoglobin, delta | 18786 | -0.108 | -0.1559 | Yes |
| 182 | MBOAT2 | Memczak2013 ALT\_ACCEPTOR, ALT\_DONOR, coding, INTERNAL, intronic best transcript NM\_138799 | 18819 | -0.111 | -0.1449 | Yes |
| 183 | ASNS | asparagine synthetase (glutamine-hydrolyzing) | 18864 | -0.115 | -0.1342 | Yes |
| 184 | RNF19A | ring finger protein 19A, RBR E3 ubiquitin protein ligase | 18982 | -0.125 | -0.1260 | Yes |
| 185 | KEL | Kell blood group, metallo-endopeptidase | 19048 | -0.133 | -0.1143 | Yes |
| 186 | ERMAP | erythroblast membrane-associated protein (Scianna blood group) | 19160 | -0.151 | -0.1029 | Yes |
| 187 | ATG4A | autophagy related 4A, cysteine peptidase | 19164 | -0.152 | -0.0858 | Yes |
| 188 | RAP1GAP | RAP1 GTPase activating protein | 19313 | -0.196 | -0.0712 | Yes |
| 189 | TYR | tyrosinase | 19357 | -0.216 | -0.0489 | Yes |
| 190 | CA1 | carbonic anhydrase I | 19390 | -0.233 | -0.0241 | Yes |
| 191 | CLIC2 | chloride intracellular channel 2 | 19436 | -0.273 | 0.0046 | Yes |
Table: GSEA details [plain text format]

  

Fig 2: HALLMARK\_HEME\_METABOLISM      
 Blue-Pink O' Gram in the Space of the Analyzed GeneSet

  

Fig 3: HALLMARK\_HEME\_METABOLISM: Random ES distribution      
 Gene set null distribution of ES for **HALLMARK\_HEME\_METABOLISM**

  
