## Supplementary material for "Somatic hypomethylation of pericentromeric SST1 repeats and tetraploidization in human colorectal cancer cells": GSEA results: HALLMARK_IL2_STAT5_SIGNALING.html

Details for gene set HALLMARK\_IL2\_STAT5\_SIGNALING[GSEA]

|  || Dataset | eset\_byprobe\_collapsed\_to\_symbols.Diploid\_vs\_Tetraploid.cls #Tetraploid\_versus\_Diploid.Diploid\_vs\_Tetraploid.cls #Tetraploid\_versus\_Diploid\_repos |
| Phenotype | Diploid\_vs\_Tetraploid.cls#Tetraploid\_versus\_Diploid\_repos |
| Upregulated in class | Tetraploid |
| GeneSet | HALLMARK\_IL2\_STAT5\_SIGNALING |
| Enrichment Score (ES) | 0.27802828 |
| Normalized Enrichment Score (NES) | 1.2449241 |
| Nominal p-value | 0.06849315 |
| FDR q-value | 0.19353276 |
| FWER p-Value | 0.809 |
Table: GSEA Results Summary

  

Fig 1: Enrichment plot: HALLMARK\_IL2\_STAT5\_SIGNALING      
 Profile of the Running ES Score & Positions of GeneSet Members on the Rank Ordered List

  

| SYMBOL | TITLE | RANK IN GENE LIST | RANK METRIC SCORE | RUNNING ES | CORE ENRICHMENT || 1 | CCND2 | cyclin D2 | 34 | 0.313 | 0.0275 | Yes |
| 2 | NT5E | 5-nucleotidase, ecto (CD73) | 37 | 0.307 | 0.0561 | Yes |
| 3 | RRAGD | Ras-related GTP binding D | 54 | 0.280 | 0.0814 | Yes |
| 4 | CA2 | carbonic anhydrase II | 69 | 0.250 | 0.1040 | Yes |
| 5 | ST3GAL4 | ST3 beta-galactoside alpha-2,3-sialyltransferase 4 | 104 | 0.207 | 0.1216 | Yes |
| 6 | AGER | advanced glycosylation end product-specific receptor | 219 | 0.150 | 0.1297 | Yes |
| 7 | GSTO1 | glutathione S-transferase omega 1 | 280 | 0.136 | 0.1393 | Yes |
| 8 | MYO1E | myosin IE | 330 | 0.128 | 0.1488 | Yes |
| 9 | PDCD2L | programmed cell death 2-like | 421 | 0.117 | 0.1550 | Yes |
| 10 | LIF | leukemia inhibitory factor | 423 | 0.117 | 0.1659 | Yes |
| 11 | IRF8 | interferon regulatory factor 8 | 434 | 0.115 | 0.1761 | Yes |
| 12 | CD44 | CD44 molecule (Indian blood group) | 577 | 0.104 | 0.1786 | Yes |
| 13 | GALM | galactose mutarotase (aldose 1-epimerase) | 674 | 0.098 | 0.1828 | Yes |
| 14 | TGM2 | transglutaminase 2 | 714 | 0.096 | 0.1897 | Yes |
| 15 | PTGER2 | prostaglandin E receptor 2 | 737 | 0.095 | 0.1975 | Yes |
| 16 | CCNE1 | cyclin E1 | 738 | 0.095 | 0.2064 | Yes |
| 17 | PHLDA1 | pleckstrin homology-like domain, family A, member 1 | 740 | 0.095 | 0.2152 | Yes |
| 18 | ALCAM | activated leukocyte cell adhesion molecule | 762 | 0.094 | 0.2229 | Yes |
| 19 | AHNAK | AHNAK nucleoprotein | 775 | 0.093 | 0.2310 | Yes |
| 20 | DRC1 | dynein regulatory complex subunit 1 | 932 | 0.087 | 0.2310 | Yes |
| 21 | EMP1 | epithelial membrane protein 1 | 967 | 0.085 | 0.2373 | Yes |
| 22 | TNFSF10 | tumor necrosis factor (ligand) superfamily, member 10 | 1006 | 0.084 | 0.2432 | Yes |
| 23 | IL1R2 | interleukin 1 receptor, type II | 1012 | 0.084 | 0.2507 | Yes |
| 24 | SLC29A2 | solute carrier family 29 (equilibrative nucleoside transporter), member 2 | 1017 | 0.084 | 0.2584 | Yes |
| 25 | CSF1 | colony stimulating factor 1 (macrophage) | 1040 | 0.083 | 0.2650 | Yes |
| 26 | IL4R | interleukin 4 receptor | 1066 | 0.082 | 0.2713 | Yes |
| 27 | NRP1 | neuropilin 1 | 1363 | 0.075 | 0.2630 | Yes |
| 28 | BATF | basic leucine zipper transcription factor, ATF-like | 1397 | 0.074 | 0.2682 | Yes |
| 29 | PNP | purine nucleoside phosphorylase | 1405 | 0.074 | 0.2748 | Yes |
| 30 | SHE | Src homology 2 domain containing E | 1566 | 0.070 | 0.2730 | Yes |
| 31 | MAP6 | microtubule associated protein 6 | 1598 | 0.069 | 0.2778 | Yes |
| 32 | SPRY4 | sprouty RTK signaling antagonist 4 | 1732 | 0.066 | 0.2771 | Yes |
| 33 | UCK2 | Jeck2013 ALT\_ACCEPTOR, ALT\_DONOR, coding, INTERNAL, intronic best transcript NM\_012474 | 1906 | 0.063 | 0.2741 | Yes |
| 34 | FAH | fumarylacetoacetate hydrolase (fumarylacetoacetase) | 2119 | 0.060 | 0.2687 | Yes |
| 35 | CDC6 | cell division cycle 6 | 2205 | 0.058 | 0.2697 | Yes |
| 36 | TNFRSF8 | tumor necrosis factor receptor superfamily, member 8 | 2322 | 0.056 | 0.2690 | Yes |
| 37 | TIAM1 | T-cell lymphoma invasion and metastasis 1 | 2375 | 0.056 | 0.2715 | Yes |
| 38 | POU2F1 | POU class 2 homeobox 1 | 2465 | 0.054 | 0.2720 | Yes |
| 39 | BCL2L1 | BCL2-like 1 | 2466 | 0.054 | 0.2771 | Yes |
| 40 | ITGAE | integrin alpha E | 2544 | 0.053 | 0.2780 | Yes |
| 41 | UMPS | uridine monophosphate synthetase | 2899 | 0.049 | 0.2643 | No |
| 42 | CCR4 | chemokine (C-C motif) receptor 4 | 2907 | 0.048 | 0.2684 | No |
| 43 | ADAM19 | ADAM metallopeptidase domain 19 | 2980 | 0.048 | 0.2692 | No |
| 44 | MAFF | v-maf avian musculoaponeurotic fibrosarcoma oncogene homolog F | 3075 | 0.047 | 0.2687 | No |
| 45 | MYC | v-myc avian myelocytomatosis viral oncogene homolog | 3194 | 0.046 | 0.2668 | No |
| 46 | IL10RA | interleukin 10 receptor, alpha | 3223 | 0.045 | 0.2696 | No |
| 47 | CD81 | CD81 molecule | 3320 | 0.044 | 0.2688 | No |
| 48 | PUS1 | pseudouridylate synthase 1 | 3333 | 0.044 | 0.2723 | No |
| 49 | COCH | cochlin | 3477 | 0.043 | 0.2689 | No |
| 50 | GATA1 | GATA binding protein 1 (globin transcription factor 1) | 3535 | 0.042 | 0.2699 | No |
| 51 | FLT3LG | fms-related tyrosine kinase 3 ligand | 3591 | 0.042 | 0.2710 | No |
| 52 | WLS | wntless Wnt ligand secretion mediator | 3854 | 0.039 | 0.2611 | No |
| 53 | DENND5A | DENN/MADD domain containing 5A | 4458 | 0.033 | 0.2330 | No |
| 54 | LTB | lymphotoxin beta (TNF superfamily, member 3) | 4496 | 0.033 | 0.2342 | No |
| 55 | F2RL2 | coagulation factor II (thrombin) receptor-like 2 | 4564 | 0.033 | 0.2338 | No |
| 56 | CASP3 | caspase 3 | 4699 | 0.032 | 0.2298 | No |
| 57 | NCS1 | neuronal calcium sensor 1 | 4902 | 0.030 | 0.2222 | No |
| 58 | TRAF1 | TNF receptor-associated factor 1 | 5030 | 0.029 | 0.2183 | No |
| 59 | GPR65 | G protein-coupled receptor 65 | 5142 | 0.028 | 0.2152 | No |
| 60 | SCN9A | sodium channel, voltage gated, type IX alpha subunit | 5147 | 0.028 | 0.2177 | No |
| 61 | CAPG | capping protein (actin filament), gelsolin-like | 5399 | 0.026 | 0.2071 | No |
| 62 | SWAP70 | SWAP switching B-cell complex 70kDa subunit | 5447 | 0.026 | 0.2071 | No |
| 63 | ITGA6 | integrin alpha 6 | 5546 | 0.025 | 0.2044 | No |
| 64 | P2RX4 | purinergic receptor P2X, ligand gated ion channel, 4 | 5563 | 0.025 | 0.2059 | No |
| 65 | GADD45B | growth arrest and DNA-damage-inducible, beta | 5568 | 0.025 | 0.2080 | No |
| 66 | CD86 | CD86 molecule | 5650 | 0.024 | 0.2061 | No |
| 67 | PLSCR1 | phospholipid scramblase 1 | 5829 | 0.023 | 0.1991 | No |
| 68 | IL10 | interleukin 10 | 6071 | 0.022 | 0.1886 | No |
| 69 | GPX4 | glutathione peroxidase 4 | 6109 | 0.021 | 0.1887 | No |
| 70 | NOP2 | NOP2 nucleolar protein | 6188 | 0.021 | 0.1866 | No |
| 71 | IL18R1 | interleukin 18 receptor 1 | 6256 | 0.021 | 0.1851 | No |
| 72 | PTRH2 | peptidyl-tRNA hydrolase 2 | 6300 | 0.020 | 0.1847 | No |
| 73 | EOMES | eomesodermin | 6303 | 0.020 | 0.1865 | No |
| 74 | CYFIP1 | cytoplasmic FMR1 interacting protein 1 | 6348 | 0.020 | 0.1861 | No |
| 75 | PLPP1 | phospholipid phosphatase 1 | 6641 | 0.018 | 0.1727 | No |
| 76 | ICOS | inducible T-cell co-stimulator | 6767 | 0.017 | 0.1678 | No |
| 77 | SMPDL3A | sphingomyelin phosphodiesterase, acid-like 3A | 6861 | 0.017 | 0.1646 | No |
| 78 | SYNGR2 | synaptogyrin 2 | 6958 | 0.016 | 0.1611 | No |
| 79 | PHTF2 | putative homeodomain transcription factor 2 | 6984 | 0.016 | 0.1613 | No |
| 80 | BMPR2 | bone morphogenetic protein receptor type II | 6988 | 0.016 | 0.1626 | No |
| 81 | IRF4 | interferon regulatory factor 4 | 7069 | 0.015 | 0.1600 | No |
| 82 | IL1RL1 | interleukin 1 receptor-like 1 | 7132 | 0.015 | 0.1582 | No |
| 83 | PLEC | plectin | 7170 | 0.015 | 0.1576 | No |
| 84 | CDC42SE2 | CDC42 small effector 2 | 7498 | 0.013 | 0.1419 | No |
| 85 | IFITM3 | interferon induced transmembrane protein 3 | 7579 | 0.012 | 0.1389 | No |
| 86 | CSF2 | colony stimulating factor 2 (granulocyte-macrophage) | 7825 | 0.011 | 0.1273 | No |
| 87 | ECM1 | extracellular matrix protein 1 | 7942 | 0.010 | 0.1222 | No |
| 88 | NFKBIZ | nuclear factor of kappa light polypeptide gene enhancer in B-cells inhibitor, zeta | 7959 | 0.010 | 0.1224 | No |
| 89 | PENK | proenkephalin | 8055 | 0.010 | 0.1183 | No |
| 90 | CXCL10 | chemokine (C-X-C motif) ligand 10 | 8286 | 0.008 | 0.1072 | No |
| 91 | IFNGR1 | interferon gamma receptor 1 | 8413 | 0.007 | 0.1014 | No |
| 92 | IKZF4 | IKAROS family zinc finger 4 | 8439 | 0.007 | 0.1008 | No |
| 93 | S100A1 | S100 calcium binding protein A1 | 8573 | 0.006 | 0.0945 | No |
| 94 | HK2 | hexokinase 2 | 8575 | 0.006 | 0.0950 | No |
| 95 | MYO1C | myosin IC | 8827 | 0.005 | 0.0825 | No |
| 96 | AHCY | adenosylhomocysteinase | 8841 | 0.005 | 0.0823 | No |
| 97 | TWSG1 | twisted gastrulation BMP signaling modulator 1 | 8844 | 0.005 | 0.0827 | No |
| 98 | TLR7 | toll-like receptor 7 | 8896 | 0.005 | 0.0805 | No |
| 99 | HOPX | HOP homeobox | 8924 | 0.004 | 0.0795 | No |
| 100 | SNX14 | sorting nexin 14 | 9002 | 0.004 | 0.0759 | No |
| 101 | RHOH | ras homolog family member H | 9007 | 0.004 | 0.0761 | No |
| 102 | ANXA4 | Salzman2013 ANTISENSE, coding, INTERNAL, intronic best transcript NM\_001153 | 9010 | 0.004 | 0.0763 | No |
| 103 | PLIN2 | perilipin 2 | 9014 | 0.004 | 0.0765 | No |
| 104 | CD79B | CD79b molecule, immunoglobulin-associated beta | 9020 | 0.004 | 0.0766 | No |
| 105 | NDRG1 | N-myc downstream regulated 1 | 9078 | 0.004 | 0.0740 | No |
| 106 | DCPS | decapping enzyme, scavenger | 9185 | 0.003 | 0.0688 | No |
| 107 | IL13 | interleukin 13 | 9330 | 0.002 | 0.0616 | No |
| 108 | CTLA4 | cytotoxic T-lymphocyte-associated protein 4 | 9842 | -0.000 | 0.0352 | No |
| 109 | TNFRSF1B | tumor necrosis factor receptor superfamily, member 1B | 9919 | -0.001 | 0.0313 | No |
| 110 | SYT11 | synaptotagmin XI | 9931 | -0.001 | 0.0308 | No |
| 111 | IL2RB | interleukin 2 receptor, beta | 10056 | -0.001 | 0.0246 | No |
| 112 | TNFRSF18 | tumor necrosis factor receptor superfamily, member 18 | 10199 | -0.002 | 0.0174 | No |
| 113 | SELP | selectin P (granule membrane protein 140kDa, antigen CD62) | 10228 | -0.002 | 0.0162 | No |
| 114 | ENO3 | enolase 3 (beta, muscle) | 10237 | -0.003 | 0.0160 | No |
| 115 | PTCH1 | patched 1 | 10420 | -0.004 | 0.0070 | No |
| 116 | RGS16 | regulator of G-protein signaling 16 | 11032 | -0.007 | -0.0240 | No |
| 117 | CDKN1C | cyclin-dependent kinase inhibitor 1C (p57, Kip2) | 11246 | -0.008 | -0.0342 | No |
| 118 | IKZF2 | IKAROS family zinc finger 2 | 11363 | -0.009 | -0.0394 | No |
| 119 | PIM1 | Pim-1 proto-oncogene, serine/threonine kinase | 11480 | -0.010 | -0.0444 | No |
| 120 | SNX9 | sorting nexin 9 | 11520 | -0.010 | -0.0455 | No |
| 121 | SPP1 | secreted phosphoprotein 1 | 11575 | -0.010 | -0.0474 | No |
| 122 | CTSZ | cathepsin Z | 11627 | -0.011 | -0.0490 | No |
| 123 | DHRS3 | dehydrogenase/reductase (SDR family) member 3 | 11753 | -0.011 | -0.0544 | No |
| 124 | LCLAT1 | lysocardiolipin acyltransferase 1 | 11870 | -0.012 | -0.0593 | No |
| 125 | RNH1 | ribonuclease/angiogenin inhibitor 1 | 11959 | -0.013 | -0.0627 | No |
| 126 | MUC1 | mucin 1, cell surface associated | 11994 | -0.013 | -0.0632 | No |
| 127 | ITIH5 | inter-alpha-trypsin inhibitor heavy chain family, member 5 | 12048 | -0.013 | -0.0648 | No |
| 128 | GLIPR2 | GLI pathogenesis-related 2 | 12366 | -0.015 | -0.0797 | No |
| 129 | APLP1 | amyloid beta (A4) precursor-like protein 1 | 12392 | -0.015 | -0.0796 | No |
| 130 | SELL | selectin L | 12417 | -0.016 | -0.0794 | No |
| 131 | SOCS1 | suppressor of cytokine signaling 1 | 12611 | -0.017 | -0.0878 | No |
| 132 | IRF6 | interferon regulatory factor 6 | 12694 | -0.017 | -0.0904 | No |
| 133 | CD48 | CD48 molecule | 12879 | -0.018 | -0.0982 | No |
| 134 | KLF6 | Kruppel-like factor 6 | 12945 | -0.019 | -0.0998 | No |
| 135 | CDCP1 | CUB domain containing protein 1 | 12988 | -0.019 | -0.1002 | No |
| 136 | CAPN3 | calpain 3 | 13021 | -0.019 | -0.1000 | No |
| 137 | AHR | aryl hydrocarbon receptor | 13095 | -0.020 | -0.1020 | No |
| 138 | ODC1 | ornithine decarboxylase 1 | 13133 | -0.020 | -0.1020 | No |
| 139 | IGF1R | insulin-like growth factor 1 receptor | 13395 | -0.022 | -0.1135 | No |
| 140 | ENPP1 | ectonucleotide pyrophosphatase/phosphodiesterase 1 | 13430 | -0.022 | -0.1132 | No |
| 141 | IL3RA | interleukin 3 receptor, alpha (low affinity) | 13504 | -0.023 | -0.1148 | No |
| 142 | P4HA1 | prolyl 4-hydroxylase, alpha polypeptide I | 13604 | -0.023 | -0.1178 | No |
| 143 | FAM126B | family with sequence similarity 126, member B | 14215 | -0.028 | -0.1467 | No |
| 144 | GBP4 | guanylate binding protein 4 | 14250 | -0.028 | -0.1458 | No |
| 145 | GPR83 | G protein-coupled receptor 83 | 14328 | -0.029 | -0.1471 | No |
| 146 | BCL2 | B-cell CLL/lymphoma 2 | 14503 | -0.030 | -0.1533 | No |
| 147 | TTC39B | Transcript Identified by AceView, Entrez Gene ID(s) 158219 | 14687 | -0.032 | -0.1597 | No |
| 148 | NCOA3 | nuclear receptor coactivator 3 | 14803 | -0.033 | -0.1626 | No |
| 149 | CCND3 | cyclin D3 | 14958 | -0.034 | -0.1674 | No |
| 150 | BATF3 | basic leucine zipper transcription factor, ATF-like 3 | 15029 | -0.035 | -0.1678 | No |
| 151 | MAP3K8 | mitogen-activated protein kinase kinase kinase 8 | 15311 | -0.037 | -0.1789 | No |
| 152 | COL6A1 | collagen, type VI, alpha 1 | 15384 | -0.038 | -0.1790 | No |
| 153 | ETV4 | ets variant 4 | 15403 | -0.038 | -0.1764 | No |
| 154 | HUWE1 | HECT, UBA and WWE domain containing 1, E3 ubiquitin protein ligase | 15418 | -0.038 | -0.1736 | No |
| 155 | HIPK2 | homeodomain interacting protein kinase 2 | 15466 | -0.038 | -0.1724 | No |
| 156 | XBP1 | X-box binding protein 1 | 15622 | -0.040 | -0.1767 | No |
| 157 | CISH | cytokine inducible SH2-containing protein | 15667 | -0.040 | -0.1752 | No |
| 158 | CD83 | CD83 molecule | 15767 | -0.041 | -0.1765 | No |
| 159 | MAPKAPK2 | mitogen-activated protein kinase-activated protein kinase 2 | 15801 | -0.042 | -0.1743 | No |
| 160 | TNFRSF21 | tumor necrosis factor receptor superfamily, member 21 | 15802 | -0.042 | -0.1704 | No |
| 161 | TNFRSF9 | tumor necrosis factor receptor superfamily, member 9 | 15868 | -0.042 | -0.1698 | No |
| 162 | TNFSF11 | tumor necrosis factor (ligand) superfamily, member 11 | 16018 | -0.044 | -0.1734 | No |
| 163 | SLC2A3 | solute carrier family 2 (facilitated glucose transporter), member 3 | 16073 | -0.045 | -0.1720 | No |
| 164 | ITGAV | integrin alpha V | 16105 | -0.045 | -0.1694 | No |
| 165 | PRKCH | protein kinase C, eta | 16154 | -0.046 | -0.1676 | No |
| 166 | LRRC8C | leucine rich repeat containing 8 family, member C | 16228 | -0.046 | -0.1671 | No |
| 167 | IL2RA | interleukin 2 receptor, alpha | 16231 | -0.046 | -0.1628 | No |
| 168 | BMP2 | bone morphogenetic protein 2 | 16350 | -0.048 | -0.1645 | No |
| 169 | SERPINC1 | serpin peptidase inhibitor, clade C (antithrombin), member 1 | 16408 | -0.048 | -0.1629 | No |
| 170 | AMACR | alpha-methylacyl-CoA racemase | 16704 | -0.053 | -0.1733 | No |
| 171 | CST7 | cystatin F (leukocystatin) | 17019 | -0.057 | -0.1842 | No |
| 172 | CKAP4 | cytoskeleton-associated protein 4 | 17085 | -0.058 | -0.1821 | No |
| 173 | SPRED2 | sprouty-related, EVH1 domain containing 2 | 17266 | -0.061 | -0.1857 | No |
| 174 | RABGAP1L | RAB GTPase activating protein 1-like | 17616 | -0.068 | -0.1974 | No |
| 175 | IGF2R | insulin-like growth factor 2 receptor | 17794 | -0.072 | -0.1999 | No |
| 176 | TNFRSF4 | tumor necrosis factor receptor superfamily, member 4 | 17866 | -0.074 | -0.1967 | No |
| 177 | PRNP | prion protein | 17960 | -0.076 | -0.1944 | No |
| 178 | FURIN | furin (paired basic amino acid cleaving enzyme) | 18023 | -0.077 | -0.1903 | No |
| 179 | ABCB1 | ATP binding cassette subfamily B member 1 | 18030 | -0.078 | -0.1834 | No |
| 180 | SH3BGRL2 | Transcript Identified by AceView, Entrez Gene ID(s) 83699 | 18091 | -0.079 | -0.1791 | No |
| 181 | MXD1 | Memczak2013 ANTISENSE, coding, INTERNAL, intronic best transcript NM\_001202514 | 18336 | -0.086 | -0.1836 | No |
| 182 | PTH1R | parathyroid hormone 1 receptor | 18370 | -0.088 | -0.1771 | No |
| 183 | SLC1A5 | solute carrier family 1 (neutral amino acid transporter), member 5 | 18428 | -0.090 | -0.1717 | No |
| 184 | NFIL3 | nuclear factor, interleukin 3 regulated | 18475 | -0.092 | -0.1655 | No |
| 185 | FGL2 | fibrinogen-like 2 | 18506 | -0.093 | -0.1584 | No |
| 186 | RHOB | ras homolog family member B | 18827 | -0.112 | -0.1645 | No |
| 187 | SERPINB6 | serpin peptidase inhibitor, clade B (ovalbumin), member 6 | 19076 | -0.138 | -0.1644 | No |
| 188 | RORA | RAR-related orphan receptor A | 19240 | -0.171 | -0.1569 | No |
| 189 | ARL4A | ADP-ribosylation factor like GTPase 4A | 19243 | -0.172 | -0.1409 | No |
| 190 | GABARAPL1 | GABA(A) receptor-associated protein like 1 | 19244 | -0.172 | -0.1248 | No |
| 191 | BHLHE40 | basic helix-loop-helix family, member e40 | 19252 | -0.175 | -0.1088 | No |
| 192 | LRIG1 | leucine-rich repeats and immunoglobulin-like domains 1 | 19335 | -0.203 | -0.0940 | No |
| 193 | SOCS2 | suppressor of cytokine signaling 2 | 19368 | -0.220 | -0.0751 | No |
| 194 | PLAGL1 | pleiomorphic adenoma gene-like 1 | 19471 | -0.328 | -0.0498 | No |
| 195 | SLC39A8 | solute carrier family 39 (zinc transporter), member 8 | 19523 | -0.562 | 0.0001 | No |
Table: GSEA details [plain text format]

  

Fig 2: HALLMARK\_IL2\_STAT5\_SIGNALING      
 Blue-Pink O' Gram in the Space of the Analyzed GeneSet

  

Fig 3: HALLMARK\_IL2\_STAT5\_SIGNALING: Random ES distribution      
 Gene set null distribution of ES for **HALLMARK\_IL2\_STAT5\_SIGNALING**

  
