## Supplementary material for "Somatic hypomethylation of pericentromeric SST1 repeats and tetraploidization in human colorectal cancer cells": GSEA results: HALLMARK_IL6_JAK_STAT3_SIGNALING.html

Details for gene set HALLMARK\_IL6\_JAK\_STAT3\_SIGNALING[GSEA]

|  || Dataset | eset\_byprobe\_collapsed\_to\_symbols.Diploid\_vs\_Tetraploid.cls #Tetraploid\_versus\_Diploid.Diploid\_vs\_Tetraploid.cls #Tetraploid\_versus\_Diploid\_repos |
| Phenotype | Diploid\_vs\_Tetraploid.cls#Tetraploid\_versus\_Diploid\_repos |
| Upregulated in class | Diploid |
| GeneSet | HALLMARK\_IL6\_JAK\_STAT3\_SIGNALING |
| Enrichment Score (ES) | -0.2723899 |
| Normalized Enrichment Score (NES) | -0.9914638 |
| Nominal p-value | 0.47716534 |
| FDR q-value | 0.5695585 |
| FWER p-Value | 1.0 |
Table: GSEA Results Summary

  

Fig 1: Enrichment plot: HALLMARK\_IL6\_JAK\_STAT3\_SIGNALING      
 Profile of the Running ES Score & Positions of GeneSet Members on the Rank Ordered List

  

| SYMBOL | TITLE | RANK IN GENE LIST | RANK METRIC SCORE | RUNNING ES | CORE ENRICHMENT || 1 | FAS | Fas cell surface death receptor | 186 | 0.161 | 0.0284 | No |
| 2 | IL2RG | interleukin 2 receptor, gamma | 268 | 0.137 | 0.0567 | No |
| 3 | CD44 | CD44 molecule (Indian blood group) | 577 | 0.104 | 0.0655 | No |
| 4 | CXCL1 | chemokine (C-X-C motif) ligand 1 (melanoma growth stimulating activity, alpha) | 672 | 0.098 | 0.0839 | No |
| 5 | CD38 | CD38 molecule | 689 | 0.097 | 0.1061 | No |
| 6 | CNTFR | ciliary neurotrophic factor receptor | 920 | 0.087 | 0.1149 | No |
| 7 | IL1R2 | interleukin 1 receptor, type II | 1012 | 0.084 | 0.1301 | No |
| 8 | CSF1 | colony stimulating factor 1 (macrophage) | 1040 | 0.083 | 0.1483 | No |
| 9 | CXCL3 | chemokine (C-X-C motif) ligand 3 | 1048 | 0.083 | 0.1674 | No |
| 10 | IL4R | interleukin 4 receptor | 1066 | 0.082 | 0.1859 | No |
| 11 | IL7 | interleukin 7 | 1267 | 0.077 | 0.1938 | No |
| 12 | A2M | alpha-2-macroglobulin | 1412 | 0.074 | 0.2038 | No |
| 13 | PDGFC | platelet derived growth factor C | 1516 | 0.071 | 0.2153 | No |
| 14 | IL9R | interleukin 9 receptor | 1543 | 0.070 | 0.2305 | No |
| 15 | ITGA4 | integrin alpha 4 | 2062 | 0.061 | 0.2182 | No |
| 16 | CSF2RB | colony stimulating factor 2 receptor, beta, low-affinity (granulocyte-macrophage) | 2475 | 0.054 | 0.2098 | No |
| 17 | IL15RA | interleukin 15 receptor, alpha | 2659 | 0.051 | 0.2125 | No |
| 18 | IL17RB | interleukin 17 receptor B | 2765 | 0.050 | 0.2190 | No |
| 19 | PF4 | platelet factor 4 | 2778 | 0.050 | 0.2302 | No |
| 20 | PTPN2 | protein tyrosine phosphatase, non-receptor type 2 | 2895 | 0.049 | 0.2357 | No |
| 21 | TNF | tumor necrosis factor | 3210 | 0.045 | 0.2303 | No |
| 22 | HMOX1 | heme oxygenase 1 | 3502 | 0.043 | 0.2254 | No |
| 23 | TYK2 | tyrosine kinase 2 | 4270 | 0.035 | 0.1942 | No |
| 24 | LTB | lymphotoxin beta (TNF superfamily, member 3) | 4496 | 0.033 | 0.1905 | No |
| 25 | CXCL11 | chemokine (C-X-C motif) ligand 11 | 6249 | 0.021 | 0.1052 | No |
| 26 | IL18R1 | interleukin 18 receptor 1 | 6256 | 0.021 | 0.1097 | No |
| 27 | CRLF2 | cytokine receptor-like factor 2 | 6690 | 0.018 | 0.0916 | No |
| 28 | TNFRSF12A | tumor necrosis factor receptor superfamily, member 12A | 6787 | 0.017 | 0.0907 | No |
| 29 | CD9 | CD9 molecule | 6905 | 0.016 | 0.0886 | No |
| 30 | PIK3R5 | phosphoinositide-3-kinase, regulatory subunit 5 | 6911 | 0.016 | 0.0922 | No |
| 31 | TNFRSF1A | tumor necrosis factor receptor superfamily, member 1A | 7027 | 0.016 | 0.0900 | No |
| 32 | CCL7 | chemokine (C-C motif) ligand 7 | 7135 | 0.015 | 0.0881 | No |
| 33 | LTBR | lymphotoxin beta receptor (TNFR superfamily, member 3) | 7385 | 0.014 | 0.0785 | No |
| 34 | CSF2 | colony stimulating factor 2 (granulocyte-macrophage) | 7825 | 0.011 | 0.0584 | No |
| 35 | ACVRL1 | activin A receptor type IL | 7900 | 0.010 | 0.0571 | No |
| 36 | CXCL10 | chemokine (C-X-C motif) ligand 10 | 8286 | 0.008 | 0.0392 | No |
| 37 | IFNGR1 | interferon gamma receptor 1 | 8413 | 0.007 | 0.0345 | No |
| 38 | ACVR1B | activin A receptor type IB | 8737 | 0.006 | 0.0191 | No |
| 39 | IL12RB1 | interleukin 12 receptor, beta 1 | 8957 | 0.004 | 0.0089 | No |
| 40 | IFNGR2 | interferon gamma receptor 2 (interferon gamma transducer 1) | 9041 | 0.004 | 0.0055 | No |
| 41 | CXCL9 | chemokine (C-X-C motif) ligand 9 | 9517 | 0.001 | -0.0186 | No |
| 42 | TNFRSF1B | tumor necrosis factor receptor superfamily, member 1B | 9919 | -0.001 | -0.0391 | No |
| 43 | IL17RA | interleukin 17 receptor A | 9996 | -0.001 | -0.0427 | No |
| 44 | CSF3R | Memczak2013 ANTISENSE, CDS, coding, INTERNAL best transcript NM\_156039 | 10082 | -0.002 | -0.0467 | No |
| 45 | OSMR | oncostatin M receptor | 10294 | -0.003 | -0.0569 | No |
| 46 | MYD88 | myeloid differentiation primary response 88 | 10440 | -0.004 | -0.0634 | No |
| 47 | PTPN11 | protein tyrosine phosphatase, non-receptor type 11 | 10486 | -0.004 | -0.0648 | No |
| 48 | SOCS3 | suppressor of cytokine signaling 3 | 10714 | -0.005 | -0.0752 | No |
| 49 | IFNAR1 | interferon (alpha, beta and omega) receptor 1 | 10839 | -0.006 | -0.0801 | No |
| 50 | PIM1 | Pim-1 proto-oncogene, serine/threonine kinase | 11480 | -0.010 | -0.1107 | No |
| 51 | CBL | Cbl proto-oncogene, E3 ubiquitin protein ligase | 11535 | -0.010 | -0.1111 | No |
| 52 | BAK1 | BCL2-antagonist/killer 1 | 11698 | -0.011 | -0.1169 | No |
| 53 | IL6 | interleukin 6 | 11825 | -0.012 | -0.1206 | No |
| 54 | GRB2 | growth factor receptor bound protein 2 | 11880 | -0.012 | -0.1205 | No |
| 55 | TGFB1 | transforming growth factor beta 1 | 12464 | -0.016 | -0.1467 | No |
| 56 | SOCS1 | suppressor of cytokine signaling 1 | 12611 | -0.017 | -0.1503 | No |
| 57 | ITGB3 | integrin beta 3 | 12821 | -0.018 | -0.1567 | No |
| 58 | STAT1 | signal transducer and activator of transcription 1 | 12830 | -0.018 | -0.1529 | No |
| 59 | CD36 | CD36 molecule (thrombospondin receptor) | 13245 | -0.021 | -0.1693 | No |
| 60 | IL3RA | interleukin 3 receptor, alpha (low affinity) | 13504 | -0.023 | -0.1772 | No |
| 61 | CD14 | CD14 molecule | 13724 | -0.024 | -0.1827 | No |
| 62 | LEPR | leptin receptor | 13887 | -0.026 | -0.1850 | No |
| 63 | STAT3 | signal transducer and activator of transcription 3 (acute-phase response factor) | 14182 | -0.028 | -0.1936 | No |
| 64 | IL1R1 | interleukin 1 receptor, type I | 14863 | -0.033 | -0.2207 | No |
| 65 | REG1A | regenerating islet-derived 1 alpha | 15058 | -0.035 | -0.2225 | No |
| 66 | MAP3K8 | mitogen-activated protein kinase kinase kinase 8 | 15311 | -0.037 | -0.2267 | No |
| 67 | TNFRSF21 | tumor necrosis factor receptor superfamily, member 21 | 15802 | -0.042 | -0.2420 | No |
| 68 | IL2RA | interleukin 2 receptor, alpha | 16231 | -0.046 | -0.2531 | No |
| 69 | TLR2 | toll-like receptor 2 | 16448 | -0.049 | -0.2526 | No |
| 70 | CXCL13 | chemokine (C-X-C motif) ligand 13 | 16578 | -0.051 | -0.2472 | No |
| 71 | STAM2 | signal transducing adaptor molecule (SH3 domain and ITAM motif) 2 | 16637 | -0.052 | -0.2380 | No |
| 72 | PTPN1 | protein tyrosine phosphatase, non-receptor type 1 | 17307 | -0.062 | -0.2578 | Yes |
| 73 | PLA2G2A | phospholipase A2, group IIA (platelets, synovial fluid) | 17371 | -0.063 | -0.2461 | Yes |
| 74 | HAX1 | HCLS1 associated protein X-1 | 17593 | -0.067 | -0.2417 | Yes |
| 75 | IL10RB | interleukin 10 receptor, beta | 17739 | -0.071 | -0.2325 | Yes |
| 76 | IL13RA1 | interleukin 13 receptor, alpha 1 | 18154 | -0.081 | -0.2346 | Yes |
| 77 | CSF2RA | colony stimulating factor 2 receptor, alpha, low-affinity (granulocyte-macrophage) | 18316 | -0.086 | -0.2225 | Yes |
| 78 | DNTT | DNA nucleotidylexotransferase | 18359 | -0.087 | -0.2041 | Yes |
| 79 | EBI3 | Epstein-Barr virus induced 3 | 18431 | -0.090 | -0.1866 | Yes |
| 80 | STAT2 | signal transducer and activator of transcription 2 | 18572 | -0.095 | -0.1712 | Yes |
| 81 | CCR1 | chemokine (C-C motif) receptor 1 | 18734 | -0.105 | -0.1547 | Yes |
| 82 | IRF1 | interferon regulatory factor 1 | 18746 | -0.106 | -0.1302 | Yes |
| 83 | IRF9 | interferon regulatory factor 9 | 18860 | -0.114 | -0.1090 | Yes |
| 84 | IL1B | interleukin 1 beta | 19023 | -0.130 | -0.0867 | Yes |
| 85 | JUN | jun proto-oncogene | 19037 | -0.132 | -0.0562 | Yes |
| 86 | IL6ST | interleukin 6 signal transducer | 19134 | -0.147 | -0.0264 | Yes |
| 87 | INHBE | inhibin beta E | 19316 | -0.197 | 0.0107 | Yes |
Table: GSEA details [plain text format]

  

Fig 2: HALLMARK\_IL6\_JAK\_STAT3\_SIGNALING      
 Blue-Pink O' Gram in the Space of the Analyzed GeneSet

  

Fig 3: HALLMARK\_IL6\_JAK\_STAT3\_SIGNALING: Random ES distribution      
 Gene set null distribution of ES for **HALLMARK\_IL6\_JAK\_STAT3\_SIGNALING**

  
