## Supplementary material for "Somatic hypomethylation of pericentromeric SST1 repeats and tetraploidization in human colorectal cancer cells": GSEA results: HALLMARK_INFLAMMATORY_RESPONSE.html

Details for gene set HALLMARK\_INFLAMMATORY\_RESPONSE[GSEA]

|  || Dataset | eset\_byprobe\_collapsed\_to\_symbols.Diploid\_vs\_Tetraploid.cls #Tetraploid\_versus\_Diploid.Diploid\_vs\_Tetraploid.cls #Tetraploid\_versus\_Diploid\_repos |
| Phenotype | Diploid\_vs\_Tetraploid.cls#Tetraploid\_versus\_Diploid\_repos |
| Upregulated in class | Tetraploid |
| GeneSet | HALLMARK\_INFLAMMATORY\_RESPONSE |
| Enrichment Score (ES) | 0.35824633 |
| Normalized Enrichment Score (NES) | 1.5927532 |
| Nominal p-value | 0.0 |
| FDR q-value | 0.018129347 |
| FWER p-Value | 0.069 |
Table: GSEA Results Summary

  

Fig 1: Enrichment plot: HALLMARK\_INFLAMMATORY\_RESPONSE      
 Profile of the Running ES Score & Positions of GeneSet Members on the Rank Ordered List

  

| SYMBOL | TITLE | RANK IN GENE LIST | RANK METRIC SCORE | RUNNING ES | CORE ENRICHMENT || 1 | TNFSF15 | tumor necrosis factor (ligand) superfamily, member 15 | 8 | 0.444 | 0.0374 | Yes |
| 2 | CCL20 | chemokine (C-C motif) ligand 20 | 12 | 0.406 | 0.0717 | Yes |
| 3 | CD70 | CD70 molecule | 28 | 0.349 | 0.1007 | Yes |
| 4 | KCNJ2 | potassium channel, inwardly rectifying subfamily J, member 2 | 35 | 0.311 | 0.1268 | Yes |
| 5 | ITGB8 | integrin beta 8 | 52 | 0.283 | 0.1500 | Yes |
| 6 | PTGER4 | prostaglandin E receptor 4 (subtype EP4) | 59 | 0.272 | 0.1728 | Yes |
| 7 | IL15 | interleukin 15 | 64 | 0.261 | 0.1948 | Yes |
| 8 | F3 | coagulation factor III (thromboplastin, tissue factor) | 119 | 0.191 | 0.2083 | Yes |
| 9 | TNFSF9 | tumor necrosis factor (ligand) superfamily, member 9 | 223 | 0.150 | 0.2157 | Yes |
| 10 | IRF7 | interferon regulatory factor 7 | 230 | 0.148 | 0.2279 | Yes |
| 11 | IL1A | interleukin 1 alpha | 237 | 0.146 | 0.2401 | Yes |
| 12 | IRAK2 | interleukin 1 receptor associated kinase 2 | 250 | 0.142 | 0.2515 | Yes |
| 13 | TIMP1 | TIMP metallopeptidase inhibitor 1 | 254 | 0.141 | 0.2633 | Yes |
| 14 | BTG2 | BTG family, member 2 | 281 | 0.136 | 0.2736 | Yes |
| 15 | CXCL8 | chemokine (C-X-C motif) ligand 8 | 350 | 0.125 | 0.2807 | Yes |
| 16 | EDN1 | endothelin 1 | 371 | 0.123 | 0.2901 | Yes |
| 17 | ADGRE1 | adhesion G protein-coupled receptor E1 | 386 | 0.121 | 0.2997 | Yes |
| 18 | LIF | leukemia inhibitory factor | 423 | 0.117 | 0.3077 | Yes |
| 19 | ADORA2B | adenosine A2b receptor | 596 | 0.104 | 0.3076 | Yes |
| 20 | CDKN1A | cyclin-dependent kinase inhibitor 1A (p21, Cip1) | 690 | 0.097 | 0.3111 | Yes |
| 21 | PTGER2 | prostaglandin E receptor 2 | 737 | 0.095 | 0.3168 | Yes |
| 22 | HRH1 | histamine receptor H1 | 765 | 0.094 | 0.3234 | Yes |
| 23 | APLNR | apelin receptor | 896 | 0.088 | 0.3242 | Yes |
| 24 | BDKRB1 | bradykinin receptor B1 | 999 | 0.084 | 0.3261 | Yes |
| 25 | GP1BA | glycoprotein Ib (platelet), alpha polypeptide | 1000 | 0.084 | 0.3333 | Yes |
| 26 | TNFSF10 | tumor necrosis factor (ligand) superfamily, member 10 | 1006 | 0.084 | 0.3402 | Yes |
| 27 | CSF1 | colony stimulating factor 1 (macrophage) | 1040 | 0.083 | 0.3455 | Yes |
| 28 | IFITM1 | interferon induced transmembrane protein 1 | 1056 | 0.082 | 0.3517 | Yes |
| 29 | IL4R | interleukin 4 receptor | 1066 | 0.082 | 0.3582 | Yes |
| 30 | PLAUR | plasminogen activator, urokinase receptor | 1355 | 0.075 | 0.3497 | No |
| 31 | SPHK1 | sphingosine kinase 1 | 1575 | 0.070 | 0.3443 | No |
| 32 | ICAM1 | intercellular adhesion molecule 1 | 1606 | 0.069 | 0.3486 | No |
| 33 | CX3CL1 | chemokine (C-X3-C motif) ligand 1 | 1648 | 0.068 | 0.3522 | No |
| 34 | CD82 | CD82 molecule | 2272 | 0.057 | 0.3249 | No |
| 35 | IL15RA | interleukin 15 receptor, alpha | 2659 | 0.051 | 0.3093 | No |
| 36 | CYBB | cytochrome b-245, beta polypeptide | 2945 | 0.048 | 0.2986 | No |
| 37 | BST2 | bone marrow stromal cell antigen 2 | 2954 | 0.048 | 0.3023 | No |
| 38 | MYC | v-myc avian myelocytomatosis viral oncogene homolog | 3194 | 0.046 | 0.2938 | No |
| 39 | IL10RA | interleukin 10 receptor, alpha | 3223 | 0.045 | 0.2962 | No |
| 40 | PROK2 | prokineticin 2 | 3307 | 0.045 | 0.2957 | No |
| 41 | SEMA4D | Memczak2013 ANTISENSE, coding, INTERNAL, UTR3 best transcript NM\_006378 | 3437 | 0.043 | 0.2927 | No |
| 42 | NMUR1 | neuromedin U receptor 1 | 3453 | 0.043 | 0.2956 | No |
| 43 | IL7R | interleukin 7 receptor | 3650 | 0.041 | 0.2890 | No |
| 44 | NFKB1 | nuclear factor of kappa light polypeptide gene enhancer in B-cells 1 | 3856 | 0.039 | 0.2817 | No |
| 45 | DCBLD2 | discoidin, CUB and LCCL domain containing 2 | 3862 | 0.039 | 0.2847 | No |
| 46 | FFAR2 | free fatty acid receptor 2 | 3956 | 0.038 | 0.2831 | No |
| 47 | OPRK1 | opioid receptor, kappa 1 | 4023 | 0.037 | 0.2829 | No |
| 48 | HPN | hepsin | 4274 | 0.035 | 0.2729 | No |
| 49 | CCR7 | chemokine (C-C motif) receptor 7 | 4390 | 0.034 | 0.2699 | No |
| 50 | TACR3 | tachykinin receptor 3 | 4600 | 0.032 | 0.2618 | No |
| 51 | ICAM4 | intercellular adhesion molecule 4 (Landsteiner-Wiener blood group) | 4895 | 0.030 | 0.2492 | No |
| 52 | NDP | Norrie disease (pseudoglioma) | 5007 | 0.029 | 0.2459 | No |
| 53 | SLC31A2 | solute carrier family 31 (copper transporter), member 2 | 5143 | 0.028 | 0.2413 | No |
| 54 | LCP2 | lymphocyte cytosolic protein 2 | 5232 | 0.027 | 0.2391 | No |
| 55 | SLC11A2 | solute carrier family 11 (proton-coupled divalent metal ion transporter), member 2 | 5392 | 0.026 | 0.2331 | No |
| 56 | CCL2 | chemokine (C-C motif) ligand 2 | 5479 | 0.026 | 0.2309 | No |
| 57 | RASGRP1 | RAS guanyl releasing protein 1 (calcium and DAG-regulated) | 5558 | 0.025 | 0.2289 | No |
| 58 | P2RX4 | purinergic receptor P2X, ligand gated ion channel, 4 | 5563 | 0.025 | 0.2308 | No |
| 59 | EREG | epiregulin | 5902 | 0.023 | 0.2153 | No |
| 60 | NLRP3 | NLR family, pyrin domain containing 3 | 6017 | 0.022 | 0.2113 | No |
| 61 | CHST2 | carbohydrate (N-acetylglucosamine-6-O) sulfotransferase 2 | 6022 | 0.022 | 0.2129 | No |
| 62 | LCK | LCK proto-oncogene, Src family tyrosine kinase | 6061 | 0.022 | 0.2128 | No |
| 63 | IL10 | interleukin 10 | 6071 | 0.022 | 0.2142 | No |
| 64 | CCL24 | chemokine (C-C motif) ligand 24 | 6077 | 0.022 | 0.2157 | No |
| 65 | MET | MET proto-oncogene, receptor tyrosine kinase | 6216 | 0.021 | 0.2103 | No |
| 66 | CXCL11 | chemokine (C-X-C motif) ligand 11 | 6249 | 0.021 | 0.2104 | No |
| 67 | IL18R1 | interleukin 18 receptor 1 | 6256 | 0.021 | 0.2119 | No |
| 68 | NOD2 | nucleotide-binding oligomerization domain containing 2 | 6453 | 0.019 | 0.2034 | No |
| 69 | GPR132 | G protein-coupled receptor 132 | 6591 | 0.018 | 0.1979 | No |
| 70 | RTP4 | receptor (chemosensory) transporter protein 4 | 6759 | 0.017 | 0.1907 | No |
| 71 | PIK3R5 | phosphoinositide-3-kinase, regulatory subunit 5 | 6911 | 0.016 | 0.1843 | No |
| 72 | CCL7 | chemokine (C-C motif) ligand 7 | 7135 | 0.015 | 0.1740 | No |
| 73 | RAF1 | Raf-1 proto-oncogene, serine/threonine kinase | 7383 | 0.014 | 0.1624 | No |
| 74 | SGMS2 | sphingomyelin synthase 2 | 7491 | 0.013 | 0.1579 | No |
| 75 | CSF3 | colony stimulating factor 3 | 7735 | 0.011 | 0.1463 | No |
| 76 | MSR1 | macrophage scavenger receptor 1 | 7748 | 0.011 | 0.1467 | No |
| 77 | SLC1A2 | solute carrier family 1 (glial high affinity glutamate transporter), member 2 | 7798 | 0.011 | 0.1451 | No |
| 78 | IL18 | interleukin 18 | 7966 | 0.010 | 0.1373 | No |
| 79 | TLR1 | toll-like receptor 1 | 8229 | 0.008 | 0.1245 | No |
| 80 | CXCL10 | chemokine (C-X-C motif) ligand 10 | 8286 | 0.008 | 0.1223 | No |
| 81 | OLR1 | oxidized low density lipoprotein (lectin-like) receptor 1 | 8290 | 0.008 | 0.1228 | No |
| 82 | LTA | lymphotoxin alpha | 8541 | 0.007 | 0.1104 | No |
| 83 | CCL5 | chemokine (C-C motif) ligand 5 | 8596 | 0.006 | 0.1082 | No |
| 84 | ITGA5 | integrin alpha 5 | 8613 | 0.006 | 0.1079 | No |
| 85 | ACVR1B | activin A receptor type IB | 8737 | 0.006 | 0.1020 | No |
| 86 | PTPRE | protein tyrosine phosphatase, receptor type, E | 8781 | 0.005 | 0.1002 | No |
| 87 | SLAMF1 | signaling lymphocytic activation molecule family member 1 | 8858 | 0.005 | 0.0967 | No |
| 88 | LY6E | lymphocyte antigen 6 complex, locus E | 8909 | 0.005 | 0.0945 | No |
| 89 | IFNGR2 | interferon gamma receptor 2 (interferon gamma transducer 1) | 9041 | 0.004 | 0.0880 | No |
| 90 | MMP14 | matrix metallopeptidase 14 (membrane-inserted) | 9110 | 0.003 | 0.0848 | No |
| 91 | SCARF1 | scavenger receptor class F, member 1 | 9200 | 0.003 | 0.0805 | No |
| 92 | PTAFR | platelet-activating factor receptor | 9251 | 0.003 | 0.0781 | No |
| 93 | CD40 | CD40 molecule, TNF receptor superfamily member 5 | 9416 | 0.002 | 0.0698 | No |
| 94 | LDLR | low density lipoprotein receptor | 9447 | 0.002 | 0.0684 | No |
| 95 | CXCL9 | chemokine (C-X-C motif) ligand 9 | 9517 | 0.001 | 0.0649 | No |
| 96 | NFKBIA | nuclear factor of kappa light polypeptide gene enhancer in B-cells inhibitor, alpha | 9745 | 0.000 | 0.0532 | No |
| 97 | KCNMB2 | potassium channel subfamily M regulatory beta subunit 2 | 9846 | -0.000 | 0.0480 | No |
| 98 | PTGIR | prostaglandin I2 (prostacyclin) receptor (IP) | 9847 | -0.000 | 0.0481 | No |
| 99 | IL18RAP | interleukin 18 receptor accessory protein | 9873 | -0.000 | 0.0468 | No |
| 100 | TNFRSF1B | tumor necrosis factor receptor superfamily, member 1B | 9919 | -0.001 | 0.0446 | No |
| 101 | IL2RB | interleukin 2 receptor, beta | 10056 | -0.001 | 0.0377 | No |
| 102 | CSF3R | Memczak2013 ANTISENSE, CDS, coding, INTERNAL best transcript NM\_156039 | 10082 | -0.002 | 0.0365 | No |
| 103 | ADRM1 | adhesion regulating molecule 1 | 10136 | -0.002 | 0.0339 | No |
| 104 | OSMR | oncostatin M receptor | 10294 | -0.003 | 0.0260 | No |
| 105 | FPR1 | formyl peptide receptor 1 | 10329 | -0.003 | 0.0245 | No |
| 106 | OSM | oncostatin M | 10331 | -0.003 | 0.0248 | No |
| 107 | RHOG | ras homolog family member G | 10457 | -0.004 | 0.0186 | No |
| 108 | GABBR1 | gamma-aminobutyric acid (GABA) B receptor, 1 | 10502 | -0.004 | 0.0167 | No |
| 109 | PSEN1 | presenilin 1 | 10566 | -0.004 | 0.0138 | No |
| 110 | GNAI3 | guanine nucleotide binding protein (G protein), alpha inhibiting activity polypeptide 3 | 10833 | -0.006 | 0.0006 | No |
| 111 | IFNAR1 | interferon (alpha, beta and omega) receptor 1 | 10839 | -0.006 | 0.0008 | No |
| 112 | RGS16 | regulator of G-protein signaling 16 | 11032 | -0.007 | -0.0085 | No |
| 113 | KCNA3 | potassium channel, voltage gated shaker related subfamily A, member 3 | 11304 | -0.009 | -0.0218 | No |
| 114 | FZD5 | frizzled class receptor 5 | 11350 | -0.009 | -0.0233 | No |
| 115 | CALCRL | calcitonin receptor like receptor | 11374 | -0.009 | -0.0237 | No |
| 116 | CCL17 | chemokine (C-C motif) ligand 17 | 11513 | -0.010 | -0.0300 | No |
| 117 | ATP2C1 | ATPase, Ca++ transporting, type 2C, member 1 | 11578 | -0.010 | -0.0325 | No |
| 118 | ATP2A2 | ATPase, Ca++ transporting, cardiac muscle, slow twitch 2 | 11662 | -0.011 | -0.0358 | No |
| 119 | IL6 | interleukin 6 | 11825 | -0.012 | -0.0432 | No |
| 120 | CXCL6 | chemokine (C-X-C motif) ligand 6 | 11900 | -0.012 | -0.0460 | No |
| 121 | AXL | AXL receptor tyrosine kinase | 11942 | -0.012 | -0.0471 | No |
| 122 | CD55 | CD55 molecule, decay accelerating factor for complement (Cromer blood group) | 12082 | -0.013 | -0.0531 | No |
| 123 | SELL | selectin L | 12417 | -0.016 | -0.0691 | No |
| 124 | CMKLR1 | chemerin chemokine-like receptor 1 | 12531 | -0.016 | -0.0736 | No |
| 125 | RIPK2 | receptor-interacting serine-threonine kinase 2 | 12627 | -0.017 | -0.0770 | No |
| 126 | GCH1 | GTP cyclohydrolase 1 | 12793 | -0.018 | -0.0841 | No |
| 127 | ITGB3 | integrin beta 3 | 12821 | -0.018 | -0.0839 | No |
| 128 | CD48 | CD48 molecule | 12879 | -0.018 | -0.0853 | No |
| 129 | KLF6 | Kruppel-like factor 6 | 12945 | -0.019 | -0.0871 | No |
| 130 | SERPINE1 | serpin peptidase inhibitor, clade E (nexin, plasminogen activator inhibitor type 1), member 1 | 12974 | -0.019 | -0.0869 | No |
| 131 | AHR | aryl hydrocarbon receptor | 13095 | -0.020 | -0.0914 | No |
| 132 | MEFV | Mediterranean fever | 13146 | -0.020 | -0.0923 | No |
| 133 | SCN1B | sodium channel, voltage gated, type I beta subunit | 13217 | -0.021 | -0.0941 | No |
| 134 | CD69 | CD69 molecule | 13361 | -0.022 | -0.0997 | No |
| 135 | ABI1 | abl-interactor 1 | 13706 | -0.024 | -0.1154 | No |
| 136 | CD14 | CD14 molecule | 13724 | -0.024 | -0.1143 | No |
| 137 | GNA15 | guanine nucleotide binding protein (G protein), alpha 15 (Gq class) | 13912 | -0.026 | -0.1217 | No |
| 138 | ACVR2A | activin A receptor type IIA | 14013 | -0.026 | -0.1247 | No |
| 139 | TAPBP | TAP binding protein (tapasin) | 14083 | -0.027 | -0.1260 | No |
| 140 | PVR | poliovirus receptor | 14143 | -0.027 | -0.1267 | No |
| 141 | SRI | sorcin | 14400 | -0.030 | -0.1374 | No |
| 142 | LYN | LYN proto-oncogene, Src family tyrosine kinase | 14438 | -0.030 | -0.1368 | No |
| 143 | HBEGF | heparin-binding EGF-like growth factor | 14753 | -0.032 | -0.1503 | No |
| 144 | MARCO | macrophage receptor with collagenous structure | 14821 | -0.033 | -0.1509 | No |
| 145 | IL1R1 | interleukin 1 receptor, type I | 14863 | -0.033 | -0.1502 | No |
| 146 | NPFFR2 | neuropeptide FF receptor 2 | 14882 | -0.033 | -0.1483 | No |
| 147 | SLC7A2 | solute carrier family 7 (cationic amino acid transporter, y+ system), member 2 | 14954 | -0.034 | -0.1491 | No |
| 148 | INHBA | inhibin beta A | 15087 | -0.035 | -0.1529 | No |
| 149 | STAB1 | stabilin 1 | 15306 | -0.037 | -0.1610 | No |
| 150 | RELA | v-rel avian reticuloendotheliosis viral oncogene homolog A | 15578 | -0.039 | -0.1717 | No |
| 151 | HAS2 | hyaluronan synthase 2 | 15583 | -0.040 | -0.1686 | No |
| 152 | RGS1 | regulator of G-protein signaling 1 | 15765 | -0.041 | -0.1744 | No |
| 153 | SLC4A4 | solute carrier family 4 (sodium bicarbonate cotransporter), member 4 | 15861 | -0.042 | -0.1757 | No |
| 154 | TNFRSF9 | tumor necrosis factor receptor superfamily, member 9 | 15868 | -0.042 | -0.1724 | No |
| 155 | PDPN | podoplanin | 15902 | -0.043 | -0.1705 | No |
| 156 | ABCA1 | ATP binding cassette subfamily A member 1 | 15924 | -0.043 | -0.1679 | No |
| 157 | ADM | adrenomedullin | 16402 | -0.048 | -0.1885 | No |
| 158 | TACR1 | tachykinin receptor 1 | 16439 | -0.049 | -0.1862 | No |
| 159 | TLR2 | toll-like receptor 2 | 16448 | -0.049 | -0.1824 | No |
| 160 | EIF2AK2 | eukaryotic translation initiation factor 2-alpha kinase 2 | 16462 | -0.049 | -0.1789 | No |
| 161 | KIF1B | kinesin family member 1B | 16464 | -0.049 | -0.1748 | No |
| 162 | GPC3 | glypican 3 | 16605 | -0.051 | -0.1777 | No |
| 163 | NAMPT | nicotinamide phosphoribosyltransferase | 16683 | -0.052 | -0.1772 | No |
| 164 | C5AR1 | complement component 5a receptor 1 | 16693 | -0.052 | -0.1732 | No |
| 165 | EMP3 | epithelial membrane protein 3 | 16695 | -0.052 | -0.1688 | No |
| 166 | VIP | vasoactive intestinal peptide | 16902 | -0.055 | -0.1747 | No |
| 167 | ICOSLG | inducible T-cell co-stimulator ligand | 16929 | -0.056 | -0.1714 | No |
| 168 | HIF1A | hypoxia inducible factor 1, alpha subunit (basic helix-loop-helix transcription factor) | 16936 | -0.056 | -0.1669 | No |
| 169 | C3AR1 | complement component 3a receptor 1 | 17045 | -0.058 | -0.1676 | No |
| 170 | TPBG | trophoblast glycoprotein | 17065 | -0.058 | -0.1637 | No |
| 171 | P2RY2 | purinergic receptor P2Y, G-protein coupled, 2 | 17123 | -0.059 | -0.1616 | No |
| 172 | ATP2B1 | ATPase, Ca++ transporting, plasma membrane 1 | 17206 | -0.060 | -0.1608 | No |
| 173 | LPAR1 | lysophosphatidic acid receptor 1 | 17303 | -0.062 | -0.1605 | No |
| 174 | RNF144B | ring finger protein 144B | 17334 | -0.062 | -0.1567 | No |
| 175 | SLC7A1 | solute carrier family 7 (cationic amino acid transporter, y+ system), member 1 | 17491 | -0.065 | -0.1593 | No |
| 176 | GPR183 | G protein-coupled receptor 183 | 17802 | -0.072 | -0.1692 | No |
| 177 | PCDH7 | protocadherin 7 | 17860 | -0.073 | -0.1659 | No |
| 178 | SLC31A1 | solute carrier family 31 (copper transporter), member 1 | 18145 | -0.081 | -0.1737 | No |
| 179 | SELE | selectin E | 18155 | -0.081 | -0.1672 | No |
| 180 | TNFAIP6 | tumor necrosis factor, alpha-induced protein 6 | 18183 | -0.082 | -0.1616 | No |
| 181 | CCL22 | chemokine (C-C motif) ligand 22 | 18217 | -0.083 | -0.1563 | No |
| 182 | AQP9 | aquaporin 9 | 18219 | -0.083 | -0.1493 | No |
| 183 | MXD1 | Memczak2013 ANTISENSE, coding, INTERNAL, intronic best transcript NM\_001202514 | 18336 | -0.086 | -0.1479 | No |
| 184 | IL12B | interleukin 12B | 18360 | -0.087 | -0.1417 | No |
| 185 | EBI3 | Epstein-Barr virus induced 3 | 18431 | -0.090 | -0.1377 | No |
| 186 | MEP1A | meprin A, alpha (PABA peptide hydrolase) | 18452 | -0.091 | -0.1310 | No |
| 187 | NMI | N-myc (and STAT) interactor | 18617 | -0.097 | -0.1312 | No |
| 188 | IRF1 | interferon regulatory factor 1 | 18746 | -0.106 | -0.1288 | No |
| 189 | LAMP3 | lysosomal-associated membrane protein 3 | 18788 | -0.109 | -0.1217 | No |
| 190 | CLEC5A | C-type lectin domain family 5, member A | 18845 | -0.113 | -0.1149 | No |
| 191 | IL1B | interleukin 1 beta | 19023 | -0.130 | -0.1131 | No |
| 192 | CXCR6 | chemokine (C-X-C motif) receptor 6 | 19046 | -0.133 | -0.1029 | No |
| 193 | PDE4B | phosphodiesterase 4B, cAMP-specific | 19061 | -0.135 | -0.0921 | No |
| 194 | P2RX7 | purinergic receptor P2X, ligand gated ion channel, 7 | 19094 | -0.141 | -0.0818 | No |
| 195 | ROS1 | ROS proto-oncogene 1 , receptor tyrosine kinase | 19112 | -0.143 | -0.0705 | No |
| 196 | CCRL2 | chemokine (C-C motif) receptor-like 2 | 19232 | -0.168 | -0.0624 | No |
| 197 | SLC28A2 | solute carrier family 28 (concentrative nucleoside transporter), member 2 | 19341 | -0.205 | -0.0506 | No |
| 198 | BEST1 | bestrophin 1 | 19444 | -0.294 | -0.0309 | No |
| 199 | TLR3 | toll-like receptor 3 | 19502 | -0.411 | 0.0011 | No |
Table: GSEA details [plain text format]

  

Fig 2: HALLMARK\_INFLAMMATORY\_RESPONSE      
 Blue-Pink O' Gram in the Space of the Analyzed GeneSet

  

Fig 3: HALLMARK\_INFLAMMATORY\_RESPONSE: Random ES distribution      
 Gene set null distribution of ES for **HALLMARK\_INFLAMMATORY\_RESPONSE**

  
