## Supplementary material for "Somatic hypomethylation of pericentromeric SST1 repeats and tetraploidization in human colorectal cancer cells": GSEA results: HALLMARK_INTERFERON_ALPHA_RESPONSE.html

Details for gene set HALLMARK\_INTERFERON\_ALPHA\_RESPONSE[GSEA]

|  || Dataset | eset\_byprobe\_collapsed\_to\_symbols.Diploid\_vs\_Tetraploid.cls #Tetraploid\_versus\_Diploid.Diploid\_vs\_Tetraploid.cls #Tetraploid\_versus\_Diploid\_repos |
| Phenotype | Diploid\_vs\_Tetraploid.cls#Tetraploid\_versus\_Diploid\_repos |
| Upregulated in class | Diploid |
| GeneSet | HALLMARK\_INTERFERON\_ALPHA\_RESPONSE |
| Enrichment Score (ES) | -0.33017758 |
| Normalized Enrichment Score (NES) | -1.2390056 |
| Nominal p-value | 0.14471544 |
| FDR q-value | 0.27929345 |
| FWER p-Value | 0.967 |
Table: GSEA Results Summary

  

Fig 1: Enrichment plot: HALLMARK\_INTERFERON\_ALPHA\_RESPONSE      
 Profile of the Running ES Score & Positions of GeneSet Members on the Rank Ordered List

  

| SYMBOL | TITLE | RANK IN GENE LIST | RANK METRIC SCORE | RUNNING ES | CORE ENRICHMENT || 1 | IL15 | interleukin 15 | 64 | 0.261 | 0.0495 | No |
| 2 | IRF7 | interferon regulatory factor 7 | 230 | 0.148 | 0.0709 | No |
| 3 | GBP2 | guanylate binding protein 2, interferon-inducible | 302 | 0.133 | 0.0943 | No |
| 4 | CD47 | CD47 molecule | 349 | 0.125 | 0.1173 | No |
| 5 | PROCR | protein C receptor, endothelial | 384 | 0.121 | 0.1400 | No |
| 6 | IFI27 | interferon, alpha-inducible protein 27 | 1013 | 0.084 | 0.1247 | No |
| 7 | IRF2 | interferon regulatory factor 2 | 1032 | 0.083 | 0.1405 | No |
| 8 | CSF1 | colony stimulating factor 1 (macrophage) | 1040 | 0.083 | 0.1570 | No |
| 9 | IFITM1 | interferon induced transmembrane protein 1 | 1056 | 0.082 | 0.1728 | No |
| 10 | IL4R | interleukin 4 receptor | 1066 | 0.082 | 0.1889 | No |
| 11 | IL7 | interleukin 7 | 1267 | 0.077 | 0.1943 | No |
| 12 | TRIM21 | tripartite motif containing 21 | 1810 | 0.065 | 0.1794 | No |
| 13 | OASL | 2-5-oligoadenylate synthetase-like | 2077 | 0.060 | 0.1779 | No |
| 14 | BST2 | bone marrow stromal cell antigen 2 | 2954 | 0.048 | 0.1426 | No |
| 15 | CMPK2 | cytidine monophosphate (UMP-CMP) kinase 2, mitochondrial | 3051 | 0.047 | 0.1471 | No |
| 16 | HELZ2 | helicase with zinc finger 2, transcriptional coactivator | 3945 | 0.038 | 0.1089 | No |
| 17 | TRIM26 | tripartite motif containing 26 | 4319 | 0.035 | 0.0967 | No |
| 18 | TRIM14 | tripartite motif containing 14 | 4463 | 0.033 | 0.0961 | No |
| 19 | PSME1 | proteasome activator subunit 1 | 4737 | 0.031 | 0.0884 | No |
| 20 | OGFR | opioid growth factor receptor | 4790 | 0.031 | 0.0919 | No |
| 21 | PNPT1 | polyribonucleotide nucleotidyltransferase 1 | 5539 | 0.025 | 0.0585 | No |
| 22 | HLA-C | major histocompatibility complex, class I, C | 5587 | 0.025 | 0.0611 | No |
| 23 | MVB12A | multivesicular body subunit 12A | 5743 | 0.024 | 0.0579 | No |
| 24 | PLSCR1 | phospholipid scramblase 1 | 5829 | 0.023 | 0.0583 | No |
| 25 | CD74 | CD74 molecule, major histocompatibility complex, class II invariant chain | 5852 | 0.023 | 0.0618 | No |
| 26 | SLC25A28 | solute carrier family 25 (mitochondrial iron transporter), member 28 | 5910 | 0.023 | 0.0634 | No |
| 27 | IFI44L | interferon-induced protein 44-like | 5931 | 0.022 | 0.0669 | No |
| 28 | TAP1 | transporter 1, ATP-binding cassette, sub-family B (MDR/TAP) | 6169 | 0.021 | 0.0590 | No |
| 29 | CXCL11 | chemokine (C-X-C motif) ligand 11 | 6249 | 0.021 | 0.0591 | No |
| 30 | RTP4 | receptor (chemosensory) transporter protein 4 | 6759 | 0.017 | 0.0364 | No |
| 31 | B2M | beta-2-microglobulin | 7058 | 0.015 | 0.0242 | No |
| 32 | IFITM2 | interferon induced transmembrane protein 2 | 7142 | 0.015 | 0.0229 | No |
| 33 | GMPR | guanosine monophosphate reductase | 7328 | 0.014 | 0.0162 | No |
| 34 | IFITM3 | interferon induced transmembrane protein 3 | 7579 | 0.012 | 0.0059 | No |
| 35 | RNF31 | ring finger protein 31 | 7656 | 0.012 | 0.0044 | No |
| 36 | ELF1 | E74-like factor 1 (ets domain transcription factor) | 8011 | 0.010 | -0.0118 | No |
| 37 | OAS1 | 2-5-oligoadenylate synthetase 1 | 8073 | 0.009 | -0.0131 | No |
| 38 | PSMA3 | proteasome subunit alpha 3 | 8134 | 0.009 | -0.0143 | No |
| 39 | PSMB9 | proteasome subunit beta 9 | 8144 | 0.009 | -0.0129 | No |
| 40 | CXCL10 | chemokine (C-X-C motif) ligand 10 | 8286 | 0.008 | -0.0186 | No |
| 41 | PSME2 | proteasome activator subunit 2 | 8352 | 0.008 | -0.0204 | No |
| 42 | RSAD2 | radical S-adenosyl methionine domain containing 2 | 8715 | 0.006 | -0.0378 | No |
| 43 | CNP | 2,3-cyclic nucleotide 3 phosphodiesterase | 8879 | 0.005 | -0.0453 | No |
| 44 | LY6E | lymphocyte antigen 6 complex, locus E | 8909 | 0.005 | -0.0458 | No |
| 45 | LGALS3BP | lectin, galactoside-binding, soluble, 3 binding protein | 9239 | 0.003 | -0.0622 | No |
| 46 | TRIM25 | tripartite motif containing 25 | 9786 | -0.000 | -0.0903 | No |
| 47 | DDX60 | DEAD (Asp-Glu-Ala-Asp) box polypeptide 60 | 10356 | -0.003 | -0.1189 | No |
| 48 | PARP9 | poly(ADP-ribose) polymerase family member 9 | 10477 | -0.004 | -0.1243 | No |
| 49 | LPAR6 | lysophosphatidic acid receptor 6 | 10501 | -0.004 | -0.1246 | No |
| 50 | MX1 | MX dynamin-like GTPase 1 | 10754 | -0.006 | -0.1365 | No |
| 51 | IFIT2 | interferon-induced protein with tetratricopeptide repeats 2 | 11183 | -0.008 | -0.1568 | No |
| 52 | CASP8 | caspase 8, apoptosis-related cysteine peptidase | 11632 | -0.011 | -0.1777 | No |
| 53 | IFI35 | interferon-induced protein 35 | 11915 | -0.012 | -0.1898 | No |
| 54 | PARP14 | poly(ADP-ribose) polymerase family member 14 | 12214 | -0.014 | -0.2022 | No |
| 55 | LAP3 | leucine aminopeptidase 3 | 12270 | -0.015 | -0.2021 | No |
| 56 | BATF2 | basic leucine zipper transcription factor, ATF-like 2 | 12411 | -0.016 | -0.2062 | No |
| 57 | SELL | selectin L | 12417 | -0.016 | -0.2033 | No |
| 58 | ISG15 | ISG15 ubiquitin-like modifier | 12602 | -0.017 | -0.2094 | No |
| 59 | ADAR | adenosine deaminase, RNA-specific | 12610 | -0.017 | -0.2063 | No |
| 60 | RIPK2 | receptor-interacting serine-threonine kinase 2 | 12627 | -0.017 | -0.2037 | No |
| 61 | PSMB8 | proteasome subunit beta 8 | 12759 | -0.018 | -0.2069 | No |
| 62 | MOV10 | Mov10 RISC complex RNA helicase | 13247 | -0.021 | -0.2277 | No |
| 63 | C1S | complement component 1, s subcomponent | 13614 | -0.023 | -0.2418 | No |
| 64 | IFIT3 | interferon-induced protein with tetratricopeptide repeats 3 | 13619 | -0.023 | -0.2373 | No |
| 65 | GBP4 | guanylate binding protein 4 | 14250 | -0.028 | -0.2640 | No |
| 66 | CMTR1 | cap methyltransferase 1 | 14622 | -0.031 | -0.2767 | No |
| 67 | SAMD9L | sterile alpha motif domain containing 9-like | 14841 | -0.033 | -0.2812 | No |
| 68 | HERC6 | HECT and RLD domain containing E3 ubiquitin protein ligase family member 6 | 14995 | -0.034 | -0.2822 | No |
| 69 | UBA7 | Memczak2013 ANTISENSE, coding, INTERNAL, intronic best transcript NM\_003335 | 15871 | -0.042 | -0.3186 | No |
| 70 | TXNIP | thioredoxin interacting protein | 16097 | -0.045 | -0.3211 | Yes |
| 71 | USP18 | ubiquitin specific peptidase 18 | 16182 | -0.046 | -0.3161 | Yes |
| 72 | SP110 | SP110 nuclear body protein | 16263 | -0.047 | -0.3108 | Yes |
| 73 | ISG20 | interferon stimulated exonuclease gene 20kDa | 16332 | -0.047 | -0.3047 | Yes |
| 74 | IFIH1 | interferon induced, with helicase C domain 1 | 16404 | -0.048 | -0.2986 | Yes |
| 75 | EIF2AK2 | eukaryotic translation initiation factor 2-alpha kinase 2 | 16462 | -0.049 | -0.2915 | Yes |
| 76 | NUB1 | negative regulator of ubiquitin-like proteins 1 | 16772 | -0.053 | -0.2966 | Yes |
| 77 | EPSTI1 | epithelial stromal interaction 1 (breast) | 17199 | -0.060 | -0.3064 | Yes |
| 78 | TMEM140 | transmembrane protein 140 | 17324 | -0.062 | -0.3002 | Yes |
| 79 | TDRD7 | tudor domain containing 7 | 17494 | -0.065 | -0.2957 | Yes |
| 80 | TRAFD1 | TRAF-type zinc finger domain containing 1 | 17666 | -0.069 | -0.2906 | Yes |
| 81 | TRIM5 | tripartite motif containing 5 | 17675 | -0.069 | -0.2771 | Yes |
| 82 | NCOA7 | nuclear receptor coactivator 7 | 17806 | -0.072 | -0.2692 | Yes |
| 83 | PARP12 | poly(ADP-ribose) polymerase family member 12 | 18125 | -0.081 | -0.2692 | Yes |
| 84 | UBE2L6 | ubiquitin-conjugating enzyme E2L 6 | 18293 | -0.085 | -0.2606 | Yes |
| 85 | DHX58 | DEXH (Asp-Glu-X-His) box polypeptide 58 | 18439 | -0.090 | -0.2499 | Yes |
| 86 | STAT2 | signal transducer and activator of transcription 2 | 18572 | -0.095 | -0.2374 | Yes |
| 87 | NMI | N-myc (and STAT) interactor | 18617 | -0.097 | -0.2200 | Yes |
| 88 | IRF1 | interferon regulatory factor 1 | 18746 | -0.106 | -0.2051 | Yes |
| 89 | LAMP3 | lysosomal-associated membrane protein 3 | 18788 | -0.109 | -0.1852 | Yes |
| 90 | IRF9 | interferon regulatory factor 9 | 18860 | -0.114 | -0.1657 | Yes |
| 91 | CCRL2 | chemokine (C-C motif) receptor-like 2 | 19232 | -0.168 | -0.1508 | Yes |
| 92 | IFI44 | interferon-induced protein 44 | 19319 | -0.197 | -0.1154 | Yes |
| 93 | CASP1 | caspase 1 | 19457 | -0.309 | -0.0600 | Yes |
| 94 | SAMD9 | sterile alpha motif domain containing 9 | 19462 | -0.313 | 0.0032 | Yes |
Table: GSEA details [plain text format]

  

Fig 2: HALLMARK\_INTERFERON\_ALPHA\_RESPONSE      
 Blue-Pink O' Gram in the Space of the Analyzed GeneSet

  

Fig 3: HALLMARK\_INTERFERON\_ALPHA\_RESPONSE: Random ES distribution      
 Gene set null distribution of ES for **HALLMARK\_INTERFERON\_ALPHA\_RESPONSE**

  
