## Supplementary material for "Somatic hypomethylation of pericentromeric SST1 repeats and tetraploidization in human colorectal cancer cells": GSEA results: HALLMARK_INTERFERON_GAMMA_RESPONSE.html

Details for gene set HALLMARK\_INTERFERON\_GAMMA\_RESPONSE[GSEA]

|  || Dataset | eset\_byprobe\_collapsed\_to\_symbols.Diploid\_vs\_Tetraploid.cls #Tetraploid\_versus\_Diploid.Diploid\_vs\_Tetraploid.cls #Tetraploid\_versus\_Diploid\_repos |
| Phenotype | Diploid\_vs\_Tetraploid.cls#Tetraploid\_versus\_Diploid\_repos |
| Upregulated in class | Diploid |
| GeneSet | HALLMARK\_INTERFERON\_GAMMA\_RESPONSE |
| Enrichment Score (ES) | -0.2776694 |
| Normalized Enrichment Score (NES) | -1.1524676 |
| Nominal p-value | 0.16591929 |
| FDR q-value | 0.3462616 |
| FWER p-Value | 0.997 |
Table: GSEA Results Summary

  

Fig 1: Enrichment plot: HALLMARK\_INTERFERON\_GAMMA\_RESPONSE      
 Profile of the Running ES Score & Positions of GeneSet Members on the Rank Ordered List

  

| SYMBOL | TITLE | RANK IN GENE LIST | RANK METRIC SCORE | RUNNING ES | CORE ENRICHMENT || 1 | HLA-DMA | major histocompatibility complex, class II, DM alpha | 15 | 0.392 | 0.0382 | No |
| 2 | IL15 | interleukin 15 | 64 | 0.261 | 0.0617 | No |
| 3 | CFB | complement factor B | 164 | 0.167 | 0.0731 | No |
| 4 | FAS | Fas cell surface death receptor | 186 | 0.161 | 0.0880 | No |
| 5 | IRF7 | interferon regulatory factor 7 | 230 | 0.148 | 0.1005 | No |
| 6 | EIF4E3 | eukaryotic translation initiation factor 4E family member 3 | 239 | 0.146 | 0.1145 | No |
| 7 | IRF8 | interferon regulatory factor 8 | 434 | 0.115 | 0.1160 | No |
| 8 | SSPN | sarcospan | 483 | 0.112 | 0.1246 | No |
| 9 | PELI1 | pellino E3 ubiquitin protein ligase 1 | 593 | 0.104 | 0.1292 | No |
| 10 | CD38 | CD38 molecule | 689 | 0.097 | 0.1340 | No |
| 11 | CDKN1A | cyclin-dependent kinase inhibitor 1A (p21, Cip1) | 690 | 0.097 | 0.1437 | No |
| 12 | MX2 | MX dynamin-like GTPase 2 | 784 | 0.093 | 0.1481 | No |
| 13 | SLAMF7 | SLAM family member 7 | 978 | 0.085 | 0.1466 | No |
| 14 | TNFSF10 | tumor necrosis factor (ligand) superfamily, member 10 | 1006 | 0.084 | 0.1535 | No |
| 15 | IFI27 | interferon, alpha-inducible protein 27 | 1013 | 0.084 | 0.1616 | No |
| 16 | IRF2 | interferon regulatory factor 2 | 1032 | 0.083 | 0.1689 | No |
| 17 | IL4R | interleukin 4 receptor | 1066 | 0.082 | 0.1753 | No |
| 18 | TNFAIP2 | tumor necrosis factor, alpha-induced protein 2 | 1223 | 0.078 | 0.1750 | No |
| 19 | IL7 | interleukin 7 | 1267 | 0.077 | 0.1805 | No |
| 20 | PNP | purine nucleoside phosphorylase | 1405 | 0.074 | 0.1807 | No |
| 21 | P2RY14 | purinergic receptor P2Y, G-protein coupled, 14 | 1605 | 0.069 | 0.1773 | No |
| 22 | ICAM1 | intercellular adhesion molecule 1 | 1606 | 0.069 | 0.1841 | No |
| 23 | MVP | major vault protein | 1742 | 0.066 | 0.1837 | No |
| 24 | TRIM21 | tripartite motif containing 21 | 1810 | 0.065 | 0.1866 | No |
| 25 | LATS2 | large tumor suppressor kinase 2 | 1913 | 0.063 | 0.1876 | No |
| 26 | GBP6 | guanylate binding protein family, member 6 | 2042 | 0.061 | 0.1870 | No |
| 27 | OASL | 2-5-oligoadenylate synthetase-like | 2077 | 0.060 | 0.1913 | No |
| 28 | FCGR1A | Fc fragment of IgG, high affinity Ia, receptor (CD64) | 2126 | 0.060 | 0.1947 | No |
| 29 | SERPING1 | serpin peptidase inhibitor, clade G (C1 inhibitor), member 1 | 2131 | 0.059 | 0.2004 | No |
| 30 | RAPGEF6 | Rap guanine nucleotide exchange factor 6 | 2294 | 0.057 | 0.1977 | No |
| 31 | NUP93 | nucleoporin 93kDa | 2429 | 0.055 | 0.1962 | No |
| 32 | CSF2RB | colony stimulating factor 2 receptor, beta, low-affinity (granulocyte-macrophage) | 2475 | 0.054 | 0.1992 | No |
| 33 | IL15RA | interleukin 15 receptor, alpha | 2659 | 0.051 | 0.1949 | No |
| 34 | PTPN2 | protein tyrosine phosphatase, non-receptor type 2 | 2895 | 0.049 | 0.1876 | No |
| 35 | BST2 | bone marrow stromal cell antigen 2 | 2954 | 0.048 | 0.1893 | No |
| 36 | CMPK2 | cytidine monophosphate (UMP-CMP) kinase 2, mitochondrial | 3051 | 0.047 | 0.1890 | No |
| 37 | IL10RA | interleukin 10 receptor, alpha | 3223 | 0.045 | 0.1847 | No |
| 38 | CFH | complement factor H | 3276 | 0.045 | 0.1865 | No |
| 39 | SECTM1 | secreted and transmembrane 1 | 3545 | 0.042 | 0.1768 | No |
| 40 | IFNAR2 | interferon (alpha, beta and omega) receptor 2 | 3642 | 0.041 | 0.1759 | No |
| 41 | NFKB1 | nuclear factor of kappa light polypeptide gene enhancer in B-cells 1 | 3856 | 0.039 | 0.1688 | No |
| 42 | HELZ2 | helicase with zinc finger 2, transcriptional coactivator | 3945 | 0.038 | 0.1680 | No |
| 43 | SAMHD1 | SAM domain and HD domain 1 | 4186 | 0.036 | 0.1591 | No |
| 44 | TRIM26 | tripartite motif containing 26 | 4319 | 0.035 | 0.1557 | No |
| 45 | TRIM14 | tripartite motif containing 14 | 4463 | 0.033 | 0.1517 | No |
| 46 | CASP3 | caspase 3 | 4699 | 0.032 | 0.1426 | No |
| 47 | PSME1 | proteasome activator subunit 1 | 4737 | 0.031 | 0.1438 | No |
| 48 | OGFR | opioid growth factor receptor | 4790 | 0.031 | 0.1442 | No |
| 49 | RNF213 | ring finger protein 213 | 4842 | 0.030 | 0.1446 | No |
| 50 | ISOC1 | isochorismatase domain containing 1 | 5132 | 0.028 | 0.1325 | No |
| 51 | LCP2 | lymphocyte cytosolic protein 2 | 5232 | 0.027 | 0.1301 | No |
| 52 | ARID5B | AT rich interactive domain 5B (MRF1-like) | 5331 | 0.027 | 0.1276 | No |
| 53 | CCL2 | chemokine (C-C motif) ligand 2 | 5479 | 0.026 | 0.1226 | No |
| 54 | PNPT1 | polyribonucleotide nucleotidyltransferase 1 | 5539 | 0.025 | 0.1220 | No |
| 55 | CD86 | CD86 molecule | 5650 | 0.024 | 0.1188 | No |
| 56 | PLSCR1 | phospholipid scramblase 1 | 5829 | 0.023 | 0.1119 | No |
| 57 | CD74 | CD74 molecule, major histocompatibility complex, class II invariant chain | 5852 | 0.023 | 0.1130 | No |
| 58 | SLC25A28 | solute carrier family 25 (mitochondrial iron transporter), member 28 | 5910 | 0.023 | 0.1123 | No |
| 59 | IFI44L | interferon-induced protein 44-like | 5931 | 0.022 | 0.1135 | No |
| 60 | HLA-A | major histocompatibility complex, class I, A | 5955 | 0.022 | 0.1145 | No |
| 61 | CD274 | CD274 molecule | 6035 | 0.022 | 0.1126 | No |
| 62 | HLA-DQA1 | major histocompatibility complex, class II, DQ alpha 1 | 6155 | 0.021 | 0.1085 | No |
| 63 | TAP1 | transporter 1, ATP-binding cassette, sub-family B (MDR/TAP) | 6169 | 0.021 | 0.1100 | No |
| 64 | MT2A | metallothionein 2A | 6247 | 0.021 | 0.1080 | No |
| 65 | CXCL11 | chemokine (C-X-C motif) ligand 11 | 6249 | 0.021 | 0.1100 | No |
| 66 | HLA-G | major histocompatibility complex, class I, G | 6286 | 0.020 | 0.1102 | No |
| 67 | RTP4 | receptor (chemosensory) transporter protein 4 | 6759 | 0.017 | 0.0875 | No |
| 68 | CASP7 | caspase 7 | 6811 | 0.017 | 0.0865 | No |
| 69 | B2M | beta-2-microglobulin | 7058 | 0.015 | 0.0753 | No |
| 70 | IRF4 | interferon regulatory factor 4 | 7069 | 0.015 | 0.0763 | No |
| 71 | CCL7 | chemokine (C-C motif) ligand 7 | 7135 | 0.015 | 0.0745 | No |
| 72 | IFITM2 | interferon induced transmembrane protein 2 | 7142 | 0.015 | 0.0757 | No |
| 73 | BANK1 | B-cell scaffold protein with ankyrin repeats 1 | 7400 | 0.014 | 0.0637 | No |
| 74 | IFITM3 | interferon induced transmembrane protein 3 | 7579 | 0.012 | 0.0557 | No |
| 75 | RNF31 | ring finger protein 31 | 7656 | 0.012 | 0.0530 | No |
| 76 | BTG1 | B-cell translocation gene 1, anti-proliferative | 7702 | 0.012 | 0.0518 | No |
| 77 | PSMA3 | proteasome subunit alpha 3 | 8134 | 0.009 | 0.0304 | No |
| 78 | PSMB9 | proteasome subunit beta 9 | 8144 | 0.009 | 0.0309 | No |
| 79 | VAMP5 | vesicle associated membrane protein 5 | 8263 | 0.008 | 0.0256 | No |
| 80 | CXCL10 | chemokine (C-X-C motif) ligand 10 | 8286 | 0.008 | 0.0252 | No |
| 81 | PFKP | phosphofructokinase, platelet | 8326 | 0.008 | 0.0240 | No |
| 82 | PSME2 | proteasome activator subunit 2 | 8352 | 0.008 | 0.0235 | No |
| 83 | TOR1B | torsin family 1, member B (torsin B) | 8438 | 0.007 | 0.0198 | No |
| 84 | SOD2 | superoxide dismutase 2, mitochondrial | 8450 | 0.007 | 0.0199 | No |
| 85 | METTL7B | methyltransferase like 7B | 8550 | 0.007 | 0.0155 | No |
| 86 | CCL5 | chemokine (C-C motif) ligand 5 | 8596 | 0.006 | 0.0138 | No |
| 87 | RSAD2 | radical S-adenosyl methionine domain containing 2 | 8715 | 0.006 | 0.0082 | No |
| 88 | LY6E | lymphocyte antigen 6 complex, locus E | 8909 | 0.005 | -0.0013 | No |
| 89 | C1R | complement component 1, r subcomponent | 9028 | 0.004 | -0.0070 | No |
| 90 | HLA-B | major histocompatibility complex, class I, B | 9206 | 0.003 | -0.0159 | No |
| 91 | LGALS3BP | lectin, galactoside-binding, soluble, 3 binding protein | 9239 | 0.003 | -0.0173 | No |
| 92 | CD40 | CD40 molecule, TNF receptor superfamily member 5 | 9416 | 0.002 | -0.0262 | No |
| 93 | CXCL9 | chemokine (C-X-C motif) ligand 9 | 9517 | 0.001 | -0.0312 | No |
| 94 | NFKBIA | nuclear factor of kappa light polypeptide gene enhancer in B-cells inhibitor, alpha | 9745 | 0.000 | -0.0429 | No |
| 95 | TRIM25 | tripartite motif containing 25 | 9786 | -0.000 | -0.0450 | No |
| 96 | NOD1 | nucleotide-binding oligomerization domain containing 1 | 9844 | -0.000 | -0.0479 | No |
| 97 | PSMB2 | proteasome subunit beta 2 | 9881 | -0.001 | -0.0497 | No |
| 98 | IL2RB | interleukin 2 receptor, beta | 10056 | -0.001 | -0.0586 | No |
| 99 | CIITA | class II, major histocompatibility complex, transactivator | 10088 | -0.002 | -0.0600 | No |
| 100 | IRF5 | interferon regulatory factor 5 | 10108 | -0.002 | -0.0608 | No |
| 101 | SELP | selectin P (granule membrane protein 140kDa, antigen CD62) | 10228 | -0.002 | -0.0668 | No |
| 102 | FPR1 | formyl peptide receptor 1 | 10329 | -0.003 | -0.0716 | No |
| 103 | DDX60 | DEAD (Asp-Glu-Ala-Asp) box polypeptide 60 | 10356 | -0.003 | -0.0726 | No |
| 104 | MYD88 | myeloid differentiation primary response 88 | 10440 | -0.004 | -0.0766 | No |
| 105 | XCL1 | chemokine (C motif) ligand 1 | 10695 | -0.005 | -0.0892 | No |
| 106 | SOCS3 | suppressor of cytokine signaling 3 | 10714 | -0.005 | -0.0896 | No |
| 107 | MX1 | MX dynamin-like GTPase 1 | 10754 | -0.006 | -0.0910 | No |
| 108 | PML | promyelocytic leukemia | 10973 | -0.007 | -0.1016 | No |
| 109 | IFIT2 | interferon-induced protein with tetratricopeptide repeats 2 | 11183 | -0.008 | -0.1116 | No |
| 110 | PIM1 | Pim-1 proto-oncogene, serine/threonine kinase | 11480 | -0.010 | -0.1259 | No |
| 111 | CASP8 | caspase 8, apoptosis-related cysteine peptidase | 11632 | -0.011 | -0.1327 | No |
| 112 | OAS3 | 2-5-oligoadenylate synthetase 3 | 11646 | -0.011 | -0.1323 | No |
| 113 | OAS2 | 2-5-oligoadenylate synthetase 2 | 11677 | -0.011 | -0.1328 | No |
| 114 | IL6 | interleukin 6 | 11825 | -0.012 | -0.1392 | No |
| 115 | ZBP1 | Z-DNA binding protein 1 | 11855 | -0.012 | -0.1395 | No |
| 116 | IFI35 | interferon-induced protein 35 | 11915 | -0.012 | -0.1414 | No |
| 117 | PSMA2 | proteasome subunit alpha 2 | 11937 | -0.012 | -0.1412 | No |
| 118 | PARP14 | poly(ADP-ribose) polymerase family member 14 | 12214 | -0.014 | -0.1541 | No |
| 119 | LAP3 | leucine aminopeptidase 3 | 12270 | -0.015 | -0.1555 | No |
| 120 | BATF2 | basic leucine zipper transcription factor, ATF-like 2 | 12411 | -0.016 | -0.1612 | No |
| 121 | CMKLR1 | chemerin chemokine-like receptor 1 | 12531 | -0.016 | -0.1657 | No |
| 122 | ISG15 | ISG15 ubiquitin-like modifier | 12602 | -0.017 | -0.1677 | No |
| 123 | ADAR | adenosine deaminase, RNA-specific | 12610 | -0.017 | -0.1664 | No |
| 124 | SOCS1 | suppressor of cytokine signaling 1 | 12611 | -0.017 | -0.1647 | No |
| 125 | RIPK2 | receptor-interacting serine-threonine kinase 2 | 12627 | -0.017 | -0.1638 | No |
| 126 | PSMB8 | proteasome subunit beta 8 | 12759 | -0.018 | -0.1688 | No |
| 127 | GCH1 | GTP cyclohydrolase 1 | 12793 | -0.018 | -0.1688 | No |
| 128 | STAT1 | signal transducer and activator of transcription 1 | 12830 | -0.018 | -0.1688 | No |
| 129 | JAK2 | Janus kinase 2 | 13031 | -0.019 | -0.1772 | No |
| 130 | ZNFX1 | zinc finger, NFX1-type containing 1 | 13292 | -0.021 | -0.1886 | No |
| 131 | IL18BP | interleukin 18 binding protein | 13325 | -0.021 | -0.1881 | No |
| 132 | CD69 | CD69 molecule | 13361 | -0.022 | -0.1878 | No |
| 133 | TNFAIP3 | tumor necrosis factor, alpha-induced protein 3 | 13363 | -0.022 | -0.1857 | No |
| 134 | VAMP8 | vesicle associated membrane protein 8 | 13409 | -0.022 | -0.1858 | No |
| 135 | C1S | complement component 1, s subcomponent | 13614 | -0.023 | -0.1940 | No |
| 136 | IFIT3 | interferon-induced protein with tetratricopeptide repeats 3 | 13619 | -0.023 | -0.1919 | No |
| 137 | PTPN6 | protein tyrosine phosphatase, non-receptor type 6 | 13725 | -0.024 | -0.1949 | No |
| 138 | TAPBP | TAP binding protein (tapasin) | 14083 | -0.027 | -0.2107 | No |
| 139 | SPPL2A | signal peptide peptidase like 2A | 14124 | -0.027 | -0.2101 | No |
| 140 | STAT3 | signal transducer and activator of transcription 3 (acute-phase response factor) | 14182 | -0.028 | -0.2103 | No |
| 141 | GBP4 | guanylate binding protein 4 | 14250 | -0.028 | -0.2109 | No |
| 142 | VCAM1 | vascular cell adhesion molecule 1 | 14397 | -0.030 | -0.2155 | No |
| 143 | SRI | sorcin | 14400 | -0.030 | -0.2127 | No |
| 144 | CMTR1 | cap methyltransferase 1 | 14622 | -0.031 | -0.2210 | No |
| 145 | NCOA3 | nuclear receptor coactivator 3 | 14803 | -0.033 | -0.2271 | No |
| 146 | HLA-DRB1 | major histocompatibility complex, class II, DR beta 1 | 14826 | -0.033 | -0.2249 | No |
| 147 | SAMD9L | sterile alpha motif domain containing 9-like | 14841 | -0.033 | -0.2224 | No |
| 148 | DDX58 | DEAD (Asp-Glu-Ala-Asp) box polypeptide 58 | 14926 | -0.034 | -0.2233 | No |
| 149 | HERC6 | HECT and RLD domain containing E3 ubiquitin protein ligase family member 6 | 14995 | -0.034 | -0.2234 | No |
| 150 | PSMB10 | proteasome subunit beta 10 | 15143 | -0.036 | -0.2275 | No |
| 151 | ITGB7 | integrin beta 7 | 15643 | -0.040 | -0.2493 | No |
| 152 | BPGM | 2,3-bisphosphoglycerate mutase | 15645 | -0.040 | -0.2454 | No |
| 153 | RIPK1 | receptor (TNFRSF)-interacting serine-threonine kinase 1 | 16084 | -0.045 | -0.2636 | No |
| 154 | TXNIP | thioredoxin interacting protein | 16097 | -0.045 | -0.2597 | No |
| 155 | USP18 | ubiquitin specific peptidase 18 | 16182 | -0.046 | -0.2595 | No |
| 156 | SP110 | SP110 nuclear body protein | 16263 | -0.047 | -0.2590 | No |
| 157 | ISG20 | interferon stimulated exonuclease gene 20kDa | 16332 | -0.047 | -0.2578 | No |
| 158 | IFIH1 | interferon induced, with helicase C domain 1 | 16404 | -0.048 | -0.2567 | No |
| 159 | EIF2AK2 | eukaryotic translation initiation factor 2-alpha kinase 2 | 16462 | -0.049 | -0.2548 | No |
| 160 | ST3GAL5 | ST3 beta-galactoside alpha-2,3-sialyltransferase 5 | 16581 | -0.051 | -0.2558 | No |
| 161 | NAMPT | nicotinamide phosphoribosyltransferase | 16683 | -0.052 | -0.2558 | No |
| 162 | RBCK1 | RanBP-type and C3HC4-type zinc finger containing 1 | 16750 | -0.053 | -0.2539 | No |
| 163 | HIF1A | hypoxia inducible factor 1, alpha subunit (basic helix-loop-helix transcription factor) | 16936 | -0.056 | -0.2580 | No |
| 164 | EPSTI1 | epithelial stromal interaction 1 (breast) | 17199 | -0.060 | -0.2656 | No |
| 165 | PTPN1 | protein tyrosine phosphatase, non-receptor type 1 | 17307 | -0.062 | -0.2649 | No |
| 166 | TDRD7 | tudor domain containing 7 | 17494 | -0.065 | -0.2681 | No |
| 167 | IDO1 | indoleamine 2,3-dioxygenase 1 | 17540 | -0.066 | -0.2639 | No |
| 168 | TRAFD1 | TRAF-type zinc finger domain containing 1 | 17666 | -0.069 | -0.2635 | No |
| 169 | GPR18 | G protein-coupled receptor 18 | 17941 | -0.076 | -0.2702 | Yes |
| 170 | IFIT1 | interferon-induced protein with tetratricopeptide repeats 1 | 17946 | -0.076 | -0.2628 | Yes |
| 171 | STAT4 | signal transducer and activator of transcription 4 | 17951 | -0.076 | -0.2555 | Yes |
| 172 | AUTS2 | autism susceptibility candidate 2 | 17980 | -0.076 | -0.2494 | Yes |
| 173 | PARP12 | poly(ADP-ribose) polymerase family member 12 | 18125 | -0.081 | -0.2488 | Yes |
| 174 | TNFAIP6 | tumor necrosis factor, alpha-induced protein 6 | 18183 | -0.082 | -0.2436 | Yes |
| 175 | MTHFD2 | methylenetetrahydrofolate dehydrogenase (NADP+ dependent) 2, methenyltetrahydrofolate cyclohydrolase | 18255 | -0.084 | -0.2389 | Yes |
| 176 | UBE2L6 | ubiquitin-conjugating enzyme E2L 6 | 18293 | -0.085 | -0.2323 | Yes |
| 177 | UPP1 | uridine phosphorylase 1 | 18404 | -0.089 | -0.2292 | Yes |
| 178 | DHX58 | DEXH (Asp-Glu-X-His) box polypeptide 58 | 18439 | -0.090 | -0.2220 | Yes |
| 179 | FGL2 | fibrinogen-like 2 | 18506 | -0.093 | -0.2162 | Yes |
| 180 | STAT2 | signal transducer and activator of transcription 2 | 18572 | -0.095 | -0.2101 | Yes |
| 181 | XAF1 | XIAP associated factor 1 | 18584 | -0.096 | -0.2011 | Yes |
| 182 | NMI | N-myc (and STAT) interactor | 18617 | -0.097 | -0.1931 | Yes |
| 183 | IRF1 | interferon regulatory factor 1 | 18746 | -0.106 | -0.1892 | Yes |
| 184 | IRF9 | interferon regulatory factor 9 | 18860 | -0.114 | -0.1837 | Yes |
| 185 | LYSMD2 | LysM, putative peptidoglycan-binding, domain containing 2 | 18959 | -0.123 | -0.1766 | Yes |
| 186 | CASP4 | caspase 4 | 19027 | -0.131 | -0.1671 | Yes |
| 187 | PDE4B | phosphodiesterase 4B, cAMP-specific | 19061 | -0.135 | -0.1553 | Yes |
| 188 | NLRC5 | NLR family, CARD domain containing 5 | 19109 | -0.143 | -0.1436 | Yes |
| 189 | ST8SIA4 | ST8 alpha-N-acetyl-neuraminide alpha-2,8-sialyltransferase 4 | 19177 | -0.154 | -0.1317 | Yes |
| 190 | APOL6 | apolipoprotein L, 6 | 19187 | -0.158 | -0.1165 | Yes |
| 191 | ARL4A | ADP-ribosylation factor like GTPase 4A | 19243 | -0.172 | -0.1022 | Yes |
| 192 | IFI44 | interferon-induced protein 44 | 19319 | -0.197 | -0.0865 | Yes |
| 193 | GZMA | granzyme A | 19327 | -0.200 | -0.0670 | Yes |
| 194 | PTGS2 | prostaglandin-endoperoxide synthase 2 (prostaglandin G/H synthase and cyclooxygenase) | 19364 | -0.218 | -0.0472 | Yes |
| 195 | PLA2G4A | phospholipase A2, group IVA (cytosolic, calcium-dependent) | 19411 | -0.248 | -0.0249 | Yes |
| 196 | CASP1 | caspase 1 | 19457 | -0.309 | 0.0035 | Yes |
Table: GSEA details [plain text format]

  

Fig 2: HALLMARK\_INTERFERON\_GAMMA\_RESPONSE      
 Blue-Pink O' Gram in the Space of the Analyzed GeneSet

  

Fig 3: HALLMARK\_INTERFERON\_GAMMA\_RESPONSE: Random ES distribution      
 Gene set null distribution of ES for **HALLMARK\_INTERFERON\_GAMMA\_RESPONSE**

  
