## Supplementary material for "Somatic hypomethylation of pericentromeric SST1 repeats and tetraploidization in human colorectal cancer cells": GSEA results: HALLMARK_KRAS_SIGNALING_DN.html

Details for gene set HALLMARK\_KRAS\_SIGNALING\_DN[GSEA]

|  || Dataset | eset\_byprobe\_collapsed\_to\_symbols.Diploid\_vs\_Tetraploid.cls #Tetraploid\_versus\_Diploid.Diploid\_vs\_Tetraploid.cls #Tetraploid\_versus\_Diploid\_repos |
| Phenotype | Diploid\_vs\_Tetraploid.cls#Tetraploid\_versus\_Diploid\_repos |
| Upregulated in class | Tetraploid |
| GeneSet | HALLMARK\_KRAS\_SIGNALING\_DN |
| Enrichment Score (ES) | 0.27305692 |
| Normalized Enrichment Score (NES) | 1.2398108 |
| Nominal p-value | 0.038674034 |
| FDR q-value | 0.18521672 |
| FWER p-Value | 0.82 |
Table: GSEA Results Summary

  

Fig 1: Enrichment plot: HALLMARK\_KRAS\_SIGNALING\_DN      
 Profile of the Running ES Score & Positions of GeneSet Members on the Rank Ordered List

  

| SYMBOL | TITLE | RANK IN GENE LIST | RANK METRIC SCORE | RUNNING ES | CORE ENRICHMENT || 1 | SLC6A14 | solute carrier family 6 (amino acid transporter), member 14 | 26 | 0.352 | 0.0352 | Yes |
| 2 | KRT4 | keratin 4, type II | 63 | 0.266 | 0.0611 | Yes |
| 3 | HTR1D | 5-hydroxytryptamine (serotonin) receptor 1D, G protein-coupled | 98 | 0.210 | 0.0812 | Yes |
| 4 | SIDT1 | SID1 transmembrane family, member 1 | 100 | 0.209 | 0.1028 | Yes |
| 5 | NPY4R | Homo sapiens neuropeptide Y receptor Y4 (NPY4R), transcript variant 2, mRNA. | 125 | 0.187 | 0.1211 | Yes |
| 6 | THRB | thyroid hormone receptor, beta | 181 | 0.162 | 0.1351 | Yes |
| 7 | RIBC2 | RIB43A domain with coiled-coils 2 | 189 | 0.159 | 0.1513 | Yes |
| 8 | DTNB | dystrobrevin beta | 279 | 0.136 | 0.1608 | Yes |
| 9 | BTG2 | BTG family, member 2 | 281 | 0.136 | 0.1749 | Yes |
| 10 | SLC5A5 | solute carrier family 5 (sodium/iodide cotransporter), member 5 | 345 | 0.126 | 0.1847 | Yes |
| 11 | EDN1 | endothelin 1 | 371 | 0.123 | 0.1962 | Yes |
| 12 | NR4A2 | nuclear receptor subfamily 4, group A, member 2 | 499 | 0.110 | 0.2011 | Yes |
| 13 | PAX4 | paired box 4 | 560 | 0.106 | 0.2089 | Yes |
| 14 | RGS11 | regulator of G-protein signaling 11 | 632 | 0.101 | 0.2158 | Yes |
| 15 | ABCG4 | ATP binding cassette subfamily G member 4 | 791 | 0.093 | 0.2172 | Yes |
| 16 | PCDHB1 | protocadherin beta 1 | 820 | 0.091 | 0.2253 | Yes |
| 17 | CNTFR | ciliary neurotrophic factor receptor | 920 | 0.087 | 0.2293 | Yes |
| 18 | GP1BA | glycoprotein Ib (platelet), alpha polypeptide | 1000 | 0.084 | 0.2339 | Yes |
| 19 | ITIH3 | inter-alpha-trypsin inhibitor heavy chain 3 | 1033 | 0.083 | 0.2409 | Yes |
| 20 | PTGFR | prostaglandin F receptor (FP) | 1110 | 0.081 | 0.2454 | Yes |
| 21 | ZC2HC1C | zinc finger, C2HC-type containing 1C | 1230 | 0.078 | 0.2473 | Yes |
| 22 | KCNN1 | potassium channel, calcium activated intermediate/small conductance subfamily N alpha, member 1 | 1247 | 0.077 | 0.2545 | Yes |
| 23 | MSH5 | mutS homolog 5 | 1384 | 0.074 | 0.2552 | Yes |
| 24 | SPRR3 | small proline-rich protein 3 | 1551 | 0.070 | 0.2539 | Yes |
| 25 | IFNG | interferon, gamma | 1797 | 0.065 | 0.2480 | Yes |
| 26 | ATP6V1B1 | ATPase, H+ transporting, lysosomal 56/58kDa, V1 subunit B1 | 1823 | 0.064 | 0.2534 | Yes |
| 27 | KLK7 | kallikrein related peptidase 7 | 1856 | 0.064 | 0.2584 | Yes |
| 28 | FGFR3 | fibroblast growth factor receptor 3 | 1953 | 0.062 | 0.2599 | Yes |
| 29 | KRT13 | keratin 13, type I | 2215 | 0.058 | 0.2524 | Yes |
| 30 | MYOT | myotilin | 2247 | 0.058 | 0.2568 | Yes |
| 31 | CLDN8 | claudin 8 | 2308 | 0.057 | 0.2596 | Yes |
| 32 | BARD1 | BRCA1 associated RING domain 1 | 2585 | 0.053 | 0.2507 | Yes |
| 33 | SLC16A7 | solute carrier family 16 (monocarboxylate transporter), member 7 | 2682 | 0.051 | 0.2511 | Yes |
| 34 | CCR8 | chemokine (C-C motif) receptor 8 | 2705 | 0.051 | 0.2553 | Yes |
| 35 | HNF1A | HNF1 homeobox A | 2751 | 0.051 | 0.2582 | Yes |
| 36 | LGALS7 | lectin, galactoside-binding, soluble, 7 | 2820 | 0.050 | 0.2598 | Yes |
| 37 | EGF | epidermal growth factor | 2875 | 0.049 | 0.2621 | Yes |
| 38 | EDN2 | endothelin 2 | 2912 | 0.048 | 0.2653 | Yes |
| 39 | CACNA1F | calcium channel, voltage-dependent, L type, alpha 1F subunit | 3167 | 0.046 | 0.2569 | Yes |
| 40 | P2RY4 | pyrimidinergic receptor P2Y, G-protein coupled, 4 | 3186 | 0.046 | 0.2607 | Yes |
| 41 | TSHB | thyroid stimulating hormone, beta | 3228 | 0.045 | 0.2633 | Yes |
| 42 | PTPRJ | protein tyrosine phosphatase, receptor type, J | 3248 | 0.045 | 0.2670 | Yes |
| 43 | CAPN9 | calpain 9 | 3271 | 0.045 | 0.2705 | Yes |
| 44 | PRODH | Homo sapiens proline dehydrogenase (oxidase) 1 (PRODH), transcript variant 2, mRNA. | 3313 | 0.045 | 0.2731 | Yes |
| 45 | FGF22 | fibroblast growth factor 22 | 3480 | 0.043 | 0.2689 | No |
| 46 | DCC | DCC netrin 1 receptor | 3629 | 0.041 | 0.2656 | No |
| 47 | TGFB2 | transforming growth factor beta 2 | 3684 | 0.041 | 0.2670 | No |
| 48 | SPHK2 | sphingosine kinase 2 | 3765 | 0.040 | 0.2670 | No |
| 49 | KCNMB1 | potassium channel subfamily M regulatory beta subunit 1 | 3781 | 0.040 | 0.2704 | No |
| 50 | SLC25A23 | solute carrier family 25 (mitochondrial carrier; phosphate carrier), member 23 | 3888 | 0.039 | 0.2689 | No |
| 51 | CD207 | CD207 molecule, langerin | 4009 | 0.037 | 0.2666 | No |
| 52 | ARPP21 | cAMP-regulated phosphoprotein 21kDa | 4161 | 0.036 | 0.2625 | No |
| 53 | SNN | stannin | 4231 | 0.035 | 0.2626 | No |
| 54 | VPREB1 | pre-B lymphocyte 1 | 4341 | 0.034 | 0.2606 | No |
| 55 | BRDT | bromodomain, testis-specific | 4486 | 0.033 | 0.2566 | No |
| 56 | SCN10A | sodium channel, voltage gated, type X alpha subunit | 4546 | 0.033 | 0.2569 | No |
| 57 | IRS4 | insulin receptor substrate 4 | 4582 | 0.032 | 0.2585 | No |
| 58 | TFAP2B | transcription factor AP-2 beta (activating enhancer binding protein 2 beta) | 4606 | 0.032 | 0.2607 | No |
| 59 | KLHDC8A | kelch domain containing 8A | 4766 | 0.031 | 0.2557 | No |
| 60 | SCGB1A1 | secretoglobin, family 1A, member 1 (uteroglobin) | 4791 | 0.031 | 0.2576 | No |
| 61 | CALCB | calcitonin-related polypeptide beta | 4863 | 0.030 | 0.2571 | No |
| 62 | COPZ2 | coatomer protein complex subunit zeta 2 | 4946 | 0.030 | 0.2560 | No |
| 63 | RYR1 | ryanodine receptor 1 (skeletal) | 4983 | 0.029 | 0.2571 | No |
| 64 | CHRNG | cholinergic receptor, nicotinic gamma | 5104 | 0.029 | 0.2539 | No |
| 65 | EFHD1 | EF-hand domain family member D1 | 5159 | 0.028 | 0.2540 | No |
| 66 | SOX10 | SRY box 10 | 5362 | 0.027 | 0.2463 | No |
| 67 | THNSL2 | threonine synthase-like 2 | 5421 | 0.026 | 0.2461 | No |
| 68 | SLC38A3 | solute carrier family 38, member 3 | 5646 | 0.024 | 0.2370 | No |
| 69 | KCNQ2 | potassium channel, voltage gated KQT-like subfamily Q, member 2 | 5656 | 0.024 | 0.2391 | No |
| 70 | GTF3C5 | general transcription factor IIIC subunit 5 | 5714 | 0.024 | 0.2386 | No |
| 71 | NPHS1 | nephrosis 1, congenital, Finnish type (nephrin) | 5715 | 0.024 | 0.2411 | No |
| 72 | IFI44L | interferon-induced protein 44-like | 5931 | 0.022 | 0.2323 | No |
| 73 | CHST2 | carbohydrate (N-acetylglucosamine-6-O) sulfotransferase 2 | 6022 | 0.022 | 0.2299 | No |
| 74 | SERPINB2 | serpin peptidase inhibitor, clade B (ovalbumin), member 2 | 6047 | 0.022 | 0.2310 | No |
| 75 | PNMT | phenylethanolamine N-methyltransferase | 6164 | 0.021 | 0.2271 | No |
| 76 | KLK8 | kallikrein related peptidase 8 | 6224 | 0.021 | 0.2262 | No |
| 77 | SGK1 | serum/glucocorticoid regulated kinase 1 | 6290 | 0.020 | 0.2250 | No |
| 78 | COL2A1 | collagen, type II, alpha 1 | 6341 | 0.020 | 0.2245 | No |
| 79 | SLC6A3 | solute carrier family 6 (neurotransmitter transporter), member 3 | 6367 | 0.020 | 0.2253 | No |
| 80 | CLPS | colipase, pancreatic | 7015 | 0.016 | 0.1934 | No |
| 81 | KRT5 | keratin 5, type II | 7175 | 0.015 | 0.1867 | No |
| 82 | ZBTB16 | zinc finger and BTB domain containing 16 | 7433 | 0.013 | 0.1748 | No |
| 83 | NTF3 | neurotrophin 3 | 7543 | 0.013 | 0.1705 | No |
| 84 | NRIP2 | nuclear receptor interacting protein 2 | 7578 | 0.012 | 0.1700 | No |
| 85 | HSD11B2 | hydroxysteroid (11-beta) dehydrogenase 2 | 7753 | 0.011 | 0.1622 | No |
| 86 | GPRC5C | G protein-coupled receptor, class C, group 5, member C | 7760 | 0.011 | 0.1631 | No |
| 87 | TCL1A | T-cell leukemia/lymphoma 1A | 7777 | 0.011 | 0.1634 | No |
| 88 | EPHA5 | EPH receptor A5 | 8023 | 0.010 | 0.1518 | No |
| 89 | SYNPO | synaptopodin | 8121 | 0.009 | 0.1477 | No |
| 90 | DLK2 | delta-like 2 homolog (Drosophila) | 8230 | 0.008 | 0.1430 | No |
| 91 | MYH7 | myosin, heavy chain 7, cardiac muscle, beta | 8257 | 0.008 | 0.1425 | No |
| 92 | OXT | oxytocin/neurophysin I prepropeptide | 8357 | 0.008 | 0.1382 | No |
| 93 | TCF7L1 | transcription factor 7-like 1 (T-cell specific, HMG-box) | 8662 | 0.006 | 0.1231 | No |
| 94 | RSAD2 | radical S-adenosyl methionine domain containing 2 | 8715 | 0.006 | 0.1210 | No |
| 95 | ASB7 | ankyrin repeat and SOCS box containing 7 | 8803 | 0.005 | 0.1170 | No |
| 96 | MAST3 | microtubule associated serine/threonine kinase 3 | 9088 | 0.004 | 0.1027 | No |
| 97 | SSTR4 | somatostatin receptor 4 | 9182 | 0.003 | 0.0982 | No |
| 98 | TGM1 | transglutaminase 1 | 9354 | 0.002 | 0.0896 | No |
| 99 | NR6A1 | nuclear receptor subfamily 6, group A, member 1 | 9516 | 0.001 | 0.0814 | No |
| 100 | TEX15 | testis expressed 15 | 9574 | 0.001 | 0.0785 | No |
| 101 | CALML5 | calmodulin-like 5 | 9667 | 0.001 | 0.0738 | No |
| 102 | CPA2 | carboxypeptidase A2 (pancreatic) | 9669 | 0.001 | 0.0739 | No |
| 103 | FSHB | follicle stimulating hormone, beta polypeptide | 9790 | -0.000 | 0.0677 | No |
| 104 | CACNG1 | calcium channel, voltage-dependent, gamma subunit 1 | 9810 | -0.000 | 0.0667 | No |
| 105 | FGF16 | fibroblast growth factor 16 | 9815 | -0.000 | 0.0665 | No |
| 106 | STAG3 | stromal antigen 3 | 9831 | -0.000 | 0.0658 | No |
| 107 | CDH16 | cadherin 16, KSP-cadherin | 10337 | -0.003 | 0.0400 | No |
| 108 | SPTBN2 | spectrin, beta, non-erythrocytic 2 | 10405 | -0.004 | 0.0369 | No |
| 109 | PDCD1 | programmed cell death 1 | 10491 | -0.004 | 0.0329 | No |
| 110 | TNNI3 | troponin I type 3 (cardiac) | 10545 | -0.004 | 0.0306 | No |
| 111 | KCND1 | potassium channel, voltage gated Shal related subfamily D, member 1 | 10599 | -0.005 | 0.0284 | No |
| 112 | NGB | neuroglobin | 10689 | -0.005 | 0.0243 | No |
| 113 | MX1 | MX dynamin-like GTPase 1 | 10754 | -0.006 | 0.0216 | No |
| 114 | SNCB | synuclein beta | 10952 | -0.007 | 0.0121 | No |
| 115 | ATP4A | ATPase, H+/K+ exchanging, alpha polypeptide | 11105 | -0.008 | 0.0050 | No |
| 116 | FGGY | FGGY carbohydrate kinase domain containing | 11125 | -0.008 | 0.0049 | No |
| 117 | CD80 | CD80 molecule | 11264 | -0.009 | -0.0014 | No |
| 118 | KRT15 | keratin 15, type I | 11281 | -0.009 | -0.0013 | No |
| 119 | SHOX2 | short stature homeobox 2 | 11484 | -0.010 | -0.0107 | No |
| 120 | ZNF112 | zinc finger protein 112 | 11693 | -0.011 | -0.0204 | No |
| 121 | CELSR2 | cadherin, EGF LAG seven-pass G-type receptor 2 | 12052 | -0.013 | -0.0375 | No |
| 122 | ABCB11 | ATP binding cassette subfamily B member 11 | 12174 | -0.014 | -0.0423 | No |
| 123 | CYP11B2 | cytochrome P450, family 11, subfamily B, polypeptide 2 | 12210 | -0.014 | -0.0427 | No |
| 124 | CD40LG | CD40 ligand | 12277 | -0.015 | -0.0445 | No |
| 125 | GDNF | glial cell derived neurotrophic factor | 12324 | -0.015 | -0.0454 | No |
| 126 | HTR1B | 5-hydroxytryptamine (serotonin) receptor 1B, G protein-coupled | 12326 | -0.015 | -0.0439 | No |
| 127 | UPK3B | uroplakin 3B | 12523 | -0.016 | -0.0523 | No |
| 128 | MYO15A | myosin XVA | 12525 | -0.016 | -0.0507 | No |
| 129 | SLC29A3 | solute carrier family 29 (equilibrative nucleoside transporter), member 3 | 12609 | -0.017 | -0.0532 | No |
| 130 | CPB1 | carboxypeptidase B1 (tissue) | 12986 | -0.019 | -0.0707 | No |
| 131 | CAMK1D | calcium/calmodulin-dependent protein kinase ID | 13122 | -0.020 | -0.0756 | No |
| 132 | MEFV | Mediterranean fever | 13146 | -0.020 | -0.0747 | No |
| 133 | NOS1 | nitric oxide synthase 1 (neuronal) | 13175 | -0.020 | -0.0740 | No |
| 134 | IL5 | interleukin 5 | 13284 | -0.021 | -0.0774 | No |
| 135 | CDKAL1 | CDK5 regulatory subunit associated protein 1-like 1 | 13533 | -0.023 | -0.0879 | No |
| 136 | SKIL | SKI-like proto-oncogene | 13624 | -0.024 | -0.0901 | No |
| 137 | CCDC106 | coiled-coil domain containing 106 | 13646 | -0.024 | -0.0887 | No |
| 138 | YBX2 | Y box binding protein 2 | 13662 | -0.024 | -0.0870 | No |
| 139 | KMT2D | lysine (K)-specific methyltransferase 2D | 13730 | -0.024 | -0.0880 | No |
| 140 | SERPINA10 | serpin peptidase inhibitor, clade A (alpha-1 antiproteinase, antitrypsin), member 10 | 13741 | -0.024 | -0.0859 | No |
| 141 | MAGIX | MAGI family member, X-linked | 13748 | -0.024 | -0.0837 | No |
| 142 | ACTC1 | actin, alpha, cardiac muscle 1 | 13790 | -0.025 | -0.0832 | No |
| 143 | CCNA1 | cyclin A1 | 13813 | -0.025 | -0.0818 | No |
| 144 | SLC30A3 | solute carrier family 30 (zinc transporter), member 3 | 13839 | -0.025 | -0.0805 | No |
| 145 | LFNG | LFNG O-fucosylpeptide 3-beta-N-acetylglucosaminyltransferase | 14482 | -0.030 | -0.1105 | No |
| 146 | GRID2 | glutamate receptor, ionotropic, delta 2 | 14837 | -0.033 | -0.1254 | No |
| 147 | NUDT11 | nudix hydrolase 11 | 14839 | -0.033 | -0.1220 | No |
| 148 | CLSTN3 | calsyntenin 3 | 15151 | -0.036 | -0.1344 | No |
| 149 | TFF2 | trefoil factor 2 | 15205 | -0.036 | -0.1334 | No |
| 150 | CKM | creatine kinase, muscle | 15324 | -0.037 | -0.1356 | No |
| 151 | PDE6B | phosphodiesterase 6B, cGMP-specific, rod, beta | 15350 | -0.037 | -0.1330 | No |
| 152 | AKR1B10 | aldo-keto reductase family 1, member B10 (aldose reductase) | 15462 | -0.038 | -0.1348 | No |
| 153 | PKP1 | plakophilin 1 | 15607 | -0.040 | -0.1381 | No |
| 154 | IDUA | iduronidase, alpha-L- | 15706 | -0.041 | -0.1389 | No |
| 155 | GPR3 | G protein-coupled receptor 3 | 15712 | -0.041 | -0.1349 | No |
| 156 | UGT2B17 | UDP glucuronosyltransferase 2 family, polypeptide B17 | 15883 | -0.043 | -0.1393 | No |
| 157 | ITGB1BP2 | integrin beta 1 binding protein (melusin) 2 | 16031 | -0.044 | -0.1423 | No |
| 158 | VPS50 | VPS50 EARP/GARPII complex subunit | 16037 | -0.044 | -0.1380 | No |
| 159 | ADRA2C | adrenoceptor alpha 2C | 16225 | -0.046 | -0.1429 | No |
| 160 | LYPD3 | LY6/PLAUR domain containing 3 | 16257 | -0.047 | -0.1396 | No |
| 161 | AMBN | ameloblastin | 16328 | -0.047 | -0.1383 | No |
| 162 | PDK2 | pyruvate dehydrogenase kinase, isozyme 2 | 16370 | -0.048 | -0.1355 | No |
| 163 | P2RX6 | purinergic receptor P2X, ligand gated ion channel, 6 | 16472 | -0.049 | -0.1355 | No |
| 164 | MFSD6 | major facilitator superfamily domain containing 6 | 16550 | -0.051 | -0.1343 | No |
| 165 | TFCP2L1 | transcription factor CP2-like 1 | 16585 | -0.051 | -0.1307 | No |
| 166 | PAX3 | paired box 3 | 16713 | -0.053 | -0.1318 | No |
| 167 | GAMT | guanidinoacetate N-methyltransferase | 16799 | -0.054 | -0.1306 | No |
| 168 | GP2 | glycoprotein 2 (zymogen granule membrane) | 16900 | -0.055 | -0.1300 | No |
| 169 | BMPR1B | bone morphogenetic protein receptor type IB | 16952 | -0.056 | -0.1269 | No |
| 170 | ALOX12B | arachidonate 12-lipoxygenase, 12R type | 17025 | -0.057 | -0.1246 | No |
| 171 | TAS2R4 | taste receptor, type 2, member 4 | 17131 | -0.059 | -0.1239 | No |
| 172 | RYR2 | ryanodine receptor 2 (cardiac) | 17244 | -0.061 | -0.1234 | No |
| 173 | PROP1 | PROP paired-like homeobox 1 | 17260 | -0.061 | -0.1179 | No |
| 174 | KRT1 | keratin 1, type II | 17418 | -0.064 | -0.1193 | No |
| 175 | EDAR | ectodysplasin A receptor | 17532 | -0.066 | -0.1183 | No |
| 176 | WNT16 | wingless-type MMTV integration site family, member 16 | 17628 | -0.068 | -0.1162 | No |
| 177 | KCNE2 | potassium channel, voltage gated subfamily E regulatory beta subunit 2 | 17887 | -0.074 | -0.1218 | No |
| 178 | INSL5 | insulin-like 5 | 17922 | -0.075 | -0.1158 | No |
| 179 | TENM2 | teneurin transmembrane protein 2 | 18022 | -0.077 | -0.1129 | No |
| 180 | C5 | complement component 5 | 18126 | -0.081 | -0.1098 | No |
| 181 | MTHFR | methylenetetrahydrofolate reductase (NAD(P)H) | 18240 | -0.084 | -0.1069 | No |
| 182 | IL12B | interleukin 12B | 18360 | -0.087 | -0.1040 | No |
| 183 | TG | thyroglobulin | 18368 | -0.088 | -0.0953 | No |
| 184 | GPR19 | G protein-coupled receptor 19 | 18408 | -0.089 | -0.0881 | No |
| 185 | CLDN16 | claudin 16 | 18414 | -0.089 | -0.0790 | No |
| 186 | ARHGDIG | Rho GDP dissociation inhibitor (GDI) gamma | 18433 | -0.090 | -0.0706 | No |
| 187 | SMPX | small muscle protein, X-linked | 18498 | -0.093 | -0.0643 | No |
| 188 | TLX1 | T-cell leukemia homeobox 1 | 18577 | -0.096 | -0.0584 | No |
| 189 | SLC12A3 | solute carrier family 12 (sodium/chloride transporter), member 3 | 18814 | -0.111 | -0.0591 | No |
| 190 | ENTPD7 | ectonucleoside triphosphate diphosphohydrolase 7 | 18922 | -0.119 | -0.0523 | No |
| 191 | CPEB3 | Transcript Identified by AceView, Entrez Gene ID(s) 22849 | 19083 | -0.139 | -0.0461 | No |
| 192 | IGFBP2 | insulin like growth factor binding protein 2 | 19180 | -0.155 | -0.0349 | No |
| 193 | CYP39A1 | cytochrome P450, family 39, subfamily A, polypeptide 1 | 19213 | -0.165 | -0.0194 | No |
| 194 | YPEL1 | yippee like 1 | 19218 | -0.165 | -0.0025 | No |
| 195 | PLAG1 | pleiomorphic adenoma gene 1 | 19255 | -0.176 | 0.0139 | No |
Table: GSEA details [plain text format]

  

Fig 2: HALLMARK\_KRAS\_SIGNALING\_DN      
 Blue-Pink O' Gram in the Space of the Analyzed GeneSet

  

Fig 3: HALLMARK\_KRAS\_SIGNALING\_DN: Random ES distribution      
 Gene set null distribution of ES for **HALLMARK\_KRAS\_SIGNALING\_DN**

  
