## Supplementary material for "Somatic hypomethylation of pericentromeric SST1 repeats and tetraploidization in human colorectal cancer cells": GSEA results: HALLMARK_KRAS_SIGNALING_UP.html

Details for gene set HALLMARK\_KRAS\_SIGNALING\_UP[GSEA]

|  || Dataset | eset\_byprobe\_collapsed\_to\_symbols.Diploid\_vs\_Tetraploid.cls #Tetraploid\_versus\_Diploid.Diploid\_vs\_Tetraploid.cls #Tetraploid\_versus\_Diploid\_repos |
| Phenotype | Diploid\_vs\_Tetraploid.cls#Tetraploid\_versus\_Diploid\_repos |
| Upregulated in class | Diploid |
| GeneSet | HALLMARK\_KRAS\_SIGNALING\_UP |
| Enrichment Score (ES) | -0.26733392 |
| Normalized Enrichment Score (NES) | -1.1047868 |
| Nominal p-value | 0.24700598 |
| FDR q-value | 0.37778786 |
| FWER p-Value | 1.0 |
Table: GSEA Results Summary

  

Fig 1: Enrichment plot: HALLMARK\_KRAS\_SIGNALING\_UP      
 Profile of the Running ES Score & Positions of GeneSet Members on the Rank Ordered List

  

| SYMBOL | TITLE | RANK IN GENE LIST | RANK METRIC SCORE | RUNNING ES | CORE ENRICHMENT || 1 | CCL20 | chemokine (C-C motif) ligand 20 | 12 | 0.406 | 0.0347 | No |
| 2 | CCND2 | cyclin D2 | 34 | 0.313 | 0.0609 | No |
| 3 | CA2 | carbonic anhydrase II | 69 | 0.250 | 0.0809 | No |
| 4 | BTC | betacellulin | 130 | 0.186 | 0.0939 | No |
| 5 | CFB | complement factor B | 164 | 0.167 | 0.1068 | No |
| 6 | IL33 | interleukin 33 | 207 | 0.153 | 0.1180 | No |
| 7 | ANO1 | anoctamin 1, calcium activated chloride channel | 263 | 0.139 | 0.1272 | No |
| 8 | IL2RG | interleukin 2 receptor, gamma | 268 | 0.137 | 0.1390 | No |
| 9 | PPBP | pro-platelet basic protein | 416 | 0.117 | 0.1416 | No |
| 10 | LIF | leukemia inhibitory factor | 423 | 0.117 | 0.1514 | No |
| 11 | IRF8 | interferon regulatory factor 8 | 434 | 0.115 | 0.1610 | No |
| 12 | IGFBP3 | insulin like growth factor binding protein 3 | 582 | 0.104 | 0.1624 | No |
| 13 | DUSP6 | dual specificity phosphatase 6 | 638 | 0.101 | 0.1684 | No |
| 14 | FCER1G | Fc fragment of IgE, high affinity I, receptor for; gamma polypeptide | 654 | 0.100 | 0.1763 | No |
| 15 | BTBD3 | BTB (POZ) domain containing 3 | 751 | 0.095 | 0.1796 | No |
| 16 | SLPI | secretory leukocyte peptidase inhibitor | 894 | 0.088 | 0.1799 | No |
| 17 | PRDM1 | PR domain containing 1, with ZNF domain | 898 | 0.088 | 0.1874 | No |
| 18 | EMP1 | epithelial membrane protein 1 | 967 | 0.085 | 0.1914 | No |
| 19 | PTCD2 | pentatricopeptide repeat domain 2 | 1057 | 0.082 | 0.1939 | No |
| 20 | MALL | mal, T-cell differentiation protein-like | 1059 | 0.082 | 0.2010 | No |
| 21 | ITGBL1 | integrin beta like 1 | 1148 | 0.080 | 0.2034 | No |
| 22 | ID2 | inhibitor of DNA binding 2, dominant negative helix-loop-helix protein | 1150 | 0.080 | 0.2103 | No |
| 23 | TNNT2 | troponin T type 2 (cardiac) | 1198 | 0.079 | 0.2147 | No |
| 24 | PLAUR | plasminogen activator, urokinase receptor | 1355 | 0.075 | 0.2132 | No |
| 25 | NRP1 | neuropilin 1 | 1363 | 0.075 | 0.2193 | No |
| 26 | BIRC3 | Memczak2013 ANTISENSE, CDS, coding, INTERNAL best transcript NM\_182962 | 1400 | 0.074 | 0.2239 | No |
| 27 | ACE | angiotensin I converting enzyme | 1448 | 0.073 | 0.2278 | No |
| 28 | SNAP91 | synaptosome associated protein 91kDa | 1590 | 0.069 | 0.2265 | No |
| 29 | CBR4 | carbonyl reductase 4 | 1723 | 0.066 | 0.2255 | No |
| 30 | ST6GAL1 | ST6 beta-galactosamide alpha-2,6-sialyltranferase 1 | 1800 | 0.065 | 0.2272 | No |
| 31 | ANGPTL4 | angiopoietin like 4 | 1873 | 0.064 | 0.2290 | No |
| 32 | KCNN4 | potassium channel, calcium activated intermediate/small conductance subfamily N alpha, member 4 | 2000 | 0.061 | 0.2278 | No |
| 33 | LAPTM5 | lysosomal protein transmembrane 5 | 2124 | 0.060 | 0.2266 | No |
| 34 | GPRC5B | G protein-coupled receptor, class C, group 5, member B | 2159 | 0.059 | 0.2300 | No |
| 35 | SPON1 | spondin 1, extracellular matrix protein | 2271 | 0.057 | 0.2293 | No |
| 36 | RBP4 | retinol binding protein 4, plasma | 2424 | 0.055 | 0.2262 | No |
| 37 | MAP7 | microtubule-associated protein 7 | 2428 | 0.055 | 0.2308 | No |
| 38 | NR1H4 | nuclear receptor subfamily 1, group H, member 4 | 3094 | 0.047 | 0.2005 | No |
| 39 | SCG3 | secretogranin III | 3098 | 0.047 | 0.2044 | No |
| 40 | CLEC4A | C-type lectin domain family 4, member A | 3189 | 0.046 | 0.2037 | No |
| 41 | IL10RA | interleukin 10 receptor, alpha | 3223 | 0.045 | 0.2059 | No |
| 42 | PLVAP | plasmalemma vesicle associated protein | 3264 | 0.045 | 0.2078 | No |
| 43 | CFH | complement factor H | 3276 | 0.045 | 0.2111 | No |
| 44 | IL7R | interleukin 7 receptor | 3650 | 0.041 | 0.1954 | No |
| 45 | VWA5A | von Willebrand factor A domain containing 5A | 3657 | 0.041 | 0.1986 | No |
| 46 | MMP9 | matrix metallopeptidase 9 | 3774 | 0.040 | 0.1961 | No |
| 47 | DCBLD2 | discoidin, CUB and LCCL domain containing 2 | 3862 | 0.039 | 0.1950 | No |
| 48 | DNMBP | dynamin binding protein | 4000 | 0.038 | 0.1912 | No |
| 49 | CPE | carboxypeptidase E | 4003 | 0.038 | 0.1943 | No |
| 50 | NAP1L2 | Transcript Identified by AceView, Entrez Gene ID(s) 4674 | 4139 | 0.036 | 0.1905 | No |
| 51 | TSPAN1 | tetraspanin 1 | 4155 | 0.036 | 0.1929 | No |
| 52 | GYPC | glycophorin C (Gerbich blood group) | 4447 | 0.034 | 0.1808 | No |
| 53 | ADGRL4 | adhesion G protein-coupled receptor L4 | 4814 | 0.031 | 0.1645 | No |
| 54 | ADAM8 | Memczak2013 ANTISENSE, CDS, coding, INTERNAL best transcript NM\_001109 | 4934 | 0.030 | 0.1609 | No |
| 55 | TRAF1 | TNF receptor-associated factor 1 | 5030 | 0.029 | 0.1586 | No |
| 56 | SATB1 | SATB homeobox 1 | 5090 | 0.029 | 0.1580 | No |
| 57 | MPZL2 | myelin protein zero-like 2 | 5117 | 0.028 | 0.1591 | No |
| 58 | ANXA10 | annexin A10 | 5136 | 0.028 | 0.1607 | No |
| 59 | EPHB2 | EPH receptor B2 | 5161 | 0.028 | 0.1619 | No |
| 60 | F13A1 | coagulation factor XIII, A1 polypeptide | 5186 | 0.028 | 0.1631 | No |
| 61 | PLAU | plasminogen activator, urokinase | 5343 | 0.027 | 0.1573 | No |
| 62 | NGF | nerve growth factor (beta polypeptide) | 5632 | 0.025 | 0.1445 | No |
| 63 | EREG | epiregulin | 5902 | 0.023 | 0.1326 | No |
| 64 | ADGRA2 | adhesion G protein-coupled receptor A2 | 5996 | 0.022 | 0.1297 | No |
| 65 | LCP1 | lymphocyte cytosolic protein 1 (L-plastin) | 6241 | 0.021 | 0.1189 | No |
| 66 | MMD | monocyte to macrophage differentiation-associated | 6413 | 0.020 | 0.1117 | No |
| 67 | CD37 | CD37 molecule | 7012 | 0.016 | 0.0822 | No |
| 68 | GNG11 | guanine nucleotide binding protein (G protein), gamma 11 | 7017 | 0.016 | 0.0833 | No |
| 69 | HDAC9 | histone deacetylase 9 | 7032 | 0.016 | 0.0840 | No |
| 70 | MAP3K1 | mitogen-activated protein kinase kinase kinase 1, E3 ubiquitin protein ligase | 7078 | 0.015 | 0.0830 | No |
| 71 | ATG10 | autophagy related 10 | 7357 | 0.014 | 0.0698 | No |
| 72 | PLEK2 | pleckstrin 2 | 7366 | 0.014 | 0.0706 | No |
| 73 | RBM4 | RNA binding motif protein 4 | 7370 | 0.014 | 0.0716 | No |
| 74 | CFHR2 | complement factor H-related 2 | 7586 | 0.012 | 0.0616 | No |
| 75 | WDR33 | WD repeat domain 33 | 7738 | 0.011 | 0.0548 | No |
| 76 | CSF2 | colony stimulating factor 2 (granulocyte-macrophage) | 7825 | 0.011 | 0.0513 | No |
| 77 | CXCL10 | chemokine (C-X-C motif) ligand 10 | 8286 | 0.008 | 0.0282 | No |
| 78 | CXCR4 | chemokine (C-X-C motif) receptor 4 | 8789 | 0.005 | 0.0026 | No |
| 79 | NIN | ninein (GSK3B interacting protein) | 8837 | 0.005 | 0.0007 | No |
| 80 | APOD | apolipoprotein D | 9033 | 0.004 | -0.0091 | No |
| 81 | LY96 | lymphocyte antigen 96 | 9061 | 0.004 | -0.0102 | No |
| 82 | PCSK1N | proprotein convertase subtilisin/kexin type 1 inhibitor | 9104 | 0.003 | -0.0121 | No |
| 83 | SPRY2 | sprouty RTK signaling antagonist 2 | 9142 | 0.003 | -0.0137 | No |
| 84 | RELN | reelin | 9550 | 0.001 | -0.0346 | No |
| 85 | IGF2 | insulin-like growth factor 2 | 9551 | 0.001 | -0.0345 | No |
| 86 | F2RL1 | coagulation factor II (thrombin) receptor-like 1 | 9798 | -0.000 | -0.0472 | No |
| 87 | TNFRSF1B | tumor necrosis factor receptor superfamily, member 1B | 9919 | -0.001 | -0.0534 | No |
| 88 | PTBP2 | polypyrimidine tract binding protein 2 | 10020 | -0.001 | -0.0584 | No |
| 89 | SCG5 | secretogranin V | 10169 | -0.002 | -0.0659 | No |
| 90 | SOX9 | SRY box 9 | 10277 | -0.003 | -0.0712 | No |
| 91 | ITGB2 | Memczak2013 ANTISENSE, CDS, coding, INTERNAL best transcript NM\_001127491 | 10383 | -0.003 | -0.0764 | No |
| 92 | MAFB | v-maf avian musculoaponeurotic fibrosarcoma oncogene homolog B | 10398 | -0.003 | -0.0768 | No |
| 93 | ARG1 | arginase 1 | 10703 | -0.005 | -0.0920 | No |
| 94 | ALDH1A3 | aldehyde dehydrogenase 1 family, member A3 | 10720 | -0.005 | -0.0924 | No |
| 95 | PLAT | plasminogen activator, tissue | 10760 | -0.006 | -0.0939 | No |
| 96 | HOXD11 | homeobox D11 | 10947 | -0.007 | -0.1029 | No |
| 97 | RGS16 | regulator of G-protein signaling 16 | 11032 | -0.007 | -0.1067 | No |
| 98 | TLR8 | toll-like receptor 8 | 11049 | -0.007 | -0.1068 | No |
| 99 | MYCN | v-myc avian myelocytomatosis viral oncogene neuroblastoma derived homolog | 11075 | -0.007 | -0.1075 | No |
| 100 | SPARCL1 | SPARC like 1 | 11139 | -0.008 | -0.1100 | No |
| 101 | TMEM100 | transmembrane protein 100 | 11309 | -0.009 | -0.1180 | No |
| 102 | CBL | Cbl proto-oncogene, E3 ubiquitin protein ligase | 11535 | -0.010 | -0.1288 | No |
| 103 | SPP1 | secreted phosphoprotein 1 | 11575 | -0.010 | -0.1299 | No |
| 104 | IKZF1 | IKAROS family zinc finger 1 | 11602 | -0.010 | -0.1304 | No |
| 105 | FBXO4 | F-box protein 4 | 11607 | -0.010 | -0.1296 | No |
| 106 | IL1RL2 | interleukin 1 receptor-like 2 | 11708 | -0.011 | -0.1339 | No |
| 107 | NR0B2 | nuclear receptor subfamily 0, group B, member 2 | 11794 | -0.012 | -0.1372 | No |
| 108 | GFPT2 | glutamine-fructose-6-phosphate transaminase 2 | 11850 | -0.012 | -0.1391 | No |
| 109 | FGF9 | fibroblast growth factor 9 | 11971 | -0.013 | -0.1442 | No |
| 110 | ZNF639 | zinc finger protein 639 | 12528 | -0.016 | -0.1715 | No |
| 111 | CMKLR1 | chemerin chemokine-like receptor 1 | 12531 | -0.016 | -0.1702 | No |
| 112 | PSMB8 | proteasome subunit beta 8 | 12759 | -0.018 | -0.1804 | No |
| 113 | FLT4 | fms-related tyrosine kinase 4 | 12846 | -0.018 | -0.1833 | No |
| 114 | RETN | resistin | 12935 | -0.019 | -0.1862 | No |
| 115 | PRKG2 | protein kinase, cGMP-dependent, type II | 13033 | -0.019 | -0.1895 | No |
| 116 | FUCA1 | fucosidase, alpha-L- 1, tissue | 13114 | -0.020 | -0.1919 | No |
| 117 | PEG3 | paternally expressed 3 | 13165 | -0.020 | -0.1927 | No |
| 118 | SCN1B | sodium channel, voltage gated, type I beta subunit | 13217 | -0.021 | -0.1936 | No |
| 119 | JUP | junction plakoglobin | 13218 | -0.021 | -0.1918 | No |
| 120 | PDCD1LG2 | programmed cell death 1 ligand 2 | 13265 | -0.021 | -0.1923 | No |
| 121 | TNFAIP3 | tumor necrosis factor, alpha-induced protein 3 | 13363 | -0.022 | -0.1955 | No |
| 122 | AKAP12 | A kinase (PRKA) anchor protein 12 | 13539 | -0.023 | -0.2025 | No |
| 123 | TMEM176A | transmembrane protein 176A | 13759 | -0.025 | -0.2117 | No |
| 124 | CBX8 | chromobox homolog 8 | 13774 | -0.025 | -0.2103 | No |
| 125 | ENG | endoglin | 13817 | -0.025 | -0.2103 | No |
| 126 | MMP10 | matrix metallopeptidase 10 | 13819 | -0.025 | -0.2082 | No |
| 127 | ETS1 | v-ets avian erythroblastosis virus E26 oncogene homolog 1 | 13826 | -0.025 | -0.2063 | No |
| 128 | ERO1A | endoplasmic reticulum oxidoreductase alpha | 13862 | -0.025 | -0.2059 | No |
| 129 | ZNF277 | zinc finger protein 277 | 13909 | -0.026 | -0.2060 | No |
| 130 | AKT2 | v-akt murine thymoma viral oncogene homolog 2 | 13964 | -0.026 | -0.2066 | No |
| 131 | STRN | striatin, calmodulin binding protein | 14062 | -0.027 | -0.2093 | No |
| 132 | PRELID3B | PRELI domain containing 3B | 14290 | -0.029 | -0.2185 | No |
| 133 | DOCK2 | dedicator of cytokinesis 2 | 14338 | -0.029 | -0.2184 | No |
| 134 | SERPINA3 | serpin peptidase inhibitor, clade A (alpha-1 antiproteinase, antitrypsin), member 3 | 14750 | -0.032 | -0.2368 | No |
| 135 | HBEGF | heparin-binding EGF-like growth factor | 14753 | -0.032 | -0.2341 | No |
| 136 | CIDEA | cell death-inducing DFFA-like effector a | 14801 | -0.033 | -0.2337 | No |
| 137 | HSD11B1 | hydroxysteroid (11-beta) dehydrogenase 1 | 14833 | -0.033 | -0.2324 | No |
| 138 | TPH1 | tryptophan hydroxylase 1 | 14976 | -0.034 | -0.2368 | No |
| 139 | INHBA | inhibin beta A | 15087 | -0.035 | -0.2394 | No |
| 140 | SDCCAG8 | serologically defined colon cancer antigen 8 | 15397 | -0.038 | -0.2521 | No |
| 141 | ETV4 | ets variant 4 | 15403 | -0.038 | -0.2491 | No |
| 142 | ITGA2 | integrin, alpha 2 (CD49B, alpha 2 subunit of VLA-2 receptor) | 15498 | -0.039 | -0.2506 | No |
| 143 | BPGM | 2,3-bisphosphoglycerate mutase | 15645 | -0.040 | -0.2546 | No |
| 144 | TMEM158 | transmembrane protein 158 (gene/pseudogene) | 15662 | -0.040 | -0.2519 | No |
| 145 | TRIB1 | tribbles pseudokinase 1 | 15669 | -0.041 | -0.2487 | No |
| 146 | CCSER2 | coiled-coil serine rich protein 2 | 15896 | -0.043 | -0.2567 | No |
| 147 | USP12 | ubiquitin specific peptidase 12 | 16063 | -0.044 | -0.2614 | No |
| 148 | SNAP25 | synaptosome associated protein 25kDa | 16069 | -0.045 | -0.2578 | No |
| 149 | PECAM1 | platelet/endothelial cell adhesion molecule 1 | 16255 | -0.047 | -0.2633 | Yes |
| 150 | ANKH | ANKH inorganic pyrophosphate transport regulator | 16325 | -0.047 | -0.2627 | Yes |
| 151 | ADAMDEC1 | ADAM-like, decysin 1 | 16345 | -0.048 | -0.2595 | Yes |
| 152 | BMP2 | bone morphogenetic protein 2 | 16350 | -0.048 | -0.2556 | Yes |
| 153 | GLRX | glutaredoxin | 16461 | -0.049 | -0.2570 | Yes |
| 154 | AMMECR1 | Alport syndrome, mental retardation, midface hypoplasia and elliptocytosis chromosomal region gene 1 | 16567 | -0.051 | -0.2580 | Yes |
| 155 | YRDC | yrdC N(6)-threonylcarbamoyltransferase domain containing | 16643 | -0.052 | -0.2574 | Yes |
| 156 | CTSS | cathepsin S | 16657 | -0.052 | -0.2535 | Yes |
| 157 | ADAM17 | ADAM metallopeptidase domain 17 | 16706 | -0.053 | -0.2514 | Yes |
| 158 | GADD45G | growth arrest and DNA-damage-inducible, gamma | 17011 | -0.057 | -0.2622 | Yes |
| 159 | ALDH1A2 | aldehyde dehydrogenase 1 family, member A2 | 17022 | -0.057 | -0.2577 | Yes |
| 160 | C3AR1 | complement component 3a receptor 1 | 17045 | -0.058 | -0.2538 | Yes |
| 161 | KIF5C | kinesin family member 5C | 17209 | -0.060 | -0.2570 | Yes |
| 162 | TOR1AIP2 | torsin A interacting protein 2 | 17294 | -0.062 | -0.2560 | Yes |
| 163 | TSPAN13 | tetraspanin 13 | 17370 | -0.063 | -0.2544 | Yes |
| 164 | ETV1 | ets variant 1 | 17589 | -0.067 | -0.2598 | Yes |
| 165 | SEMA3B | sema domain, immunoglobulin domain (Ig), short basic domain, secreted, (semaphorin) 3B | 17591 | -0.067 | -0.2541 | Yes |
| 166 | RABGAP1L | RAB GTPase activating protein 1-like | 17616 | -0.068 | -0.2494 | Yes |
| 167 | GPNMB | glycoprotein (transmembrane) nmb | 17617 | -0.068 | -0.2435 | Yes |
| 168 | MTMR10 | myotubularin related protein 10 | 17774 | -0.071 | -0.2454 | Yes |
| 169 | MAP4K1 | mitogen-activated protein kinase kinase kinase kinase 1 | 17834 | -0.073 | -0.2421 | Yes |
| 170 | EVI5 | ecotropic viral integration site 5 | 17858 | -0.073 | -0.2369 | Yes |
| 171 | USH1C | Usher syndrome 1C | 17897 | -0.074 | -0.2324 | Yes |
| 172 | PTPRR | protein tyrosine phosphatase, receptor type, R | 17901 | -0.074 | -0.2260 | Yes |
| 173 | AVL9 | AVL9 homolog (S. cerevisiase) | 17944 | -0.076 | -0.2216 | Yes |
| 174 | KLF4 | Kruppel-like factor 4 (gut) | 17974 | -0.076 | -0.2165 | Yes |
| 175 | ABCB1 | ATP binding cassette subfamily B member 1 | 18030 | -0.078 | -0.2126 | Yes |
| 176 | CDADC1 | cytidine and dCMP deaminase domain containing 1 | 18041 | -0.078 | -0.2063 | Yes |
| 177 | CSF2RA | colony stimulating factor 2 receptor, alpha, low-affinity (granulocyte-macrophage) | 18316 | -0.086 | -0.2129 | Yes |
| 178 | TRIB2 | tribbles pseudokinase 2 | 18347 | -0.087 | -0.2069 | Yes |
| 179 | GALNT3 | polypeptide N-acetylgalactosaminyltransferase 3 | 18446 | -0.091 | -0.2041 | Yes |
| 180 | LAT2 | linker for activation of T-cells family member 2 | 18505 | -0.093 | -0.1990 | Yes |
| 181 | MMP11 | matrix metallopeptidase 11 | 18635 | -0.098 | -0.1971 | Yes |
| 182 | PIGR | polymeric immunoglobulin receptor | 18668 | -0.101 | -0.1900 | Yes |
| 183 | TSPAN7 | tetraspanin 7 | 18694 | -0.102 | -0.1824 | Yes |
| 184 | G0S2 | G0/G1 switch 2 | 18709 | -0.103 | -0.1741 | Yes |
| 185 | TMEM176B | transmembrane protein 176B | 18831 | -0.112 | -0.1706 | Yes |
| 186 | PPP1R15A | protein phosphatase 1, regulatory subunit 15A | 18867 | -0.115 | -0.1624 | Yes |
| 187 | HKDC1 | hexokinase domain containing 1 | 18889 | -0.117 | -0.1533 | Yes |
| 188 | PRRX1 | paired related homeobox 1 | 18904 | -0.118 | -0.1437 | Yes |
| 189 | IL1B | interleukin 1 beta | 19023 | -0.130 | -0.1386 | Yes |
| 190 | CAB39L | calcium binding protein 39-like | 19029 | -0.131 | -0.1274 | Yes |
| 191 | WNT7A | wingless-type MMTV integration site family, member 7A | 19034 | -0.132 | -0.1162 | Yes |
| 192 | GABRA3 | gamma-aminobutyric acid (GABA) A receptor, alpha 3 | 19068 | -0.137 | -0.1060 | Yes |
| 193 | PCP4 | Purkinje cell protein 4 | 19121 | -0.145 | -0.0960 | Yes |
| 194 | ETV5 | ets variant 5 | 19192 | -0.159 | -0.0857 | Yes |
| 195 | PTGS2 | prostaglandin-endoperoxide synthase 2 (prostaglandin G/H synthase and cyclooxygenase) | 19364 | -0.218 | -0.0756 | Yes |
| 196 | CROT | carnitine O-octanoyltransferase | 19375 | -0.223 | -0.0567 | Yes |
| 197 | EPB41L3 | erythrocyte membrane protein band 4.1-like 3 | 19478 | -0.335 | -0.0328 | Yes |
| 198 | TFPI | tissue factor pathway inhibitor (lipoprotein-associated coagulation inhibitor) | 19499 | -0.403 | 0.0013 | Yes |
Table: GSEA details [plain text format]

  

Fig 2: HALLMARK\_KRAS\_SIGNALING\_UP      
 Blue-Pink O' Gram in the Space of the Analyzed GeneSet

  

Fig 3: HALLMARK\_KRAS\_SIGNALING\_UP: Random ES distribution      
 Gene set null distribution of ES for **HALLMARK\_KRAS\_SIGNALING\_UP**

  
