## Supplementary material for "Somatic hypomethylation of pericentromeric SST1 repeats and tetraploidization in human colorectal cancer cells": GSEA results: HALLMARK_MTORC1_SIGNALING.html

Details for gene set HALLMARK\_MTORC1\_SIGNALING[GSEA]

|  || Dataset | eset\_byprobe\_collapsed\_to\_symbols.Diploid\_vs\_Tetraploid.cls #Tetraploid\_versus\_Diploid.Diploid\_vs\_Tetraploid.cls #Tetraploid\_versus\_Diploid\_repos |
| Phenotype | Diploid\_vs\_Tetraploid.cls#Tetraploid\_versus\_Diploid\_repos |
| Upregulated in class | Diploid |
| GeneSet | HALLMARK\_MTORC1\_SIGNALING |
| Enrichment Score (ES) | -0.2779568 |
| Normalized Enrichment Score (NES) | -1.1446766 |
| Nominal p-value | 0.2027439 |
| FDR q-value | 0.33682904 |
| FWER p-Value | 0.998 |
Table: GSEA Results Summary

  

Fig 1: Enrichment plot: HALLMARK\_MTORC1\_SIGNALING      
 Profile of the Running ES Score & Positions of GeneSet Members on the Rank Ordered List

  

| SYMBOL | TITLE | RANK IN GENE LIST | RANK METRIC SCORE | RUNNING ES | CORE ENRICHMENT || 1 | SQLE | squalene epoxidase | 167 | 0.166 | 0.0090 | No |
| 2 | BTG2 | BTG family, member 2 | 281 | 0.136 | 0.0175 | No |
| 3 | CDC25A | cell division cycle 25A | 295 | 0.134 | 0.0311 | No |
| 4 | FADS1 | fatty acid desaturase 1 | 296 | 0.134 | 0.0453 | No |
| 5 | HPRT1 | hypoxanthine phosphoribosyltransferase 1 | 311 | 0.132 | 0.0586 | No |
| 6 | FADS2 | fatty acid desaturase 2 | 315 | 0.131 | 0.0723 | No |
| 7 | HMGCS1 | 3-hydroxy-3-methylglutaryl-CoA synthase 1 (soluble) | 387 | 0.121 | 0.0814 | No |
| 8 | PSMG1 | proteasome (prosome, macropain) assembly chaperone 1 | 497 | 0.111 | 0.0875 | No |
| 9 | CDKN1A | cyclin-dependent kinase inhibitor 1A (p21, Cip1) | 690 | 0.097 | 0.0879 | No |
| 10 | CORO1A | coronin, actin binding protein, 1A | 1041 | 0.083 | 0.0786 | No |
| 11 | CACYBP | calcyclin binding protein | 1071 | 0.082 | 0.0857 | No |
| 12 | SYTL2 | synaptotagmin-like 2 | 1087 | 0.081 | 0.0936 | No |
| 13 | EBP | emopamil binding protein (sterol isomerase) | 1093 | 0.081 | 0.1019 | No |
| 14 | FDXR | ferredoxin reductase | 1184 | 0.079 | 0.1056 | No |
| 15 | TUBA4A | tubulin, alpha 4a | 1323 | 0.076 | 0.1065 | No |
| 16 | MCM4 | minichromosome maintenance complex component 4 | 1325 | 0.076 | 0.1145 | No |
| 17 | TMEM97 | transmembrane protein 97 | 1371 | 0.075 | 0.1200 | No |
| 18 | RRP9 | ribosomal RNA processing 9, small subunit (SSU) processome component, homolog (yeast) | 1374 | 0.075 | 0.1278 | No |
| 19 | TM7SF2 | transmembrane 7 superfamily member 2 | 1402 | 0.074 | 0.1343 | No |
| 20 | PNP | purine nucleoside phosphorylase | 1405 | 0.074 | 0.1420 | No |
| 21 | HSPE1 | heat shock 10kDa protein 1 | 1413 | 0.074 | 0.1494 | No |
| 22 | PPA1 | pyrophosphatase (inorganic) 1 | 1788 | 0.065 | 0.1370 | No |
| 23 | HMGCR | 3-hydroxy-3-methylglutaryl-CoA reductase | 1821 | 0.064 | 0.1422 | No |
| 24 | DHCR7 | 7-dehydrocholesterol reductase | 1877 | 0.064 | 0.1460 | No |
| 25 | IDI1 | isopentenyl-diphosphate delta isomerase 1 | 1905 | 0.063 | 0.1513 | No |
| 26 | POLR3G | polymerase (RNA) III (DNA directed) polypeptide G (32kD) | 1915 | 0.063 | 0.1575 | No |
| 27 | PSMC6 | proteasome 26S subunit, ATPase 6 | 1998 | 0.061 | 0.1598 | No |
| 28 | CYB5B | cytochrome b5 type B (outer mitochondrial membrane) | 2082 | 0.060 | 0.1619 | No |
| 29 | GLA | galactosidase, alpha | 2101 | 0.060 | 0.1673 | No |
| 30 | PRDX1 | peroxiredoxin 1 | 2171 | 0.059 | 0.1700 | No |
| 31 | CTH | cystathionine gamma-lyase | 2192 | 0.058 | 0.1751 | No |
| 32 | UNG | uracil DNA glycosylase | 2227 | 0.058 | 0.1795 | No |
| 33 | CCNG1 | cyclin G1 | 2329 | 0.056 | 0.1802 | No |
| 34 | SORD | sorbitol dehydrogenase | 2500 | 0.054 | 0.1771 | No |
| 35 | DHCR24 | 24-dehydrocholesterol reductase | 2597 | 0.052 | 0.1777 | No |
| 36 | RRM2 | ribonucleotide reductase M2 | 3128 | 0.046 | 0.1552 | No |
| 37 | ELOVL6 | ELOVL fatty acid elongase 6 | 3132 | 0.046 | 0.1599 | No |
| 38 | GMPS | guanine monophosphate synthase | 3161 | 0.046 | 0.1633 | No |
| 39 | STC1 | stanniocalcin 1 | 3174 | 0.046 | 0.1676 | No |
| 40 | SCD | stearoyl-CoA desaturase (delta-9-desaturase) | 3227 | 0.045 | 0.1697 | No |
| 41 | SLC9A3R1 | solute carrier family 9, subfamily A (NHE3, cation proton antiporter 3), member 3 regulator 1 | 3344 | 0.044 | 0.1684 | No |
| 42 | INSIG1 | insulin induced gene 1 | 3412 | 0.044 | 0.1695 | No |
| 43 | CFP | complement factor properdin | 3594 | 0.042 | 0.1646 | No |
| 44 | SLC37A4 | solute carrier family 37 (glucose-6-phosphate transporter), member 4 | 3916 | 0.038 | 0.1520 | No |
| 45 | PSMA4 | proteasome subunit alpha 4 | 3948 | 0.038 | 0.1545 | No |
| 46 | TOMM40 | translocase of outer mitochondrial membrane 40 homolog (yeast) | 3997 | 0.038 | 0.1559 | No |
| 47 | STIP1 | stress-induced phosphoprotein 1 | 4082 | 0.037 | 0.1555 | No |
| 48 | TPI1 | triosephosphate isomerase 1 | 4710 | 0.031 | 0.1264 | No |
| 49 | HSPA4 | heat shock 70kDa protein 4 | 4795 | 0.031 | 0.1253 | No |
| 50 | RPA1 | replication protein A1 | 4848 | 0.030 | 0.1258 | No |
| 51 | DHFR | dihydrofolate reductase | 4862 | 0.030 | 0.1284 | No |
| 52 | SKAP2 | src kinase associated phosphoprotein 2 | 4881 | 0.030 | 0.1307 | No |
| 53 | AURKA | aurora kinase A | 5067 | 0.029 | 0.1241 | No |
| 54 | EEF1E1 | eukaryotic translation elongation factor 1 epsilon 1 | 5137 | 0.028 | 0.1236 | No |
| 55 | EGLN3 | egl-9 family hypoxia-inducible factor 3 | 5195 | 0.028 | 0.1236 | No |
| 56 | DDX39A | DEAD (Asp-Glu-Ala-Asp) box polypeptide 39A | 5309 | 0.027 | 0.1206 | No |
| 57 | MCM2 | minichromosome maintenance complex component 2 | 5326 | 0.027 | 0.1226 | No |
| 58 | VLDLR | very low density lipoprotein receptor | 5549 | 0.025 | 0.1138 | No |
| 59 | TUBG1 | tubulin, gamma 1 | 5617 | 0.025 | 0.1129 | No |
| 60 | ACLY | ATP citrate lyase | 5633 | 0.025 | 0.1147 | No |
| 61 | PSMD13 | proteasome 26S subunit, non-ATPase 13 | 5655 | 0.024 | 0.1162 | No |
| 62 | GPI | glucose-6-phosphate isomerase | 5659 | 0.024 | 0.1186 | No |
| 63 | ELOVL5 | ELOVL fatty acid elongase 5 | 5798 | 0.023 | 0.1140 | No |
| 64 | BUB1 | BUB1 mitotic checkpoint serine/threonine kinase | 5988 | 0.022 | 0.1065 | No |
| 65 | LDHA | lactate dehydrogenase A | 6019 | 0.022 | 0.1073 | No |
| 66 | HSPD1 | heat shock 60kDa protein 1 (chaperonin) | 6186 | 0.021 | 0.1009 | No |
| 67 | CYP51A1 | cytochrome P450, family 51, subfamily A, polypeptide 1 | 6319 | 0.020 | 0.0962 | No |
| 68 | ALDOA | aldolase A, fructose-bisphosphate | 6361 | 0.020 | 0.0962 | No |
| 69 | SEC11A | SEC11 homolog A, signal peptidase complex subunit | 6770 | 0.017 | 0.0770 | No |
| 70 | PITPNB | Memczak2013 ANTISENSE, coding, INTERNAL, intronic best transcript NM\_012399 | 6845 | 0.017 | 0.0749 | No |
| 71 | PSMB5 | proteasome subunit beta 5 | 6886 | 0.017 | 0.0746 | No |
| 72 | CD9 | CD9 molecule | 6905 | 0.016 | 0.0754 | No |
| 73 | HMBS | hydroxymethylbilane synthase | 6994 | 0.016 | 0.0725 | No |
| 74 | LGMN | legumain | 7134 | 0.015 | 0.0669 | No |
| 75 | TBK1 | TANK-binding kinase 1 | 7221 | 0.015 | 0.0640 | No |
| 76 | LTA4H | leukotriene A4 hydrolase | 7229 | 0.015 | 0.0652 | No |
| 77 | CCNF | cyclin F | 7608 | 0.012 | 0.0470 | No |
| 78 | ARPC5L | actin related protein 2/3 complex subunit 5-like | 8051 | 0.010 | 0.0251 | No |
| 79 | PSMA3 | proteasome subunit alpha 3 | 8134 | 0.009 | 0.0218 | No |
| 80 | CTSC | cathepsin C | 8167 | 0.009 | 0.0211 | No |
| 81 | PSMD14 | proteasome 26S subunit, non-ATPase 14 | 8280 | 0.008 | 0.0162 | No |
| 82 | ACACA | acetyl-CoA carboxylase alpha | 8311 | 0.008 | 0.0155 | No |
| 83 | PIK3R3 | phosphoinositide-3-kinase, regulatory subunit 3 (gamma) | 8325 | 0.008 | 0.0156 | No |
| 84 | STARD4 | StAR-related lipid transfer domain containing 4 | 8485 | 0.007 | 0.0082 | No |
| 85 | HK2 | hexokinase 2 | 8575 | 0.006 | 0.0042 | No |
| 86 | PPIA | peptidylprolyl isomerase A (cyclophilin A) | 8627 | 0.006 | 0.0023 | No |
| 87 | NFYC | nuclear transcription factor Y subunit gamma | 8632 | 0.006 | 0.0027 | No |
| 88 | PFKL | phosphofructokinase, liver | 8698 | 0.006 | -0.0001 | No |
| 89 | PLK1 | polo-like kinase 1 | 8734 | 0.006 | -0.0013 | No |
| 90 | ADIPOR2 | adiponectin receptor 2 | 8746 | 0.005 | -0.0013 | No |
| 91 | CXCR4 | chemokine (C-X-C motif) receptor 4 | 8789 | 0.005 | -0.0029 | No |
| 92 | TFRC | transferrin receptor | 8948 | 0.004 | -0.0106 | No |
| 93 | HSPA9 | heat shock 70kDa protein 9 (mortalin) | 9059 | 0.004 | -0.0159 | No |
| 94 | SLC6A6 | solute carrier family 6 (neurotransmitter transporter), member 6 | 9077 | 0.004 | -0.0164 | No |
| 95 | UCHL5 | ubiquitin C-terminal hydrolase L5 | 9161 | 0.003 | -0.0204 | No |
| 96 | PSME3 | proteasome activator subunit 3 | 9271 | 0.003 | -0.0257 | No |
| 97 | GGA2 | golgi-associated, gamma adaptin ear containing, ARF binding protein 2 | 9285 | 0.003 | -0.0261 | No |
| 98 | LDLR | low density lipoprotein receptor | 9447 | 0.002 | -0.0343 | No |
| 99 | CCT6A | chaperonin containing TCP1, subunit 6A (zeta 1) | 9522 | 0.001 | -0.0380 | No |
| 100 | GAPDH | glyceraldehyde-3-phosphate dehydrogenase | 9830 | -0.000 | -0.0538 | No |
| 101 | GSR | glutathione reductase | 9874 | -0.000 | -0.0560 | No |
| 102 | SC5D | sterol-C5-desaturase | 9987 | -0.001 | -0.0617 | No |
| 103 | SLA | Src-like-adaptor | 10023 | -0.001 | -0.0633 | No |
| 104 | NMT1 | N-myristoyltransferase 1 | 10101 | -0.002 | -0.0671 | No |
| 105 | M6PR | mannose-6-phosphate receptor (cation dependent) | 10105 | -0.002 | -0.0671 | No |
| 106 | IMMT | inner membrane protein, mitochondrial | 10313 | -0.003 | -0.0775 | No |
| 107 | ITGB2 | Memczak2013 ANTISENSE, CDS, coding, INTERNAL best transcript NM\_001127491 | 10383 | -0.003 | -0.0807 | No |
| 108 | ACTR2 | ARP2 actin-related protein 2 homolog (yeast) | 10400 | -0.003 | -0.0812 | No |
| 109 | MAP2K3 | mitogen-activated protein kinase kinase 3 | 10451 | -0.004 | -0.0834 | No |
| 110 | PSMC2 | proteasome 26S subunit, ATPase 2 | 10455 | -0.004 | -0.0831 | No |
| 111 | IGFBP5 | insulin like growth factor binding protein 5 | 10518 | -0.004 | -0.0859 | No |
| 112 | GOT1 | glutamic-oxaloacetic transaminase 1, soluble | 11039 | -0.007 | -0.1120 | No |
| 113 | ACSL3 | acyl-CoA synthetase long-chain family member 3 | 11087 | -0.008 | -0.1136 | No |
| 114 | PGK1 | phosphoglycerate kinase 1 | 11128 | -0.008 | -0.1149 | No |
| 115 | SLC2A1 | solute carrier family 2 (facilitated glucose transporter), member 1 | 11178 | -0.008 | -0.1165 | No |
| 116 | RDH11 | retinol dehydrogenase 11 (all-trans/9-cis/11-cis) | 11501 | -0.010 | -0.1321 | No |
| 117 | ENO1 | Memczak2013 ANTISENSE, coding, INTERNAL, UTR3 best transcript NM\_001428 | 11613 | -0.010 | -0.1368 | No |
| 118 | ATP2A2 | ATPase, Ca++ transporting, cardiac muscle, slow twitch 2 | 11662 | -0.011 | -0.1381 | No |
| 119 | ABCF2 | ATP binding cassette subfamily F member 2 | 11747 | -0.011 | -0.1413 | No |
| 120 | PSMD12 | proteasome 26S subunit, non-ATPase 12 | 11772 | -0.011 | -0.1413 | No |
| 121 | GTF2H1 | general transcription factor IIH subunit 1 | 11860 | -0.012 | -0.1445 | No |
| 122 | PHGDH | phosphoglycerate dehydrogenase | 12063 | -0.013 | -0.1536 | No |
| 123 | ATP6V1D | ATPase, H+ transporting, lysosomal 34kDa, V1 subunit D | 12283 | -0.015 | -0.1633 | No |
| 124 | ETF1 | eukaryotic translation termination factor 1 | 12626 | -0.017 | -0.1792 | No |
| 125 | PSMC4 | proteasome 26S subunit, ATPase 4 | 12803 | -0.018 | -0.1865 | No |
| 126 | FKBP2 | FK506 binding protein 2 | 12981 | -0.019 | -0.1936 | No |
| 127 | SQSTM1 | sequestosome 1 | 13032 | -0.019 | -0.1941 | No |
| 128 | G6PD | glucose-6-phosphate dehydrogenase | 13330 | -0.021 | -0.2072 | No |
| 129 | NUFIP1 | nuclear fragile X mental retardation protein interacting protein 1 | 13394 | -0.022 | -0.2082 | No |
| 130 | RAB1A | RAB1A, member RAS oncogene family | 13574 | -0.023 | -0.2150 | No |
| 131 | P4HA1 | prolyl 4-hydroxylase, alpha polypeptide I | 13604 | -0.023 | -0.2140 | No |
| 132 | CANX | calnexin | 13700 | -0.024 | -0.2163 | No |
| 133 | ERO1A | endoplasmic reticulum oxidoreductase alpha | 13862 | -0.025 | -0.2220 | No |
| 134 | UBE2D3 | Memczak2013 ANTISENSE, coding, INTERNAL, intronic best transcript NM\_181893 | 13866 | -0.025 | -0.2194 | No |
| 135 | PNO1 | partner of NOB1 homolog | 13879 | -0.025 | -0.2174 | No |
| 136 | EIF2S2 | eukaryotic translation initiation factor 2, subunit 2 beta, 38kDa | 14130 | -0.027 | -0.2274 | No |
| 137 | MLLT11 | myeloid/lymphoid or mixed-lineage leukemia; translocated to, 11 | 14372 | -0.029 | -0.2368 | No |
| 138 | USO1 | USO1 vesicle transport factor | 14373 | -0.029 | -0.2336 | No |
| 139 | CALR | calreticulin | 14607 | -0.031 | -0.2424 | No |
| 140 | GSK3B | glycogen synthase kinase 3 beta | 14754 | -0.032 | -0.2465 | No |
| 141 | SERPINH1 | serpin peptidase inhibitor, clade H (heat shock protein 47), member 1, (collagen binding protein 1) | 14845 | -0.033 | -0.2477 | No |
| 142 | IDH1 | isocitrate dehydrogenase 1 (NADP+) | 14950 | -0.034 | -0.2494 | No |
| 143 | ACTR3 | ARP3 actin-related protein 3 homolog (yeast) | 14975 | -0.034 | -0.2470 | No |
| 144 | PSAT1 | phosphoserine aminotransferase 1 | 14978 | -0.034 | -0.2435 | No |
| 145 | SHMT2 | serine hydroxymethyltransferase 2 (mitochondrial) | 14979 | -0.034 | -0.2399 | No |
| 146 | QDPR | quinoid dihydropteridine reductase | 15173 | -0.036 | -0.2461 | No |
| 147 | HSP90B1 | heat shock protein 90kDa beta (Grp94), member 1 | 15295 | -0.037 | -0.2484 | No |
| 148 | NFKBIB | nuclear factor of kappa light polypeptide gene enhancer in B-cells inhibitor, beta | 15315 | -0.037 | -0.2454 | No |
| 149 | XBP1 | X-box binding protein 1 | 15622 | -0.040 | -0.2570 | No |
| 150 | COPS5 | COP9 signalosome subunit 5 | 15670 | -0.041 | -0.2552 | No |
| 151 | SLC2A3 | solute carrier family 2 (facilitated glucose transporter), member 3 | 16073 | -0.045 | -0.2713 | No |
| 152 | ADD3 | adducin 3 (gamma) | 16088 | -0.045 | -0.2672 | No |
| 153 | RIT1 | Ras-like without CAAX 1 | 16124 | -0.045 | -0.2643 | No |
| 154 | SDF2L1 | stromal cell-derived factor 2-like 1 | 16144 | -0.045 | -0.2604 | No |
| 155 | SSR1 | signal sequence receptor, alpha | 16290 | -0.047 | -0.2629 | No |
| 156 | GLRX | glutaredoxin | 16461 | -0.049 | -0.2665 | No |
| 157 | NAMPT | nicotinamide phosphoribosyltransferase | 16683 | -0.052 | -0.2724 | Yes |
| 158 | MTHFD2L | methylenetetrahydrofolate dehydrogenase (NADP+ dependent) 2-like | 16765 | -0.053 | -0.2709 | Yes |
| 159 | TCEA1 | transcription elongation factor A (SII), 1 | 16811 | -0.054 | -0.2675 | Yes |
| 160 | AK4 | adenylate kinase 4 | 16854 | -0.055 | -0.2639 | Yes |
| 161 | PDAP1 | PDGFA associated protein 1 | 17018 | -0.057 | -0.2663 | Yes |
| 162 | RPN1 | ribophorin I | 17081 | -0.058 | -0.2634 | Yes |
| 163 | BCAT1 | branched chain amino-acid transaminase 1, cytosolic | 17138 | -0.059 | -0.2600 | Yes |
| 164 | TXNRD1 | thioredoxin reductase 1 | 17390 | -0.063 | -0.2663 | Yes |
| 165 | DDIT4 | DNA damage inducible transcript 4 | 17557 | -0.066 | -0.2678 | Yes |
| 166 | PLOD2 | procollagen-lysine, 2-oxoglutarate 5-dioxygenase 2 | 17575 | -0.067 | -0.2617 | Yes |
| 167 | SLC7A5 | solute carrier family 7 (amino acid transporter light chain, L system), member 5 | 17849 | -0.073 | -0.2680 | Yes |
| 168 | SRD5A1 | steroid-5-alpha-reductase, alpha polypeptide 1 (3-oxo-5 alpha-steroid delta 4-dehydrogenase alpha 1) | 17911 | -0.075 | -0.2633 | Yes |
| 169 | NUPR1 | nuclear protein 1, transcriptional regulator | 17958 | -0.076 | -0.2576 | Yes |
| 170 | GCLC | glutamate-cysteine ligase, catalytic subunit | 17981 | -0.077 | -0.2506 | Yes |
| 171 | ME1 | malic enzyme 1, NADP(+)-dependent, cytosolic | 18066 | -0.079 | -0.2466 | Yes |
| 172 | EDEM1 | ER degradation enhancer, mannosidase alpha-like 1 | 18132 | -0.081 | -0.2414 | Yes |
| 173 | YKT6 | YKT6 v-SNARE homolog (S. cerevisiae) | 18191 | -0.082 | -0.2357 | Yes |
| 174 | PDK1 | pyruvate dehydrogenase kinase, isozyme 1 | 18204 | -0.083 | -0.2276 | Yes |
| 175 | MTHFD2 | methylenetetrahydrofolate dehydrogenase (NADP+ dependent) 2, methenyltetrahydrofolate cyclohydrolase | 18255 | -0.084 | -0.2212 | Yes |
| 176 | NUP205 | nucleoporin 205kDa | 18334 | -0.086 | -0.2161 | Yes |
| 177 | SLC1A5 | solute carrier family 1 (neutral amino acid transporter), member 5 | 18428 | -0.090 | -0.2114 | Yes |
| 178 | HSPA5 | heat shock 70kDa protein 5 (glucose-regulated protein, 78kDa) | 18430 | -0.090 | -0.2019 | Yes |
| 179 | PSPH | phosphoserine phosphatase | 18469 | -0.091 | -0.1942 | Yes |
| 180 | NFIL3 | nuclear factor, interleukin 3 regulated | 18475 | -0.092 | -0.1848 | Yes |
| 181 | FGL2 | fibrinogen-like 2 | 18506 | -0.093 | -0.1765 | Yes |
| 182 | SLC7A11 | solute carrier family 7 (anionic amino acid transporter light chain, xc- system), member 11 | 18650 | -0.099 | -0.1733 | Yes |
| 183 | SLC1A4 | solute carrier family 1 (glutamate/neutral amino acid transporter), member 4 | 18809 | -0.110 | -0.1698 | Yes |
| 184 | ASNS | asparagine synthetase (glutamine-hydrolyzing) | 18864 | -0.115 | -0.1604 | Yes |
| 185 | PPP1R15A | protein phosphatase 1, regulatory subunit 15A | 18867 | -0.115 | -0.1483 | Yes |
| 186 | TRIB3 | tribbles pseudokinase 3 | 18870 | -0.115 | -0.1362 | Yes |
| 187 | TES | testin LIM domain protein | 18892 | -0.117 | -0.1249 | Yes |
| 188 | UFM1 | ubiquitin-fold modifier 1 | 18980 | -0.125 | -0.1161 | Yes |
| 189 | SERP1 | stress-associated endoplasmic reticulum protein 1 | 19031 | -0.131 | -0.1048 | Yes |
| 190 | GBE1 | glucan (1,4-alpha-), branching enzyme 1 | 19141 | -0.148 | -0.0948 | Yes |
| 191 | IFRD1 | interferon-related developmental regulator 1 | 19158 | -0.151 | -0.0796 | Yes |
| 192 | BHLHE40 | basic helix-loop-helix family, member e40 | 19252 | -0.175 | -0.0659 | Yes |
| 193 | DAPP1 | dual adaptor of phosphotyrosine and 3-phosphoinositides | 19293 | -0.191 | -0.0477 | Yes |
| 194 | PGM1 | phosphoglucomutase 1 | 19438 | -0.280 | -0.0255 | Yes |
| 195 | DDIT3 | DNA-damage-inducible transcript 3 | 19440 | -0.282 | 0.0043 | Yes |
Table: GSEA details [plain text format]

  

Fig 2: HALLMARK\_MTORC1\_SIGNALING      
 Blue-Pink O' Gram in the Space of the Analyzed GeneSet

  

Fig 3: HALLMARK\_MTORC1\_SIGNALING: Random ES distribution      
 Gene set null distribution of ES for **HALLMARK\_MTORC1\_SIGNALING**

  
