## Supplementary material for "Somatic hypomethylation of pericentromeric SST1 repeats and tetraploidization in human colorectal cancer cells": GSEA results: HALLMARK_MYC_TARGETS_V1.html

Details for gene set HALLMARK\_MYC\_TARGETS\_V1[GSEA]

|  || Dataset | eset\_byprobe\_collapsed\_to\_symbols.Diploid\_vs\_Tetraploid.cls #Tetraploid\_versus\_Diploid.Diploid\_vs\_Tetraploid.cls #Tetraploid\_versus\_Diploid\_repos |
| Phenotype | Diploid\_vs\_Tetraploid.cls#Tetraploid\_versus\_Diploid\_repos |
| Upregulated in class | Tetraploid |
| GeneSet | HALLMARK\_MYC\_TARGETS\_V1 |
| Enrichment Score (ES) | 0.40434572 |
| Normalized Enrichment Score (NES) | 1.8043815 |
| Nominal p-value | 0.0 |
| FDR q-value | 0.0021375 |
| FWER p-Value | 0.004 |
Table: GSEA Results Summary

  

Fig 1: Enrichment plot: HALLMARK\_MYC\_TARGETS\_V1      
 Profile of the Running ES Score & Positions of GeneSet Members on the Rank Ordered List

  

| SYMBOL | TITLE | RANK IN GENE LIST | RANK METRIC SCORE | RUNNING ES | CORE ENRICHMENT || 1 | HPRT1 | hypoxanthine phosphoribosyltransferase 1 | 311 | 0.132 | 0.0064 | Yes |
| 2 | CDC45 | cell division cycle 45 | 603 | 0.103 | 0.0089 | Yes |
| 3 | SRSF7 | serine/arginine-rich splicing factor 7 | 617 | 0.102 | 0.0257 | Yes |
| 4 | TYMS | thymidylate synthetase | 694 | 0.097 | 0.0383 | Yes |
| 5 | CTPS1 | CTP synthase 1 | 860 | 0.090 | 0.0450 | Yes |
| 6 | CSTF2 | cleavage stimulation factor, 3 pre-RNA, subunit 2 | 943 | 0.086 | 0.0555 | Yes |
| 7 | CDC20 | cell division cycle 20 | 1022 | 0.084 | 0.0656 | Yes |
| 8 | MRPL23 | Homo sapiens mitochondrial ribosomal protein L23 (MRPL23), mRNA. | 1163 | 0.079 | 0.0719 | Yes |
| 9 | NOP16 | NOP16 nucleolar protein | 1218 | 0.078 | 0.0824 | Yes |
| 10 | MCM4 | minichromosome maintenance complex component 4 | 1325 | 0.076 | 0.0898 | Yes |
| 11 | RRP9 | ribosomal RNA processing 9, small subunit (SSU) processome component, homolog (yeast) | 1374 | 0.075 | 0.1000 | Yes |
| 12 | HSPE1 | heat shock 10kDa protein 1 | 1413 | 0.074 | 0.1106 | Yes |
| 13 | XPO1 | exportin 1 | 1444 | 0.073 | 0.1215 | Yes |
| 14 | GOT2 | glutamic-oxaloacetic transaminase 2, mitochondrial | 1538 | 0.070 | 0.1286 | Yes |
| 15 | UBA2 | ubiquitin-like modifier activating enzyme 2 | 1562 | 0.070 | 0.1394 | Yes |
| 16 | DHX15 | DEAH (Asp-Glu-Ala-His) box helicase 15 | 1690 | 0.067 | 0.1442 | Yes |
| 17 | MCM5 | minichromosome maintenance complex component 5 | 1829 | 0.064 | 0.1480 | Yes |
| 18 | SRPK1 | SRSF protein kinase 1 | 1933 | 0.062 | 0.1533 | Yes |
| 19 | PSMC6 | proteasome 26S subunit, ATPase 6 | 1998 | 0.061 | 0.1604 | Yes |
| 20 | RANBP1 | RAN binding protein 1 | 2070 | 0.060 | 0.1670 | Yes |
| 21 | MAD2L1 | MAD2 mitotic arrest deficient-like 1 (yeast) | 2149 | 0.059 | 0.1730 | Yes |
| 22 | CCT4 | chaperonin containing TCP1, subunit 4 (delta) | 2319 | 0.056 | 0.1739 | Yes |
| 23 | NAP1L1 | nucleosome assembly protein 1-like 1 | 2338 | 0.056 | 0.1825 | Yes |
| 24 | FBL | fibrillarin | 2346 | 0.056 | 0.1917 | Yes |
| 25 | NDUFAB1 | NADH dehydrogenase (ubiquinone) 1, alpha/beta subcomplex, 1, 8kDa | 2464 | 0.054 | 0.1949 | Yes |
| 26 | EXOSC7 | exosome component 7 | 2562 | 0.053 | 0.1989 | Yes |
| 27 | SSB | Sjogren syndrome antigen B (autoantigen La) | 2634 | 0.052 | 0.2040 | Yes |
| 28 | CDK2 | cyclin-dependent kinase 2 | 2666 | 0.051 | 0.2112 | Yes |
| 29 | LSM2 | LSM2 homolog, U6 small nuclear RNA and mRNA degradation associated | 2693 | 0.051 | 0.2185 | Yes |
| 30 | SRSF3 | serine/arginine-rich splicing factor 3 | 2731 | 0.051 | 0.2253 | Yes |
| 31 | PSMB3 | proteasome subunit beta 3 | 2856 | 0.049 | 0.2272 | Yes |
| 32 | NPM1 | nucleophosmin (nucleolar phosphoprotein B23, numatrin) | 2890 | 0.049 | 0.2338 | Yes |
| 33 | SRSF2 | serine/arginine-rich splicing factor 2 | 2978 | 0.048 | 0.2374 | Yes |
| 34 | IMPDH2 | IMP (inosine 5-monophosphate) dehydrogenase 2 | 2988 | 0.048 | 0.2451 | Yes |
| 35 | HNRNPD | heterogeneous nuclear ribonucleoprotein D | 3023 | 0.047 | 0.2514 | Yes |
| 36 | CCNA2 | cyclin A2 | 3183 | 0.046 | 0.2509 | Yes |
| 37 | MYC | v-myc avian myelocytomatosis viral oncogene homolog | 3194 | 0.046 | 0.2582 | Yes |
| 38 | GLO1 | glyoxalase I | 3195 | 0.046 | 0.2659 | Yes |
| 39 | PCNA | proliferating cell nuclear antigen | 3328 | 0.044 | 0.2667 | Yes |
| 40 | MCM6 | minichromosome maintenance complex component 6 | 3558 | 0.042 | 0.2619 | Yes |
| 41 | MRPL9 | mitochondrial ribosomal protein L9 | 3631 | 0.041 | 0.2653 | Yes |
| 42 | TUFM | Tu translation elongation factor, mitochondrial | 3702 | 0.041 | 0.2685 | Yes |
| 43 | SF3B3 | splicing factor 3b subunit 3 | 3758 | 0.040 | 0.2725 | Yes |
| 44 | PWP1 | PWP1 homolog, endonuclein | 3832 | 0.039 | 0.2754 | Yes |
| 45 | NOP56 | NOP56 ribonucleoprotein | 3937 | 0.038 | 0.2765 | Yes |
| 46 | PSMA4 | proteasome subunit alpha 4 | 3948 | 0.038 | 0.2825 | Yes |
| 47 | ILF2 | interleukin enhancer binding factor 2 | 3963 | 0.038 | 0.2882 | Yes |
| 48 | PTGES3 | prostaglandin E synthase 3 (cytosolic) | 4011 | 0.037 | 0.2921 | Yes |
| 49 | CCT7 | chaperonin containing TCP1, subunit 7 (eta) | 4143 | 0.036 | 0.2915 | Yes |
| 50 | TCP1 | t-complex 1 | 4283 | 0.035 | 0.2903 | Yes |
| 51 | PRPS2 | phosphoribosyl pyrophosphate synthetase 2 | 4302 | 0.035 | 0.2952 | Yes |
| 52 | HNRNPA2B1 | heterogeneous nuclear ribonucleoprotein A2/B1 | 4338 | 0.034 | 0.2993 | Yes |
| 53 | TRA2B | transformer 2 beta homolog (Drosophila) | 4395 | 0.034 | 0.3022 | Yes |
| 54 | NME1 | NME/NM23 nucleoside diphosphate kinase 1 | 4477 | 0.033 | 0.3037 | Yes |
| 55 | APEX1 | APEX nuclease (multifunctional DNA repair enzyme) 1 | 4609 | 0.032 | 0.3024 | Yes |
| 56 | CCT2 | chaperonin containing TCP1, subunit 2 (beta) | 4704 | 0.031 | 0.3029 | Yes |
| 57 | SRM | spermidine synthase | 4706 | 0.031 | 0.3082 | Yes |
| 58 | UBE2E1 | ubiquitin conjugating enzyme E2E 1 | 4733 | 0.031 | 0.3122 | Yes |
| 59 | CCT3 | chaperonin containing TCP1, subunit 3 (gamma) | 4768 | 0.031 | 0.3157 | Yes |
| 60 | RSL1D1 | ribosomal L1 domain containing 1 | 4811 | 0.031 | 0.3187 | Yes |
| 61 | PSMD1 | proteasome 26S subunit, non-ATPase 1 | 4885 | 0.030 | 0.3201 | Yes |
| 62 | TRIM28 | tripartite motif containing 28 | 4901 | 0.030 | 0.3245 | Yes |
| 63 | MRPS18B | mitochondrial ribosomal protein S18B | 5040 | 0.029 | 0.3223 | Yes |
| 64 | PRDX3 | peroxiredoxin 3 | 5128 | 0.028 | 0.3226 | Yes |
| 65 | EIF3B | eukaryotic translation initiation factor 3, subunit B | 5164 | 0.028 | 0.3256 | Yes |
| 66 | TARDBP | TAR DNA binding protein | 5183 | 0.028 | 0.3294 | Yes |
| 67 | PHB | prohibitin | 5208 | 0.028 | 0.3329 | Yes |
| 68 | GSPT1 | G1 to S phase transition 1 | 5217 | 0.028 | 0.3371 | Yes |
| 69 | RPL14 | ribosomal protein L14 | 5269 | 0.027 | 0.3391 | Yes |
| 70 | PSMA6 | proteasome subunit alpha 6 | 5287 | 0.027 | 0.3429 | Yes |
| 71 | MCM2 | minichromosome maintenance complex component 2 | 5326 | 0.027 | 0.3455 | Yes |
| 72 | VDAC1 | voltage-dependent anion channel 1 | 5360 | 0.027 | 0.3483 | Yes |
| 73 | PRPF31 | pre-mRNA processing factor 31 | 5408 | 0.026 | 0.3503 | Yes |
| 74 | PSMA1 | proteasome subunit alpha 1 | 5435 | 0.026 | 0.3534 | Yes |
| 75 | RRM1 | ribonucleotide reductase M1 | 5510 | 0.025 | 0.3539 | Yes |
| 76 | SMARCC1 | SWI/SNF related, matrix associated, actin dependent regulator of chromatin, subfamily c, member 1 | 5534 | 0.025 | 0.3570 | Yes |
| 77 | SNRPD2 | small nuclear ribonucleoprotein D2 polypeptide | 5575 | 0.025 | 0.3591 | Yes |
| 78 | RUVBL2 | RuvB-like AAA ATPase 2 | 5599 | 0.025 | 0.3621 | Yes |
| 79 | NHP2 | NHP2 ribonucleoprotein | 5638 | 0.024 | 0.3643 | Yes |
| 80 | CCT5 | chaperonin containing TCP1, subunit 5 (epsilon) | 5644 | 0.024 | 0.3682 | Yes |
| 81 | EIF4E | eukaryotic translation initiation factor 4E | 5726 | 0.024 | 0.3681 | Yes |
| 82 | PRDX4 | peroxiredoxin 4 | 5730 | 0.024 | 0.3720 | Yes |
| 83 | PPM1G | protein phosphatase, Mg2+/Mn2+ dependent, 1G | 5742 | 0.024 | 0.3755 | Yes |
| 84 | VDAC3 | voltage-dependent anion channel 3 | 5771 | 0.024 | 0.3781 | Yes |
| 85 | RFC4 | replication factor C subunit 4 | 5802 | 0.023 | 0.3805 | Yes |
| 86 | PHB2 | prohibitin 2 | 5827 | 0.023 | 0.3832 | Yes |
| 87 | HNRNPA3 | heterogeneous nuclear ribonucleoprotein A3 | 5895 | 0.023 | 0.3836 | Yes |
| 88 | SERBP1 | SERPINE1 mRNA binding protein 1 | 5961 | 0.022 | 0.3840 | Yes |
| 89 | NOLC1 | nucleolar and coiled-body phosphoprotein 1 | 5963 | 0.022 | 0.3878 | Yes |
| 90 | HSP90AB1 | heat shock protein 90kDa alpha (cytosolic), class B member 1 | 5978 | 0.022 | 0.3908 | Yes |
| 91 | SRSF1 | serine/arginine-rich splicing factor 1 | 6000 | 0.022 | 0.3935 | Yes |
| 92 | LDHA | lactate dehydrogenase A | 6019 | 0.022 | 0.3963 | Yes |
| 93 | HSPD1 | heat shock 60kDa protein 1 (chaperonin) | 6186 | 0.021 | 0.3912 | Yes |
| 94 | KPNA2 | karyopherin alpha 2 (RAG cohort 1, importin alpha 1) | 6200 | 0.021 | 0.3941 | Yes |
| 95 | SNRPD3 | small nuclear ribonucleoprotein D3 polypeptide | 6248 | 0.021 | 0.3952 | Yes |
| 96 | PA2G4 | proliferation-associated 2G4 | 6269 | 0.020 | 0.3976 | Yes |
| 97 | RPS10 | ribosomal protein S10 | 6297 | 0.020 | 0.3997 | Yes |
| 98 | SNRPA | small nuclear ribonucleoprotein polypeptide A | 6305 | 0.020 | 0.4028 | Yes |
| 99 | SNRPG | small nuclear ribonucleoprotein polypeptide G | 6375 | 0.020 | 0.4026 | Yes |
| 100 | SNRPD1 | small nuclear ribonucleoprotein D1 polypeptide | 6481 | 0.019 | 0.4004 | Yes |
| 101 | HNRNPR | heterogeneous nuclear ribonucleoprotein R | 6539 | 0.019 | 0.4006 | Yes |
| 102 | NCBP1 | nuclear cap binding protein subunit 1 | 6575 | 0.019 | 0.4020 | Yes |
| 103 | TFDP1 | transcription factor Dp-1 | 6725 | 0.018 | 0.3972 | Yes |
| 104 | EIF1AX | eukaryotic translation initiation factor 1A, X-linked | 6738 | 0.017 | 0.3996 | Yes |
| 105 | RPL6 | ribosomal protein L6 | 6771 | 0.017 | 0.4008 | Yes |
| 106 | SNRPA1 | small nuclear ribonucleoprotein polypeptide A | 6780 | 0.017 | 0.4033 | Yes |
| 107 | RPL34 | ribosomal protein L34 | 6896 | 0.016 | 0.4002 | Yes |
| 108 | ERH | enhancer of rudimentary homolog (Drosophila) | 6916 | 0.016 | 0.4020 | Yes |
| 109 | SLC25A3 | solute carrier family 25 (mitochondrial carrier; phosphate carrier), member 3 | 7114 | 0.015 | 0.3944 | Yes |
| 110 | RNPS1 | RNA binding protein S1, serine-rich domain | 7115 | 0.015 | 0.3970 | Yes |
| 111 | PSMD8 | proteasome 26S subunit, non-ATPase 8 | 7157 | 0.015 | 0.3974 | Yes |
| 112 | RAN | RAN, member RAS oncogene family | 7181 | 0.015 | 0.3988 | Yes |
| 113 | CDK4 | cyclin-dependent kinase 4 | 7198 | 0.015 | 0.4004 | Yes |
| 114 | SYNCRIP | synaptotagmin binding, cytoplasmic RNA interacting protein | 7248 | 0.015 | 0.4004 | Yes |
| 115 | CBX3 | chromobox homolog 3 | 7262 | 0.014 | 0.4022 | Yes |
| 116 | G3BP1 | GTPase activating protein (SH3 domain) binding protein 1 | 7286 | 0.014 | 0.4034 | Yes |
| 117 | YWHAQ | tyrosine 3-monooxygenase/tryptophan 5-monooxygenase activation protein, theta | 7315 | 0.014 | 0.4043 | Yes |
| 118 | POLD2 | polymerase (DNA directed), delta 2, accessory subunit | 7486 | 0.013 | 0.3978 | No |
| 119 | ABCE1 | ATP binding cassette subfamily E member 1 | 7513 | 0.013 | 0.3986 | No |
| 120 | SET | SET nuclear proto-oncogene | 7577 | 0.012 | 0.3974 | No |
| 121 | RPL22 | ribosomal protein L22 | 7665 | 0.012 | 0.3950 | No |
| 122 | C1QBP | complement component 1, q subcomponent binding protein | 7708 | 0.012 | 0.3948 | No |
| 123 | KPNB1 | karyopherin (importin) beta 1 | 7709 | 0.012 | 0.3968 | No |
| 124 | EEF1B2 | eukaryotic translation elongation factor 1 beta 2 | 7761 | 0.011 | 0.3960 | No |
| 125 | DEK | DEK proto-oncogene | 7764 | 0.011 | 0.3979 | No |
| 126 | HNRNPC | heterogeneous nuclear ribonucleoprotein C (C1/C2) | 7972 | 0.010 | 0.3889 | No |
| 127 | MCM7 | minichromosome maintenance complex component 7 | 7985 | 0.010 | 0.3899 | No |
| 128 | RPS2 | ribosomal protein S2 | 8092 | 0.009 | 0.3860 | No |
| 129 | CLNS1A | chloride channel, nucleotide-sensitive, 1A | 8102 | 0.009 | 0.3871 | No |
| 130 | POLE3 | polymerase (DNA directed), epsilon 3, accessory subunit | 8126 | 0.009 | 0.3875 | No |
| 131 | EIF4A1 | eukaryotic translation initiation factor 4A1 | 8146 | 0.009 | 0.3881 | No |
| 132 | RPS3 | ribosomal protein S3 | 8225 | 0.008 | 0.3855 | No |
| 133 | NCBP2 | nuclear cap binding protein subunit 2 | 8274 | 0.008 | 0.3844 | No |
| 134 | PSMD14 | proteasome 26S subunit, non-ATPase 14 | 8280 | 0.008 | 0.3855 | No |
| 135 | XRCC6 | X-ray repair complementing defective repair in Chinese hamster cells 6 | 8395 | 0.007 | 0.3809 | No |
| 136 | YWHAE | tyrosine 3-monooxygenase/tryptophan 5-monooxygenase activation protein, epsilon | 8419 | 0.007 | 0.3809 | No |
| 137 | HNRNPU | heterogeneous nuclear ribonucleoprotein U (scaffold attachment factor A) | 8561 | 0.007 | 0.3748 | No |
| 138 | PPIA | peptidylprolyl isomerase A (cyclophilin A) | 8627 | 0.006 | 0.3724 | No |
| 139 | HNRNPA1 | heterogeneous nuclear ribonucleoprotein A1 | 8814 | 0.005 | 0.3637 | No |
| 140 | COX5A | cytochrome c oxidase subunit Va | 8864 | 0.005 | 0.3620 | No |
| 141 | PSMD7 | proteasome 26S subunit, non-ATPase 7 | 8885 | 0.005 | 0.3618 | No |
| 142 | LSM7 | LSM7 homolog, U6 small nuclear RNA and mRNA degradation associated | 8914 | 0.005 | 0.3611 | No |
| 143 | PSMD3 | proteasome 26S subunit, non-ATPase 3 | 9044 | 0.004 | 0.3551 | No |
| 144 | STARD7 | StAR-related lipid transfer domain containing 7 | 9344 | 0.002 | 0.3400 | No |
| 145 | EIF3D | eukaryotic translation initiation factor 3, subunit D | 9586 | 0.001 | 0.3277 | No |
| 146 | DDX18 | DEAD (Asp-Glu-Ala-Asp) box polypeptide 18 | 9630 | 0.001 | 0.3256 | No |
| 147 | HDDC2 | HD domain containing 2 | 9770 | 0.000 | 0.3184 | No |
| 148 | PSMB2 | proteasome subunit beta 2 | 9881 | -0.001 | 0.3128 | No |
| 149 | RPS6 | ribosomal protein S6 | 10058 | -0.001 | 0.3040 | No |
| 150 | RPS5 | ribosomal protein S5 | 10127 | -0.002 | 0.3008 | No |
| 151 | BUB3 | BUB3 mitotic checkpoint protein | 10144 | -0.002 | 0.3003 | No |
| 152 | HDGF | hepatoma-derived growth factor | 10182 | -0.002 | 0.2987 | No |
| 153 | PSMA7 | proteasome subunit alpha 7 | 10203 | -0.002 | 0.2981 | No |
| 154 | GNL3 | guanine nucleotide binding protein-like 3 (nucleolar) | 10216 | -0.002 | 0.2979 | No |
| 155 | EIF2S1 | eukaryotic translation initiation factor 2, subunit 1 alpha, 35kDa | 10272 | -0.003 | 0.2955 | No |
| 156 | HDAC2 | histone deacetylase 2 | 10584 | -0.005 | 0.2802 | No |
| 157 | RPLP0 | ribosomal protein, large, P0 | 10652 | -0.005 | 0.2776 | No |
| 158 | PCBP1 | poly(rC) binding protein 1 | 10753 | -0.006 | 0.2734 | No |
| 159 | EIF4G2 | eukaryotic translation initiation factor 4 gamma, 2 | 10924 | -0.007 | 0.2657 | No |
| 160 | RPL18 | ribosomal protein L18 | 11076 | -0.008 | 0.2592 | No |
| 161 | PGK1 | phosphoglycerate kinase 1 | 11128 | -0.008 | 0.2579 | No |
| 162 | UBE2L3 | ubiquitin conjugating enzyme E2L 3 | 11130 | -0.008 | 0.2592 | No |
| 163 | SNRPB2 | small nuclear ribonucleoprotein polypeptide B | 11785 | -0.012 | 0.2273 | No |
| 164 | SF3A1 | splicing factor 3a, subunit 1, 120kDa | 11859 | -0.012 | 0.2256 | No |
| 165 | PSMA2 | proteasome subunit alpha 2 | 11937 | -0.012 | 0.2237 | No |
| 166 | DDX21 | DEAD (Asp-Glu-Ala-Asp) box helicase 21 | 12006 | -0.013 | 0.2224 | No |
| 167 | CYC1 | cytochrome c-1 | 12040 | -0.013 | 0.2229 | No |
| 168 | TXNL4A | thioredoxin-like 4A | 12202 | -0.014 | 0.2170 | No |
| 169 | PABPC1 | poly(A) binding protein, cytoplasmic 1 | 12244 | -0.014 | 0.2173 | No |
| 170 | RAD23B | RAD23 homolog B, nucleotide excision repair protein | 12313 | -0.015 | 0.2163 | No |
| 171 | U2AF1 | U2 small nuclear RNA auxiliary factor 1 | 12604 | -0.017 | 0.2042 | No |
| 172 | ETF1 | eukaryotic translation termination factor 1 | 12626 | -0.017 | 0.2060 | No |
| 173 | USP1 | ubiquitin specific peptidase 1 | 12752 | -0.018 | 0.2025 | No |
| 174 | PSMC4 | proteasome 26S subunit, ATPase 4 | 12803 | -0.018 | 0.2030 | No |
| 175 | FAM120A | family with sequence similarity 120A | 13066 | -0.020 | 0.1928 | No |
| 176 | ODC1 | ornithine decarboxylase 1 | 13133 | -0.020 | 0.1928 | No |
| 177 | CNBP | CCHC-type zinc finger, nucleic acid binding protein | 13274 | -0.021 | 0.1891 | No |
| 178 | EIF3J | eukaryotic translation initiation factor 3, subunit J | 13279 | -0.021 | 0.1925 | No |
| 179 | PABPC4 | poly(A) binding protein, cytoplasmic 4 (inducible form) | 13319 | -0.021 | 0.1941 | No |
| 180 | CANX | calnexin | 13700 | -0.024 | 0.1786 | No |
| 181 | SSBP1 | single-stranded DNA binding protein 1, mitochondrial | 14095 | -0.027 | 0.1628 | No |
| 182 | EIF2S2 | eukaryotic translation initiation factor 2, subunit 2 beta, 38kDa | 14130 | -0.027 | 0.1657 | No |
| 183 | CAD | carbamoyl-phosphate synthetase 2, aspartate transcarbamylase, and dihydroorotase | 14416 | -0.030 | 0.1560 | No |
| 184 | ACP1 | acid phosphatase 1, soluble | 14417 | -0.030 | 0.1610 | No |
| 185 | VBP1 | von Hippel-Lindau binding protein 1 | 14451 | -0.030 | 0.1644 | No |
| 186 | AIMP2 | aminoacyl tRNA synthetase complex-interacting multifunctional protein 2 | 15543 | -0.039 | 0.1147 | No |
| 187 | EIF4H | eukaryotic translation initiation factor 4H | 15610 | -0.040 | 0.1180 | No |
| 188 | COPS5 | COP9 signalosome subunit 5 | 15670 | -0.041 | 0.1219 | No |
| 189 | CUL1 | cullin 1 | 15794 | -0.042 | 0.1226 | No |
| 190 | AP3S1 | adaptor-related protein complex 3, sigma 1 subunit | 16602 | -0.051 | 0.0896 | No |
| 191 | ORC2 | origin recognition complex subunit 2 | 17197 | -0.060 | 0.0691 | No |
| 192 | XPOT | exportin, tRNA | 17359 | -0.063 | 0.0715 | No |
| 193 | DUT | deoxyuridine triphosphatase | 18343 | -0.087 | 0.0354 | No |
| 194 | IFRD1 | interferon-related developmental regulator 1 | 19158 | -0.151 | 0.0189 | No |
Table: GSEA details [plain text format]

  

Fig 2: HALLMARK\_MYC\_TARGETS\_V1      
 Blue-Pink O' Gram in the Space of the Analyzed GeneSet

  

Fig 3: HALLMARK\_MYC\_TARGETS\_V1: Random ES distribution      
 Gene set null distribution of ES for **HALLMARK\_MYC\_TARGETS\_V1**

  
