## Supplementary material for "Somatic hypomethylation of pericentromeric SST1 repeats and tetraploidization in human colorectal cancer cells": GSEA results: HALLMARK_P53_PATHWAY.html

Details for gene set HALLMARK\_P53\_PATHWAY[GSEA]

|  || Dataset | eset\_byprobe\_collapsed\_to\_symbols.Diploid\_vs\_Tetraploid.cls #Tetraploid\_versus\_Diploid.Diploid\_vs\_Tetraploid.cls #Tetraploid\_versus\_Diploid\_repos |
| Phenotype | Diploid\_vs\_Tetraploid.cls#Tetraploid\_versus\_Diploid\_repos |
| Upregulated in class | Tetraploid |
| GeneSet | HALLMARK\_P53\_PATHWAY |
| Enrichment Score (ES) | 0.35002476 |
| Normalized Enrichment Score (NES) | 1.5265676 |
| Nominal p-value | 0.0 |
| FDR q-value | 0.029029533 |
| FWER p-Value | 0.126 |
Table: GSEA Results Summary

  

Fig 1: Enrichment plot: HALLMARK\_P53\_PATHWAY      
 Profile of the Running ES Score & Positions of GeneSet Members on the Rank Ordered List

  

| SYMBOL | TITLE | RANK IN GENE LIST | RANK METRIC SCORE | RUNNING ES | CORE ENRICHMENT || 1 | S100A4 | S100 calcium binding protein A4 | 5 | 0.471 | 0.0413 | Yes |
| 2 | SERPINB5 | serpin peptidase inhibitor, clade B (ovalbumin), member 5 | 31 | 0.322 | 0.0684 | Yes |
| 3 | CCND2 | cyclin D2 | 34 | 0.313 | 0.0959 | Yes |
| 4 | TM4SF1 | transmembrane 4 L six family member 1 | 81 | 0.225 | 0.1134 | Yes |
| 5 | PLK2 | polo-like kinase 2 | 109 | 0.202 | 0.1297 | Yes |
| 6 | SESN1 | sestrin 1 | 121 | 0.190 | 0.1459 | Yes |
| 7 | GLS2 | glutaminase 2 (liver, mitochondrial) | 149 | 0.172 | 0.1597 | Yes |
| 8 | FAS | Fas cell surface death receptor | 186 | 0.161 | 0.1721 | Yes |
| 9 | DRAM1 | DNA-damage regulated autophagy modulator 1 | 191 | 0.158 | 0.1858 | Yes |
| 10 | TNFSF9 | tumor necrosis factor (ligand) superfamily, member 9 | 223 | 0.150 | 0.1974 | Yes |
| 11 | IL1A | interleukin 1 alpha | 237 | 0.146 | 0.2096 | Yes |
| 12 | BTG2 | BTG family, member 2 | 281 | 0.136 | 0.2194 | Yes |
| 13 | TOB1 | transducer of ERBB2, 1 | 343 | 0.127 | 0.2274 | Yes |
| 14 | PROCR | protein C receptor, endothelial | 384 | 0.121 | 0.2360 | Yes |
| 15 | LIF | leukemia inhibitory factor | 423 | 0.117 | 0.2444 | Yes |
| 16 | MDM2 | MDM2 proto-oncogene, E3 ubiquitin protein ligase | 433 | 0.116 | 0.2541 | Yes |
| 17 | DGKA | diacylglycerol kinase alpha | 470 | 0.112 | 0.2622 | Yes |
| 18 | ZMAT3 | zinc finger, matrin-type 3 | 525 | 0.108 | 0.2689 | Yes |
| 19 | POLH | polymerase (DNA directed), eta | 545 | 0.106 | 0.2773 | Yes |
| 20 | CYFIP2 | cytoplasmic FMR1 interacting protein 2 | 557 | 0.106 | 0.2861 | Yes |
| 21 | IER5 | immediate early response 5 | 657 | 0.100 | 0.2898 | Yes |
| 22 | STEAP3 | STEAP family member 3, metalloreductase | 677 | 0.098 | 0.2974 | Yes |
| 23 | CDKN1A | cyclin-dependent kinase inhibitor 1A (p21, Cip1) | 690 | 0.097 | 0.3054 | Yes |
| 24 | S100A10 | S100 calcium binding protein A10 | 886 | 0.089 | 0.3031 | Yes |
| 25 | HSPA4L | heat shock 70kDa protein 4-like | 1070 | 0.082 | 0.3009 | Yes |
| 26 | CTSF | cathepsin F | 1146 | 0.080 | 0.3041 | Yes |
| 27 | FDXR | ferredoxin reductase | 1184 | 0.079 | 0.3091 | Yes |
| 28 | NOTCH1 | notch 1 | 1193 | 0.079 | 0.3156 | Yes |
| 29 | CDKN2A | cyclin-dependent kinase inhibitor 2A | 1257 | 0.077 | 0.3192 | Yes |
| 30 | ABCC5 | ATP binding cassette subfamily C member 5 | 1283 | 0.077 | 0.3246 | Yes |
| 31 | PITPNC1 | phosphatidylinositol transfer protein, cytoplasmic 1 | 1540 | 0.070 | 0.3176 | Yes |
| 32 | RPS27L | ribosomal protein S27-like | 1564 | 0.070 | 0.3226 | Yes |
| 33 | SPHK1 | sphingosine kinase 1 | 1575 | 0.070 | 0.3282 | Yes |
| 34 | PHLDA3 | pleckstrin homology-like domain, family A, member 3 | 1617 | 0.068 | 0.3321 | Yes |
| 35 | TRIAP1 | TP53 regulated inhibitor of apoptosis 1 | 1623 | 0.068 | 0.3379 | Yes |
| 36 | DDB2 | damage-specific DNA binding protein 2 | 1815 | 0.065 | 0.3337 | Yes |
| 37 | ADA | adenosine deaminase | 1881 | 0.063 | 0.3360 | Yes |
| 38 | TP63 | tumor protein p63 | 1907 | 0.063 | 0.3402 | Yes |
| 39 | PPM1D | protein phosphatase, Mg2+/Mn2+ dependent, 1D | 1919 | 0.063 | 0.3452 | Yes |
| 40 | SFN | stratifin | 2005 | 0.061 | 0.3462 | Yes |
| 41 | NHLH2 | nescient helix-loop-helix 2 | 2036 | 0.061 | 0.3500 | Yes |
| 42 | CD82 | CD82 molecule | 2272 | 0.057 | 0.3429 | No |
| 43 | CCNG1 | cyclin G1 | 2329 | 0.056 | 0.3450 | No |
| 44 | RAD51C | RAD51 paralog C | 2687 | 0.051 | 0.3310 | No |
| 45 | BLCAP | bladder cancer associated protein | 2822 | 0.050 | 0.3285 | No |
| 46 | ITGB4 | integrin beta 4 | 3024 | 0.047 | 0.3223 | No |
| 47 | BAX | BCL2-associated X protein | 3143 | 0.046 | 0.3202 | No |
| 48 | RB1 | retinoblastoma 1 | 3205 | 0.046 | 0.3211 | No |
| 49 | SLC19A2 | solute carrier family 19 (thiamine transporter), member 2 | 3211 | 0.045 | 0.3248 | No |
| 50 | EPS8L2 | EPS8-like 2 | 3294 | 0.045 | 0.3245 | No |
| 51 | NOL8 | nucleolar protein 8 | 3299 | 0.045 | 0.3283 | No |
| 52 | CD81 | CD81 molecule | 3320 | 0.044 | 0.3312 | No |
| 53 | PCNA | proliferating cell nuclear antigen | 3328 | 0.044 | 0.3347 | No |
| 54 | HMOX1 | heme oxygenase 1 | 3502 | 0.043 | 0.3295 | No |
| 55 | TCHH | trichohyalin | 3503 | 0.043 | 0.3333 | No |
| 56 | EI24 | etoposide induced 2.4 | 3581 | 0.042 | 0.3330 | No |
| 57 | VWA5A | von Willebrand factor A domain containing 5A | 3657 | 0.041 | 0.3327 | No |
| 58 | SLC35D1 | solute carrier family 35 (UDP-GlcA/UDP-GalNAc transporter), member D1 | 3925 | 0.038 | 0.3223 | No |
| 59 | AEN | apoptosis enhancing nuclease | 4089 | 0.037 | 0.3171 | No |
| 60 | EPHA2 | EPH receptor A2 | 4100 | 0.037 | 0.3198 | No |
| 61 | ISCU | iron-sulfur cluster assembly enzyme | 4109 | 0.037 | 0.3226 | No |
| 62 | AK1 | adenylate kinase 1 | 4140 | 0.036 | 0.3243 | No |
| 63 | HRAS | Harvey rat sarcoma viral oncogene homolog | 4211 | 0.035 | 0.3238 | No |
| 64 | RAP2B | RAP2B, member of RAS oncogene family | 4263 | 0.035 | 0.3242 | No |
| 65 | TGFA | transforming growth factor alpha | 4449 | 0.034 | 0.3176 | No |
| 66 | FBXW7 | F-box and WD repeat domain containing 7, E3 ubiquitin protein ligase | 4459 | 0.033 | 0.3201 | No |
| 67 | TPD52L1 | tumor protein D52-like 1 | 4623 | 0.032 | 0.3145 | No |
| 68 | PIDD1 | p53-induced death domain protein 1 | 4690 | 0.032 | 0.3139 | No |
| 69 | TPRKB | TP53RK binding protein | 4879 | 0.030 | 0.3068 | No |
| 70 | PMM1 | phosphomannomutase 1 | 4930 | 0.030 | 0.3069 | No |
| 71 | KIF13B | kinesin family member 13B | 4976 | 0.029 | 0.3072 | No |
| 72 | LDHB | lactate dehydrogenase B | 5017 | 0.029 | 0.3077 | No |
| 73 | CDK5R1 | cyclin-dependent kinase 5, regulatory subunit 1 (p35) | 5166 | 0.028 | 0.3025 | No |
| 74 | SDC1 | syndecan 1 | 5305 | 0.027 | 0.2977 | No |
| 75 | DCXR | dicarbonyl/L-xylulose reductase | 5337 | 0.027 | 0.2985 | No |
| 76 | PDGFA | platelet-derived growth factor alpha polypeptide | 5403 | 0.026 | 0.2974 | No |
| 77 | TRAF4 | TNF receptor-associated factor 4 | 5464 | 0.026 | 0.2966 | No |
| 78 | DNTTIP2 | deoxynucleotidyltransferase, terminal, interacting protein 2 | 5636 | 0.024 | 0.2899 | No |
| 79 | TAP1 | transporter 1, ATP-binding cassette, sub-family B (MDR/TAP) | 6169 | 0.021 | 0.2643 | No |
| 80 | KLK8 | kallikrein related peptidase 8 | 6224 | 0.021 | 0.2633 | No |
| 81 | APP | amyloid beta (A4) precursor protein | 6318 | 0.020 | 0.2603 | No |
| 82 | HINT1 | histidine triad nucleotide binding protein 1 | 6495 | 0.019 | 0.2528 | No |
| 83 | SERTAD3 | SERTA domain containing 3 | 6579 | 0.018 | 0.2502 | No |
| 84 | GPX2 | glutathione peroxidase 2 | 6587 | 0.018 | 0.2514 | No |
| 85 | TM7SF3 | transmembrane 7 superfamily member 3 | 6597 | 0.018 | 0.2526 | No |
| 86 | WRAP73 | WD repeat containing, antisense to TP73 | 6707 | 0.018 | 0.2485 | No |
| 87 | ABAT | 4-aminobutyrate aminotransferase | 6739 | 0.017 | 0.2484 | No |
| 88 | PLK3 | polo-like kinase 3 | 6907 | 0.016 | 0.2413 | No |
| 89 | KRT17 | keratin 17, type I | 7096 | 0.015 | 0.2329 | No |
| 90 | RCHY1 | ring finger and CHY zinc finger domain containing 1, E3 ubiquitin protein ligase | 7271 | 0.014 | 0.2252 | No |
| 91 | GADD45A | growth arrest and DNA-damage-inducible, alpha | 7290 | 0.014 | 0.2255 | No |
| 92 | ZBTB16 | zinc finger and BTB domain containing 16 | 7433 | 0.013 | 0.2193 | No |
| 93 | XPC | xeroderma pigmentosum, complementation group C | 7484 | 0.013 | 0.2179 | No |
| 94 | BTG1 | B-cell translocation gene 1, anti-proliferative | 7702 | 0.012 | 0.2077 | No |
| 95 | RXRA | retinoid X receptor alpha | 7732 | 0.011 | 0.2072 | No |
| 96 | CDKN2AIP | CDKN2A interacting protein | 7775 | 0.011 | 0.2060 | No |
| 97 | FOXO3 | forkhead box O3 | 8135 | 0.009 | 0.1882 | No |
| 98 | RHBDF2 | rhomboid 5 homolog 2 (Drosophila) | 8277 | 0.008 | 0.1817 | No |
| 99 | VDR | vitamin D (1,25- dihydroxyvitamin D3) receptor | 8293 | 0.008 | 0.1816 | No |
| 100 | RALGDS | ral guanine nucleotide dissociation stimulator | 8453 | 0.007 | 0.1740 | No |
| 101 | ACVR1B | activin A receptor type IB | 8737 | 0.006 | 0.1598 | No |
| 102 | PTPRE | protein tyrosine phosphatase, receptor type, E | 8781 | 0.005 | 0.1581 | No |
| 103 | ZNF365 | zinc finger protein 365 | 8808 | 0.005 | 0.1572 | No |
| 104 | NUDT15 | nudix hydrolase 15 | 8869 | 0.005 | 0.1545 | No |
| 105 | NDRG1 | N-myc downstream regulated 1 | 9078 | 0.004 | 0.1441 | No |
| 106 | CGRRF1 | cell growth regulator with ring finger domain 1 | 9361 | 0.002 | 0.1297 | No |
| 107 | HEXIM1 | hexamethylene bis-acetamide inducible 1 | 9408 | 0.002 | 0.1275 | No |
| 108 | BAIAP2 | BAI1-associated protein 2 | 9661 | 0.001 | 0.1145 | No |
| 109 | RNF19B | Memczak2013 ANTISENSE, CDS, coding, INTERNAL best transcript NM\_153341 | 9730 | 0.000 | 0.1110 | No |
| 110 | JAG2 | jagged 2 | 10208 | -0.002 | 0.0865 | No |
| 111 | RPS12 | ribosomal protein S12 | 10377 | -0.003 | 0.0782 | No |
| 112 | HDAC3 | histone deacetylase 3 | 10633 | -0.005 | 0.0654 | No |
| 113 | FAM162A | family with sequence similarity 162, member A | 10663 | -0.005 | 0.0644 | No |
| 114 | NINJ1 | ninjurin 1 | 10805 | -0.006 | 0.0576 | No |
| 115 | DEF6 | DEF6 guanine nucleotide exchange factor | 10849 | -0.006 | 0.0559 | No |
| 116 | RGS16 | regulator of G-protein signaling 16 | 11032 | -0.007 | 0.0471 | No |
| 117 | RPL18 | ribosomal protein L18 | 11076 | -0.008 | 0.0456 | No |
| 118 | PRKAB1 | protein kinase, AMP-activated, beta 1 non-catalytic subunit | 11678 | -0.011 | 0.0155 | No |
| 119 | BAK1 | BCL2-antagonist/killer 1 | 11698 | -0.011 | 0.0155 | No |
| 120 | SP1 | Sp1 transcription factor | 11717 | -0.011 | 0.0155 | No |
| 121 | RAD9A | RAD9 checkpoint clamp component A | 11909 | -0.012 | 0.0067 | No |
| 122 | RRP8 | Transcript Identified by AceView, Entrez Gene ID(s) 23378 | 11966 | -0.013 | 0.0049 | No |
| 123 | INHBB | inhibin beta B | 12019 | -0.013 | 0.0034 | No |
| 124 | PTPN14 | protein tyrosine phosphatase, non-receptor type 14 | 12097 | -0.013 | 0.0006 | No |
| 125 | WWP1 | WW domain containing E3 ubiquitin protein ligase 1 | 12204 | -0.014 | -0.0037 | No |
| 126 | ZFP36L1 | ZFP36 ring finger protein-like 1 | 12249 | -0.014 | -0.0047 | No |
| 127 | IP6K2 | inositol hexakisphosphate kinase 2 | 12320 | -0.015 | -0.0070 | No |
| 128 | TGFB1 | transforming growth factor beta 1 | 12464 | -0.016 | -0.0129 | No |
| 129 | SOCS1 | suppressor of cytokine signaling 1 | 12611 | -0.017 | -0.0190 | No |
| 130 | ERCC5 | excision repair cross-complementation group 5 | 12684 | -0.017 | -0.0212 | No |
| 131 | GM2A | GM2 ganglioside activator | 12726 | -0.017 | -0.0218 | No |
| 132 | EPHX1 | epoxide hydrolase 1, microsomal (xenobiotic) | 12904 | -0.019 | -0.0293 | No |
| 133 | FUCA1 | fucosidase, alpha-L- 1, tissue | 13114 | -0.020 | -0.0384 | No |
| 134 | F2R | coagulation factor II (thrombin) receptor | 13235 | -0.021 | -0.0427 | No |
| 135 | VAMP8 | vesicle associated membrane protein 8 | 13409 | -0.022 | -0.0497 | No |
| 136 | TSC22D1 | TSC22 domain family, member 1 | 14022 | -0.026 | -0.0791 | No |
| 137 | TP53 | tumor protein p53 | 14048 | -0.027 | -0.0780 | No |
| 138 | IER3 | immediate early response 3 | 14076 | -0.027 | -0.0770 | No |
| 139 | ST14 | suppression of tumorigenicity 14 (colon carcinoma) | 14096 | -0.027 | -0.0756 | No |
| 140 | IRAK1 | interleukin 1 receptor associated kinase 1 | 14157 | -0.028 | -0.0763 | No |
| 141 | PERP | PERP, TP53 apoptosis effector | 14176 | -0.028 | -0.0748 | No |
| 142 | RRAD | Ras-related associated with diabetes | 14315 | -0.029 | -0.0794 | No |
| 143 | CEBPA | CCAAT/enhancer binding protein (C/EBP), alpha | 14543 | -0.031 | -0.0884 | No |
| 144 | TCN2 | transcobalamin II | 14548 | -0.031 | -0.0859 | No |
| 145 | HBEGF | heparin-binding EGF-like growth factor | 14753 | -0.032 | -0.0936 | No |
| 146 | PLXNB2 | plexin B2 | 14771 | -0.033 | -0.0916 | No |
| 147 | MXD4 | MAX dimerization protein 4 | 14872 | -0.033 | -0.0938 | No |
| 148 | CCND3 | cyclin D3 | 14958 | -0.034 | -0.0952 | No |
| 149 | CDKN2B | cyclin-dependent kinase inhibitor 2B (p15, inhibits CDK4) | 15064 | -0.035 | -0.0976 | No |
| 150 | RAB40C | RAB40C, member RAS oncogene family | 15204 | -0.036 | -0.1016 | No |
| 151 | CLCA2 | chloride channel accessory 2 | 15293 | -0.037 | -0.1029 | No |
| 152 | POM121 | POM121 transmembrane nucleoporin | 15476 | -0.039 | -0.1089 | No |
| 153 | TNNI1 | troponin I type 1 (skeletal, slow) | 15521 | -0.039 | -0.1077 | No |
| 154 | MAPKAPK3 | mitogen-activated protein kinase-activated protein kinase 3 | 15722 | -0.041 | -0.1144 | No |
| 155 | TXNIP | thioredoxin interacting protein | 16097 | -0.045 | -0.1298 | No |
| 156 | APAF1 | apoptotic peptidase activating factor 1 | 16221 | -0.046 | -0.1321 | No |
| 157 | BMP2 | bone morphogenetic protein 2 | 16350 | -0.048 | -0.1345 | No |
| 158 | TAX1BP3 | Tax1 (human T-cell leukemia virus type I) binding protein 3 | 16466 | -0.049 | -0.1361 | No |
| 159 | ALOX15B | arachidonate 15-lipoxygenase, type B | 16555 | -0.051 | -0.1362 | No |
| 160 | MKNK2 | MAP kinase interacting serine/threonine kinase 2 | 17133 | -0.059 | -0.1608 | No |
| 161 | TSPYL2 | TSPY-like 2 | 17146 | -0.059 | -0.1562 | No |
| 162 | OSGIN1 | oxidative stress induced growth inhibitor 1 | 17246 | -0.061 | -0.1560 | No |
| 163 | CSRNP2 | cysteine-serine-rich nuclear protein 2 | 17338 | -0.062 | -0.1552 | No |
| 164 | FGF13 | fibroblast growth factor 13 | 17417 | -0.064 | -0.1536 | No |
| 165 | FOS | FBJ murine osteosarcoma viral oncogene homolog | 17528 | -0.066 | -0.1535 | No |
| 166 | DDIT4 | DNA damage inducible transcript 4 | 17557 | -0.066 | -0.1491 | No |
| 167 | TRAFD1 | TRAF-type zinc finger domain containing 1 | 17666 | -0.069 | -0.1486 | No |
| 168 | RETSAT | retinol saturase (all-trans-retinol 13,14-reductase) | 17683 | -0.069 | -0.1433 | No |
| 169 | CCP110 | centriolar coiled coil protein 110kDa | 17778 | -0.071 | -0.1419 | No |
| 170 | PRMT2 | protein arginine methyltransferase 2 | 17836 | -0.073 | -0.1384 | No |
| 171 | CCNK | cyclin K | 17948 | -0.076 | -0.1375 | No |
| 172 | NUPR1 | nuclear protein 1, transcriptional regulator | 17958 | -0.076 | -0.1312 | No |
| 173 | KLF4 | Kruppel-like factor 4 (gut) | 17974 | -0.076 | -0.1253 | No |
| 174 | ANKRA2 | ankyrin repeat, family A (RFXANK-like), 2 | 18100 | -0.080 | -0.1247 | No |
| 175 | SAT1 | spermidine/spermine N1-acetyltransferase 1 | 18295 | -0.085 | -0.1272 | No |
| 176 | MXD1 | Memczak2013 ANTISENSE, coding, INTERNAL, intronic best transcript NM\_001202514 | 18336 | -0.086 | -0.1217 | No |
| 177 | CDH13 | cadherin 13 | 18362 | -0.087 | -0.1153 | No |
| 178 | UPP1 | uridine phosphorylase 1 | 18404 | -0.089 | -0.1095 | No |
| 179 | CTSD | cathepsin D | 18557 | -0.095 | -0.1090 | No |
| 180 | SLC7A11 | solute carrier family 7 (anionic amino acid transporter light chain, xc- system), member 11 | 18650 | -0.099 | -0.1050 | No |
| 181 | SLC3A2 | solute carrier family 3 (amino acid transporter heavy chain), member 2 | 18787 | -0.109 | -0.1025 | No |
| 182 | SEC61A1 | Sec61 translocon alpha 1 subunit | 18812 | -0.111 | -0.0940 | No |
| 183 | PPP1R15A | protein phosphatase 1, regulatory subunit 15A | 18867 | -0.115 | -0.0866 | No |
| 184 | TRIB3 | tribbles pseudokinase 3 | 18870 | -0.115 | -0.0765 | No |
| 185 | JUN | jun proto-oncogene | 19037 | -0.132 | -0.0734 | No |
| 186 | STOM | stomatin | 19039 | -0.132 | -0.0618 | No |
| 187 | ABHD4 | abhydrolase domain containing 4 | 19071 | -0.138 | -0.0513 | No |
| 188 | ATF3 | activating transcription factor 3 | 19421 | -0.254 | -0.0469 | No |
| 189 | DDIT3 | DNA-damage-inducible transcript 3 | 19440 | -0.282 | -0.0229 | No |
| 190 | CASP1 | caspase 1 | 19457 | -0.309 | 0.0035 | No |
Table: GSEA details [plain text format]

  

Fig 2: HALLMARK\_P53\_PATHWAY      
 Blue-Pink O' Gram in the Space of the Analyzed GeneSet

  

Fig 3: HALLMARK\_P53\_PATHWAY: Random ES distribution      
 Gene set null distribution of ES for **HALLMARK\_P53\_PATHWAY**

  
