## Supplementary material for "Somatic hypomethylation of pericentromeric SST1 repeats and tetraploidization in human colorectal cancer cells": GSEA results: HALLMARK_PI3K_AKT_MTOR_SIGNALING.html

Details for gene set HALLMARK\_PI3K\_AKT\_MTOR\_SIGNALING[GSEA]

|  || Dataset | eset\_byprobe\_collapsed\_to\_symbols.Diploid\_vs\_Tetraploid.cls #Tetraploid\_versus\_Diploid.Diploid\_vs\_Tetraploid.cls #Tetraploid\_versus\_Diploid\_repos |
| Phenotype | Diploid\_vs\_Tetraploid.cls#Tetraploid\_versus\_Diploid\_repos |
| Upregulated in class | Diploid |
| GeneSet | HALLMARK\_PI3K\_AKT\_MTOR\_SIGNALING |
| Enrichment Score (ES) | -0.34884182 |
| Normalized Enrichment Score (NES) | -1.316761 |
| Nominal p-value | 0.06482982 |
| FDR q-value | 0.26437563 |
| FWER p-Value | 0.853 |
Table: GSEA Results Summary

  

Fig 1: Enrichment plot: HALLMARK\_PI3K\_AKT\_MTOR\_SIGNALING      
 Profile of the Running ES Score & Positions of GeneSet Members on the Rank Ordered List

  

| SYMBOL | TITLE | RANK IN GENE LIST | RANK METRIC SCORE | RUNNING ES | CORE ENRICHMENT || 1 | PRKAA2 | protein kinase, AMP-activated, alpha 2 catalytic subunit | 57 | 0.275 | 0.0567 | No |
| 2 | IL2RG | interleukin 2 receptor, gamma | 268 | 0.137 | 0.0758 | No |
| 3 | CDKN1A | cyclin-dependent kinase inhibitor 1A (p21, Cip1) | 690 | 0.097 | 0.0752 | No |
| 4 | TRAF2 | TNF receptor-associated factor 2 | 907 | 0.088 | 0.0832 | No |
| 5 | FASLG | Fas ligand (TNF superfamily, member 6) | 1039 | 0.083 | 0.0944 | No |
| 6 | EGFR | epidermal growth factor receptor | 1396 | 0.074 | 0.0922 | No |
| 7 | ECSIT | ECSIT signalling integrator | 1813 | 0.065 | 0.0848 | No |
| 8 | SFN | stratifin | 2005 | 0.061 | 0.0883 | No |
| 9 | TIAM1 | T-cell lymphoma invasion and metastasis 1 | 2375 | 0.056 | 0.0813 | No |
| 10 | FGF17 | fibroblast growth factor 17 | 2615 | 0.052 | 0.0804 | No |
| 11 | CDK2 | cyclin-dependent kinase 2 | 2666 | 0.051 | 0.0889 | No |
| 12 | MKNK1 | MAP kinase interacting serine/threonine kinase 1 | 2996 | 0.048 | 0.0823 | No |
| 13 | FGF22 | fibroblast growth factor 22 | 3480 | 0.043 | 0.0667 | No |
| 14 | HRAS | Harvey rat sarcoma viral oncogene homolog | 4211 | 0.035 | 0.0369 | No |
| 15 | PLCG1 | phospholipase C, gamma 1 | 4485 | 0.033 | 0.0300 | No |
| 16 | RPS6KA1 | ribosomal protein S6 kinase, 90kDa, polypeptide 1 | 4554 | 0.033 | 0.0336 | No |
| 17 | CDK1 | cyclin-dependent kinase 1 | 4823 | 0.031 | 0.0265 | No |
| 18 | UBE2N | ubiquitin conjugating enzyme E2N | 4992 | 0.029 | 0.0242 | No |
| 19 | ATF1 | activating transcription factor 1 | 5471 | 0.026 | 0.0052 | No |
| 20 | NGF | nerve growth factor (beta polypeptide) | 5632 | 0.025 | 0.0023 | No |
| 21 | AKT1S1 | Memczak2013 ANTISENSE, coding, INTERNAL, intronic best transcript NM\_001098633 | 5709 | 0.024 | 0.0036 | No |
| 22 | EIF4E | eukaryotic translation initiation factor 4E | 5726 | 0.024 | 0.0080 | No |
| 23 | RPTOR | regulatory associated protein of MTOR, complex 1 | 5738 | 0.024 | 0.0126 | No |
| 24 | LCK | LCK proto-oncogene, Src family tyrosine kinase | 6061 | 0.022 | 0.0007 | No |
| 25 | PIN1 | peptidylprolyl cis/trans isomerase, NIMA-interacting 1 | 6262 | 0.020 | -0.0052 | No |
| 26 | MAPK9 | mitogen-activated protein kinase 9 | 6304 | 0.020 | -0.0029 | No |
| 27 | MAP3K7 | mitogen-activated protein kinase kinase kinase 7 | 6582 | 0.018 | -0.0131 | No |
| 28 | PPP2R1B | protein phosphatase 2, regulatory subunit A, beta | 6781 | 0.017 | -0.0196 | No |
| 29 | ARPC3 | actin related protein 2/3 complex subunit 3 | 6792 | 0.017 | -0.0164 | No |
| 30 | ADCY2 | Transcript Identified by AceView, Entrez Gene ID(s) 108 | 6915 | 0.016 | -0.0191 | No |
| 31 | TNFRSF1A | tumor necrosis factor receptor superfamily, member 1A | 7027 | 0.016 | -0.0214 | No |
| 32 | CDK4 | cyclin-dependent kinase 4 | 7198 | 0.015 | -0.0270 | No |
| 33 | TBK1 | TANK-binding kinase 1 | 7221 | 0.015 | -0.0249 | No |
| 34 | PFN1 | profilin 1 | 7240 | 0.015 | -0.0227 | No |
| 35 | RAF1 | Raf-1 proto-oncogene, serine/threonine kinase | 7383 | 0.014 | -0.0271 | No |
| 36 | RALB | v-ral simian leukemia viral oncogene homolog B | 7970 | 0.010 | -0.0551 | No |
| 37 | NCK1 | NCK adaptor protein 1 | 8042 | 0.010 | -0.0566 | No |
| 38 | ACACA | acetyl-CoA carboxylase alpha | 8311 | 0.008 | -0.0687 | No |
| 39 | PIK3R3 | phosphoinositide-3-kinase, regulatory subunit 3 (gamma) | 8325 | 0.008 | -0.0676 | No |
| 40 | CFL1 | cofilin 1 (non-muscle) | 8519 | 0.007 | -0.0761 | No |
| 41 | CXCR4 | chemokine (C-X-C motif) receptor 4 | 8789 | 0.005 | -0.0888 | No |
| 42 | ITPR2 | inositol 1,4,5-trisphosphate receptor, type 2 | 8809 | 0.005 | -0.0887 | No |
| 43 | SMAD2 | SMAD family member 2 | 8933 | 0.004 | -0.0941 | No |
| 44 | CAMK4 | calcium/calmodulin-dependent protein kinase IV | 9650 | 0.001 | -0.1308 | No |
| 45 | NOD1 | nucleotide-binding oligomerization domain containing 1 | 9844 | -0.000 | -0.1406 | No |
| 46 | E2F1 | Memczak2013 ANTISENSE, coding, INTERNAL, intronic best transcript NM\_005225 | 9936 | -0.001 | -0.1451 | No |
| 47 | SLA | Src-like-adaptor | 10023 | -0.001 | -0.1493 | No |
| 48 | CAB39 | calcium binding protein 39 | 10079 | -0.002 | -0.1518 | No |
| 49 | RAC1 | ras-related C3 botulinum toxin substrate 1 (rho family, small GTP binding protein Rac1) | 10275 | -0.003 | -0.1612 | No |
| 50 | GNA14 | guanine nucleotide binding protein (G protein), alpha 14 | 10304 | -0.003 | -0.1620 | No |
| 51 | ACTR2 | ARP2 actin-related protein 2 homolog (yeast) | 10400 | -0.003 | -0.1661 | No |
| 52 | MYD88 | myeloid differentiation primary response 88 | 10440 | -0.004 | -0.1673 | No |
| 53 | MAP2K3 | mitogen-activated protein kinase kinase 3 | 10451 | -0.004 | -0.1670 | No |
| 54 | PTPN11 | protein tyrosine phosphatase, non-receptor type 11 | 10486 | -0.004 | -0.1679 | No |
| 55 | AKT1 | v-akt murine thymoma viral oncogene homolog 1 | 10680 | -0.005 | -0.1767 | No |
| 56 | THEM4 | thioesterase superfamily member 4 | 10693 | -0.005 | -0.1761 | No |
| 57 | YWHAB | tyrosine 3-monooxygenase/tryptophan 5-monooxygenase activation protein, beta | 10762 | -0.006 | -0.1784 | No |
| 58 | RPS6KA3 | ribosomal protein S6 kinase, 90kDa, polypeptide 3 | 10766 | -0.006 | -0.1773 | No |
| 59 | PRKCB | protein kinase C, beta | 10864 | -0.006 | -0.1809 | No |
| 60 | PPP1CA | protein phosphatase 1, catalytic subunit, alpha isozyme | 11025 | -0.007 | -0.1876 | No |
| 61 | SLC2A1 | solute carrier family 2 (facilitated glucose transporter), member 1 | 11178 | -0.008 | -0.1936 | No |
| 62 | MAPKAP1 | mitogen-activated protein kinase associated protein 1 | 11339 | -0.009 | -0.1999 | No |
| 63 | PTEN | phosphatase and tensin homolog | 11389 | -0.009 | -0.2005 | No |
| 64 | GRB2 | growth factor receptor bound protein 2 | 11880 | -0.012 | -0.2231 | No |
| 65 | ARHGDIA | Rho GDP dissociation inhibitor (GDI) alpha | 11975 | -0.013 | -0.2251 | No |
| 66 | FGF6 | fibroblast growth factor 6 | 12410 | -0.016 | -0.2441 | No |
| 67 | AP2M1 | adaptor-related protein complex 2, mu 1 subunit | 12582 | -0.017 | -0.2493 | No |
| 68 | PRKAG1 | protein kinase, AMP-activated, gamma 1 non-catalytic subunit | 12940 | -0.019 | -0.2636 | No |
| 69 | SQSTM1 | sequestosome 1 | 13032 | -0.019 | -0.2641 | No |
| 70 | CLTC | clathrin, heavy chain (Hc) | 13075 | -0.020 | -0.2619 | No |
| 71 | MAPK8 | mitogen-activated protein kinase 8 | 13286 | -0.021 | -0.2682 | No |
| 72 | IL4 | interleukin 4 | 13374 | -0.022 | -0.2679 | No |
| 73 | PIKFYVE | phosphoinositide kinase, FYVE finger containing | 13389 | -0.022 | -0.2639 | No |
| 74 | MAPK1 | mitogen-activated protein kinase 1 | 13405 | -0.022 | -0.2599 | No |
| 75 | UBE2D3 | Memczak2013 ANTISENSE, coding, INTERNAL, intronic best transcript NM\_181893 | 13866 | -0.025 | -0.2781 | No |
| 76 | GNGT1 | guanine nucleotide binding protein (G protein), gamma transducing activity polypeptide 1 | 14145 | -0.027 | -0.2865 | No |
| 77 | CALR | calreticulin | 14607 | -0.031 | -0.3034 | No |
| 78 | TSC2 | tuberous sclerosis 2 | 14696 | -0.032 | -0.3010 | No |
| 79 | GSK3B | glycogen synthase kinase 3 beta | 14754 | -0.032 | -0.2969 | No |
| 80 | ACTR3 | ARP3 actin-related protein 3 homolog (yeast) | 14975 | -0.034 | -0.3008 | No |
| 81 | HSP90B1 | heat shock protein 90kDa beta (Grp94), member 1 | 15295 | -0.037 | -0.3092 | No |
| 82 | NFKBIB | nuclear factor of kappa light polypeptide gene enhancer in B-cells inhibitor, beta | 15315 | -0.037 | -0.3021 | No |
| 83 | IRAK4 | interleukin 1 receptor associated kinase 4 | 15372 | -0.038 | -0.2968 | No |
| 84 | ARF1 | ADP-ribosylation factor 1 | 15527 | -0.039 | -0.2963 | No |
| 85 | RIPK1 | receptor (TNFRSF)-interacting serine-threonine kinase 1 | 16084 | -0.045 | -0.3152 | No |
| 86 | RIT1 | Ras-like without CAAX 1 | 16124 | -0.045 | -0.3074 | No |
| 87 | CDKN1B | cyclin-dependent kinase inhibitor 1B (p27, Kip1) | 16336 | -0.047 | -0.3079 | No |
| 88 | PAK4 | p21 protein (Cdc42/Rac)-activated kinase 4 | 16829 | -0.054 | -0.3215 | No |
| 89 | MKNK2 | MAP kinase interacting serine/threonine kinase 2 | 17133 | -0.059 | -0.3243 | No |
| 90 | PLA2G12A | Memczak2013 ANTISENSE, CDS, coding, INTERNAL best transcript NM\_030821 | 17611 | -0.068 | -0.3342 | Yes |
| 91 | DUSP3 | dual specificity phosphatase 3 | 17693 | -0.069 | -0.3233 | Yes |
| 92 | PRKAR2A | protein kinase, cAMP-dependent, regulatory, type II, alpha | 17801 | -0.072 | -0.3131 | Yes |
| 93 | MAPK10 | mitogen-activated protein kinase 10 | 18059 | -0.079 | -0.3093 | Yes |
| 94 | PDK1 | pyruvate dehydrogenase kinase, isozyme 1 | 18204 | -0.083 | -0.2988 | Yes |
| 95 | STAT2 | signal transducer and activator of transcription 2 | 18572 | -0.095 | -0.2970 | Yes |
| 96 | TRIB3 | tribbles pseudokinase 3 | 18870 | -0.115 | -0.2872 | Yes |
| 97 | MAP2K6 | mitogen-activated protein kinase kinase 6 | 18927 | -0.119 | -0.2641 | Yes |
| 98 | CAB39L | calcium binding protein 39-like | 19029 | -0.131 | -0.2409 | Yes |
| 99 | DAPP1 | dual adaptor of phosphotyrosine and 3-phosphoinositides | 19293 | -0.191 | -0.2130 | Yes |
| 100 | PITX2 | paired-like homeodomain 2 | 19326 | -0.200 | -0.1712 | Yes |
| 101 | PLCB1 | Memczak2013 ANTISENSE, coding, INTERNAL, intronic best transcript NM\_182734 | 19365 | -0.220 | -0.1255 | Yes |
| 102 | DDIT3 | DNA-damage-inducible transcript 3 | 19440 | -0.282 | -0.0680 | Yes |
| 103 | VAV3 | vav 3 guanine nucleotide exchange factor | 19476 | -0.333 | 0.0025 | Yes |
Table: GSEA details [plain text format]

  

Fig 2: HALLMARK\_PI3K\_AKT\_MTOR\_SIGNALING      
 Blue-Pink O' Gram in the Space of the Analyzed GeneSet

  

Fig 3: HALLMARK\_PI3K\_AKT\_MTOR\_SIGNALING: Random ES distribution      
 Gene set null distribution of ES for **HALLMARK\_PI3K\_AKT\_MTOR\_SIGNALING**

  
