## Supplementary material for "Somatic hypomethylation of pericentromeric SST1 repeats and tetraploidization in human colorectal cancer cells": GSEA results: HALLMARK_SPERMATOGENESIS.html

Details for gene set HALLMARK\_SPERMATOGENESIS[GSEA]

|  || Dataset | eset\_byprobe\_collapsed\_to\_symbols.Diploid\_vs\_Tetraploid.cls #Tetraploid\_versus\_Diploid.Diploid\_vs\_Tetraploid.cls #Tetraploid\_versus\_Diploid\_repos |
| Phenotype | Diploid\_vs\_Tetraploid.cls#Tetraploid\_versus\_Diploid\_repos |
| Upregulated in class | Tetraploid |
| GeneSet | HALLMARK\_SPERMATOGENESIS |
| Enrichment Score (ES) | 0.22505696 |
| Normalized Enrichment Score (NES) | 0.94126743 |
| Nominal p-value | 0.58870965 |
| FDR q-value | 0.812543 |
| FWER p-Value | 1.0 |
Table: GSEA Results Summary

  

Fig 1: Enrichment plot: HALLMARK\_SPERMATOGENESIS      
 Profile of the Running ES Score & Positions of GeneSet Members on the Rank Ordered List

  

| SYMBOL | TITLE | RANK IN GENE LIST | RANK METRIC SCORE | RUNNING ES | CORE ENRICHMENT || 1 | DMC1 | DNA meiotic recombinase 1 | 96 | 0.211 | 0.0333 | Yes |
| 2 | TLE4 | transducin-like enhancer of split 4 | 179 | 0.163 | 0.0585 | Yes |
| 3 | PSMG1 | proteasome (prosome, macropain) assembly chaperone 1 | 497 | 0.111 | 0.0622 | Yes |
| 4 | DMRT1 | doublesex and mab-3 related transcription factor 1 | 868 | 0.090 | 0.0594 | Yes |
| 5 | GPR182 | G protein-coupled receptor 182 | 998 | 0.084 | 0.0681 | Yes |
| 6 | HSPA4L | heat shock 70kDa protein 4-like | 1070 | 0.082 | 0.0792 | Yes |
| 7 | CFTR | cystic fibrosis transmembrane conductance regulator | 1189 | 0.079 | 0.0874 | Yes |
| 8 | TTK | TTK protein kinase | 1229 | 0.078 | 0.0995 | Yes |
| 9 | ZC2HC1C | zinc finger, C2HC-type containing 1C | 1230 | 0.078 | 0.1137 | Yes |
| 10 | ACE | angiotensin I converting enzyme | 1448 | 0.073 | 0.1157 | Yes |
| 11 | AGFG1 | ArfGAP with FG repeats 1 | 1550 | 0.070 | 0.1232 | Yes |
| 12 | SHE | Src homology 2 domain containing E | 1566 | 0.070 | 0.1351 | Yes |
| 13 | SNAP91 | synaptosome associated protein 91kDa | 1590 | 0.069 | 0.1464 | Yes |
| 14 | NPY5R | neuropeptide Y receptor Y5 | 1610 | 0.069 | 0.1579 | Yes |
| 15 | PARP2 | poly(ADP-ribose) polymerase 2 | 1614 | 0.068 | 0.1701 | Yes |
| 16 | PGS1 | phosphatidylglycerophosphate synthase 1 | 1846 | 0.064 | 0.1698 | Yes |
| 17 | HSPA2 | heat shock 70kDa protein 2 | 1901 | 0.063 | 0.1785 | Yes |
| 18 | SLC12A2 | solute carrier family 12 (sodium/potassium/chloride transporter), member 2 | 2023 | 0.061 | 0.1833 | Yes |
| 19 | ACTL7B | actin-like 7B | 2113 | 0.060 | 0.1896 | Yes |
| 20 | DBF4 | DBF4 zinc finger | 2187 | 0.058 | 0.1964 | Yes |
| 21 | IL12RB2 | interleukin 12 receptor, beta 2 | 2268 | 0.057 | 0.2027 | Yes |
| 22 | MAP7 | microtubule-associated protein 7 | 2428 | 0.055 | 0.2044 | Yes |
| 23 | TUBA3C | tubulin, alpha 3c | 2438 | 0.055 | 0.2138 | Yes |
| 24 | DPEP3 | dipeptidase 3 | 2717 | 0.051 | 0.2087 | Yes |
| 25 | THEG | theg spermatid protein | 2727 | 0.051 | 0.2175 | Yes |
| 26 | HBZ | hemoglobin, zeta | 3017 | 0.047 | 0.2111 | Yes |
| 27 | SCG3 | secretogranin III | 3098 | 0.047 | 0.2155 | Yes |
| 28 | GSTM3 | glutathione S-transferase mu 3 (brain) | 3265 | 0.045 | 0.2150 | Yes |
| 29 | DDX25 | DEAD (Asp-Glu-Ala-Asp) box helicase 25 | 3277 | 0.045 | 0.2226 | Yes |
| 30 | CLVS1 | clavesin 1 | 3449 | 0.043 | 0.2216 | Yes |
| 31 | DCC | DCC netrin 1 receptor | 3629 | 0.041 | 0.2199 | Yes |
| 32 | OAZ3 | ornithine decarboxylase antizyme 3 | 3673 | 0.041 | 0.2251 | Yes |
| 33 | ACRBP | acrosin binding protein | 4402 | 0.034 | 0.1937 | No |
| 34 | MLLT10 | myeloid/lymphoid or mixed-lineage leukemia; translocated to, 10 | 4594 | 0.032 | 0.1897 | No |
| 35 | ELOVL3 | ELOVL fatty acid elongase 3 | 4675 | 0.032 | 0.1913 | No |
| 36 | CDK1 | cyclin-dependent kinase 1 | 4823 | 0.031 | 0.1893 | No |
| 37 | PHKG2 | phosphorylase kinase, gamma 2 (testis) | 4880 | 0.030 | 0.1919 | No |
| 38 | AURKA | aurora kinase A | 5067 | 0.029 | 0.1875 | No |
| 39 | POMC | proopiomelanocortin | 5172 | 0.028 | 0.1872 | No |
| 40 | PAPOLB | poly(A) polymerase beta | 5218 | 0.028 | 0.1899 | No |
| 41 | TOPBP1 | topoisomerase (DNA) II binding protein 1 | 5259 | 0.027 | 0.1928 | No |
| 42 | MLF1 | myeloid leukemia factor 1 | 5320 | 0.027 | 0.1946 | No |
| 43 | HOXB1 | homeobox B1 | 5406 | 0.026 | 0.1949 | No |
| 44 | GMCL1 | germ cell-less, spermatogenesis associated 1 | 5621 | 0.025 | 0.1883 | No |
| 45 | PACRG | PARK2 co-regulated | 5665 | 0.024 | 0.1905 | No |
| 46 | HTR5A | 5-hydroxytryptamine (serotonin) receptor 5A, G protein-coupled | 5770 | 0.024 | 0.1895 | No |
| 47 | VDAC3 | voltage-dependent anion channel 3 | 5771 | 0.024 | 0.1937 | No |
| 48 | RFC4 | replication factor C subunit 4 | 5802 | 0.023 | 0.1964 | No |
| 49 | PHF7 | PHD finger protein 7 | 5823 | 0.023 | 0.1996 | No |
| 50 | BUB1 | BUB1 mitotic checkpoint serine/threonine kinase | 5988 | 0.022 | 0.1951 | No |
| 51 | EZH2 | enhancer of zeste 2 polycomb repressive complex 2 subunit | 5989 | 0.022 | 0.1991 | No |
| 52 | HSPA1L | heat shock 70kDa protein 1-like | 6184 | 0.021 | 0.1929 | No |
| 53 | TULP2 | tubby like protein 2 | 6384 | 0.020 | 0.1862 | No |
| 54 | ODF1 | outer dense fiber of sperm tails 1 | 6441 | 0.019 | 0.1869 | No |
| 55 | ZNRF4 | zinc and ring finger 4 | 6545 | 0.019 | 0.1849 | No |
| 56 | NCAPH | non-SMC condensin I complex subunit H | 6995 | 0.016 | 0.1646 | No |
| 57 | TNP2 | transition protein 2 (during histone to protamine replacement) | 7100 | 0.015 | 0.1620 | No |
| 58 | COIL | coilin | 7186 | 0.015 | 0.1604 | No |
| 59 | NEFH | neurofilament, heavy polypeptide | 7329 | 0.014 | 0.1556 | No |
| 60 | PGK2 | phosphoglycerate kinase 2 | 7365 | 0.014 | 0.1563 | No |
| 61 | GAPDHS | glyceraldehyde-3-phosphate dehydrogenase, spermatogenic | 7644 | 0.012 | 0.1441 | No |
| 62 | DNAJB8 | DnaJ (Hsp40) homolog, subfamily B, member 8 | 7689 | 0.012 | 0.1440 | No |
| 63 | PRM2 | protamine 2 | 7954 | 0.010 | 0.1322 | No |
| 64 | CCNB2 | cyclin B2 | 8020 | 0.010 | 0.1306 | No |
| 65 | NPHP1 | nephronophthisis 1 (juvenile) | 8117 | 0.009 | 0.1273 | No |
| 66 | CRISP2 | cysteine-rich secretory protein 2 | 8207 | 0.009 | 0.1243 | No |
| 67 | LDHC | lactate dehydrogenase C | 8359 | 0.008 | 0.1179 | No |
| 68 | TALDO1 | transaldolase 1 | 8396 | 0.007 | 0.1174 | No |
| 69 | PCSK4 | proprotein convertase subtilisin/kexin type 4 | 8729 | 0.006 | 0.1013 | No |
| 70 | IP6K1 | inositol hexakisphosphate kinase 1 | 8801 | 0.005 | 0.0986 | No |
| 71 | MTNR1A | melatonin receptor 1A | 8961 | 0.004 | 0.0911 | No |
| 72 | ALOX15 | arachidonate 15-lipoxygenase | 9076 | 0.004 | 0.0859 | No |
| 73 | PCSK1N | proprotein convertase subtilisin/kexin type 1 inhibitor | 9104 | 0.003 | 0.0851 | No |
| 74 | ZPBP | zona pellucida binding protein | 9314 | 0.002 | 0.0748 | No |
| 75 | CDKN3 | cyclin-dependent kinase inhibitor 3 | 9369 | 0.002 | 0.0724 | No |
| 76 | CAMK4 | calcium/calmodulin-dependent protein kinase IV | 9650 | 0.001 | 0.0581 | No |
| 77 | PIAS2 | protein inhibitor of activated STAT 2 | 9854 | -0.000 | 0.0477 | No |
| 78 | CHRM4 | cholinergic receptor, muscarinic 4 | 9969 | -0.001 | 0.0420 | No |
| 79 | PEBP1 | phosphatidylethanolamine binding protein 1 | 9974 | -0.001 | 0.0420 | No |
| 80 | SCG5 | secretogranin V | 10169 | -0.002 | 0.0323 | No |
| 81 | DDX4 | DEAD (Asp-Glu-Ala-Asp) box polypeptide 4 | 10279 | -0.003 | 0.0272 | No |
| 82 | ACRV1 | acrosomal vesicle protein 1 | 10323 | -0.003 | 0.0256 | No |
| 83 | TNNI3 | troponin I type 3 (cardiac) | 10545 | -0.004 | 0.0150 | No |
| 84 | GFI1 | growth factor independent 1 transcription repressor | 10648 | -0.005 | 0.0106 | No |
| 85 | CLGN | calmegin | 11000 | -0.007 | -0.0062 | No |
| 86 | ADAD1 | adenosine deaminase domain containing 1 | 11012 | -0.007 | -0.0055 | No |
| 87 | ADCYAP1 | adenylate cyclase activating polypeptide 1 (pituitary) | 11129 | -0.008 | -0.0100 | No |
| 88 | KIF2C | kinesin family member 2C | 11587 | -0.010 | -0.0317 | No |
| 89 | NEK2 | NIMA-related kinase 2 | 11616 | -0.011 | -0.0313 | No |
| 90 | CLPB | ClpB homolog, mitochondrial AAA ATPase chaperonin | 11672 | -0.011 | -0.0321 | No |
| 91 | ZC3H14 | zinc finger CCCH-type containing 14 | 11756 | -0.011 | -0.0344 | No |
| 92 | SYCP1 | synaptonemal complex protein 1 | 11766 | -0.011 | -0.0327 | No |
| 93 | TSSK2 | testis-specific serine kinase 2 | 11920 | -0.012 | -0.0384 | No |
| 94 | JAM3 | junctional adhesion molecule 3 | 12131 | -0.014 | -0.0468 | No |
| 95 | MTOR | mechanistic target of rapamycin (serine/threonine kinase) | 12446 | -0.016 | -0.0601 | No |
| 96 | IDE | insulin-degrading enzyme | 12755 | -0.018 | -0.0728 | No |
| 97 | STRBP | spermatid perinuclear RNA binding protein | 12795 | -0.018 | -0.0715 | No |
| 98 | NOS1 | nitric oxide synthase 1 (neuronal) | 13175 | -0.020 | -0.0874 | No |
| 99 | GAD1 | glutamate decarboxylase 1 | 13246 | -0.021 | -0.0872 | No |
| 100 | YBX2 | Y box binding protein 2 | 13662 | -0.024 | -0.1043 | No |
| 101 | ADAM2 | ADAM metallopeptidase domain 2 | 13744 | -0.024 | -0.1041 | No |
| 102 | CCNA1 | cyclin A1 | 13813 | -0.025 | -0.1030 | No |
| 103 | MAST2 | microtubule associated serine/threonine kinase 2 | 13921 | -0.026 | -0.1039 | No |
| 104 | TNP1 | transition protein 1 (during histone to protamine replacement) | 14768 | -0.033 | -0.1416 | No |
| 105 | TSN | translin | 14905 | -0.034 | -0.1425 | No |
| 106 | PDHA2 | pyruvate dehydrogenase (lipoamide) alpha 2 | 15177 | -0.036 | -0.1500 | No |
| 107 | LPIN1 | Transcript Identified by AceView, Entrez Gene ID(s) 23175 | 15296 | -0.037 | -0.1494 | No |
| 108 | CSNK2A2 | casein kinase 2, alpha prime polypeptide | 15362 | -0.038 | -0.1459 | No |
| 109 | NAA11 | N(alpha)-acetyltransferase 11, NatA catalytic subunit | 15536 | -0.039 | -0.1478 | No |
| 110 | SIRT1 | sirtuin 1 | 15838 | -0.042 | -0.1557 | No |
| 111 | RAD17 | RAD17 checkpoint clamp loader component | 15985 | -0.044 | -0.1553 | No |
| 112 | GSG1 | germ cell associated 1 | 16068 | -0.045 | -0.1515 | No |
| 113 | STAM2 | signal transducing adaptor molecule (SH3 domain and ITAM motif) 2 | 16637 | -0.052 | -0.1714 | No |
| 114 | ART3 | ADP-ribosyltransferase 3 | 16849 | -0.055 | -0.1724 | No |
| 115 | SPATA6 | spermatogenesis associated 6 | 16932 | -0.056 | -0.1665 | No |
| 116 | TEKT2 | tektin 2 (testicular) | 17035 | -0.057 | -0.1613 | No |
| 117 | CNIH2 | cornichon family AMPA receptor auxiliary protein 2 | 17121 | -0.059 | -0.1551 | No |
| 118 | NF2 | neurofibromin 2 (merlin) | 17161 | -0.059 | -0.1463 | No |
| 119 | TCP11 | t-complex 11, testis-specific | 17184 | -0.060 | -0.1366 | No |
| 120 | CST8 | cystatin 8 (cystatin-related epididymal specific) | 17330 | -0.062 | -0.1328 | No |
| 121 | IL13RA2 | interleukin 13 receptor, alpha 2 | 17427 | -0.064 | -0.1262 | No |
| 122 | CCT6B | chaperonin containing TCP1, subunit 6B (zeta 2) | 17752 | -0.071 | -0.1301 | No |
| 123 | CHFR | checkpoint with forkhead and ring finger domains, E3 ubiquitin protein ligase | 17781 | -0.072 | -0.1185 | No |
| 124 | TKTL1 | transketolase-like 1 | 17783 | -0.072 | -0.1056 | No |
| 125 | PRKAR2A | protein kinase, cAMP-dependent, regulatory, type II, alpha | 17801 | -0.072 | -0.0934 | No |
| 126 | SLC2A5 | solute carrier family 2 (facilitated glucose/fructose transporter), member 5 | 17838 | -0.073 | -0.0820 | No |
| 127 | AKAP4 | A kinase (PRKA) anchor protein 4 | 18147 | -0.081 | -0.0832 | No |
| 128 | IFT88 | intraflagellar transport 88 | 18324 | -0.086 | -0.0767 | No |
| 129 | BRAF | B-Raf proto-oncogene, serine/threonine kinase | 18407 | -0.089 | -0.0648 | No |
| 130 | MEP1B | meprin A, beta | 19195 | -0.160 | -0.0763 | No |
| 131 | RPL39L | ribosomal protein L39-like | 19202 | -0.163 | -0.0471 | No |
| 132 | ARL4A | ADP-ribosylation factor like GTPase 4A | 19243 | -0.172 | -0.0180 | No |
| 133 | GRM8 | glutamate receptor, metabotropic 8 | 19266 | -0.179 | 0.0133 | No |
Table: GSEA details [plain text format]

  

Fig 2: HALLMARK\_SPERMATOGENESIS      
 Blue-Pink O' Gram in the Space of the Analyzed GeneSet

  

Fig 3: HALLMARK\_SPERMATOGENESIS: Random ES distribution      
 Gene set null distribution of ES for **HALLMARK\_SPERMATOGENESIS**

  
