## Supplementary material for "Somatic hypomethylation of pericentromeric SST1 repeats and tetraploidization in human colorectal cancer cells": GSEA results: HALLMARK_TGF_BETA_SIGNALING.html

Details for gene set HALLMARK\_TGF\_BETA\_SIGNALING[GSEA]

|  || Dataset | eset\_byprobe\_collapsed\_to\_symbols.Diploid\_vs\_Tetraploid.cls #Tetraploid\_versus\_Diploid.Diploid\_vs\_Tetraploid.cls #Tetraploid\_versus\_Diploid\_repos |
| Phenotype | Diploid\_vs\_Tetraploid.cls#Tetraploid\_versus\_Diploid\_repos |
| Upregulated in class | Tetraploid |
| GeneSet | HALLMARK\_TGF\_BETA\_SIGNALING |
| Enrichment Score (ES) | 0.31173033 |
| Normalized Enrichment Score (NES) | 1.1286685 |
| Nominal p-value | 0.23893805 |
| FDR q-value | 0.33881408 |
| FWER p-Value | 0.981 |
Table: GSEA Results Summary

  

Fig 1: Enrichment plot: HALLMARK\_TGF\_BETA\_SIGNALING      
 Profile of the Running ES Score & Positions of GeneSet Members on the Rank Ordered List

  

| SYMBOL | TITLE | RANK IN GENE LIST | RANK METRIC SCORE | RUNNING ES | CORE ENRICHMENT || 1 | SMAD6 | SMAD family member 6 | 132 | 0.186 | 0.0893 | Yes |
| 2 | THBS1 | thrombospondin 1 | 192 | 0.158 | 0.1680 | Yes |
| 3 | SMAD7 | SMAD family member 7 | 372 | 0.123 | 0.2224 | Yes |
| 4 | ID3 | inhibitor of DNA binding 3, dominant negative helix-loop-helix protein | 1001 | 0.084 | 0.2338 | Yes |
| 5 | ID1 | inhibitor of DNA binding 1, dominant negative helix-loop-helix protein | 1068 | 0.082 | 0.2729 | Yes |
| 6 | ID2 | inhibitor of DNA binding 2, dominant negative helix-loop-helix protein | 1150 | 0.080 | 0.3100 | Yes |
| 7 | LEFTY2 | left-right determination factor 2 | 1775 | 0.065 | 0.3117 | Yes |
| 8 | TGIF1 | TGFB-induced factor homeobox 1 | 3674 | 0.041 | 0.2354 | No |
| 9 | SMURF2 | SMAD specific E3 ubiquitin protein ligase 2 | 4014 | 0.037 | 0.2374 | No |
| 10 | SKI | SKI proto-oncogene | 5564 | 0.025 | 0.1708 | No |
| 11 | RAB31 | Transcript Identified by AceView, Entrez Gene ID(s) 11031 | 6174 | 0.021 | 0.1503 | No |
| 12 | PPM1A | protein phosphatase, Mg2+/Mn2+ dependent, 1A | 6498 | 0.019 | 0.1436 | No |
| 13 | MAP3K7 | mitogen-activated protein kinase kinase kinase 7 | 6582 | 0.018 | 0.1489 | No |
| 14 | ARID4B | AT rich interactive domain 4B (RBP1-like) | 6960 | 0.016 | 0.1378 | No |
| 15 | BMPR2 | bone morphogenetic protein receptor type II | 6988 | 0.016 | 0.1447 | No |
| 16 | CDH1 | cadherin 1, type 1 | 7180 | 0.015 | 0.1426 | No |
| 17 | SMAD3 | SMAD family member 3 | 7263 | 0.014 | 0.1458 | No |
| 18 | SPTBN1 | spectrin, beta, non-erythrocytic 1 | 7325 | 0.014 | 0.1500 | No |
| 19 | IFNGR2 | interferon gamma receptor 2 (interferon gamma transducer 1) | 9041 | 0.004 | 0.0638 | No |
| 20 | NOG | noggin | 9209 | 0.003 | 0.0568 | No |
| 21 | HDAC1 | histone deacetylase 1 | 9475 | 0.002 | 0.0440 | No |
| 22 | PMEPA1 | prostate transmembrane protein, androgen induced 1 | 9736 | 0.000 | 0.0307 | No |
| 23 | FKBP1A | FK506 binding protein 1A | 9775 | 0.000 | 0.0288 | No |
| 24 | RHOA | ras homolog family member A | 10006 | -0.001 | 0.0176 | No |
| 25 | TRIM33 | tripartite motif containing 33 | 10432 | -0.004 | -0.0023 | No |
| 26 | CDK9 | cyclin-dependent kinase 9 | 10556 | -0.004 | -0.0063 | No |
| 27 | SMURF1 | SMAD specific E3 ubiquitin protein ligase 1 | 10807 | -0.006 | -0.0160 | No |
| 28 | JUNB | jun B proto-oncogene | 10813 | -0.006 | -0.0131 | No |
| 29 | SLC20A1 | solute carrier family 20 (phosphate transporter), member 1 | 10841 | -0.006 | -0.0113 | No |
| 30 | PPP1CA | protein phosphatase 1, catalytic subunit, alpha isozyme | 11025 | -0.007 | -0.0170 | No |
| 31 | CDKN1C | cyclin-dependent kinase inhibitor 1C (p57, Kip2) | 11246 | -0.008 | -0.0239 | No |
| 32 | SMAD1 | SMAD family member 1 | 11250 | -0.008 | -0.0197 | No |
| 33 | WWTR1 | WW domain containing transcription regulator 1 | 11345 | -0.009 | -0.0198 | No |
| 34 | CTNNB1 | catenin (cadherin-associated protein), beta 1 | 11452 | -0.010 | -0.0203 | No |
| 35 | TGFBR1 | transforming growth factor, beta receptor 1 | 12290 | -0.015 | -0.0556 | No |
| 36 | NCOR2 | nuclear receptor corepressor 2 | 12339 | -0.015 | -0.0503 | No |
| 37 | TGFB1 | transforming growth factor beta 1 | 12464 | -0.016 | -0.0484 | No |
| 38 | SERPINE1 | serpin peptidase inhibitor, clade E (nexin, plasminogen activator inhibitor type 1), member 1 | 12974 | -0.019 | -0.0647 | No |
| 39 | SKIL | SKI-like proto-oncogene | 13624 | -0.024 | -0.0859 | No |
| 40 | ENG | endoglin | 13817 | -0.025 | -0.0828 | No |
| 41 | UBE2D3 | Memczak2013 ANTISENSE, coding, INTERNAL, intronic best transcript NM\_181893 | 13866 | -0.025 | -0.0721 | No |
| 42 | BMPR1A | bone morphogenetic protein receptor type IA | 14258 | -0.028 | -0.0775 | No |
| 43 | HIPK2 | homeodomain interacting protein kinase 2 | 15466 | -0.038 | -0.1196 | No |
| 44 | FNTA | farnesyltransferase, CAAX box, alpha | 15473 | -0.039 | -0.0999 | No |
| 45 | TJP1 | tight junction protein 1 | 15540 | -0.039 | -0.0830 | No |
| 46 | XIAP | X-linked inhibitor of apoptosis, E3 ubiquitin protein ligase | 15992 | -0.044 | -0.0836 | No |
| 47 | APC | adenomatous polyposis coli | 16107 | -0.045 | -0.0662 | No |
| 48 | BMP2 | bone morphogenetic protein 2 | 16350 | -0.048 | -0.0539 | No |
| 49 | LTBP2 | latent transforming growth factor beta binding protein 2 | 16467 | -0.049 | -0.0343 | No |
| 50 | BCAR3 | breast cancer anti-estrogen resistance 3 | 16807 | -0.054 | -0.0237 | No |
| 51 | ACVR1 | activin A receptor type I | 16819 | -0.054 | 0.0037 | No |
| 52 | KLF10 | Kruppel-like factor 10 | 17645 | -0.068 | -0.0033 | No |
| 53 | FURIN | furin (paired basic amino acid cleaving enzyme) | 18023 | -0.077 | 0.0174 | No |
| 54 | PPP1R15A | protein phosphatase 1, regulatory subunit 15A | 18867 | -0.115 | 0.0337 | No |
Table: GSEA details [plain text format]

  

Fig 2: HALLMARK\_TGF\_BETA\_SIGNALING      
 Blue-Pink O' Gram in the Space of the Analyzed GeneSet

  

Fig 3: HALLMARK\_TGF\_BETA\_SIGNALING: Random ES distribution      
 Gene set null distribution of ES for **HALLMARK\_TGF\_BETA\_SIGNALING**

  
