## Supplementary material for "Somatic hypomethylation of pericentromeric SST1 repeats and tetraploidization in human colorectal cancer cells": GSEA results: HALLMARK_TNFA_SIGNALING_VIA_NFKB.html

Details for gene set HALLMARK\_TNFA\_SIGNALING\_VIA\_NFKB[GSEA]

|  || Dataset | eset\_byprobe\_collapsed\_to\_symbols.Diploid\_vs\_Tetraploid.cls #Tetraploid\_versus\_Diploid.Diploid\_vs\_Tetraploid.cls #Tetraploid\_versus\_Diploid\_repos |
| Phenotype | Diploid\_vs\_Tetraploid.cls#Tetraploid\_versus\_Diploid\_repos |
| Upregulated in class | Tetraploid |
| GeneSet | HALLMARK\_TNFA\_SIGNALING\_VIA\_NFKB |
| Enrichment Score (ES) | 0.2608101 |
| Normalized Enrichment Score (NES) | 1.1686003 |
| Nominal p-value | 0.13725491 |
| FDR q-value | 0.27180895 |
| FWER p-Value | 0.952 |
Table: GSEA Results Summary

  

Fig 1: Enrichment plot: HALLMARK\_TNFA\_SIGNALING\_VIA\_NFKB      
 Profile of the Running ES Score & Positions of GeneSet Members on the Rank Ordered List

  

| SYMBOL | TITLE | RANK IN GENE LIST | RANK METRIC SCORE | RUNNING ES | CORE ENRICHMENT || 1 | CCL20 | chemokine (C-C motif) ligand 20 | 12 | 0.406 | 0.0340 | Yes |
| 2 | PTGER4 | prostaglandin E receptor 4 (subtype EP4) | 59 | 0.272 | 0.0549 | Yes |
| 3 | PLK2 | polo-like kinase 2 | 109 | 0.202 | 0.0696 | Yes |
| 4 | F3 | coagulation factor III (thromboplastin, tissue factor) | 119 | 0.191 | 0.0854 | Yes |
| 5 | DRAM1 | DNA-damage regulated autophagy modulator 1 | 191 | 0.158 | 0.0952 | Yes |
| 6 | TNFSF9 | tumor necrosis factor (ligand) superfamily, member 9 | 223 | 0.150 | 0.1064 | Yes |
| 7 | IL1A | interleukin 1 alpha | 237 | 0.146 | 0.1182 | Yes |
| 8 | BCL6 | B-cell CLL/lymphoma 6 | 241 | 0.145 | 0.1304 | Yes |
| 9 | BTG2 | BTG family, member 2 | 281 | 0.136 | 0.1400 | Yes |
| 10 | ZBTB10 | zinc finger and BTB domain containing 10 | 326 | 0.129 | 0.1488 | Yes |
| 11 | EDN1 | endothelin 1 | 371 | 0.123 | 0.1570 | Yes |
| 12 | FOSL1 | FOS-like antigen 1 | 393 | 0.120 | 0.1661 | Yes |
| 13 | LIF | leukemia inhibitory factor | 423 | 0.117 | 0.1746 | Yes |
| 14 | ACKR3 | atypical chemokine receptor 3 | 455 | 0.114 | 0.1827 | Yes |
| 15 | BTG3 | BTG family, member 3 | 465 | 0.113 | 0.1919 | Yes |
| 16 | NR4A2 | nuclear receptor subfamily 4, group A, member 2 | 499 | 0.110 | 0.1996 | Yes |
| 17 | CD44 | CD44 molecule (Indian blood group) | 577 | 0.104 | 0.2045 | Yes |
| 18 | TRIP10 | thyroid hormone receptor interactor 10 | 585 | 0.104 | 0.2131 | Yes |
| 19 | IER5 | immediate early response 5 | 657 | 0.100 | 0.2179 | Yes |
| 20 | CXCL1 | chemokine (C-X-C motif) ligand 1 (melanoma growth stimulating activity, alpha) | 672 | 0.098 | 0.2256 | Yes |
| 21 | CDKN1A | cyclin-dependent kinase inhibitor 1A (p21, Cip1) | 690 | 0.097 | 0.2330 | Yes |
| 22 | PHLDA1 | pleckstrin homology-like domain, family A, member 1 | 740 | 0.095 | 0.2386 | Yes |
| 23 | JAG1 | jagged 1 | 962 | 0.085 | 0.2345 | Yes |
| 24 | CSF1 | colony stimulating factor 1 (macrophage) | 1040 | 0.083 | 0.2376 | Yes |
| 25 | IRS2 | insulin receptor substrate 2 | 1045 | 0.083 | 0.2444 | Yes |
| 26 | CXCL3 | chemokine (C-X-C motif) ligand 3 | 1048 | 0.083 | 0.2514 | Yes |
| 27 | ID2 | inhibitor of DNA binding 2, dominant negative helix-loop-helix protein | 1150 | 0.080 | 0.2529 | Yes |
| 28 | TNFAIP2 | tumor necrosis factor, alpha-induced protein 2 | 1223 | 0.078 | 0.2559 | Yes |
| 29 | PLAUR | plasminogen activator, urokinase receptor | 1355 | 0.075 | 0.2555 | Yes |
| 30 | BIRC3 | Memczak2013 ANTISENSE, CDS, coding, INTERNAL best transcript NM\_182962 | 1400 | 0.074 | 0.2595 | Yes |
| 31 | SPHK1 | sphingosine kinase 1 | 1575 | 0.070 | 0.2565 | Yes |
| 32 | ICAM1 | intercellular adhesion molecule 1 | 1606 | 0.069 | 0.2608 | Yes |
| 33 | DUSP2 | dual specificity phosphatase 2 | 2355 | 0.056 | 0.2269 | No |
| 34 | ETS2 | v-ets avian erythroblastosis virus E26 oncogene homolog 2 | 2456 | 0.054 | 0.2264 | No |
| 35 | FJX1 | four jointed box 1 | 2459 | 0.054 | 0.2309 | No |
| 36 | IL15RA | interleukin 15 receptor, alpha | 2659 | 0.051 | 0.2250 | No |
| 37 | PTX3 | pentraxin 3, long | 2893 | 0.049 | 0.2171 | No |
| 38 | MAFF | v-maf avian musculoaponeurotic fibrosarcoma oncogene homolog F | 3075 | 0.047 | 0.2117 | No |
| 39 | CXCL2 | chemokine (C-X-C motif) ligand 2 | 3085 | 0.047 | 0.2153 | No |
| 40 | MYC | v-myc avian myelocytomatosis viral oncogene homolog | 3194 | 0.046 | 0.2136 | No |
| 41 | TNF | tumor necrosis factor | 3210 | 0.045 | 0.2167 | No |
| 42 | RELB | v-rel avian reticuloendotheliosis viral oncogene homolog B | 3521 | 0.042 | 0.2043 | No |
| 43 | IL7R | interleukin 7 receptor | 3650 | 0.041 | 0.2011 | No |
| 44 | TGIF1 | TGFB-induced factor homeobox 1 | 3674 | 0.041 | 0.2034 | No |
| 45 | PHLDA2 | pleckstrin homology-like domain, family A, member 2 | 3707 | 0.041 | 0.2053 | No |
| 46 | B4GALT5 | UDP-Gal:betaGlcNAc beta 1,4- galactosyltransferase, polypeptide 5 | 3744 | 0.040 | 0.2068 | No |
| 47 | NFKB1 | nuclear factor of kappa light polypeptide gene enhancer in B-cells 1 | 3856 | 0.039 | 0.2044 | No |
| 48 | SLC2A6 | solute carrier family 2 (facilitated glucose transporter), member 6 | 3972 | 0.038 | 0.2017 | No |
| 49 | GEM | GTP binding protein overexpressed in skeletal muscle | 4213 | 0.035 | 0.1923 | No |
| 50 | SNN | stannin | 4231 | 0.035 | 0.1944 | No |
| 51 | DENND5A | DENN/MADD domain containing 5A | 4458 | 0.033 | 0.1856 | No |
| 52 | PFKFB3 | 6-phosphofructo-2-kinase/fructose-2,6-biphosphatase 3 | 4583 | 0.032 | 0.1820 | No |
| 53 | CCND1 | cyclin D1 | 4585 | 0.032 | 0.1847 | No |
| 54 | NFAT5 | nuclear factor of activated T-cells 5, tonicity-responsive | 4618 | 0.032 | 0.1858 | No |
| 55 | TUBB2A | tubulin, beta 2A class IIa | 4954 | 0.030 | 0.1710 | No |
| 56 | TRAF1 | TNF receptor-associated factor 1 | 5030 | 0.029 | 0.1696 | No |
| 57 | NFKBIE | nuclear factor of kappa light polypeptide gene enhancer in B-cells inhibitor, epsilon | 5273 | 0.027 | 0.1594 | No |
| 58 | PLAU | plasminogen activator, urokinase | 5343 | 0.027 | 0.1581 | No |
| 59 | HES1 | hes family bHLH transcription factor 1 | 5409 | 0.026 | 0.1570 | No |
| 60 | CCL2 | chemokine (C-C motif) ligand 2 | 5479 | 0.026 | 0.1556 | No |
| 61 | GADD45B | growth arrest and DNA-damage-inducible, beta | 5568 | 0.025 | 0.1532 | No |
| 62 | SERPINB2 | serpin peptidase inhibitor, clade B (ovalbumin), member 2 | 6047 | 0.022 | 0.1303 | No |
| 63 | TAP1 | transporter 1, ATP-binding cassette, sub-family B (MDR/TAP) | 6169 | 0.021 | 0.1258 | No |
| 64 | CXCL11 | chemokine (C-X-C motif) ligand 11 | 6249 | 0.021 | 0.1235 | No |
| 65 | SGK1 | serum/glucocorticoid regulated kinase 1 | 6290 | 0.020 | 0.1231 | No |
| 66 | NFE2L2 | nuclear factor, erythroid 2-like 2 | 6594 | 0.018 | 0.1090 | No |
| 67 | PDLIM5 | PDZ and LIM domain 5 | 6773 | 0.017 | 0.1013 | No |
| 68 | TNIP1 | TNFAIP3 interacting protein 1 | 6826 | 0.017 | 0.1000 | No |
| 69 | SMAD3 | SMAD family member 3 | 7263 | 0.014 | 0.0787 | No |
| 70 | GADD45A | growth arrest and DNA-damage-inducible, alpha | 7290 | 0.014 | 0.0786 | No |
| 71 | TNIP2 | TNFAIP3 interacting protein 2 | 7420 | 0.013 | 0.0731 | No |
| 72 | MCL1 | myeloid cell leukemia 1 | 7519 | 0.013 | 0.0691 | No |
| 73 | FOSL2 | FOS-like antigen 2 | 7696 | 0.012 | 0.0610 | No |
| 74 | BTG1 | B-cell translocation gene 1, anti-proliferative | 7702 | 0.012 | 0.0617 | No |
| 75 | EHD1 | EH domain containing 1 | 7793 | 0.011 | 0.0580 | No |
| 76 | CSF2 | colony stimulating factor 2 (granulocyte-macrophage) | 7825 | 0.011 | 0.0573 | No |
| 77 | ZC3H12A | zinc finger CCCH-type containing 12A | 7845 | 0.011 | 0.0573 | No |
| 78 | IL18 | interleukin 18 | 7966 | 0.010 | 0.0519 | No |
| 79 | CXCL10 | chemokine (C-X-C motif) ligand 10 | 8286 | 0.008 | 0.0361 | No |
| 80 | OLR1 | oxidized low density lipoprotein (lectin-like) receptor 1 | 8290 | 0.008 | 0.0366 | No |
| 81 | TNFAIP8 | tumor necrosis factor, alpha-induced protein 8 | 8321 | 0.008 | 0.0358 | No |
| 82 | NFKB2 | nuclear factor of kappa light polypeptide gene enhancer in B-cells 2 (p49/p100) | 8341 | 0.008 | 0.0354 | No |
| 83 | SOD2 | superoxide dismutase 2, mitochondrial | 8450 | 0.007 | 0.0305 | No |
| 84 | CCL5 | chemokine (C-C motif) ligand 5 | 8596 | 0.006 | 0.0235 | No |
| 85 | PTPRE | protein tyrosine phosphatase, receptor type, E | 8781 | 0.005 | 0.0144 | No |
| 86 | PER1 | period circadian clock 1 | 8800 | 0.005 | 0.0139 | No |
| 87 | CCNL1 | cyclin L1 | 9034 | 0.004 | 0.0022 | No |
| 88 | IFNGR2 | interferon gamma receptor 2 (interferon gamma transducer 1) | 9041 | 0.004 | 0.0022 | No |
| 89 | CFLAR | CASP8 and FADD like apoptosis regulator | 9278 | 0.003 | -0.0098 | No |
| 90 | LDLR | low density lipoprotein receptor | 9447 | 0.002 | -0.0183 | No |
| 91 | KYNU | kynureninase | 9602 | 0.001 | -0.0262 | No |
| 92 | RNF19B | Memczak2013 ANTISENSE, CDS, coding, INTERNAL best transcript NM\_153341 | 9730 | 0.000 | -0.0327 | No |
| 93 | PMEPA1 | prostate transmembrane protein, androgen induced 1 | 9736 | 0.000 | -0.0330 | No |
| 94 | NFKBIA | nuclear factor of kappa light polypeptide gene enhancer in B-cells inhibitor, alpha | 9745 | 0.000 | -0.0334 | No |
| 95 | F2RL1 | coagulation factor II (thrombin) receptor-like 1 | 9798 | -0.000 | -0.0361 | No |
| 96 | PLEK | pleckstrin | 10164 | -0.002 | -0.0548 | No |
| 97 | MAP2K3 | mitogen-activated protein kinase kinase 3 | 10451 | -0.004 | -0.0692 | No |
| 98 | SOCS3 | suppressor of cytokine signaling 3 | 10714 | -0.005 | -0.0823 | No |
| 99 | NINJ1 | ninjurin 1 | 10805 | -0.006 | -0.0865 | No |
| 100 | JUNB | jun B proto-oncogene | 10813 | -0.006 | -0.0863 | No |
| 101 | TIPARP | TCDD-inducible poly(ADP-ribose) polymerase | 11003 | -0.007 | -0.0955 | No |
| 102 | NR4A1 | nuclear receptor subfamily 4, group A, member 1 | 11095 | -0.008 | -0.0996 | No |
| 103 | IFIT2 | interferon-induced protein with tetratricopeptide repeats 2 | 11183 | -0.008 | -0.1034 | No |
| 104 | CD80 | CD80 molecule | 11264 | -0.009 | -0.1068 | No |
| 105 | IL6 | interleukin 6 | 11825 | -0.012 | -0.1347 | No |
| 106 | GFPT2 | glutamine-fructose-6-phosphate transaminase 2 | 11850 | -0.012 | -0.1350 | No |
| 107 | CXCL6 | chemokine (C-X-C motif) ligand 6 | 11900 | -0.012 | -0.1365 | No |
| 108 | RIPK2 | receptor-interacting serine-threonine kinase 2 | 12627 | -0.017 | -0.1726 | No |
| 109 | PANX1 | pannexin 1 | 12693 | -0.017 | -0.1745 | No |
| 110 | GCH1 | GTP cyclohydrolase 1 | 12793 | -0.018 | -0.1781 | No |
| 111 | FUT4 | fucosyltransferase 4 (alpha (1,3) fucosyltransferase, myeloid-specific) | 12820 | -0.018 | -0.1779 | No |
| 112 | STAT5A | signal transducer and activator of transcription 5A | 12939 | -0.019 | -0.1824 | No |
| 113 | KLF6 | Kruppel-like factor 6 | 12945 | -0.019 | -0.1810 | No |
| 114 | SERPINE1 | serpin peptidase inhibitor, clade E (nexin, plasminogen activator inhibitor type 1), member 1 | 12974 | -0.019 | -0.1808 | No |
| 115 | SQSTM1 | sequestosome 1 | 13032 | -0.019 | -0.1821 | No |
| 116 | DUSP4 | dual specificity phosphatase 4 | 13155 | -0.020 | -0.1867 | No |
| 117 | MSC | musculin | 13223 | -0.021 | -0.1884 | No |
| 118 | TNC | tenascin C | 13324 | -0.021 | -0.1918 | No |
| 119 | CD69 | CD69 molecule | 13361 | -0.022 | -0.1918 | No |
| 120 | TNFAIP3 | tumor necrosis factor, alpha-induced protein 3 | 13363 | -0.022 | -0.1900 | No |
| 121 | SIK1 | salt-inducible kinase 1 | 13968 | -0.026 | -0.2190 | No |
| 122 | TSC22D1 | TSC22 domain family, member 1 | 14022 | -0.026 | -0.2195 | No |
| 123 | IER3 | immediate early response 3 | 14076 | -0.027 | -0.2199 | No |
| 124 | EIF1 | eukaryotic translation initiation factor 1 | 14163 | -0.028 | -0.2220 | No |
| 125 | CEBPD | CCAAT/enhancer binding protein (C/EBP), delta | 14173 | -0.028 | -0.2201 | No |
| 126 | KDM6B | lysine (K)-specific demethylase 6B | 14199 | -0.028 | -0.2190 | No |
| 127 | CEBPB | CCAAT/enhancer binding protein (C/EBP), beta | 14355 | -0.029 | -0.2245 | No |
| 128 | BIRC2 | baculoviral IAP repeat containing 2 | 14552 | -0.031 | -0.2321 | No |
| 129 | HBEGF | heparin-binding EGF-like growth factor | 14753 | -0.032 | -0.2396 | No |
| 130 | EGR3 | early growth response 3 | 14802 | -0.033 | -0.2393 | No |
| 131 | DDX58 | DEAD (Asp-Glu-Ala-Asp) box polypeptide 58 | 14926 | -0.034 | -0.2428 | No |
| 132 | DUSP5 | dual specificity phosphatase 5 | 15010 | -0.035 | -0.2441 | No |
| 133 | TANK | TRAF family member-associated NFKB activator | 15072 | -0.035 | -0.2443 | No |
| 134 | INHBA | inhibin beta A | 15087 | -0.035 | -0.2420 | No |
| 135 | MAP3K8 | mitogen-activated protein kinase kinase kinase 8 | 15311 | -0.037 | -0.2504 | No |
| 136 | PNRC1 | proline-rich nuclear receptor coactivator 1 | 15332 | -0.037 | -0.2483 | No |
| 137 | DNAJB4 | DnaJ (Hsp40) homolog, subfamily B, member 4 | 15505 | -0.039 | -0.2538 | No |
| 138 | SPSB1 | splA/ryanodine receptor domain and SOCS box containing 1 | 15528 | -0.039 | -0.2516 | No |
| 139 | RELA | v-rel avian reticuloendotheliosis viral oncogene homolog A | 15578 | -0.039 | -0.2508 | No |
| 140 | REL | v-rel avian reticuloendotheliosis viral oncogene homolog | 15642 | -0.040 | -0.2506 | No |
| 141 | TRIB1 | tribbles pseudokinase 1 | 15669 | -0.041 | -0.2485 | No |
| 142 | CD83 | CD83 molecule | 15767 | -0.041 | -0.2500 | No |
| 143 | EGR2 | early growth response 2 | 15770 | -0.041 | -0.2466 | No |
| 144 | CLCF1 | cardiotrophin-like cytokine factor 1 | 15783 | -0.042 | -0.2436 | No |
| 145 | BCL3 | B-cell CLL/lymphoma 3 | 15800 | -0.042 | -0.2409 | No |
| 146 | TNFRSF9 | tumor necrosis factor receptor superfamily, member 9 | 15868 | -0.042 | -0.2407 | No |
| 147 | ABCA1 | ATP binding cassette subfamily A member 1 | 15924 | -0.043 | -0.2399 | No |
| 148 | BCL2A1 | BCL2-related protein A1 | 16007 | -0.044 | -0.2404 | No |
| 149 | KLF2 | Kruppel-like factor 2 | 16040 | -0.044 | -0.2383 | No |
| 150 | SLC2A3 | solute carrier family 2 (facilitated glucose transporter), member 3 | 16073 | -0.045 | -0.2362 | No |
| 151 | BMP2 | bone morphogenetic protein 2 | 16350 | -0.048 | -0.2464 | No |
| 152 | KLF9 | Kruppel-like factor 9 | 16381 | -0.048 | -0.2438 | No |
| 153 | IFIH1 | interferon induced, with helicase C domain 1 | 16404 | -0.048 | -0.2408 | No |
| 154 | B4GALT1 | UDP-Gal:betaGlcNAc beta 1,4- galactosyltransferase, polypeptide 1 | 16445 | -0.049 | -0.2387 | No |
| 155 | TLR2 | toll-like receptor 2 | 16448 | -0.049 | -0.2346 | No |
| 156 | AREG | amphiregulin | 16601 | -0.051 | -0.2381 | No |
| 157 | YRDC | yrdC N(6)-threonylcarbamoyltransferase domain containing | 16643 | -0.052 | -0.2358 | No |
| 158 | VEGFA | vascular endothelial growth factor A | 16659 | -0.052 | -0.2321 | No |
| 159 | NAMPT | nicotinamide phosphoribosyltransferase | 16683 | -0.052 | -0.2288 | No |
| 160 | LAMB3 | laminin, beta 3 | 16853 | -0.055 | -0.2329 | No |
| 161 | ICOSLG | inducible T-cell co-stimulator ligand | 16929 | -0.056 | -0.2320 | No |
| 162 | IER2 | immediate early response 2 | 17193 | -0.060 | -0.2405 | No |
| 163 | ATP2B1 | ATPase, Ca++ transporting, plasma membrane 1 | 17206 | -0.060 | -0.2360 | No |
| 164 | SLC16A6 | solute carrier family 16, member 6 | 17237 | -0.060 | -0.2324 | No |
| 165 | FOS | FBJ murine osteosarcoma viral oncogene homolog | 17528 | -0.066 | -0.2418 | No |
| 166 | KLF10 | Kruppel-like factor 10 | 17645 | -0.068 | -0.2420 | No |
| 167 | GPR183 | G protein-coupled receptor 183 | 17802 | -0.072 | -0.2439 | No |
| 168 | ZFP36 | ZFP36 ring finger protein | 17953 | -0.076 | -0.2452 | No |
| 169 | KLF4 | Kruppel-like factor 4 (gut) | 17974 | -0.076 | -0.2397 | No |
| 170 | IL23A | interleukin 23, alpha subunit p19 | 18037 | -0.078 | -0.2362 | No |
| 171 | SDC4 | syndecan 4 | 18151 | -0.081 | -0.2351 | No |
| 172 | MARCKS | myristoylated alanine-rich protein kinase C substrate | 18167 | -0.082 | -0.2289 | No |
| 173 | TNFAIP6 | tumor necrosis factor, alpha-induced protein 6 | 18183 | -0.082 | -0.2227 | No |
| 174 | SAT1 | spermidine/spermine N1-acetyltransferase 1 | 18295 | -0.085 | -0.2212 | No |
| 175 | MXD1 | Memczak2013 ANTISENSE, coding, INTERNAL, intronic best transcript NM\_001202514 | 18336 | -0.086 | -0.2159 | No |
| 176 | IL12B | interleukin 12B | 18360 | -0.087 | -0.2096 | No |
| 177 | SERPINB8 | serpin peptidase inhibitor, clade B (ovalbumin), member 8 | 18374 | -0.088 | -0.2028 | No |
| 178 | NFIL3 | nuclear factor, interleukin 3 regulated | 18475 | -0.092 | -0.2001 | No |
| 179 | EFNA1 | ephrin-A1 | 18705 | -0.103 | -0.2032 | No |
| 180 | G0S2 | G0/G1 switch 2 | 18709 | -0.103 | -0.1945 | No |
| 181 | IRF1 | interferon regulatory factor 1 | 18746 | -0.106 | -0.1873 | No |
| 182 | RCAN1 | regulator of calcineurin 1 | 18765 | -0.107 | -0.1791 | No |
| 183 | RHOB | ras homolog family member B | 18827 | -0.112 | -0.1727 | No |
| 184 | PLPP3 | phospholipid phosphatase 3 | 18866 | -0.115 | -0.1649 | No |
| 185 | PPP1R15A | protein phosphatase 1, regulatory subunit 15A | 18867 | -0.115 | -0.1551 | No |
| 186 | LITAF | lipopolysaccharide-induced TNF factor | 18924 | -0.119 | -0.1478 | No |
| 187 | FOSB | FBJ murine osteosarcoma viral oncogene homolog B | 18940 | -0.120 | -0.1383 | No |
| 188 | IL1B | interleukin 1 beta | 19023 | -0.130 | -0.1314 | No |
| 189 | JUN | jun proto-oncogene | 19037 | -0.132 | -0.1208 | No |
| 190 | PDE4B | phosphodiesterase 4B, cAMP-specific | 19061 | -0.135 | -0.1105 | No |
| 191 | IL6ST | interleukin 6 signal transducer | 19134 | -0.147 | -0.1017 | No |
| 192 | CCRL2 | chemokine (C-C motif) receptor-like 2 | 19232 | -0.168 | -0.0924 | No |
| 193 | BHLHE40 | basic helix-loop-helix family, member e40 | 19252 | -0.175 | -0.0784 | No |
| 194 | DUSP1 | dual specificity phosphatase 1 | 19260 | -0.177 | -0.0636 | No |
| 195 | EGR1 | early growth response 1 | 19334 | -0.202 | -0.0501 | No |
| 196 | PTGS2 | prostaglandin-endoperoxide synthase 2 (prostaglandin G/H synthase and cyclooxygenase) | 19364 | -0.218 | -0.0330 | No |
| 197 | NR4A3 | nuclear receptor subfamily 4, group A, member 3 | 19383 | -0.228 | -0.0145 | No |
| 198 | ATF3 | activating transcription factor 3 | 19421 | -0.254 | 0.0053 | No |
Table: GSEA details [plain text format]

  

Fig 2: HALLMARK\_TNFA\_SIGNALING\_VIA\_NFKB      
 Blue-Pink O' Gram in the Space of the Analyzed GeneSet

  

Fig 3: HALLMARK\_TNFA\_SIGNALING\_VIA\_NFKB: Random ES distribution      
 Gene set null distribution of ES for **HALLMARK\_TNFA\_SIGNALING\_VIA\_NFKB**

  
