## Supplementary material for "Somatic hypomethylation of pericentromeric SST1 repeats and tetraploidization in human colorectal cancer cells": GSEA results: HALLMARK_UNFOLDED_PROTEIN_RESPONSE.html

Details for gene set HALLMARK\_UNFOLDED\_PROTEIN\_RESPONSE[GSEA]

|  || Dataset | eset\_byprobe\_collapsed\_to\_symbols.Diploid\_vs\_Tetraploid.cls #Tetraploid\_versus\_Diploid.Diploid\_vs\_Tetraploid.cls #Tetraploid\_versus\_Diploid\_repos |
| Phenotype | Diploid\_vs\_Tetraploid.cls#Tetraploid\_versus\_Diploid\_repos |
| Upregulated in class | Diploid |
| GeneSet | HALLMARK\_UNFOLDED\_PROTEIN\_RESPONSE |
| Enrichment Score (ES) | -0.46685314 |
| Normalized Enrichment Score (NES) | -1.776545 |
| Nominal p-value | 0.0 |
| FDR q-value | 0.0069990745 |
| FWER p-Value | 0.01 |
Table: GSEA Results Summary

  

Fig 1: Enrichment plot: HALLMARK\_UNFOLDED\_PROTEIN\_RESPONSE      
 Profile of the Running ES Score & Positions of GeneSet Members on the Rank Ordered List

  

| SYMBOL | TITLE | RANK IN GENE LIST | RANK METRIC SCORE | RUNNING ES | CORE ENRICHMENT || 1 | WFS1 | Wolfram syndrome 1 (wolframin) | 150 | 0.172 | 0.0239 | No |
| 2 | GEMIN4 | gem nuclear organelle associated protein 4 | 802 | 0.092 | 0.0074 | No |
| 3 | NFYB | nuclear transcription factor Y subunit beta | 835 | 0.091 | 0.0225 | No |
| 4 | RRP9 | ribosomal RNA processing 9, small subunit (SSU) processome component, homolog (yeast) | 1374 | 0.075 | 0.0085 | No |
| 5 | EXOSC2 | exosome component 2 | 1485 | 0.072 | 0.0160 | No |
| 6 | BAG3 | BCL2-associated athanogene 3 | 1721 | 0.066 | 0.0162 | No |
| 7 | ZBTB17 | zinc finger and BTB domain containing 17 | 1843 | 0.064 | 0.0217 | No |
| 8 | IGFBP1 | insulin like growth factor binding protein 1 | 2762 | 0.050 | -0.0163 | No |
| 9 | NPM1 | nucleophosmin (nucleolar phosphoprotein B23, numatrin) | 2890 | 0.049 | -0.0138 | No |
| 10 | DDX10 | DEAD (Asp-Glu-Ala-Asp) box polypeptide 10 | 3120 | 0.046 | -0.0171 | No |
| 11 | DNAJA4 | DnaJ (Hsp40) homolog, subfamily A, member 4 | 3634 | 0.041 | -0.0359 | No |
| 12 | TATDN2 | TatD DNase domain containing 2 | 3688 | 0.041 | -0.0311 | No |
| 13 | NOP56 | NOP56 ribonucleoprotein | 3937 | 0.038 | -0.0369 | No |
| 14 | DCP2 | decapping mRNA 2 | 4517 | 0.033 | -0.0606 | No |
| 15 | DKC1 | dyskeratosis congenita 1, dyskerin | 4547 | 0.033 | -0.0561 | No |
| 16 | TUBB2A | tubulin, beta 2A class IIa | 4954 | 0.030 | -0.0715 | No |
| 17 | NOP14 | NOP14 nucleolar protein | 5088 | 0.029 | -0.0731 | No |
| 18 | EIF4A3 | eukaryotic translation initiation factor 4A3 | 5227 | 0.028 | -0.0751 | No |
| 19 | EDC4 | enhancer of mRNA decapping 4 | 5438 | 0.026 | -0.0812 | No |
| 20 | CCL2 | chemokine (C-C motif) ligand 2 | 5479 | 0.026 | -0.0785 | No |
| 21 | NHP2 | NHP2 ribonucleoprotein | 5638 | 0.024 | -0.0821 | No |
| 22 | EIF4E | eukaryotic translation initiation factor 4E | 5726 | 0.024 | -0.0822 | No |
| 23 | NOLC1 | nucleolar and coiled-body phosphoprotein 1 | 5963 | 0.022 | -0.0903 | No |
| 24 | CNOT6 | CCR4-NOT transcription complex subunit 6 | 6137 | 0.021 | -0.0953 | No |
| 25 | EXOSC9 | exosome component 9 | 6260 | 0.020 | -0.0978 | No |
| 26 | NABP1 | nucleic acid binding protein 1 | 6323 | 0.020 | -0.0973 | No |
| 27 | NFYA | nuclear transcription factor Y subunit alpha | 6480 | 0.019 | -0.1018 | No |
| 28 | EIF4A2 | eukaryotic translation initiation factor 4A2 | 6507 | 0.019 | -0.0996 | No |
| 29 | LSM4 | LSM4 homolog, U6 small nuclear RNA and mRNA degradation associated | 6745 | 0.017 | -0.1086 | No |
| 30 | SEC11A | SEC11 homolog A, signal peptidase complex subunit | 6770 | 0.017 | -0.1067 | No |
| 31 | KIF5B | Memczak2013 ANTISENSE, CDS, coding, INTERNAL, UTR3 best transcript NM\_004521 | 6866 | 0.017 | -0.1085 | No |
| 32 | YIF1A | Yip1 interacting factor homolog A (S. cerevisiae) | 7125 | 0.015 | -0.1190 | No |
| 33 | ATP6V0D1 | ATPase, H+ transporting, lysosomal 38kDa, V0 subunit d1 | 7524 | 0.013 | -0.1372 | No |
| 34 | EXOC2 | exocyst complex component 2 | 7660 | 0.012 | -0.1419 | No |
| 35 | EIF4A1 | eukaryotic translation initiation factor 4A1 | 8146 | 0.009 | -0.1653 | No |
| 36 | EXOSC5 | exosome component 5 | 8270 | 0.008 | -0.1701 | No |
| 37 | SPCS1 | signal peptidase complex subunit 1 | 8316 | 0.008 | -0.1710 | No |
| 38 | EXOSC4 | exosome component 4 | 8674 | 0.006 | -0.1883 | No |
| 39 | ALDH18A1 | aldehyde dehydrogenase 18 family, member A1 | 8962 | 0.004 | -0.2023 | No |
| 40 | HSPA9 | heat shock 70kDa protein 9 (mortalin) | 9059 | 0.004 | -0.2065 | No |
| 41 | KHSRP | KH-type splicing regulatory protein | 9258 | 0.003 | -0.2162 | No |
| 42 | PDIA6 | protein disulfide isomerase family A, member 6 | 9484 | 0.002 | -0.2275 | No |
| 43 | IMP3 | IMP3, U3 small nucleolar ribonucleoprotein | 9618 | 0.001 | -0.2342 | No |
| 44 | POP4 | POP4 homolog, ribonuclease P/MRP subunit | 9765 | 0.000 | -0.2417 | No |
| 45 | DCTN1 | dynactin 1 | 10021 | -0.001 | -0.2546 | No |
| 46 | CKS1B | CDC28 protein kinase regulatory subunit 1B | 10258 | -0.003 | -0.2663 | No |
| 47 | EIF2S1 | eukaryotic translation initiation factor 2, subunit 1 alpha, 35kDa | 10272 | -0.003 | -0.2664 | No |
| 48 | YWHAZ | tyrosine 3-monooxygenase/tryptophan 5-monooxygenase activation protein, zeta | 10747 | -0.006 | -0.2898 | No |
| 49 | FUS | FUS RNA binding protein | 10761 | -0.006 | -0.2894 | No |
| 50 | EEF2 | eukaryotic translation elongation factor 2 | 10886 | -0.006 | -0.2946 | No |
| 51 | LSM1 | LSM1 homolog, mRNA degradation associated | 11160 | -0.008 | -0.3072 | No |
| 52 | PREB | prolactin regulatory element binding | 11555 | -0.010 | -0.3256 | No |
| 53 | BANF1 | barrier to autointegration factor 1 | 11935 | -0.012 | -0.3429 | No |
| 54 | PDIA5 | protein disulfide isomerase family A, member 5 | 12042 | -0.013 | -0.3459 | No |
| 55 | RPS14 | ribosomal protein S14 | 12084 | -0.013 | -0.3456 | No |
| 56 | CNOT2 | CCR4-NOT transcription complex subunit 2 | 12493 | -0.016 | -0.3636 | No |
| 57 | PAIP1 | poly(A) binding protein interacting protein 1 | 12517 | -0.016 | -0.3618 | No |
| 58 | EIF4G1 | eukaryotic translation initiation factor 4 gamma, 1 | 12951 | -0.019 | -0.3807 | No |
| 59 | SLC30A5 | solute carrier family 30 (zinc transporter), member 5 | 13230 | -0.021 | -0.3912 | No |
| 60 | CNOT4 | CCR4-NOT transcription complex subunit 4 | 13794 | -0.025 | -0.4156 | No |
| 61 | ERO1A | endoplasmic reticulum oxidoreductase alpha | 13862 | -0.025 | -0.4144 | No |
| 62 | SDAD1 | SDA1 domain containing 1 | 14294 | -0.029 | -0.4313 | No |
| 63 | CEBPB | CCAAT/enhancer binding protein (C/EBP), beta | 14355 | -0.029 | -0.4290 | No |
| 64 | CALR | calreticulin | 14607 | -0.031 | -0.4362 | No |
| 65 | EXOSC10 | exosome component 10 | 14693 | -0.032 | -0.4347 | No |
| 66 | DCP1A | decapping mRNA 1A | 14807 | -0.033 | -0.4344 | No |
| 67 | PSAT1 | phosphoserine aminotransferase 1 | 14978 | -0.034 | -0.4369 | No |
| 68 | HSP90B1 | heat shock protein 90kDa beta (Grp94), member 1 | 15295 | -0.037 | -0.4463 | No |
| 69 | TTC37 | tetratricopeptide repeat domain 37 | 15361 | -0.038 | -0.4428 | No |
| 70 | XBP1 | X-box binding protein 1 | 15622 | -0.040 | -0.4488 | No |
| 71 | PARN | poly(A)-specific ribonuclease | 15672 | -0.041 | -0.4439 | No |
| 72 | EXOSC1 | exosome component 1 | 15721 | -0.041 | -0.4388 | No |
| 73 | ARFGAP1 | ADP-ribosylation factor GTPase activating protein 1 | 16267 | -0.047 | -0.4582 | Yes |
| 74 | SSR1 | signal sequence receptor, alpha | 16290 | -0.047 | -0.4507 | Yes |
| 75 | CXXC1 | CXXC finger protein 1 | 16296 | -0.047 | -0.4423 | Yes |
| 76 | ATF4 | activating transcription factor 4 | 16446 | -0.049 | -0.4409 | Yes |
| 77 | SPCS3 | signal peptidase complex subunit 3 | 16529 | -0.050 | -0.4359 | Yes |
| 78 | VEGFA | vascular endothelial growth factor A | 16659 | -0.052 | -0.4330 | Yes |
| 79 | EIF2AK3 | eukaryotic translation initiation factor 2-alpha kinase 3 | 16768 | -0.053 | -0.4287 | Yes |
| 80 | ATF6 | activating transcription factor 6 | 17060 | -0.058 | -0.4330 | Yes |
| 81 | TSPYL2 | TSPY-like 2 | 17146 | -0.059 | -0.4265 | Yes |
| 82 | STC2 | stanniocalcin 2 | 17326 | -0.062 | -0.4243 | Yes |
| 83 | XPOT | exportin, tRNA | 17359 | -0.063 | -0.4144 | Yes |
| 84 | SHC1 | SHC (Src homology 2 domain containing) transforming protein 1 | 17544 | -0.066 | -0.4116 | Yes |
| 85 | DDIT4 | DNA damage inducible transcript 4 | 17557 | -0.066 | -0.4000 | Yes |
| 86 | KDELR3 | KDEL (Lys-Asp-Glu-Leu) endoplasmic reticulum protein retention receptor 3 | 17705 | -0.070 | -0.3948 | Yes |
| 87 | EIF4EBP1 | eukaryotic translation initiation factor 4E binding protein 1 | 17800 | -0.072 | -0.3863 | Yes |
| 88 | SLC7A5 | solute carrier family 7 (amino acid transporter light chain, L system), member 5 | 17849 | -0.073 | -0.3753 | Yes |
| 89 | IFIT1 | interferon-induced protein with tetratricopeptide repeats 1 | 17946 | -0.076 | -0.3663 | Yes |
| 90 | SRPRB | signal recognition particle receptor, B subunit | 18026 | -0.078 | -0.3561 | Yes |
| 91 | EDEM1 | ER degradation enhancer, mannosidase alpha-like 1 | 18132 | -0.081 | -0.3466 | Yes |
| 92 | MTHFD2 | methylenetetrahydrofolate dehydrogenase (NADP+ dependent) 2, methenyltetrahydrofolate cyclohydrolase | 18255 | -0.084 | -0.3374 | Yes |
| 93 | CEBPG | CCAAT/enhancer binding protein (C/EBP), gamma | 18337 | -0.086 | -0.3257 | Yes |
| 94 | SEC31A | SEC31 homolog A, COPII coat complex component | 18367 | -0.088 | -0.3110 | Yes |
| 95 | HSPA5 | heat shock 70kDa protein 5 (glucose-regulated protein, 78kDa) | 18430 | -0.090 | -0.2977 | Yes |
| 96 | HYOU1 | hypoxia up-regulated 1 | 18453 | -0.091 | -0.2821 | Yes |
| 97 | FKBP14 | FK506 binding protein 14 | 18802 | -0.110 | -0.2798 | Yes |
| 98 | SLC1A4 | solute carrier family 1 (glutamate/neutral amino acid transporter), member 4 | 18809 | -0.110 | -0.2597 | Yes |
| 99 | ASNS | asparagine synthetase (glutamine-hydrolyzing) | 18864 | -0.115 | -0.2413 | Yes |
| 100 | HERPUD1 | homocysteine-inducible, endoplasmic reticulum stress-inducible, ubiquitin-like domain member 1 | 18948 | -0.121 | -0.2233 | Yes |
| 101 | DNAJC3 | DnaJ (Hsp40) homolog, subfamily C, member 3 | 19028 | -0.131 | -0.2033 | Yes |
| 102 | SERP1 | stress-associated endoplasmic reticulum protein 1 | 19031 | -0.131 | -0.1792 | Yes |
| 103 | GOSR2 | golgi SNAP receptor complex member 2 | 19041 | -0.132 | -0.1553 | Yes |
| 104 | ERN1 | endoplasmic reticulum to nucleus signaling 1 | 19173 | -0.153 | -0.1338 | Yes |
| 105 | CHAC1 | ChaC glutathione-specific gamma-glutamylcyclotransferase 1 | 19193 | -0.159 | -0.1054 | Yes |
| 106 | WIPI1 | WD repeat domain, phosphoinositide interacting 1 | 19291 | -0.190 | -0.0754 | Yes |
| 107 | DNAJB9 | DnaJ (Hsp40) homolog, subfamily B, member 9 | 19366 | -0.220 | -0.0387 | Yes |
| 108 | ATF3 | activating transcription factor 3 | 19421 | -0.254 | 0.0053 | Yes |
Table: GSEA details [plain text format]

  

Fig 2: HALLMARK\_UNFOLDED\_PROTEIN\_RESPONSE      
 Blue-Pink O' Gram in the Space of the Analyzed GeneSet

  

Fig 3: HALLMARK\_UNFOLDED\_PROTEIN\_RESPONSE: Random ES distribution      
 Gene set null distribution of ES for **HALLMARK\_UNFOLDED\_PROTEIN\_RESPONSE**

  
