## Supplementary material for "Somatic hypomethylation of pericentromeric SST1 repeats and tetraploidization in human colorectal cancer cells": GSEA results: HALLMARK_UV_RESPONSE_UP.html

Details for gene set HALLMARK\_UV\_RESPONSE\_UP[GSEA]

|  || Dataset | eset\_byprobe\_collapsed\_to\_symbols.Diploid\_vs\_Tetraploid.cls #Tetraploid\_versus\_Diploid.Diploid\_vs\_Tetraploid.cls #Tetraploid\_versus\_Diploid\_repos |
| Phenotype | Diploid\_vs\_Tetraploid.cls#Tetraploid\_versus\_Diploid\_repos |
| Upregulated in class | Diploid |
| GeneSet | HALLMARK\_UV\_RESPONSE\_UP |
| Enrichment Score (ES) | -0.26635388 |
| Normalized Enrichment Score (NES) | -1.0732045 |
| Nominal p-value | 0.28139904 |
| FDR q-value | 0.4224834 |
| FWER p-Value | 1.0 |
Table: GSEA Results Summary

  

Fig 1: Enrichment plot: HALLMARK\_UV\_RESPONSE\_UP      
 Profile of the Running ES Score & Positions of GeneSet Members on the Rank Ordered List

  

| SYMBOL | TITLE | RANK IN GENE LIST | RANK METRIC SCORE | RUNNING ES | CORE ENRICHMENT || 1 | CA2 | carbonic anhydrase II | 69 | 0.250 | 0.0306 | No |
| 2 | BTG2 | BTG family, member 2 | 281 | 0.136 | 0.0383 | No |
| 3 | CHKA | choline kinase alpha | 287 | 0.135 | 0.0566 | No |
| 4 | CHRNA5 | cholinergic receptor, nicotinic alpha 5 | 303 | 0.133 | 0.0740 | No |
| 5 | DNAJB1 | DnaJ (Hsp40) homolog, subfamily B, member 1 | 338 | 0.127 | 0.0897 | No |
| 6 | BTG3 | BTG family, member 3 | 465 | 0.113 | 0.0986 | No |
| 7 | GGH | gamma-glutamyl hydrolase (conjugase, folylpolygammaglutamyl hydrolase) | 487 | 0.111 | 0.1128 | No |
| 8 | GAL | galanin/GMAP prepropeptide | 581 | 0.104 | 0.1223 | No |
| 9 | CCNE1 | cyclin E1 | 738 | 0.095 | 0.1272 | No |
| 10 | FKBP4 | FK506 binding protein 4 | 973 | 0.085 | 0.1268 | No |
| 11 | MRPL23 | Homo sapiens mitochondrial ribosomal protein L23 (MRPL23), mRNA. | 1163 | 0.079 | 0.1279 | No |
| 12 | TUBA4A | tubulin, alpha 4a | 1323 | 0.076 | 0.1300 | No |
| 13 | DNAJA1 | DnaJ (Hsp40) homolog, subfamily A, member 1 | 1418 | 0.074 | 0.1353 | No |
| 14 | ICAM1 | intercellular adhesion molecule 1 | 1606 | 0.069 | 0.1350 | No |
| 15 | PARP2 | poly(ADP-ribose) polymerase 2 | 1614 | 0.068 | 0.1441 | No |
| 16 | HSPA2 | heat shock 70kDa protein 2 | 1901 | 0.063 | 0.1379 | No |
| 17 | NTRK3 | neurotrophic tyrosine kinase, receptor, type 3 | 1988 | 0.062 | 0.1419 | No |
| 18 | CYB5B | cytochrome b5 type B (outer mitochondrial membrane) | 2082 | 0.060 | 0.1454 | No |
| 19 | FGF18 | fibroblast growth factor 18 | 2138 | 0.059 | 0.1506 | No |
| 20 | CDC34 | cell division cycle 34 | 2318 | 0.057 | 0.1491 | No |
| 21 | NXF1 | nuclear RNA export factor 1 | 2344 | 0.056 | 0.1555 | No |
| 22 | CTSV | cathepsin V | 2384 | 0.056 | 0.1611 | No |
| 23 | CDK2 | cyclin-dependent kinase 2 | 2666 | 0.051 | 0.1536 | No |
| 24 | POLR2H | polymerase (RNA) II (DNA directed) polypeptide H | 2849 | 0.049 | 0.1510 | No |
| 25 | AMD1 | adenosylmethionine decarboxylase 1 | 3049 | 0.047 | 0.1471 | No |
| 26 | CXCL2 | chemokine (C-X-C motif) ligand 2 | 3085 | 0.047 | 0.1517 | No |
| 27 | ENO2 | enolase 2 (gamma, neuronal) | 3086 | 0.047 | 0.1581 | No |
| 28 | E2F5 | E2F transcription factor 5, p130-binding | 3444 | 0.043 | 0.1456 | No |
| 29 | HMOX1 | heme oxygenase 1 | 3502 | 0.043 | 0.1485 | No |
| 30 | TCHH | trichohyalin | 3503 | 0.043 | 0.1543 | No |
| 31 | BID | BH3 interacting domain death agonist | 3546 | 0.042 | 0.1579 | No |
| 32 | HTR7 | 5-hydroxytryptamine (serotonin) receptor 7, adenylate cyclase-coupled | 3764 | 0.040 | 0.1522 | No |
| 33 | STIP1 | stress-induced phosphoprotein 1 | 4082 | 0.037 | 0.1408 | No |
| 34 | TACR3 | tachykinin receptor 3 | 4600 | 0.032 | 0.1186 | No |
| 35 | GRPEL1 | GrpE-like 1, mitochondrial (E. coli) | 4601 | 0.032 | 0.1230 | No |
| 36 | STK25 | serine/threonine kinase 25 | 4635 | 0.032 | 0.1257 | No |
| 37 | PSMC3 | proteasome 26S subunit, ATPase 3 | 4643 | 0.032 | 0.1297 | No |
| 38 | CASP3 | caspase 3 | 4699 | 0.032 | 0.1312 | No |
| 39 | NAT1 | N-acetyltransferase 1 (arylamine N-acetyltransferase) | 4763 | 0.031 | 0.1322 | No |
| 40 | RET | ret proto-oncogene | 4931 | 0.030 | 0.1276 | No |
| 41 | PPT1 | palmitoyl-protein thioesterase 1 | 5049 | 0.029 | 0.1255 | No |
| 42 | KLHDC3 | kelch domain containing 3 | 5243 | 0.027 | 0.1193 | No |
| 43 | TGFBRAP1 | transforming growth factor beta receptor associated protein 1 | 5271 | 0.027 | 0.1217 | No |
| 44 | PLCL1 | phospholipase C-like 1 | 5493 | 0.026 | 0.1137 | No |
| 45 | RASGRP1 | RAS guanyl releasing protein 1 (calcium and DAG-regulated) | 5558 | 0.025 | 0.1139 | No |
| 46 | CDC5L | cell division cycle 5-like | 5697 | 0.024 | 0.1100 | No |
| 47 | AGO2 | argonaute RISC catalytic component 2 | 5699 | 0.024 | 0.1133 | No |
| 48 | RFC4 | replication factor C subunit 4 | 5802 | 0.023 | 0.1112 | No |
| 49 | SLC6A12 | solute carrier family 6 (neurotransmitter transporter), member 12 | 6101 | 0.021 | 0.0988 | No |
| 50 | SULT1A1 | sulfotransferase family 1A member 1 | 6114 | 0.021 | 0.1011 | No |
| 51 | TAP1 | transporter 1, ATP-binding cassette, sub-family B (MDR/TAP) | 6169 | 0.021 | 0.1011 | No |
| 52 | COL2A1 | collagen, type II, alpha 1 | 6341 | 0.020 | 0.0951 | No |
| 53 | OLFM1 | olfactomedin 1 | 6358 | 0.020 | 0.0970 | No |
| 54 | ALDOA | aldolase A, fructose-bisphosphate | 6361 | 0.020 | 0.0996 | No |
| 55 | FEN1 | flap structure-specific endonuclease 1 | 6460 | 0.019 | 0.0972 | No |
| 56 | BCL2L11 | BCL2-like 11 (apoptosis facilitator) | 6584 | 0.018 | 0.0933 | No |
| 57 | ACAA1 | acetyl-CoA acyltransferase 1 | 6834 | 0.017 | 0.0828 | No |
| 58 | DGAT1 | diacylglycerol O-acyltransferase 1 | 6889 | 0.017 | 0.0823 | No |
| 59 | SIGMAR1 | sigma non-opioid intracellular receptor 1 | 6912 | 0.016 | 0.0834 | No |
| 60 | CYP1A1 | cytochrome P450, family 1, subfamily A, polypeptide 1 | 6941 | 0.016 | 0.0841 | No |
| 61 | POLG2 | polymerase (DNA directed), gamma 2, accessory subunit | 6948 | 0.016 | 0.0860 | No |
| 62 | UROD | uroporphyrinogen decarboxylase | 7213 | 0.015 | 0.0744 | No |
| 63 | PPIF | peptidylprolyl isomerase F | 7224 | 0.015 | 0.0759 | No |
| 64 | BTG1 | B-cell translocation gene 1, anti-proliferative | 7702 | 0.012 | 0.0529 | No |
| 65 | PRPF3 | pre-mRNA processing factor 3 | 7780 | 0.011 | 0.0504 | No |
| 66 | POLE3 | polymerase (DNA directed), epsilon 3, accessory subunit | 8126 | 0.009 | 0.0339 | No |
| 67 | SOD2 | superoxide dismutase 2, mitochondrial | 8450 | 0.007 | 0.0182 | No |
| 68 | NKX2-5 | NK2 homeobox 5 | 8499 | 0.007 | 0.0167 | No |
| 69 | HNRNPU | heterogeneous nuclear ribonucleoprotein U (scaffold attachment factor A) | 8561 | 0.007 | 0.0144 | No |
| 70 | HLA-F | major histocompatibility complex, class I, F | 8584 | 0.006 | 0.0141 | No |
| 71 | ONECUT1 | one cut homeobox 1 | 8771 | 0.005 | 0.0053 | No |
| 72 | CLTB | clathrin, light chain B | 8783 | 0.005 | 0.0054 | No |
| 73 | AQP3 | aquaporin 3 (Gill blood group) | 8812 | 0.005 | 0.0047 | No |
| 74 | CNP | 2,3-cyclic nucleotide 3 phosphodiesterase | 8879 | 0.005 | 0.0019 | No |
| 75 | HYAL2 | hyaluronoglucosaminidase 2 | 8921 | 0.005 | 0.0004 | No |
| 76 | TFRC | transferrin receptor | 8948 | 0.004 | -0.0003 | No |
| 77 | EPCAM | epithelial cell adhesion molecule | 8997 | 0.004 | -0.0022 | No |
| 78 | MMP14 | matrix metallopeptidase 14 (membrane-inserted) | 9110 | 0.003 | -0.0076 | No |
| 79 | SPR | sepiapterin reductase (7,8-dihydrobiopterin:NADP+ oxidoreductase) | 9651 | 0.001 | -0.0353 | No |
| 80 | NFKBIA | nuclear factor of kappa light polypeptide gene enhancer in B-cells inhibitor, alpha | 9745 | 0.000 | -0.0401 | No |
| 81 | ARRB2 | arrestin, beta 2 | 9889 | -0.001 | -0.0474 | No |
| 82 | GLS | glutaminase | 10027 | -0.001 | -0.0543 | No |
| 83 | PPAT | phosphoribosyl pyrophosphate amidotransferase | 10084 | -0.002 | -0.0570 | No |
| 84 | AP2S1 | adaptor-related protein complex 2 sigma 1 subunit | 10452 | -0.004 | -0.0754 | No |
| 85 | ALAS1 | 5-aminolevulinate synthase 1 | 10706 | -0.005 | -0.0877 | No |
| 86 | SLC6A8 | solute carrier family 6 (neurotransmitter transporter), member 8 | 10735 | -0.006 | -0.0884 | No |
| 87 | EIF5 | eukaryotic translation initiation factor 5 | 10768 | -0.006 | -0.0893 | No |
| 88 | JUNB | jun B proto-oncogene | 10813 | -0.006 | -0.0907 | No |
| 89 | WIZ | widely interspaced zinc finger motifs | 10825 | -0.006 | -0.0904 | No |
| 90 | NR4A1 | nuclear receptor subfamily 4, group A, member 1 | 11095 | -0.008 | -0.1033 | No |
| 91 | CDKN1C | cyclin-dependent kinase inhibitor 1C (p57, Kip2) | 11246 | -0.008 | -0.1099 | No |
| 92 | SHOX2 | short stature homeobox 2 | 11484 | -0.010 | -0.1208 | No |
| 93 | PRKACA | protein kinase, cAMP-dependent, catalytic, alpha | 11532 | -0.010 | -0.1218 | No |
| 94 | BAK1 | BCL2-antagonist/killer 1 | 11698 | -0.011 | -0.1288 | No |
| 95 | RAB27A | RAB27A, member RAS oncogene family | 11762 | -0.011 | -0.1305 | No |
| 96 | IL6 | interleukin 6 | 11825 | -0.012 | -0.1321 | No |
| 97 | EIF2S3 | eukaryotic translation initiation factor 2, subunit 3 gamma, 52kDa | 11933 | -0.012 | -0.1359 | No |
| 98 | DDX21 | DEAD (Asp-Glu-Ala-Asp) box helicase 21 | 12006 | -0.013 | -0.1379 | No |
| 99 | TMBIM6 | transmembrane BAX inhibitor motif containing 6 | 12126 | -0.014 | -0.1422 | No |
| 100 | CCK | cholecystokinin | 12181 | -0.014 | -0.1430 | No |
| 101 | GRINA | glutamate receptor, ionotropic, N-methyl D-aspartate-associated protein 1 (glutamate binding) | 12443 | -0.016 | -0.1544 | No |
| 102 | CLCN2 | chloride channel, voltage-sensitive 2 | 12570 | -0.017 | -0.1586 | No |
| 103 | ATP6V1F | ATPase, H+ transporting, lysosomal 14kDa, V1 subunit F | 12691 | -0.017 | -0.1624 | No |
| 104 | C4BPB | complement component 4 binding protein, beta | 12745 | -0.018 | -0.1628 | No |
| 105 | GCH1 | GTP cyclohydrolase 1 | 12793 | -0.018 | -0.1628 | No |
| 106 | EPHX1 | epoxide hydrolase 1, microsomal (xenobiotic) | 12904 | -0.019 | -0.1659 | No |
| 107 | SLC25A4 | solute carrier family 25 (mitochondrial carrier; adenine nucleotide translocator), member 4 | 12970 | -0.019 | -0.1666 | No |
| 108 | STARD3 | StAR-related lipid transfer domain containing 3 | 12983 | -0.019 | -0.1646 | No |
| 109 | SQSTM1 | sequestosome 1 | 13032 | -0.019 | -0.1645 | No |
| 110 | MARK2 | MAP/microtubule affinity-regulating kinase 2 | 13306 | -0.021 | -0.1756 | No |
| 111 | PDLIM3 | PDZ and LIM domain 3 | 13450 | -0.022 | -0.1800 | No |
| 112 | PRKCD | protein kinase C, delta | 13658 | -0.024 | -0.1874 | No |
| 113 | GPX3 | glutathione peroxidase 3 | 13735 | -0.024 | -0.1880 | No |
| 114 | MAPK8IP2 | mitogen-activated protein kinase 8 interacting protein 2 | 13766 | -0.025 | -0.1862 | No |
| 115 | TYRO3 | TYRO3 protein tyrosine kinase | 14020 | -0.026 | -0.1956 | No |
| 116 | RRAD | Ras-related associated with diabetes | 14315 | -0.029 | -0.2069 | No |
| 117 | LYN | LYN proto-oncogene, Src family tyrosine kinase | 14438 | -0.030 | -0.2091 | No |
| 118 | CYB5R1 | cytochrome b5 reductase 1 | 14673 | -0.032 | -0.2168 | No |
| 119 | CCND3 | cyclin D3 | 14958 | -0.034 | -0.2268 | No |
| 120 | BSG | basigin (Ok blood group) | 14969 | -0.034 | -0.2226 | No |
| 121 | CDKN2B | cyclin-dependent kinase inhibitor 2B (p15, inhibits CDK4) | 15064 | -0.035 | -0.2227 | No |
| 122 | DLG4 | discs, large homolog 4 (Drosophila) | 15378 | -0.038 | -0.2337 | No |
| 123 | NUP58 | nucleoporin 58kDa | 15532 | -0.039 | -0.2362 | No |
| 124 | TST | thiosulfate sulfurtransferase (rhodanese) | 15640 | -0.040 | -0.2363 | No |
| 125 | RXRB | retinoid X receptor beta | 15723 | -0.041 | -0.2349 | No |
| 126 | NPTX2 | neuronal pentraxin II | 15942 | -0.043 | -0.2402 | No |
| 127 | CREG1 | cellular repressor of E1A-stimulated genes 1 | 16096 | -0.045 | -0.2420 | No |
| 128 | MSX1 | msh homeobox 1 | 16206 | -0.046 | -0.2413 | No |
| 129 | FMO1 | flavin containing monooxygenase 1 | 16324 | -0.047 | -0.2409 | No |
| 130 | BMP2 | bone morphogenetic protein 2 | 16350 | -0.048 | -0.2356 | No |
| 131 | NPTXR | neuronal pentraxin receptor | 16426 | -0.049 | -0.2328 | No |
| 132 | ATP6V1C1 | ATPase, H+ transporting, lysosomal 42kDa, V1 subunit C1 | 16733 | -0.053 | -0.2414 | No |
| 133 | LHX2 | LIM homeobox 2 | 16861 | -0.055 | -0.2405 | No |
| 134 | PDAP1 | PDGFA associated protein 1 | 17018 | -0.057 | -0.2407 | No |
| 135 | RPN1 | ribophorin I | 17081 | -0.058 | -0.2359 | No |
| 136 | FOS | FBJ murine osteosarcoma viral oncogene homolog | 17528 | -0.066 | -0.2499 | No |
| 137 | APOM | apolipoprotein M | 17542 | -0.066 | -0.2416 | No |
| 138 | FURIN | furin (paired basic amino acid cleaving enzyme) | 18023 | -0.077 | -0.2558 | Yes |
| 139 | ABCB1 | ATP binding cassette subfamily B member 1 | 18030 | -0.078 | -0.2454 | Yes |
| 140 | PPP1R2 | protein phosphatase 1, regulatory (inhibitor) subunit 2 | 18051 | -0.078 | -0.2358 | Yes |
| 141 | CDO1 | cysteine dioxygenase type 1 | 18137 | -0.081 | -0.2291 | Yes |
| 142 | YKT6 | YKT6 v-SNARE homolog (S. cerevisiae) | 18191 | -0.082 | -0.2206 | Yes |
| 143 | CEBPG | CCAAT/enhancer binding protein (C/EBP), gamma | 18337 | -0.086 | -0.2162 | Yes |
| 144 | IRF1 | interferon regulatory factor 1 | 18746 | -0.106 | -0.2228 | Yes |
| 145 | RHOB | ras homolog family member B | 18827 | -0.112 | -0.2116 | Yes |
| 146 | ASNS | asparagine synthetase (glutamine-hydrolyzing) | 18864 | -0.115 | -0.1978 | Yes |
| 147 | FOSB | FBJ murine osteosarcoma viral oncogene homolog B | 18940 | -0.120 | -0.1852 | Yes |
| 148 | MGAT1 | mannosyl (alpha-1,3-)-glycoprotein beta-1,2-N-acetylglucosaminyltransferase | 19066 | -0.136 | -0.1730 | Yes |
| 149 | HSPA13 | heat shock protein 70kDa family, member 13 | 19126 | -0.146 | -0.1561 | Yes |
| 150 | IL6ST | interleukin 6 signal transducer | 19134 | -0.147 | -0.1363 | Yes |
| 151 | MAOA | monoamine oxidase A | 19140 | -0.148 | -0.1163 | Yes |
| 152 | IGFBP2 | insulin like growth factor binding protein 2 | 19180 | -0.155 | -0.0971 | Yes |
| 153 | PTPRD | protein tyrosine phosphatase, receptor type, D | 19216 | -0.165 | -0.0763 | Yes |
| 154 | ATF3 | activating transcription factor 3 | 19421 | -0.254 | -0.0520 | Yes |
| 155 | KCNH2 | potassium channel, voltage gated eag related subfamily H, member 2 | 19505 | -0.419 | 0.0010 | Yes |
Table: GSEA details [plain text format]

  

Fig 2: HALLMARK\_UV\_RESPONSE\_UP      
 Blue-Pink O' Gram in the Space of the Analyzed GeneSet

  

Fig 3: HALLMARK\_UV\_RESPONSE\_UP: Random ES distribution      
 Gene set null distribution of ES for **HALLMARK\_UV\_RESPONSE\_UP**

  
