## Supplementary material for "Somatic hypomethylation of pericentromeric SST1 repeats and tetraploidization in human colorectal cancer cells": GSEA results: HALLMARK_WNT_BETA_CATENIN_SIGNALING.html

Details for gene set HALLMARK\_WNT\_BETA\_CATENIN\_SIGNALING[GSEA]

|  || Dataset | eset\_byprobe\_collapsed\_to\_symbols.Diploid\_vs\_Tetraploid.cls #Tetraploid\_versus\_Diploid.Diploid\_vs\_Tetraploid.cls #Tetraploid\_versus\_Diploid\_repos |
| Phenotype | Diploid\_vs\_Tetraploid.cls#Tetraploid\_versus\_Diploid\_repos |
| Upregulated in class | Diploid |
| GeneSet | HALLMARK\_WNT\_BETA\_CATENIN\_SIGNALING |
| Enrichment Score (ES) | -0.44881976 |
| Normalized Enrichment Score (NES) | -1.4505308 |
| Nominal p-value | 0.060763888 |
| FDR q-value | 0.10252717 |
| FWER p-Value | 0.437 |
Table: GSEA Results Summary

  

Fig 1: Enrichment plot: HALLMARK\_WNT\_BETA\_CATENIN\_SIGNALING      
 Profile of the Running ES Score & Positions of GeneSet Members on the Rank Ordered List

  

| SYMBOL | TITLE | RANK IN GENE LIST | RANK METRIC SCORE | RUNNING ES | CORE ENRICHMENT || 1 | CCND2 | cyclin D2 | 34 | 0.313 | 0.1031 | No |
| 2 | DKK1 | dickkopf WNT signaling pathway inhibitor 1 | 106 | 0.204 | 0.1677 | No |
| 3 | JAG1 | jagged 1 | 962 | 0.085 | 0.1525 | No |
| 4 | NOTCH1 | notch 1 | 1193 | 0.079 | 0.1670 | No |
| 5 | AXIN2 | axin 2 | 1385 | 0.074 | 0.1821 | No |
| 6 | FRAT1 | frequently rearranged in advanced T-cell lymphomas 1 | 1769 | 0.065 | 0.1844 | No |
| 7 | WNT1 | wingless-type MMTV integration site family, member 1 | 2270 | 0.057 | 0.1779 | No |
| 8 | WNT5B | wingless-type MMTV integration site family, member 5B | 3027 | 0.047 | 0.1549 | No |
| 9 | MYC | v-myc avian myelocytomatosis viral oncogene homolog | 3194 | 0.046 | 0.1617 | No |
| 10 | SKP2 | S-phase kinase-associated protein 2, E3 ubiquitin protein ligase | 7073 | 0.015 | -0.0322 | No |
| 11 | AXIN1 | axin 1 | 7356 | 0.014 | -0.0420 | No |
| 12 | NOTCH4 | notch 4 | 8040 | 0.010 | -0.0738 | No |
| 13 | MAML1 | mastermind-like transcriptional coactivator 1 | 8044 | 0.010 | -0.0707 | No |
| 14 | FZD8 | frizzled class receptor 8 | 8074 | 0.009 | -0.0691 | No |
| 15 | HDAC11 | histone deacetylase 11 | 8563 | 0.007 | -0.0919 | No |
| 16 | KAT2A | Memczak2013 ALT\_DONOR, coding, INTERNAL, intronic best transcript NM\_021078 | 9189 | 0.003 | -0.1230 | No |
| 17 | PPARD | peroxisome proliferator-activated receptor delta | 9547 | 0.001 | -0.1409 | No |
| 18 | JAG2 | jagged 2 | 10208 | -0.002 | -0.1740 | No |
| 19 | PTCH1 | patched 1 | 10420 | -0.004 | -0.1836 | No |
| 20 | HDAC2 | histone deacetylase 2 | 10584 | -0.005 | -0.1904 | No |
| 21 | WNT6 | wingless-type MMTV integration site family, member 6 | 11094 | -0.008 | -0.2140 | No |
| 22 | CTNNB1 | catenin (cadherin-associated protein), beta 1 | 11452 | -0.010 | -0.2291 | No |
| 23 | DVL2 | dishevelled segment polarity protein 2 | 11675 | -0.011 | -0.2368 | No |
| 24 | NUMB | numb homolog (Drosophila) | 12111 | -0.014 | -0.2546 | No |
| 25 | NCOR2 | nuclear receptor corepressor 2 | 12339 | -0.015 | -0.2612 | No |
| 26 | TCF7 | transcription factor 7 (T-cell specific, HMG-box) | 12391 | -0.015 | -0.2587 | No |
| 27 | CSNK1E | casein kinase 1, epsilon | 13231 | -0.021 | -0.2948 | No |
| 28 | HDAC5 | histone deacetylase 5 | 13792 | -0.025 | -0.3152 | No |
| 29 | TP53 | tumor protein p53 | 14048 | -0.027 | -0.3194 | No |
| 30 | NCSTN | nicastrin | 14058 | -0.027 | -0.3109 | No |
| 31 | RBPJ | recombination signal binding protein for immunoglobulin kappa J region | 14632 | -0.031 | -0.3298 | No |
| 32 | FZD1 | frizzled class receptor 1 | 15704 | -0.041 | -0.3711 | No |
| 33 | CUL1 | cullin 1 | 15794 | -0.042 | -0.3617 | No |
| 34 | ADAM17 | ADAM metallopeptidase domain 17 | 16706 | -0.053 | -0.3908 | No |
| 35 | PSEN2 | presenilin 2 | 17539 | -0.066 | -0.4114 | No |
| 36 | HEY1 | hes-related family bHLH transcription factor with YRPW motif 1 | 18270 | -0.085 | -0.4205 | Yes |
| 37 | NKD1 | naked cuticle homolog 1 (Drosophila) | 18631 | -0.098 | -0.4061 | Yes |
| 38 | DLL1 | delta-like 1 (Drosophila) | 18868 | -0.115 | -0.3797 | Yes |
| 39 | HEY2 | hes-related family bHLH transcription factor with YRPW motif 2 | 18894 | -0.118 | -0.3416 | Yes |
| 40 | LEF1 | lymphoid enhancer-binding factor 1 | 19413 | -0.250 | -0.2843 | Yes |
| 41 | GNAI1 | guanine nucleotide binding protein (G protein), alpha inhibiting activity polypeptide 1 | 19420 | -0.254 | -0.1996 | Yes |
| 42 | DKK4 | dickkopf WNT signaling pathway inhibitor 4 | 19524 | -0.611 | 0.0000 | Yes |
Table: GSEA details [plain text format]

  

Fig 2: HALLMARK\_WNT\_BETA\_CATENIN\_SIGNALING      
 Blue-Pink O' Gram in the Space of the Analyzed GeneSet

  

Fig 3: HALLMARK\_WNT\_BETA\_CATENIN\_SIGNALING: Random ES distribution      
 Gene set null distribution of ES for **HALLMARK\_WNT\_BETA\_CATENIN\_SIGNALING**

  
