## Supplementary material for "Somatic hypomethylation of pericentromeric SST1 repeats and tetraploidization in human colorectal cancer cells": GSEA results: heat_map_corr_plot.html

Heat map and correlation plot for eset\_byprobe\_collapsed\_to\_symbols.Diploid\_vs\_Tetraploid.cls#Tetraploid\_versus\_Diploid  

Fig 1: heat\_map      
 Heat Map of the top 50 features for each phenotype in eset\_byprobe\_collapsed\_to\_symbols.Diploid\_vs\_Tetraploid.cls#Tetraploid\_versus\_Diploid

  
  

Fig 2: Ranked Gene List Correlation Profile      
 Ranked list correlations for eset\_byprobe\_collapsed\_to\_symbols.Diploid\_vs\_Tetraploid.cls#Tetraploid\_versus\_Diploid

  
  
    
