## Supplementary material for "Somatic hypomethylation of pericentromeric SST1 repeats and tetraploidization in human colorectal cancer cells": GSEA results: index.html

Index for xtools.gsea.Gsea my\_analysis.Gsea.1608738186968

### GSEA Report for Dataset eset\_byprobe

#### Enrichment in phenotype: **Tetraploid (4 samples)**

- 27 / 50 gene sets are upregulated in phenotype **Tetraploid**- 14 gene sets are significant at FDR < 25%- 7 gene sets are significantly enriched at nominal pvalue < 1%- 11 gene sets are significantly enriched at nominal pvalue < 5%- Snapshot of enrichment results- Detailed enrichment results in html format- Detailed enrichment results in TSV format (tab delimited text)- Guide to interpret results

#### Enrichment in phenotype: **Diploid (4 samples)**

- 23 / 50 gene sets are upregulated in phenotype **Diploid**- 4 gene sets are significantly enriched at FDR < 25%- 3 gene sets are significantly enriched at nominal pvalue < 1%- 4 gene sets are significantly enriched at nominal pvalue < 5%- Snapshot of enrichment results- Detailed enrichment results in html format- Detailed enrichment results in TSV format (tab delimited text)- Guide to interpret results

#### Dataset details

- The dataset has 21448 native features- After collapsing features into gene symbols, there are: 19525 genes

#### Gene set details

- Gene set size filters (min=10, max=500) resulted in filtering out 0 / 50 gene sets- The remaining 50 gene sets were used in the analysis- List of gene sets used and their sizes (restricted to features in the specified dataset)

#### Gene markers for the **Tetraploid** *versus* **Diploid** comparison

- The dataset has 19525 features (genes)- # of markers for phenotype **Tetraploid**: 9779 (50.1% ) with correlation area 47.6%- # of markers for phenotype **Diploid**: 9746 (49.9% ) with correlation area 52.4%- Detailed rank ordered gene list for all features in the dataset- Heat map and gene list correlation  profile for all features in the dataset

#### Global statistics and plots

- Plot of p-values *vs.* NES- Global ES histogram

#### Other

- Parameters used for this analysis

#### Comments

- Timestamp used as random seed: 1608738189173

---

Report: my\_analysis.Gsea.1608738186968.rpt   by user: salonso

xtools.gsea.Gsea [Wed, Dec 23, '20 4 PM 43]

Website: www.gsea-msigdb.org/gsea
Questions & Suggestions: Contact page
