## Supplementary figures and images for "Somatic hypomethylation of pericentromeric SST1 repeats and tetraploidization in human colorectal cancer cells"

### enplot_HALLMARK_DNA_REPAIR_301.png

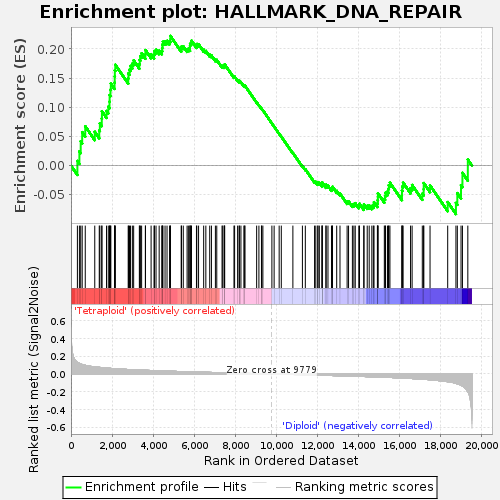

### enplot_HALLMARK_ESTROGEN_RESPONSE_LATE_268.png

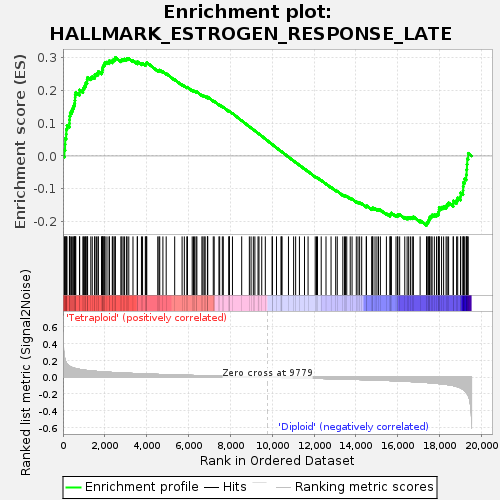

### enplot_HALLMARK_HEDGEHOG_SIGNALING_298.png

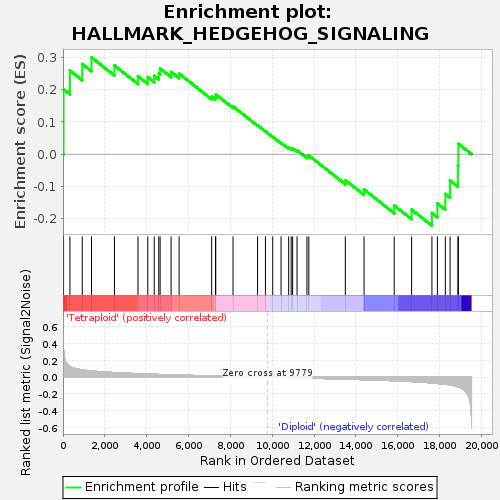

### enplot_HALLMARK_IL6_JAK_STAT3_SIGNALING_361.png

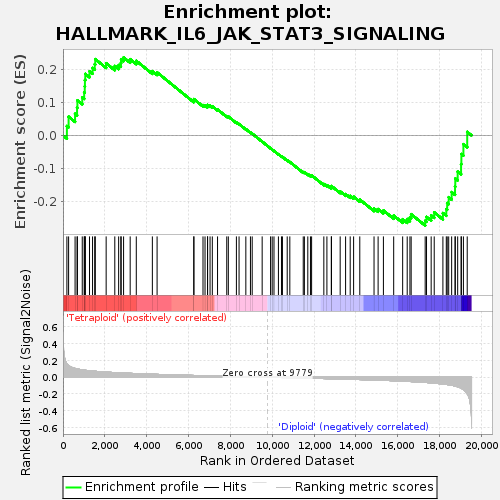

### enplot_HALLMARK_INTERFERON_ALPHA_RESPONSE_331.png

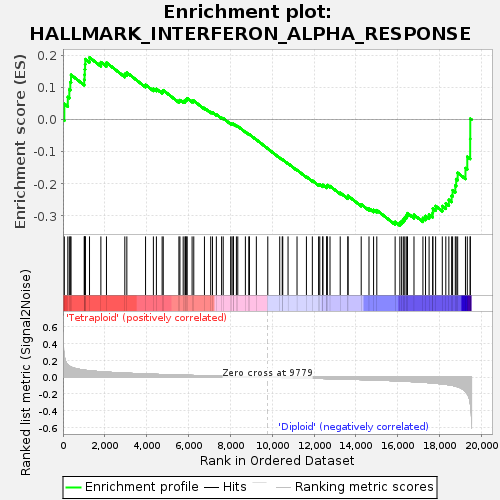

### enplot_HALLMARK_PROTEIN_SECRETION_310.png

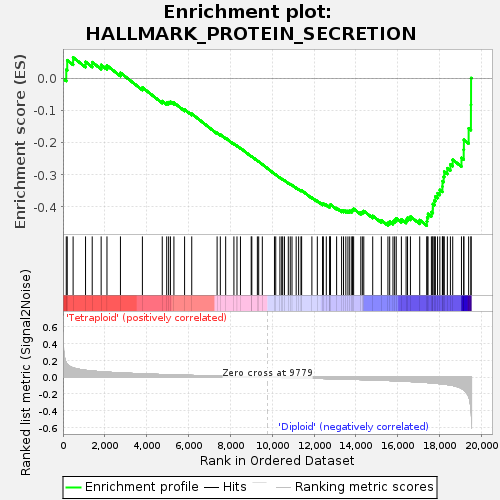

### gset_rnd_es_dist_273.png

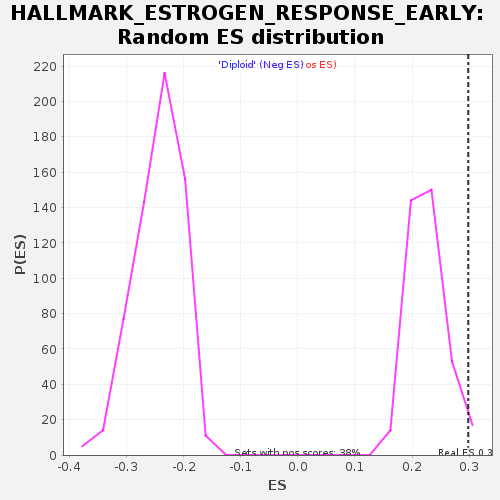

### gset_rnd_es_dist_288.png

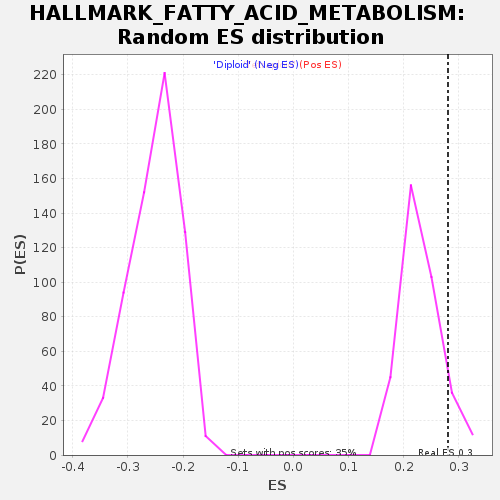

### HALLMARK_ANDROGEN_RESPONSE_365.png

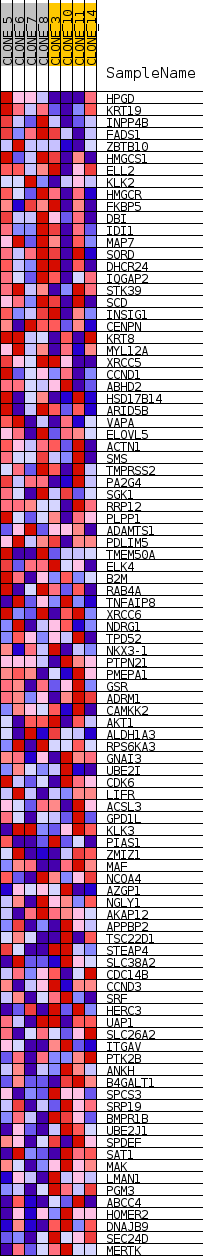

### HALLMARK_ESTROGEN_RESPONSE_EARLY_272.png

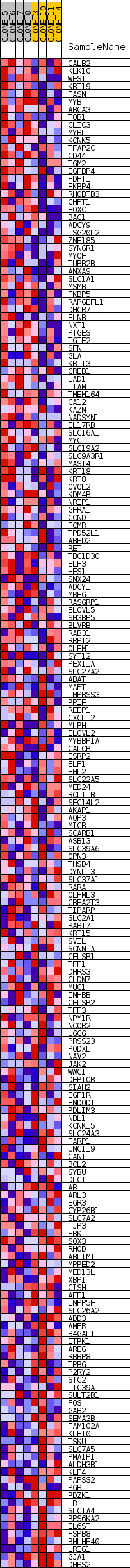

### HALLMARK_P53_PATHWAY_266.png

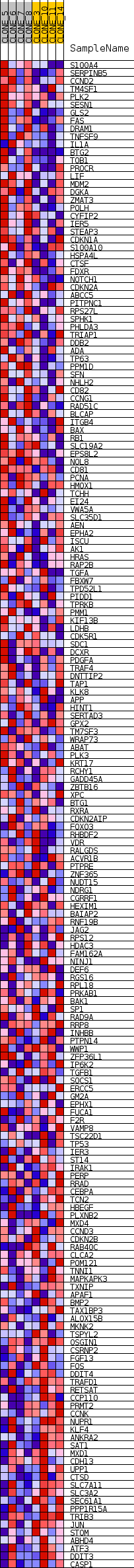

### HALLMARK_SPERMATOGENESIS_305.png

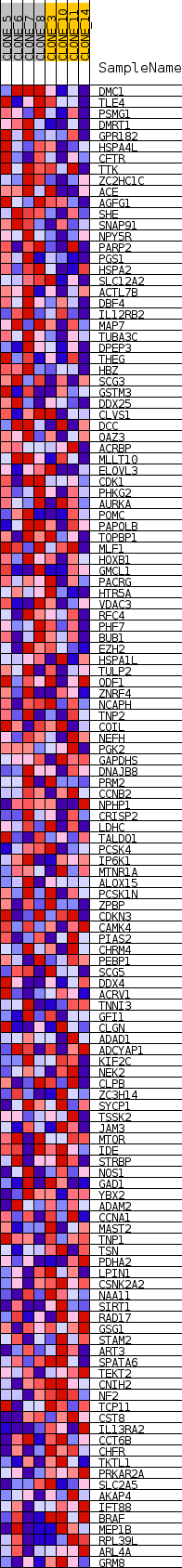
