## Supplementary figures and images for "Somatic hypomethylation of pericentromeric SST1 repeats and tetraploidization in human colorectal cancer cells"

### enplot_HALLMARK_ALLOGRAFT_REJECTION_277.png

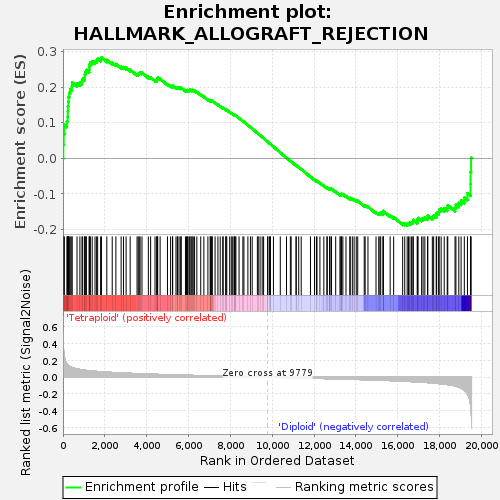

### enplot_HALLMARK_ANDROGEN_RESPONSE_364.png

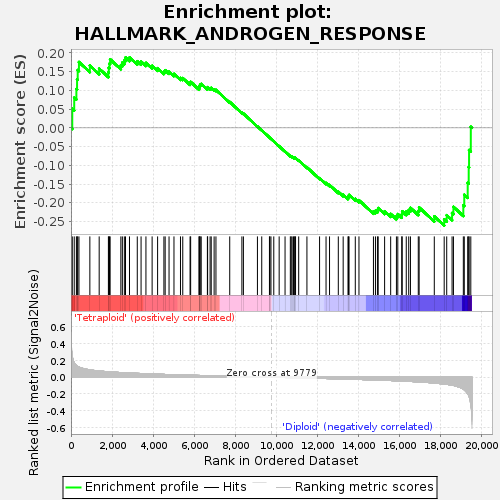

### enplot_HALLMARK_ANGIOGENESIS_274.png

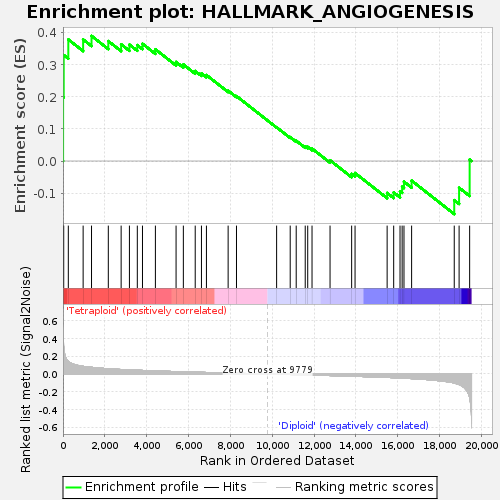

### enplot_HALLMARK_APICAL_SURFACE_295.png

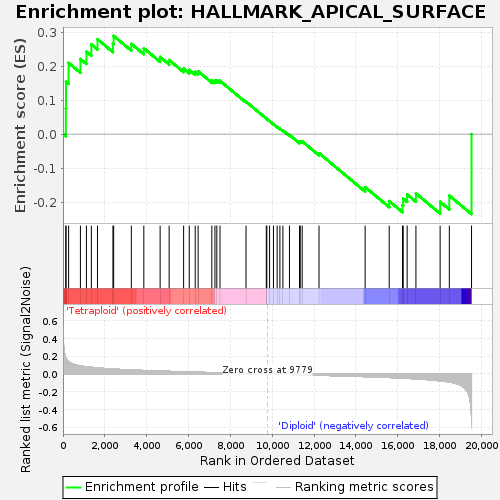

### enplot_HALLMARK_APOPTOSIS_337.png

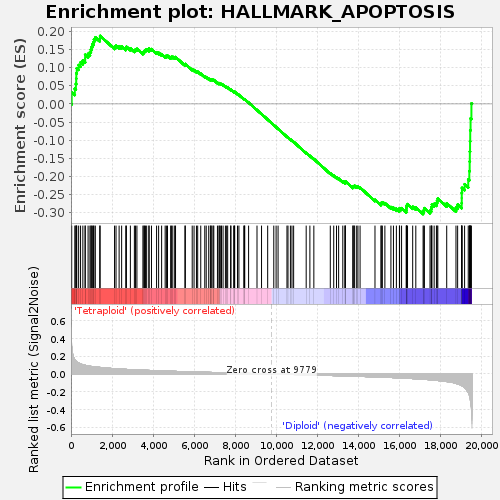

### enplot_HALLMARK_BILE_ACID_METABOLISM_358.png

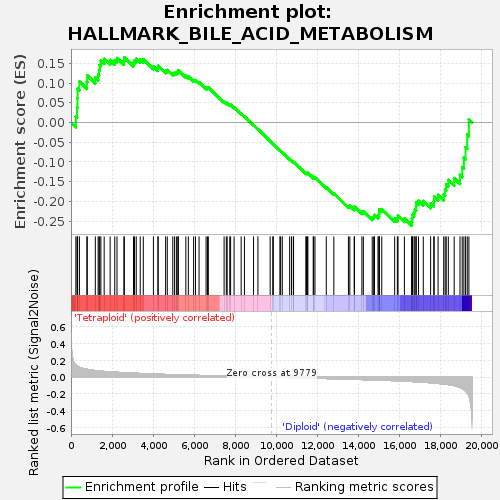

### enplot_HALLMARK_CHOLESTEROL_HOMEOSTASIS_259.png

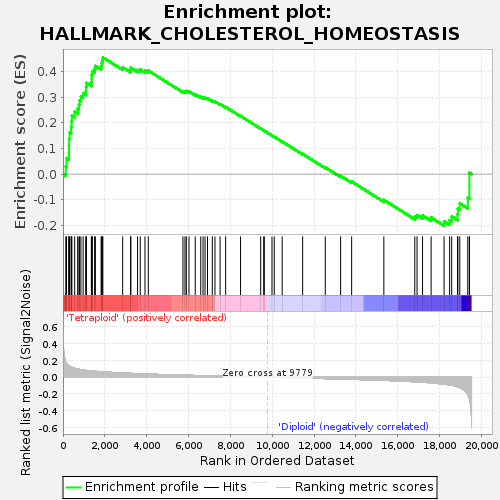

### enplot_HALLMARK_COMPLEMENT_313.png

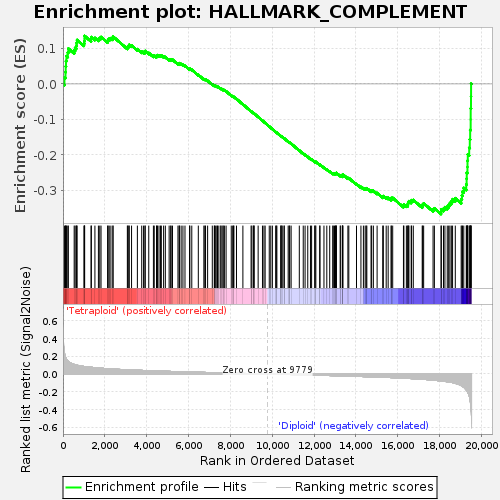

### enplot_HALLMARK_E2F_TARGETS_247.png

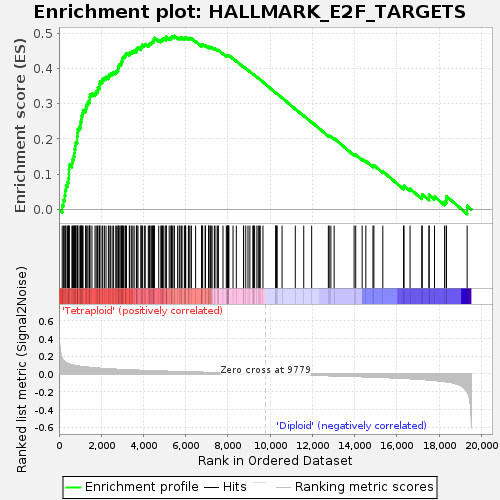

### enplot_HALLMARK_EPITHELIAL_MESENCHYMAL_TRANSITION_322.png

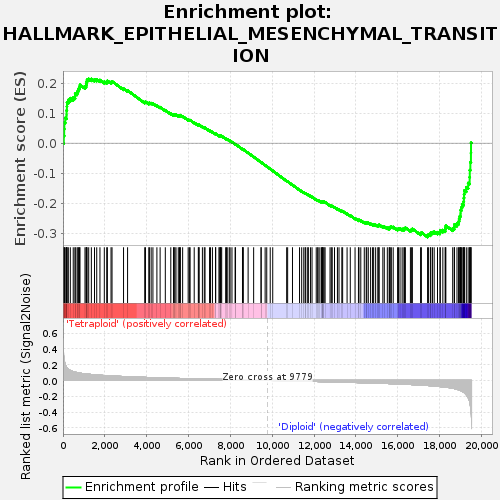

### enplot_HALLMARK_ESTROGEN_RESPONSE_EARLY_271.png

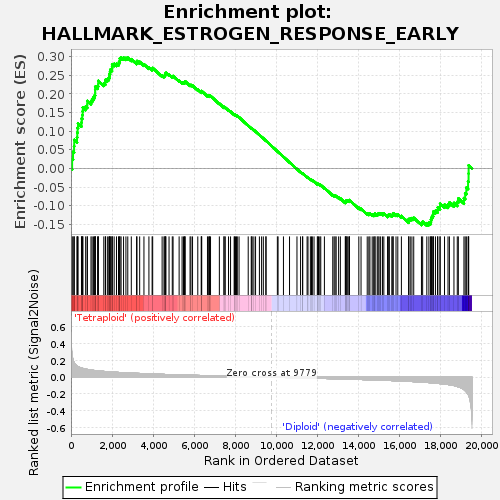

### enplot_HALLMARK_FATTY_ACID_METABOLISM_286.png

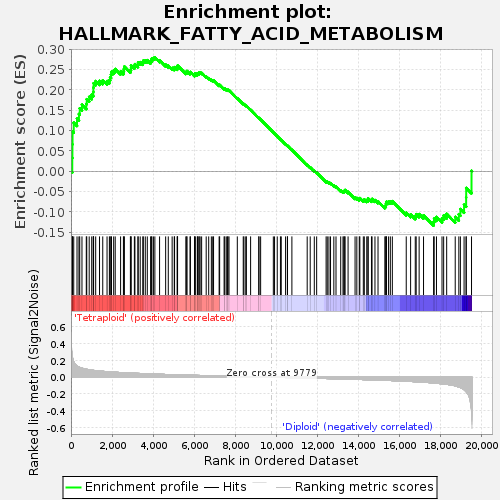

### enplot_HALLMARK_G2M_CHECKPOINT_256.png

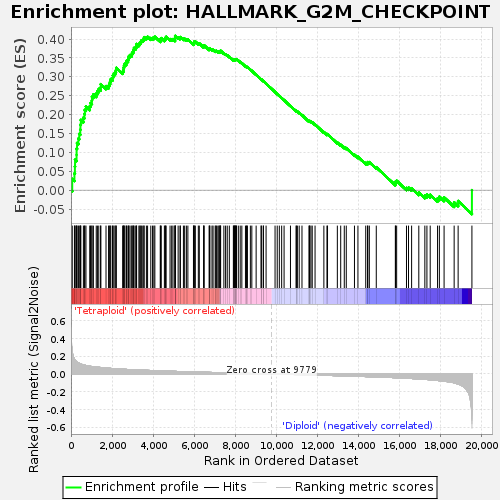

### enplot_HALLMARK_HEME_METABOLISM_340.png

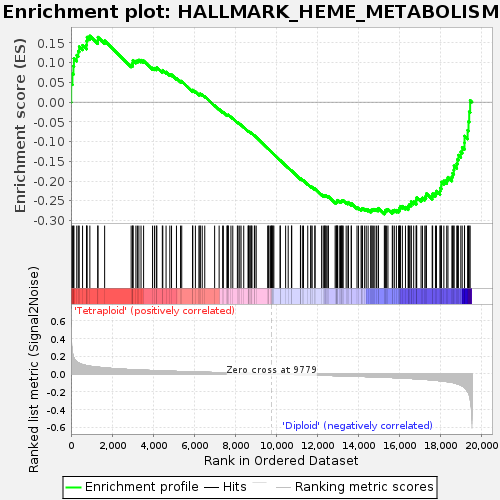

### enplot_HALLMARK_HYPOXIA_328.png

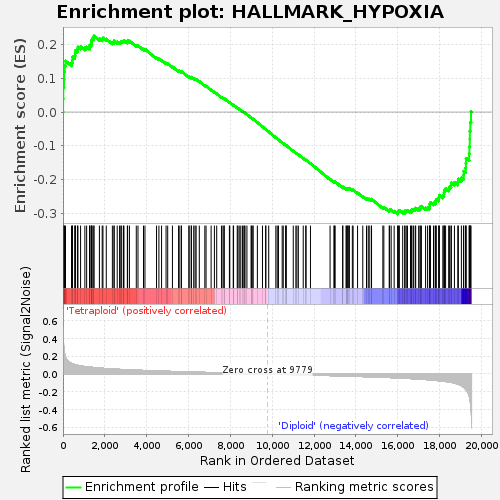

### enplot_HALLMARK_IL2_STAT5_SIGNALING_280.png

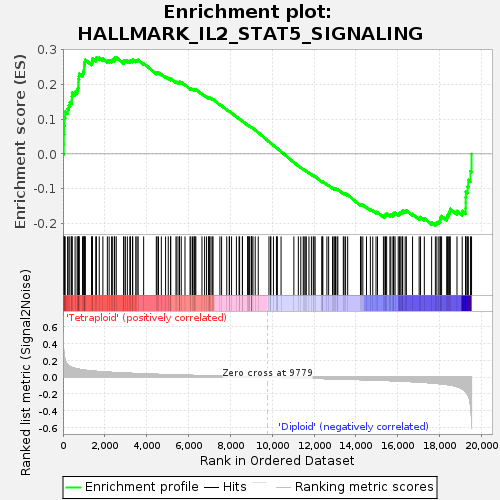

### enplot_HALLMARK_INFLAMMATORY_RESPONSE_262.png

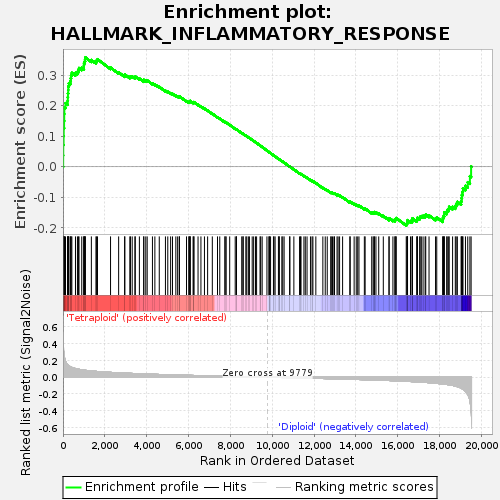

### enplot_HALLMARK_INTERFERON_GAMMA_RESPONSE_343.png

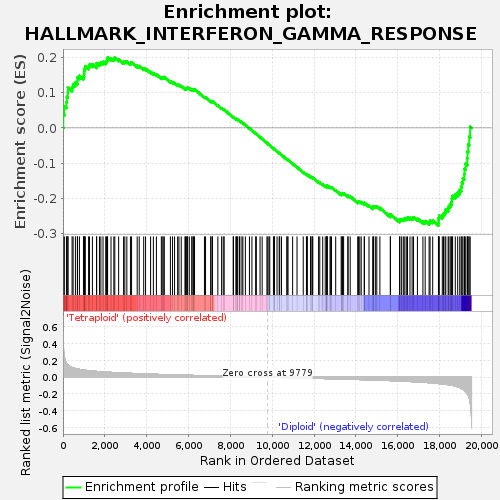

### enplot_HALLMARK_KRAS_SIGNALING_DN_283.png

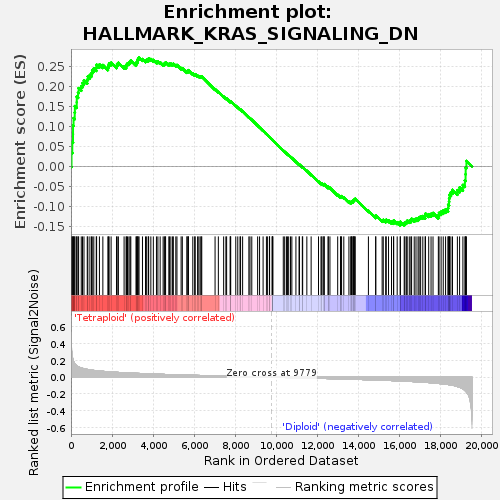

### enplot_HALLMARK_KRAS_SIGNALING_UP_352.png

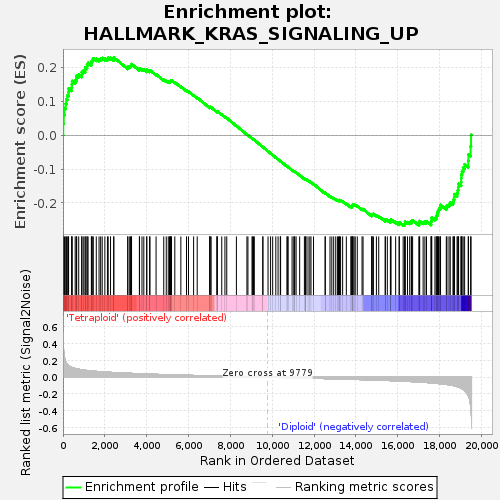

### enplot_HALLMARK_MTORC1_SIGNALING_346.png

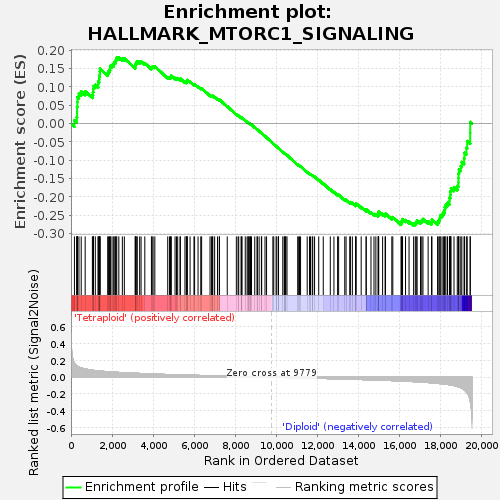

### enplot_HALLMARK_MYC_TARGETS_V2_250.png

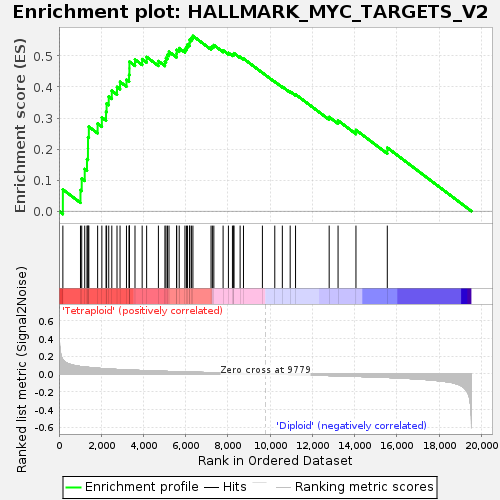

### enplot_HALLMARK_P53_PATHWAY_265.png

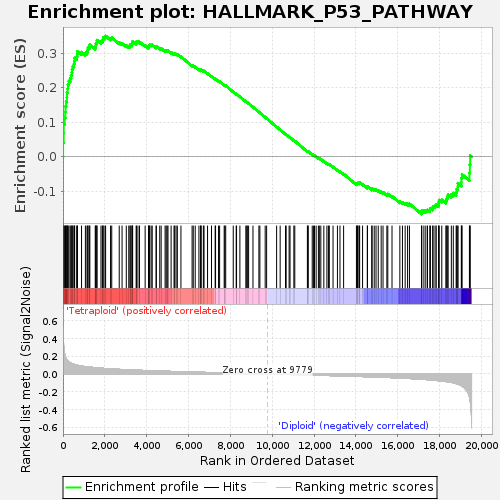

### enplot_HALLMARK_PI3K_AKT_MTOR_SIGNALING_319.png

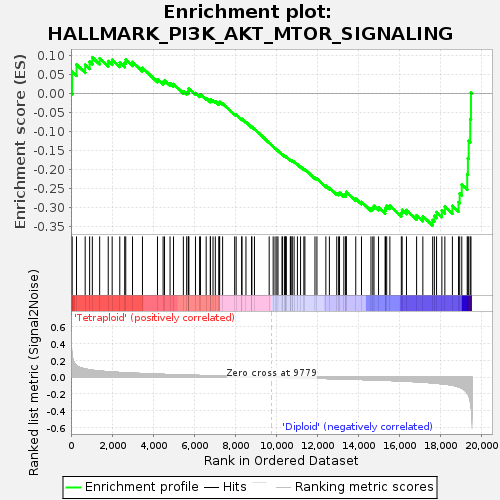

### enplot_HALLMARK_SPERMATOGENESIS_304.png

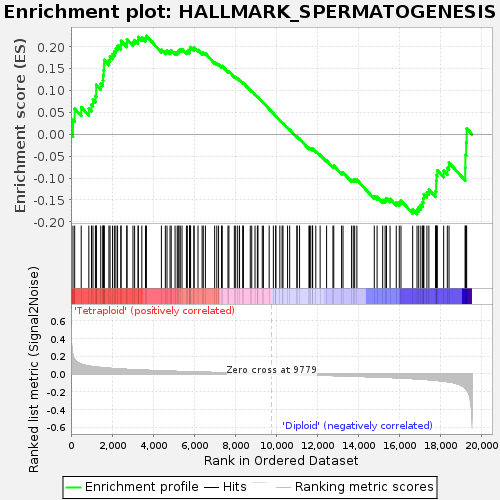

### enplot_HALLMARK_TGF_BETA_SIGNALING_292.png

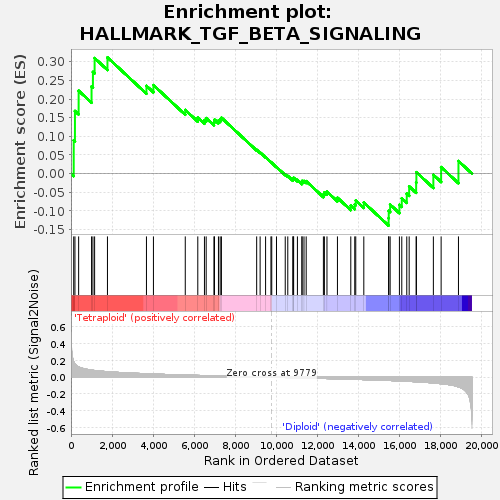

### enplot_HALLMARK_TNFA_SIGNALING_VIA_NFKB_289.png

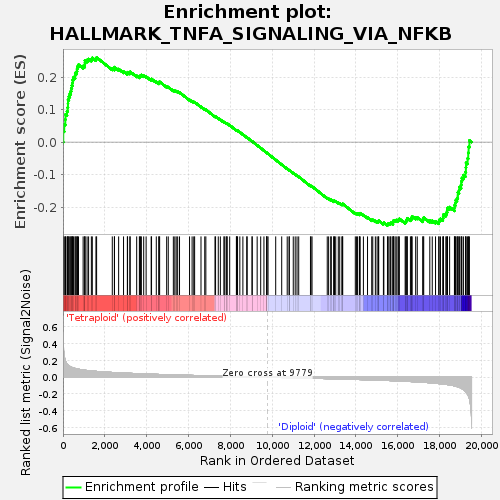

### enplot_HALLMARK_UNFOLDED_PROTEIN_RESPONSE_307.png

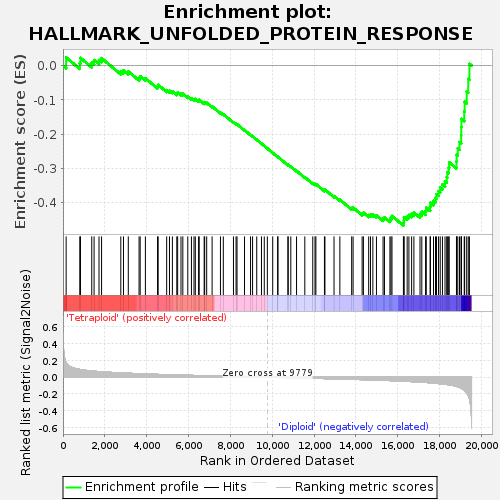

### enplot_HALLMARK_UV_RESPONSE_UP_355.png

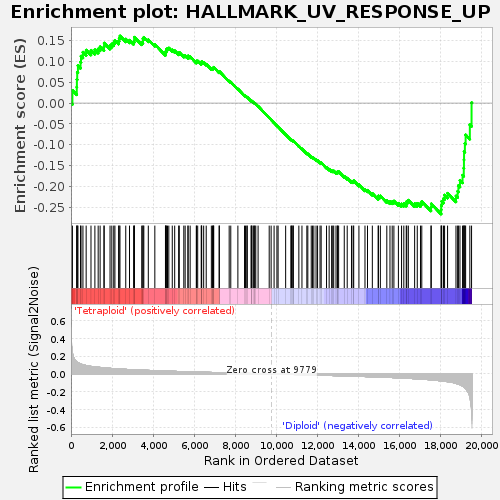

### enplot_HALLMARK_WNT_BETA_CATENIN_SIGNALING_316.png

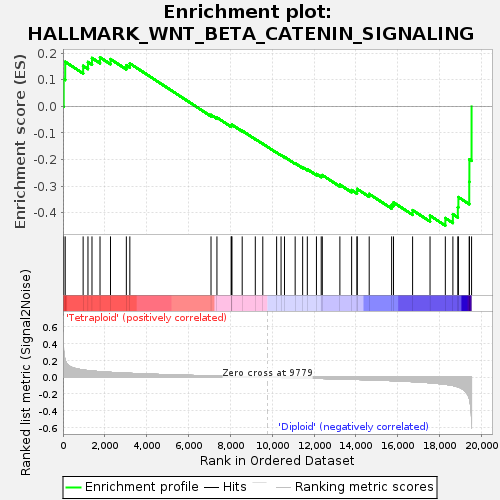

### global_es_histogram.png

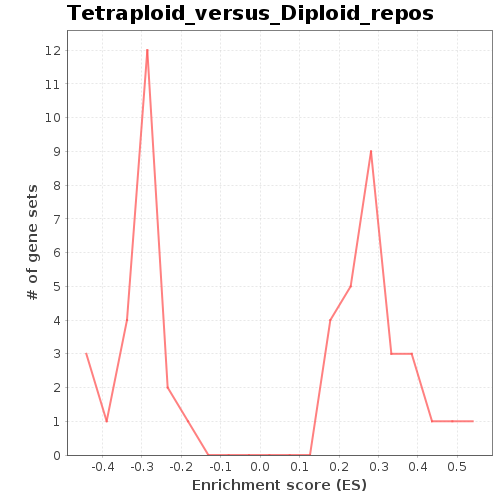

### gset_rnd_es_dist_249.png

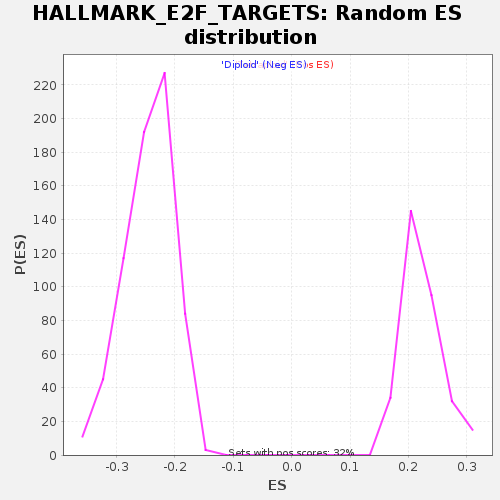

### gset_rnd_es_dist_252.png

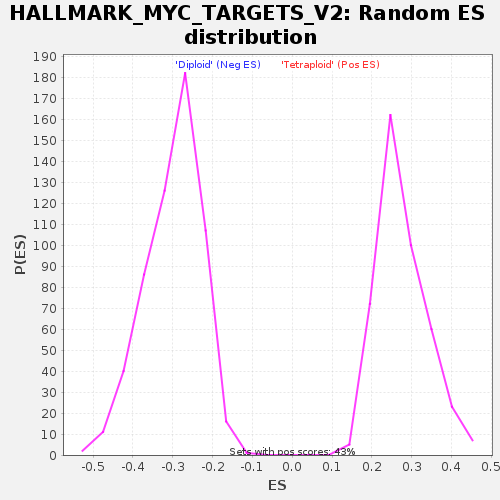

### gset_rnd_es_dist_255.png

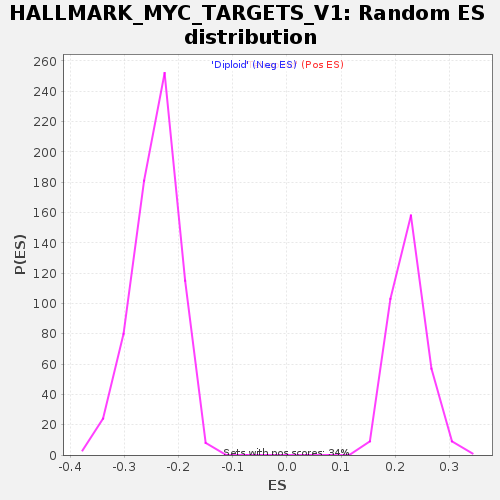

### gset_rnd_es_dist_258.png

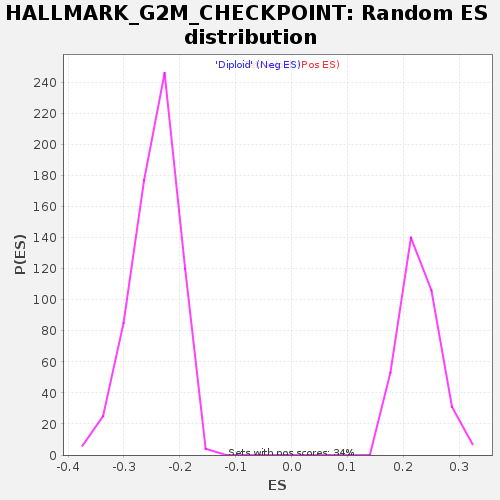

### gset_rnd_es_dist_261.png

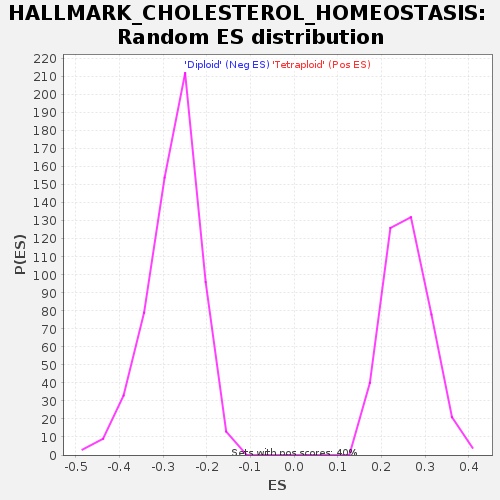

### gset_rnd_es_dist_264.png

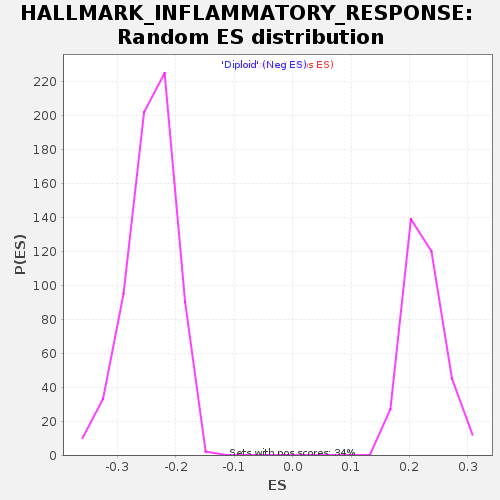

### gset_rnd_es_dist_267.png

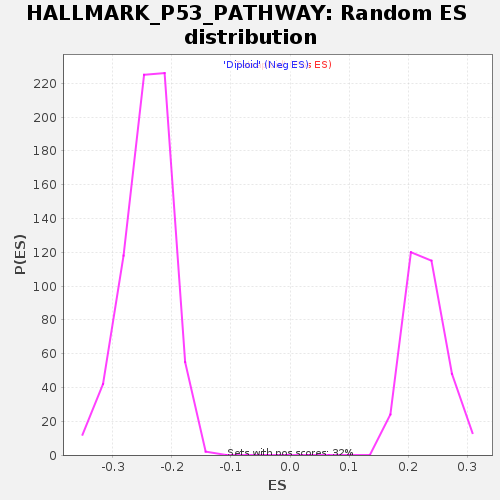

### gset_rnd_es_dist_270.png

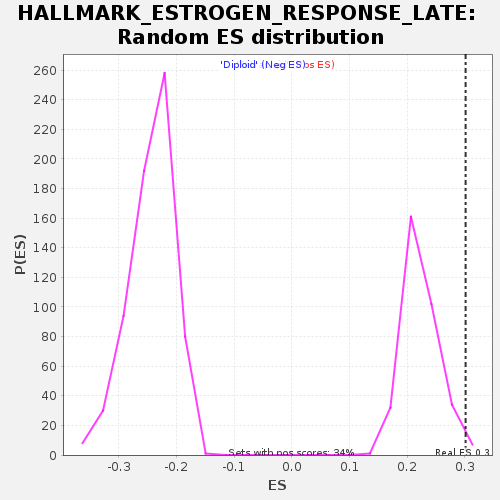

### gset_rnd_es_dist_276.png

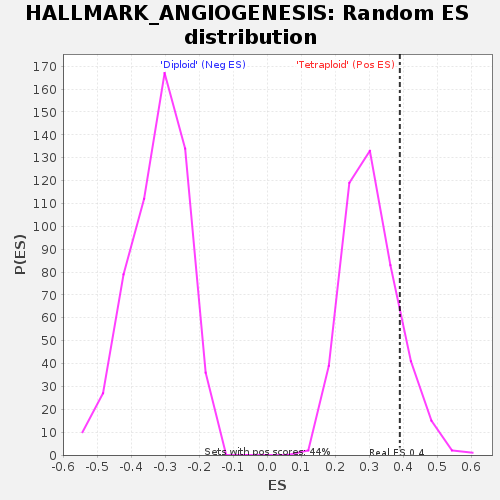

### gset_rnd_es_dist_279.png

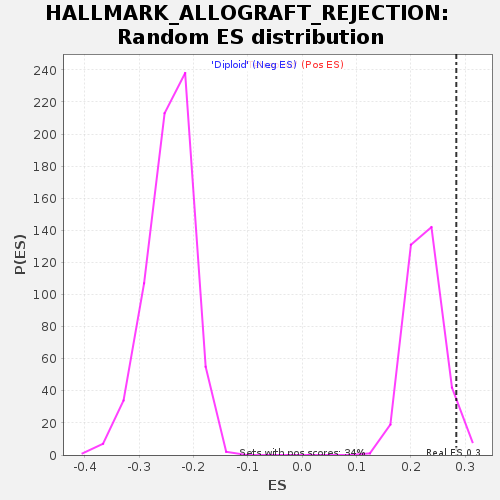

### gset_rnd_es_dist_282.png

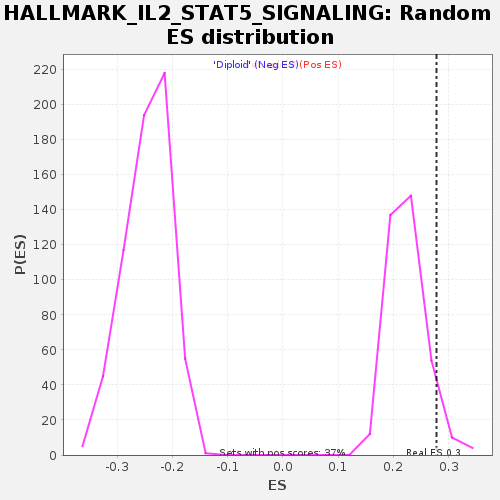

### gset_rnd_es_dist_285.png

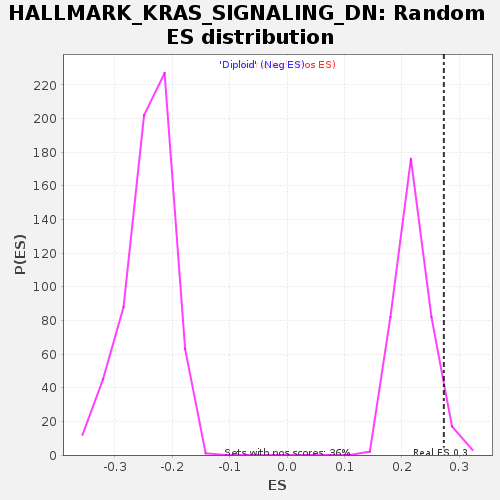

### gset_rnd_es_dist_291.png

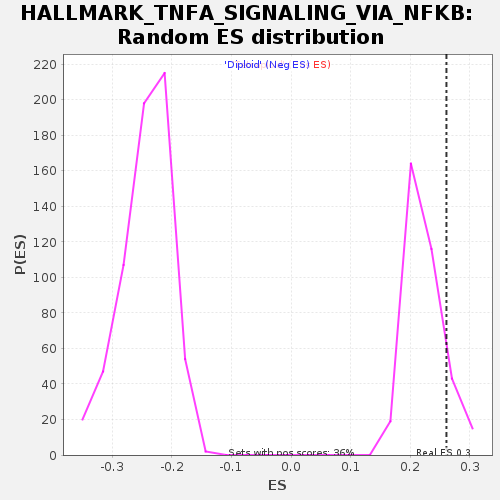

### gset_rnd_es_dist_294.png

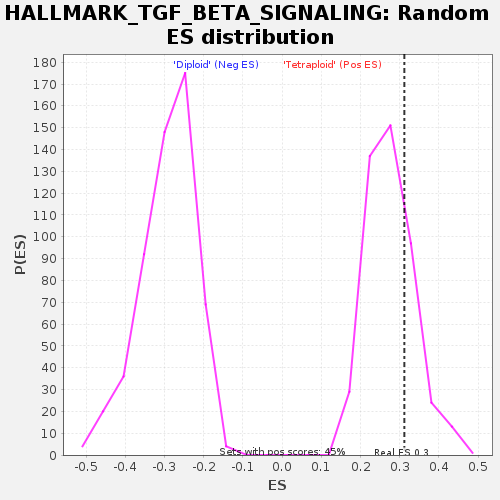

### gset_rnd_es_dist_297.png

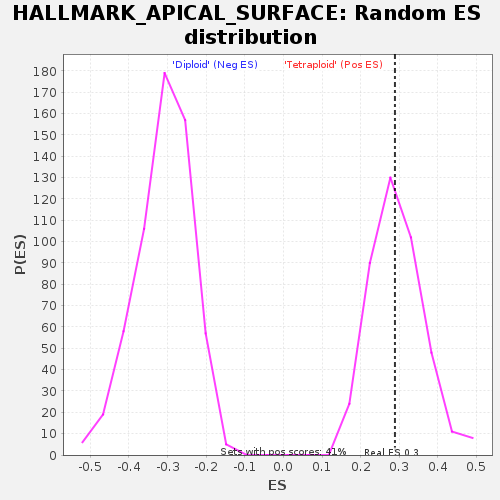

### gset_rnd_es_dist_300.png

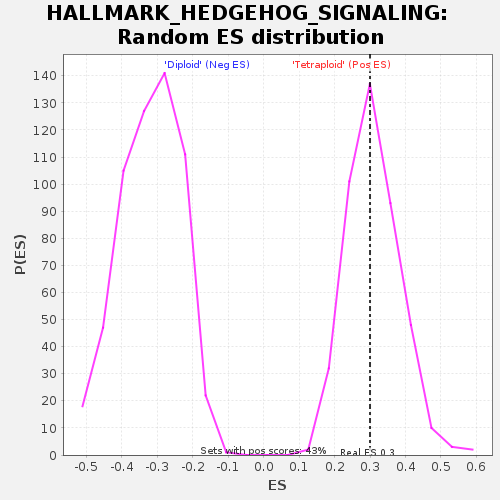

### gset_rnd_es_dist_303.png

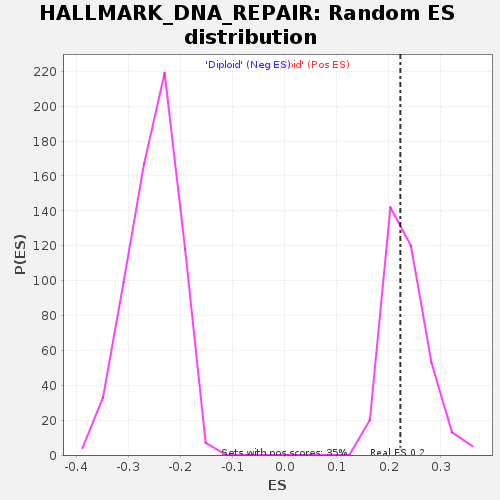

### gset_rnd_es_dist_306.png

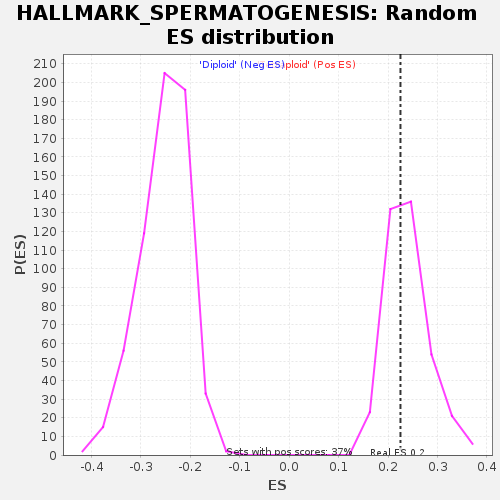

### gset_rnd_es_dist_312.png

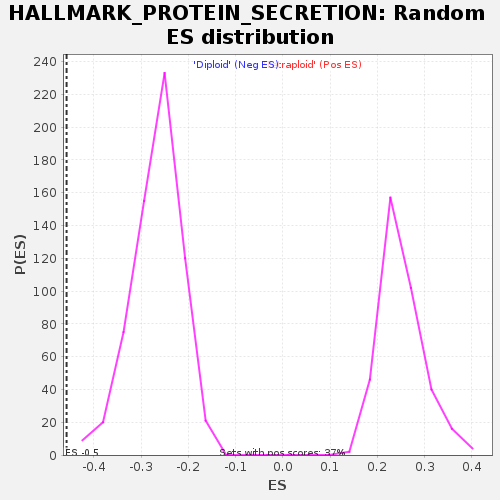

### gset_rnd_es_dist_315.png

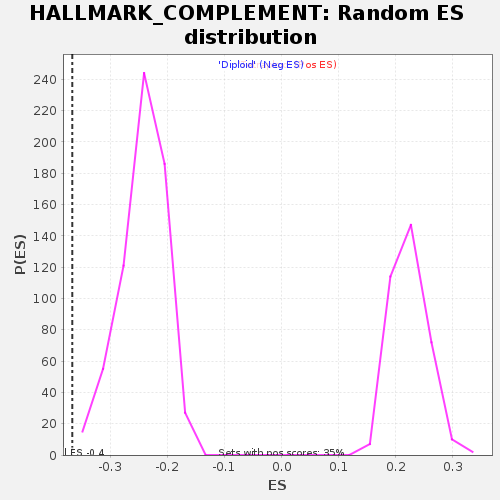

### gset_rnd_es_dist_318.png

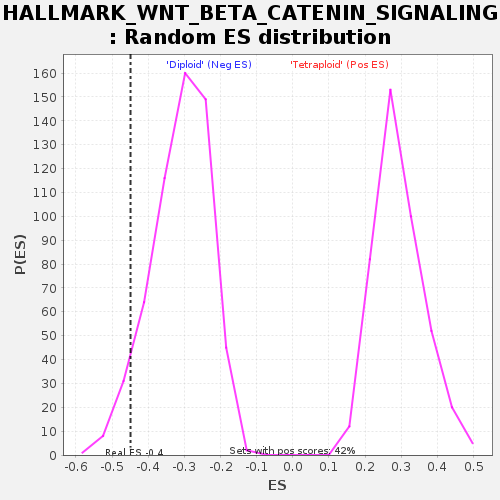

### gset_rnd_es_dist_324.png

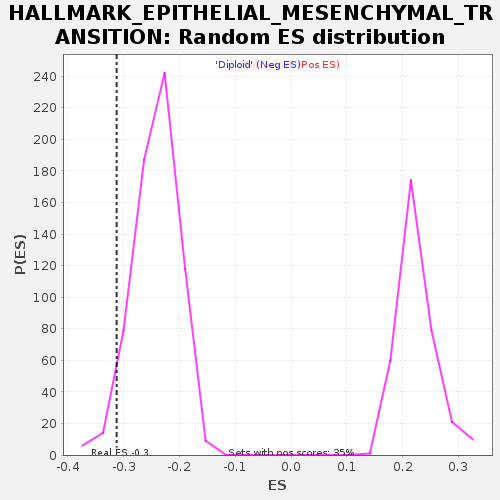

### gset_rnd_es_dist_327.png

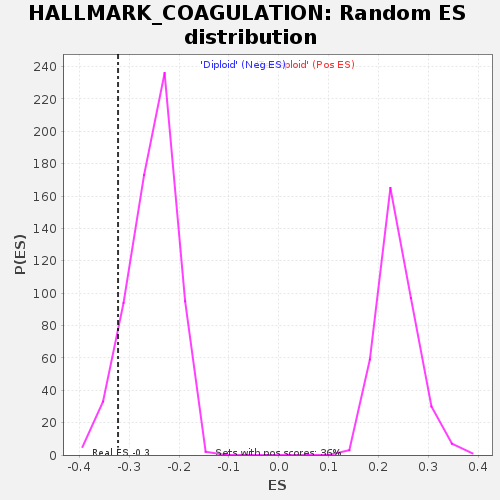

### gset_rnd_es_dist_330.png

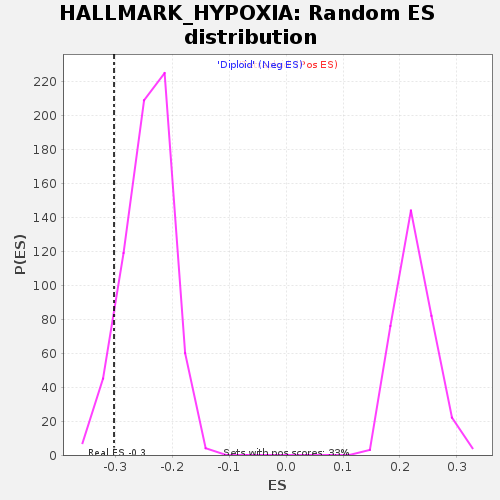

### gset_rnd_es_dist_336.png

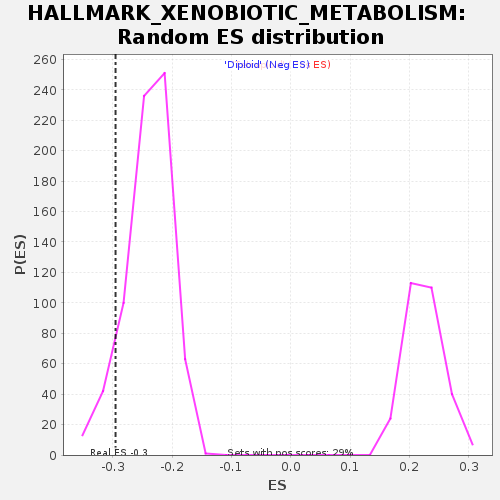

### gset_rnd_es_dist_339.png

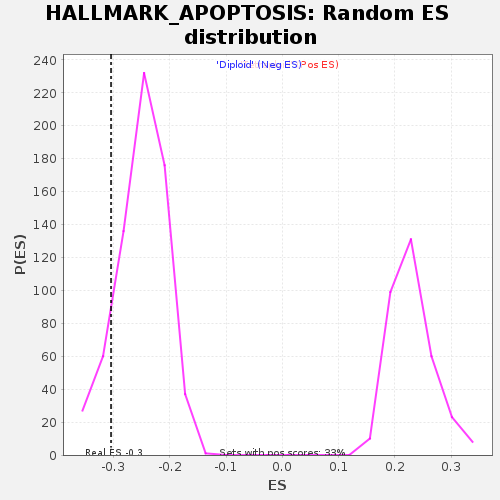

### gset_rnd_es_dist_342.png

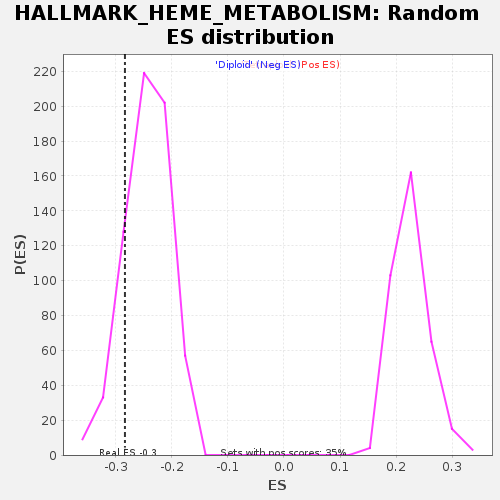

### gset_rnd_es_dist_345.png

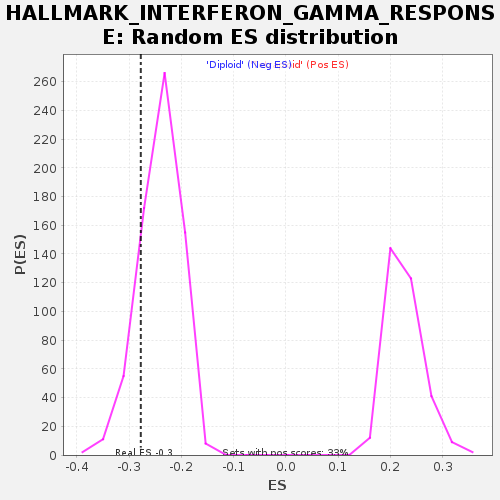

### gset_rnd_es_dist_348.png

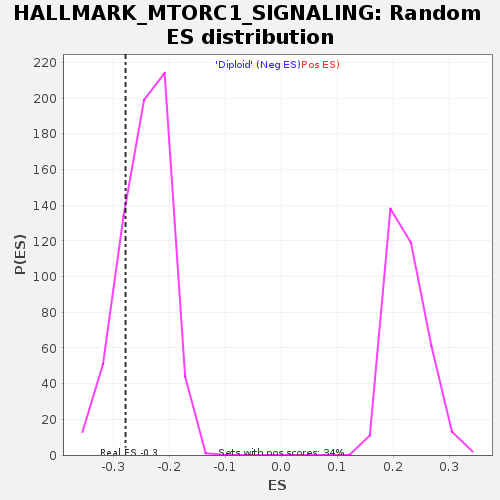

### gset_rnd_es_dist_351.png

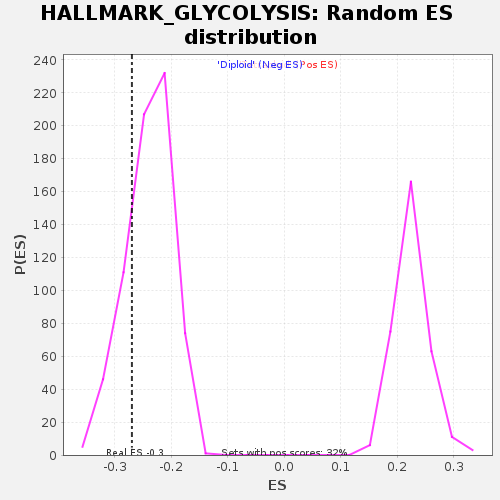

### gset_rnd_es_dist_354.png

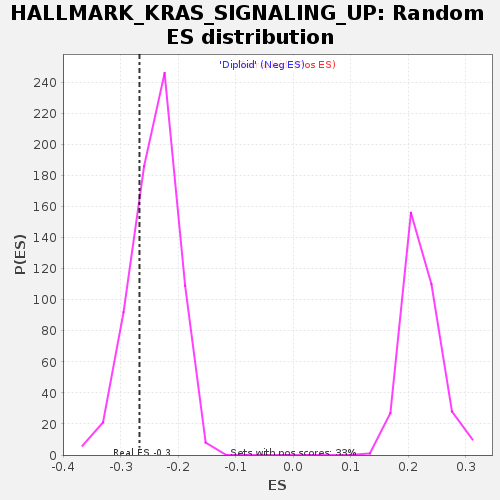

### gset_rnd_es_dist_357.png

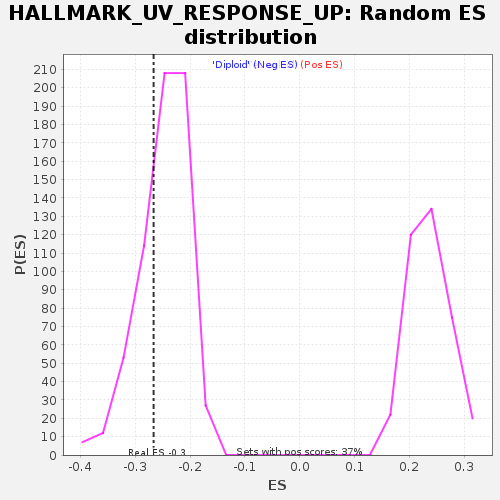

### gset_rnd_es_dist_363.png

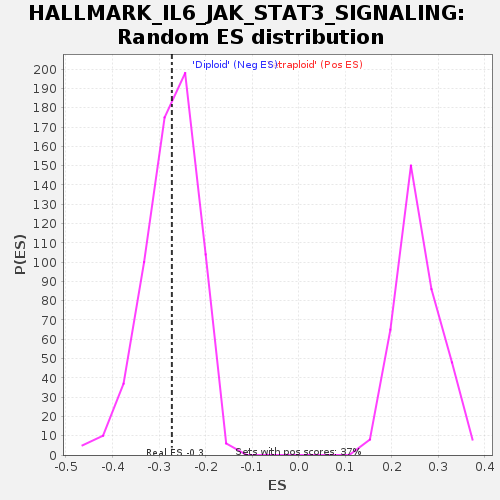

### gset_rnd_es_dist_366.png

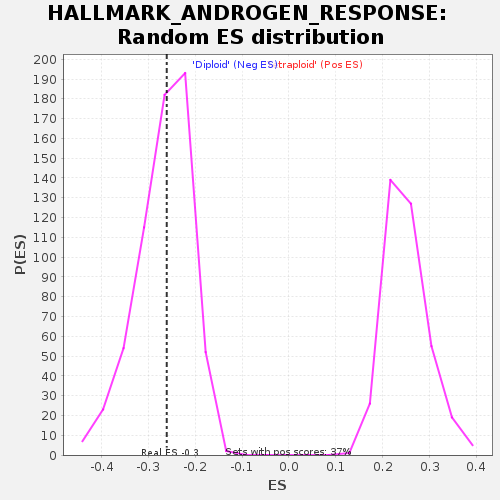

### HALLMARK_ALLOGRAFT_REJECTION_278.png

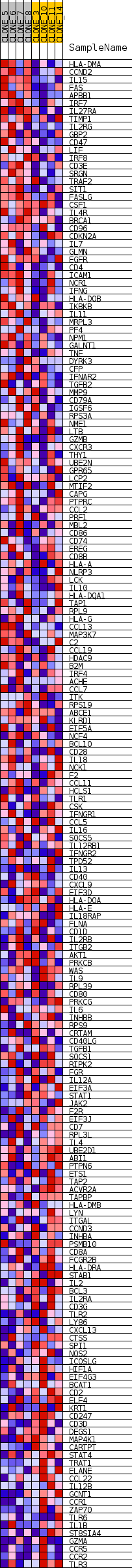

### HALLMARK_ANGIOGENESIS_275.png

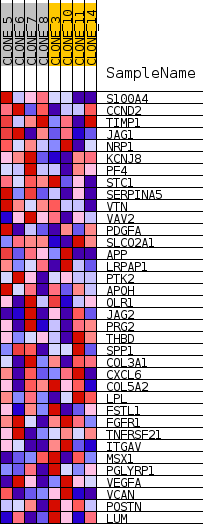

### HALLMARK_APICAL_SURFACE_296.png

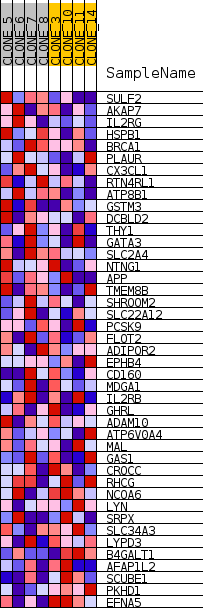

### HALLMARK_APOPTOSIS_338.png

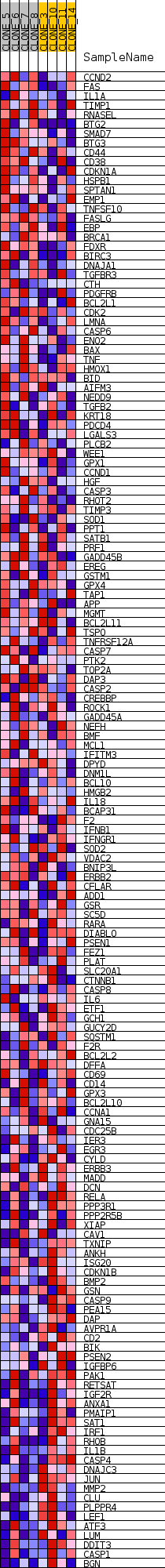

### HALLMARK_BILE_ACID_METABOLISM_359.png

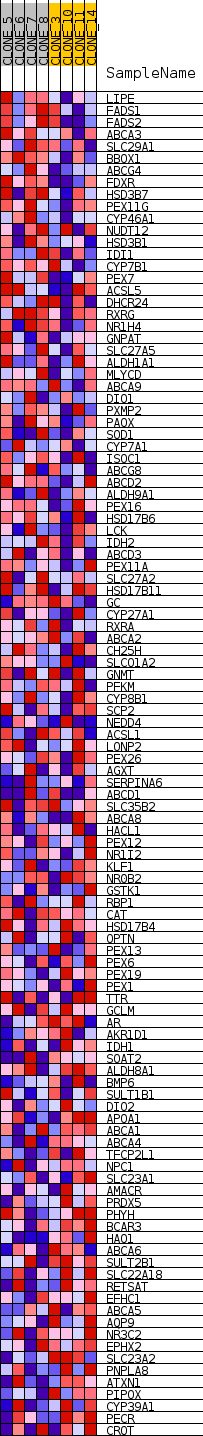

### HALLMARK_CHOLESTEROL_HOMEOSTASIS_260.png

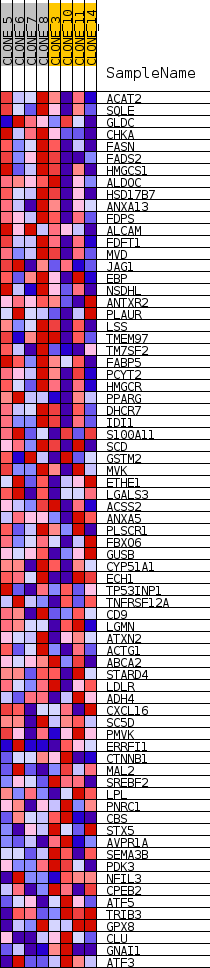

### HALLMARK_DNA_REPAIR_302.png

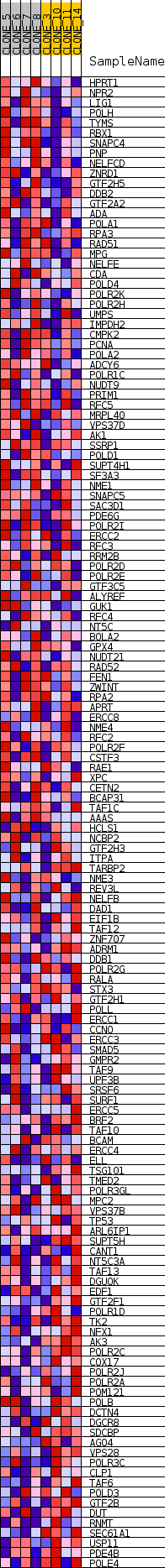

### HALLMARK_E2F_TARGETS_248.png

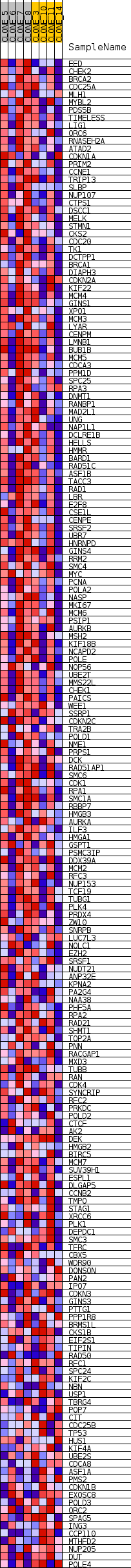

### HALLMARK_EPITHELIAL_MESENCHYMAL_TRANSITION_323.png

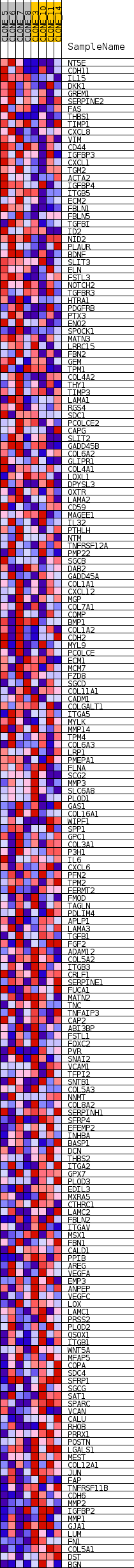

### HALLMARK_ESTROGEN_RESPONSE_LATE_269.png

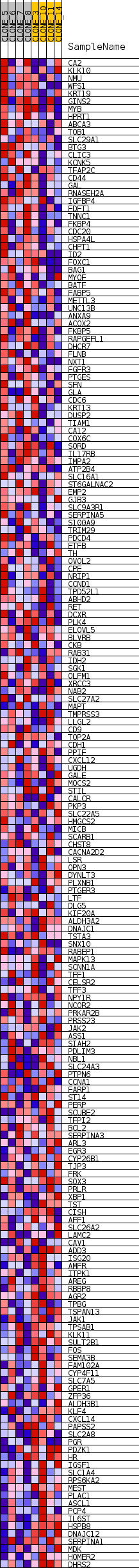

### HALLMARK_FATTY_ACID_METABOLISM_287.png

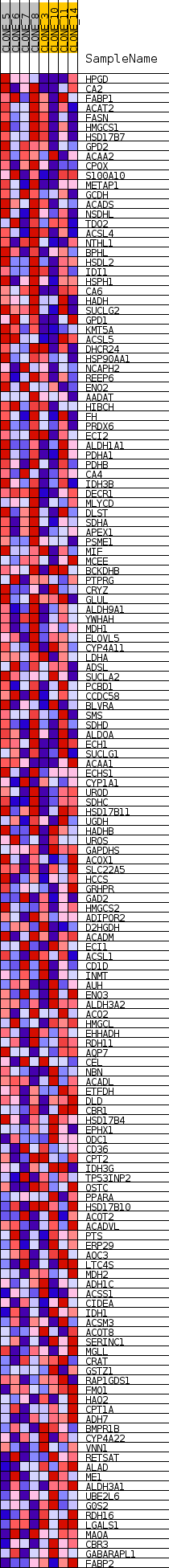

### HALLMARK_G2M_CHECKPOINT_257.png

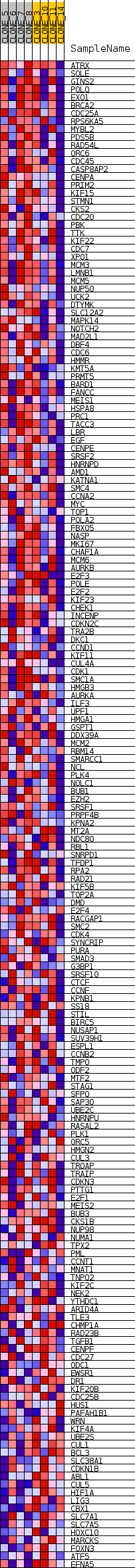

### HALLMARK_GLYCOLYSIS_350.png

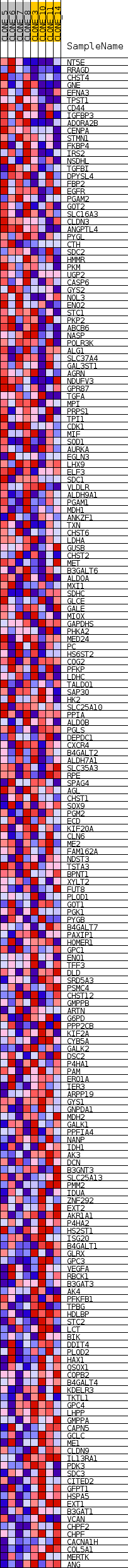

### HALLMARK_HEDGEHOG_SIGNALING_299.png

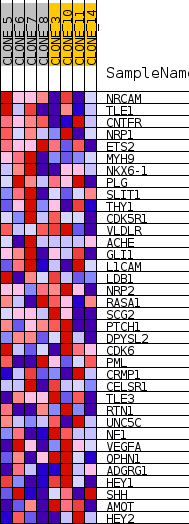

### HALLMARK_HEME_METABOLISM_341.png

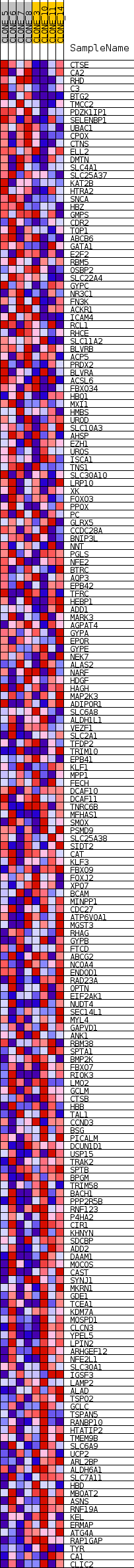

### HALLMARK_HYPOXIA_329.png

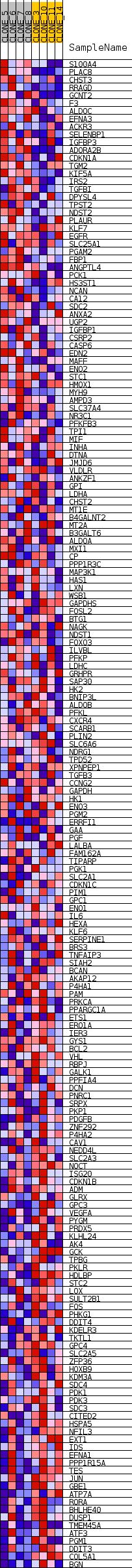

### HALLMARK_IL2_STAT5_SIGNALING_281.png

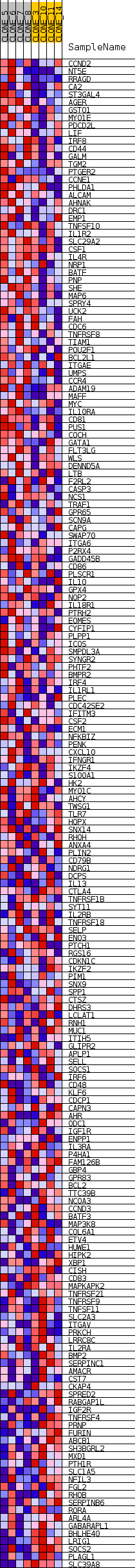

### HALLMARK_IL6_JAK_STAT3_SIGNALING_362.png

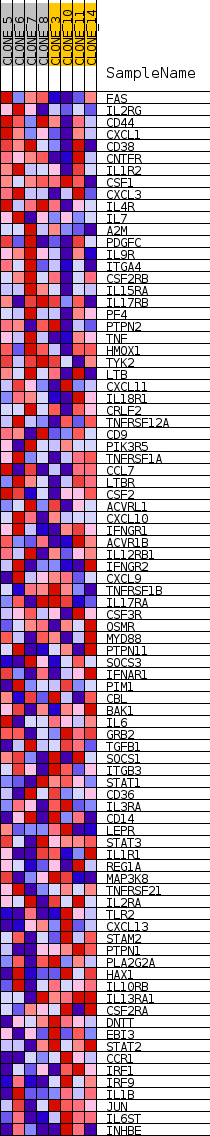

### HALLMARK_INFLAMMATORY_RESPONSE_263.png

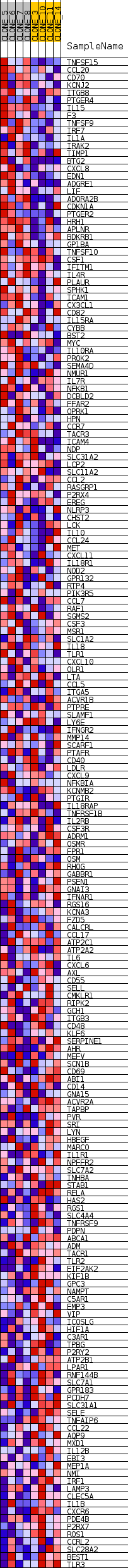

### HALLMARK_INTERFERON_ALPHA_RESPONSE_332.png

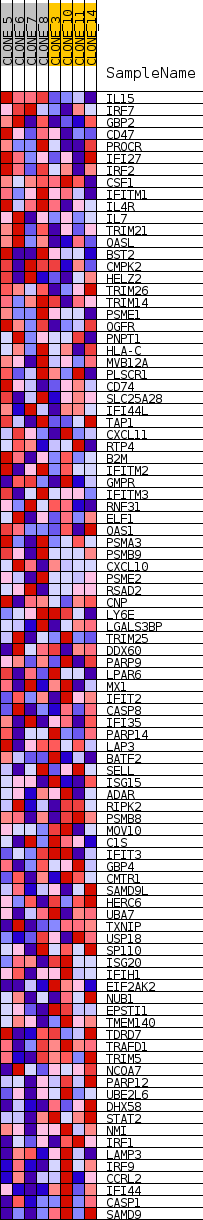

### HALLMARK_INTERFERON_GAMMA_RESPONSE_344.png

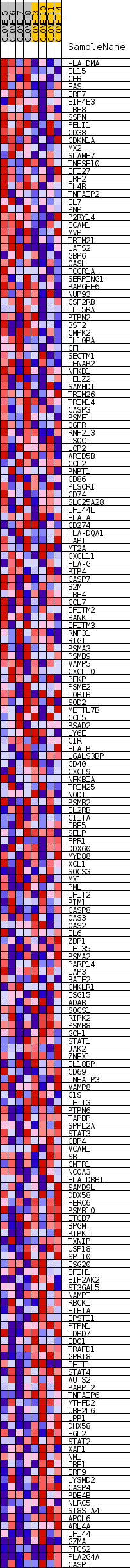

### HALLMARK_KRAS_SIGNALING_DN_284.png

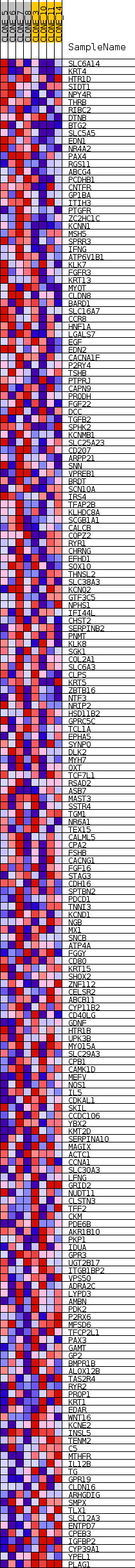

### HALLMARK_KRAS_SIGNALING_UP_353.png

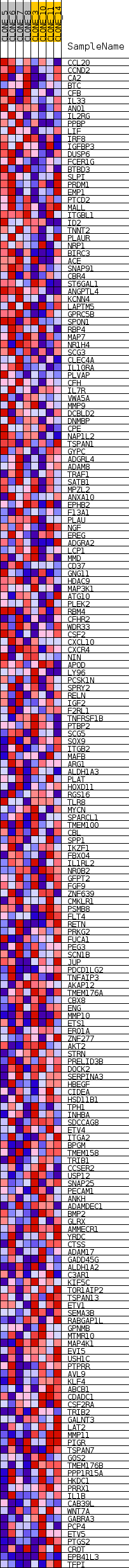

### HALLMARK_MYC_TARGETS_V1_254.png

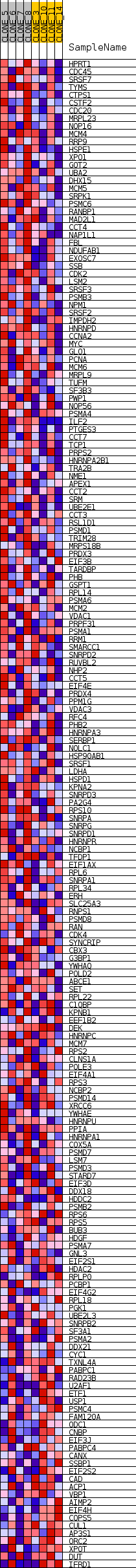

### HALLMARK_MYC_TARGETS_V2_251.png

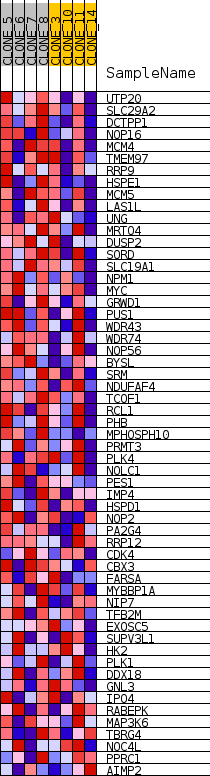

### HALLMARK_PI3K_AKT_MTOR_SIGNALING_320.png

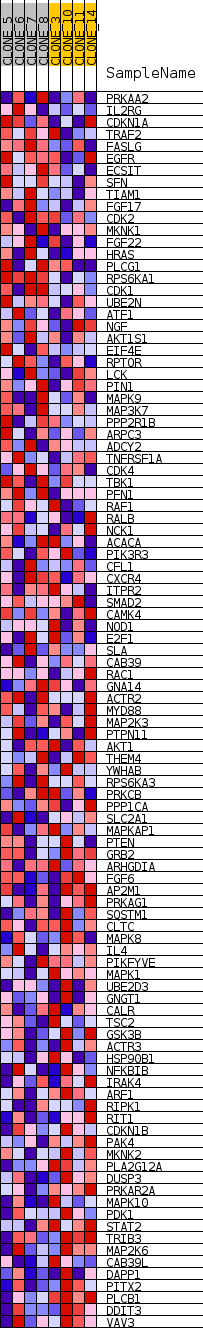

### HALLMARK_PROTEIN_SECRETION_311.png

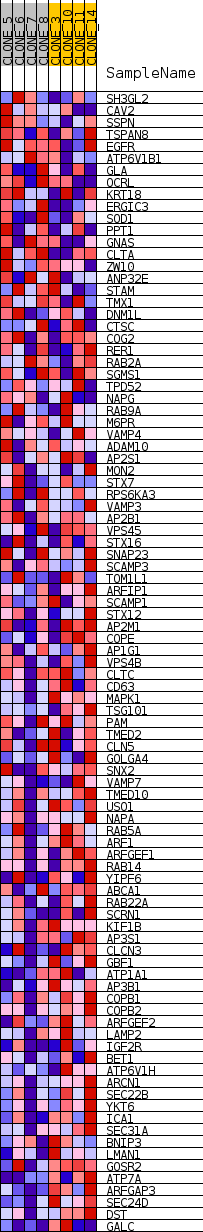

### HALLMARK_TGF_BETA_SIGNALING_293.png

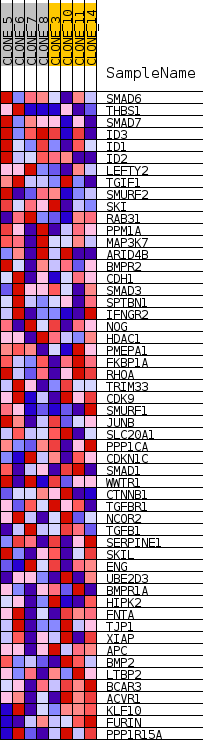

### HALLMARK_TNFA_SIGNALING_VIA_NFKB_290.png

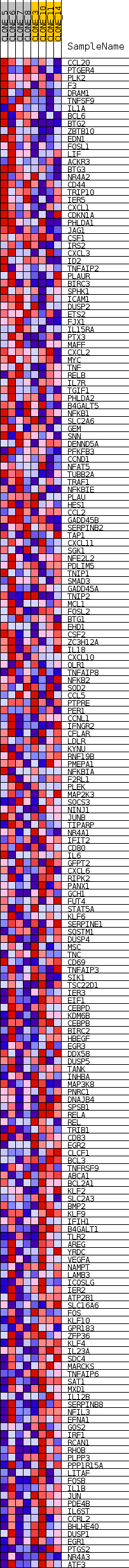

### HALLMARK_UV_RESPONSE_UP_356.png

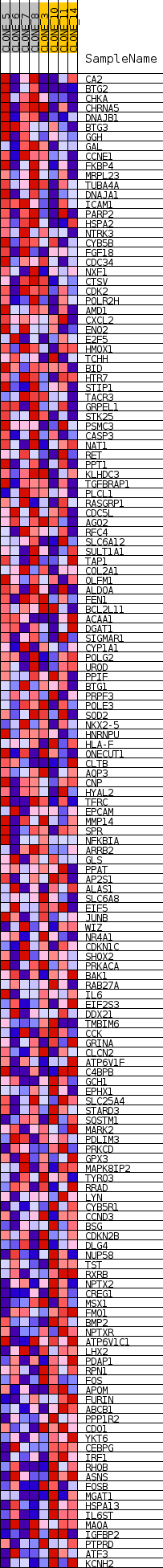

### HALLMARK_WNT_BETA_CATENIN_SIGNALING_317.png

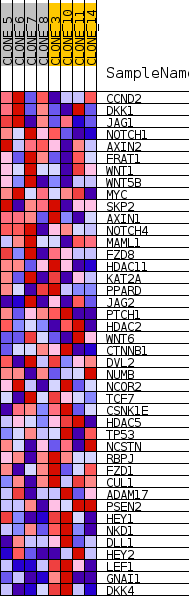

### HALLMARK_XENOBIOTIC_METABOLISM_335.png

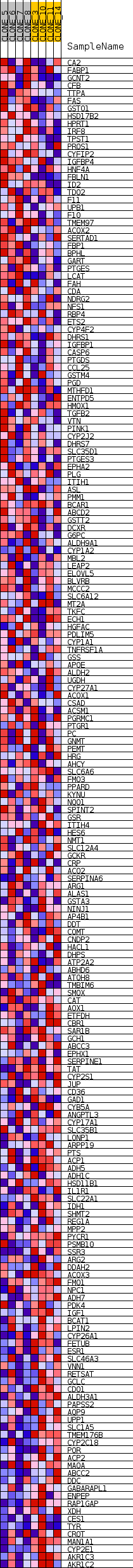

### heat_map_245.png

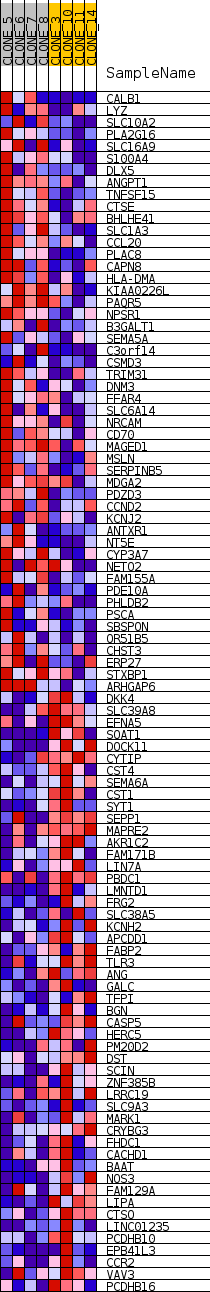

### pvalues_vs_nes_plot.png

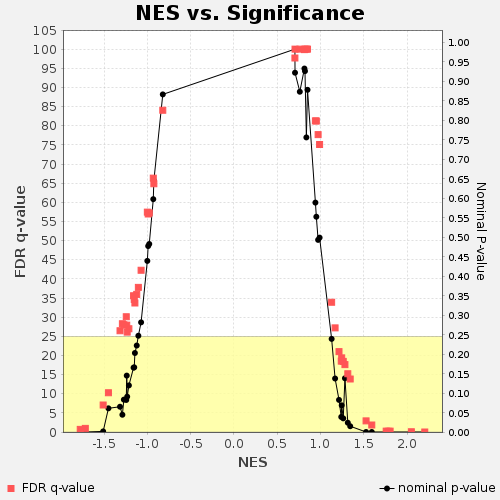

### ranked_list_corr_246.png

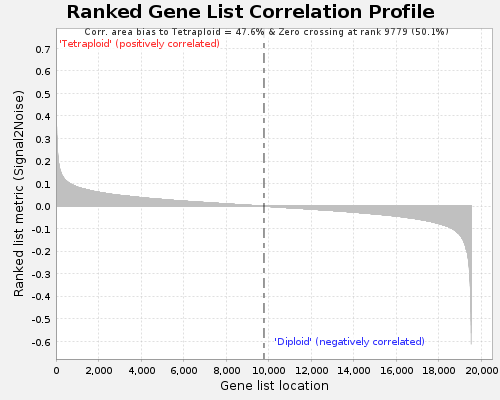
